# SEA: The small RNA Expression Atlas

**DOI:** 10.1101/133199

**Authors:** Raza-Ur Rahman, Vikas Bansal, Maksims Fiosins, Anna-Maria Liebhoff, Ashish Rajput, Abdul Sattar, Daniel Sumner Magruder, Sumit Madan, Ting Sun, Abhivyakti Gautam, Sven Heins, Timur Liwinski, Jörn Bethune, Claudia Trenkwalder, Juliane Fluck, Brit Mollenhauer, Stefan Bonn

**Affiliations:** Institute of Medical Systems Biology, Center for Molecular Neurobiology, University Medical Center Hamburg-Eppendorf, Hamburg, Germany; German Center for Neurodegenerative Diseases, Tübingen, Germany; Genevention GmbH, Göttingen, Germany; Fraunhofer Institute for Algorithms and Scientific Computing, Schloss Birlinghoven, Sankt Augustin, Germany; University Medical Center Goettingen; Rheinische Friedrich-Wilhelms-Universität Bonn, Bonn 53113, Germany; German National Library of Medicine (ZB MED) - Information Centre for Life Sciences, Bonn, Germany; Institute of Geodesy and Geoinformation, University of Bonn, Germany; Department of Neurology, University Medical Center Goettingen, Robert-Koch Strasse 40, 37075 Germany; Paracelsus-Elena-Klinik, Klinikstraße 16, 34119 Kassel, Germany; Department of Neurosurgery, University Medical Center Goettingen, Robert-Koch Strasse 40, 37075 Germany; Department of Neurogenetics, Max Planck Institute of Experimental Medicine, Göttingen, Germany; 1st Department of Medicine, University Medical Center Hamburg-Eppendorf, Hamburg, Germany

**Keywords:** small RNA, miRNA, differential expression, classification, prediction, biomarker, virus, bacteria, pathogens, disease, atlas, database, Parkinson’s disease

## Abstract

We present the Small RNA Expression Atlas (SEA), a web application that allows for the interactive querying, visualization, and analysis of known and novel small RNAs across ten organisms. It contains sRNA and pathogen expression information for over 4,200 published samples with standardized search terms and ontologies. In addition, SEA allows for the interactive visualization and re-analysis of 879 differential expression and 514 classification comparisons. SEA’s user model enables sRNA researchers to compare and re-analyze user-specific and published datasets, highlighting common and distinct sRNA expression patterns.

We provide evidence for SEA’s fidelity by (i) generating a set of 591 tissue specific miRNAs across 30 tissues, (ii) finding known and novel bacterial and viral infections across diseases, and (iii) determining a Parkinson’s disease-specific blood biomarker signature using novel data.

We believe that SEA’s simple semantic search interface, the flexible interactive reports, and the user model with rich analysis capabilities will enable researchers to better understand the potential function and diagnostic value of sRNAs or pathogens across tissues, diseases, and organisms.

**Availability and Implementation:** SEA is implemented in Java, J2EE, spring, Django, html5, css3, JavaScript, Bootstrap, Vue.js, D3, mongodb and neo4j. It is freely available at http://sea.ims.bio/.

## 1 Background

Small RNAs (sRNAs) are a class of short, non-coding RNAs with important biological functions in nearly all aspects of organismal development in health and disease. Especially in diagnostic and therapeutic research, sRNAs such as miRNAs and piRNAs received recent attention (Witwer, 2015). The increasing number of deep sequencing sRNA studies (sRNA-seq) is reflecting the importance of sRNAs in biological processes as well as disease diagnosis and therapy. To harvest the true potential of existing data, it is important to allow for querying, visualization, and analysis of sRNA-seq data across organisms, tissues, cell types, and disease states. This would allow researchers, for example, to search for disease-specific sRNA biomarker signatures across all disease entities investigated. Data integration and interoperability require (i) a streamlined analysis workflow to reduce analysis bias between experiments (ii) also necessitates standardized annotation using ontologies to search and retrieve relevant samples and (iii) flexible and interactive visualization of the data.

To date, several web-based sRNA-seq expression profile databases are available that differ in their level of information, portfolio, performance, and user-friendliness. Recent additions to sRNA web based databases include miRmine (Panwar et al., 2017), provides expression of a single or multiple miRNAs for a specific tissue, cell-line or disease. Results are displayed in multiple interactive, graphical and downloadable formats; DASHR2 (Kuksa et al., 2019) supports sRNA expression profiles across different genome versions of the same species across tissues and cell types. Results are provided in an interactive manner, such as sncRNA locus sorting and filtering by biological features. All annotation and expression information are downloadable and accessible as UCSC genome browser tracks; miratlas (Vitsios et al., 2017) allows for searching miRNA expression profiles as well as sRNA-seq experiments and provides information on the miRNA modification analysis; and YM500v3 (Chung et al., 2017) provides interactive web reports on sRNA expression profiles, sRNA differential expression and miRNA gene targets.

Although many good web platforms for the sRNA-seq data exist, some important aspects for storing and searching have yet to be integrated. For example, no current web application allows for the ontology based search of sRNA-seq experiments. Current tools lack an important association of miRNAs with disease. miRNA disease associations are provided by HMDD (Li et al., 2014), but it does not provide miRNA expression information. Except YM500v3 (Chung et al., 2017), current tools do not provide miRNAs and gene targets, of note YM500v3 is only limited to cancer miRNome studies. Also, there is currently no web application that allows for the identification of biomarkers of disease via machine-learning. The above mentioned web platforms do not provide expression of novel miRNAs in known disease state or tissues, including the structure and probability of the novel miRNA prediction. To our knowledge no other tool provides pathogenic signatures from sRNA-seq data including their differential expression in healthy and diseased condition. Moreover, in current tools users can only search for the results that are stored, there is no option for the users to reanalyze data with the samples of their choice. Finally, current sRNA-seq web services do not allow for the user data upload, a feature that would greatly facilitate researchers to compare their in-house sRNA-seq experimental data with the publicly available data. In the end, these functionalities should be paired with a flexible and interactive visualization of the sRNA-seq data supporting more species and cross study comparisons.

In order to address the above mentioned limitations, we hereby present the **<u>s</u>**mall-RNA **<u>E</u>**xpression **<u>A</u>**tlas (SEA), a web application that allows for querying, visualization, and analysis of over 4200 published sRNA-seq expression samples. SEA automatically downloads and re-analyzes published data using Oasis 2 (Rahman et al., 2018), semantically annotates relevant meta-information using standardized terms (the annotations are later checked and corrected manually), synchronizes sRNA information with other databases, allows for the querying of terms across ontological graphs, and presents quality curated sRNA expression information as interactive web reports. In addition, SEA stores sRNA differential expression, sRNA based classification, pathogenic sRNA signatures from bacteria and viruses and pathogen differential expression. Gene targets and disease associations for miRNAs are also incorporated into SEA.

One of the most useful features of SEA is to enable users to upload their analysis results of differential expression and classification from Oasis 2. This allow users to compare their data to over 4,200 experimental samples across different conditions. Using SEA’s interactive visualizations, users can upload their data into their own workspace, select the published datasets to compare to, and define if differential expression or classification results should be compared. SEA also provides users with an option to perform on the fly analysis such as overlapping differentially expressed sRNAs or pathogens across different studies or the most important features (sRNAs) identified with classification. Lastly, SEA enables end users to re-submit samples from interactive plots for differential expression or classification, this helps users to choose samples of their choice from an experiment. It currently supports 10 organisms (Table 1) and is continuously updated with novel published sRNA-seq datasets and relevant sRNA information from various online resources. A detailed comparison of SEA to other existing sRNA expression databases (Table 2) highlights that SEA is superior in terms of supported organism, annotations, diseases, tissues, sRNA based classification, pathogen k-mer DE, known miRNA disease associations, user specific experimental data upload, cross study comparisons and re-analysis with selected samples. SEA contains over 4200 samples in its database, which is considerably less than YM500v3 (Chung et al., 2017), which hosts over 8000 cancer samples. It is to be noted, however, that the YM500v3 database only supports cancer datasets and no other disease types (Table 2). Additionally, SEA also stores in-house data (for a month) from the end users to enable comparison with the data in SEA.

**Table 1.**
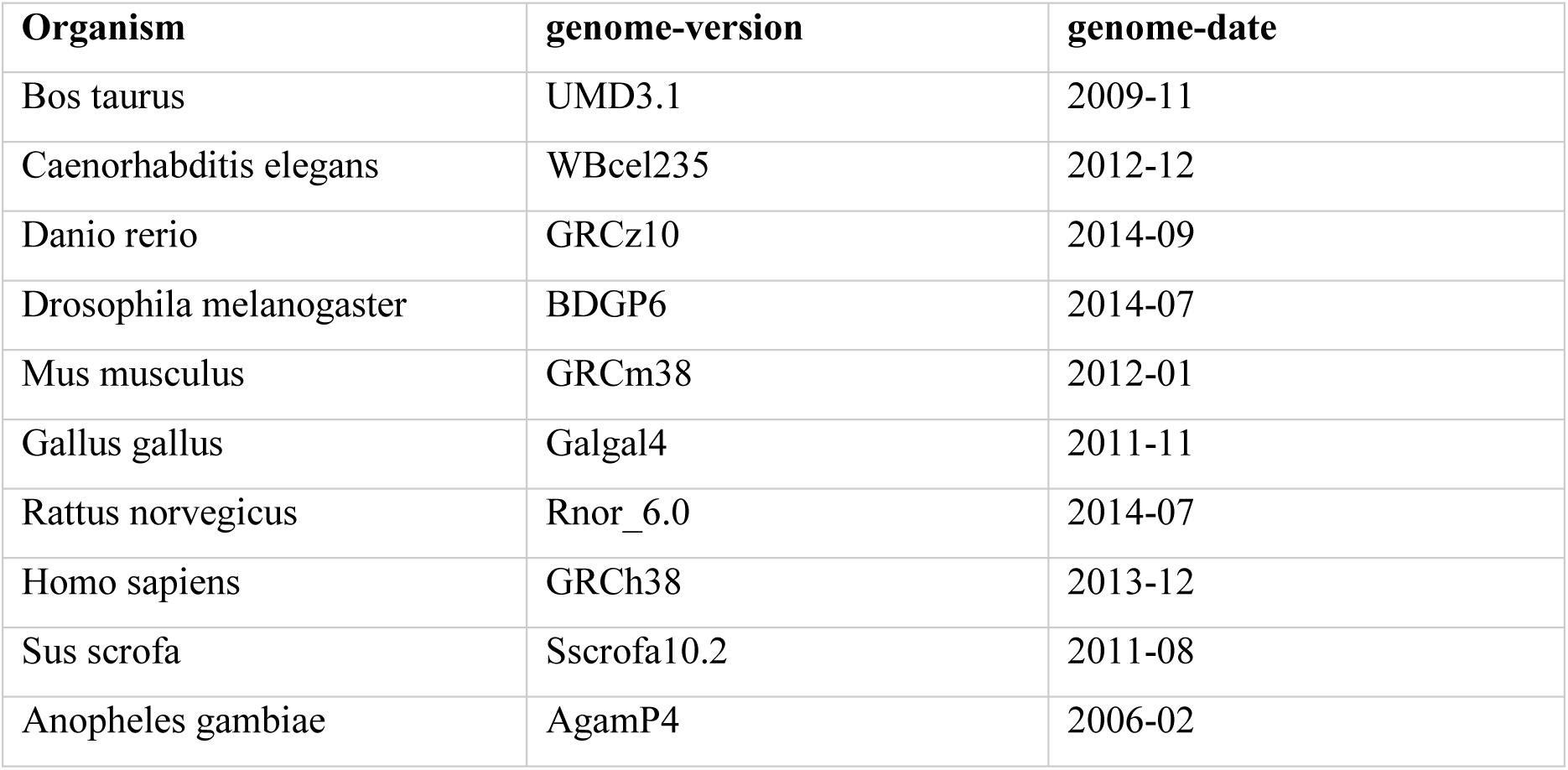
Supported SEA organisms and their corresponding genome versions.

**Table 2.**
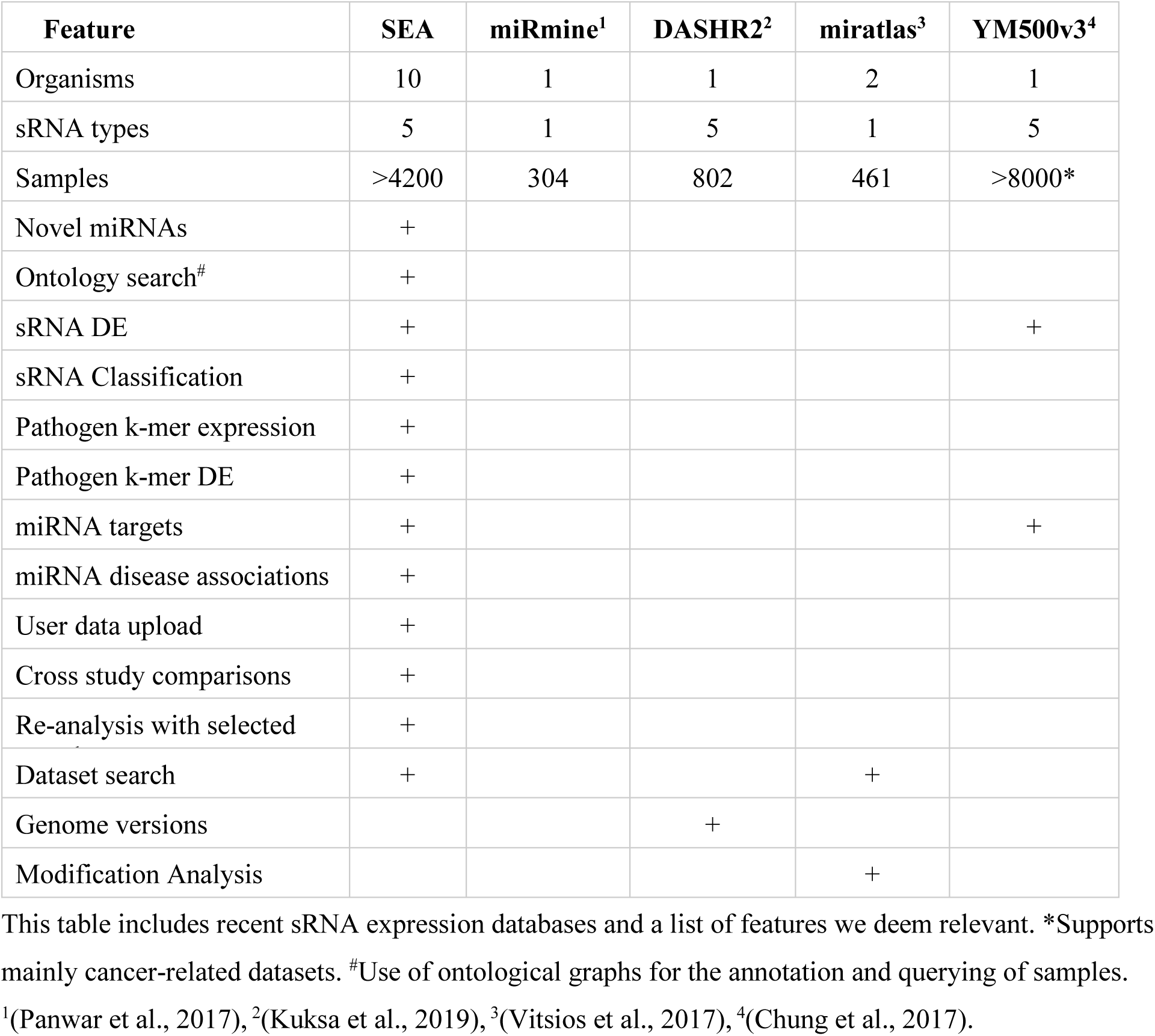
Comparison of sRNA expression databases.

| Feature | SEA | miRmine<sup>1</sup> | DASHR2<sup>2</sup> | miratlas<sup>3</sup> | YM500v3<sup>4</sup> |
| --- | --- | --- | --- | --- | --- |
| Organisms | 10 | 1 | 1 | 2 | 1 |
| sRNA types | 5 | 1 | 5 | 1 | 5 |
| Samples | >4200 | 304 | 802 | 461 | >8000* |
| Novel miRNAs | + |  |  |  |  |
| Ontology search<sup>#</sup> | + |  |  |  |  |
| sRNA DE | + |  |  |  | + |
| sRNA Classification | + |  |  |  |  |
| Pathogen k-mer expression | + |  |  |  |  |
| Pathogen k-mer DE | + |  |  |  |  |
| miRNA targets | + |  |  |  | + |
| miRNA disease associations | + |  |  |  |  |
| User data upload | + |  |  |  |  |
| Cross study comparisons | + |  |  |  |  |
| Re-analysis with selected | + |  |  |  |  |
| Dataset search | + |  |  | + |  |
| Genome versions |  |  | + |  |  |
| Modification Analysis |  |  |  | + |  |

This table includes recent sRNA expression databases and a list of features we deem relevant. \*Supports mainly cancer-related datasets. <sup>#</sup>Use of ontological graphs for the annotation and querying of samples.
<sup>1</sup>(Panwar et al., 2017), <sup>2</sup>(Kuksa et al., 2019), <sup>3</sup>(Vitsios et al., 2017), <sup>4</sup>(Chung et al., 2017).

## 2 System Design

SEA stores sRNA expression information, sRNA differential expression, sRNA based classification, pathogenic sRNA signatures from bacteria and viruses, pathogen differential expression, miRNA gene targets and disease association as well as deep and standardized metadata on the samples, analysis workflows, and databases used. Metadata information is normalized using ontologies to allow for standardized search and retrieval across ontological hierarchies (section ‘Semantic data layer’ & supplementary material). The following sections will detail the system design of SEA (Fig. 1).

**Figure 1.**
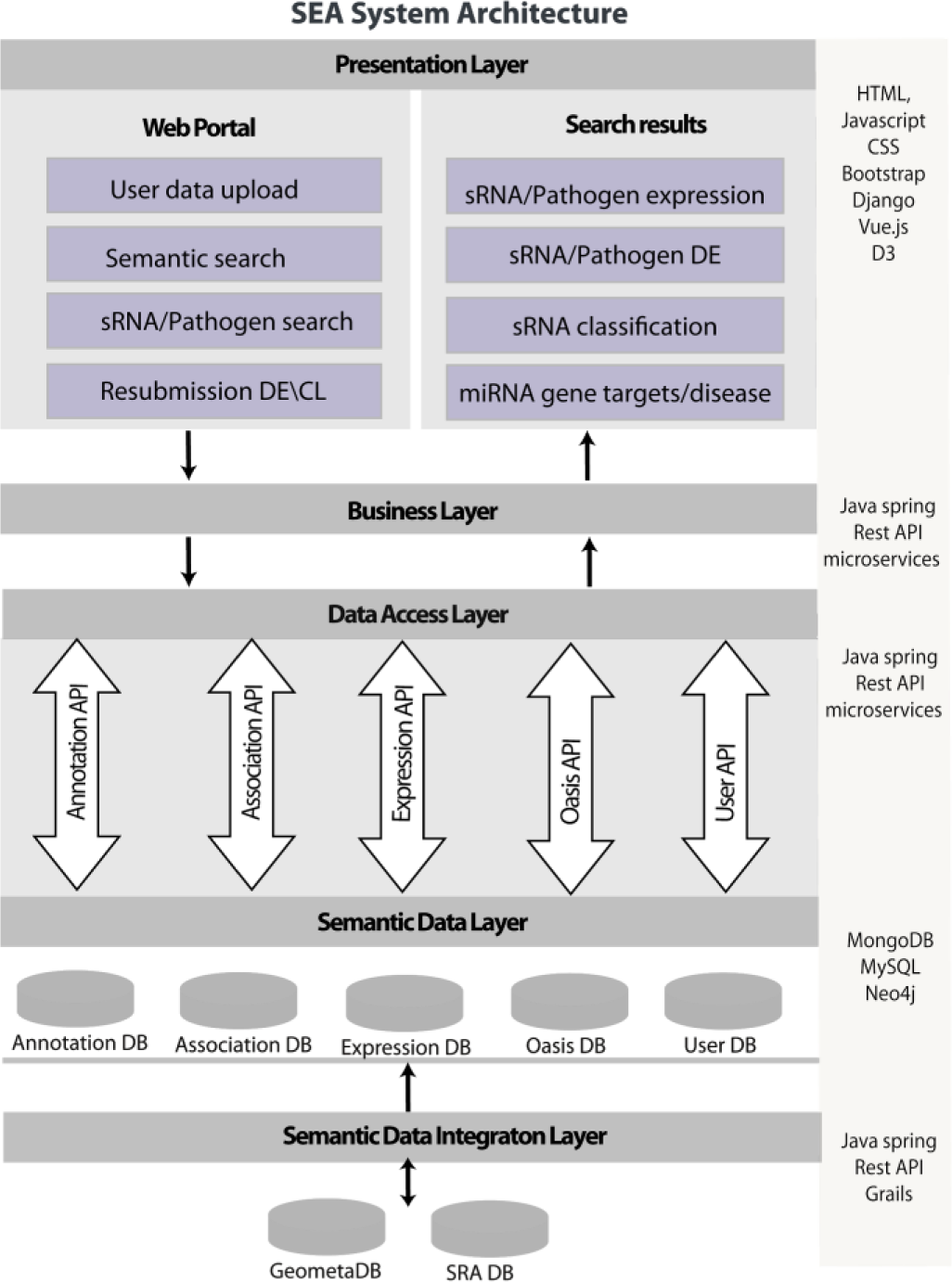
SEA system architecture. SEA system was developed using the modular system design approach (model-view-controller). The system has a **presentation layer** for user interface and visualization of search results. Presentation layer is followed by a **business layer** which transform complex user queries and distribute particular requests to the data access layer REST API services. There is a **semantic data layer**, to store and access primary and derived data together with annotations and links to secondary data. **Annotation-DB** stores metadata for experiments, samples, corresponding ontological terms as well as relations between dataset/sample, sample/term and term/term. **Association-DB** contains information about sRNAs and genes chromosomal locations, miRNA target genes and miRNA disease associations. **Expression-DB** stores sRNA expression profiles, sRNA differential expression; sRNA based classification as well as pathogen detection and pathogen differential expression. It also store details about dataset processing pipeline and parameters. **Oasis-DB** was used to store novel predicted miRNA information. **User DB** contains in-house data uploaded by the end users from Oasis 2 pipeline. **Semantic data integration layer** integrates primary and secondary data into the mentioned databases. Microservices were implemented in order to achieve strong encapsulation and well-defined interfaces via REST APIs.

### 2.1 Acquisition and analysis of sRNA datasets

SEA acquires raw published sRNA-seq datasets and their primary annotation from Gene Expression Omnibus (GEO) (Barrett et al., 2013) and NCBI’s Sequence Reads Archive (SRA) repository (Leinonen et al., 2011) (supplementary material). Novel datasets are downloaded and stored in SEA’s raw data repository while corresponding annotations are stored in SEA’s annotation database and are manually curated. Raw data is subsequently processed automatically by SEA’s sRNA analysis workflow using Oasis 2.0 (Rahman et al., 2018) (http://oasis.ims.bio/) (supplementary material). Subsequently, sRNA counts of high-quality samples are stored in the sRNA expression database. For all the experiments with samples from different conditions such as disease, tissue, cell line or cell type; sRNA differential expression and classification was performed within the experiment using Oasis 2. All possible comparisons for an experiment were taken into account such as healthy vs disease stage 1, healthy vs disease stage 2, disease stage 1 vs disease stage 2 as explained in supplementary section 3.4. Additionally, differential expression analysis of detected pathogens was performed using DESeq2 package (Love et al., 2014). In order to reduce bias that could be introduced into the data by using different analysis routines, every sample in SEA has been analyzed by identical analysis workflows using identical databases and genome versions. SEA additionally stores versioning information about the software and databases used for analysis. In case of changes in databases or analysis routines, we completely re-analyze all SEA’ data for consistency.

Additionally, sample annotations are processed automatically with SEA’s annotation workflow. Processed files and annotations are subsequently semi-automatically curated (section 2.3 and supplementary section 3).

### 2.2 Data storage

Once the raw sequencing data is analyzed, the next step is to store the analysis results to the database for downstream analysis and querying. Most metadata is quite different between experiments. Some experiments may have information such as disease, tissue, cell line, gender, age of patient while others may completely lack this. Due to this sparse nature of the biological experimental data, we opted to use NoSQL database management systems such as MongoDB and Neo4J for hierarchical (connected) normalized data. A multi-database management system architecture was used to store different types of data:

In brief, **Expression-DB** is created to save sRNA expression profiles, sRNA differential expression, sRNA based classification as well as pathogen detection and pathogen differential expression. This database stores the identification and description of the experiment (dataset), information about dataset processing (pipeline information and parameters), information about samples. **Association-DB** is used to store genomic coordinates for sRNAs, miRNA gene targets and miRNA diseases association. It contains information about sRNA’s and gene’s chromosomal locations, miRNA target genes and miRNA disease associations. Chromosomal coordinates were obtained from miRBase version 21 (Kozomara et al., 2014), ensemble version 84 (Zerbino et al., 2018) and piRNA bank (Sai Lakshmi et al., 2008), miRNA gene targets were obtained from mirTarBase version 7.0 (Chou et al., 2018) as well as from BELIEF text mining pipeline (Madan et al., 2016) (material and methods), miRNA disease associations were obtained from HMDD database version 2.0 (Li et al., 2014). In order to enable search by ontological terms, **Annotation-DB** is created using the Neo4J database management system. Neo4J is a graph database, representing elements as graph nodes or vertices. **Annotation-DB** (supplementary Fig. S1) stores the following three node types: (i) Experiments (datasets), this type of node stores information about the experiment such as description of the experiment, reference to database, experimental design, and any global level information, which is common amongst all the samples. (ii) Sample node type is used to store information about individual sample, such as description of a sample, reference to database, sample-specific processing parameters. (iii) Annotation term node type stores annotation term information of samples such as organism, disease, tissue, cell type, cell line, age, gender, condition (treated\untreated) and extracted molecule for sequencing etc. If the annotation term is normalized, it stores ontology reference (term identifier and preferred level). The nodes are connected if they have a relation (dataset/sample, sample/term and term/term) (supplementary Fig. S1). To allow for fast ontological search, all parents of a term in the ontology are also stored in the database and connected with their corresponding annotation terms (section ‘Semantic data layer’ & supplementary material)**. User-DB** stores in-house sRNA-seq data (differential expression and classification from Oasis) uploaded by the users. This database allows users to compare their own data to the huge and diverse sRNA-seq published data.

In addition, SEA contains information about the GEO series accession (GSE) and sample accession (GSM) identifiers along with the sample identifier from the Sequence Read Archive (SRA) database (SRR). We optimize search and retrieval times by indexing for the most common queries and most relevant terms.

### 2.3 Semantic data layer

Given the diversity of the biological data, users of the SEA system are given a possibility to interpret data independently using common terminologies. In order to enable users to browse data autonomously using common well-structured terminology, a standardized semantic layer for data retrieval is developed (Fig. 1). It includes semantic annotations of data and semantic search, linking data with semantic lookup platform (OLS), as well as storing primary and derived data together with provenance information and references to secondary data.

One of the most important aspects of semantic layer are ontology-based data annotations. They enable interoperability of the data, as well as using of standard terminologies for data retrieval. It is important to standardize annotations using ontologies and semantic mappings (Schuurman et al., 2008). Ontologies define not only standard classes, but also the relations between terms, which enables semantic search by term hierarchies, for example, by parent terms. In SEA, we connect (normalize) annotations with ontologies in a semi-automatic way, i.e. first automatically extract possible annotation terms from GEO descriptions and normalize them, and later curate annotations manually (supplementary section 3). The Ontologies and the number of normalized terms in SEA are listed in Table 3. To enable the search across ontological hierarchies we integrated data with the relevant ontologies into the graph database Neo4J (supplementary Fig. S1). Ontology Lookup Service (OLS) is a service which allows to extract relevant terms from ontologies together with term information. SEA uses OLS for annotation normalization and accesses ontologies via the OLS REST interface, which supports complex and compound queries and query auto-completion (Côté et al., 2010). Details about annotation criteria, processing and group annotation are described in supplementary Section 3.2.

**Table 3.** SEA keys and used ontologies (as of June 2019).

| Key | Ontology(s) | # Annotations | # Terms |
| --- | --- | --- | --- |
| Organism | NCBI Taxonomy<sup>1</sup> | 4235 | 126 |
| Tissue | BRENDA tissue / enzyme source<sup>2</sup> | 3021 | 190 |
| Disease | Human Disease Ontology<sup>3</sup> | 1951 | 287 |
| Cell type | Cell Ontology<sup>4</sup> | 732 | 304 |
| Cell line | Cell Line Ontology<sup>5</sup> | 663 | 132 |
|  | Experimental Factor Ontology<sup>6</sup> | 134 | 76 |

Another aspect of the semantic layer is storing of the primary and the derived data together with provenance information. For SEA, primary data are FASTQ files, retrieved from the NCBI SRR database. This data is not stored after Oasis analysis, only provenance data about source and analysis details is saved. So for SEA, primary data are sRNA counts. Based on those counts, DE and classification results are obtained and are also saved to allow data interpretation. From derived data, the provenance information allows to retrieve raw counts and check how those results are obtained.

### 2.4 Querying and visualization

Application programming interfaces (APIs) are developed to access data in SEA databases (supplementary section 3.5). The APIs help to use the multi-database system components independently as well as in combination. In brief, we extend the SEA backend application with RESTful web services, such as Annotation-API, Association-API, Expression-API, User-Expression-API, Predicted miRNA-API to access Annotation-DB, Association-DB, Expression-DB, User-Expression-DB and Oasis-DB respectively. Additionally the SEA business logic API is created in order to combine all those APIs and make necessary data transformations between frontend and other APIs. As a result, the user can make queries to answer biological questions like; what is the expression of hsa-miR-488-5p across all human tissues? Is hsa-miR-488-5p expressed higher in adenocarcinomas as compared to other cancer types? Is a particular sRNA/pathogen differentially expressed in Alzheimer’s disease? What are common differentially expressed sRNAs/pathogens or potential sRNA based biomarkers in a particular disease or tissue? What is the expression of a novel miRNA for known disease states? All API calls are described in supplementary Section 3.5.

In brief, the SEA system is developed using the modular system design approach (Fig. 1). We build micro services to achieve strong encapsulation and well-defined interfaces via REST APIs. An object oriented programming approach is used to build the SEA application using the spring framework and Java 8. The SEA user interface (UI) is developed in Django framework version 2.0, HTML version 5, D3 and CSS 3. SEA visualizes the results depending on the user query, such as a violin plot for the expression of sRNAs or pathogens. Upset plots are shown for the overlap of sRNAs or pathogens (based on DE or classification) across experiments. SEA enables the download of search results in the form of CSV files. The functionality is tested on all major browsers (Table 4).

**Table 4.** SEA browser compatibility.

| Browser | Version |
| --- | --- |
| Chrome | 61.0.3163.100, 62.0.3202.62 |
| Mozilla Firefox | 55.0.3, 56.0 (64-bit), 57.0 (64-bit) |
| Chromium | 62.0.3202.75 |
| Safari | 11.0.1 |
| Internet explorer | 11 |

Browsers that are used to test SEA functionalities.

## 3 Applications of SEA

In this section, we describe a few examples that illustrate how SEA can be employed to answer biological questions and to uncover unappreciated properties of sRNA data integration with interactive result visualization. First, we took advantage of the diverse and massive sRNA-seq data in SEA to present the most comprehensive set of tissue specific miRNAs till date. Second, we utilized the pathogenic reads in sRNA-seq to find their association to diseases. Finally, we show a use case of SEA by comparing an in-house Parkinson’s disease (PD) sRNA-ome to other neurodegenerative diseases sRNA expression profiles available in SEA.

### 1. miRNA tissue specificity

Several studies have shown tissue specificity for miRNAs. Recently, (Ludwig et al., 2016), analyzed several human tissue biopsies of different organs from two individuals to define the distribution of miRNAs using tissue specificity index (TSI) and found several groups of miRNAs with tissue-specific expression. Similarly, (Lee et al., 2008) provides the expression of 201 miRNAs across 9 human tissues to find tissue specificity of miRNAs. miRNAs whose expression is 20-fold or higher in a certain tissue compared with the mean of all the other tissues were characterized as tissue specific. According to (Lee et al., 2008), skeletal muscle, brain, heart, and pancreas are the tissues expressing the most specific miRNAs. Moreover, (Guo et al., 2014) manually extracted 116 tissue specific miRNAs across 12 human tissues. We used Shannon entropy to calculate TSI for each miRNA across tissues in SEA (material and methods). We used very stringent criteria: miRNAs with Shannon entropy score more than 0.8 were considered as tissue specific and less than equal to 0.2 were considered as ubiquitous miRNAs (Fig. 2, Supplementary Table 1). We were able to provide by far the most comprehensive set of 591 distinct tissue specific miRNAs across 30 tissues; skin, liver, testes, blood, semen, prefrontal cortex, peripheral blood, brain, renal cortex, bladder, embryo, colon, placenta, tongue, breast, tonsils, bone marrow, blood plasma, lymph node, heart, lung, neocortex, blood serum, serum, cornea, cerebellum, skeletal muscle, kidney, muscle and thyroid gland (Fig. 2, Supplementary Table 1). In order to compare the TSI for miRNAs in SEA with the existing findings, we merged the list of miRNAs from the above studies and retained all the 12 tissues. Out of 12 tissues, we did not have sequencing data for four of them: thymus, pancreas, spleen and bone.

**Figure 2.**
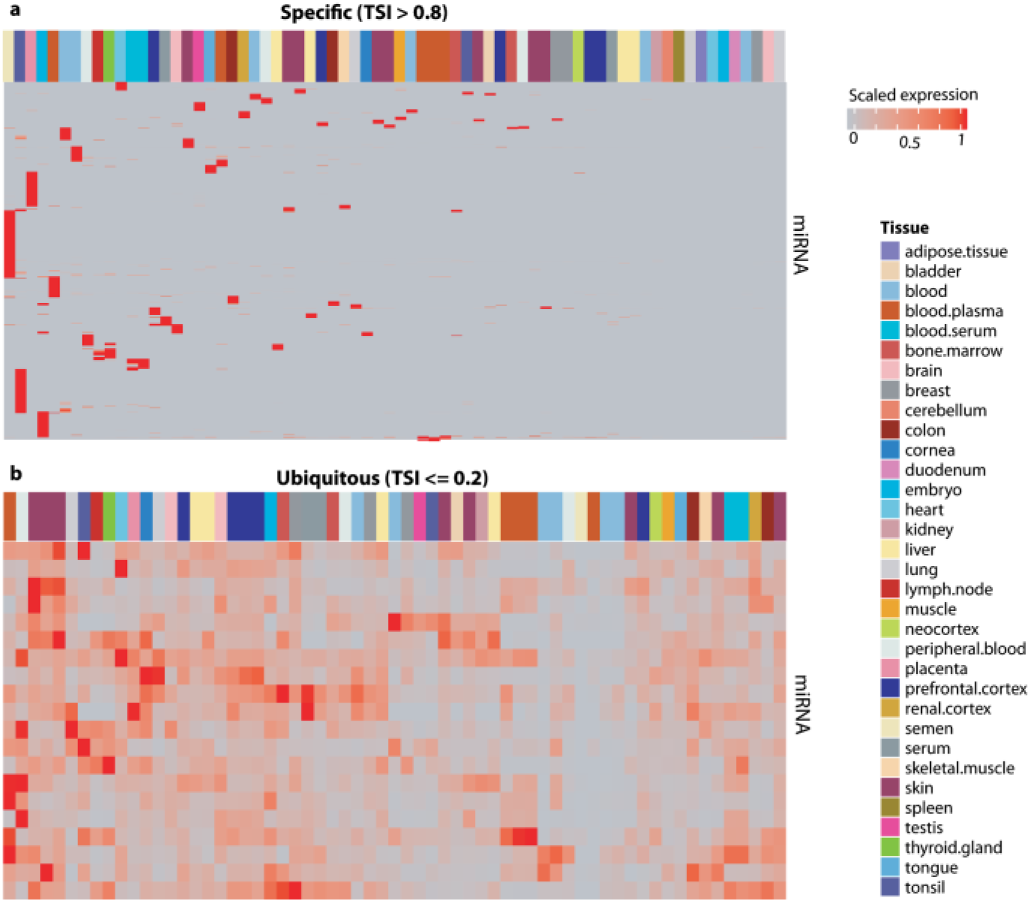
Tissue specific miRNAs. Heatmaps show scaled expression (0-1) for each miRNA across all the tissues. 1 is maximum expression of a miRNA in that tissue compared to all other tissues. (**a) Tissue specific miRNAs**. miRNA across all tissues with TSI > 0.8 (n=591). (**b) Ubiquitous miRNAs.** miRNA expression across all tissues with TSI <= 0.2 (n=20). miRNAs names are omitted for simplicity. A complete list can be found in Supplementary Table 1.

We were able to detect all the three **heart** specific miRNAs (miR-1, miR-133a, miR-302d) from (Lee et al., 2008) study, and 7 out of 10 heart specific miRNAs (hsa-miR-1-5p, hsa-miR-208a-3p, hsa-miR-208b-5p, hsa-miR-208b-3p, hsa-miR-302d-3p, hsa-miR-133b, hsa-miR-302a-3p, hsa-miR-302a-5p, hsa-miR-133a-5p, hsa-miR-302b-3p) from the manually curated list of (Guo et al., 2014). miR-208 is obtained from an old annotation, because the latest release of miRBase has more specific annotation like miR-208a-3p, miR-208b-3/5p. Interestingly we were able to find the whole family of miR-208 as heart specific. We were not able to detect miR-126, miR-302c, miR-367 in heart. Of note, none of these three is heart specific in the (Lee et al., 2008) study.

**Muscle** and **brain** were the only two tissues covered by all the three above mentioned studies. In muscle, we were able to detect all the three muscle specific miRNAs (miR-133b, miR-133a-3p, miR-1-3p) from (Ludwig et al., 2016), 3 out of 4 (miR-95 was not found to be muscle specific) from (Lee et al., 2008), and 6 out of 10 for (Guo et al., 2014) compilation. We were not able to detect miR-134, miR-193a, miR-95 and miR-128a. Note that from the same study miR-134 is mentioned as muscle as well as testis specific and miR-128a as muscle as well as brain specific. Moreover miR-95 is the only miRNA that is muscle specific in all of the three studies.

Another tissue covered by all of the three studies is the brain. In total 30 miRNAs were shown to be brain specific, only 1 out of 30 (miR-7) is common among all the three studies and only three in two studies (miR-124, miR-9, miR-218) one of which is in the curated list. In our study, we found 35 miRNAs brain specific but only two from the known ones (hsa-miR-125b-2-3p, hsa-miR-125b-2-3p).

Tissue with the most number (n= 43) of known specific miRNA was **placenta** provided by (Guo et al., 2014). Interestingly, miRNAs associated with placenta were mostly evolutionary related. We were able to detect these evolutionary related miRNAs to be placenta specific as well. In short, we detected **517**a/b/c, **518**a/b/c/d/e/f, **519**a/b/c/d/e, **520**a/d/e/f/g (not detecting **520**b/c/h). Moreover we were also able to detect miR-371, miR-372, miR-512, miR-522, miR-523, miR-524, miR-525, miR-526b and miR-527. Out of 43, we detected 35 and did not detected miR-377, miR-526a, miR-184, miR-154, miR-381, miR-503, miR-450, and miR-136. We detected only 3 (miR-513c-5p, miR-202-3p, miR-34c-5p) out of 15 for **testes**. There were two tissues, **lung** and **liver**; mentioned only in one study (Guo et al., 2014), we could not detect the only miRNA miR-126 for lung. Interestingly this miRNA is also mentioned as heart specific in the same study. We also did not find the four liver specific miRNAs miR-122, miR-483, miR-92a, miR-192; two (miR-483, miR-92a) of which are shown as bone specific in the same study. In **kidney** we were able to detect only 1 miR-200a out of 8 kidney specific in (Guo et al., 2014). Of note (Lee et al., 2008), also found only one miRNA miR-204 to be kidney specific and does not have any evidence for the rest of the seven miRNAs.

As (Ludwig et al., 2016) used only two individual’s tissues, (Lee et al., 2008) also performed own experiments in a control (same laboratory, same protocols) environment and used different statistical methods compared to ours, we were still able to get a reasonable overlap with tissue specific miRNAs considering diverse (different laboratories, different protocols) and massive data. Therefore, we think that this work provides the most comprehensive set of tissue specific miRNAs till date (n = 591 miRNAs) (Supplementary Table 1).

### 2. Known and novel bacterial or viral infections

We validated our approach of pathogen detection using seven datasets with known infection status. The samples in these datasets are known to be infected with seven bacterial pathogens and three viral pathogens. Of note, we focused on within-dataset comparison in order to avoid technical confounders (Supplementary Table 2). For each sample, k-mer counts were calculated for all infectious species present in Kraken database (4336 viral and 2784 bacterial/archaeal genomes) and differential abundance analysis was carried out for those species that have at least 3 counts (baseMean) in a particular comparison. As expected, in all comparisons the known pathogen represented the best hit (i.e., smallest adjusted p-value) except Vaccinia virus (Fig. 3a). However, Vaccinia virus has the highest log2 fold change as expected within the dataset (GSE54235) comparison. It is worthy to note that Chlamydia trachomatis detection is based on sRNA-seq performed on conjunctival tissue from children with follicular trachoma and children with healthy conjunctivae, indicating a good performance of our pathogen detection pipeline from tissues.

**Figure 3.**
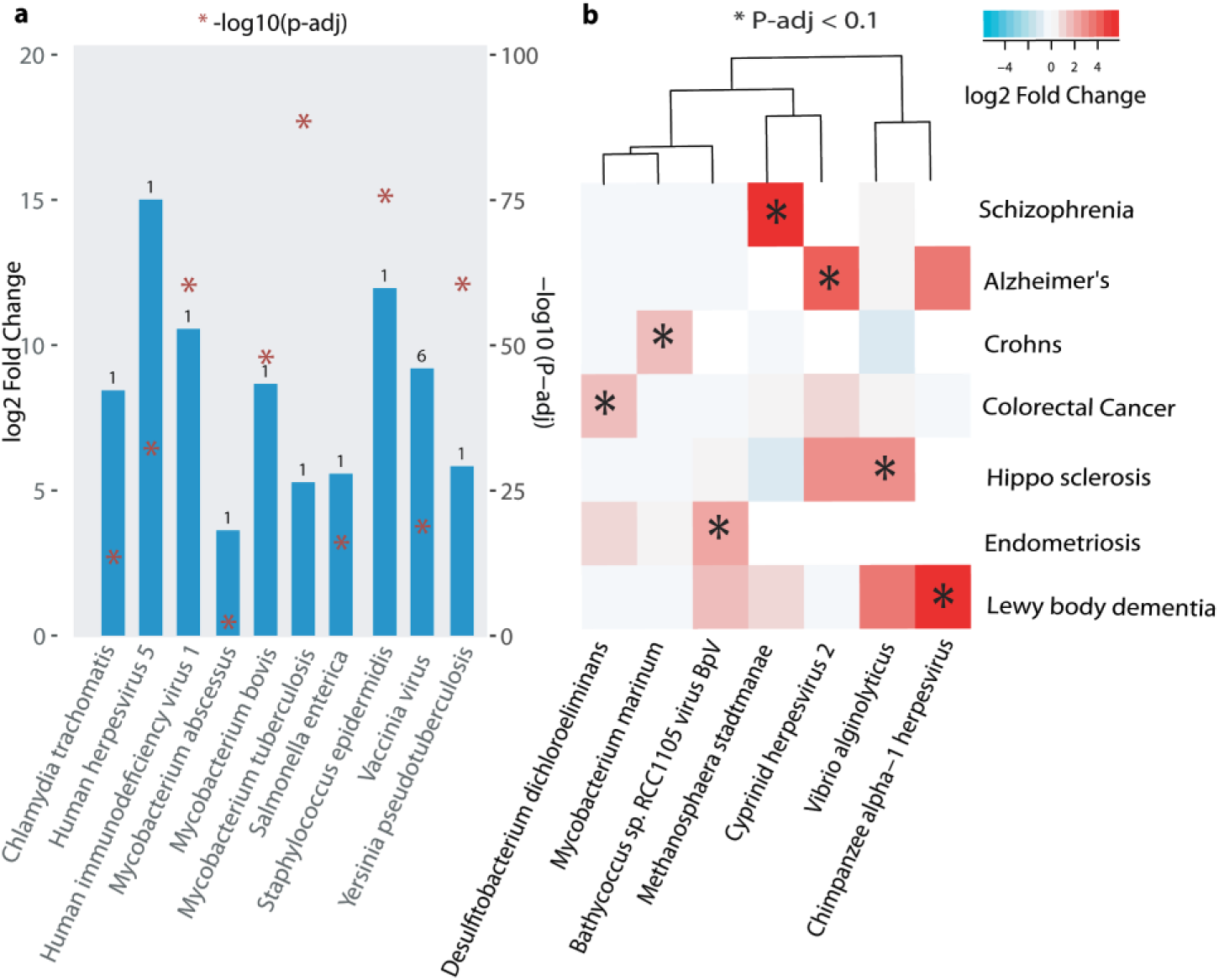
Known and Novel bacterial or viral infections. **(a) Known associations.** Pathogen detection using seven datasets known to be infected with seven bacterial and three viral pathogens. Bar represents pathogen log2fold difference between the uninfected and infected state (Supplementary Table 2). Number on top of the bar denotes rank of the pathogen compared to all the other DE pathogens within the comparison (i.e., smallest adjusted p-value). **(b) Novel associations.** Heatmap shows log2fold difference of pathogens significantly upregulated in disease as compared to healthy (fold change > 1 and padj < 0.1) (Supplementary Table 2). Comparisons that have less than 6 pathogens significantly differentially expressed are selected for specificity. Details about dataset, comparison groups, log2fold and padj for both (a,b) are provided in (Supplementary Table 2).

Next, we aimed to find novel associations of pathogens with disease. We took all the comparisons, which has “healthy” and at least a disease state annotation (Supplementary Table 2). In order to achieve more specificity we took only comparisons that have less than 6 pathogens significantly up-regulated in disease as compared to healthy (FC > 1 and padj < 0.1). There were a total of 8 comparisons but we removed “GSE69837” as this was a known case (Chlamydia trachomatis already shown in Fig. 3a). It was interesting to find viruses and bacteria significantly upregulated in sRNA-seq data in certain disease compared to healthy patients (Fig. 3b). Some of the most interesting cases are highlighted in this section below.

#### Mycobacterium marinum in patients with ileal Crohn’s disease

In the original study, expression of microRNAs in mucosae of patients with a normal pouch after colectomy for intractable ulcerative colitis was compared to several control cohorts, among them was a cohort of patients with Crohn’s disease (CD) of the terminal ileum (Ben-Shachar et al., 2016). CD patients were previously not exposed to immunosuppression. Compared to patients with non-inflamed ileal pouch, patients with ileal CD showed an increased mucosal expression of *Mycobacterium marinum*. The bacterial genus *Mycobacterium* causes diverse diseases in humans, of which Tuberculosis is the most serious with around one-quarter of the world population latently infected and approximately 1.6 million deaths in 2017 on a global scale. *M. marinum* is a non-tuberculous (also termed “atypical”) *Mycobacterium* species, which is ubiquitously abundant in aquatic environments (Johnson et al., 2015). Infection of humans is well known, but it is considered a rare event. It typically occurs after exposure to contaminated water or infected marine animals, and it is more common in immunosuppressed individuals. The most commonly affected organ is the skin, in more severe cases involvement of muscles, bones or joints is reported (Johnson et al., 2015). Opportunistic infection with *M. marinum* in CD is recognized in those patients receiving anti-tumour necrosis factor therapy (e. g. infliximab) (Ferreira et al., 2012). However, to the best of our knowledge, enteric super-infection with *M. marinum* has not been reported in the literature so far. Interestingly, due to the resemblance of the granulomatous intestinal inflammation in CD with enteric infection caused by other Mycobacteria, it has been hypothesized that Mycobacterial infection is involved in the pathogenesis of CD, with much focus on *M. avium paratuberculosis* (McMullen et al., 2015). However, the aetiological significance of this pathogen in CD remains uncertain. Hence, the gut mucosal prevalence of *M. marinum* and its potential pathophysiologic significance in patients with CD should be further explored.

#### Methanosphaera stadtmanae in patients with schizophrenia

We detected an overabundance of *Methanosphaera stadtmanae* in neurons derived from induced pluripotent stem cells (iPSC) of patients with schizophrenia, compared to healthy controls. *M. stadtmanae* is an Archaeal microorganism which is frequently detected in the healthy human gut microbiota (Dridi et al., 2009). It is involved in intestinal methanogenesis and associated fermentative dynamics. *M. stadtmanae* is recognised by the innate immune system, therefore it can induce inflammatory cytokine responses and could have diverse immunomodulatory functions (Bang et al., 2014). Interestingly, *M. stadtmanae* was found with an increased prevalence in faecal samples of patients with inflammatory bowel diseases (IBD) Crohn’s disease (CD) and ulcerative colitis with antigen-specific IgG-responses (Blais Lecours et al., 2014). Immune system processes have been proposed to be involved in the pathogenesis of schizophrenia (Pouget, 2018). Regarding the immune-genetic basis of schizophrenia, genome-wide pleiotropy has been reported between schizophrenia and CD as well as an increased prevalence of schizophrenia in patients with IBD (Bernstein et al., 2019). Therefore, the potential immunogenic importance of *M. stadtmanae* in schizophrenia should be investigated.

#### Chimpanzee herpesvirus in Lewy body dementia

We detected an increased abundance of a viral pathogen identified as *chimpanzee herpesvirus* (ChHV) in the cerebral cortex of patients with lewy body dementia (LBD) compared to non-demented controls (Hébert et al., 2013). ChHV is an alphaherpesvirus closely related to human herpes simplex virus type 2 (HSV-2) (Severini et al., 2013). LBD is a neurodegenerative disorder, which underlies 4.2% of all dementia cases, second only to Alzheimer’s dementia (AD) (Vann Jones et al., 2014). The aetiology of LBD is obscure, but growing evidence points towards neuro inflammation as a key pathophysiologic factor, analogous to the pathogenesis of AD (Surendranathan et al., 2015). In AD it is assumed that multiple pathogens infecting the brain are key triggers of neural dysfunctional protein accumulation and neuro inflammation in genetically vulnerable individuals (Harris et al., 2015). Among the pathogens detected in brains of AD patients, multiple lines of evidence point at herpes simplex virus type 1 (HSV-1) and HSV-2 as two of the main drivers of AD neurodegeneration (Harris et al., 2015; Sochocka et al., 2017). Given the close phylogenetic relationship between ChHV and HSV-2, ChHV might play a role in inflammatory neurodegenerative processes in LBD similar to the other herpesviruses in AD. Therefore, the association detected in the present study should be further elaborated.

### 3. Analyzing in-house data and comparing with SEA data

One of the key features available in SEA is uploading the in-house data and comparing it with the already integrated data. Mostly, researchers use different analysis pipelines to carry out differential expression or classification, which makes it very hard to compare the results with the publicly available data. Therefore, we require a database with interactive visualizations that has all the publicly available data analyzed using the same pipeline with same parameters. For SEA, we have analyzed and integrated all the data using Oasis 2 pipeline. We expect that comparing the in-house data with the data in SEA will yield disease-specific signatures, in this case a sRNA or group of sRNAs. Note that uploading to SEA requires the output of Oasis 2 (supplementary material).

In order to test this feature, we uploaded in-house sRNA-seq data from well characterized 47 Parkinson disease (PD) and 53 frequency-matched healthy controls, which is a baseline data from the longitudinal de novo Parkinson disease (DeNoPa) cohort (Supplementary Table 3) and available as “demo user data” in SEA. SEA gives us a unique opportunity to identify PD-specific biomarkers associated with early-stage PD that can eventually help us in early diagnosis, therefore, better treatment of the disease. Below we describe the differential expression and classification results from PD data and an approach in order to identify PD-specific biomarkers that do not overlap with other neurodegenerative diseases.

We found four significantly differentially expressed (DE) miRNAs with adjusted p-value less than 0.1. Out of these, two are up-regulated in PD (hsa-miR-502-3p and hsa-miR-532-5p) and two are down-regulated in PD (hsa-miR-30d-5p and hsa-miR-22-5p) (Supplementary Table 3). Next, we overlapped these four DE miRNAs with all the neurodegenerative disease-related datasets integrated in SEA. We focused on nine comparisons (from five datasets) in which one of the conditions is a healthy state and the other is a diseased condition (Alzheimer’s disease (AD), Lewy body dementia, tangle-predominant dementia, Huntington’s disease (HD), Frontotemporal dementia or Hippocampal sclerosis of aging). Out of the two up-regulated miRNAs in PD, one (hsa-miR-502-3p) is up-regulated in Alzheimer’s disease and one (hsa-miR-532-5p) is up-regulated in both Alzheimer’s and Huntington’s disease (Fig. 4a). In contrast, none of the down-regulated miRNAs in PD were found to be significantly down in any of these nine comparisons. Interestingly, it has been shown that the expression of miR-22 is down-regulated in a 6-hydroxydopamine-induced cell model of PD using RT-PCR (Yang et al., 2016). Moreover, Margis et al., found that hsa-miR-22 has reduced expression in the blood of de novo PD patients (Margis et al., 2011). Furthermore, family members of hsa-miR-30d-5p are known to be deregulated in PD (Leggio et al., 2017) and putatively target the PD-related gene, **LRRK2 (PARK8)** (Heman-Ackah et al., 2013). These results confirms, the potential role of hsa-miR-30d-5p and hsa-miR-22-5p in PD. To explore the mechanism by which these two miRNA are involved in PD, we performed gene ontology (GO) analysis of the validated and predicted targets using webgestalt (Wang et al., 2017). The top ten terms ranked according to FDR adjusted p-value are shown in the (Fig. 4b). The top significant hit (FDR < 0.1) is axon development. Recent publications (Bolam et al., 2012; Pissadaki et al., 2013; Surmeier et al., 2017) have suggested the role of massive and unmyelinated axonal arbor in PD. In substantia nigra pars compacta (SNc), the axonal arbor of dopamine neurons is very large as compared to other neuronal types. This leads to the hypothesis that these dopamine neurons have selective and exceptional vulnerability in PD, and have a higher energy demand that may play a crucial role in cell death (Pissadaki et al., 2013).

**Figure 4.**
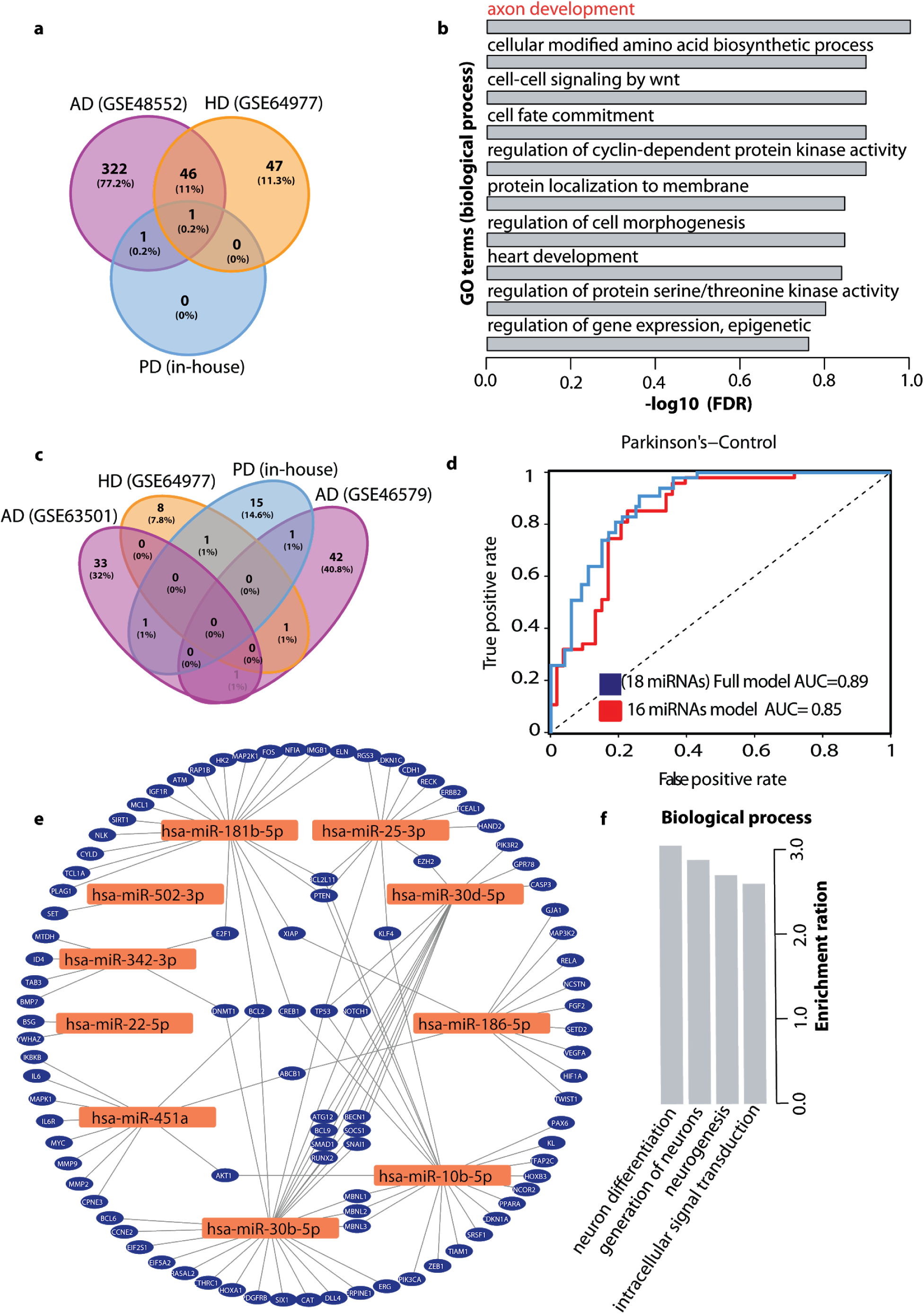
In-house de novo Parkinson disease (DeNoPa). **(a) sRNA DE Overlap.** Overlap of upregulated sRNAs between in-house denopa (blue), AD (purple) and HD (orange). Overall nine neurodegenerative disease comparisons were considered and overlap was found with these two datasets. **(b) Gene Ontology (GO) terms.** Top 10 GO terms associated with the target genes of the two down regulated sRNAs. **(c) sRNA classification Overlap.** Overlap of classification features (sRNAs) between in-house denopa (blue), AD (two datasets) (purple) and HD (orange). **(d) DeNoPa classification.** Receiver-operating characteristic (ROC) curve showing true- and false-positive rates for **DeNoPa** disease prediction based on sRNA expression profile using 18 sRNAs in full model (blue) and 16 unique (not found in other neurodegenerative diseases) sRNAs (red). **(e) PD associated genes.** Network of PD associated genes and 13 known miRNAs from the classification. **(f) GO terms for novel miRNAs.** GO terms associated with the target genes of the three novel miRNAs from the classification.

To further gain insights of unique PD biomarkers, we explored the classification results integrated in SEA. PD and healthy were classified with an AUC of 0.89 (Fig. 4d). Interestingly, the classifier used only 18 sRNAs to separate the two states (Supplementary Table 3). We overlapped these 18 sRNAs with the classification results from other neurodegenerative diseases integrated in SEA (Fig. 4c). There are only three sRNA that are also found in AD or HD but they have opposite change of expression. This suggests the specificity of these sRNAs to PD as compared to other neurodegenerative diseases. Furthermore, to filter out sRNAs known to be associated with other neurodegenerative diseases, we used the association database of sRNA-disease association available in SEA. The results showed that hsa-miR-342-3p has been associated with other neurodegenerative diseases (Montag et al., 2009; Saba et al., 2008). Next, we also filtered out sRNAs if the base mean read count is less than 5 and also, hsa-miR-502-3p that was found to be up-regulated in AD (Fig. 4a). Then we run a random forest classifier using the normalized counts for the remaining 15 sRNAs and hsa-miR-22-5p that is down-regulated in our data. (Fig. 4d) shows that using 16 sRNAs to classify PD and controls, yielded 85% area under the curve (AUC) with 83% recall and 77% of precision. Furthermore, to find the relevance of the 13 known miRNAs (out of 16 sRNAs) in PD, we obtained their target genes from SEA (only 10 miRNAs out of 13 have targets supported by strong evidence) and overlapped with the targets genes of PD associated miRNAs in SEA. Interestingly, these 10 known miRNAs targets 96 genes, which are known to be associated with PD (Fig. 4e, Supplementary Table 3). The list includes **TP53** (Alves da Costa et al., 2011) that contributes to the apoptotic deterioration taking place in PD, **PTEN** (Ogino et al., 2016) that has been linked to PD via DNA damage and DNA repair machinery, **SMAD1** (Hegarty et al., 2018) is an important regulator required for neurite growth, **EZH2** (Södersten et al., 2014) is a lysine methyltransferase component of polycomb repressive complex 2 that has been associated with PD and **BCL2** (van der Heide et al., 2013) is required for proper development of the dopaminergic system and has been implicated in the pathogenesis of PD. To gain further insights into the 3 novel predicted miRNAs (out of 16 sRNAs) used to classify PD and controls, we performed gene enrichment analysis on their target genes using webgestalt (Zhang et al., 2005). The novel miRNAs were p-hsa-miR-113, p-hsa-miR-247, and p-hsa-miR-235-**1/2/3** (supplementary material). We used miRDB (Wong et al., 2015) to get target genes for the mature sequences of these predicted miRNAs (material and methods). Interestingly the GO terms for these miRNAs were **neuron differentiation, generation of neurons**, **neurogenesis** and **regulation of intracellular signal transduction** (Fig. 4f). All these processes are highly related to PD, and hence we think these novel miRNAs should further be explored and validated in the laboratory. Predicted structure of these miRNAs can be found in supplementary material.

All together, these results make a strong case in favor of using SEA in order to retrieve disease-specific biomarkers.

## Conclusions

SEA is designed for the biological or medical end-user that is interested to define where and when a sRNA of interest is expressed. Prototypical questions that can be addressed with SEA are: What is the expression of hsa-miR-488-5p across all human tissues? Is hsa-miR-488-5p expressed higher in adenocarcinomas as compared to other cancer types? Is the tissue-specific expression of hsa-miR-488-5p conserved in mice? Its unique selling points are the deep and standardized annotation of meta-information, the re-analysis of published data with Oasis 2 to reduce analysis bias, a user-friendly search interface that supports complex queries, and the fast and interactive visualization of analysis results across 10 organisms (Table 1) and various sRNA-species. SEA also contains information on the expression of currently 769 high-quality predicted miRNAs, across organisms and tissues.

In addition, SEA also stores sRNA differential expression, sRNA based classification, pathogenic sRNA signatures from bacteria and viruses and pathogen differential expression. Furthermore, SEA can be used to search gene targets or diseases associated with a miRNA. Moreover, SEA allows end users to upload their analysis results of differential expression and classification from Oasis 2. This will allow users to compare their data to over 4200 experimental samples across different conditions. SEA also provides users with an option to perform on the fly analysis such as overlapping differentially expressed sRNAs or pathogens across different studies or the most important features (sRNAs) identified with classification. SEA enables end users to re-submit samples from interactive plots for differential expression or classification, this will help users to choose samples of their choice from an experiment.

Moreover, SEA is continuously growing and aims to eventually encompass all sRNA-seq datasets across all organisms deposited in GEO and other repositories. Genome versions will be updated with every major release of SEA. SEA will be backwards compatible in the future by allowing users to choose previous genome versions and annotations. A detailed comparison of SEA to other existing sRNA expression databases highlights that SEA is superior in terms of supported organism, annotations, diseases, tissues, sRNA based classification, pathogen k-mer DE, known miRNA disease associations, user specific experimental data upload, cross study comparisons and re-analysis with selected samples (Table 2).

As far as we are aware, SEA is the only sRNA-seq database that supports ontology-based queries, supporting single or combined searches for five predefined keys (organism, tissue, disease, cell type, and cell line) across all datasets. However, the SEA database system contains additional (meta)-information including age, gender, developmental stage, genotype as well as technical experimental details such as the sequencing instrument and protocol details (e.g., library kit, RNA extraction procedure). We plan to normalize most of this additional information in future versions of SEA. This will allow users, for example, to query and analyze sRNA expression effects that are introduced by library kit or sequencing platform differences (both of these features can introduce large biases in the detection and expression of sRNAs). Other future developments will include information on sRNA editing, modifications, and mutation events.

In summary, SEA supports interactive result visualization on all levels, from querying and displaying of sRNA expression information to the mapping and quality information for each of the over 4200 samples. SEA is a fast, flexible, and fully interactive web application for the investigation of sRNA and pathogen expression across cell lines, tissues, diseases, organisms, and sRNA-species. As such, SEA should be a valuable addition to the landscape of sRNA expression databases.

Additionally, we presented the most comprehensive set of tissue specific miRNAs till date. We were able to provide by far the most complete set of 591 distinct tissue specific miRNAs across 30 tissues. To our knowledge this is by far the most comprehensive analysis (set) of tissue specific miRNAs.

In the current work, we also found pathogen signatures from sRNA-seq data. We found signatures of pathogens in severe diseases like dementia. In brief, we found differential regulation of mycobacterium marinum in patients with ileal crohn’s disease, methanosphaera stadtmanae in patients with schizophrenia and chimpanzee herpesvirus in Lewy body dementia.

From our in house Parkinson’s disease data, we were able to find potential biomarkers based on differential expression and classification for the early detection of PD. The top term for the gene ontology (GO) analysis of the two down-regulated miRNAs is axon development, suggesting their role in PD. Moreover, gene targets of the sRNAs for the top important features (potential biomarkers) for PD using classification were overlapping with the targets of the known PD miRNAs. Additionally, GO analysis for the targets of the three novel miRNAs are neuron differentiation, generation of neurons, neurogenesis and regulation of intracellular signal transduction (Fig. 4f). We think these novel miRNAs should be further explored and validated in the laboratory.

Lastly, researchers have used massive sRNA data from SEA for other tasks, for example, it enables to use deep learning for data augmentation problem such as predicting sex and tissue based on sRNA expression profiles (Fiosina et al., 2019). As such, SEA should be a valuable addition to the landscape of sRNA-seq web applications.

## Material and methods

### miRNA tissue specificity

To compute tissue specificity index (TSI) for all the miRNAs in SEA, we calculated median of reads per million (RPM) for each miRNAs within a dataset having the same tissue. Healthy and diseased samples were mixed for tissues within the same dataset. Shannon entropy from BioQC R package was used to calculate TSI for each miRNA across tissues.

### Novel miRNA gene targets

miRDB (Wong et al., 2015) was used to obtain targets of the novel miRNAs. We restricted the analysis to highly probable gene targets having a score of 70 or more.

### Text mining pipeline

To extract miRNA-gene targets, a dedicated text mining pipeline that reads unstructured text data and outputs structured data that includes the detected and normalized genes and miRNAs as well as the relations between them. Named entity recognition software ProMiner (Hanisch et al., 2005, Fluck et al. 2007) and MiRNADetector (Bagewadi et al. 2015) are used to detect and normalize genes and miRNAs, respectively. Both detectors are incorporated in the BELIEF text mining pipeline (Madan et al., 2016) that contains machine learning models to detect specific relations from the complete Medline abstracts.

### Gene enrichment analysis

Gene enrichment analysis was performed using webgestalt R package version 0.3.0.

### In house parkinson disease data

#### Isolation of total RNA from peripheral blood sample

Peripheral blood samples were collected into PAXgene Blood RNA tube (PreAnalytiX) from consenting patients and healthy controls, the tubes were gently inverted for multiple times, incubated for 20-24 hours under room temperature and stored under −80 °C until processing. Total RNA was isolated using the PAXgene Blood RNA kit (PreAnalytiX) according to the manufacturer’s protocol. The purity and concentration of isolated RNA were measured with NanoDrop™ 2000 spectrophotometer (Thermo Fisher Scientific). The RNA integrity was determined by Agilent RNA 6000 Nanochip (Agilent Technologies) using the 2100 Bioanalyzer (Agilent Technologies).

#### small RNA library preparation

Small RNA libraries were prepared using 1 ug high-quality RNA following the protocol of Illumina TrueSeq small RNA library kit (Illumina). In brief, 3’adapter was denatured for 2 minutes under 70°C, and ligated to the RNA with T4 RNA Ligase 2 deletion mutant for 1 hour at 28°C. Then the reaction was stopped with stop solution for 15 minutes under 28°C. Subsequently, 5’ adapter was denatured for 2 minutes at 70°C, then added to the RNA with ATP and T4 DNA ligase for 1 hour under 28°C. After adaptors ligation, the RNA was reverse transcribed to complement DNA (cDNA) by using SuperScript II Reverse Transcriptase (Thermo Fisher Scientific) and dNTPs for 1 hour at 50°C. Then, the cDNA was indexed and amplified with PCR mix and primers supplied in the kit for 12 cycles (denaturing at 98°C for 30 s, annealing at 60°C for 30 s, extension at 72°C for 15 s, with a final extension at 72°C for 10 min). Amplified and indexed cDNAs were then pooled together, mixed with DNA loading dye and loaded on a 5% TBE acrylamide gels (Bio-Rad). After 57 minutes electrophoresis under 145 V, the gel was stained with Midori Green for 5 minutes and viewed under the UV transilluminator, fragments between Illumina’s custom ladder 145 bp to 160 bp were cut out for library preparation. The gel was centrifuged at 20,000 x g for 2 minutes through a Gel Breaker tube (Bio-Cat). Then cDNA was eluted from the homogenized gel by adding 300ul UltraPure water and shaking under 800 x rpm for 2 hours. Then the gel was transferred on a 5 um filter tube (Bio-Cat) and centrifuged for 10 seconds under 600 x g and the gel debris was separated. Afterward, 2ul Glycoblue, 30ul of 3M sodium acetate and 975ul 100% ethanol (pre-chilled under −20°C) were added and well mixed to the sample, following an immediate centrifuge at 20,000 x g for 20 minutes under 4°C. After remove and discard the supernatant, the pellet was washed with 500ul 70% pre-chilled ethanol. The supernatant was discarded after sample being centrifuged at 20,000 x g for 2 minutes under room temperature, and the pellet was dried in a 37°C heat block for 10 minutes with open lid. Finally, the pellet was resuspended in 10ul 10mM Tris-HCL (pH 8.5) and the sample quality was checked using Agilent High Sensitivity DNA chip (Agilent Technologies) using the 2100 Bioanalyzer (Agilent Technologies). All high quality libraries were then sequenced on Illumina HiSeq 2000 Sequencer.

## Supporting information

p-hsa-miR-113

p-hsa-miR-247

p-hsa-miR-235-1

p-hsa-miR-235-2

p-hsa-miR-235-3

## List of abbreviations

SRA: Sequence Read Archive
GEO: Gene Expression Omnibus
sRNA: Small RNA
miRNA: MicroRNA
DB: Database

## Declarations

### Acknowledgements

We would like to thank Mariah Snyder, Yu Zhao, the ZMNH IT, and all of the SEA users for helpful suggestions.

### Funding

This work was supported by the DFG (BO4224/4–1), the Network of Centres of Excellence in Neurodegeneration (CoEN) initiative, the Volkswagen Stiftung (Az88705), iMed – the Helmholtz Initiative on Personalized Medicine, and the BMBF grant Integrative Data Semantics in Neurodegeneration (031L0029B, IDSN).

### Availability of data and materials

SEA is freely available at http://sea.ims.bio/.

### Authors’ contributions

**SB** initiated the study and designed the web application as well as analyses together with **RR** and **VB**. **RR**, **AS**, **AL** designed and implemented the expression database. **RR** and **MF** designed and implemented association database. **MF**, **SM**, and **JF** designed and developed the semantic integration service. **RR**, **AS, AL**, **MF** implemented the APIs. **RR** and **AG** implemented the pipeline to automatically download and submit sRNA-seq data to Oasis 2. **RR** and **AG** implemented the predicted miRNA API. **SH** designed the development and deployment system infrastructures. **TS** annotated the experiments and samples. **DSM** and **AL** developed the interactive user interface. **AL, JB, VB** and **RR** made the user manual and tutorials. **VB**, **RR, AL** and **TL** analyzed the sRNA-seq data mentioned in the manuscript. **AR, CT** and **BM** provided and sequenced the Denopa sRNA samples. **SB**, **RR, VB** and **TL** wrote the manuscript. All authors read and approved the final manuscript.

### Ethics approval and consent to participate

NA

### Consent for publication

N/A

### Competing interests

The authors declare that they have no competing interests.

## Supplementary material

### 1. Acquisition and analysis of sRNA datasets

SEA acquires raw published sRNA-seq datasets and their primary annotation from Gene Expression Omnibus (GEO) and NCBI’s Sequence Reads Archive (SRA) repository. GEO makes two databases in SQLite format available for download: GEOmetadb for annotations and SRAdb for SRA sequences. An automated data acquisition pipeline searches for new sRNA data bi-weekly, keeping SEA continuously updated. Following the acquisition of sRNA datasets, the SEA analysis workflow automatically analyzes new files using the Oasis 2.0 (http://oasis.ims.bio/). The SEA analysis workflow determines data quality and detects and quantifies sRNAs, including the prediction of novel, high-quality miRNAs. Low quality files are flagged automatically and subjected to manual curation. Any files not passing manual curation are removed from SEA. Subsequently, sRNA counts of high-quality samples are stored in the sRNA expression database while corresponding quality information is saved in the data quality repository. Pathogenic signatures from bacteria and viruses, supplying information on potentially infected samples is also incorporated into SEA. Additionally, SEA stores expression information of high-quality predicted miRNAs including the ID, organism, chromosomal location, precursor and mature sequences, structure, read counts, prediction scores, and detailed information on the software and its versions used to predict the miRNA. SEA’s primary analysis results including per sample quality and expression information can be examined and downloaded as interactive web reports. Detailed information on the primary analysis of sRNAs and predicted miRNAs can be found in the Oasis 2 manuscript (Rahman et al.).

### 2. Differential expression and Classification

sRNA data differential expression (DE) and classification are obtained by passing obtained sRNA counts to DE and classification modules of Oasis. As biological conditions for comparisons, group annotations (section 2.3) are used. This means that several comparisons inside of one dataset are possible. The results of DE analysis are mean value, fold change, p value and p adjusted value for each sRNA. The results of classification analysis are Gini index decrease for each sRNA.

Those results show importance of the corresponding sRNA in distinguishing between two biological conditions. SEA stores above mentioned results together with group information, including annotations of conditions that were compared, list of samples in each group for provenance as well as DE and classification module version and initial settings for reproducibility.

### 3. Semantic data layer

Given diversity of biological data, the users of the corresponding information systems should be given a possibility to interpret data independently using common terminologies. For this purpose we developed a semantic data layer, which provides unified access to data together with metadata and interpretations. Metadata is normalized using ontologies, which makes annotations standardized and hierarchically searchable. Data interpretations are represented by DE and classification result, described in Section 2.

#### 3.1. Ontologies

In biological research, controlled biological vocabularies such as terminologies, ontologies, taxonomies play an important role for annotation, integration, and analysis of biological data. In order to allow for the interoperability of data, one important step is to standardize annotations using ontologies and semantic mapping (Schuurman and Leszczynski, 2008). Ontologies define standard terms, their properties, and the relations between them. Each term in the ontology has a unique identifier, and annotation terms that are connected to ontologies are called ‘normalized’. Another advantage of ontology use is a possibility to search by parent terms. Ontology is a hierarchical structure, where children are more specific as parents. Usually parents are connected to children using one of the two relations: “**is-a**” or “**part-of**”. “is-a” relation means that a child represents more specific term as a parent, for example Alzheimer’s disease **is a** tauopathy, tauopathy **is a** neurodegenerative disease. “part-of” relation means that a child is part of a parent, for example neocortex is **part of** a cerebral cortex, cerebral cortex is **part of** a brain.

In order to enable the search across ontological hierarchies we integrated the relevant ontologies into the graph database Neo4j. Graph databases are NoSQL databases which support storage of objects and connections between them, as is the case for ontologies. Following the manual curation, sample annotations are uploaded to the SEA annotation graph database including all ontological parent terms (having an ‘is-a’ or ‘part of’ relation to it). This allows search by ontology terms, as well as by their parents, which are in fact groups of terms (e.g. ‘cancer’ or ‘neurodegenerative disease’). SEA accesses ontologies via the Ontology Lookup Service using a REST interface, supporting complex and compound queries and query auto-completion.

#### 3.2. Annotation Process

SEA’s sRNA annotation workflow maps free-text GEO annotations to standardized terms in three consecutive steps. In general, GEO data annotations are free text that can be parsed into key-value pairs. In a first fully automated step the annotation workflow extracts key-value relations and stores them in the annotation database. These key-value relations are extracted from an experiment or sample description text. It is unstructured and contains variety of information: therefore we opted for a NoSql annotation database with an optimized indexing for prototypical questions (supplementary section 2.4). The second fully automated step normalizes the extracted keys and values using ontologies as standard dictionaries. SEA has a list of predefined keys, five of which (organism, tissue, disease, cell type, and cell line) can be currently queried for in SEA. Each extracted key is compared to predefined keys. For values, the ontologies are used as standard terminology dictionaries. For each pre-defined key, one or several corresponding ontologies are used. Each extracted value is searched in the corresponding ontologies and, if the same or a similar term is found, we normalize the value by it.

Automatic annotation is followed by semi-automatic manual curation. For that purpose, we developed an internal curation Web interface (Supplementary Fig 1) using Groovy/Grails^7^, which allows browsing and editing of annotations from the annotation database as well as manual normalization of keys and values in annotations, searching among pre-defined keys and corresponding ontologies. Thus, curators examine all keys and values for consistency and update missing or additional information with standardized terms where necessary (e.g. organism, tissue, cell type, cell line, disease). At the moment, all SEA annotations are manually curated, a quality standard that we intend to keep for every future SEA entry.

#### 3.3. Annotation criteria

Some basic rules for annotation of sRNA-seq samples are discussed below:

1. Annotations can be defined as experiment-level or sample-level annotations. Experiment level annotations are exactly the same for all samples of the dataset. Sample-level annotations differ among the samples of the dataset.
2. In exceptional situations, when experiment-level and sample-level annotations have the same key, the corresponding sample will have two annotations with that key (local annotation does not override the global one).
3. In cases, where alternatives exist for an annotation term, we tried to be as specific as possible. For example, in case the sample is from breast fibrosarcoma and the term “breast fibrosarcoma” is available we will try to annotate it with the same term, although it can be annotated with breast cancer as well i.e. we choose the term deepest in the ontology tree.
4. In cases where the relevant term cannot be found in any ontology, we tried to normalize the terms with synonyms or slightly less specific. The aim was to annotate as much as possible to have standard terms rather than just textual information

#### 3.4. Annotation of groups

Apart from sRNA expression, SEA contains also results of differential expression and classification between different biological conditions. In order to make these comparisons automatically possible, biologically meaningful comparisons are also annotated manually. This annotation belongs to a dataset and we call it “sample group”. Sample group annotation includes annotation fields, which are important to define groups to compare. For example:

1. In the case an experiment has healthy and diseased patients. The desired comparison will be to perform differential expression or classification based on these two groups of samples. In this case, sample group is annotated as disease. The field disease contains two distinct values – disease name (for example, Alzheimer’s disease) and “healthy”. And correspondingly, one comparison will be made.
2. In the case an experiment contains two diseases and healthy controls. Sample group is annotated as disease. The field disease contains three values – disease 1 name (for example, Alzheimer’s disease) disease 2 name (for example, Frontotemporal Dementia) and “healthy”. Correspondingly, three comparisons will be made: healthy vs. disease 1, healthy vs. disease 2 and disease 1 vs. disease 2.
3. In the case an experiment contains one disease with two stages and healthy controls. In this case, sample group is annotated as disease + disease details. Correspondingly, three groups will be created: disease+stage1, disease+stage2, healthy. Correspondingly, three comparisons will be made: healthy vs. disease stage1, healthy vs. disease stage2 and disease stage1 vs. disease stage2.

#### 3.5. Querying and visualization

Application programming interfaces (APIs) were developed to access data in SEA databases. We developed **annotation-API** to access annotation-DB, **association-API** to access association-DB, **expression-API** to access expression-DB, **predictedMirna-API** to access Oasis-DB and **SEA business logic API** to call all these APIs based on the end user request and then combine their results (responses) back to the end user. Each API answer particular queries as explained below.

1. **Annotation-API** to access Annotation-DB, and answer questions like

a. Get all the experiments for term (will return experiments and samples along with annotation details for term as well as for the sub-type of term). Term can be disease, tissue, cell line, cell type, organism or their combination
2. **Association-API** to access Association-DB, and respond to questions like

a. Get all gene targets for a miRNA.
b. Get all diseases that are associated with a miRNA from literature.
c. Get all miRNAs that are associated with a disease or it’s sub-types.
d. Get genomic coordinates for sRNA or target genes.
3. **Expression-API** to access Expression-DB and shows

a. Get expression of one or more sRNAs in a particular or all experiments.
b. Get list of experiments where a particular sRNA is differentially expressed.
c. Get list of experiments where a particular sRNA is identified as potential biomarker via classification.
d. Get expression of a pathogen in a particular or all experiments.
e. Get list of all experiments where a particular pathogen is differentially expressed.
4. **PredictedMirna-API** to access Oasis-DB (Rahman et al.) and shows

a. Get list of all novel predicted miRNAs from Oasis-DB
b. Get genomic coordinates and sequence for predicted miRNA(s).
5. **User-API** to access User-DB and gets

a. Get list of all user specific experiments (uploaded by the user).
b. Get list of experiments where a particular sRNA is differentially expressed in their own uploaded data.
c. Get list of experiments where a particular sRNA is identified as potential biomarker via classification in their own uploaded data.
6. **SEA business logic API:** was built in order to put all those APIs together and make necessary data transformations between frontend and other APIs. As a result, the user can make queries to answer biological questions like;

a. What is expression of one more sRNAs in specific cell types or tissues?
b. Is a particular sRNA differentially expressed in Alzheimer’s disease?
c. Compare sRNAs across different studies?
d. What are differentially expressed sRNAs in breast cancer and healthy women?
e. Common differentially expressed sRNAs or potential sRNAs based biomarker across particular disease or tissue.
f. Expression of one or more novel miRNAs for known diseased states.
g. Analysis results with all the quality information for 350 datasets and over 4200 samples.

---

**Supplementary Fig.1.**
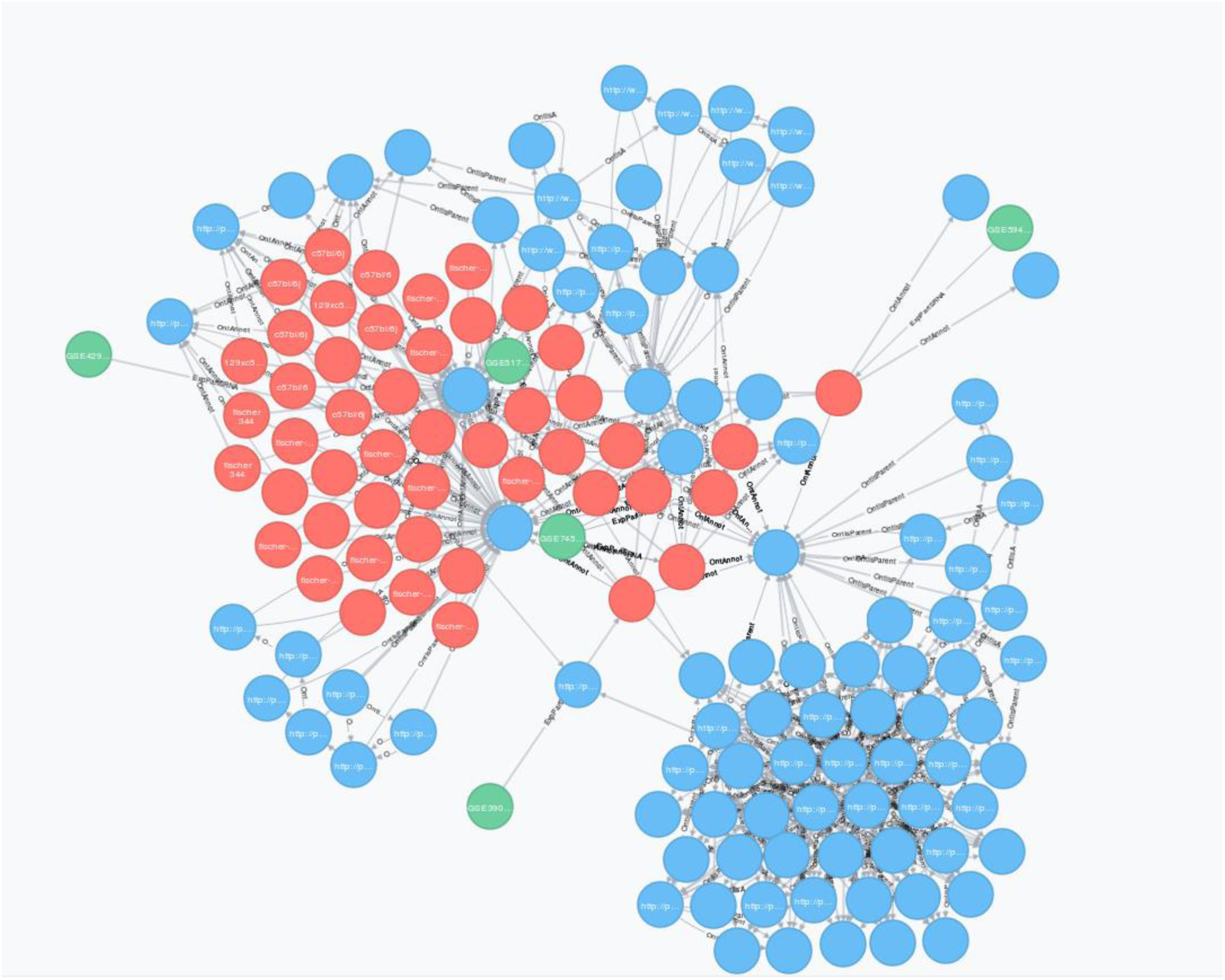
Annotation-DB. Objects in the SEA graph database (Neo4j). A fragment of the SEA graph database is visualized, where green nodes represent datasets, red nodes represent samples and blue nodes represent ontology terms. Grey edges represent ‘is a’ relations between the different datasets, samples, and ontology terms.

## Footnotes

7 https://grails.org/

## Notes

#### Summary of Updates

To remove wrong ORCIDs and correct spelling of one author name.

