## Supplementary material for "SEA: The small RNA Expression Atlas": p-hsa-miR-113

| | | |
|---|---|---|
| Provisional ID | : | chr2_1671 |
| Score total | : | 526.3 |
| Score for star read(s) | : | 3.9 |
| Score for read counts | : | 520.1 |
| Score for mfe | : | 1.3 |
| Score for randfold | : | 1.6 |
| Score for cons. seed | : | -0.6 |
| Total read count | : | 1032 |
| Mature read count | : | 1012 |
| Loop read count | : | 1 |
| Star read count | : | 19 |

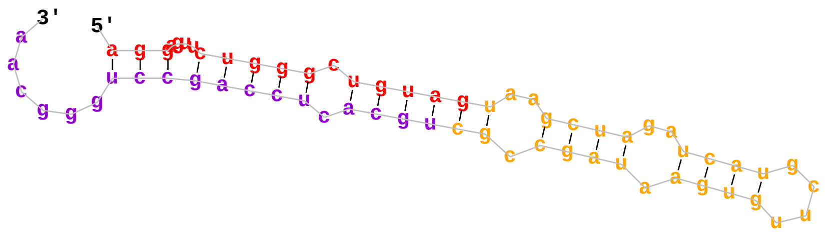


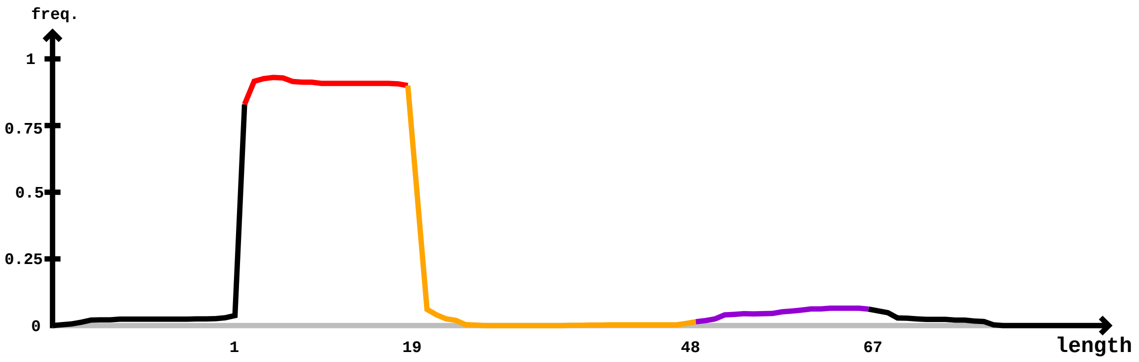


Mature Star

```
5'- ggaggauugcuugaaccuaggaguucugggcuguaguaagcuagaucaugcuugugaauagccgcugcacuccagccugggcaacagaaggagacuaugucucuaaaa  -3'  obs
    ggaggauugcuugaaccuaggaguucugggcuguaguaagcuagaucaugcuugugaauagccgcugcacuccagccugggcaacagaaggagacuaugucucuaaaa        exp
    ......(((.(((..((((((....(((((.((((((..((((..((((....)))).)))).)))))).))))))))))))))))..((((((...))))))....   reads   mm   sample
    ggagAauugcuugaaccu..........................................................................................   2       1    aou
    ggagAauugcuugaaccuaggag.....................................................................................   2       1    aou
    ggaggauugcuugaaccuaggagAuc..................................................................................   3       1    aou
    .gaggauugcuugaUccuagga......................................................................................   1       1    aou
    .gagAauugcuugaaccuaggag.....................................................................................   1       1    aou
    .gaggauugcuugaaccuaggagu....................................................................................   3       0    aou
    .gaggauugcuugaaccuaggagAuc..................................................................................   2       1    aou
    ..aggauugcuugaaccuagga......................................................................................   1       0    aou
    ..agAauugcuugaaccuaggag.....................................................................................   5       1    aou
    ..aggauugcuugaaccuaggag.....................................................................................   3       0    aou
    ...gAauugcuugaaccuaggag.....................................................................................   1       1    aou
    .....auugcuugaaccuGggag.....................................................................................   3       1    aou
    .............aGccuaggaguucugggcugu..........................................................................   1       1    aou
    ...............ccCaggaguucugggcuguaguaa.....................................................................   1       1    aou
    ................cNaggaguucugggcugu..........................................................................   1       1    aou
    ................cNaggaguucugggcugua.........................................................................   1       1    aou
    ................cAaggaguucugggcuguag........................................................................   1       1    aou
    ................cCaggaguucugggcuguaguaagc...................................................................   1       1    aou
    .................uaggaguucugggcugua.........................................................................   5       0    aou
    .................Gaggaguucugggcugua.........................................................................   1       1    aou
    .................uaggaguucugggcuguag........................................................................   1       0    aou
    .................uaggaguucugggcugGag........................................................................   1       1    aou
    .................Caggaguucugggcuguaguaa.....................................................................   1       1    aou
    ..................aggaguucugggcuguag........................................................................   507     0    aou
    ..................aggaguucugggcuUuag........................................................................   4       1    aou
    ..................aggNguucugggcuguag........................................................................   1       1    aou
    ..................aggaguucugggGuguag........................................................................   1       1    aou
    ..................agNaguucugggcuguag........................................................................   1       1    aou
    ..................aggaguucugggcuguCg........................................................................   1       1    aou
    ..................aggaguucugggcuguaC........................................................................   1       1    aou
    ..................aggaguucugggUuguag........................................................................   7       1    aou
    ..................aggaguucugggcugGag........................................................................   1       1    aou
    ..................aggCguucugggcuguag........................................................................   1       1    aou
```

Mature                                             Star

```
ggaggauugcuugaaccuaggaguucugggcuguaguaagcuagaucaugcuugugaauagccgcugcacuccagccugggcaacagaaggagacuaugucucuaaaa
..................aggaguucugCgcuguag........................................................................    4     1     aou
..................aggaguucugggcuAuag........................................................................    1     1     aou
..................aggaguucuUggcuguag........................................................................    6     1     aou
..................aggaguucuggUcuguag........................................................................    10    1     aou
..................aggaguucugUgcuguag........................................................................    9     1     aou
..................aggaguucuggCcuguag........................................................................    1     1     aou
..................aggaguucugggcuguGg........................................................................    2     1     aou
..................aggGguucugggcuguag........................................................................    2     1     aou
..................aggaguucugggcuguaA........................................................................    2     1     aou
..................aggaguucuggAcuguag........................................................................    1     1     aou
..................aggaguucugggAuguag........................................................................    1     1     aou
..................Nggaguucugggcuguag........................................................................    1     1     aou
..................aggaguucugggcugAag........................................................................    2     1     aou
..................aggaguuAugggcuguag........................................................................    3     1     aou
..................aggagCucugggcuguag........................................................................    3     1     aou
..................aNgaguucugggcuguag........................................................................    1     1     aou
..................aggaguuAugggcuguagu.......................................................................    2     1     aou
..................aggaguucugUgcuguagu.......................................................................    10    1     aou
..................aggaguucugggcuguNgu.......................................................................    1     1     aou
..................aggaguucugggcuguaAu.......................................................................    3     1     aou
..................aggCguucugggcuguagu.......................................................................    2     1     aou
..................Nggaguucugggcuguagu.......................................................................    1     1     aou
..................aggaguucugggUuguagu.......................................................................    5     1     aou
..................aggaguucugggcugGagu.......................................................................    3     1     aou
..................aggaguucugCgcuguagu.......................................................................    3     1     aou
..................aggaguucugggcuguCgu.......................................................................    1     1     aou
..................aggaAuucugggcuguagu.......................................................................    1     1     aou
..................aggagCucugggcuguagu.......................................................................    1     1     aou
..................aggaguucugggcuguagC.......................................................................    3     1     aou
..................aggaguucuggCcuguagu.......................................................................    4     1     aou
..................agCaguucugggcuguagu.......................................................................    1     1     aou
..................aggaguucuggUcuguagu.......................................................................    13    1     aou
..................aggaguucugggcuguagu.......................................................................    218   0     aou
..................aggGguucugggcuguagu.......................................................................    1     1     aou
..................aggaguucugggcuNuagu.......................................................................    1     1     aou
..................Cggaguucugggcuguagu.......................................................................    1     1     aou
..................agUaguucugggcuguagu.......................................................................    1     1     aou
..................agAaguucugggcuguagu.......................................................................    2     1     aou
..................aggaguucugggcugCagu.......................................................................    3     1     aou
..................aggaguucugggcuUuagu.......................................................................    4     1     aou
..................aggaguCcugggcuguagu.......................................................................    1     1     aou
..................aggaguucugAgcuguagu.......................................................................    1     1     aou
..................aggaguucugggcuguagua......................................................................    3     0     aou
..................aggaguucugggcuguaguC......................................................................    11    1     aou
..................aggaguucugggcuguaAua......................................................................    2     1     aou
..................aggaguucugggcuguaguU......................................................................    1     1     aou
..................aggaguucugggcuguaguGa.....................................................................    9     1     aou
..................aggaguucugggcuguaguaaC....................................................................    1     1     aou
..................aggaguucugggcuguaguaaAc...................................................................    1     1     aou
..................aggaguucugggcuguaguaaUc...................................................................    1     1     aou
...................gUaguucugggcuguagu.......................................................................    1     1     aou
...................ggaguCcugggcuguagu.......................................................................    1     1     aou
...................ggCguucugggcuguagu.......................................................................    2     1     aou
...................ggaguucugggcuguaCu.......................................................................    1     1     aou
...................ggaguucugggcuguagu.......................................................................    70    0     aou
...................ggaguucugggAuguagu.......................................................................    1     1     aou
...................Ugaguucugggcuguagu.......................................................................    1     1     aou
...................ggaguucugggcuguaguC......................................................................    1     1     aou
...................ggaguucugggcuguaguU......................................................................    1     1     aou
...................ggaguucugggcuguagua......................................................................    2     0     aou
...................ggaguucugggcuguaAua......................................................................    1     1     aou
...................ggaguucugggcuguaguGa.....................................................................    5     1     aou
...................ggaguucugggcuguaguaaC....................................................................    4     1     aou
...................ggaguucugggcuguaguCagc...................................................................    1     1     aou
...................ggaguucugggcuguaguUagc...................................................................    2     1     aou
...................ggaguucugggcuguaguGagc...................................................................    1     1     aou
...................ggaguucuAggcuguaguaagc...................................................................    1     1     aou
...................ggaguucugggcuguaguGagcu..................................................................    1     1     aou
....................gaguucugggcuguaguC......................................................................    1     1     aou
....................gaguucugggcuguaguCa.....................................................................    1     1     aou
```

**Mature** **Star**

```
ggaggauugcuugaaccuaggaguucugggcuguaguaagcuagaucaugcuugugaauagccgcugcacuccagccugggcaacagaaggagacuaugucucuaaaa
....................gaguucugggcuguaguGag....................................................................	1	1	aou
....................gaguucugggcuguaguaaUc...................................................................	1	1	aou
....................gaguucugggcuguaguaaAc...................................................................	1	1	aou
....................gaguucugggcuguaguaagc...................................................................	3	0	aou
....................gaguucugggcuguaguUagc...................................................................	1	1	aou
....................gaguucugggcuguaguaCgcua.................................................................	2	1	aou
.....................aguucugggcuguaguaaAc...................................................................	2	1	aou
.....................aguucugggcuguaguGagc...................................................................	2	1	aou
.....................aguucugggcuguaguGagcu..................................................................	1	1	aou
....................................................ugugaauagcGgcugcacu.....................................	1	1	aou
......................................................ugaauagccCcugcacucca..................................	1	1	aou
........................................................aauagccCcugcacuccag.................................	1	1	aou
................................................................cugcacuccagccCgggca.........................	1	1	aou
................................................................cugcacCccagccugggca.........................	1	1	aou
................................................................cugcacuccaUccugggca.........................	1	1	aou
................................................................cugcacucNagccugggcaa........................	1	1	aou
................................................................cugcacuccaNccugggcaa........................	1	1	aou
................................................................cugcacuccaNccugggcaac.......................	1	1	aou
.................................................................ugcacuccagAcugggca.........................	1	1	aou
.................................................................ugcacucNagccugggcaa........................	1	1	aou
.................................................................ugcacuccagAcugggcaa........................	1	1	aou
.................................................................ugcacuccagNcugggcaa........................	1	1	aou
.................................................................ugcaUuccagccugggcaa........................	1	1	aou
.................................................................ugcacuccagNcugggcaaca......................	2	1	aou
..................................................................gcacuccagcUugggcaa........................	1	1	aou
..................................................................gcacuGcagccugggcaa........................	2	1	aou
..................................................................gcacuUcagccugggcaac.......................	1	1	aou
..................................................................gcacuGcagccugggcaac.......................	1	1	aou
...................................................................cacuccaNccugggcaac.......................	1	1	aou
...................................................................cacuccagccuggUcaac.......................	1	1	aou
...................................................................cacuGcagccugggcaac.......................	1	1	aou
...................................................................cacuccagcUugggcaaca......................	2	1	aou
...................................................................cacuccagccugUgcaaca......................	1	1	aou
...................................................................cacuGcagccugggcaaca......................	1	1	aou
....................................................................acuccagcAugggcaaca......................	1	1	aou
....................................................................acuccagccugAgcaaca......................	12	1	aou
....................................................................acuccagAcugggcaaca......................	1	1	aou
....................................................................acuccagccAgggcaaca......................	1	1	aou
....................................................................acuccagcUugggcaaca......................	1	1	aou
....................................................................acuccagcGugggcaacag.....................	1	1	aou
.....................................................................cuccagcAugggcaacaga....................	1	1	aou
.....................................................................cuccagGcugggcaacaga....................	1	1	aou
......................................................................uccagccuggNcaacaga....................	1	1	aou
......................................................................uccagccugggcaCcagaa...................	1	1	aou
......................................................................uccagAcugggcaacagaa...................	1	1	aou
........................................................................cagccuggAcaacagaagga................	1	1	aou
.........................................................................Cgccugggcaacagaaggagacu............	1	1	aou
..........................................................................gccugggcaacagaaUga................	1	1	aou
..........................................................................gccugggcaacagaaCga................	1	1	aou
..........................................................................gccugggcaacagaaAgagac.............	1	1	aou
..........................................................................gccugggcaacagaagUagacu............	1	1	aou
..........................................................................Accugggcaacagaaggagacu............	3	1	aou
..........................................................................gccugggcaacagaaggagacu............	1	0	aou
...........................................................................ccugggcaacagaaCgaga..............	1	1	aou
...........................................................................ccugggcaacagaaUgaga..............	1	1	aou
...........................................................................ccugggcaacagaaUgagac.............	1	1	aou
...........................................................................ccugggcaacagaagUagacu............	1	1	aou
............................................................................cugggcaacagaGggaga..............	2	1	aou
............................................................................cugggcaacagaaAgagacu............	1	1	aou
............................................................................cugggcaacagaaggagacu............	1	0	aou
.............................................................................ugggcaacagaGggagacu............	3	1	aou
.............................................................................ugggcaacagaaggagacu............	2	0	aou
...............................................................................ggcaacagaaCgagacua...........	3	1	aou
```
