## Supplementary material for "SEA: The small RNA Expression Atlas": p-hsa-miR-247

| | | |
|---|---|---|
| Provisional ID | : | chr14_823 |
| Score total | : | 510487.8 |
| Score for star read(s) | : | 3.9 |
| Score for read counts | : | 510477.3 |
| Score for mfe | : | 2 |
| Score for randfold | : | 1.6 |
| Score for cons. seed | : | 3 |
| Total read count | : | 1001291 |
| Mature read count | : | 1001289 |
| Loop read count | : | 1 |
| Star read count | : | 1 |

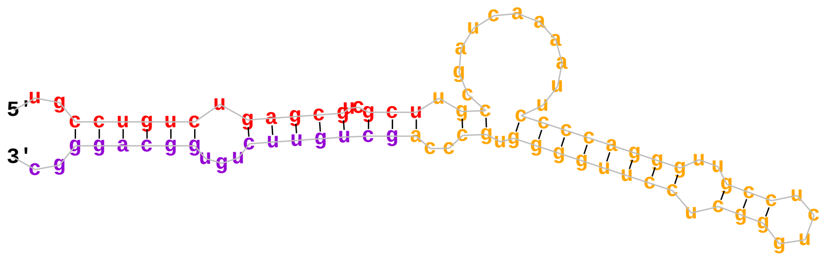


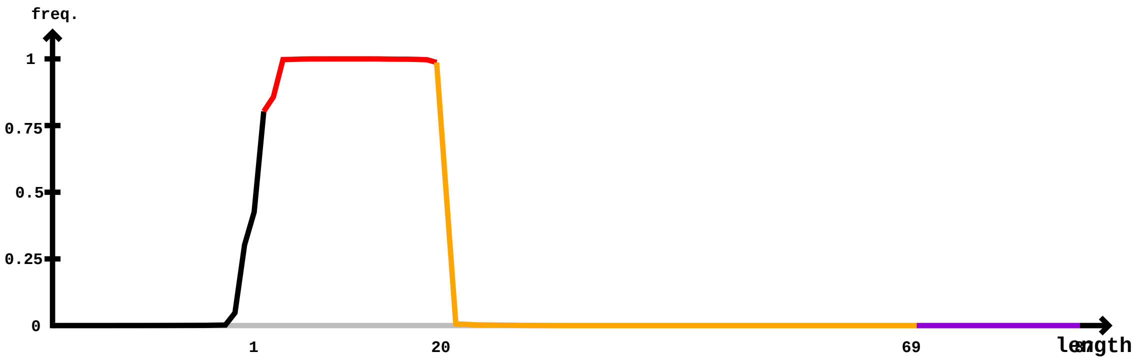


```
                        Mature                                                        Star
5'- cuggguuccucccagggcuaugccugucugagcgucgcuugccgaucaaaauuccccaggguugccucugggcuccuuggggugcccagcuguucuguggcagggccc  -3'   obs
    cuggguuccucccagggcuaugccugucugagcgucgcuugccgaucaaaauuccccaggguugccucugggcuccuuggggugcccagcuguucuguggcagggccc        exp
    (((((.....)))))((((.((((.....(((((..(((.((..........((((((((..(((....))).)))))))).))..))))))))...)))).)))).   reads   mm   sample
    ..ggguuccucccGgggcua........................................................................................   2       1    mv7
    ....guuccucccGgggcuaug......................................................................................   1       1    mv7
    ....guuccucccGgggcuaugccugu.................................................................................   1       1    mv7
    ....guuccucccGgggcuaugccuguc................................................................................   1       1    mv7
    ....guuccucccGgggcuaugccugucug..............................................................................   1       1    mv7
    ....guuccucccagggcuaCgccugucugagc...........................................................................   2       1    mv7
    ....guuccucccagggcuaCgccugucugagcguc........................................................................   1       1    mv7
    .....uuccucccGgggcuaugccugucuga.............................................................................   1       1    mv7
    .....uuccucccagggcuaCgccugucugag............................................................................   1       1    mv7
    .....uuccucccagggcuaCgccugucugagc...........................................................................   9       1    mv7
    .....uuccucccGgggcuaugccugucugagc...........................................................................   20      1    mv7
    .....uuccucccGgggcuaugccugucugagcg..........................................................................   1       1    mv7
    .....uuccucccagggcuaCgccugucugagcgu.........................................................................   2       1    mv7
    .....uuccucccagggcuaCgccugucugagcguc........................................................................   2       1    mv7
    .....uuccucccGgggcuaugccugucugagcguc........................................................................   3       1    mv7
    .....uuccucccGgggcuaugccugucugagcgucg.......................................................................   4       1    mv7
    .....uuccucccagggcuaCgccugucugagcgucg.......................................................................   2       1    mv7
    ......uccucccGgggcuaugccugucug..............................................................................   1       1    mv7
    ......uccucccagggcuaCgccugucugag............................................................................   1       1    mv7
    ......uccucccGgggcuaugccugucugag............................................................................   1       1    mv7
    ......uccucccGgggcuaugccugucugagc...........................................................................   14      1    mv7
    ......uccucccagggcuaCgccugucugagc...........................................................................   5       1    mv7
    ......uccucccGgggcuaugccugucugagcg..........................................................................   1       1    mv7
    ......uccucccagggcuaCgccugucugagcgu.........................................................................   1       1    mv7
    ......uccucccGgggcuaugccugucugagcgu.........................................................................   1       1    mv7
    ......uccucccGgggcuaugccugucugagcguc........................................................................   5       1    mv7
    ......uccucccagggcuaCgccugucugagcgucgc......................................................................   2       1    mv7
    ......uccucccGgggcuaugccugucugagcgucgc......................................................................   1       1    mv7
    .......ccucccagggcuaCgccugucuga.............................................................................   2       1    mv7
    .......ccucccGgggcuaugccugucugag............................................................................   1       1    mv7
    .......ccucccagggcuaCgccugucugagc...........................................................................   2       1    mv7
    .......ccucccGgggcuaugccugucugagc...........................................................................   2       1    mv7
    .......ccucccagggcuaCgccugucugagcgu.........................................................................   2       1    mv7
```

```
                    Mature                                                            Star
cuggguuccucccagggcuaugccugucugagcgucgcuugccgaucaaaauuccccaggguugccucugggcuccuuggggugcccagcuguucuguggcagggccc
.......ccucccagggcuaCgccugucugagcguc........................................................................ 2   1   mv7
.......ccucccGgggcuaugccugucugagcgucgcu..................................................................... 43  1   mv7
.......ccucccagggcuaCgccugucugagcgucgcu..................................................................... 7   1   mv7
........cucccGgggcuaugccugu................................................................................. 1   1   mv7
........cucccGgggcuaugccugucugagc........................................................................... 2   1   mv7
........cucccagggcuaCgccugucugagcgu......................................................................... 1   1   mv7
........cucccGgggcuaugccugucugagcgucgcu..................................................................... 41  1   mv7
........cucccCgggcuaugccugucugagcgucgcu..................................................................... 1   1   mv7
........cucccagggcuaCgccugucugagcgucgcu..................................................................... 6   1   mv7
........cucccagggcuaCgccugucugagcgucgcuu.................................................................... 1   1   mv7
.........ucccGgggcuaugccugucugag............................................................................ 1   1   mv7
.........ucccGgggcuaugccugucugagc........................................................................... 1   1   mv7
.........ucccagggcuaCgccugucugagc........................................................................... 1   1   mv7
.........ucccGgggcuaugccugucugagcgu......................................................................... 1   1   mv7
.........ucccagggcuaCgccugucugagcgu......................................................................... 1   1   mv7
.........ucccGgggcuaugccugucugagcgucg....................................................................... 2   1   mv7
.........ucccagggcuaCgccugucugagcgucgc...................................................................... 1   1   mv7
.........ucccGgggcuaugccugucugagcgucgc...................................................................... 1   1   mv7
.........ucccGgggcuaugccugucugagcgucgcu..................................................................... 19  1   mv7
.........ucccagggcuaCgccugucugagcgucgcu..................................................................... 2   1   mv7
.........ucccGgggcuaugccugucugagcgucgcuu.................................................................... 1   1   mv7
..........cccGgggcuaugccugucuga............................................................................. 1   1   mv7
..........cccGgggcuaugccugucugag............................................................................ 2   1   mv7
..........cccagggcuaCgccugucugag............................................................................ 5   1   mv7
..........cccagggcuaCgccugucugagc........................................................................... 10  1   mv7
..........cccGgggcuaugccugucugagc........................................................................... 6   1   mv7
..........cccagggcuaCgccugucugagcgu......................................................................... 8   1   mv7
..........cccGgggcuaugccugucugagcgu......................................................................... 1   1   mv7
..........cccGgggcuaugccugucugagcguc........................................................................ 5   1   mv7
..........cccagggcuaCgccugucugagcguc........................................................................ 8   1   mv7
..........cccagggcuaCgccugucugagcgucg....................................................................... 7   1   mv7
..........cccagggcuaugccugucAgagcgucg....................................................................... 1   1   mv7
..........cccagggcuaCgccugucugagcgucgc...................................................................... 1   1   mv7
..........cccGgggcuaugccugucugagcgucgcu..................................................................... 18  1   mv7
..........cccCgggcuaugccugucugagcgucgcu..................................................................... 1   1   mv7
..........cccagggcuaCgccugucugagcgucgcu..................................................................... 29  1   mv7
..........cccGgggcuaugccugucugagcgucgcuu.................................................................... 6   1   mv7
...........ccGgggcuaugccugucu............................................................................... 2   1   mv7
...........ccagggcuaCgccugucugag............................................................................ 2   1   mv7
...........ccGgggcuaugccugucugagc........................................................................... 1   1   mv7
...........ccagggcuaCgccugucugagc........................................................................... 3   1   mv7
...........ccagggcuaCgccugucugagcg.......................................................................... 8   1   mv7
...........ccagggcuaCgccugucugagcgu......................................................................... 5   1   mv7
...........ccUgggcuaugccugucugagcgu......................................................................... 1   1   mv7
...........ccGgggcuaugccugucugagcgu......................................................................... 4   1   mv7
...........ccagggcuaCgccugucugagcguc........................................................................ 3   1   mv7
...........ccGgggcuaugccugucugagcguc........................................................................ 1   1   mv7
...........ccagggcuaCgccugucugagcgucg....................................................................... 1   1   mv7
...........ccGgggcuaugccugucugagcgucg....................................................................... 2   1   mv7
...........ccagggcuaCgccugucugagcgucgc...................................................................... 6   1   mv7
...........ccagggcuaCgccugucugagcgucgcu..................................................................... 25  1   mv7
...........ccCgggcuaugccugucugagcgucgcu..................................................................... 1   1   mv7
...........ccGgggcuaugccugucugagcgucgcu..................................................................... 22  1   mv7
...........Ucagggcuaugccugucugagcgucgcu..................................................................... 1   1   mv7
...........ccagggcuaugccugucugagcgucgcuC.................................................................... 1   1   mv7
...........ccagggcuaCgccugucugagcgucgcuu.................................................................... 2   1   mv7
...........ccGgggcuaugccugucugagcgucgcuu.................................................................... 8   1   mv7
............cagggcuaCgccugucugag............................................................................ 22  1   mv7
............cGgggcuaugccugucugag............................................................................ 16  1   mv7
............cGgggcuaugccugucugagc........................................................................... 5   1   mv7
............cagggcuaCgccugucugagc........................................................................... 2   1   mv7
............cGgggcuaugccugucugagcg.......................................................................... 8   1   mv7
............cagggcuaCgccugucugagcg.......................................................................... 7   1   mv7
............cagggcuaCgccugucugagcgu......................................................................... 2   1   mv7
............cGgggcuaugccugucugagcgu......................................................................... 2   1   mv7
............cUgggcuaugccugucugagcgu......................................................................... 1   1   mv7
............cagggcuaCgccugucugagcguc........................................................................ 1   1   mv7
............cagggcuaCgccugucugagcgucg....................................................................... 1   1   mv7
............cGgggcuaugccugucugagcgucg....................................................................... 1   1   mv7
............cGgggcuaugccugucugagcgucgc...................................................................... 2   1   mv7
```

Mature Star

| | | | |
|---|---|---|---|
| cuggguuccucccagggcuaugccugucugagcgucgcuugccgaucaaaauuccccaggguugccucugggcuccuuggggugcccagcuguucuguggcagggccc | | | |
| ............cGgggcuaugccugucugagcgucgcu..................................................................... | 21 | 1 | mv7 |
| ............cagggcuaCgccugucugagcgucgcu..................................................................... | 12 | 1 | mv7 |
| ............cGgggcuaugccugucugagcgucgcuu.................................................................... | 8 | 1 | mv7 |
| ............cCgggcuaugccugucugagcgucgcuu.................................................................... | 1 | 1 | mv7 |
| .............Ggggcuaugccugucuga............................................................................. | 1 | 1 | mv7 |
| .............agggcuaCgccugucuga............................................................................. | 1 | 1 | mv7 |
| .............agggcuaCgccugucugag............................................................................ | 6 | 1 | mv7 |
| .............Ggggcuaugccugucugag............................................................................ | 1 | 1 | mv7 |
| .............agggcuaAgccugucugagc........................................................................... | 1 | 1 | mv7 |
| .............Ggggcuaugccugucugagc........................................................................... | 4 | 1 | mv7 |
| .............agggcuaCgccugucugagc........................................................................... | 5 | 1 | mv7 |
| .............Ggggcuaugccugucugagcg.......................................................................... | 1 | 1 | mv7 |
| .............agggcuaCgccugucugagcg.......................................................................... | 1 | 1 | mv7 |
| .............agggcuaCgccugucugagcgu......................................................................... | 3 | 1 | mv7 |
| .............agggcuaCgccugucugagcgucgcu..................................................................... | 1 | 1 | mv7 |
| .............Ggggcuaugccugucugagcgucgcu..................................................................... | 15 | 1 | mv7 |
| ..............gggcuaCgccugucugag............................................................................ | 23 | 1 | mv7 |
| ..............gggcuaAgccugucugag............................................................................ | 1 | 1 | mv7 |
| ..............gggcuaCgccugucugagc........................................................................... | 43 | 1 | mv7 |
| ..............gggcuaCgccugucugagcg.......................................................................... | 16 | 1 | mv7 |
| ..............gggcuaCgccugucugagcgu......................................................................... | 15 | 1 | mv7 |
| ..............gggcuaCgccugucugagcguc........................................................................ | 9 | 1 | mv7 |
| ..............gggcuaCgccugucugagcgucg....................................................................... | 13 | 1 | mv7 |
| ..............gggcuaCgccugucugagcgucgc...................................................................... | 1 | 1 | mv7 |
| ..............gggcuaugccugucugagcguUgcu..................................................................... | 1 | 1 | mv7 |
| ..............gggcuaugccugucugagcgucgcu..................................................................... | 4 | 0 | mv7 |
| ..............gggcuaCgccugucugagcgucgcu..................................................................... | 49 | 1 | mv7 |
| ..............gggcuaCgccugucugagcgucgcuu.................................................................... | 12 | 1 | mv7 |
| ...............ggcuaCgccugucugagc........................................................................... | 575 | 1 | mv7 |
| ...............ggcuaGgccugucugagc........................................................................... | 1 | 1 | mv7 |
| ...............ggcuaugccugucugagU........................................................................... | 1 | 1 | mv7 |
| ...............ggcuaugccugucugagc........................................................................... | 2 | 0 | mv7 |
| ...............ggcuaugccugucGgagc........................................................................... | 1 | 1 | mv7 |
| ...............ggcuaCgccugucugagcg.......................................................................... | 28 | 1 | mv7 |
| ...............ggcuaCgccugucugagcgu......................................................................... | 8 | 1 | mv7 |
| ...............ggcuaCgccugucugagcguc........................................................................ | 11 | 1 | mv7 |
| ...............ggcuaCgccugucugagcgucg....................................................................... | 13 | 1 | mv7 |
| ...............ggcuaCgccugucugagcgucgc...................................................................... | 5 | 1 | mv7 |
| ...............ggcuaCgccugucugagcgucgcu..................................................................... | 88 | 1 | mv7 |
| ...............ggcuaCgccugucugagcgucgcuu.................................................................... | 8 | 1 | mv7 |
| ...............ggcuaugccugucugagcgucgcuu.................................................................... | 2 | 0 | mv7 |
| ................gcuaugccugucugagcA.......................................................................... | 1 | 1 | mv7 |
| ................gcuaCgccugucugagcg.......................................................................... | 218 | 1 | mv7 |
| ................gcuaCgccugucugagcgu......................................................................... | 29 | 1 | mv7 |
| ................gcuaCgccugucugagcguc........................................................................ | 33 | 1 | mv7 |
| ................gcuaCgccugucugagcgucg....................................................................... | 3 | 1 | mv7 |
| ................gcuaCgccugucugagcgucgc...................................................................... | 1 | 1 | mv7 |
| ................gcuaugccugucugagcgucgcC..................................................................... | 2 | 1 | mv7 |
| ................gcuaugccugucugagcgucgcu..................................................................... | 13 | 0 | mv7 |
| ................Ucuaugccugucugagcgucgcu..................................................................... | 1 | 1 | mv7 |
| ................gcuaCgccugucugagcgucgcu..................................................................... | 156 | 1 | mv7 |
| ................gcuaugccCgucugagcgucgcu..................................................................... | 1 | 1 | mv7 |
| ................gcuaCgccugucugagcgucgcuu.................................................................... | 10 | 1 | mv7 |
| ................gcuaCgccugucugagcgucgcuugc.................................................................. | 2 | 1 | mv7 |
| ................gcuaCgccugucugagcgucgcuugccg................................................................ | 1 | 1 | mv7 |
| .................cuaAgccugucugagcgu......................................................................... | 2 | 1 | mv7 |
| .................cuaugccugucugagcgu......................................................................... | 2 | 0 | mv7 |
| .................cuaCgccugucugagcgu......................................................................... | 76 | 1 | mv7 |
| .................cuaCgccugucugagcguc........................................................................ | 132 | 1 | mv7 |
| .................cuaAgccugucugagcguc........................................................................ | 2 | 1 | mv7 |
| .................cuauAccugucugagcguc........................................................................ | 1 | 1 | mv7 |
| .................Guaugccugucugagcguc........................................................................ | 1 | 1 | mv7 |
| .................cuaCgccugucugagcgucg....................................................................... | 16 | 1 | mv7 |
| .................cuaAgccugucugagcgucg....................................................................... | 1 | 1 | mv7 |
| .................cuaugccugucugagcgucgc...................................................................... | 1 | 0 | mv7 |
| .................cuaugccugucugagcgCcgc...................................................................... | 1 | 1 | mv7 |
| .................cuaGgccugucugagcgucgc...................................................................... | 1 | 1 | mv7 |
| .................cuaCgccugucugagcgucgc...................................................................... | 327 | 1 | mv7 |
| .................cuaAgccugucugagcgucgc...................................................................... | 5 | 1 | mv7 |
| .................cuaAgccugucugagcgucgcu..................................................................... | 1051 | 1 | mv7 |

```
                    Mature                                                      Star
cuggguuccucccagggcuaugccugucugagcgucgcuugccgaucaaaauuccccagggguugccucugggcuccuuggggugcccagcuguucuguggcagggccc
.................cuaGgccugucugagcgucgcu......................................................................  85      1  mv7
.................cuaugAcugucugagcgucgcu......................................................................  1       1  mv7
.................Guaugccugucugagcgucgcu......................................................................  1       1  mv7
.................cuaugccugucuUagcgucgcu......................................................................  1       1  mv7
.................cuaugccugucugagcgucgcu......................................................................  156     0  mv7
.................cuaugccuAucugagcgucgcu......................................................................  1       1  mv7
.................cAaugccugucugagcgucgcu......................................................................  1       1  mv7
.................cuaugccuguUugagcgucgcu......................................................................  1       1  mv7
.................cuaugccugucugagcgucgcC......................................................................  18      1  mv7
.................cCaugccugucugagcgucgcu......................................................................  1       1  mv7
.................cuaugccugAcugagcgucgcu......................................................................  1       1  mv7
.................cuaugccugucugagcgucgcG......................................................................  4       1  mv7
.................Uuaugccugucugagcgucgcu......................................................................  8       1  mv7
.................cuaugccugucugGgcgucgcu......................................................................  1       1  mv7
.................cuGugccugucugagcgucgcu......................................................................  2       1  mv7
.................cuaugccugCcugagcgucgcu......................................................................  1       1  mv7
.................cuaCgccugucugagcgucgcu......................................................................  41664   1  mv7
.................Auaugccugucugagcgucgcu......................................................................  10      1  mv7
.................cuaugccugucugagcgucgcA......................................................................  6       1  mv7
.................cuaugccugucugagUgucgcu......................................................................  1       1  mv7
.................cuaAgccugucugagcgucgcuu.....................................................................  83      1  mv7
.................cuaugccugucugagcguUgcuu.....................................................................  1       1  mv7
.................cuaugccugucAgagcgucgcuu.....................................................................  1       1  mv7
.................cCaugccugucugagcgucgcuu.....................................................................  1       1  mv7
.................cuaugccugucugagcgucgcuC.....................................................................  3       1  mv7
.................Auaugccugucugagcgucgcuu.....................................................................  1       1  mv7
.................cuaugccugucugagcgucgcuu.....................................................................  27      0  mv7
.................cuaugccugucugagcgucgcuA.....................................................................  1       1  mv7
.................cuaugccugucugagcgucgcCu.....................................................................  1       1  mv7
.................cuaugccugucugagcgCcgcuu.....................................................................  1       1  mv7
.................cuaugccugucugagcgucgcuG.....................................................................  2       1  mv7
.................cuaCgccugucugagcgucgcuu.....................................................................  4288    1  mv7
.................cuaugccuguAugagcgucgcuu.....................................................................  1       1  mv7
.................cuaGgccugucugagcgucgcuu.....................................................................  10      1  mv7
.................cuaCgccugucugagcgucgcuug....................................................................  9       1  mv7
.................cuaCgccugucugagcgucgcuugc...................................................................  48      1  mv7
.................cuaGgccugucugagcgucgcuugc...................................................................  1       1  mv7
.................cuaCgccugucugagcgucgcuugcc..................................................................  4       1  mv7
.................cuaCgccugucugagcgucgcuugccg.................................................................  11      1  mv7
.................cuaAgccugucugagcgucgcuugccg.................................................................  1       1  mv7
.................cuaCgccugucugagcgucgcuugccga................................................................  16      1  mv7
.................cuaugccugucugagcgucgcuugccga................................................................  1       0  mv7
.................cuaCgccugucugagcgucgcuugccgau...............................................................  15      1  mv7
.................cuaCgccugucugagcgucgcuugccgauc..............................................................  10      1  mv7
.................cuaAgccugucugagcgucgcuugccgauc..............................................................  1       1  mv7
.................cuaCgccugucugagcgucgcuugccgauca.............................................................  7       1  mv7
.................cuaCgccugucugagcgucgcuugccgaucaa............................................................  5       1  mv7
.................cuaAgccugucugagcgucgcuugccgaucaa............................................................  1       1  mv7
..................uaCgccugucugagcguc.........................................................................  624     1  mv7
..................uaugccugucugagcguc.........................................................................  4       0  mv7
..................uaAgccugucugagcguc.........................................................................  1       1  mv7
..................uaugccugucugagcgCc.........................................................................  1       1  mv7
..................Caugccugucugagcguc.........................................................................  1       1  mv7
..................uaGgccugucugagcguc.........................................................................  9       1  mv7
..................uaugccugucugagcguU.........................................................................  1       1  mv7
..................uaugccugucugGgcgucg........................................................................  1       1  mv7
..................uaugccugucugagcgucg........................................................................  5       0  mv7
..................uaGgccugucugagcgucg........................................................................  1       1  mv7
..................uaAgccugucugagcgucg........................................................................  1       1  mv7
..................uaCgccugucugagcgucg........................................................................  563     1  mv7
..................uaGgccugucugagcgucgc.......................................................................  10      1  mv7
..................uaugccugucugagcgucgU.......................................................................  2       1  mv7
..................uaugccugucugagcgucgc.......................................................................  4       0  mv7
..................uaCgccugucugagcgucgc.......................................................................  1939    1  mv7
..................uaugcUugucugagcgucgc.......................................................................  1       1  mv7
..................uaAgccugucugagcgucgc.......................................................................  10      1  mv7
..................uaNgccugucugagcgucgc.......................................................................  1       1  mv7
..................uaCgccugucugagcgucgcu......................................................................  236239  1  mv7
..................uaugccugucugagGgucgcu......................................................................  1       1  mv7
..................uaNgccugucugagcgucgcu......................................................................  9       1  mv7
```

```
                    Mature                                                 Star
cuggguuccucccagggcuaugccugucugagcgucgcuugccgaucaaaauuccccaggguugccucugggcuccuuggggugcccagcuguucuguggcagggccc
..................uaugccugucugagcguUgcu.....................................................................  2      1  mv7
..................uauCccugucugagcgucgcu.....................................................................  9      1  mv7
..................uaugccugucugUgcgucgcu.....................................................................  2      1  mv7
..................uaugccugucugagcgucgcG.....................................................................  23     1  mv7
..................uaugccuAucugagcgucgcu.....................................................................  2      1  mv7
..................uaugcUugucugagcgucgcu.....................................................................  3      1  mv7
..................uaugccugucugagcgCcgcu.....................................................................  1      1  mv7
..................uaugccCgucugagcgucgcu.....................................................................  1      1  mv7
..................Aaugccugucugagcgucgcu.....................................................................  5      1  mv7
..................uaugccugucugagcUucgcu.....................................................................  2      1  mv7
..................uaugccugucugagcgucgUu.....................................................................  2      1  mv7
..................uaugccuguUugagcgucgcu.....................................................................  2      1  mv7
..................uaugccugucugagcAucgcu.....................................................................  2      1  mv7
..................uaugccugucCgagcgucgcu.....................................................................  1      1  mv7
..................uCugccugucugagcgucgcu.....................................................................  3      1  mv7
..................uaugccugucugagcgucgcu.....................................................................  739    0  mv7
..................uaugccugucugagcCucgcu.....................................................................  1      1  mv7
..................uauAccugucugagcgucgcu.....................................................................  9      1  mv7
..................uaugccGgucugagcgucgcu.....................................................................  1      1  mv7
..................uaugccugucugagcguAgcu.....................................................................  1      1  mv7
..................uaugccugucugagUgucgcu.....................................................................  3      1  mv7
..................uaAgccugucugagcgucgcu.....................................................................  653    1  mv7
..................uaGgccugucugagcgucgcu.....................................................................  1513   1  mv7
..................uaugccugucugCgcgucgcu.....................................................................  7      1  mv7
..................uaugccugucugagcgucUcu.....................................................................  2      1  mv7
..................Caugccugucugagcgucgcu.....................................................................  3      1  mv7
..................uaugccugucugagcgucgAu.....................................................................  1      1  mv7
..................uaugccugucugagAgucgcu.....................................................................  5      1  mv7
..................uaugccugucugaUcgucgcu.....................................................................  1      1  mv7
..................uaugccugucugaAcgucgcu.....................................................................  2      1  mv7
..................uaugccugucAgagcgucgcu.....................................................................  1      1  mv7
..................uaugccugCcugagcgucgcu.....................................................................  1      1  mv7
..................uaugccugucugagcgucgcA.....................................................................  36     1  mv7
..................Naugccugucugagcgucgcu.....................................................................  1      1  mv7
..................uGugccugucugagcgucgcu.....................................................................  8      1  mv7
..................uaugccugucugagcguGgcu.....................................................................  3      1  mv7
..................uaugccugucugaCcgucgcu.....................................................................  1      1  mv7
..................uaugGcugucugagcgucgcu.....................................................................  1      1  mv7
..................uaugccugucugagcgucgcC.....................................................................  91     1  mv7
..................uaugccugucugUgcgucgcuu....................................................................  1      1  mv7
..................uaugccugucugagcgucUcuu....................................................................  1      1  mv7
..................uaugccugucugaUcgucgcuu....................................................................  1      1  mv7
..................uaugccugucugagcgucgcuG....................................................................  1      1  mv7
..................uCugccugucugagcgucgcuu....................................................................  1      1  mv7
..................uaCgccugucugagcgucgcuu....................................................................  24995  1  mv7
..................uaugccugucugaAcgucgcuu....................................................................  1      1  mv7
..................uauCccugucugagcgucgcuu....................................................................  1      1  mv7
..................Aaugccugucugagcgucgcuu....................................................................  1      1  mv7
..................uaugccugucugCgcgucgcuu....................................................................  1      1  mv7
..................uaGgccugucugagcgucgcuu....................................................................  136    1  mv7
..................uauAccugucugagcgucgcuu....................................................................  1      1  mv7
..................uaugccugucugagcgucgcuu....................................................................  110    0  mv7
..................uaugccugucugagcgucgcCu....................................................................  1      1  mv7
..................uaugccugucugagcgucgcuA....................................................................  3      1  mv7
..................Caugccugucugagcgucgcuu....................................................................  2      1  mv7
..................uaugccugucugagcgucgcuC....................................................................  16     1  mv7
..................uaugccuguAugagcgucgcuu....................................................................  1      1  mv7
..................uaugcUugucugagcgucgcuu....................................................................  3      1  mv7
..................uaAgccugucugagcgucgcuu....................................................................  66     1  mv7
..................uaugccugucugagcgucgcuAg...................................................................  1      1  mv7
..................uaugccugucugagcgucgcuuU...................................................................  1      1  mv7
..................uaNgccugucugagcgucgcuug...................................................................  1      1  mv7
..................uaGgccugucugagcgucgcuug...................................................................  1      1  mv7
..................uaCgccugucugagcgucgcuug...................................................................  115    1  mv7
..................uaCgccugucugagcgucgcuugc..................................................................  238    1  mv7
..................uaAgccugucugagcgucgcuugc..................................................................  1      1  mv7
..................uaCgccugucugagcgucgcuugcc.................................................................  34     1  mv7
..................uaCgccugucugagcgucgcuugccg................................................................  44     1  mv7
..................uaugccugucugagcgucgcuugccga...............................................................  1      0  mv7
..................uaCgccugucugagcgucgcuugccga...............................................................  74     1  mv7
```

```
                    Mature                                                           Star
cuggguuccucccagggcuaugccugucugagcgucgcuugccgaucaaaauuccccaggguugccucugggcuccuuggggugcccagcuguucuguggcagggccc
..................uaGgccugucugagcgucgcuugccgau..............................................................   1        1    mv7
..................uaCgccugucugagcgucgcuugccgau..............................................................   85       1    mv7
..................uaAgccugucugagcgucgcuugccgau..............................................................   2        1    mv7
..................uaCgccugucugagcgucgcuugccgauc.............................................................   50       1    mv7
..................uaCgccugucugagcgucgcuugccgauca............................................................   12       1    mv7
..................uaCgccugucugagcgucgcuugccgaucaa...........................................................   13       1    mv7
..................uaGgccugucugagcgucgcuugccgaucaaa..........................................................   1        1    mv7
..................uaCgccugucugagcgucgcuugccgaucaaa..........................................................   9        1    mv7
...................augccugucugagcgucg.......................................................................   2        0    mv7
...................aCgccugucugagcgucg.......................................................................   625      1    mv7
...................Cugccugucugagcgucg.......................................................................   1        1    mv7
...................aAgccugucugagcgucg.......................................................................   2        1    mv7
...................aGgccugucugagcgucgc......................................................................   2        1    mv7
...................aNgccugucugagcgucgc......................................................................   1        1    mv7
...................aAgccugucugagcgucgc......................................................................   1        1    mv7
...................augccugucugagcgucgU......................................................................   1        1    mv7
...................augccugucugagcgucgc......................................................................   2        0    mv7
...................aCgccugucugagcgucgc......................................................................   1111     1    mv7
...................augccuguUugagcgucgcu.....................................................................   2        1    mv7
...................auAccugucugagcgucgcu.....................................................................   3        1    mv7
...................augccugucugagAgucgcu.....................................................................   1        1    mv7
...................augccugucCgagcgucgcu.....................................................................   1        1    mv7
...................augccugucugagcUucgcu.....................................................................   3        1    mv7
...................Cugccugucugagcgucgcu.....................................................................   22       1    mv7
...................augccugucugagcgucUcu.....................................................................   2        1    mv7
...................augccugucugagcgucgcu.....................................................................   383      0    mv7
...................augccugucugagcgucgcC.....................................................................   53       1    mv7
...................augccugucugagcgucgUu.....................................................................   1        1    mv7
...................auCccugucugagcgucgcu.....................................................................   3        1    mv7
...................augccugCcugagcgucgcu.....................................................................   2        1    mv7
...................augccugucugagcguUgcu.....................................................................   1        1    mv7
...................augccugucugagcgCcgcu.....................................................................   2        1    mv7
...................augccugucugGgcgucgcu.....................................................................   1        1    mv7
...................augccugucugagcgucgcA.....................................................................   17       1    mv7
...................augccGgucugagcgucgcu.....................................................................   1        1    mv7
...................aAgccugucugagcgucgcu.....................................................................   278      1    mv7
...................aNgccugucugagcgucgcu.....................................................................   1        1    mv7
...................augccugucugagcAucgcu.....................................................................   3        1    mv7
...................auUccugucugagcgucgcu.....................................................................   5        1    mv7
...................aCgccugucugagcgucgcu.....................................................................   113087   1    mv7
...................augccuAucugagcgucgcu.....................................................................   1        1    mv7
...................augccugucugagUgucgcu.....................................................................   2        1    mv7
...................aGgccugucugagcgucgcu.....................................................................   152      1    mv7
...................augccCgucugagcgucgcu.....................................................................   2        1    mv7
...................augccugucuUagcgucgcu.....................................................................   1        1    mv7
...................augAcugucugagcgucgcu.....................................................................   21       1    mv7
...................Gugccugucugagcgucgcu.....................................................................   2        1    mv7
...................augcUugucugagcgucgcu.....................................................................   1        1    mv7
...................augccugucugagcgucgcG.....................................................................   6        1    mv7
...................augccuCucugagcgucgcu.....................................................................   1        1    mv7
...................Gugccugucugagcgucgcuu....................................................................   2        1    mv7
...................aGgccugucugagcgucgcuu....................................................................   11       1    mv7
...................auCccugucugagcgucgcuu....................................................................   1        1    mv7
...................augccCgucugagcgucgcuu....................................................................   2        1    mv7
...................auUccugucugagcgucgcuu....................................................................   3        1    mv7
...................augccugucugagcgucgcuu....................................................................   65       0    mv7
...................augAcugucugagcgucgcuu....................................................................   2        1    mv7
...................augccugucugagcgucgcuA....................................................................   1        1    mv7
...................auAccugucugagcgucgcuu....................................................................   1        1    mv7
...................augccGgucugagcgucgcuu....................................................................   1        1    mv7
...................aCgccugucugagcgucgcuu....................................................................   13635    1    mv7
...................augccAgucugagcgucgcuu....................................................................   1        1    mv7
...................augccugucCgagcgucgcuu....................................................................   1        1    mv7
...................aAgccugucugagcgucgcuu....................................................................   25       1    mv7
...................augccugucugagcgucgcuC....................................................................   6        1    mv7
...................aCgccugucugagcgucgcuug...................................................................   43       1    mv7
...................Cugccugucugagcgucgcuugc..................................................................   1        1    mv7
...................aCgccugucugagcgucgcuugc..................................................................   114      1    mv7
...................augccugucugagcgucgcuugc..................................................................   1        0    mv7
...................aCgccugucugagcgucgcuugcc.................................................................   17       1    mv7
```

Mature  Star

| | | | |
|---|---|---|---|
| cuggguuccucccagggcuaugccugucugagcgucgcuugccgaucaaaauuccccaggguugccucugggcuccuuggggugcccagcuguucuguggcagggccc | | | |
| ...................aCgccugucugagcgucgcuugccg................................................................ | 20 | 1 | mv7 |
| ...................aCgccugucugagcgucgcuugccga............................................................... | 21 | 1 | mv7 |
| ...................aCgccugucugagcgucgcuugccgau.............................................................. | 25 | 1 | mv7 |
| ...................aCgccugucugagcgucgcuugccgauc............................................................. | 15 | 1 | mv7 |
| ...................aCgccugucugagcgucgcuugccgauca............................................................ | 5 | 1 | mv7 |
| ...................aCgccugucugagcgucgcuugccgaucaa........................................................... | 6 | 1 | mv7 |
| ...................aCgccugucugagcgucgcuugccgaucaaa.......................................................... | 7 | 1 | mv7 |
| ...................aCgccugucugagcgucgcuugccgaucaaaa......................................................... | 3 | 1 | mv7 |
| ....................ugccugucugGgcgucgc...................................................................... | 1 | 1 | mv7 |
| ....................ugccugucugagAgucgc...................................................................... | 2 | 1 | mv7 |
| ....................ugccugucugagUgucgc...................................................................... | 3 | 1 | mv7 |
| ....................ugcGugucugagcgucgc...................................................................... | 3 | 1 | mv7 |
| ....................ugccuguUugagcgucgc...................................................................... | 1 | 1 | mv7 |
| ....................ugccugucugagcgucgU...................................................................... | 35 | 1 | mv7 |
| ....................ugccugucugagcgucgA...................................................................... | 6 | 1 | mv7 |
| ....................ugccugucugagcgGcgc...................................................................... | 1 | 1 | mv7 |
| ....................Ggccugucugagcgucgc...................................................................... | 14 | 1 | mv7 |
| ....................ugGcugucugagcgucgc...................................................................... | 2 | 1 | mv7 |
| ....................ugcUugucugagcgucgc...................................................................... | 1 | 1 | mv7 |
| ....................uUccugucugagcgucgc...................................................................... | 4 | 1 | mv7 |
| ....................Cgccugucugagcgucgc...................................................................... | 6803 | 1 | mv7 |
| ....................ugccugucugagcgucgG...................................................................... | 1 | 1 | mv7 |
| ....................Ngccugucugagcgucgc...................................................................... | 4 | 1 | mv7 |
| ....................ugcAugucugagcgucgc...................................................................... | 13 | 1 | mv7 |
| ....................Agccugucugagcgucgc...................................................................... | 157 | 1 | mv7 |
| ....................ugccugucugagcgucgc...................................................................... | 119 | 0 | mv7 |
| ....................ugccugucugagcgCcgc...................................................................... | 2 | 1 | mv7 |
| ....................ugccuguGugagcgucgcu..................................................................... | 3 | 1 | mv7 |
| ....................ugccugucugagcguUgcu..................................................................... | 29 | 1 | mv7 |
| ....................ugccuCucugagcgucgcu..................................................................... | 4 | 1 | mv7 |
| ....................ugccugucugagcgucgcC..................................................................... | 859 | 1 | mv7 |
| ....................ugccugucugagcUucgcu..................................................................... | 6 | 1 | mv7 |
| ....................ugccugucCgagcgucgcu..................................................................... | 41 | 1 | mv7 |
| ....................ugccugucugagcgucgcG..................................................................... | 164 | 1 | mv7 |
| ....................ugccugucugagcgucUcu..................................................................... | 7 | 1 | mv7 |
| ....................ugccugucugGgcgucgcu..................................................................... | 40 | 1 | mv7 |
| ....................ugccugucugagcgucgAu..................................................................... | 18 | 1 | mv7 |
| ....................ugAcugucugagcgucgcu..................................................................... | 13 | 1 | mv7 |
| ....................ugccugGcugagcgucgcu..................................................................... | 6 | 1 | mv7 |
| ....................ugccugucugagAgucgcu..................................................................... | 112 | 1 | mv7 |
| ....................ugccGgucugagcgucgcu..................................................................... | 3 | 1 | mv7 |
| ....................uUccugucugagcgucgcu..................................................................... | 18 | 1 | mv7 |
| ....................ugccugucuUagcgucgcu..................................................................... | 13 | 1 | mv7 |
| ....................ugccAgucugagcgucgcu..................................................................... | 7 | 1 | mv7 |
| ....................ugccCgucugagcgucgcu..................................................................... | 14 | 1 | mv7 |
| ....................uAccugucugagcgucgcu..................................................................... | 43 | 1 | mv7 |
| ....................ugccugCcugagcgucgcu..................................................................... | 19 | 1 | mv7 |
| ....................ugccugucugagcgucgcu..................................................................... | 6368 | 0 | mv7 |
| ....................ugccugucugagcgCcgcu..................................................................... | 30 | 1 | mv7 |
| ....................ugccugucugagUgucgcu..................................................................... | 39 | 1 | mv7 |
| ....................ugccugucugagcgucgGu..................................................................... | 14 | 1 | mv7 |
| ....................Ngccugucugagcgucgcu..................................................................... | 179 | 1 | mv7 |
| ....................ugccugucugUgcgucgcu..................................................................... | 11 | 1 | mv7 |
| ....................uCccugucugagcgucgcu..................................................................... | 24 | 1 | mv7 |
| ....................ugccugucugaCcgucgcu..................................................................... | 2 | 1 | mv7 |
| ....................ugccugucuCagcgucgcu..................................................................... | 3 | 1 | mv7 |
| ....................ugcUugucugagcgucgcu..................................................................... | 43 | 1 | mv7 |
| ....................ugccugucugagcgucgUu..................................................................... | 21 | 1 | mv7 |
| ....................ugccugucugagcgucgcA..................................................................... | 310 | 1 | mv7 |
| ....................Ggccugucugagcgucgcu..................................................................... | 721 | 1 | mv7 |
| ....................ugccugucugCgcgucgcu..................................................................... | 16 | 1 | mv7 |
| ....................ugcGugucugagcgucgcu..................................................................... | 143 | 1 | mv7 |
| ....................ugccuguUugagcgucgcu..................................................................... | 12 | 1 | mv7 |
| ....................ugGcugucugagcgucgcu..................................................................... | 166 | 1 | mv7 |
| ....................ugccuAucugagcgucgcu..................................................................... | 7 | 1 | mv7 |
| ....................ugccugucugaAcgucgcu..................................................................... | 9 | 1 | mv7 |
| ....................ugccugucugagcCucgcu..................................................................... | 3 | 1 | mv7 |
| ....................ugccugucugagcAucgcu..................................................................... | 8 | 1 | mv7 |
| ....................ugccugucGgagcgucgcu..................................................................... | 4 | 1 | mv7 |
| ....................Agccugucugagcgucgcu..................................................................... | 8881 | 1 | mv7 |

```
                   Mature                                                         Star
cuggguuccucccagggcuaugccugucugagcgucgcuugccgaucaaaauuccccaggguugccucugggcuccuuggggugcccagcuguucuguggcagggccc
....................Cgccugucugagcgucgcu.....................................................................  319253  1  mv7
....................ugcAugucugagcgucgcu.....................................................................  556     1  mv7
....................ugccugucugagcgucCcu.....................................................................  3       1  mv7
....................ugccuUucugagcgucgcu.....................................................................  27      1  mv7
....................ugccugucugagcguGgcu.....................................................................  3       1  mv7
....................ugccugucAgagcgucgcu.....................................................................  5       1  mv7
....................ugccuguAugagcgucgcu.....................................................................  23      1  mv7
....................ugccugucugaUcgucgcu.....................................................................  9       1  mv7
....................ugccugAcugagcgucgcu.....................................................................  6       1  mv7
....................ugccugucugagcgGcgcu.....................................................................  14      1  mv7
....................ugccugucugCgcgucgcuu....................................................................  1       1  mv7
....................ugccugucugagcgGcgcuu....................................................................  4       1  mv7
....................ugAcugucugagcgucgcuu....................................................................  4       1  mv7
....................ugcAugucugagcgucgcuu....................................................................  77      1  mv7
....................ugccugucugagcgucgcGu....................................................................  2       1  mv7
....................Agccugucugagcgucgcuu....................................................................  1327    1  mv7
....................ugccugucugagcgucgcuA....................................................................  28      1  mv7
....................ugccuguAugagcgucgcuu....................................................................  2       1  mv7
....................ugcGugucugagcgucgcuu....................................................................  17      1  mv7
....................uAccugucugagcgucgcuu....................................................................  9       1  mv7
....................uCccugucugagcgucgcuu....................................................................  6       1  mv7
....................ugccugucugagcUucgcuu....................................................................  1       1  mv7
....................ugccugucCgagcgucgcuu....................................................................  6       1  mv7
....................ugccCgucugagcgucgcuu....................................................................  1       1  mv7
....................ugccugucugagcgCcgcuu....................................................................  1       1  mv7
....................ugccugucugagcguUgcuu....................................................................  5       1  mv7
....................ugccugucugagcgucgUuu....................................................................  1       1  mv7
....................ugccugucugagcAucgcuu....................................................................  1       1  mv7
....................ugccugucugUgcgucgcuu....................................................................  3       1  mv7
....................ugGcugucugagcgucgcuu....................................................................  19      1  mv7
....................ugccugAcugagcgucgcuu....................................................................  1       1  mv7
....................uUccugucugagcgucgcuu....................................................................  6       1  mv7
....................ugccugucugaUcgucgcuu....................................................................  1       1  mv7
....................ugccuguGugagcgucgcuu....................................................................  2       1  mv7
....................ugccugucugagcgucgcuu....................................................................  968     0  mv7
....................ugccuUucugagcgucgcuu....................................................................  4       1  mv7
....................Ngccugucugagcgucgcuu....................................................................  25      1  mv7
....................ugccugucugagcgucgcuC....................................................................  119     1  mv7
....................ugcUugucugagcgucgcuu....................................................................  3       1  mv7
....................ugccugucugagcgucgGuu....................................................................  2       1  mv7
....................ugccugucuCagcgucgcuu....................................................................  1       1  mv7
....................ugccugucugagcguAgcuu....................................................................  3       1  mv7
....................ugccugucugagcgucgcCu....................................................................  2       1  mv7
....................ugccugucugaAcgucgcuu....................................................................  1       1  mv7
....................ugccugucugagcgucgcuG....................................................................  37      1  mv7
....................Cgccugucugagcgucgcuu....................................................................  48360   1  mv7
....................Ggccugucugagcgucgcuu....................................................................  100     1  mv7
....................ugccugucugagcgucUcuu....................................................................  2       1  mv7
....................ugccugucGgagcgucgcuu....................................................................  1       1  mv7
....................ugccugCcugagcgucgcuu....................................................................  6       1  mv7
....................ugccugucugagcguGgcuu....................................................................  1       1  mv7
....................ugccugucugGgcgucgcuu....................................................................  7       1  mv7
....................ugccugucuUagcgucgcuu....................................................................  2       1  mv7
....................ugccugucugagUgucgcuu....................................................................  6       1  mv7
....................ugccugucugagAgucgcuu....................................................................  17      1  mv7
....................ugccuAucugagcgucgcuu....................................................................  1       1  mv7
....................ugccugucugagcgucgcuug...................................................................  4       0  mv7
....................Cgccugucugagcgucgcuug...................................................................  164     1  mv7
....................Agccugucugagcgucgcuug...................................................................  9       1  mv7
....................Cgccugucugagcgucgcuugc..................................................................  343     1  mv7
....................ugccugucugagAgucgcuugc..................................................................  1       1  mv7
....................ugccugucugagcgucgcuugA..................................................................  1       1  mv7
....................Ggccugucugagcgucgcuugc..................................................................  2       1  mv7
....................Agccugucugagcgucgcuugc..................................................................  12      1  mv7
....................ugccugucugagcgucgcuugc..................................................................  1       0  mv7
....................ugccugucugagcgucgcuuCc..................................................................  1       1  mv7
....................ugccugucugagcgucgcuugcA.................................................................  1       1  mv7
....................ugccugucugagcgucgcuugcc.................................................................  2       0  mv7
....................ugcAugucugagcgucgcuugcc.................................................................  1       1  mv7
....................Cgccugucugagcgucgcuugcc.................................................................  32      1  mv7
```

Mature                                                              Star

| | | | |
|---|---|---|---|
| cuggguuccucccagggcuaugccugucugagcgucgcuugccgaucaaaauuccccaggguugccucugggcuccuuggggugcccagcuguucuguggcagggccc | | | |
| ....................Agccugucugagcgucgcuugcc................................................................. | 1 | 1 | mv7 |
| ....................ugccugucugagcgucgcuugcU................................................................. | 1 | 1 | mv7 |
| ....................Cgccugucugagcgucgcuugccg................................................................ | 36 | 1 | mv7 |
| ....................Agccugucugagcgucgcuugccg................................................................ | 2 | 1 | mv7 |
| ....................ugccugucugagcgucgcuugccg................................................................ | 2 | 0 | mv7 |
| ....................Agccugucugagcgucgcuugccga............................................................... | 1 | 1 | mv7 |
| ....................Cgccugucugagcgucgcuugccga............................................................... | 43 | 1 | mv7 |
| ....................ugccugucugagcgucgcuugccgaA.............................................................. | 1 | 1 | mv7 |
| ....................Agccugucugagcgucgcuugccgau.............................................................. | 1 | 1 | mv7 |
| ....................Cgccugucugagcgucgcuugccgau.............................................................. | 39 | 1 | mv7 |
| ....................ugGcugucugagcgucgcuugccgau.............................................................. | 1 | 1 | mv7 |
| ....................Agccugucugagcgucgcuugccgauc............................................................. | 1 | 1 | mv7 |
| ....................Cgccugucugagcgucgcuugccgauc............................................................. | 37 | 1 | mv7 |
| ....................ugccugucugagcgucgcuugccgauA............................................................. | 1 | 1 | mv7 |
| ....................Agccugucugagcgucgcuugccgauca............................................................ | 2 | 1 | mv7 |
| ....................Cgccugucugagcgucgcuugccgauca............................................................ | 17 | 1 | mv7 |
| ....................Cgccugucugagcgucgcuugccgaucaa........................................................... | 10 | 1 | mv7 |
| ....................Cgccugucugagcgucgcuugccgaucaaa.......................................................... | 18 | 1 | mv7 |
| ....................Cgccugucugagcgucgcuugccgaucaaaa......................................................... | 7 | 1 | mv7 |
| ....................Agccugucugagcgucgcuugccgaucaaaa......................................................... | 2 | 1 | mv7 |
| .....................gccugGcugagcgucgcu..................................................................... | 143 | 1 | mv7 |
| .....................gccuAucugagcgucgcu..................................................................... | 32 | 1 | mv7 |
| .....................gccAgucugagcgucgcu..................................................................... | 234 | 1 | mv7 |
| .....................gccugucugagcguUgcu..................................................................... | 81 | 1 | mv7 |
| .....................gccugucugagcgucgcu..................................................................... | 35057 | 0 | mv7 |
| .....................gccugucCgagcgucgcu..................................................................... | 120 | 1 | mv7 |
| .....................gccugucugagcgGcgcu..................................................................... | 47 | 1 | mv7 |
| .....................gccGgucugagcgucgcu..................................................................... | 4 | 1 | mv7 |
| .....................gccugucugagcgucUcu..................................................................... | 86 | 1 | mv7 |
| .....................gcGugucugagcgucgcu..................................................................... | 299 | 1 | mv7 |
| .....................gccugucugaUcgucgcu..................................................................... | 31 | 1 | mv7 |
| .....................gccugucugagcgucgAu..................................................................... | 159 | 1 | mv7 |
| .....................gccuguGugagcgucgcu..................................................................... | 34 | 1 | mv7 |
| .....................gcNugucugagcgucgcu..................................................................... | 1 | 1 | mv7 |
| .....................gccCgucugagcgucgcu..................................................................... | 238 | 1 | mv7 |
| .....................gccugucuUagcgucgcu..................................................................... | 24 | 1 | mv7 |
| .....................gccugucugagcCucgcu..................................................................... | 168 | 1 | mv7 |
| .....................gccuNucugagcgucgcu..................................................................... | 3 | 1 | mv7 |
| .....................gcAugucugagcgucgcu..................................................................... | 124 | 1 | mv7 |
| .....................gccuguAugagcgucgcu..................................................................... | 96 | 1 | mv7 |
| .....................gccugucugagcgucgcA..................................................................... | 1552 | 1 | mv7 |
| .....................gccugucugGgcgucgcu..................................................................... | 136 | 1 | mv7 |
| .....................gccugucugaCcgucgcu..................................................................... | 18 | 1 | mv7 |
| .....................gccugucugagcUucgcu..................................................................... | 41 | 1 | mv7 |
| .....................gccugucugagcgucgUu..................................................................... | 133 | 1 | mv7 |
| .....................gccugAcugagcgucgcu..................................................................... | 50 | 1 | mv7 |
| .....................gccugucugagcgucCcu..................................................................... | 15 | 1 | mv7 |
| .....................gccuCucugagcgucgcu..................................................................... | 5 | 1 | mv7 |
| .....................gccuguUugagcgucgcu..................................................................... | 89 | 1 | mv7 |
| .....................gccugucugagUgucgcu..................................................................... | 118 | 1 | mv7 |
| .....................gccugucugagcgucgcG..................................................................... | 861 | 1 | mv7 |
| .....................Cccugucugagcgucgcu..................................................................... | 598 | 1 | mv7 |
| .....................gccugucugagGgucgcu..................................................................... | 8 | 1 | mv7 |
| .....................gccuUucugagcgucgcu..................................................................... | 15 | 1 | mv7 |
| .....................gccugucGgagcgucgcu..................................................................... | 11 | 1 | mv7 |
| .....................gccugucugCgcgucgcu..................................................................... | 62 | 1 | mv7 |
| .....................gccugucuCagcgucgcu..................................................................... | 69 | 1 | mv7 |
| .....................gccugucAgagcgucgcu..................................................................... | 50 | 1 | mv7 |
| .....................gccugucugagcAucgcu..................................................................... | 54 | 1 | mv7 |
| .....................gccugucugagcgucgcC..................................................................... | 4787 | 1 | mv7 |
| .....................Accugucugagcgucgcu..................................................................... | 348 | 1 | mv7 |
| .....................gAcugucugagcgucgcu..................................................................... | 115 | 1 | mv7 |
| .....................gccugucugagcgucgGu..................................................................... | 80 | 1 | mv7 |
| .....................gccugucugUgcgucgcu..................................................................... | 56 | 1 | mv7 |
| .....................Nccugucugagcgucgcu..................................................................... | 19 | 1 | mv7 |
| .....................gccugucugagAgucgcu..................................................................... | 24 | 1 | mv7 |
| .....................gcUugucugagcgucgcu..................................................................... | 100 | 1 | mv7 |
| .....................gccugucugaAcgucgcu..................................................................... | 52 | 1 | mv7 |
| .....................gGcugucugagcgucgcu..................................................................... | 83 | 1 | mv7 |
| .....................gccugucugagcguGgcu..................................................................... | 134 | 1 | mv7 |

**Mature** **Star**

| | | | |
|---|---|---|---|
| cuggguuccucccagggcuaugccugucugagcgucgcuugccgaucaaaauuccccaggguugccucugggcuccuuggggugcccagcuguucuguggcagggccc | | | |
| .....................gccugCcugagcgucgcu..................................................................... | 133 | 1 | mv7 |
| .....................Uccugucugagcgucgcu..................................................................... | 486 | 1 | mv7 |
| .....................gccugucugagUgucgcuu.................................................................... | 9 | 1 | mv7 |
| .....................gAcugucugagcgucgcuu.................................................................... | 33 | 1 | mv7 |
| .....................gccugGcugagcgucgcuu.................................................................... | 29 | 1 | mv7 |
| .....................gccugucugagcUucgcuu.................................................................... | 4 | 1 | mv7 |
| .....................gccugucugagcgucgUuu.................................................................... | 14 | 1 | mv7 |
| .....................gccugucugagcgucgcuA.................................................................... | 166 | 1 | mv7 |
| .....................gcAugucugagcgucgcuu.................................................................... | 23 | 1 | mv7 |
| .....................gccugucugaCcgucgcuu.................................................................... | 2 | 1 | mv7 |
| .....................gcGugucugagcgucgcuu.................................................................... | 57 | 1 | mv7 |
| .....................gccugCcugagcgucgcuu.................................................................... | 20 | 1 | mv7 |
| .....................gccugucugagcgucCcuu.................................................................... | 5 | 1 | mv7 |
| .....................gccugucugUgcgucgcuu.................................................................... | 15 | 1 | mv7 |
| .....................gccugucugagcgCcgcuu.................................................................... | 27 | 1 | mv7 |
| .....................gccNgucugagcgucgcuu.................................................................... | 2 | 1 | mv7 |
| .....................gccugucugagAgucgcuu.................................................................... | 6 | 1 | mv7 |
| .....................gccuAucugagcgucgcuu.................................................................... | 9 | 1 | mv7 |
| .....................gccugucGgagcgucgcuu.................................................................... | 5 | 1 | mv7 |
| .....................gccugucugagcgucgcuu.................................................................... | 7432 | 0 | mv7 |
| .....................gccugucugCgcgucgcuu.................................................................... | 17 | 1 | mv7 |
| .....................gccugucugagcgucgcGu.................................................................... | 44 | 1 | mv7 |
| .....................gccugucugagcgucgcuG.................................................................... | 91 | 1 | mv7 |
| .....................gccugucugaAcgucgcuu.................................................................... | 9 | 1 | mv7 |
| .....................gccugucugGgcgucgcuu.................................................................... | 26 | 1 | mv7 |
| .....................gccugucugagcgucgAuu.................................................................... | 36 | 1 | mv7 |
| .....................gccugucugagcgGcgcuu.................................................................... | 10 | 1 | mv7 |
| .....................gccuguAugagcgucgcuu.................................................................... | 23 | 1 | mv7 |
| .....................gccugucugagcgucgcAu.................................................................... | 12 | 1 | mv7 |
| .....................gccugucugagGgucgcuu.................................................................... | 2 | 1 | mv7 |
| .....................gccugucugaUcgucgcuu.................................................................... | 2 | 1 | mv7 |
| .....................gccugucCgagcgucgcuu.................................................................... | 26 | 1 | mv7 |
| .....................gccugucugagcguGgcuu.................................................................... | 23 | 1 | mv7 |
| .....................gccugucuUagcgucgcuu.................................................................... | 5 | 1 | mv7 |
| .....................gccAgucugagcgucgcuu.................................................................... | 55 | 1 | mv7 |
| .....................gccugucugagcgucUcuu.................................................................... | 17 | 1 | mv7 |
| .....................gccugucugagcguUgcuu.................................................................... | 16 | 1 | mv7 |
| .....................gccCgucugagcgucgcuu.................................................................... | 48 | 1 | mv7 |
| .....................gccugucuCagcgucgcuu.................................................................... | 12 | 1 | mv7 |
| .....................gccugucugagcguAgcuu.................................................................... | 40 | 1 | mv7 |
| .....................gccGgucugagcgucgcuu.................................................................... | 2 | 1 | mv7 |
| .....................gccugAcugagcgucgcuu.................................................................... | 10 | 1 | mv7 |
| .....................gccugucugagcCucgcuu.................................................................... | 37 | 1 | mv7 |
| .....................gccugucugagcgucgGuu.................................................................... | 11 | 1 | mv7 |
| .....................gGcugucugagcgucgcuu.................................................................... | 11 | 1 | mv7 |
| .....................Accugucugagcgucgcuu.................................................................... | 75 | 1 | mv7 |
| .....................gccugucugagcgucgcuC.................................................................... | 826 | 1 | mv7 |
| .....................Cccugucugagcgucgcuu.................................................................... | 108 | 1 | mv7 |
| .....................gccugucAgagcgucgcuu.................................................................... | 12 | 1 | mv7 |
| .....................gccugucugagcAucgcuu.................................................................... | 5 | 1 | mv7 |
| .....................Nccugucugagcgucgcuu.................................................................... | 3 | 1 | mv7 |
| .....................gcUugucugagcgucgcuu.................................................................... | 31 | 1 | mv7 |
| .....................gccuguGugagcgucgcuu.................................................................... | 12 | 1 | mv7 |
| .....................gccuguUugagcgucgcuu.................................................................... | 8 | 1 | mv7 |
| .....................Uccugucugagcgucgcuu.................................................................... | 101 | 1 | mv7 |
| .....................gccugucugagcgucgcCu.................................................................... | 49 | 1 | mv7 |
| .....................gccuUucugagcgucgcuu.................................................................... | 1 | 1 | mv7 |
| .....................gccugucugagcgucgcAug................................................................... | 1 | 1 | mv7 |
| .....................gccugucugagcgucgcuug................................................................... | 14 | 0 | mv7 |
| .....................gccugucugagcgucgcuuA................................................................... | 2 | 1 | mv7 |
| .....................gccugCcugagcgucgcuug................................................................... | 1 | 1 | mv7 |
| .....................gAcugucugagcgucgcuug................................................................... | 1 | 1 | mv7 |
| .....................gccugucugagcgucgcuuU................................................................... | 29 | 1 | mv7 |
| .....................gccugucugagcgucgcuAg................................................................... | 2 | 1 | mv7 |
| .....................gccugucugagcgucgcuuC................................................................... | 4 | 1 | mv7 |
| .....................gccugucugagcgucgcuGg................................................................... | 1 | 1 | mv7 |
| .....................gccCgucugagcgucgcuugc.................................................................. | 1 | 1 | mv7 |
| .....................gccugucugagcgucgcuugA.................................................................. | 4 | 1 | mv7 |
| .....................gccugucugagcgucgcuuUc.................................................................. | 1 | 1 | mv7 |
| .....................gccugucugagcgucgcuugU.................................................................. | 6 | 1 | mv7 |

**Mature** **Star**

| cuggguuccucccagggcuaugccugucugagcgucgcuugccgaucaaaauuccccaggguugccucugggcuccuugggggugcccagcuguucuguggcagggccc | | | |
|---|---|---|---|
| .....................gccugucugagcgucgcuugc.................................................................. | 18 | 0 | mv7 |
| .....................gccugucugagcCucgcuugc.................................................................. | 1 | 1 | mv7 |
| .....................gccugucugagcgucgcuugcc................................................................. | 6 | 0 | mv7 |
| .....................gccugucugagcgucgcuugcA................................................................. | 2 | 1 | mv7 |
| .....................gccugucugagcgucgAuugcc................................................................. | 1 | 1 | mv7 |
| .....................gccugucugagcgucgcuugccA................................................................ | 1 | 1 | mv7 |
| .....................gccugucugagcgucgcuugccgG............................................................... | 1 | 1 | mv7 |
| .....................gccugucugagcgucgcuugccga............................................................... | 1 | 0 | mv7 |
| .....................gccugucugCgcgucgcuugccga............................................................... | 1 | 1 | mv7 |
| .....................gccugucugagcgucgcuugccgau.............................................................. | 8 | 0 | mv7 |
| .....................gccuguGugagcgucgcuugccgau.............................................................. | 1 | 1 | mv7 |
| .....................gccugucugagcgucgcuugccgaA.............................................................. | 1 | 1 | mv7 |
| .....................gccugucugGgcgucgcuugccgauc............................................................. | 1 | 1 | mv7 |
| .....................gccugucugagcgucgcuugccgauc............................................................. | 6 | 0 | mv7 |
| .....................gccugucugagcgucgcuugccgauA............................................................. | 1 | 1 | mv7 |
| .....................gccugucugagcgucgGuugccgauc............................................................. | 1 | 1 | mv7 |
| .....................gccugucugagcgucgcuugccgauAa............................................................ | 1 | 1 | mv7 |
| .....................gccCgucugagcgucgcuugccgauca............................................................ | 1 | 1 | mv7 |
| .....................gcGugucugagcgucgcuugccgauca............................................................ | 1 | 1 | mv7 |
| .....................gccugucugagcgucgcuugccgauca............................................................ | 3 | 0 | mv7 |
| .....................gccugucugagcgucgcuugccgaucaa........................................................... | 1 | 0 | mv7 |
| .....................gccugucugagcgucgcuugccgaucaaU.......................................................... | 3 | 1 | mv7 |
| .....................gcGugucugagcgucgcuugccgaucaaaa......................................................... | 1 | 1 | mv7 |
| ......................ccugucugagcgucgcuu.................................................................... | 113818 | 0 | mv7 |
| ......................ccugucuNagcgucgcuu.................................................................... | 1 | 1 | mv7 |
| ......................ccCgucugagcgucgcuu.................................................................... | 394 | 1 | mv7 |
| ......................ccGgucugagcgucgcuu.................................................................... | 816 | 1 | mv7 |
| ......................ccugucugaAcgucgcuu.................................................................... | 148 | 1 | mv7 |
| ......................cAugucugagcgucgcuu.................................................................... | 260 | 1 | mv7 |
| ......................ccugucugagUgucgcuu.................................................................... | 258 | 1 | mv7 |
| ......................Gcugucugagcgucgcuu.................................................................... | 263 | 1 | mv7 |
| ......................ccugucugagcgucgGuu.................................................................... | 67 | 1 | mv7 |
| ......................ccugucugagcgucgcuC.................................................................... | 12936 | 1 | mv7 |
| ......................cNugucugagcgucgcuu.................................................................... | 1 | 1 | mv7 |
| ......................ccugucugagcgCcgcuu.................................................................... | 727 | 1 | mv7 |
| ......................ccugucugagcgucUcuu.................................................................... | 556 | 1 | mv7 |
| ......................ccugAcugagcgucgcuu.................................................................... | 93 | 1 | mv7 |
| ......................Acugucugagcgucgcuu.................................................................... | 3098 | 1 | mv7 |
| ......................ccugNcugagcgucgcuu.................................................................... | 7 | 1 | mv7 |
| ......................ccugucAgagcgucgcuu.................................................................... | 260 | 1 | mv7 |
| ......................cUugucugagcgucgcuu.................................................................... | 189 | 1 | mv7 |
| ......................ccuguGugagcgucgcuu.................................................................... | 16 | 1 | mv7 |
| ......................ccugucugUgcgucgcuu.................................................................... | 184 | 1 | mv7 |
| ......................ccuguNugagcgucgcuu.................................................................... | 1 | 1 | mv7 |
| ......................ccugucuUagcgucgcuu.................................................................... | 44 | 1 | mv7 |
| ......................Ncugucugagcgucgcuu.................................................................... | 54 | 1 | mv7 |
| ......................ccugGcugagcgucgcuu.................................................................... | 27 | 1 | mv7 |
| ......................ccuguAugagcgucgcuu.................................................................... | 87 | 1 | mv7 |
| ......................ccuAucugagcgucgcuu.................................................................... | 855 | 1 | mv7 |
| ......................ccuNucugagcgucgcuu.................................................................... | 4 | 1 | mv7 |
| ......................ccugucugagcgucgUuu.................................................................... | 187 | 1 | mv7 |
| ......................ccugucuCagcgucgcuu.................................................................... | 41 | 1 | mv7 |
| ......................ccuCucugagcgucgcuu.................................................................... | 374 | 1 | mv7 |
| ......................ccugucugagGgucgcuu.................................................................... | 63 | 1 | mv7 |
| ......................ccugCcugagcgucgcuu.................................................................... | 334 | 1 | mv7 |
| ......................ccugucugagcgucgcuG.................................................................... | 1617 | 1 | mv7 |
| ......................ccugucugGgcgucgcuu.................................................................... | 459 | 1 | mv7 |
| ......................ccNgucugagcgucgcuu.................................................................... | 4 | 1 | mv7 |
| ......................ccugucugagcgucgcuA.................................................................... | 3059 | 1 | mv7 |
| ......................ccugucGgagcgucgcuu.................................................................... | 125 | 1 | mv7 |
| ......................ccugucugagcUucgcuu.................................................................... | 32 | 1 | mv7 |
| ......................ccugucugCgcgucgcuu.................................................................... | 446 | 1 | mv7 |
| ......................ccugucugagcAucgcuu.................................................................... | 111 | 1 | mv7 |
| ......................ccugucugaUcgucgcuu.................................................................... | 81 | 1 | mv7 |
| ......................ccugucugagcgucgcGu.................................................................... | 687 | 1 | mv7 |
| ......................ccuUucugagcgucgcuu.................................................................... | 25 | 1 | mv7 |
| ......................ccugucugagAgucgcuu.................................................................... | 178 | 1 | mv7 |
| ......................ccugucugagcgucgAuu.................................................................... | 248 | 1 | mv7 |
| ......................ccugucugagcguUgcuu.................................................................... | 175 | 1 | mv7 |
| ......................cGugucugagcgucgcuu.................................................................... | 160 | 1 | mv7 |

**Mature** **Star**

| | | | |
|---|---|---|---|
| cuggguuccucccagggcuaugccugucugagcgucgcuugccgaucaaaauuccccaggguugccucugggcuccuuggggugcccagcuguucuguggcagggccc | | | |
| ......................ccugucugagcgGcgcuu.................................................................... | 72 | 1 | mv7 |
| ......................ccugucugagcgucgcAu.................................................................... | 239 | 1 | mv7 |
| ......................ccugucugagcguAgcuu.................................................................... | 41 | 1 | mv7 |
| ......................ccuguUugagcgucgcuu.................................................................... | 190 | 1 | mv7 |
| ......................ccugucCgagcgucgcuu.................................................................... | 359 | 1 | mv7 |
| ......................ccugucugagcguGgcuu.................................................................... | 29 | 1 | mv7 |
| ......................ccugucugagcCucgcuu.................................................................... | 15 | 1 | mv7 |
| ......................ccugucugagcgucgcCu.................................................................... | 720 | 1 | mv7 |
| ......................ccAgucugagcgucgcuu.................................................................... | 392 | 1 | mv7 |
| ......................ccugucugagcgucCcuu.................................................................... | 216 | 1 | mv7 |
| ......................ccugucugaCcgucgcuu.................................................................... | 56 | 1 | mv7 |
| ......................ccugucugGgcgucgcuug................................................................... | 2 | 1 | mv7 |
| ......................ccAgucugagcgucgcuug................................................................... | 6 | 1 | mv7 |
| ......................ccugucugagUgucgcuug................................................................... | 1 | 1 | mv7 |
| ......................ccugucugagcgucgcuGg................................................................... | 2 | 1 | mv7 |
| ......................Ncugucugagcgucgcuug................................................................... | 1 | 1 | mv7 |
| ......................ccGgucugagcgucgcuug................................................................... | 2 | 1 | mv7 |
| ......................ccugucugagcguAgcuug................................................................... | 1 | 1 | mv7 |
| ......................ccugucugagcgucgcuCg................................................................... | 7 | 1 | mv7 |
| ......................ccugucugUgcgucgcuug................................................................... | 2 | 1 | mv7 |
| ......................ccuguUugagcgucgcuug................................................................... | 1 | 1 | mv7 |
| ......................cGugucugagcgucgcuug................................................................... | 1 | 1 | mv7 |
| ......................ccuAucugagcgucgcuug................................................................... | 1 | 1 | mv7 |
| ......................ccugucugagcgucgUuug................................................................... | 1 | 1 | mv7 |
| ......................ccugucugagcgAcgcuug................................................................... | 1 | 1 | mv7 |
| ......................cAugucugagcgucgcuug................................................................... | 1 | 1 | mv7 |
| ......................ccugCcugagcgucgcuug................................................................... | 1 | 1 | mv7 |
| ......................Gcugucugagcgucgcuug................................................................... | 1 | 1 | mv7 |
| ......................ccugucugagcgucgcuuA................................................................... | 38 | 1 | mv7 |
| ......................ccugucugagcgucgcuAg................................................................... | 1 | 1 | mv7 |
| ......................ccugucugagcgucCcuug................................................................... | 1 | 1 | mv7 |
| ......................ccuguAugagcgucgcuug................................................................... | 1 | 1 | mv7 |
| ......................Acugucugagcgucgcuug................................................................... | 14 | 1 | mv7 |
| ......................ccugucugagcgucgcuuU................................................................... | 185 | 1 | mv7 |
| ......................ccugucugagcgucgcuuC................................................................... | 22 | 1 | mv7 |
| ......................ccugucugaUcgucgcuug................................................................... | 1 | 1 | mv7 |
| ......................ccugucugagcguUgcuug................................................................... | 1 | 1 | mv7 |
| ......................ccugucugagAgucgcuug................................................................... | 1 | 1 | mv7 |
| ......................ccugucuUagcgucgcuug................................................................... | 1 | 1 | mv7 |
| ......................ccugucugagcgucgcuug................................................................... | 335 | 0 | mv7 |
| ......................ccugucugagcgucUcuug................................................................... | 1 | 1 | mv7 |
| ......................ccugucCgagcgucgcuug................................................................... | 2 | 1 | mv7 |
| ......................Gcugucugagcgucgcuugc.................................................................. | 2 | 1 | mv7 |
| ......................cUugucugagcgucgcuugc.................................................................. | 1 | 1 | mv7 |
| ......................ccugCcugagcgucgcuugc.................................................................. | 1 | 1 | mv7 |
| ......................ccugucugagcgucgcuAgc.................................................................. | 1 | 1 | mv7 |
| ......................ccugucugagcgucgcuCgc.................................................................. | 2 | 1 | mv7 |
| ......................ccugucugUgcgucgcuugc.................................................................. | 1 | 1 | mv7 |
| ......................ccugucugagcgucgcCugc.................................................................. | 1 | 1 | mv7 |
| ......................ccugucugagcgucgcuugU.................................................................. | 84 | 1 | mv7 |
| ......................ccugucugaUcgucgcuugc.................................................................. | 1 | 1 | mv7 |
| ......................ccugucugagcAucgcuugc.................................................................. | 1 | 1 | mv7 |
| ......................ccugucugagcgucUcuugc.................................................................. | 1 | 1 | mv7 |
| ......................ccugucugagcgucgcuuUc.................................................................. | 1 | 1 | mv7 |
| ......................ccugucugagcgucgcuGgc.................................................................. | 1 | 1 | mv7 |
| ......................ccugucugagcgCcgcuugc.................................................................. | 2 | 1 | mv7 |
| ......................ccuAucugagcgucgcuugc.................................................................. | 3 | 1 | mv7 |
| ......................ccGgucugagcgucgcuugc.................................................................. | 2 | 1 | mv7 |
| ......................ccugucugagcgucgcuugc.................................................................. | 311 | 0 | mv7 |
| ......................ccugucugagcgucgcAugc.................................................................. | 1 | 1 | mv7 |
| ......................ccugucugagcgucgcuugG.................................................................. | 2 | 1 | mv7 |
| ......................Acugucugagcgucgcuugc.................................................................. | 8 | 1 | mv7 |
| ......................ccugucugagcgucgcuugA.................................................................. | 34 | 1 | mv7 |
| ......................ccugucugagcgucCcuugcc................................................................. | 1 | 1 | mv7 |
| ......................ccugucugagcgucgcuugcc................................................................. | 68 | 0 | mv7 |
| ......................Acugucugagcgucgcuugcc................................................................. | 4 | 1 | mv7 |
| ......................ccugucugagcgucgcuuCcc................................................................. | 1 | 1 | mv7 |
| ......................ccugucugagcgCcgcuugcc................................................................. | 1 | 1 | mv7 |
| ......................ccugucugagcgucgcuugcA................................................................. | 36 | 1 | mv7 |
| ......................ccuguUugagcgucgcuugcc................................................................. | 1 | 1 | mv7 |

```
                     Mature                                                          Star
cuggguuccucccagggcuaugccugucugagcgucgcuugccgaucaaaauuccccaggguugccucugggcuccuuggggugcccagcuguucuguggcagggccc
......................ccGgucugagcgucgcuugcc.................................................................	1	1	mv7
......................ccugucugagcgucgcuuAcc.................................................................	1	1	mv7
......................ccugucugagAgucgcuugcc.................................................................	1	1	mv7
......................ccugucugagcguUgcuugcc.................................................................	1	1	mv7
......................ccugucugagcgucgcuugcU.................................................................	18	1	mv7
......................ccugucugagcgucgcuAgccg................................................................	2	1	mv7
......................Acugucugagcgucgcuugccg................................................................	1	1	mv7
......................ccugucugagcgucgAuugccg................................................................	1	1	mv7
......................ccugucugagUgucgcuugccg................................................................	1	1	mv7
......................ccugucugagcgucgcuCgccg................................................................	1	1	mv7
......................ccugucugagcgucgcuugccg................................................................	65	0	mv7
......................ccugCcugagcgucgcuugccg................................................................	2	1	mv7
......................ccugucugagcgucgcuugccC................................................................	1	1	mv7
......................ccugucugagcgucgcuugccA................................................................	8	1	mv7
......................ccugucugagcgucgcuugcAg................................................................	1	1	mv7
......................ccugucuUagcgucgcuugccga...............................................................	1	1	mv7
......................ccugucugagcgucgcuugccgC...............................................................	1	1	mv7
......................ccugucugagcgucgcuugccgU...............................................................	2	1	mv7
......................ccuAucugagcgucgcuugccga...............................................................	1	1	mv7
......................ccugucugagcgucgcuugccga...............................................................	30	0	mv7
......................ccugucugagcgucAcuugccga...............................................................	2	1	mv7
......................ccugucugagcgucgcuugccgG...............................................................	1	1	mv7
......................ccugucugagcgucgcuugccgaG..............................................................	1	1	mv7
......................ccugucugagcgucgcuugccgaA..............................................................	10	1	mv7
......................Gcugucugagcgucgcuugccgau..............................................................	1	1	mv7
......................ccugucugagcgucgcuugccgaC..............................................................	2	1	mv7
......................Acugucugagcgucgcuugccgau..............................................................	1	1	mv7
......................ccugucugagcgucCcuugccgau..............................................................	1	1	mv7
......................ccugCcugagcgucgcuugccgau..............................................................	1	1	mv7
......................ccugucugagcgucAcuugccgau..............................................................	1	1	mv7
......................ccugucugagcgucgcuugccgau..............................................................	56	0	mv7
......................ccugucugagcgucgcuCgccgauc.............................................................	1	1	mv7
......................ccugucCgagcgucgcuugccgauc.............................................................	1	1	mv7
......................ccugucugagcgucgcuugccgGuc.............................................................	1	1	mv7
......................ccugucugagcgucgcuugccgauU.............................................................	3	1	mv7
......................ccuAucugagcgucgcuugccgauc.............................................................	1	1	mv7
......................ccugucugagcgucAcuugccgauc.............................................................	2	1	mv7
......................cAugucugagcgucgcuugccgauc.............................................................	1	1	mv7
......................ccugucugagcgucgcuugccgauc.............................................................	57	0	mv7
......................ccugucugagcgucgcuugccgauA.............................................................	16	1	mv7
......................Acugucugagcgucgcuugccgauc.............................................................	3	1	mv7
......................ccugucugagcgucgcuugccgauG.............................................................	1	1	mv7
......................ccugucugaUcgucgcuugccgauc.............................................................	1	1	mv7
......................ccugucugagcgucgcuugccgauca............................................................	9	0	mv7
......................ccugucugagcgucgcuugccgaucU............................................................	1	1	mv7
......................ccugucugagcgucgcuugccgauAa............................................................	3	1	mv7
......................ccugucugagcgucgcuugccgauGaa...........................................................	1	1	mv7
......................ccugucugagcgucgcuugccgaucaa...........................................................	10	0	mv7
......................ccugucugagcgucgcuugccgauAaa...........................................................	7	1	mv7
......................ccugucugagcgucgcuugccgaucaaU..........................................................	6	1	mv7
......................ccugucugagUgucgcuugccgaucaaa..........................................................	1	1	mv7
......................ccugucugagcgucgcuugccgaucaaa..........................................................	21	0	mv7
......................ccugucugagcgucgcuugccgauAaaa..........................................................	7	1	mv7
......................ccugucugagcgucgcuugccgaucaaUa.........................................................	12	1	mv7
......................ccugCcugagcgucgcuugccgaucaaaa.........................................................	1	1	mv7
......................ccugucugagcgucgcuugccgaucaaaa.........................................................	8	0	mv7
......................ccugucugagcgucgcuugccgaucaaaG.........................................................	1	1	mv7
......................ccugucugagcgucgcuugccgauAaaaa.........................................................	5	1	mv7
......................ccugucugagcgucgcuugccgaucaaaaA........................................................	12	1	mv7
......................ccugucugagcgucgcuugccgaucaaaauA.......................................................	2	1	mv7
.......................Augucugagcgucgcuug...................................................................	7	1	mv7
.......................cugucugagcgCcgcuug...................................................................	2	1	mv7
.......................cugucugagcgucgcuAg...................................................................	2	1	mv7
.......................cugucCgagcgucgcuug...................................................................	1	1	mv7
.......................cugucugCgcgucgcuug...................................................................	2	1	mv7
.......................cugucuCagcgucgcuug...................................................................	1	1	mv7
.......................cugucugagcgucgcuuU...................................................................	54	1	mv7
.......................cugAcugagcgucgcuug...................................................................	2	1	mv7
.......................cugucugagcgucgcCug...................................................................	1	1	mv7
.......................cugucugagcgucgcuug...................................................................	168	0	mv7
```

```
                     Mature                                          Star
cuggguuccucccagggcuaugccugucugagcgucgcuugccgaucaaaauuccccagggguugccucugggcuccuuggggugcccagcuguucuguggcagggccc
......................cugucugagcguUgcuug.....................................................................  1    1  mv7
......................Gugucugagcgucgcuug.....................................................................  1    1  mv7
......................cugucugagcgucgcuGg.....................................................................  3    1  mv7
......................cGgucugagcgucgcuug.....................................................................  1    1  mv7
......................cuUucugagcgucgcuug.....................................................................  6    1  mv7
......................cugucugagcgucgcuuC.....................................................................  11   1  mv7
......................cugucugagcgucgcuuA.....................................................................  21   1  mv7
......................cugucugagcgucgcGug.....................................................................  2    1  mv7
......................Uugucugagcgucgcuug.....................................................................  4    1  mv7
......................cugucugagcgucgcuCg.....................................................................  4    1  mv7
......................cugucugagcgucgcCugc....................................................................  2    1  mv7
......................cCgucugagcgucgcuugc....................................................................  2    1  mv7
......................Uugucugagcgucgcuugc....................................................................  1    1  mv7
......................cugAcugagcgucgcuugc....................................................................  4    1  mv7
......................cugCcugagcgucgcuugc....................................................................  1    1  mv7
......................cuguUugagcgucgcuugc....................................................................  1    1  mv7
......................cugucugagcgCcgcuugc....................................................................  1    1  mv7
......................cugucugagcgucgcuGgc....................................................................  1    1  mv7
......................cugucugagcgucgcuugU....................................................................  43   1  mv7
......................cuUucugagcgucgcuugc....................................................................  5    1  mv7
......................cugucAgagcgucgcuugc....................................................................  1    1  mv7
......................cugucugagcgucgcuugc....................................................................  190  0  mv7
......................cugucugagcgucgcuuCc....................................................................  1    1  mv7
......................cuCucugagcgucgcuugc....................................................................  1    1  mv7
......................cugucugagcCucgcuugc....................................................................  1    1  mv7
......................cugucugagcgucgAuugc....................................................................  3    1  mv7
......................cugucugagcgucgcuugA....................................................................  17   1  mv7
......................Augucugagcgucgcuugc....................................................................  4    1  mv7
......................cugucCgagcgucgcuugc....................................................................  2    1  mv7
......................cugucugagcgucgcGugc....................................................................  1    1  mv7
......................cugucugagcgucgAuugcc...................................................................  1    1  mv7
......................cCgucugagcgucgcuugcc...................................................................  1    1  mv7
......................cugucugagcgucgcuugcG...................................................................  1    1  mv7
......................cugucugagcgucgcuugAc...................................................................  1    1  mv7
......................cugucugagcgucgcuAgcc...................................................................  1    1  mv7
......................cGgucugagcgucgcuugcc...................................................................  1    1  mv7
......................cuUucugagcgucgcuugcc...................................................................  1    1  mv7
......................cugucuUagcgucgcuugcc...................................................................  1    1  mv7
......................cugucugagcgucgcuugcU...................................................................  6    1  mv7
......................cuAucugagcgucgcuugcc...................................................................  1    1  mv7
......................cugucugagcgucgcuugcA...................................................................  17   1  mv7
......................cugucugagcgucgcuugcc...................................................................  33   0  mv7
......................cugucugagcgucgcCugccg..................................................................  1    1  mv7
......................cugucugagcgucgcuugccA..................................................................  5    1  mv7
......................cugucugagcgucgcuugccg..................................................................  12   0  mv7
......................cugucugagcgucgcuugccU..................................................................  1    1  mv7
......................cugucugagcgucAcuugccg..................................................................  1    1  mv7
......................Augucugagcgucgcuugccg..................................................................  1    1  mv7
......................cugucugagcgucgcuAgccga.................................................................  1    1  mv7
......................cugucugagcguUgcuugccga.................................................................  1    1  mv7
......................cugucugagcgucgcuugccga.................................................................  24   0  mv7
......................Uugucugagcgucgcuugccgau................................................................  1    1  mv7
......................cugucugagcguAgcuugccgau................................................................  1    1  mv7
......................cugucugagcgucgcuugccgaA................................................................  8    1  mv7
......................cugucugagcgucgcuugccgau................................................................  30   0  mv7
......................cugCcugagcgucgcuugccgau................................................................  1    1  mv7
......................cugucugagcAucgcuugccgau................................................................  1    1  mv7
......................Augucugagcgucgcuugccgau................................................................  1    1  mv7
......................cugucugagcgucgcuuAccgau................................................................  1    1  mv7
......................cugAcugagcgucgcuugccgau................................................................  1    1  mv7
......................cugucugagcgucgcuugccgaC................................................................  4    1  mv7
......................Uugucugagcgucgcuugccgauc...............................................................  1    1  mv7
......................cugucugagcgucgcuugccgauU...............................................................  1    1  mv7
......................cugucGgagcgucgcuugccgauc...............................................................  1    1  mv7
......................cugucugagcgucgcuugccgauA...............................................................  16   1  mv7
......................cugucugagcgucgcuugccgauc...............................................................  33   0  mv7
......................cugucugCgcgucgcuugccgauc...............................................................  1    1  mv7
......................cugucugagcgucgGuugccgauc...............................................................  1    1  mv7
......................cugucugagcgucgcuugccgauca..............................................................  5    0  mv7
......................Uugucugagcgucgcuugccgauca..............................................................  1    1  mv7
```

**Mature** **Star**

```
cuggguuccucccagggcuaugccugucugagcgucgcuugccgaucaaaauuccccaggguugccucugggcuccuuggggugcccagcuguucuguggcagggccc
.......................cugucugagcguUgcuugccgauca............................................................ 1 1 mv7
.......................cugucugGgcgucgcuugccgauca............................................................ 1 1 mv7
.......................cugAcugagcgucgcuugccgauca............................................................ 1 1 mv7
.......................cugucugagcgucgcuGgccgaucaa........................................................... 1 1 mv7
.......................cugucugagcgucgcuugccgaucaa........................................................... 6 0 mv7
.......................cugucugagcgucgcuugccgauAaa........................................................... 2 1 mv7
.......................cugucugagcgucgcuugccgaucaaa.......................................................... 3 0 mv7
.......................cugucugagcgucgcuugccgauAaaa.......................................................... 3 1 mv7
.......................cugucugagcgucgcuugccgaucaaC.......................................................... 2 1 mv7
.......................cugucugagcgucgcuugccgaucaaaG......................................................... 2 1 mv7
.......................cugucugagcgucgcuugccgaucaaaa......................................................... 5 0 mv7
.......................cugucugagcgucgcuugccgaucaaaC......................................................... 1 1 mv7
.......................cugucugagcgucgcuugccgauAaaaa......................................................... 4 1 mv7
.......................cugucugagcgucgcuugccgaucaaaaA........................................................ 1 1 mv7
........................ugAcugagcgucgcuugc.................................................................. 2 1 mv7
........................ugucugagcgucgcuuCc.................................................................. 1 1 mv7
........................uguAugagcgucgcuugc.................................................................. 12 1 mv7
........................ugCcugagcgucgcuugc.................................................................. 2 1 mv7
........................ugGcugagcgucgcuugc.................................................................. 3 1 mv7
........................ugucugagcgucgcuugU.................................................................. 92 1 mv7
........................ugucugGgcgucgcuugc.................................................................. 6 1 mv7
........................ugucugagcgucgcuuAc.................................................................. 1 1 mv7
........................Ngucugagcgucgcuugc.................................................................. 1 1 mv7
........................ugucuUagcgucgcuugc.................................................................. 2 1 mv7
........................ugucugagcgucCcuugc.................................................................. 1 1 mv7
........................Cgucugagcgucgcuugc.................................................................. 2 1 mv7
........................uAucugagcgucgcuugc.................................................................. 1 1 mv7
........................ugucugagcgucgcuCgc.................................................................. 2 1 mv7
........................ugucugagcgucgcGugc.................................................................. 2 1 mv7
........................ugucugagcgucgcAugc.................................................................. 2 1 mv7
........................Agucugagcgucgcuugc.................................................................. 1 1 mv7
........................ugucugagcgucgcuugc.................................................................. 360 0 mv7
........................ugucugagcgucgcuugG.................................................................. 4 1 mv7
........................ugucugagUgucgcuugc.................................................................. 1 1 mv7
........................ugucugagcgucgcuugA.................................................................. 42 1 mv7
........................ugucugagcgCcgcuugcc................................................................. 1 1 mv7
........................ugucugagcgucgcuugcc................................................................. 95 0 mv7
........................ugucugagcgGcgcuugcc................................................................. 1 1 mv7
........................Agucugagcgucgcuugcc................................................................. 2 1 mv7
........................ugucugagcgucCcuugcc................................................................. 1 1 mv7
........................ugGcugagcgucgcuugcc................................................................. 2 1 mv7
........................ugucugagcgucgcuugcA................................................................. 30 1 mv7
........................ugucugagcgucgcuugUc................................................................. 1 1 mv7
........................ugucugagcgucgcuCgcc................................................................. 2 1 mv7
........................ugucugagcgucgcuuAcc................................................................. 1 1 mv7
........................uguAugagcgucgcuugcc................................................................. 3 1 mv7
........................ugucugagcgucgcuugcU................................................................. 22 1 mv7
........................ugucugagcgucgcuugccC................................................................ 2 1 mv7
........................ugGcugagcgucgcuugccg................................................................ 1 1 mv7
........................Cgucugagcgucgcuugccg................................................................ 2 1 mv7
........................uguAugagcgucgcuugccg................................................................ 2 1 mv7
........................ugucugUgcgucgcuugccg................................................................ 1 1 mv7
........................ugucugGgcgucgcuugccg................................................................ 1 1 mv7
........................ugucugagcgucgcCugccg................................................................ 1 1 mv7
........................ugucugagcgucgcuugccA................................................................ 4 1 mv7
........................ugucugagcgucgcuugccg................................................................ 53 0 mv7
........................ugucugagcgucgcuugAcg................................................................ 1 1 mv7
........................ugucugagcgucgcuugccAa............................................................... 1 1 mv7
........................ugucugagcgucgcuugcAga............................................................... 3 1 mv7
........................uguAugagcgucgcuugccga............................................................... 1 1 mv7
........................ugucugagcgucgcAugccga............................................................... 1 1 mv7
........................ugucugagcgucgcuugccga............................................................... 69 0 mv7
........................ugucugagcgucgcuuUccga............................................................... 1 1 mv7
........................ugGcugagcgucgcuugccga............................................................... 3 1 mv7
........................ugucugagcgucgcuCgccga............................................................... 1 1 mv7
........................ugucugagcgucgcuugccgG............................................................... 2 1 mv7
........................ugucugagcgucgcuugccgaA.............................................................. 26 1 mv7
........................ugucugCgcgucgcuugccgau.............................................................. 1 1 mv7
........................ugucugagcgucgcuugccgaG.............................................................. 2 1 mv7
........................ugucugGgcgucgcuugccgau.............................................................. 1 1 mv7
```

Mature Star

| | | | |
|---|---|---|---|
| cuggguuccucccagggcuaugccugucugagcgucgcuugccgaucaaaauuccccaggguugccucugggcuccuuggggugcccagcuguucuguggcagggccc | | | |
| ........................Cgucugagcgucgcuugccgau.............................................................. | 1 | 1 | mv7 |
| ........................ugucugagcgucgcuugccgau.............................................................. | 61 | 0 | mv7 |
| ........................ugAcugagcgucgcuugccgau.............................................................. | 1 | 1 | mv7 |
| ........................ugucugagcgucgcuugccgaC.............................................................. | 11 | 1 | mv7 |
| ........................ugGcugagcgucgcuugccgau.............................................................. | 1 | 1 | mv7 |
| ........................ugucuUagcgucgcuugccgauc............................................................. | 1 | 1 | mv7 |
| ........................ugucugagcgucgcuugccgaCc............................................................. | 1 | 1 | mv7 |
| ........................ugucugagcgucgcuugccgauG............................................................. | 2 | 1 | mv7 |
| ........................ugucugagcgucgcGugccgauc............................................................. | 1 | 1 | mv7 |
| ........................ugucugagcgucgcuugccgauA............................................................. | 16 | 1 | mv7 |
| ........................Agucugagcgucgcuugccgauc............................................................. | 3 | 1 | mv7 |
| ........................ugucugagcgucgcuugAcgauc............................................................. | 1 | 1 | mv7 |
| ........................uguGugagcgucgcuugccgauc............................................................. | 1 | 1 | mv7 |
| ........................ugucugagcgucgcuugccgauc............................................................. | 120 | 0 | mv7 |
| ........................uAucugagcgucgcuugccgauc............................................................. | 2 | 1 | mv7 |
| ........................ugAcugagcgucgcuugccgauc............................................................. | 2 | 1 | mv7 |
| ........................ugCcugagcgucgcuugccgauc............................................................. | 1 | 1 | mv7 |
| ........................ugGcugagcgucgcuugccgauc............................................................. | 8 | 1 | mv7 |
| ........................ugucugUgcgucgcuugccgauc............................................................. | 1 | 1 | mv7 |
| ........................ugucugagcgucCcuugccgauc............................................................. | 3 | 1 | mv7 |
| ........................ugucugagcgucgcuugGcgauc............................................................. | 1 | 1 | mv7 |
| ........................ugucugGgcgucgcuugccgauc............................................................. | 1 | 1 | mv7 |
| ........................ugucugagcgucgcuugccAauc............................................................. | 1 | 1 | mv7 |
| ........................ugucCgagcgucgcuugccgauc............................................................. | 1 | 1 | mv7 |
| ........................ugucugagcgucgcuugccgauU............................................................. | 8 | 1 | mv7 |
| ........................uguAugagcgucgcuugccgauc............................................................. | 6 | 1 | mv7 |
| ........................ugucugagcgucgcuGgccgauc............................................................. | 2 | 1 | mv7 |
| ........................Cgucugagcgucgcuugccgauca............................................................ | 1 | 1 | mv7 |
| ........................ugucugagcgucgcuuCccgauca............................................................ | 1 | 1 | mv7 |
| ........................ugucugagcgucgcuugccgauAa............................................................ | 1 | 1 | mv7 |
| ........................ugucugagcgucgcuugccgauca............................................................ | 16 | 0 | mv7 |
| ........................ugucugagcgucgcuugccgauGaa........................................................... | 1 | 1 | mv7 |
| ........................ugucugagcgucgcuugccgauAaa........................................................... | 6 | 1 | mv7 |
| ........................ugucugagcgucgcuugccgaucaa........................................................... | 26 | 0 | mv7 |
| ........................ugucugagcgucgcuugccgaucGa........................................................... | 1 | 1 | mv7 |
| ........................Agucugagcgucgcuugccgaucaa........................................................... | 1 | 1 | mv7 |
| ........................ugucugagcgucgcuugccgaucaCa.......................................................... | 1 | 1 | mv7 |
| ........................ugucugagcgucgcCugccgaucaaa.......................................................... | 1 | 1 | mv7 |
| ........................ugucugagcgucgcuugccgaucaaG.......................................................... | 2 | 1 | mv7 |
| ........................ugGcugagcgucgcuugccgaucaaa.......................................................... | 1 | 1 | mv7 |
| ........................ugucugagcgucgcuugccgaucaaC.......................................................... | 2 | 1 | mv7 |
| ........................ugucugagcgucgcuugccgaucaaa.......................................................... | 31 | 0 | mv7 |
| ........................ugucugagcgucgcuugccgauAaaa.......................................................... | 11 | 1 | mv7 |
| ........................ugucugagcgucgcuugccgauGaaa.......................................................... | 1 | 1 | mv7 |
| ........................ugucugagAgucgcuugccgaucaaaa......................................................... | 1 | 1 | mv7 |
| ........................Agucugagcgucgcuugccgaucaaaa......................................................... | 1 | 1 | mv7 |
| ........................ugucugagcgucgcuugccgaucaaaa......................................................... | 23 | 0 | mv7 |
| ........................ugGcugagcgucgcuugccgaucaaaa......................................................... | 1 | 1 | mv7 |
| ........................ugucugagcgucgcuugccgaucGaaa......................................................... | 1 | 1 | mv7 |
| ........................ugucugagcgucgcuugccgaucaaaC......................................................... | 1 | 1 | mv7 |
| ........................ugucugagcgucgcuugccgaucaaaG......................................................... | 1 | 1 | mv7 |
| ........................ugucugagcgucgcuugccgauAaaaa......................................................... | 6 | 1 | mv7 |
| ........................ugucugagcgucgcuugccgaucaaaaA........................................................ | 10 | 1 | mv7 |
| .........................gucugagcgucgcuugcU................................................................. | 12 | 1 | mv7 |
| .........................gucugagcgCcgcuugcc................................................................. | 1 | 1 | mv7 |
| .........................gucugagcgucgcuAgcc................................................................. | 1 | 1 | mv7 |
| .........................gucugagcgucgcuugcA................................................................. | 41 | 1 | mv7 |
| .........................gucugagcgucgcuCgcc................................................................. | 2 | 1 | mv7 |
| .........................gucugagcgucgcuugcc................................................................. | 37 | 0 | mv7 |
| .........................gucugagcgucgcuugcG................................................................. | 1 | 1 | mv7 |
| .........................gucCgagcgucgcuugcc................................................................. | 1 | 1 | mv7 |
| .........................gucugagcgucgcAugcc................................................................. | 1 | 1 | mv7 |
| .........................gucugagcgucgcuGgcc................................................................. | 1 | 1 | mv7 |
| .........................gucugagcgucgcCugcc................................................................. | 1 | 1 | mv7 |
| .........................gucugagcgucgAuugccg................................................................ | 1 | 1 | mv7 |
| .........................gucugaAcgucgcuugccg................................................................ | 1 | 1 | mv7 |
| .........................gucugagcgucUcuugccg................................................................ | 1 | 1 | mv7 |
| .........................gucugagcgucgcuugccA................................................................ | 11 | 1 | mv7 |
| .........................gucugagcgucgcuugcUg................................................................ | 1 | 1 | mv7 |
| .........................gucugagcgucgcuAgccg................................................................ | 1 | 1 | mv7 |

```
                 Mature                                                           Star
cuggguuccucccagggcuaugccugucugagcgucgcuugccgaucaaaauuccccaggguugccucugggcuccuuggggugcccagcuguucuguggcagggccc
.........................gucugagcgucgcuugccg................................................................  36    0    mv7
.........................Cucugagcgucgcuugccg................................................................  1     1    mv7
.........................gucugagcgucgcuugcAg................................................................  1     1    mv7
.........................gucugGgcgucgcuugccg................................................................  1     1    mv7
.........................gucugagcgucgcuugccU................................................................  1     1    mv7
.........................gucugUgcgucgcuugccga...............................................................  1     1    mv7
.........................gucugagcgucgAuugccga...............................................................  1     1    mv7
.........................gucugagcgucgcuugGcga...............................................................  1     1    mv7
.........................gucugagcgucgcuugccgU...............................................................  1     1    mv7
.........................gucugagcgucgcuugccga...............................................................  57    0    mv7
.........................Cucugagcgucgcuugccga...............................................................  1     1    mv7
.........................gucugagcgucgcuCgccga...............................................................  1     1    mv7
.........................gCcugagcgucgcuugccga...............................................................  1     1    mv7
.........................gucugagcgucgcuugccgaG..............................................................  1     1    mv7
.........................gucugagcguGgcuugccgau..............................................................  1     1    mv7
.........................gucugagcgucgcuugccgaA..............................................................  24    1    mv7
.........................gucugagcgucgcuugccgau..............................................................  25    0    mv7
.........................gucugagcgucgcuugccgaC..............................................................  2     1    mv7
.........................gucugagcgucgcuugccgGuc.............................................................  1     1    mv7
.........................gucugagcgucgcGugccgauc.............................................................  1     1    mv7
.........................gucugagcgCcgcuugccgauc.............................................................  1     1    mv7
.........................gucugagcgucgcuugccgauA.............................................................  17    1    mv7
.........................Aucugagcgucgcuugccgauc.............................................................  1     1    mv7
.........................gucAgagcgucgcuugccgauc.............................................................  2     1    mv7
.........................gucugagcgucgcuugccgUuc.............................................................  1     1    mv7
.........................gucCgagcgucgcuugccgauc.............................................................  1     1    mv7
.........................gucugagcgucgAuugccgauc.............................................................  1     1    mv7
.........................guGugagcgucgcuugccgauc.............................................................  3     1    mv7
.........................gucugagcgucgcuuCccgauc.............................................................  1     1    mv7
.........................gucugagcgucgcuugccgauG.............................................................  2     1    mv7
.........................Cucugagcgucgcuugccgauc.............................................................  2     1    mv7
.........................gucugagcgucgcuugccgauc.............................................................  100   0    mv7
.........................Uucugagcgucgcuugccgauc.............................................................  1     1    mv7
.........................gucugagcgucgcuugccgaCc.............................................................  1     1    mv7
.........................gucugagcgucgcuCgccgauc.............................................................  1     1    mv7
.........................gucugagcgucgcuugcUgauc.............................................................  1     1    mv7
.........................gucugagcgucgcuugccgauU.............................................................  4     1    mv7
.........................gucugUgcgucgcuugccgauc.............................................................  1     1    mv7
.........................guAugagcgucgcuugccgauc.............................................................  1     1    mv7
.........................gucugagcgucgcuugUcgauca............................................................  1     1    mv7
.........................gucugagcgucgcuugccgauca............................................................  13    0    mv7
.........................gucugagcgucgcuugccgauAa............................................................  2     1    mv7
.........................gucugagcgucgcuugccgaucaa...........................................................  11    0    mv7
.........................gucAgagcgucgcuugccgaucaa...........................................................  1     1    mv7
.........................gucugagcgucgcuugccgauAaa...........................................................  3     1    mv7
.........................gucugagcgucgcuugccgauGaaa..........................................................  1     1    mv7
.........................gucugagcgucgcuugccgauAaaa..........................................................  10    1    mv7
.........................gucugagcgucgcuugccgaucaaa..........................................................  17    0    mv7
.........................gucugagcgucgcuugccgaucaaaa.........................................................  10    0    mv7
.........................gucugagcgucgcuAgccgaucaaaa.........................................................  1     1    mv7
.........................gucugagcgucgcuugccgaucaaaU.........................................................  1     1    mv7
.........................gucugagcgucgcuugccgaucaGaa.........................................................  3     1    mv7
.........................gucugagcgucgcuugccgaucaaaaA........................................................  13    1    mv7
.........................gucugagcgucgcuugccgaucaaaaAu.......................................................  1     1    mv7
..........................ucugagcgucgcuugccg................................................................  10    0    mv7
..........................ucugagcgucgcuugccA................................................................  1     1    mv7
..........................ucugagcgucgcuugccga...............................................................  6     0    mv7
..........................ucugagcgucgcuugccgau..............................................................  6     0    mv7
..........................ucugagcgucgcuugccgaA..............................................................  8     1    mv7
..........................ucugagcgucgcuuAccgau..............................................................  1     1    mv7
..........................ucGgagcgucgcuugccgauc.............................................................  1     1    mv7
..........................ucugagcgucgcuugccgauA.............................................................  3     1    mv7
..........................ucugagcgucgcuugccgauc.............................................................  29    0    mv7
..........................ucugagcgucgcuuAccgauc.............................................................  1     1    mv7
..........................ucugagcgucgcuugccgauca............................................................  1     0    mv7
..........................ucugagcgucgcuugccgauAaa...........................................................  1     1    mv7
..........................ucugagcgucgcuugccgauAaaa..........................................................  1     1    mv7
..........................ucugagcgucgcuugccgauAaaaa.........................................................  1     1    mv7
..........................ucugagcgucgcuugccgaucaaaa.........................................................  1     0    mv7
...........................cugCgcgucgcuugccga...............................................................  1     1    mv7
```

```
                    Mature                                                           Star
cuggguuccucccagggcuaugccugucugagcgucgcuugccgaucaaaauuccccaggguugccucugggcuccuuggggugcccagcuguucuguggcagggccc
...........................cugagcgucgcuugccga...............................................................    5    0    mv7
...........................cuUagcgucgcuugccga...............................................................    1    1    mv7
...........................Uugagcgucgcuugccgau..............................................................    1    1    mv7
...........................cugagcgucgcuugccgaA..............................................................    3    1    mv7
...........................cugagcgucgcuugccgau..............................................................    8    0    mv7
...........................cugagcgucgcuugccUauc.............................................................    1    1    mv7
...........................cugagcguAgcuugccgauc.............................................................    1    1    mv7
...........................cugagcgucgcuuAccgauc.............................................................    1    1    mv7
...........................cugCgcgucgcuugccgauc.............................................................    3    1    mv7
...........................cugagcgucgcuugccgauA.............................................................    7    1    mv7
...........................cugagcgucgcuugccgauc.............................................................    10   0    mv7
...........................Augagcgucgcuugccgauc.............................................................    1    1    mv7
...........................cugagcAucgcuugccgauc.............................................................    1    1    mv7
...........................Augagcgucgcuugccgauca............................................................    1    1    mv7
...........................cugagcgucgcuugccgaucaaa..........................................................    1    0    mv7
...........................cugagcgucgcuugUcgaucaaaa.........................................................    1    1    mv7
............................ugagcgucgcuugccgaA..............................................................    6    1    mv7
............................ugaCcgucgcuugccgauc.............................................................    1    1    mv7
............................ugagcgucgcuugccgauc.............................................................    4    0    mv7
............................ugagcgucgcuugccgauA.............................................................    1    1    mv7
............................ugagcgucgcuugccgauca............................................................    1    0    mv7
............................ugagcgucgcuugccgaucaa...........................................................    2    0    mv7
............................ugagcgucgcuugccgauAaa...........................................................    1    1    mv7
............................ugagcgucgcuugccgauAaaa..........................................................    2    1    mv7
............................ugagcgucgcuugccgauAaaaa.........................................................    1    1    mv7
............................ugagcgucgcuugccgaucaaaa.........................................................    1    0    mv7
.............................gagcgucgcuugccgauc.............................................................    1    0    mv7
.................................Cucgcuugccgaucaaaa.........................................................    1    1    mv7
.......................................................ccaggguugccucugggcuG.................................    1    1    mv7
........................................................................................gcuguucuguggcaggAc..    1    1    mv7
```
