## Supplementary material for "SEA: The small RNA Expression Atlas": p-hsa-miR-235-1

| | | |
|---|---|---|
| Provisional ID | : | chrUn_GL000220v1_21620 |
| Score total | : | 144078.7 |
| Score for star read(s) | : | 3.9 |
| Score for read counts | : | 144071.6 |
| Score for mfe | : | 2.2 |
| Score for randfold | : | 1.6 |
| Score for cons. seed | : | -0.6 |
| Total read count | : | 282602 |
| Mature read count | : | 36506 |
| Loop read count | : | 245911 |
| Star read count | : | 185 |

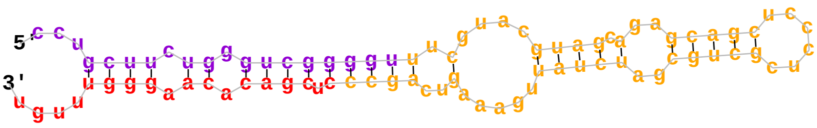

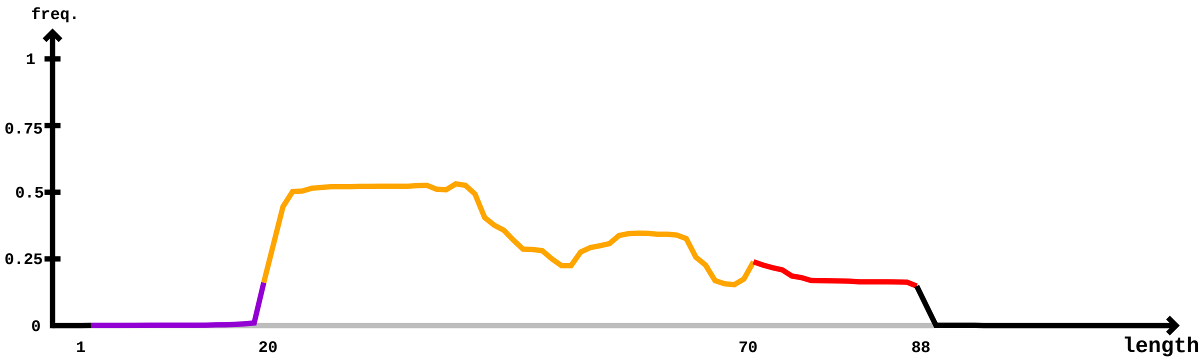


**Star** **Mature**

```
5'- gaccugcuucugggucgggguuucguacguagcagagcagcucccucgcugcgaucuauugaaagucagcccucgacacaaggguuuguccgcgcgcgcgcgcgcgcgcgcgugc  -3'  obs
    gaccugcuucugggucgggguuucguacguagcagagcagcucccucgcugcgaucuauugaaagucagcccucgacacaaggguuuguccgcgcgcgcgcgcgcgcgcgcgugc       exp
    (((..((((.((.(((((((((.(....((((.(..(((((......)))))..))))).....)..))))).)))).)).))))..))).((((((((((....))))))))))  reads  mm  sample
    ..ccugcuucugggucgggg...............................................................................................  7      0   7y1
    ..ccugcuucugggucggggG..............................................................................................  6      1   7y1
    ..ccugcuucugggucggggA..............................................................................................  1      1   7y1
    ..cAugcuucugggucggggu..............................................................................................  1      1   7y1
    ..ccugcuucugggucggggu..............................................................................................  164    0   7y1
    ...cugcuucuggguAgggguuucgu.........................................................................................  2      1   7y1
    ...cugcuucugUgucgggguuucgu.........................................................................................  1      1   7y1
    ...cugcuucugggucgggguAucgu.........................................................................................  1      1   7y1
    ...cugcuucugggucgggUuuucgu.........................................................................................  2      1   7y1
    ....ugcuucAgggucgggguu.............................................................................................  1      1   7y1
    .....gcuucugggAcgggguuucgu.........................................................................................  1      1   7y1
    .....gcuucuggguAgggguuucgu.........................................................................................  1      1   7y1
    .....gcuucuggCucgggguuucgu.........................................................................................  1      1   7y1
    .....gcuucuggUucgggguuucgu.........................................................................................  1      1   7y1
    ......cuucugggucgggguGuc...........................................................................................  1      1   7y1
    ......cuucuAggucgggguuuc...........................................................................................  1      1   7y1
    ......cuucugggucgggguuuG...........................................................................................  1      1   7y1
    ......cuucugggAcgggguuuc...........................................................................................  1      1   7y1
    ......cuucugggucggCguuuc...........................................................................................  1      1   7y1
    ......cuucuUggucgggguuucgu.........................................................................................  1      1   7y1
    ......cuucuggguGgggguuucgu.........................................................................................  1      1   7y1
    ......cuucugggucgggguCucgu.........................................................................................  1      1   7y1
    ......cuucugggucUggguuucgu.........................................................................................  1      1   7y1
    ......cuucugggAcgggguuucgu.........................................................................................  2      1   7y1
    ......cuucuCggucgggguuucgu.........................................................................................  1      1   7y1
    ......cuucuggguAgggguuucgu.........................................................................................  4      1   7y1
    ......cuucugggucCggguuucgu.........................................................................................  1      1   7y1
    ......cuucugggucgggguAucgu.........................................................................................  1      1   7y1
    ......cuucugggucggggGuucgu.........................................................................................  2      1   7y1
    ......cuucugggucgggAuuucgu.........................................................................................  1      1   7y1
    ......cuucugggucggUguuucgu.........................................................................................  1      1   7y1
    ......cuucugggucggggAuucgu.........................................................................................  1      1   7y1
    ......cuucugggucgggUuuucgu.........................................................................................  1      1   7y1
```

**Star** **Mature**

```
gaccugcuucugggucgggguuucguacguagcagagcagcucccucgcugcgaucuauugaaagucagcccucgacacaaggguuuguccgcgcgcgcgcgcgcgcgugc
......cuucugggucggCguuucgu.....................................................................................	1	1	7y1
......cuucugggucgggguuucUu.....................................................................................	3	1	7y1
.......uucugggucgggguCucgu.....................................................................................	2	1	7y1
.......uucuUggucgggguuucgu.....................................................................................	2	1	7y1
.......uucugggAcgggguuucgu.....................................................................................	3	1	7y1
.......uucugggucggUguuucgu.....................................................................................	1	1	7y1
.......uucugggucgggCuuucgu.....................................................................................	4	1	7y1
.......uucugggucgUgguuucgu.....................................................................................	2	1	7y1
.......uucugggucggggCuucgu.....................................................................................	1	1	7y1
.......uucuggguAgggguuucgu.....................................................................................	6	1	7y1
.......uucugggucgggguAucgu.....................................................................................	3	1	7y1
.......uucugCgucgggguuucgu.....................................................................................	4	1	7y1
.......uucugggucggCguuucgu.....................................................................................	1	1	7y1
.......uucugggucUggguuucgu.....................................................................................	2	1	7y1
.......uucugggucggggGuucgu.....................................................................................	1	1	7y1
.......uucuggUucgggguuucgu.....................................................................................	2	1	7y1
.......uucugUgucgggguuucgu.....................................................................................	1	1	7y1
........ucuggCucgggguuucgu.....................................................................................	4	1	7y1
........ucugggucgggCuuucgu.....................................................................................	1	1	7y1
........ucugggucgggguAucgu.....................................................................................	4	1	7y1
........ucugggucgCgguuucgu.....................................................................................	8	1	7y1
........ucuAggucgggguuucgu.....................................................................................	2	1	7y1
........ucugCgucgggguuucgu.....................................................................................	1	1	7y1
........ucuggUucgggguuucgu.....................................................................................	7	1	7y1
........ucugggucCggguuucgu.....................................................................................	2	1	7y1
........ucuggAucgggguuucgu.....................................................................................	1	1	7y1
........ucugggucggUguuucgu.....................................................................................	3	1	7y1
........ucugggucUggguuucgu.....................................................................................	2	1	7y1
........ucugggucgggguCucgu.....................................................................................	1	1	7y1
........ucugggucgUgguuucgu.....................................................................................	1	1	7y1
........ucuggguAgggguuucgu.....................................................................................	4	1	7y1
........ucugggucggggAuucgu.....................................................................................	1	1	7y1
........ucugUgucgggguuucgu.....................................................................................	2	1	7y1
........ucugggAcgggguuucgu.....................................................................................	4	1	7y1
........ucugggucgggUuuucgu.....................................................................................	1	1	7y1
........ucuggguGgggguuucgu.....................................................................................	1	1	7y1
........ucugggucgggguuAcgu.....................................................................................	1	1	7y1
........ucugggucggCguuucgu.....................................................................................	6	1	7y1
........ucugggucggCguuucgua....................................................................................	2	1	7y1
........ucugUgucgggguuucgua....................................................................................	2	1	7y1
........ucugggucgCgguuucgua....................................................................................	6	1	7y1
........ucugggucgggCuuucgua....................................................................................	1	1	7y1
........ucuggAucgggguuucgua....................................................................................	1	1	7y1
........ucugggucggAguuucgua....................................................................................	1	1	7y1
........ucugggucgggUuuucgua....................................................................................	2	1	7y1
........ucugggucggggAuucgua....................................................................................	1	1	7y1
........ucugggucgUgguuucgua....................................................................................	3	1	7y1
........ucuggguGgggguuucgua....................................................................................	1	1	7y1
........ucugggucgggguuGcgua....................................................................................	1	1	7y1
........ucugggucgggguuAcgua....................................................................................	1	1	7y1
........ucugggucUggguuucgua....................................................................................	3	1	7y1
........ucugggucCggguuucgua....................................................................................	2	1	7y1
........ucugCgucgggguuucgua....................................................................................	2	1	7y1
........ucuggUucgggguuucgua....................................................................................	5	1	7y1
........ucugggAcgggguuucgua....................................................................................	12	1	7y1
........ucugggucgggguAucgua....................................................................................	2	1	7y1
........ucugggucggUguuucgua....................................................................................	4	1	7y1
........ucuggguAgggguuucgua....................................................................................	4	1	7y1
.........cugCgucgggguuucgua....................................................................................	1	1	7y1
.........cuggguAgggguuucgua....................................................................................	5	1	7y1
.........cugggucggUguuucgua....................................................................................	1	1	7y1
.........cugggucgggguuuAgua....................................................................................	5	1	7y1
.........cugggucgggUuuucgua....................................................................................	1	1	7y1
.........cugggucgAgguuucgua....................................................................................	1	1	7y1
.........cugggucgggguuuGgua....................................................................................	1	1	7y1
.........cugggucUggguuucgua....................................................................................	1	1	7y1
.........cugggucgCgguuucgua....................................................................................	1	1	7y1
.........cugggucggggGuucgua....................................................................................	1	1	7y1
.........cugggAcgggguuucguac...................................................................................	1	1	7y1
.........cuggguAgggguuucguac...................................................................................	1	1	7y1
```

## Star                                   Mature

```
gaccugcuucugggucgggguuucguacguagcagagcagcucccucgcugcgaucuauugaaagucagcccucgacacaaggguuuguccgcgcgcgcgcgcgcgcgcgcgugc
.........cugggucgggguuAcguac.......................................................................................    1    1    7y1
.........cugggucgggUuuucguac.......................................................................................    1    1    7y1
.........cuggguCggguuucguac.......................................................................................    1    1    7y1
.........cugggucgggguuuGguac.......................................................................................    1    1    7y1
.........cugggucggCguuucguac.......................................................................................    1    1    7y1
.........cugggucgggguuuAguac.......................................................................................    1    1    7y1
..............ucgggguAucguacguag...................................................................................    1    1    7y1
..............ucggggAuucguacguag...................................................................................    1    1    7y1
..............ucggggAuucguacguagc..................................................................................    1    1    7y1
..............ucggggCuucguacguagc..................................................................................    1    1    7y1
..............ucgggguAucguacguagc..................................................................................    1    1    7y1
..............ucgggUuuucguacguagc..................................................................................    1    1    7y1
..............ucggggGuucguacguagc..................................................................................    1    1    7y1
..............ucgggguuucguaUguagc..................................................................................    1    1    7y1
..............ucgggguuucAuacguagcagagc.............................................................................    2    1    7y1
..............ucgggUuuucguacguagcagagcagcuc........................................................................    4    1    7y1
..............ucgggguCucguacguagcagagcagcuc........................................................................    1    1    7y1
..............ucgggCuuucguacguagcagagcagcuc........................................................................    6    1    7y1
..............ucggggAuucguacguagcagagcagcuc........................................................................    23   1    7y1
..............ucggggCuucguacguagcagagcagcuc........................................................................    2    1    7y1
..............ucgggguGucguacguagcagagcagcuc........................................................................    1    1    7y1
...............cgggguuucgCacguagc..................................................................................    1    1    7y1
...............cggAguuucguacguagc..................................................................................    1    1    7y1
...............cgggguuAcguacguagc..................................................................................    1    1    7y1
...............cgggguuucUuacguagc..................................................................................    1    1    7y1
...............cggggAuucguacguagc..................................................................................    2    1    7y1
...............cggCguuucguacguagc..................................................................................    1    1    7y1
...............cggggu Aucguacguagc..................................................................................    3    1    7y1
...............cgggguuucguaAguagc..................................................................................    1    1    7y1
...............cgggguuucguacAuagca.................................................................................    1    1    7y1
...............cgggguuucUuacguagca.................................................................................    1    1    7y1
...............cggggAuucguacguagca.................................................................................    2    1    7y1
...............cgggguuucguaAguagca.................................................................................    1    1    7y1
...............cgggguuuAguacguagcag................................................................................    1    1    7y1
...............cgggguuucCuacguagcag................................................................................    1    1    7y1
...............cggggAuucguacguagcag................................................................................    1    1    7y1
...............cgggUuuucguacguagcag................................................................................    1    1    7y1
...............cgggguuucguacgAagcag................................................................................    1    1    7y1
...............cgggguuucguacCuagcag................................................................................    1    1    7y1
...............Agggguuucguacguagcag................................................................................    1    1    7y1
...............cgggguuucUuacguagcaga...............................................................................    1    1    7y1
...............cgggguuuUguacguagcaga...............................................................................    1    1    7y1
...............cgggCuuucguacguagcaga...............................................................................    3    1    7y1
...............cgggguGucguacguagcaga...............................................................................    1    1    7y1
...............cgggguuucguaAguagcaga...............................................................................    3    1    7y1
...............cgggguuuGguacguagcaga...............................................................................    1    1    7y1
...............cgggguuucguacgCagcaga...............................................................................    1    1    7y1
...............cgggguAucguacguagcaga...............................................................................    4    1    7y1
...............cgggguuAcguacguagcaga...............................................................................    2    1    7y1
...............cgggguuucCuacguagcaga...............................................................................    1    1    7y1
...............cgggguuucguaGguagcaga...............................................................................    1    1    7y1
...............cggggGuucguacguagcaga...............................................................................    1    1    7y1
...............cgggguuucguacUuagcaga...............................................................................    4    1    7y1
...............cgggguuuAguacguagcaga...............................................................................    1    1    7y1
...............cggggAuucguacguagcaga...............................................................................    5    1    7y1
...............cgggguuucgAacguagcagag..............................................................................    1    1    7y1
...............cgggguuucguaAguagcagag..............................................................................    3    1    7y1
...............cgggguuuAguacguagcagag..............................................................................    1    1    7y1
...............cgggguuucguacUuagcagag..............................................................................    1    1    7y1
...............cgggguuucAuacguagcagag..............................................................................    1    1    7y1
...............cgggUuuucguacguagcagag..............................................................................    1    1    7y1
...............cgggguuucUuacguagcagag..............................................................................    1    1    7y1
...............cgggguAucguacguagcagag..............................................................................    2    1    7y1
...............cgggUuuucguacguagcagagc.............................................................................    2    1    7y1
...............cgggguuucguacUuagcagagc.............................................................................    2    1    7y1
...............cgggguuucguGcguagcagagc.............................................................................    1    1    7y1
...............cgggguuucAuacguagcagagc.............................................................................    1    1    7y1
...............cgggCuuucguacguagcagagc.............................................................................    3    1    7y1
...............cgggguuuAguacguagcagagc.............................................................................    7    1    7y1
...............cgggguuucgGacguagcagagc.............................................................................    1    1    7y1
```

**Star** **Mature**

| | | | |
|---|---|---|---|
| gaccugcuucugggucgggguuucguacguagcagagcagcucccucgcugcgaucuauugaaagucagcccucgacacaaggguuuguccgcgcgcgcgcgcgcgcgcgugc | | | |
| ...............cgggguuucguacCuagcagagc........................................................................... | 1 | 1 | 7y1 |
| ...............cgggguuucCuacguagcagagc........................................................................... | 2 | 1 | 7y1 |
| ...............cgggguuucguacAuagcagagc........................................................................... | 1 | 1 | 7y1 |
| ...............cggggGuucguacguagcagagc........................................................................... | 3 | 1 | 7y1 |
| ...............cgggguuucguacgAagcagagc........................................................................... | 3 | 1 | 7y1 |
| ...............cgggguuuGguacguagcagagc........................................................................... | 1 | 1 | 7y1 |
| ...............cgggguuucguaAguagcagagc........................................................................... | 11 | 1 | 7y1 |
| ...............cgggguuucgCacguagcagagc........................................................................... | 3 | 1 | 7y1 |
| ...............cggggAuucguacguagcagagc........................................................................... | 27 | 1 | 7y1 |
| ...............cgggguuucUuacguagcagagc........................................................................... | 1 | 1 | 7y1 |
| ...............cgggguuucgAacguagcagagc........................................................................... | 3 | 1 | 7y1 |
| ...............cgggguuCcguacguagcagagc........................................................................... | 2 | 1 | 7y1 |
| ...............cgggguCucguacguagcagagc........................................................................... | 1 | 1 | 7y1 |
| ...............cgggguAucguacguagcagagc........................................................................... | 14 | 1 | 7y1 |
| ...............cgggguuAcguacguagcagagc........................................................................... | 3 | 1 | 7y1 |
| ...............cgggguuucguaUguagcagagc........................................................................... | 1 | 1 | 7y1 |
| ...............cgggguuucguacgAagcagagca.......................................................................... | 3 | 1 | 7y1 |
| ...............cggggGuucguacguagcagagca.......................................................................... | 1 | 1 | 7y1 |
| ...............cgggguuucCuacguagcagagca.......................................................................... | 2 | 1 | 7y1 |
| ...............cgggguuucguacUuagcagagca.......................................................................... | 2 | 1 | 7y1 |
| ...............cgggguuucgGacguagcagagca.......................................................................... | 1 | 1 | 7y1 |
| ...............cgggguuucguacgGagcagagca.......................................................................... | 1 | 1 | 7y1 |
| ...............cgggguuuUguacguagcagagca.......................................................................... | 1 | 1 | 7y1 |
| ...............cgggguuucUuacguagcagagca.......................................................................... | 4 | 1 | 7y1 |
| ...............cgggguuucguGcguagcagagca.......................................................................... | 2 | 1 | 7y1 |
| ...............cgggguuucgAacguagcagagca.......................................................................... | 3 | 1 | 7y1 |
| ...............cgggguuAcguacguagcagagca.......................................................................... | 6 | 1 | 7y1 |
| ...............cgggguuucguaAguagcagagca.......................................................................... | 4 | 1 | 7y1 |
| ...............cgggCuuucguacguagcagagca.......................................................................... | 3 | 1 | 7y1 |
| ...............cgggUuuucguacguagcagagca.......................................................................... | 1 | 1 | 7y1 |
| ...............cgggguAucguacguagcagagca.......................................................................... | 10 | 1 | 7y1 |
| ...............cgggAuuucguacguagcagagca.......................................................................... | 1 | 1 | 7y1 |
| ...............cgggguuucgCacguagcagagca.......................................................................... | 1 | 1 | 7y1 |
| ...............cgggguuuAguacguagcagagca.......................................................................... | 5 | 1 | 7y1 |
| ...............cggggAuucguacguagcagagca.......................................................................... | 10 | 1 | 7y1 |
| ...............cgggguCucguacguagcagagca.......................................................................... | 1 | 1 | 7y1 |
| ...............cgggguuucAuacguagcagagca.......................................................................... | 1 | 1 | 7y1 |
| ...............cgggguAucguacguagcagagcag......................................................................... | 1 | 1 | 7y1 |
| ...............cgggCuuucguacguagcagagcag......................................................................... | 1 | 1 | 7y1 |
| ...............cgggguuAcguacguagcagagcag......................................................................... | 1 | 1 | 7y1 |
| ...............cgggUuuucguacguagcagagcag......................................................................... | 2 | 1 | 7y1 |
| ...............cgggguuucguacUuagcagagcag......................................................................... | 1 | 1 | 7y1 |
| ...............cgggguuucgAacguagcagagcag......................................................................... | 1 | 1 | 7y1 |
| ...............cgggguuuGguacguagcagagcag......................................................................... | 1 | 1 | 7y1 |
| ...............cgggguuucCuacguagcagagcag......................................................................... | 1 | 1 | 7y1 |
| ...............cgggguuucguaAguagcagagcag......................................................................... | 1 | 1 | 7y1 |
| ...............cgggguuuAguacguagcagagcag......................................................................... | 4 | 1 | 7y1 |
| ...............cgggguAucguacguagcagagcagc........................................................................ | 9 | 1 | 7y1 |
| ...............cgggguuucguacgAagcagagcagc........................................................................ | 2 | 1 | 7y1 |
| ...............cgggguuucgAacguagcagagcagc........................................................................ | 4 | 1 | 7y1 |
| ...............cggggGuucguacguagcagagcagc........................................................................ | 1 | 1 | 7y1 |
| ...............cgggguGucguacguagcagagcagc........................................................................ | 1 | 1 | 7y1 |
| ...............cggggAuucguacguagcagagcagc........................................................................ | 10 | 1 | 7y1 |
| ...............cgggguuucgGacguagcagagcagc........................................................................ | 1 | 1 | 7y1 |
| ...............cgggguuucguacUuagcagagcagc........................................................................ | 2 | 1 | 7y1 |
| ...............cggggCuucguacguagcagagcagc........................................................................ | 1 | 1 | 7y1 |
| ...............cgggguuAcguacguagcagagcagc........................................................................ | 2 | 1 | 7y1 |
| ...............cgggguuucguaGguagcagagcagc........................................................................ | 1 | 1 | 7y1 |
| ...............cgggguuuAguacguagcagagcagc........................................................................ | 2 | 1 | 7y1 |
| ...............cgggguuuUguacguagcagagcagc........................................................................ | 1 | 1 | 7y1 |
| ...............cgggguuucguaAguagcagagcagc........................................................................ | 5 | 1 | 7y1 |
| ...............cgggguuCcguacguagcagagcagc........................................................................ | 1 | 1 | 7y1 |
| ...............cgggguuucAuacguagcagagcagc........................................................................ | 2 | 1 | 7y1 |
| ...............cgggguuucguacCuagcagagcagc........................................................................ | 1 | 1 | 7y1 |
| ...............cgggguAucguacguagcagagcagcu....................................................................... | 12 | 1 | 7y1 |
| ...............cggggGuucguacguagcagagcagcu....................................................................... | 3 | 1 | 7y1 |
| ...............cgggguuucguacUuagcagagcagcu....................................................................... | 2 | 1 | 7y1 |
| ...............cgggguuuAguacguagcagagcagcu....................................................................... | 1 | 1 | 7y1 |
| ...............cggggCuucguacguagcagagcagcu....................................................................... | 1 | 1 | 7y1 |
| ...............cggggAuucguacguagcagagcagcu....................................................................... | 10 | 1 | 7y1 |

```
Star                                                                   Mature
gaccugcuucugggucggggguuucguacguagcagagcagcucccucgcugcgaucuauugaaagucagccccucgacacaaggguuuguccgcgcgcgcgcgcgcgcgcgugc
...............cgggguuucguacgAagcagagcagcu..........................................................................   1    1    7y1
...............cgggguuucgAacguagcagagcagcu..........................................................................   4    1    7y1
...............cgggguCucguacguagcagagcagcu..........................................................................   1    1    7y1
...............cgggguuucguaAguagcagagcagcu..........................................................................   2    1    7y1
...............cgggguuucCuacguagcagagcagcu..........................................................................   1    1    7y1
...............cgggguuucguGcguagcagagcagcu..........................................................................   1    1    7y1
...............cgggguuucAuacguagcagagcagcu..........................................................................   2    1    7y1
...............cgggguuAcguacguagcagagcagcu..........................................................................   4    1    7y1
...............cgggUuuucguacguagcagagcagcu..........................................................................   2    1    7y1
...............cgggguuucgGacguagcagagcagcu..........................................................................   1    1    7y1
...............cgggguuuUguacguagcagagcagcu..........................................................................   1    1    7y1
...............cgggguuucUuacguagcagagcagcu..........................................................................   1    1    7y1
...............cgggguGucguacguagcagagcagcuc.........................................................................   2    1    7y1
...............cgggguuucCuacguagcagagcagcuc.........................................................................   1    1    7y1
...............cgggguuucAuacguagcagagcagcuc.........................................................................   2    1    7y1
...............cgggUuuucguacguagcagagcagcuc.........................................................................   2    1    7y1
...............cggggAuucguacguagcagagcagcuc.........................................................................   12   1    7y1
...............cgggguuAcguacguagcagagcagcuc.........................................................................   7    1    7y1
...............cggggGuucguacguagcagagcagcuc.........................................................................   4    1    7y1
...............cgggguuucgAacguagcagagcagcuc.........................................................................   2    1    7y1
...............cgggguuucUuacguagcagagcagcuc.........................................................................   1    1    7y1
...............cgggguuucguacCuagcagagcagcuc.........................................................................   1    1    7y1
...............cgggguuucgCacguagcagagcagcuc.........................................................................   4    1    7y1
...............cgggguAucguacguagcagagcagcuc.........................................................................   16   1    7y1
...............cgggCuuucguacguagcagagcagcuc.........................................................................   2    1    7y1
...............cgggguuucguaAguagcagagcagcuc.........................................................................   1    1    7y1
...............cgggguuucguacgAagcagagcagcuc.........................................................................   1    1    7y1
...............cgggguuucguaUguagcagagcagcuc.........................................................................   1    1    7y1
...............cgggguuuGguacguagcagagcagcuc.........................................................................   1    1    7y1
...............cgggguuucguacUuagcagagcagcuc.........................................................................   3    1    7y1
...............cgggguuuAguacguagcagagcagcuc.........................................................................   5    1    7y1
...............cggggAuucguacguagcagagcagcucccuc.....................................................................   1    1    7y1
................gggguuucguaAguagcaga................................................................................   1    1    7y1
................gggguuAcguacguagcaga................................................................................   2    1    7y1
................gggguuucgAacguagcagag...............................................................................   1    1    7y1
................gggguuuAguacguagcagag...............................................................................   1    1    7y1
................gggguuuAguacguagcagagc..............................................................................   2    1    7y1
................gggguuucUuacguagcagagc..............................................................................   1    1    7y1
................gggguAucguacguagcagagc..............................................................................   5    1    7y1
................gggguuAcguacguagcagagc..............................................................................   3    1    7y1
................gggUuuucguacguagcagagc..............................................................................   3    1    7y1
................gggguuucgAacguagcagagca.............................................................................   1    1    7y1
................gggguuAcguacguagcagagca.............................................................................   2    1    7y1
................gggguuucguaAguagcagagca.............................................................................   1    1    7y1
................gggguuuAguacguagcagagca.............................................................................   1    1    7y1
................gggguAucguacguagcagagca.............................................................................   2    1    7y1
................gggUuuucguacguagcagagca.............................................................................   1    1    7y1
................gggguuucguacguGgcagagcagc...........................................................................   1    1    7y1
................gggguGucguacguagcagagcagc...........................................................................   1    1    7y1
................gggguuAcguacguagcagagcagc...........................................................................   3    1    7y1
................gggguuCcguacguagcagagcagc...........................................................................   1    1    7y1
................gggguuucgAacguagcagagcagc...........................................................................   1    1    7y1
................gggguuucguacgCagcagagcagc...........................................................................   1    1    7y1
................gggguuuAguacguagcagagcagc...........................................................................   3    1    7y1
................gggguuucguUcguagcagagcagc...........................................................................   2    1    7y1
................ggggGuucguacguagcagagcagc...........................................................................   1    1    7y1
................gggguAucguacguagcagagcagc...........................................................................   3    1    7y1
................gggguuucguacgAagcagagcagc...........................................................................   1    1    7y1
................ggggAuucguacguagcagagcagcu..........................................................................   2    1    7y1
................gggguuucguacgCagcagagcagcu..........................................................................   2    1    7y1
................ggggGuucguacguagcagagcagcu..........................................................................   1    1    7y1
................gggguuucCuacguagcagagcagcu..........................................................................   1    1    7y1
................gggguuucguGcguagcagagcagcu..........................................................................   1    1    7y1
................gggguuucguacUuagcagagcagcu..........................................................................   1    1    7y1
................gggCuuucguacguagcagagcagcu..........................................................................   1    1    7y1
................gggguuucguaAguagcagagcagcu..........................................................................   1    1    7y1
................gggguAucguacguagcagagcagcu..........................................................................   6    1    7y1
................gggguuAcguacguagcagagcagcu..........................................................................   1    1    7y1
................gggguuuAguacguagcagagcagcu..........................................................................   1    1    7y1
................gggguuucguGcguagcagagcagcuc.........................................................................   1    1    7y1
```

**Star** **Mature**

```
gaccugcuucugggucgggguuucguacguagcagagcagcucccucgcugcgaucuauugaaagucagcccucgacacaaggguuuguccgcgcgcgcgcgcgcgcgcgcgugc
................gggUuuucguacguagcagagcagcuc........................................................................   1   1   7y1
................gggguuGcguacguagcagagcagcuc........................................................................   1   1   7y1
................gggguuucguacAuagcagagcagcuc........................................................................   2   1   7y1
................ggggAuucguacguagcagagcagcuc........................................................................   1   1   7y1
................gggguuAcguacguagcagagcagcuc........................................................................   4   1   7y1
................gggguuucguacguGgcagagcagcuc........................................................................   1   1   7y1
................gggguuucAuacguagcagagcagcuc........................................................................   1   1   7y1
................gggguuuAguacguagcagagcagcuc........................................................................   8   1   7y1
................gggguAucguacguagcagagcagcuc........................................................................   9   1   7y1
................gggguuucguaAguagcagagcagcuc........................................................................   3   1   7y1
................gggguuucCuacguagcagagcagcuc........................................................................   1   1   7y1
................gggguuucguacUuagcagagcagcuc........................................................................   3   1   7y1
................gggguuucguacguUgcagagcagcuc........................................................................   1   1   7y1
................gggguuucgAacguagcagagcagcuc........................................................................   2   1   7y1
................gggguCucguacguagcagagcagcuc........................................................................   2   1   7y1
.................ggguuucguacgCagcaga...............................................................................   1   1   7y1
.................ggguuucguaAguagcaga...............................................................................   1   1   7y1
.................ggguuuAguacguagcaga...............................................................................   1   1   7y1
.................ggguuAcguacguagcaga...............................................................................   1   1   7y1
.................ggguuucAuacguagcagagc.............................................................................   1   1   7y1
.................ggguuAcguacguagcagagc.............................................................................   1   1   7y1
.................ggguuucguaAguagcagagc.............................................................................   1   1   7y1
.................ggguAucguacguagcagagc.............................................................................   2   1   7y1
.................ggguuuAguacguagcagagc.............................................................................   5   1   7y1
.................gggAuucguacguagcagagc.............................................................................   1   1   7y1
.................ggguuucgAacguagcagagc.............................................................................   2   1   7y1
.................ggguuucguaUguagcagagc.............................................................................   1   1   7y1
.................gggGuucguacguagcagagc.............................................................................   5   1   7y1
.................ggguuAcguacguagcagagca............................................................................   1   1   7y1
.................ggguuucgGacguagcagagca............................................................................   1   1   7y1
.................ggguuucguacguGgcagagca............................................................................   1   1   7y1
.................ggguuucUuacguagcagagca............................................................................   1   1   7y1
.................ggguGucguacguagcagagca............................................................................   1   1   7y1
.................ggguuCcguacguagcagagca............................................................................   1   1   7y1
.................gggGuucguacguagcagagca............................................................................   1   1   7y1
.................ggguuucguacguaCcagagca............................................................................   1   1   7y1
.................ggguuuAguacguagcagagca............................................................................   3   1   7y1
.................ggCuuucguacguagcagagca............................................................................   269   1   7y1
.................ggguuucguacguaUcagagca............................................................................   1   1   7y1
.................ggguuucgAacguagcagagca............................................................................   1   1   7y1
.................ggguCucguacguagcagagca............................................................................   1   1   7y1
.................ggguAucguacguagcagagca............................................................................   1   1   7y1
.................ggguuucguaAguagcagagca............................................................................   1   1   7y1
.................ggguuuGguacguagcagagca............................................................................   1   1   7y1
.................ggguuucguaAguagcagagcagc..........................................................................   1   1   7y1
.................gggGuucguacguagcagagcagc..........................................................................   1   1   7y1
.................ggguuucgAacguagcagagcagc..........................................................................   1   1   7y1
.................ggguuuAguacguagcagagcagc..........................................................................   2   1   7y1
.................gggGuucguacguagcagagcagcu.........................................................................   2   1   7y1
.................ggguuucguacCuagcagagcagcu.........................................................................   1   1   7y1
.................ggguuAcguacguagcagagcagcu.........................................................................   5   1   7y1
.................ggguAucguacguagcagagcagcu.........................................................................   2   1   7y1
.................ggguuucguaUguagcagagcagcu.........................................................................   1   1   7y1
.................ggguuuAguacguagcagagcagcu.........................................................................   6   1   7y1
.................ggCuuucguacguagcagagcagcu.........................................................................   1   1   7y1
.................ggguuucguacguaUcagagcagcu.........................................................................   1   1   7y1
.................ggguuucCuacguagcagagcagcu.........................................................................   1   1   7y1
.................ggguuucguaAguagcagagcagcu.........................................................................   1   1   7y1
.................ggguuucgAacguagcagagcagcu.........................................................................   1   1   7y1
.................gggGuucguacguagcagagcagcuc........................................................................   2   1   7y1
.................ggguuucguaAguagcagagcagcuc........................................................................   2   1   7y1
.................ggguuCcguacguagcagagcagcuc........................................................................   1   1   7y1
.................ggguuucguacUuagcagagcagcuc........................................................................   1   1   7y1
.................ggguuAcguacguagcagagcagcuc........................................................................   6   1   7y1
.................ggguuuAguacguagcagagcagcuc........................................................................   9   1   7y1
.................ggguuucAuacguagcagagcagcuc........................................................................   2   1   7y1
.................ggguAucguacguagcagagcagcuc........................................................................   5   1   7y1
.................ggguuucguacgAagcagagcagcuc........................................................................   2   1   7y1
.................ggguuucguacguCgcagagcagcuc........................................................................   1   1   7y1
.................ggguuucguaUguagcagagcagcuc........................................................................   1   1   7y1
```

**Star** **Mature**

| | | | |
|---|---|---|---|
| gaccugcuucugggucgggguuucguacguagcagagcagcucccucgcugcgaucuauugaaagucagcccucgacacaaggguuuguccgcgcgcgcgcgcgcgcgcgcgugc | | | |
| .................ggguuucUuacguagcagagcagcuc........................................................................ | 1 | 1 | 7y1 |
| .................gggCuucguacguagcagagcagcuc........................................................................ | 1 | 1 | 7y1 |
| .................ggguuucguacCuagcagagcagcuc........................................................................ | 1 | 1 | 7y1 |
| .................ggguuucgAacguagcagagcagcuc........................................................................ | 1 | 1 | 7y1 |
| .................ggguuucguacgAagcagagcagcucc....................................................................... | 1 | 1 | 7y1 |
| .................ggguuucguacguGgcagagcagcucccu.................................................................... | 1 | 1 | 7y1 |
| .................ggguuuAguacguagcagagcagcucccu.................................................................... | 1 | 1 | 7y1 |
| .................ggguuAcguacguagcagagcagcucccu.................................................................... | 1 | 1 | 7y1 |
| .................gggGuucguacguagcagagcagcucccu.................................................................... | 1 | 1 | 7y1 |
| .................ggguuucgAacguagcagagcagcucccu.................................................................... | 1 | 1 | 7y1 |
| ..................gUuuucguacguagcagagc............................................................................. | 3 | 1 | 7y1 |
| ..................gguuucguacguagAagagc............................................................................. | 5 | 1 | 7y1 |
| ..................gguuucguacgGagcagagc............................................................................. | 1 | 1 | 7y1 |
| ..................gguuucguUcguagcagagc............................................................................. | 1 | 1 | 7y1 |
| ..................ggAuucguacguagcagagc............................................................................. | 4 | 1 | 7y1 |
| ..................gguuuGguacguagcagagc............................................................................. | 2 | 1 | 7y1 |
| ..................gguuAcguacguagcagagc............................................................................. | 1 | 1 | 7y1 |
| ..................gguGucguacguagcagagc............................................................................. | 1 | 1 | 7y1 |
| ..................gCuuucguacguagcagagc............................................................................. | 2 | 1 | 7y1 |
| ..................gguuucguaGguagcagagc............................................................................. | 1 | 1 | 7y1 |
| ..................gguuucguacguaUcagagc............................................................................. | 2 | 1 | 7y1 |
| ..................gguuucguaAguagcagagc............................................................................. | 16 | 1 | 7y1 |
| ..................gguuucgAacguagcagagc............................................................................. | 9 | 1 | 7y1 |
| ..................gguuucCuacguagcagagc............................................................................. | 4 | 1 | 7y1 |
| ..................gguCucguacguagcagagc............................................................................. | 1 | 1 | 7y1 |
| ..................gguuuAguacguagcagagc............................................................................. | 11 | 1 | 7y1 |
| ..................gguuucguacguagGagagc............................................................................. | 1 | 1 | 7y1 |
| ..................gguuucguacguUgcagagc............................................................................. | 1 | 1 | 7y1 |
| ..................gguuucguacguaCcagagc............................................................................. | 3 | 1 | 7y1 |
| ..................gguuucAuacguagcagagc............................................................................. | 3 | 1 | 7y1 |
| ..................gguuucguacguGgcagagc............................................................................. | 1 | 1 | 7y1 |
| ..................gguuucguacUuagcagagc............................................................................. | 1 | 1 | 7y1 |
| ..................gguAucguacguagcagagc............................................................................. | 3 | 1 | 7y1 |
| ..................gguuucguacCuagcagagc............................................................................. | 2 | 1 | 7y1 |
| ..................ggGuucguacguagcagagc............................................................................. | 1 | 1 | 7y1 |
| ..................gguuucguacgAagcagagc............................................................................. | 5 | 1 | 7y1 |
| ..................gguuucCuacguagcagagca............................................................................ | 2 | 1 | 7y1 |
| ..................gCuuucguacguagcagagca............................................................................ | 1 | 1 | 7y1 |
| ..................gguuAcguacguagcagagca............................................................................ | 1 | 1 | 7y1 |
| ..................gguuucguaUguagcagagca............................................................................ | 1 | 1 | 7y1 |
| ..................gguuuAguacguagcagagca............................................................................ | 3 | 1 | 7y1 |
| ..................gguuucgGacguagcagagca............................................................................ | 1 | 1 | 7y1 |
| ..................gguuucguacguagUagagca............................................................................ | 1 | 1 | 7y1 |
| ..................gguuucguaAguagcagagca............................................................................ | 3 | 1 | 7y1 |
| ..................gguuucguacCuagcagagca............................................................................ | 1 | 1 | 7y1 |
| ..................ggAuucguacguagcagagca............................................................................ | 1 | 1 | 7y1 |
| ..................gguuucguacguaAcagagca............................................................................ | 1 | 1 | 7y1 |
| ..................gguuucgAacguagcagagca............................................................................ | 1 | 1 | 7y1 |
| ..................gguCucguacguagcagagca............................................................................ | 1 | 1 | 7y1 |
| ..................gguuucguacguaCcagagca............................................................................ | 2 | 1 | 7y1 |
| ..................gUuuucguacguagcagagcag........................................................................... | 1 | 1 | 7y1 |
| ..................gguuucgAacguagcagagcag........................................................................... | 2 | 1 | 7y1 |
| ..................gguAucguacguagcagagcag........................................................................... | 1 | 1 | 7y1 |
| ..................gguuuAguacguagcagagcag........................................................................... | 1 | 1 | 7y1 |
| ..................ggAuucguacguagcagagcag........................................................................... | 1 | 1 | 7y1 |
| ..................gguuucCuacguagcagagcag........................................................................... | 1 | 1 | 7y1 |
| ..................gguuucguaGguagcagagcagc.......................................................................... | 2 | 1 | 7y1 |
| ..................gguuucguacguagGagagcagc.......................................................................... | 3 | 1 | 7y1 |
| ..................gCuuucguacguagcagagcagc.......................................................................... | 3 | 1 | 7y1 |
| ..................gguuucguaUguagcagagcagc.......................................................................... | 1 | 1 | 7y1 |
| ..................gguuucguacguaCcagagcagc.......................................................................... | 2 | 1 | 7y1 |
| ..................gAuuucguacguagcagagcagc.......................................................................... | 1 | 1 | 7y1 |
| ..................gUuuucguacguagcagagcagc.......................................................................... | 6 | 1 | 7y1 |
| ..................gguuucguacguagUagagcagc.......................................................................... | 2 | 1 | 7y1 |
| ..................gguuucguacUuagcagagcagc.......................................................................... | 3 | 1 | 7y1 |
| ..................gguuucguacCuagcagagcagc.......................................................................... | 1 | 1 | 7y1 |
| ..................gguuucguacgCagcagagcagc.......................................................................... | 1 | 1 | 7y1 |
| ..................gguuucguacguaUcagagcagc.......................................................................... | 4 | 1 | 7y1 |
| ..................gguuucguacguagAagagcagc.......................................................................... | 4 | 1 | 7y1 |
| ..................gguAucguacguagcagagcagc.......................................................................... | 2 | 1 | 7y1 |

**Star** **Mature**

| | | | |
|---|---|---|---|
| gaccugcuucugggucggggguuucguacguagcagagcagcucccucgcugcgaucuauugaaagucagcccucgacacaaggguuuguccgcgcgcgcgcgcgcgcgcgcgugc | | | |
| .................ggGuucguacguagcagagcagc........................................................................... | 1 | 1 | 7y1 |
| .................ggAuucguacguagcagagcagc........................................................................... | 7 | 1 | 7y1 |
| .................gguuucgAacguagcagagcagc........................................................................... | 8 | 1 | 7y1 |
| .................gguuucCuacguagcagagcagc........................................................................... | 3 | 1 | 7y1 |
| .................gguuuAguacguagcagagcagc........................................................................... | 7 | 1 | 7y1 |
| .................gguCucguacguagcagagcagc........................................................................... | 2 | 1 | 7y1 |
| .................gguuucguaAguagcagagcagc........................................................................... | 5 | 1 | 7y1 |
| .................gguuucguacgAagcagagcagc........................................................................... | 6 | 1 | 7y1 |
| .................gguuucUuacguagcagagcagc........................................................................... | 4 | 1 | 7y1 |
| .................gguuucguacguGgcagagcagcu.......................................................................... | 3 | 1 | 7y1 |
| .................gguuucguGcguagcagagcagcu.......................................................................... | 1 | 1 | 7y1 |
| .................gguuucguacguagAagagcagcu.......................................................................... | 10 | 1 | 7y1 |
| .................gguuAcguacguagcagagcagcu.......................................................................... | 5 | 1 | 7y1 |
| .................gguuuAguacguagcagagcagcu.......................................................................... | 20 | 1 | 7y1 |
| .................gguuucAuacguagcagagcagcu.......................................................................... | 2 | 1 | 7y1 |
| .................gguuucguacUuagcagagcagcu.......................................................................... | 1 | 1 | 7y1 |
| .................gguuucCuacguagcagagcagcu.......................................................................... | 2 | 1 | 7y1 |
| .................gguuucgAacguagcagagcagcu.......................................................................... | 10 | 1 | 7y1 |
| .................gUuuucguacguagcagagcagcu.......................................................................... | 4 | 1 | 7y1 |
| .................gguuucguacCuagcagagcagcu.......................................................................... | 7 | 1 | 7y1 |
| .................gguuuGguacguagcagagcagcu.......................................................................... | 2 | 1 | 7y1 |
| .................gguuucguacgCagcagagcagcu.......................................................................... | 1 | 1 | 7y1 |
| .................gguuuUguacguagcagagcagcu.......................................................................... | 1 | 1 | 7y1 |
| .................gguuucguUcguagcagagcagcu.......................................................................... | 2 | 1 | 7y1 |
| .................gguuucguaAguagcagagcagcu.......................................................................... | 20 | 1 | 7y1 |
| .................gguuGcguacguagcagagcagcu.......................................................................... | 1 | 1 | 7y1 |
| .................gguuucUuacguagcagagcagcu.......................................................................... | 4 | 1 | 7y1 |
| .................gguuucguacgAagcagagcagcu.......................................................................... | 6 | 1 | 7y1 |
| .................gguuucguacguagGagagcagcu.......................................................................... | 2 | 1 | 7y1 |
| .................gguAucguacguagcagagcagcu.......................................................................... | 5 | 1 | 7y1 |
| .................gCuuucguacguagcagagcagcu.......................................................................... | 2 | 1 | 7y1 |
| .................ggAuucguacguagcagagcagcu.......................................................................... | 6 | 1 | 7y1 |
| .................gguuucguaGguagcagagcagcu.......................................................................... | 1 | 1 | 7y1 |
| .................gguuucguacAuagcagagcagcu.......................................................................... | 4 | 1 | 7y1 |
| .................gguuucguacguaUcagagcagcu.......................................................................... | 4 | 1 | 7y1 |
| .................gguuucguacguaAcagagcagcu.......................................................................... | 1 | 1 | 7y1 |
| .................ggCuucguacguagcagagcagcu.......................................................................... | 1 | 1 | 7y1 |
| .................gguuCcguacguagcagagcagcu.......................................................................... | 3 | 1 | 7y1 |
| .................gguuucguacguagUagagcagcuc......................................................................... | 1 | 1 | 7y1 |
| .................gCuuucguacguagcagagcagcuc......................................................................... | 3 | 1 | 7y1 |
| .................gguuucguacguaUcagagcagcuc......................................................................... | 1 | 1 | 7y1 |
| .................gguuucguacAuagcagagcagcuc......................................................................... | 2 | 1 | 7y1 |
| .................gguuucgCacguagcagagcagcuc......................................................................... | 1 | 1 | 7y1 |
| .................gguuucCuacguagcagagcagcuc......................................................................... | 1 | 1 | 7y1 |
| .................gguuucguaUguagcagagcagcuc......................................................................... | 1 | 1 | 7y1 |
| .................ggGuucguacguagcagagcagcuc......................................................................... | 2 | 1 | 7y1 |
| .................gguuucguaAguagcagagcagcuc......................................................................... | 6 | 1 | 7y1 |
| .................gguuucguacgAagcagagcagcuc......................................................................... | 4 | 1 | 7y1 |
| .................gguuucUuacguagcagagcagcuc......................................................................... | 1 | 1 | 7y1 |
| .................gguuucAuacguagcagagcagcuc......................................................................... | 1 | 1 | 7y1 |
| .................gguuuAguacguagcagagcagcuc......................................................................... | 3 | 1 | 7y1 |
| .................gguuucguacCuagcagagcagcuc......................................................................... | 2 | 1 | 7y1 |
| .................gguuucguacCuagcagagcagcucc........................................................................ | 2 | 1 | 7y1 |
| .................gguuucguCcguagcagagcagcucc........................................................................ | 1 | 1 | 7y1 |
| .................gguuucgCacguagcagagcagcucc........................................................................ | 1 | 1 | 7y1 |
| .................gguuucUuacguagcagagcagcucc........................................................................ | 1 | 1 | 7y1 |
| .................gguuucguacguaUcagagcagcucc........................................................................ | 1 | 1 | 7y1 |
| .................gUuuucguacguagcagagcagcucc........................................................................ | 1 | 1 | 7y1 |
| .................gguuAcguacguagcagagcagcucc........................................................................ | 2 | 1 | 7y1 |
| .................gguAucguacguagcagagcagcucc........................................................................ | 1 | 1 | 7y1 |
| .................gguuucguacguagAagagcagcucc........................................................................ | 1 | 1 | 7y1 |
| .................ggAuucguacguagcagagcagcucc........................................................................ | 4 | 1 | 7y1 |
| .................gguuucguNcguagcagagcagcucc........................................................................ | 1 | 1 | 7y1 |
| .................ggGuucguacguagcagagcagcucc........................................................................ | 2 | 1 | 7y1 |
| .................gguuuAguacguagcagagcagcucc........................................................................ | 5 | 1 | 7y1 |
| .................gguuucguacgAagcagagcagcucc........................................................................ | 2 | 1 | 7y1 |
| .................gCuuucguacguagcagagcagcucc........................................................................ | 1 | 1 | 7y1 |
| .................gguuucCuacguagcagagcagcucc........................................................................ | 1 | 1 | 7y1 |
| .................gguuucguacAuagcagagcagcucc........................................................................ | 1 | 1 | 7y1 |
| .................gguuGcguacguagcagagcagcucc........................................................................ | 1 | 1 | 7y1 |

**Star** **Mature**

| | | | |
|---|---|---|---|
| gaccugcuucugggucgggguuucguacguagcagagcagcucccucgcugcgaucuauugaaagucagcccucgacacaaggguuuguccgcgcgcgcgcgcgcgcgcgugc | | | |
| ..................gguuucguUcguagcagagcagcucc..................................................................... | 1 | 1 | 7y1 |
| ..................gguuucguaAguagcagagcagcucc..................................................................... | 2 | 1 | 7y1 |
| ..................gguuucguacAuagcagagcagcuccc.................................................................... | 1 | 1 | 7y1 |
| ..................gguAucguacguagcagagcagcuccc.................................................................... | 2 | 1 | 7y1 |
| ..................ggCuucguacguagcagagcagcuccc.................................................................... | 1 | 1 | 7y1 |
| ..................gguuucguacguaUcagagcagcuccc.................................................................... | 1 | 1 | 7y1 |
| ..................gguuucguacgAagcagagcagcuccc.................................................................... | 2 | 1 | 7y1 |
| ..................gguuuUguacguagcagagcagcuccc.................................................................... | 1 | 1 | 7y1 |
| ..................gguuucUuacguagcagagcagcuccc.................................................................... | 4 | 1 | 7y1 |
| ..................gCuuucguacguagcagagcagcuccc.................................................................... | 1 | 1 | 7y1 |
| ..................gguuucCuacguagcagagcagcuccc.................................................................... | 3 | 1 | 7y1 |
| ..................gguuucguacUuagcagagcagcuccc.................................................................... | 1 | 1 | 7y1 |
| ..................gUuuucguacguagcagagcagcuccc.................................................................... | 2 | 1 | 7y1 |
| ..................gguuGcguacguagcagagcagcuccc.................................................................... | 1 | 1 | 7y1 |
| ..................gguuucguacgGagcagagcagcuccc.................................................................... | 1 | 1 | 7y1 |
| ..................gguuucguacguagGagagcagcuccc.................................................................... | 1 | 1 | 7y1 |
| ..................ggGuucguacguagcagagcagcuccc.................................................................... | 3 | 1 | 7y1 |
| ..................gguuucgGacguagcagagcagcuccc.................................................................... | 2 | 1 | 7y1 |
| ..................gguuuAguacguagcagagcagcuccc.................................................................... | 15 | 1 | 7y1 |
| ..................gguuucguacgCagcagagcagcuccc.................................................................... | 1 | 1 | 7y1 |
| ..................ggAuucguacguagcagagcagcuccc.................................................................... | 3 | 1 | 7y1 |
| ..................gguuucguacguaCcagagcagcuccc.................................................................... | 3 | 1 | 7y1 |
| ..................gguuucguacguagAagagcagcuccc.................................................................... | 5 | 1 | 7y1 |
| ..................gguuucguaAguagcagagcagcuccc.................................................................... | 11 | 1 | 7y1 |
| ..................gguuucguacCuagcagagcagcuccc.................................................................... | 4 | 1 | 7y1 |
| ..................gguuucgAacguagcagagcagcuccc.................................................................... | 2 | 1 | 7y1 |
| ..................gguuucguaGguagcagagcagcuccc.................................................................... | 2 | 1 | 7y1 |
| ..................gguuucguacguCgcagagcagcucccu................................................................... | 1 | 1 | 7y1 |
| ..................gguuucgCacguagcagagcagcucccu................................................................... | 1 | 1 | 7y1 |
| ..................gguuucguacgAagcagagcagcucccu................................................................... | 5 | 1 | 7y1 |
| ..................gguuuUguacguagcagagcagcucccu................................................................... | 1 | 1 | 7y1 |
| ..................gguuucguacCuagcagagcagcucccu................................................................... | 2 | 1 | 7y1 |
| ..................gCuuucguacguagcagagcagcucccu................................................................... | 2 | 1 | 7y1 |
| ..................gguuucguacguGgcagagcagcucccu................................................................... | 2 | 1 | 7y1 |
| ..................ggCuucguacguagcagagcagcucccu................................................................... | 1 | 1 | 7y1 |
| ..................gAuuucguacguagcagagcagcucccu................................................................... | 1 | 1 | 7y1 |
| ..................gguuucguacguaAcagagcagcucccu................................................................... | 2 | 1 | 7y1 |
| ..................gguuucguacguagAagagcagcucccu................................................................... | 5 | 1 | 7y1 |
| ..................gguuucgAacguagcagagcagcucccu................................................................... | 7 | 1 | 7y1 |
| ..................gguuucguCcguagcagagcagcucccu................................................................... | 2 | 1 | 7y1 |
| ..................gguuucguacguagGagagcagcucccu................................................................... | 1 | 1 | 7y1 |
| ..................gguuucguacguaUcagagcagcucccu................................................................... | 3 | 1 | 7y1 |
| ..................ggGuucguacguagcagagcagcucccu................................................................... | 2 | 1 | 7y1 |
| ..................gguuucguGcguagcagagcagcucccu................................................................... | 1 | 1 | 7y1 |
| ..................gguuucUuacguagcagagcagcucccu................................................................... | 5 | 1 | 7y1 |
| ..................gguuucguaAguagcagagcagcucccu................................................................... | 12 | 1 | 7y1 |
| ..................gguuuAguacguagcagagcagcucccu................................................................... | 9 | 1 | 7y1 |
| ..................gguuucguaUguagcagagcagcucccu................................................................... | 1 | 1 | 7y1 |
| ..................gguuucAuacguagcagagcagcucccu................................................................... | 1 | 1 | 7y1 |
| ..................gUuuucguacguagcagagcagcucccu................................................................... | 1 | 1 | 7y1 |
| ..................gguuAcguacguagcagagcagcucccu................................................................... | 2 | 1 | 7y1 |
| ..................gguAucguacguagcagagcagcucccu................................................................... | 3 | 1 | 7y1 |
| ..................ggAuucguacguagcagagcagcucccu................................................................... | 4 | 1 | 7y1 |
| ..................gguuucguacUuagcagagcagcucccu................................................................... | 1 | 1 | 7y1 |
| ..................gguuucguacguagAagagcagcucccuc.................................................................. | 1 | 1 | 7y1 |
| ..................gguuucCuacguagcagagcagcucccuc.................................................................. | 2 | 1 | 7y1 |
| ..................gguuucguacgAagcagagcagcucccuc.................................................................. | 1 | 1 | 7y1 |
| ..................gguuuAguacguagcagagcagcucccuc.................................................................. | 1 | 1 | 7y1 |
| ..................gguuucgAacguagcagagcagcucccuc.................................................................. | 2 | 1 | 7y1 |
| ..................gguuucUuacguagcagagcagcucccuc.................................................................. | 1 | 1 | 7y1 |
| ..................gguuAcguacguagcagagcagcucccuc.................................................................. | 1 | 1 | 7y1 |
| ..................ggAuucguacguagcagagcagcucccuc.................................................................. | 2 | 1 | 7y1 |
| ..................gguuucguacguaCcagagcagcucccuc.................................................................. | 1 | 1 | 7y1 |
| ..................gguuucguacUuagcagagcagcucccuc.................................................................. | 2 | 1 | 7y1 |
| ..................gguuucguacguaUcagagcagcucccuc.................................................................. | 1 | 1 | 7y1 |
| ..................gguuucguaGguagcagagcagcucccuc.................................................................. | 1 | 1 | 7y1 |
| ..................gguuucguGcguagcagagcagcucccuc.................................................................. | 1 | 1 | 7y1 |
| ..................gguuucgGacguagcagagcagcucccuc.................................................................. | 1 | 1 | 7y1 |
| ..................gguuuGguacguagcagagcagcucccuc.................................................................. | 1 | 1 | 7y1 |
| ..................gguuucguacguGgcagagcagcucccuc.................................................................. | 1 | 1 | 7y1 |

**Star** **Mature**

gaccugcuucugggucgggguuucguacguagcagagcagcucccucgcugcgaucuauugaaagucagcccucgacacaaggguuuguccgcgcgcgcgcgcgcgcgcgugc

| | | | |
|---|---|---|---|
| ..................gguuucguacAuagcagagcagcucccuc.................................................................. | 2 | 1 | 7y1 |
| ..................gCuuucguacguagcagagcagcucccuc.................................................................. | 1 | 1 | 7y1 |
| ..................gguuucguaAguagcagagcagcucccuc.................................................................. | 1 | 1 | 7y1 |
| ..................ggCuucguacguagcagagcagcucccuc.................................................................. | 1 | 1 | 7y1 |
| ..................gguuucguacguagAagagcagcucccucg................................................................. | 2 | 1 | 7y1 |
| ..................gguuucguacguaCcagagcagcucccucg................................................................. | 1 | 1 | 7y1 |
| ..................gguuucgGacguagcagagcagcucccucg................................................................. | 1 | 1 | 7y1 |
| ..................gguuucguacguagUagagcagcucccucg................................................................. | 1 | 1 | 7y1 |
| ..................gguuucCuacguagcagagcagcucccucg................................................................. | 2 | 1 | 7y1 |
| ..................gguuucguacgAagcagagcagcucccucg................................................................. | 3 | 1 | 7y1 |
| ..................gguuucguacguagGagagcagcucccucg................................................................. | 1 | 1 | 7y1 |
| ..................gguuucguaAguagcagagcagcucccucg................................................................. | 1 | 1 | 7y1 |
| ..................gguuucguaUguagcagagcagcucccucg................................................................. | 2 | 1 | 7y1 |
| ..................gguuuAguacguagcagagcagcucccucg................................................................. | 3 | 1 | 7y1 |
| ..................gguuAcguacguagcagagcagcucccucg................................................................. | 1 | 1 | 7y1 |
| ..................gguAucguacguagcagagcagcucccucg................................................................. | 2 | 1 | 7y1 |
| ..................ggAuucguacguagcagagcagcucccucg................................................................. | 3 | 1 | 7y1 |
| ..................gguuucguUcguagcagagcagcucccucg................................................................. | 1 | 1 | 7y1 |
| ..................gCuuucguacguagcagagcagcucccucg................................................................. | 1 | 1 | 7y1 |
| ..................gguuucguacguaAcagagcagcucccucgc................................................................ | 1 | 1 | 7y1 |
| ..................gguCucguacguagcagagcagcucccucgc................................................................ | 1 | 1 | 7y1 |
| ..................gCuuucguacguagcagagcagcucccucgc................................................................ | 1 | 1 | 7y1 |
| ..................gUuuucguacguagcagagcagcucccucgc................................................................ | 2 | 1 | 7y1 |
| ..................gguuucguacUuagcagagcagcucccucgc................................................................ | 1 | 1 | 7y1 |
| ..................gguuucgAacguagcagagcagcucccucgc................................................................ | 4 | 1 | 7y1 |
| ..................gguuuAguacguagcagagcagcucccucgc................................................................ | 1 | 1 | 7y1 |
| ..................gguuucAuacguagcagagcagcucccucgc................................................................ | 1 | 1 | 7y1 |
| ..................gguuCcguacguagcagagcagcucccucgc................................................................ | 1 | 1 | 7y1 |
| ..................ggGuucguacguagcagagcagcucccucgc................................................................ | 2 | 1 | 7y1 |
| ..................gguuucguacguGgcagagcagcucccucgc................................................................ | 1 | 1 | 7y1 |
| ..................gguuucguacguagUagagcagcucccucgc................................................................ | 1 | 1 | 7y1 |
| ..................gguGucguacguagcagagcagcucccucgc................................................................ | 1 | 1 | 7y1 |
| ..................gguuucguacgAagcagagcagcucccucgc................................................................ | 2 | 1 | 7y1 |
| ..................gguuucguaAguagcagagcagcucccucgc................................................................ | 5 | 1 | 7y1 |
| ..................ggAuucguacguagcagagcagcucccucgc................................................................ | 2 | 1 | 7y1 |
| ..................gguAucguacguagcagagcagcucccucgc................................................................ | 1 | 1 | 7y1 |
| ..................gguuucCuacguagcagagcagcucccucgc................................................................ | 4 | 1 | 7y1 |
| ..................gguuucguacCuagcagagcagcucccucgcu............................................................... | 1 | 1 | 7y1 |
| ..................gguuucguGcguagcagagcagcucccucgcu............................................................... | 1 | 1 | 7y1 |
| ..................gguuucguacguagAagagcagcucccucgcu............................................................... | 1 | 1 | 7y1 |
| ..................gguuucUuacguagcagagcagcucccucgcu............................................................... | 1 | 1 | 7y1 |
| ..................gguuucguacguaUcagagcagcucccucgcu............................................................... | 1 | 1 | 7y1 |
| ..................gUuuucguacguagcagagcagcucccucgcu............................................................... | 1 | 1 | 7y1 |
| ..................gguuucAuacguagcagagcagcucccucgcu............................................................... | 1 | 1 | 7y1 |
| ..................ggAuucguacguagcagagcagcucccucgcu............................................................... | 1 | 1 | 7y1 |
| ..................gguuucguacguaCcagagcagcucccucgcu............................................................... | 1 | 1 | 7y1 |
| ...................guuGcguacguagcagagc........................................................................... | 1 | 1 | 7y1 |
| ...................Uuuucguacguagcagagc........................................................................... | 1 | 1 | 7y1 |
| ...................guuucgAacguagcagagc........................................................................... | 3 | 1 | 7y1 |
| ...................guuucguaAguagcagagc........................................................................... | 5 | 1 | 7y1 |
| ...................Cuuucguacguagcagagc........................................................................... | 3 | 1 | 7y1 |
| ...................guuuAguacguagcagagc........................................................................... | 1 | 1 | 7y1 |
| ...................guuucUuacguagcagagc........................................................................... | 1 | 1 | 7y1 |
| ...................guuucCuacguagcagagc........................................................................... | 1 | 1 | 7y1 |
| ...................Nuuucguacguagcagagc........................................................................... | 1 | 1 | 7y1 |
| ...................guuAcguacguagcagagca.......................................................................... | 1 | 1 | 7y1 |
| ...................guuCcguacguagcagagca.......................................................................... | 1 | 1 | 7y1 |
| ...................guuuAguacguagcagagca.......................................................................... | 1 | 1 | 7y1 |
| ...................guuucguacguagAagagca.......................................................................... | 2 | 1 | 7y1 |
| ...................gAuucguacguagcagagca.......................................................................... | 1 | 1 | 7y1 |
| ...................guuucguacCuagcagagca.......................................................................... | 2 | 1 | 7y1 |
| ...................Cuuucguacguagcagagca.......................................................................... | 2 | 1 | 7y1 |
| ...................Uuuucguacguagcagagca.......................................................................... | 1 | 1 | 7y1 |
| ...................guuucgAacguagcagagca.......................................................................... | 5 | 1 | 7y1 |
| ...................guAucguacguagcagagca.......................................................................... | 1 | 1 | 7y1 |
| ...................guuucCuacguagcagagca.......................................................................... | 2 | 1 | 7y1 |
| ...................guuucguacgAagcagagcagc........................................................................ | 1 | 1 | 7y1 |
| ...................guuucguaAguagcagagcagc........................................................................ | 2 | 1 | 7y1 |
| ...................gAuucguacguagcagagcagc........................................................................ | 5 | 1 | 7y1 |
| ...................guAucguacguagcagagcagc........................................................................ | 3 | 1 | 7y1 |

**Star** **Mature**

| | | | |
|---|---|---|---|
| gaccugcuucugggucgggguuucguacguagcagagcagcucccucgcugcgaucuauugaaagucagcccucgacacaaggguuuguccgcgcgcgcgcgcgcgcgugc | | | |
| ..................Uuuucguacguagcagagcagc....................................................................... | 286 | 1 | 7y1 |
| ..................guuucgAacguagcagagcagc....................................................................... | 2 | 1 | 7y1 |
| ..................Auuucguacguagcagagcagc....................................................................... | 1 | 1 | 7y1 |
| ..................guuucguUcguagcagagcagc....................................................................... | 1 | 1 | 7y1 |
| ..................Uuuucguacguagcagagcagcu...................................................................... | 3 | 1 | 7y1 |
| ..................guuucguacguGgcagagcagcu...................................................................... | 1 | 1 | 7y1 |
| ..................gGuucguacguagcagagcagcu...................................................................... | 1 | 1 | 7y1 |
| ..................guuAcguacguagcagagcagcu...................................................................... | 2 | 1 | 7y1 |
| ..................guuucguacguagAagagcagcu...................................................................... | 1 | 1 | 7y1 |
| ..................Cuuucguacguagcagagcagcu...................................................................... | 2 | 1 | 7y1 |
| ..................guuucgAacguagcagagcagcu...................................................................... | 3 | 1 | 7y1 |
| ..................guuucguacUuagcagagcagcu...................................................................... | 1 | 1 | 7y1 |
| ..................Nuuucguacguagcagagcagcu...................................................................... | 3 | 1 | 7y1 |
| ..................guuucUuacguagcagagcagcu...................................................................... | 1 | 1 | 7y1 |
| ..................guAucguacguagcagagcagcu...................................................................... | 4 | 1 | 7y1 |
| ..................gAuucguacguagcagagcagcu...................................................................... | 6 | 1 | 7y1 |
| ..................guuuAguacguagcagagcagcu...................................................................... | 1 | 1 | 7y1 |
| ..................guuucguacgAagcagagcagcu...................................................................... | 3 | 1 | 7y1 |
| ..................gAuucguacguagcagagcagcuc..................................................................... | 2 | 1 | 7y1 |
| ..................gGuucguacguagcagagcagcuc..................................................................... | 1 | 1 | 7y1 |
| ..................guAucguacguagcagagcagcuc..................................................................... | 3 | 1 | 7y1 |
| ..................guCucguacguagcagagcagcuc..................................................................... | 1 | 1 | 7y1 |
| ..................guuuUguacguagcagagcagcuc..................................................................... | 1 | 1 | 7y1 |
| ..................guuucguacgAagcagagcagcucc.................................................................... | 1 | 1 | 7y1 |
| ..................guuucguGcguagcagagcagcucc.................................................................... | 1 | 1 | 7y1 |
| ..................guAucguacguagcagagcagcucc.................................................................... | 1 | 1 | 7y1 |
| ..................Uuuucguacguagcagagcagcucc.................................................................... | 1 | 1 | 7y1 |
| ..................guuucguacguaAcagagcagcucc.................................................................... | 1 | 1 | 7y1 |
| ..................guuucguacguaUcagagcagcucc.................................................................... | 179 | 1 | 7y1 |
| ..................gAuucguacguagcagagcagcuccc................................................................... | 4 | 1 | 7y1 |
| ..................guuAcguacguagcagagcagcuccc................................................................... | 1 | 1 | 7y1 |
| ..................guuucgAacguagcagagcagcuccc................................................................... | 2 | 1 | 7y1 |
| ..................guuucCuacguagcagagcagcuccc................................................................... | 2 | 1 | 7y1 |
| ..................guuucguacguagAagagcagcuccc................................................................... | 5 | 1 | 7y1 |
| ..................gCuucguacguagcagagcagcuccc................................................................... | 2 | 1 | 7y1 |
| ..................guuucguacguCgcagagcagcuccc................................................................... | 1 | 1 | 7y1 |
| ..................guuucguacguaAcagagcagcuccc................................................................... | 2 | 1 | 7y1 |
| ..................guuucguacguaUcagagcagcuccc................................................................... | 1 | 1 | 7y1 |
| ..................guuucguacgGagcagagcagcuccc................................................................... | 1 | 1 | 7y1 |
| ..................guAucguacguagcagagcagcuccc................................................................... | 2 | 1 | 7y1 |
| ..................Cuuucguacguagcagagcagcuccc................................................................... | 2 | 1 | 7y1 |
| ..................guuuUguacguagcagagcagcuccc................................................................... | 1 | 1 | 7y1 |
| ..................guuucguacgAagcagagcagcuccc................................................................... | 1 | 1 | 7y1 |
| ..................guuucguacguaCcagagcagcuccc................................................................... | 3 | 1 | 7y1 |
| ..................guuucguacCuagcagagcagcuccc................................................................... | 3 | 1 | 7y1 |
| ..................guuucguaAguagcagagcagcuccc................................................................... | 2 | 1 | 7y1 |
| ..................guuuAguacguagcagagcagcuccc................................................................... | 3 | 1 | 7y1 |
| ..................guuucguacCuagcagagcagcucccu.................................................................. | 1 | 1 | 7y1 |
| ..................guuucgAacguagcagagcagcucccu.................................................................. | 7 | 1 | 7y1 |
| ..................gAuucguacguagcagagcagcucccu.................................................................. | 7 | 1 | 7y1 |
| ..................guuucguacgGagcagagcagcucccu.................................................................. | 1 | 1 | 7y1 |
| ..................guuucguacguagAagagcagcucccu.................................................................. | 3 | 1 | 7y1 |
| ..................guuuAguacguagcagagcagcucccu.................................................................. | 1 | 1 | 7y1 |
| ..................Uuuucguacguagcagagcagcucccu.................................................................. | 2 | 1 | 7y1 |
| ..................guuucguacAuagcagagcagcucccu.................................................................. | 2 | 1 | 7y1 |
| ..................guuucguacgCagcagagcagcucccu.................................................................. | 1 | 1 | 7y1 |
| ..................guuAcguacguagcagagcagcucccu.................................................................. | 2 | 1 | 7y1 |
| ..................guAucguacguagcagagcagcucccu.................................................................. | 2 | 1 | 7y1 |
| ..................gGuucguacguagcagagcagcucccu.................................................................. | 295 | 1 | 7y1 |
| ..................guuucguacguagcGgagcagcucccu.................................................................. | 2 | 1 | 7y1 |
| ..................guuucCuacguagcagagcagcucccu.................................................................. | 1 | 1 | 7y1 |
| ..................Cuuucguacguagcagagcagcucccu.................................................................. | 3 | 1 | 7y1 |
| ..................guuucguacguagcGgagcagcucccuc................................................................. | 1 | 1 | 7y1 |
| ..................guuucguacAuagcagagcagcucccuc................................................................. | 1 | 1 | 7y1 |
| ..................guuucguaAguagcagagcagcucccuc................................................................. | 1 | 1 | 7y1 |
| ..................guuucguacUuagcagagcagcucccuc................................................................. | 3 | 1 | 7y1 |
| ..................guuCcguacguagcagagcagcucccuc................................................................. | 1 | 1 | 7y1 |
| ..................guuucguGcguagcagagcagcucccuc................................................................. | 2 | 1 | 7y1 |
| ..................gGuucguacguagcagagcagcucccuc................................................................. | 1 | 1 | 7y1 |
| ..................guuucgAacguagcagagcagcucccuc................................................................. | 2 | 1 | 7y1 |

**Star** **Mature**

| | | | |
|---|---|---|---|
| gaccugcuucugggucggggguuucguacguagcagagcagcucccucgcugcgaucuauugaaagucagcccucgacacaaggguuuguccgcgcgcgcgcgcgcgcgcgugc | | | |
| ...................guuucguacCuagcagagcagcucccuc.................................................................... | 1 | 1 | 7y1 |
| ...................guAucguacguagcagagcagcucccuc.................................................................... | 1 | 1 | 7y1 |
| ...................Uuuucguacguagcagagcagcucccuc.................................................................... | 3 | 1 | 7y1 |
| ...................Cuuucguacguagcagagcagcucccuc.................................................................... | 1 | 1 | 7y1 |
| ...................guuucguacguagAagagcagcucccuc.................................................................... | 2 | 1 | 7y1 |
| ...................guuucguacgAagcagagcagcucccuc.................................................................... | 1 | 1 | 7y1 |
| ...................guuucguacguaUcagagcagcucccuc.................................................................... | 1 | 1 | 7y1 |
| ...................guuucgAacguagcagagcagcucccucg................................................................... | 4 | 1 | 7y1 |
| ...................guuucguacguagcGgagcagcucccucg................................................................... | 1 | 1 | 7y1 |
| ...................gAuucguacguagcagagcagcucccucg................................................................... | 1 | 1 | 7y1 |
| ...................guuucguacguaUcagagcagcucccucg................................................................... | 1 | 1 | 7y1 |
| ...................guAucguacguagcagagcagcucccucg................................................................... | 1 | 1 | 7y1 |
| ...................guuucguacAuagcagagcagcucccucg................................................................... | 1 | 1 | 7y1 |
| ...................guuucguacCuagcagagcagcucccucgc.................................................................. | 1 | 1 | 7y1 |
| ...................guuucguacgCagcagagcagcucccucgc.................................................................. | 1 | 1 | 7y1 |
| ...................guuucgAacguagcagagcagcucccucgc.................................................................. | 4 | 1 | 7y1 |
| ...................guuucgGacguagcagagcagcucccucgc.................................................................. | 1 | 1 | 7y1 |
| ...................gGuucguacguagcagagcagcucccucgc.................................................................. | 1 | 1 | 7y1 |
| ...................guuAcguacguagcagagcagcucccucgc.................................................................. | 2 | 1 | 7y1 |
| ...................guuucguacguaUcagagcagcucccucgc.................................................................. | 2 | 1 | 7y1 |
| ...................guuucguacUuagcagagcagcucccucgc.................................................................. | 2 | 1 | 7y1 |
| ...................guuucCuacguagcagagcagcucccucgc.................................................................. | 1 | 1 | 7y1 |
| ...................Nuuucguacguagcagagcagcucccucgc.................................................................. | 1 | 1 | 7y1 |
| ...................Cuuucguacguagcagagcagcucccucgc.................................................................. | 1 | 1 | 7y1 |
| ...................guAucguacguagcagagcagcucccucgc.................................................................. | 2 | 1 | 7y1 |
| ...................guuucUuacguagcagagcagcucccucgc.................................................................. | 1 | 1 | 7y1 |
| ...................Uuuucguacguagcagagcagcucccucgc.................................................................. | 1 | 1 | 7y1 |
| ...................guuucguacguagAagagcagcucccucgc.................................................................. | 1 | 1 | 7y1 |
| ...................guuucguaAguagcagagcagcucccucgc.................................................................. | 3 | 1 | 7y1 |
| ...................gAuucguacguagcagagcagcucccucgc.................................................................. | 1 | 1 | 7y1 |
| ...................guuucguacgAagcagagcagcucccucgc.................................................................. | 1 | 1 | 7y1 |
| ...................gAuucguacguagcagagcagcucccucgcu................................................................. | 2 | 1 | 7y1 |
| ...................Cuuucguacguagcagagcagcucccucgcu................................................................. | 2 | 1 | 7y1 |
| ...................guuucguacguagAagagcagcucccucgcu................................................................. | 1 | 1 | 7y1 |
| ...................guuucguaGguagcagagcagcucccucgcu................................................................. | 1 | 1 | 7y1 |
| ...................guuucUuacguagcagagcagcucccucgcu................................................................. | 1 | 1 | 7y1 |
| ...................guuucguacguagAagagcagcucccucgcug................................................................ | 1 | 1 | 7y1 |
| ...................guuucguacUuagcagagcagcucccucgcug................................................................ | 1 | 1 | 7y1 |
| ...................gAuucguacguagcagagcagcucccucgcug................................................................ | 1 | 1 | 7y1 |
| ....................uuucUuacguagcagagc............................................................................. | 1 | 1 | 7y1 |
| ....................uuucAuacguagcagagc............................................................................. | 1 | 1 | 7y1 |
| ....................uuucguacguagcUgagc............................................................................. | 2 | 1 | 7y1 |
| ....................uuucguacgCagcagagc............................................................................. | 1 | 1 | 7y1 |
| ....................uuucguacguagcagaUc............................................................................. | 5 | 1 | 7y1 |
| ....................uuucgAacguagcagagc............................................................................. | 8 | 1 | 7y1 |
| ....................Cuucguacguagcagagc............................................................................. | 2 | 1 | 7y1 |
| ....................uuucguacCuagcagagc............................................................................. | 4 | 1 | 7y1 |
| ....................Auucguacguagcagagc............................................................................. | 52 | 1 | 7y1 |
| ....................uuucguacguagcaUagc............................................................................. | 1 | 1 | 7y1 |
| ....................uuAcguacguagcagagc............................................................................. | 2 | 1 | 7y1 |
| ....................uAucguacguagcagagc............................................................................. | 9 | 1 | 7y1 |
| ....................uuuAguacguagcagagc............................................................................. | 5 | 1 | 7y1 |
| ....................uuucguacguagcagaCc............................................................................. | 2 | 1 | 7y1 |
| ....................uuucguacUuagcagagc............................................................................. | 1 | 1 | 7y1 |
| ....................uuucguacguGgcagagc............................................................................. | 2 | 1 | 7y1 |
| ....................uuucCuacguagcagagc............................................................................. | 192 | 1 | 7y1 |
| ....................uuucguaUguagcagagc............................................................................. | 1 | 1 | 7y1 |
| ....................uuucguacguagcagagA............................................................................. | 7 | 1 | 7y1 |
| ....................uuucguacguagAagagc............................................................................. | 4 | 1 | 7y1 |
| ....................uuucguacguaCcagagc............................................................................. | 1 | 1 | 7y1 |
| ....................uuucguacgAagcagagc............................................................................. | 11 | 1 | 7y1 |
| ....................uuucguacguagcGgagc............................................................................. | 2 | 1 | 7y1 |
| ....................uuucgGacguagcagagc............................................................................. | 1 | 1 | 7y1 |
| ....................Guucguacguagcagagc............................................................................. | 9 | 1 | 7y1 |
| ....................uuuGguacguagcagagc............................................................................. | 3 | 1 | 7y1 |
| ....................uuucguaAguagcagagc............................................................................. | 8 | 1 | 7y1 |
| ....................uuucguacguaUcagagc............................................................................. | 2 | 1 | 7y1 |
| ....................Nuucguacguagcagagc............................................................................. | 3 | 1 | 7y1 |
| ....................uuucguacguagcagagc............................................................................. | 4344 | 0 | 7y1 |
| ....................uuucguaGguagcagagca............................................................................ | 1 | 1 | 7y1 |

```
  Star                                                                  Mature
gaccugcuucugggucgggguuucguacguagcagagcagcucccucgcugcgaucuauugaaagucagcccucgacacaaggguuuguccgcgcgcgcgcgcgcgcgcgugc
....................uuucguacguaUcagagca..........................................................................    2       1       7y1
....................uuucguacguagcagagAa..........................................................................    4       1       7y1
....................uuucguacguagAagagca..........................................................................    3       1       7y1
....................uuucguacgCagcagagca..........................................................................    1       1       7y1
....................uuucguacCuagcagagca..........................................................................    2       1       7y1
....................uCucguacguagcagagca..........................................................................    1       1       7y1
....................uuucguacguagcagCgca..........................................................................    3       1       7y1
....................uuucguacgGagcagagca..........................................................................    1       1       7y1
....................uuucguacguagcGgagca..........................................................................    1       1       7y1
....................uuucCuacguagcagagca..........................................................................    1       1       7y1
....................uuuAguacguagcagagca..........................................................................    2       1       7y1
....................uuucguaAguagcagagca..........................................................................    2       1       7y1
....................Nuucguacguagcagagca..........................................................................    3       1       7y1
....................uuucguCcguagcagagca..........................................................................    1       1       7y1
....................Auucguacguagcagagca..........................................................................    24      1       7y1
....................uuucguacguagcagaUca..........................................................................    1       1       7y1
....................uuucguacguaCcagagca..........................................................................    1       1       7y1
....................uuucguacUuagcagagca..........................................................................    1       1       7y1
....................uuucguacguagcagagca..........................................................................    1926    0       7y1
....................uuucguacguagcUgagca..........................................................................    1       1       7y1
....................Guucguacguagcagagca..........................................................................    4       1       7y1
....................uuucgCacguagcagagca..........................................................................    1       1       7y1
....................uuucguGcguagcagagca..........................................................................    1       1       7y1
....................uuucguacguUgcagagca..........................................................................    2       1       7y1
....................uuucguacguagcagaCca..........................................................................    3       1       7y1
....................uuucguacguagcagagGa..........................................................................    1       1       7y1
....................uuucguacguagUagagca..........................................................................    1       1       7y1
....................uuucgAacguagcagagca..........................................................................    2       1       7y1
....................uAucguacguagcagagca..........................................................................    5       1       7y1
....................uuucguacguagcaUagca..........................................................................    1       1       7y1
....................uuucguacguagcaUagcag.........................................................................    1       1       7y1
....................uuCcguacguagcagagcag.........................................................................    1       1       7y1
....................uuucguacgAagcagagcag.........................................................................    1       1       7y1
....................uuAcguacguagcagagcag.........................................................................    1       1       7y1
....................uAucguacguagcagagcag.........................................................................    4       1       7y1
....................uuucguacguagcagagAag.........................................................................    1       1       7y1
....................uuucguacguagcGgagcag.........................................................................    1       1       7y1
....................uuucguacguagcagaUcag.........................................................................    1       1       7y1
....................uuucguacguagcagagcaU.........................................................................    2       1       7y1
....................uuucguacguagcagagcaC.........................................................................    2       1       7y1
....................Auucguacguagcagagcag.........................................................................    11      1       7y1
....................uuucgCacguagcagagcag.........................................................................    1       1       7y1
....................uuucguacguagcagagcag.........................................................................    641     0       7y1
....................uuucguacguagcagagGagc........................................................................    2       1       7y1
....................uuucguacguagcGgagcagc........................................................................    1       1       7y1
....................uuucguacUuagcagagcagc........................................................................    1       1       7y1
....................uAucguacguagcagagcagc........................................................................    5       1       7y1
....................uuucguacguagUagagcagc........................................................................    1       1       7y1
....................uuucgAacguagcagagcagc........................................................................    3       1       7y1
....................uuucgGacguagcagagcagc........................................................................    3       1       7y1
....................Nuucguacguagcagagcagc........................................................................    1       1       7y1
....................uuucguacguagcagagcaCc........................................................................    1       1       7y1
....................uuucguGcguagcagagcagc........................................................................    1       1       7y1
....................uuucguaAguagcagagcagc........................................................................    5       1       7y1
....................uuucguaGguagcagagcagc........................................................................    1       1       7y1
....................uuucguacguagcagagcaAc........................................................................    1       1       7y1
....................uuucguaUguagcagagcagc........................................................................    1       1       7y1
....................uuucguacguagcagagAagc........................................................................    3       1       7y1
....................uuucguacguagcagagcaUc........................................................................    1       1       7y1
....................uuucguacguagAagagcagc........................................................................    2       1       7y1
....................uuucguacguagGagagcagc........................................................................    1       1       7y1
....................uuucguCcguagcagagcagc........................................................................    1       1       7y1
....................uuucguacguagcagagcagc........................................................................    2127    0       7y1
....................Cuucguacguagcagagcagc........................................................................    237     1       7y1
....................uuucguacguagcaUagcagc........................................................................    1       1       7y1
....................uuucguacguagcagagcagA........................................................................    1       1       7y1
....................uuuGguacguagcagagcagc........................................................................    2       1       7y1
....................uuucgCacguagcagagcagc........................................................................    1       1       7y1
....................Guucguacguagcagagcagc........................................................................    3       1       7y1
....................Auucguacguagcagagcagc........................................................................    18      1       7y1
```

**Star** **Mature**

gaccugcuucugggucgggguuucguacguagcagagcagcucccucgcugcgaucuauugaaagucagcccucgacacaaggguuuguccgcgcgcgcgcgcgcgcgcgugc

| | | | |
|---|---|---|---|
| ....................uuucguacguagcagCgcagc........................................................................ | 5 | 1 | 7y1 |
| ....................uuucguacgAagcagagcagc........................................................................ | 2 | 1 | 7y1 |
| ....................uuucguacguagcUgagcagcu....................................................................... | 1 | 1 | 7y1 |
| ....................uuucguacguagcagagcCgcu....................................................................... | 3 | 1 | 7y1 |
| ....................uuucguacAuagcagagcagcu....................................................................... | 1 | 1 | 7y1 |
| ....................uuucguacUuagcagagcagcu....................................................................... | 1 | 1 | 7y1 |
| ....................uuucguacguaCcagagcagcu....................................................................... | 1 | 1 | 7y1 |
| ....................uuuAguacguagcagagcagcu....................................................................... | 7 | 1 | 7y1 |
| ....................uuucguacguagcagGgcagcu....................................................................... | 1 | 1 | 7y1 |
| ....................Cuucguacguagcagagcagcu....................................................................... | 3 | 1 | 7y1 |
| ....................uuucguacguagAagagcagcu....................................................................... | 2 | 1 | 7y1 |
| ....................uuucguacguagcaUagcagcu....................................................................... | 272 | 1 | 7y1 |
| ....................uuucguacguagcagagcagcG....................................................................... | 22 | 1 | 7y1 |
| ....................Auucguacguagcagagcagcu....................................................................... | 37 | 1 | 7y1 |
| ....................uuucguacguagcagaCcagcu....................................................................... | 1 | 1 | 7y1 |
| ....................uuucgAacguagcagagcagcu....................................................................... | 4 | 1 | 7y1 |
| ....................uuucguacguagcagagcagAu....................................................................... | 1 | 1 | 7y1 |
| ....................Nuucguacguagcagagcagcu....................................................................... | 2 | 1 | 7y1 |
| ....................uuucguacguagcagagcagcA....................................................................... | 3 | 1 | 7y1 |
| ....................uuucguacCuagcagagcagcu....................................................................... | 4 | 1 | 7y1 |
| ....................uuucguacguagcagCgcagcu....................................................................... | 4 | 1 | 7y1 |
| ....................uuucguacguagcagagcagcu....................................................................... | 4088 | 0 | 7y1 |
| ....................uuucguacguagcagagcagcC....................................................................... | 1 | 1 | 7y1 |
| ....................uuucguacguagcagagGagcu....................................................................... | 2 | 1 | 7y1 |
| ....................uuucguacguagcagagcaUcu....................................................................... | 4 | 1 | 7y1 |
| ....................uAucguacguagcagagcagcu....................................................................... | 10 | 1 | 7y1 |
| ....................uuucguacguaAcagagcagcu....................................................................... | 1 | 1 | 7y1 |
| ....................uuucgCacguagcagagcagcu....................................................................... | 1 | 1 | 7y1 |
| ....................uuucguacguagcagagcaCcu....................................................................... | 1 | 1 | 7y1 |
| ....................uuucguGcguagcagagcagcu....................................................................... | 1 | 1 | 7y1 |
| ....................uuAcguacguagcagagcagcu....................................................................... | 3 | 1 | 7y1 |
| ....................uuucAuacguagcagagcagcu....................................................................... | 2 | 1 | 7y1 |
| ....................uuucguaUguagcagagcagcu....................................................................... | 1 | 1 | 7y1 |
| ....................uuucguacgAagcagagcagcu....................................................................... | 11 | 1 | 7y1 |
| ....................uuucguacguagcagagAagcu....................................................................... | 9 | 1 | 7y1 |
| ....................Guucguacguagcagagcagcu....................................................................... | 7 | 1 | 7y1 |
| ....................uuucguacguagUagagcagcu....................................................................... | 1 | 1 | 7y1 |
| ....................uuuUguacguagcagagcagcu....................................................................... | 2 | 1 | 7y1 |
| ....................uuucguacguagcagUgcagcu....................................................................... | 2 | 1 | 7y1 |
| ....................uuucguaAguagcagagcagcu....................................................................... | 8 | 1 | 7y1 |
| ....................uuucguacguagcaCagcagcu....................................................................... | 2 | 1 | 7y1 |
| ....................uuucCuacguagcagagcagcu....................................................................... | 2 | 1 | 7y1 |
| ....................uuucguacguagcaCagcagcuc...................................................................... | 1 | 1 | 7y1 |
| ....................uuucguacguagcagagcagGuc...................................................................... | 1 | 1 | 7y1 |
| ....................uuuAguacguagcagagcagcuc...................................................................... | 12 | 1 | 7y1 |
| ....................uuucCuacguagcagagcagcuc...................................................................... | 2 | 1 | 7y1 |
| ....................uuucguacCuagcagagcagcuc...................................................................... | 4 | 1 | 7y1 |
| ....................uuucguacguagcagagcagcGc...................................................................... | 1 | 1 | 7y1 |
| ....................uuucguacguagcagagcagcuc...................................................................... | 7935 | 0 | 7y1 |
| ....................uuucguacguagcagagcagcuG...................................................................... | 1 | 1 | 7y1 |
| ....................uuucguacguagcCgagcagcuc...................................................................... | 1 | 1 | 7y1 |
| ....................uuucAuacguagcagagcagcuc...................................................................... | 1 | 1 | 7y1 |
| ....................uuucguacguagcagagAagcuc...................................................................... | 10 | 1 | 7y1 |
| ....................uuucguacguagcagagcUgcuc...................................................................... | 1 | 1 | 7y1 |
| ....................uuucguacAuagcagagcagcuc...................................................................... | 1 | 1 | 7y1 |
| ....................uuucUuacguagcagagcagcuc...................................................................... | 6 | 1 | 7y1 |
| ....................Guucguacguagcagagcagcuc...................................................................... | 13 | 1 | 7y1 |
| ....................uuucguacguagcUgagcagcuc...................................................................... | 1 | 1 | 7y1 |
| ....................uuucguacguagcagUgcagcuc...................................................................... | 1 | 1 | 7y1 |
| ....................uuucguaAguagcagagcagcuc...................................................................... | 9 | 1 | 7y1 |
| ....................uuucguacguagcagagcagcAc...................................................................... | 2 | 1 | 7y1 |
| ....................uuCcguacguagcagagcagcuc...................................................................... | 1 | 1 | 7y1 |
| ....................uuucgAacguagcagagcagcuc...................................................................... | 16 | 1 | 7y1 |
| ....................uuucguacguagcagagcaUcuc...................................................................... | 7 | 1 | 7y1 |
| ....................uuucguacguagcagagcCgcuc...................................................................... | 3 | 1 | 7y1 |
| ....................uuucguacguGgcagagcagcuc...................................................................... | 2 | 1 | 7y1 |
| ....................uuucguacguagcGgagcagcuc...................................................................... | 1 | 1 | 7y1 |
| ....................uuucguacguagcagagcagcuA...................................................................... | 8 | 1 | 7y1 |
| ....................uuucguacguagcagagcagcuU...................................................................... | 1 | 1 | 7y1 |
| ....................uuucgCacguagcagagcagcuc...................................................................... | 1 | 1 | 7y1 |

**Star** **Mature**

```
gaccugcuucugggucgggguuucguacguagcagagcagcucccucgcugcgaucuauugaaagucagcccucgacacaaggguuuguccgcgcgcgcgcgcgcgcgugc
....................uuucguacguagcagaUcagcuc....................................................................    3     1    7y1
....................uuucguacgAagcagagcagcuc....................................................................    11    1    7y1
....................uuucguacUuagcagagcagcuc....................................................................    3     1    7y1
....................uuucguacguagcagaCcagcuc....................................................................    4     1    7y1
....................uuucguaUguagcagagcagcuc....................................................................    1     1    7y1
....................uuucguacguagcagCgcagcuc....................................................................    7     1    7y1
....................uuuGguacguagcagagcagcuc....................................................................    3     1    7y1
....................uuucguCcguagcagagcagcuc....................................................................    2     1    7y1
....................Auucguacguagcagagcagcuc....................................................................    89    1    7y1
....................uuucguacguagcagagcGgcuc....................................................................    2     1    7y1
....................uuucguacguagcagagcagAuc....................................................................    5     1    7y1
....................uuucguacguagcagagcaCcuc....................................................................    4     1    7y1
....................uuuUguacguagcagagcagcuc....................................................................    1     1    7y1
....................Nuucguacguagcagagcagcuc....................................................................    4     1    7y1
....................uuucguacguaCcagagcagcuc....................................................................    2     1    7y1
....................uAucguacguagcagagcagcuc....................................................................    11    1    7y1
....................uuucguacguaUcagagcagcuc....................................................................    1     1    7y1
....................uuucguacguagcagagGagcuc....................................................................    1     1    7y1
....................Cuucguacguagcagagcagcuc....................................................................    175   1    7y1
....................uuucguGcguagcagagcagcuc....................................................................    1     1    7y1
....................uuucguacgCagcagagcagcuc....................................................................    1     1    7y1
....................uuucguacguagUagagcagcuc....................................................................    2     1    7y1
....................uuucguacguagcaUagcagcuc....................................................................    2     1    7y1
....................uuAcguacguagcagagcagcuc....................................................................    3     1    7y1
....................uuucguacguagcagGgcagcuc....................................................................    1     1    7y1
....................uCucguacguagcagagcagcuc....................................................................    1     1    7y1
....................uuucguacguagAagagcagcuc....................................................................    9     1    7y1
....................uuGcguacguagcagagcagcuc....................................................................    1     1    7y1
....................uuucguacguagcagagcagcucG...................................................................    1     1    7y1
....................uuucguacgAagcagagcagcucc...................................................................    7     1    7y1
....................uuucguacguagcagGgcagcucc...................................................................    4     1    7y1
....................uuucguacgGagcagagcagcucc...................................................................    1     1    7y1
....................uuucguacguagcagagcagcuUc...................................................................    1     1    7y1
....................uuucguaAguagcagagcagcucc...................................................................    7     1    7y1
....................uuucguacguagcagagcagcAcc...................................................................    1     1    7y1
....................uuucguacguagcaUagcagcucc...................................................................    1     1    7y1
....................uuucguacCuagcagagcagcucc...................................................................    4     1    7y1
....................uuucguacguagcaCagcagcucc...................................................................    1     1    7y1
....................uuucUuacguagcagagcagcucc...................................................................    2     1    7y1
....................uuucgCacguagcagagcagcucc...................................................................    2     1    7y1
....................uuAcguacguagcagagcagcucc...................................................................    1     1    7y1
....................uuCcguacguagcagagcagcucc...................................................................    2     1    7y1
....................uuucguacguaUcagagcagcucc...................................................................    3     1    7y1
....................uuucguacguagcagagcCgcucc...................................................................    1     1    7y1
....................uuucguacguagcagagcaUcucc...................................................................    3     1    7y1
....................uuucCuacguagcagagcagcucc...................................................................    1     1    7y1
....................uuGcguacguagcagagcagcucc...................................................................    1     1    7y1
....................Guucguacguagcagagcagcucc...................................................................    14    1    7y1
....................uuucgAacguagcagagcagcucc...................................................................    11    1    7y1
....................uuucguacguagcagagcagAucc...................................................................    4     1    7y1
....................uuucguacguagcagagcaCcucc...................................................................    3     1    7y1
....................uuucguacguUgcagagcagcucc...................................................................    1     1    7y1
....................uuucguacguagcagagcagcucc...................................................................    5981  0    7y1
....................uAucguacguagcagagcagcucc...................................................................    9     1    7y1
....................uuucguacguagcagaCcagcucc...................................................................    3     1    7y1
....................uuucguacguagcagagcagcucU...................................................................    2     1    7y1
....................uuucguacgCagcagagcagcucc...................................................................    1     1    7y1
....................Auucguacguagcagagcagcucc...................................................................    67    1    7y1
....................uuucguacguagcagagcGgcucc...................................................................    1     1    7y1
....................uuucguacguagcagagcagcCcc...................................................................    2     1    7y1
....................Nuucguacguagcagagcagcucc...................................................................    3     1    7y1
....................uuucguacguagcGgagcagcucc...................................................................    1     1    7y1
....................uuucguacguagcagagAagcucc...................................................................    7     1    7y1
....................uuucguaGguagcagagcagcucc...................................................................    2     1    7y1
....................uuuGguacguagcagagcagcucc...................................................................    2     1    7y1
....................uuucguaUguagcagagcagcucc...................................................................    1     1    7y1
....................uuucguacUuagcagagcagcucc...................................................................    2     1    7y1
....................uuucguacguagcagagcagcuAc...................................................................    15    1    7y1
....................Cuucguacguagcagagcagcucc...................................................................    1     1    7y1
....................uuucguUcguagcagagcagcucc...................................................................    2     1    7y1
```

```
  Star                                                              Mature
gaccugcuucugggucgggguuucguacguagcagagcagcucccucgcugcgaucuauugaaagucagcccucgacacaaggguuuguccgcgcgcgcgcgcgcgcgcgugc
....................uuuAguacguagcagagcagcucc.....................................................................    6      1    7y1
....................uNucguacguagcagagcagcucc.....................................................................    1      1    7y1
....................uuucguacguagAagagcagcucc.....................................................................    9      1    7y1
....................uuucguacguagcagaUcagcucc.....................................................................    1      1    7y1
....................uuucguacguagcagCgcagcucc.....................................................................    5      1    7y1
....................uuuUguacguagcagagcagcucc.....................................................................    1      1    7y1
....................uuucguacguagcagagcaAcucc.....................................................................    1      1    7y1
....................uuucguacguagcagUgcagcucc.....................................................................    3      1    7y1
....................uuucguacguagcagagcagcucA.....................................................................    2      1    7y1
....................uuucguacguagcagagcagAuccc....................................................................    1      1    7y1
....................Auucguacguagcagagcagcuccc....................................................................    17     1    7y1
....................uuucUuacguagcagagcagcuccc....................................................................    1      1    7y1
....................uuucguacguagcagagcagGuccc....................................................................    1      1    7y1
....................uuucguacguagcagCgcagcuccc....................................................................    4      1    7y1
....................uuucguacCuagcagagcagcuccc....................................................................    1      1    7y1
....................uuucguacguagcagUgcagcuccc....................................................................    2      1    7y1
....................uuucguacguagcagagcaUcuccc....................................................................    3      1    7y1
....................uuucCuacguagcagagcagcuccc....................................................................    1      1    7y1
....................uAucguacguagcagagcagcuccc....................................................................    3      1    7y1
....................uuucguacguagcagagcagcuccA....................................................................    4      1    7y1
....................uuucguacguagUagagcagcuccc....................................................................    277    1    7y1
....................uuuGguacguagcagagcagcuccc....................................................................    3      1    7y1
....................uuucguacguagcagagcagcuccG....................................................................    1      1    7y1
....................uuucguacUuagcagagcagcuccc....................................................................    1      1    7y1
....................uuucgCacguagcagagcagcuccc....................................................................    1      1    7y1
....................uuucguacguagcagagcagcuAcc....................................................................    11     1    7y1
....................uuucguacguagcagagAagcuccc....................................................................    2      1    7y1
....................uuucguacguagcagagcaCcuccc....................................................................    2      1    7y1
....................Guucguacguagcagagcagcuccc....................................................................    2      1    7y1
....................uuucguaGguagcagagcagcuccc....................................................................    1      1    7y1
....................uuucguacguagAagagcagcuccc....................................................................    2      1    7y1
....................uuucguacguagcagagcagcuccc....................................................................    2072   0    7y1
....................uuucguacguagcagaCcagcuccc....................................................................    1      1    7y1
....................uuucguacgAagcagagcagcuccc....................................................................    3      1    7y1
....................uuucguacguagcagagcagcAccc....................................................................    1      1    7y1
....................uuucguacguagcGgagcagcuccc....................................................................    1      1    7y1
....................uuucguacguaUcagagcagcuccc....................................................................    1      1    7y1
....................uuucguaAguagcagagcagcuccc....................................................................    3      1    7y1
....................uuuAguacguagcagagcagcuccc....................................................................    2      1    7y1
....................uuucAuacguagcagagcagcuccc....................................................................    1      1    7y1
....................uuAcguacguagcagagcagcuccc....................................................................    2      1    7y1
....................uuucguacguagGagagcagcuccc....................................................................    1      1    7y1
....................uuucgAacguagcagagcagcuccc....................................................................    4      1    7y1
....................uuucguacguaCcagagcagcucccu...................................................................    1      1    7y1
....................uuucguacguagcagGgcagcucccu...................................................................    2      1    7y1
....................uuucguacguagcaUagcagcucccu...................................................................    1      1    7y1
....................uuucguacguagcagagcagAucccu...................................................................    2      1    7y1
....................uuucguacguagcagagcagcucccu...................................................................    4551   0    7y1
....................uuucguacgAagcagagcagcucccu...................................................................    7      1    7y1
....................uuAcguacguagcagagcagcucccu...................................................................    4      1    7y1
....................uuucguacguagcagagcagcAcccu...................................................................    1      1    7y1
....................uuucguGcguagcagagcagcucccu...................................................................    1      1    7y1
....................uuucguacguagcagagcagcCcccu...................................................................    1      1    7y1
....................uuucguacguagcagagcCgcucccu...................................................................    1      1    7y1
....................uuucguacAuagcagagcagcucccu...................................................................    3      1    7y1
....................uuucUuacguagcagagcagcucccu...................................................................    3      1    7y1
....................Guucguacguagcagagcagcucccu...................................................................    14     1    7y1
....................uuucguacguagcagaUcagcucccu...................................................................    1      1    7y1
....................uuucguacguagcaAagcagcucccu...................................................................    1      1    7y1
....................uuucguaUguagcagagcagcucccu...................................................................    2      1    7y1
....................uuucguacguagcagagcagcuccAu...................................................................    4      1    7y1
....................uAucguacguagcagagcagcucccu...................................................................    9      1    7y1
....................uuucguacguagcagagcagcucccG...................................................................    17     1    7y1
....................uuucgAacguagcagagcagcucccu...................................................................    9      1    7y1
....................uuucguacguagcagagcagcucccC...................................................................    152    1    7y1
....................uuucguacguagcagagcGgcucccu...................................................................    1      1    7y1
....................uuGcguacguagcagagcagcucccu...................................................................    1      1    7y1
....................uuucguacguagcagagcagcucccA...................................................................    8      1    7y1
....................uCucguacguagcagagcagcucccu...................................................................    1      1    7y1
....................uuucguacguagcUgagcagcucccu...................................................................    1      1    7y1
```

**Star** **Mature**

| | | | |
|---|---|---|---|
| gaccugcuucugggucggggguuucguacguagcagagcagcucccucgcugcgaucuauugaaagucagcccucgacacaaggguuuguccgcgcgcgcgcgcgcgcgcgcgugc | | | |
| ....................uuucguacguagcGgagcagcucccu................................................................... | 1 | 1 | 7y1 |
| ....................uuucguacguagcagagAagcucccu................................................................... | 6 | 1 | 7y1 |
| ....................uuucguaAguagcagagcagcucccu................................................................... | 4 | 1 | 7y1 |
| ....................uuucguacguagUagagcagcucccu................................................................... | 225 | 1 | 7y1 |
| ....................uuucguacguagcagagcagcucAcu................................................................... | 3 | 1 | 7y1 |
| ....................uuucguacguagAagagcagcucccu................................................................... | 2 | 1 | 7y1 |
| ....................Nuucguacguagcagagcagcucccu................................................................... | 1 | 1 | 7y1 |
| ....................uuucguacguagcagCgcagcucccu................................................................... | 10 | 1 | 7y1 |
| ....................Cuucguacguagcagagcagcucccu................................................................... | 2 | 1 | 7y1 |
| ....................uGucguacguagcagagcagcucccu................................................................... | 1 | 1 | 7y1 |
| ....................uuucguacUuagcagagcagcucccu................................................................... | 7 | 1 | 7y1 |
| ....................uuucguacguagcagaCcagcucccu................................................................... | 2 | 1 | 7y1 |
| ....................uuucguacguaAcagagcagcucccu................................................................... | 2 | 1 | 7y1 |
| ....................uuucCuacguagcagagcagcucccu................................................................... | 1 | 1 | 7y1 |
| ....................uuucguacCuagcagagcagcucccu................................................................... | 1 | 1 | 7y1 |
| ....................uuucguacguGgcagagcagcucccu................................................................... | 2 | 1 | 7y1 |
| ....................Auucguacguagcagagcagcucccu................................................................... | 42 | 1 | 7y1 |
| ....................uuucguacguaUcagagcagcucccu................................................................... | 1 | 1 | 7y1 |
| ....................uuucguacguagcagagcagcuAccu................................................................... | 10 | 1 | 7y1 |
| ....................uuuAguacguagcagagcagcucccu................................................................... | 6 | 1 | 7y1 |
| ....................Auucguacguagcagagcagcucccuc.................................................................. | 39 | 1 | 7y1 |
| ....................uuAcguacguagcagagcagcucccuc.................................................................. | 3 | 1 | 7y1 |
| ....................uuucCuacguagcagagcagcucccuc.................................................................. | 2 | 1 | 7y1 |
| ....................uuucgAacguagcagagcagcucccuc.................................................................. | 9 | 1 | 7y1 |
| ....................uuucguacgAagcagagcagcucccuc.................................................................. | 7 | 1 | 7y1 |
| ....................uuuGguacguagcagagcagcucccuc.................................................................. | 1 | 1 | 7y1 |
| ....................uuucguaAguagcagagcagcucccuc.................................................................. | 3 | 1 | 7y1 |
| ....................uuucguacguagcagagcagcucccuA.................................................................. | 3 | 1 | 7y1 |
| ....................uuucguacguagcagagAagcucccuc.................................................................. | 6 | 1 | 7y1 |
| ....................uuucguacguagcagagcaCcucccuc.................................................................. | 1 | 1 | 7y1 |
| ....................Cuucguacguagcagagcagcucccuc.................................................................. | 1 | 1 | 7y1 |
| ....................uuucguacguagcagagGagcucccuc.................................................................. | 1 | 1 | 7y1 |
| ....................uuucguacguagcagCgcagcucccuc.................................................................. | 10 | 1 | 7y1 |
| ....................uuucguacguagcagagcagcuAccuc.................................................................. | 7 | 1 | 7y1 |
| ....................uuucguacguagcagagcCgcucccuc.................................................................. | 1 | 1 | 7y1 |
| ....................Nuucguacguagcagagcagcucccuc.................................................................. | 6 | 1 | 7y1 |
| ....................uuucguacguagcagagcagcCcccuc.................................................................. | 1 | 1 | 7y1 |
| ....................uuucguacguaAcagagcagcucccuc.................................................................. | 1 | 1 | 7y1 |
| ....................uuucguacguagcagUgcagcucccuc.................................................................. | 2 | 1 | 7y1 |
| ....................uuucguacguagcagagcaUcucccuc.................................................................. | 2 | 1 | 7y1 |
| ....................uuucguacguagcagagcagcuccAuc.................................................................. | 5 | 1 | 7y1 |
| ....................uuucguacguagcagagcagcucUcuc.................................................................. | 1 | 1 | 7y1 |
| ....................uuucguacguagcaUagcagcucccuc.................................................................. | 2 | 1 | 7y1 |
| ....................uuucgCacguagcagagcagcucccuc.................................................................. | 1 | 1 | 7y1 |
| ....................uuucguacguagcaCagcagcucccuc.................................................................. | 2 | 1 | 7y1 |
| ....................uuucguacguagcagagcGgcucccuc.................................................................. | 2 | 1 | 7y1 |
| ....................uuucguaUguagcagagcagcucccuc.................................................................. | 1 | 1 | 7y1 |
| ....................uuucguacguagcagagcagcuGccuc.................................................................. | 1 | 1 | 7y1 |
| ....................uuucguacguagcagagcagcucccAc.................................................................. | 4 | 1 | 7y1 |
| ....................uuucguacguagcagagcagcucAcuc.................................................................. | 6 | 1 | 7y1 |
| ....................uuucguacguagcagagcagcAcccuc.................................................................. | 2 | 1 | 7y1 |
| ....................uuuAguacguagcagagcagcucccuc.................................................................. | 11 | 1 | 7y1 |
| ....................uuucguacguagcagagcagcuUccuc.................................................................. | 1 | 1 | 7y1 |
| ....................uuucguacguaUcagagcagcucccuc.................................................................. | 1 | 1 | 7y1 |
| ....................uuucguacguagcagagcagcucGcuc.................................................................. | 2 | 1 | 7y1 |
| ....................uuucguacCuagcagagcagcucccuc.................................................................. | 1 | 1 | 7y1 |
| ....................uuucguacguagcagagcagAucccuc.................................................................. | 4 | 1 | 7y1 |
| ....................uuucguacguagcagagcagcucccuc.................................................................. | 4379 | 0 | 7y1 |
| ....................uuucUuacguagcagagcagcucccuc.................................................................. | 2 | 1 | 7y1 |
| ....................uuucguacguagcagagcUgcucccuc.................................................................. | 1 | 1 | 7y1 |
| ....................uuucguacguagAagagcagcucccuc.................................................................. | 5 | 1 | 7y1 |
| ....................uAucguacguagcagagcagcucccuc.................................................................. | 9 | 1 | 7y1 |
| ....................uuucguacguagUagagcagcucccuc.................................................................. | 1 | 1 | 7y1 |
| ....................Guucguacguagcagagcagcucccuc.................................................................. | 12 | 1 | 7y1 |
| ....................uuucguacUuagcagagcagcucccuc.................................................................. | 2 | 1 | 7y1 |
| ....................uuucguacguagcagagAagcucccucgc................................................................ | 1 | 1 | 7y1 |
| ....................uuuGguacguagcagagcagcucccucgc................................................................ | 1 | 1 | 7y1 |
| ....................uuucguacguagcagagcagcucccAcgc................................................................ | 1 | 1 | 7y1 |
| ....................uuucguacguagcagagcUgcucccucgc................................................................ | 1 | 1 | 7y1 |
| ....................Guucguacguagcagagcagcucccucgc................................................................ | 2 | 1 | 7y1 |

**Star** **Mature**

```
gaccugcuucuggucggggauucguacguagcagagcagcucccucgcugcgaucuauugaaagucagcccucgacacaaggguuuguccgcgcgcgcgcgcgcgcgcgugc
...................uuucguacguagcagagcaCcucccucgc................................................................ 1     1  7y1
...................uuucguacguagcagagcagcucccucgc................................................................ 614   0  7y1
...................uuucguacgAagcagagcagcucccucgc................................................................ 1     1  7y1
...................uuAcguacguagcagagcagcucccucgc................................................................ 1     1  7y1
...................uAucguacguagcagagcagcucccucgc................................................................ 1     1  7y1
...................uuucguacguagcagCgcagcucccucgc................................................................ 1     1  7y1
...................uuucgAacguagcagagcagcucccucgc................................................................ 1     1  7y1
...................uuuAguacguagcagagcagcucccucgc................................................................ 1     1  7y1
...................uuucguacguagUagagcagcucccucgc................................................................ 1     1  7y1
...................uuucguacguagcagagcagcucAcucgc................................................................ 1     1  7y1
...................uuucguacCuagcagagcagcucccucgc................................................................ 1     1  7y1
...................uuucguacguagcagaUcagcucccucgc................................................................ 1     1  7y1
...................uuucguacguagcagagcagcuAccucgc................................................................ 1     1  7y1
...................Auucguacguagcagagcagcucccucgc................................................................ 4     1  7y1
...................uuucguacguagcagGgcagcucccucgcu............................................................... 2     1  7y1
...................uuucguacguGgcagagcagcucccucgcu............................................................... 1     1  7y1
...................uuucgAacguagcagagcagcucccucgcu............................................................... 4     1  7y1
...................uuucguacguagcagagcagcucccucgcu............................................................... 2277  0  7y1
...................uuuAguacguagcagagcagcucccucgcu............................................................... 4     1  7y1
...................uuucguacguagGagagcagcucccucgcu............................................................... 1     1  7y1
...................uuucguacguagcagagcagcucccuGgcu............................................................... 1     1  7y1
...................uuucguaAguagcagagcagcucccucgcu............................................................... 2     1  7y1
...................uuucguacguagcagaCcagcucccucgcu............................................................... 1     1  7y1
...................uuucguacguaCcagagcagcucccucgcu............................................................... 1     1  7y1
...................Guucguacguagcagagcagcucccucgcu............................................................... 178   1  7y1
...................uuucguacguagcagagcagcCcccucgcu............................................................... 1     1  7y1
...................uuucguacguagAagagcagcucccucgcu............................................................... 3     1  7y1
...................uuucguacCuagcagagcagcucccucgcu............................................................... 1     1  7y1
...................uuucguacguagcCgagcagcucccucgcu............................................................... 1     1  7y1
...................uAucguacguagcagagcagcucccucgcu............................................................... 1     1  7y1
...................uuucguacguagcagagcagcucGcucgcu............................................................... 1     1  7y1
...................uuucguacguagcagagcCgcucccucgcu............................................................... 2     1  7y1
...................Auucguacguagcagagcagcucccucgcu............................................................... 20    1  7y1
...................uuGcguacguagcagagcagcucccucgcu............................................................... 1     1  7y1
...................uuAcguacguagcagagcagcucccucgcu............................................................... 1     1  7y1
...................uuucguacguagcagCgcagcucccucgcu............................................................... 2     1  7y1
...................uuucguacgCagcagagcagcucccucgcu............................................................... 1     1  7y1
...................uuucguacgAagcagagcagcucccucgcu............................................................... 6     1  7y1
...................uuucguacguagcagagcUgcucccucgcu............................................................... 1     1  7y1
...................uuucguacguagcagagcagcucUcucgcu............................................................... 2     1  7y1
...................uuucguUcguagcagagcagcucccucgcu............................................................... 1     1  7y1
...................uuucguacguagcagagAagcucccucgcu............................................................... 2     1  7y1
...................uuucguacguagcaCagcagcucccucgcu............................................................... 2     1  7y1
...................uuucguacUuagcagagcagcucccucgcu............................................................... 1     1  7y1
...................uuucguacguagcagagcagUucccucgcu............................................................... 1     1  7y1
...................uuucguacguagcagagcGgcucccucgcu............................................................... 1     1  7y1
...................uuucguacguagcagagcagcucccucAcu............................................................... 2     1  7y1
...................uuucAuacguagcagagcagcucccucgcu............................................................... 2     1  7y1
...................uuucguacguagcagagcaUcucccucgcu............................................................... 1     1  7y1
...................uuucguacguagcagUgcagcucccucgcu............................................................... 1     1  7y1
...................uuucguacguagcagagcagcucccucgAu............................................................... 3     1  7y1
...................uuucguacguagcagagcagcucccucgcG............................................................... 20    1  7y1
...................uuucguacguagcagaUcagcucccucgcu............................................................... 1     1  7y1
...................Nuucguacguagcagagcagcucccucgcu............................................................... 2     1  7y1
...................uuucguacguagcagagcagcucccAcgcu............................................................... 2     1  7y1
...................uuucguacguagcagagcagcuAccucgcu............................................................... 7     1  7y1
...................uuucguacguagcagagcagcucccuAgcu............................................................... 3     1  7y1
...................uuucguacguagcagagcagcucAcucgcu............................................................... 1     1  7y1
...................uuucguacCuagcagagcagcucccucgcug.............................................................. 1     1  7y1
...................uuucAuacguagcagagcagcucccucgcug.............................................................. 1     1  7y1
...................uuucguacguagcUgagcagcucccucgcug.............................................................. 1     1  7y1
...................uuucguacguagcagagcGgcucccucgcug.............................................................. 1     1  7y1
...................uuucguacguagcagagcagcuAccucgcug.............................................................. 2     1  7y1
...................uuucguacguagcagagAagcucccucgcug.............................................................. 2     1  7y1
...................uuucguacguagcagagcagcucccucCcug.............................................................. 1     1  7y1
...................uAucguacguagcagagcagcucccucgcug.............................................................. 2     1  7y1
...................uuucguacguagcagagcagcucccucgcGg.............................................................. 9     1  7y1
...................Auucguacguagcagagcagcucccucgcug.............................................................. 6     1  7y1
...................uuucguCcguagcagagcagcucccucgcug.............................................................. 1     1  7y1
...................uuucguacguagcagagcagcucccucgGug.............................................................. 1     1  7y1
```

**Star** **Mature**

```
gaccugcuucugggucgggguuucguacguagcagagcagcucccucgcugcgaucuauugaaagucagcccucgacacaaggguuuguccgcgcgcgcgcgcgcgcgcgugc
....................uuucguacguagcagagcagcucccAcgcug..............................................................   1      1   7y1
....................uuucguUcguagcagagcagcucccucgcug..............................................................   1      1   7y1
....................uuucgAacguagcagagcagcucccucgcug..............................................................   2      1   7y1
....................uuucguacguagcagagcagcucccucgcug..............................................................   855    0   7y1
....................uuucguacguagAagagcagcucccucgcug..............................................................   1      1   7y1
....................uuucguacguagcagagcagcucccucUcug..............................................................   1      1   7y1
....................uuucguacgAagcagagcagcucccucgcug..............................................................   1      1   7y1
....................uuucguacguagcagagcaAcucccucgcug..............................................................   1      1   7y1
....................uuucguaAguagcagagcagcucccucgcug..............................................................   1      1   7y1
....................uuucUuacguagcagagcagcucccucgcug..............................................................   1      1   7y1
....................Auucguacguagcagagcagcucccucgcugc.............................................................   5      1   7y1
....................uuucguacgAagcagagcagcucccucgcugc.............................................................   1      1   7y1
....................uAucguacguagcagagcagcucccucgcugc.............................................................   1      1   7y1
....................uuuAguacguagcagagcagcucccucgcugc.............................................................   1      1   7y1
....................uuucguacguagcagagcagcuAccucgcugc.............................................................   1      1   7y1
....................uuucguacguagcagagcagcAcccucgcugc.............................................................   1      1   7y1
....................uuucguacCuagcagagcagcucccucgcugc.............................................................   1      1   7y1
....................Guucguacguagcagagcagcucccucgcugc.............................................................   1      1   7y1
....................Nuucguacguagcagagcagcucccucgcugc.............................................................   2      1   7y1
....................uuucguacguagcagagcagcucccucgcugc.............................................................   499    0   7y1
....................uuucguacguagcagagcagAucccucgcugc.............................................................   1      1   7y1
....................uuucguacguagcagagcagcucccucgAugc.............................................................   1      1   7y1
....................uuucguacguagcagagcagcucccucgcuCc.............................................................   1      1   7y1
.....................uucguacguagcaUagca..........................................................................   1      1   7y1
.....................uucguacguagcaCagca..........................................................................   1      1   7y1
.....................uucguacguagcagagAa..........................................................................   3      1   7y1
.....................uucguacguaCcagagca..........................................................................   1      1   7y1
.....................uucguacguaAcagagca..........................................................................   1      1   7y1
.....................uucguacguagcagGgca..........................................................................   1      1   7y1
.....................uucguacguagcagagGa..........................................................................   1      1   7y1
.....................uucguacguagAagagca..........................................................................   1      1   7y1
.....................uucguacgAagcagagca..........................................................................   3      1   7y1
.....................uucAuacguagcagagca..........................................................................   1      1   7y1
.....................uAcguacguagcagagca..........................................................................   4      1   7y1
.....................Nucguacguagcagagca..........................................................................   1      1   7y1
.....................uucguaAguagcagagca..........................................................................   8      1   7y1
.....................Gucguacguagcagagca..........................................................................   2      1   7y1
.....................uuAguacguagcagagca..........................................................................   4      1   7y1
.....................uucguacguagcagaAca..........................................................................   1      1   7y1
.....................uucguacguagcagagcG..........................................................................   1      1   7y1
.....................uNcguacguagcagagca..........................................................................   1      1   7y1
.....................Aucguacguagcagagca..........................................................................   12     1   7y1
.....................uucguacCuagcagagca..........................................................................   2      1   7y1
.....................uucguacguagcagagca..........................................................................   1212   0   7y1
.....................Cucguacguagcagagca..........................................................................   1      1   7y1
.....................uucguacguagcagagcC..........................................................................   1      1   7y1
.....................uucguacguagcagaUca..........................................................................   2      1   7y1
.....................uucguacAuagcagagcag.........................................................................   1      1   7y1
.....................uucguacguagcaCagcag.........................................................................   1      1   7y1
.....................Aucguacguagcagagcag.........................................................................   16     1   7y1
.....................uAcguacguagcagagcag.........................................................................   3      1   7y1
.....................Gucguacguagcagagcag.........................................................................   2      1   7y1
.....................uucguacguagcagagcag.........................................................................   1378   0   7y1
.....................uucguacguagAagagcag.........................................................................   1      1   7y1
.....................uucguaAguagcagagcag.........................................................................   4      1   7y1
.....................uucgCacguagcagagcag.........................................................................   1      1   7y1
.....................uucguacgAagcagagcag.........................................................................   2      1   7y1
.....................uucguacguagcagagAag.........................................................................   1      1   7y1
.....................uucguacguGgcagagcag.........................................................................   1      1   7y1
.....................uucguacguagcagagcaU.........................................................................   2      1   7y1
.....................uucguacCuagcagagcag.........................................................................   1      1   7y1
.....................uucguacguagcagagcUg.........................................................................   1      1   7y1
.....................uucguacguaUcagagcag.........................................................................   1      1   7y1
.....................uuAguacguagcagagcag.........................................................................   4      1   7y1
.....................uucguUcguagcagagcagc........................................................................   2      1   7y1
.....................uucguacgAagcagagcagc........................................................................   1      1   7y1
.....................uucguacAuagcagagcagc........................................................................   1      1   7y1
.....................Nucguacguagcagagcagc........................................................................   1      1   7y1
.....................uucguaGguagcagagcagc........................................................................   2      1   7y1
.....................uAcguacguagcagagcagc........................................................................   2      1   7y1
```

```
  Star                                        Mature
gaccugcuucugggucggggguuucguacguagcagagcagcucccucgcugcgaucuauugaaagucagcccucgacacaaggguuuguccgcgcgcgcgcgcgcgcgcgcgugc
.....................uucguacguagcaCagcagc..........................................................................    1     1    7y1
.....................uucCuacguagcagagcagc..........................................................................    1     1    7y1
.....................uucguacCuagcagagcagc..........................................................................    1     1    7y1
.....................uucguacguagcagagcaUc..........................................................................    3     1    7y1
.....................uucguacguagcagaUcagc..........................................................................    1     1    7y1
.....................uucguacguagcagaCcagc..........................................................................    1     1    7y1
.....................Aucguacguagcagagcagc..........................................................................    33    1    7y1
.....................uucguacguGgcagagcagc..........................................................................    2     1    7y1
.....................uucguacguagcagagcagA..........................................................................    2     1    7y1
.....................uuUguacguagcagagcagc..........................................................................    1     1    7y1
.....................uucguacguagcaUagcagc..........................................................................    1     1    7y1
.....................uucgAacguagcagagcagc..........................................................................    1     1    7y1
.....................uucguacguagcUgagcagc..........................................................................    1     1    7y1
.....................uucguaAguagcagagcagc..........................................................................    2     1    7y1
.....................uucguacUuagcagagcagc..........................................................................    2     1    7y1
.....................uucguacguaAcagagcagc..........................................................................    1     1    7y1
.....................uCcguacguagcagagcagc..........................................................................    1     1    7y1
.....................uucguacgCagcagagcagc..........................................................................    1     1    7y1
.....................uucguacguagcaAagcagc..........................................................................    1     1    7y1
.....................uucguacguagcagagcagc..........................................................................    1790  0    7y1
.....................uucguacguagcagGgcagc..........................................................................    1     1    7y1
.....................uuAguacguagcagagcagc..........................................................................    5     1    7y1
.....................uucguacguagcagagcGgcu.........................................................................    5     1    7y1
.....................uucguacguagcagagAagcu.........................................................................    9     1    7y1
.....................uucguacguagcaUagcagcu.........................................................................    5     1    7y1
.....................uucguacguagUagagcagcu.........................................................................    185   1    7y1
.....................uucguacguagcagagcagcA.........................................................................    4     1    7y1
.....................uucguacguagcagagcagAu.........................................................................    4     1    7y1
.....................uucguacguagcGgagcagcu.........................................................................    1     1    7y1
.....................Nucguacguagcagagcagcu.........................................................................    2     1    7y1
.....................uucguGcguagcagagcagcu.........................................................................    1     1    7y1
.....................uucguacguaUcagagcagcu.........................................................................    3     1    7y1
.....................uucguacguagcUgagcagcu.........................................................................    2     1    7y1
.....................uucguacguagcagagcUgcu.........................................................................    1     1    7y1
.....................uucCuacguagcagagcagcu.........................................................................    1     1    7y1
.....................uucguacgAagcagagcagcu.........................................................................    8     1    7y1
.....................uucUuacguagcagagcagcu.........................................................................    5     1    7y1
.....................uucguacguagcagagcaUcu.........................................................................    3     1    7y1
.....................uucguacguagcagagcaAcu.........................................................................    1     1    7y1
.....................uucguacguagcagaCcagcu.........................................................................    5     1    7y1
.....................uuGguacguagcagagcagcu.........................................................................    2     1    7y1
.....................uucguacguagcagagcagGu.........................................................................    2     1    7y1
.....................uucguacguagcagGgcagcu.........................................................................    1     1    7y1
.....................uucguacguagcagagcaCcu.........................................................................    3     1    7y1
.....................uucgGacguagcagagcagcu.........................................................................    1     1    7y1
.....................uucguacguagcagagcagcu.........................................................................    7198  0    7y1
.....................Aucguacguagcagagcagcu.........................................................................    80    1    7y1
.....................uucguacguagcagagcagcG.........................................................................    40    1    7y1
.....................uucgCacguagcagagcagcu.........................................................................    3     1    7y1
.....................uucguacguagAagagcagcu.........................................................................    3     1    7y1
.....................uucguacguagcagCgcagcu.........................................................................    7     1    7y1
.....................uucguacguGgcagagcagcu.........................................................................    2     1    7y1
.....................uucgAacguagcagagcagcu.........................................................................    9     1    7y1
.....................uucguacguagcagagcagcC.........................................................................    1     1    7y1
.....................uucguacguUgcagagcagcu.........................................................................    1     1    7y1
.....................uucguaUguagcagagcagcu.........................................................................    3     1    7y1
.....................Gucguacguagcagagcagcu.........................................................................    8     1    7y1
.....................uucguacCuagcagagcagcu.........................................................................    4     1    7y1
.....................uucguacguagcaAagcagcu.........................................................................    1     1    7y1
.....................uuAguacguagcagagcagcu.........................................................................    24    1    7y1
.....................uucguUcguagcagagcagcu.........................................................................    1     1    7y1
.....................uucguacguaCcagagcagcu.........................................................................    3     1    7y1
.....................uAcguacguagcagagcagcu.........................................................................    18    1    7y1
.....................uucguacguagcagagUagcu.........................................................................    1     1    7y1
.....................uucguacguagcagagGagcu.........................................................................    4     1    7y1
.....................uucguaGguagcagagcagcu.........................................................................    1     1    7y1
.....................uucguacguagcaCagcagcu.........................................................................    1     1    7y1
.....................uucguacAuagcagagcagcu.........................................................................    1     1    7y1
.....................uucguaAguagcagagcagcu.........................................................................    22    1    7y1
.....................uucguacguagcagagcCgcu.........................................................................    1     1    7y1
```

**Star** **Mature**

```
gaccugcuucugggucggggguucguacguagcagagcagcucccucgcugcgaucuauugaaagucagcccucgacacaaggguuuguccgcgcgcgcgcgcgcgcgcgugc
.....................uucguacguagGagagcagcu.........................................................................   1       1   7y1
.....................uucguacUuagcagagcagcu.........................................................................   2       1   7y1
.....................Cucguacguagcagagcagcu.........................................................................   2       1   7y1
.....................uucguacguagcagagcagcuU........................................................................   1       1   7y1
.....................uucguaUguagcagagcagcuc........................................................................   3       1   7y1
.....................uucguacguagcagagcagAuc........................................................................   9       1   7y1
.....................uucguacguagcagagcagcAc........................................................................   4       1   7y1
.....................uucguacguagcagaUcagcuc........................................................................   6       1   7y1
.....................uucAuacguagcagagcagcuc........................................................................   6       1   7y1
.....................uucguacguaUcagagcagcuc........................................................................   11      1   7y1
.....................uucguacgCagcagagcagcuc........................................................................   1       1   7y1
.....................uucguaGguagcagagcagcuc........................................................................   1       1   7y1
.....................uucguacguagcagagcUgcuc........................................................................   1       1   7y1
.....................uucguacguagcagagcGgcuc........................................................................   3       1   7y1
.....................uAcguacguagcagagcagcuc........................................................................   27      1   7y1
.....................uucgCacguagcagagcagcuc........................................................................   2       1   7y1
.....................uucguacguagcagagcCgcuc........................................................................   1       1   7y1
.....................uucCuacguagcagagcagcuc........................................................................   2       1   7y1
.....................uucguacguagUagagcagcuc........................................................................   1       1   7y1
.....................uucguacguagcagagcaCcuc........................................................................   5       1   7y1
.....................uucguacguagcagagAagcuc........................................................................   19      1   7y1
.....................uucguacUuagcagagcagcuc........................................................................   5       1   7y1
.....................uucgAacguagcagagcagcuc........................................................................   8       1   7y1
.....................uucguacguagcaUagcagcuc........................................................................   5       1   7y1
.....................uuUguacguagcagagcagcuc........................................................................   2       1   7y1
.....................Aucguacguagcagagcagcuc........................................................................   123     1   7y1
.....................uucguacguagGagagcagcuc........................................................................   3       1   7y1
.....................uucguacguagcagagcagcuc........................................................................   12187   0   7y1
.....................uucguGcguagcagagcagcuc........................................................................   1       1   7y1
.....................uucguacguagAagagcagcuc........................................................................   13      1   7y1
.....................uucguacguaCcagagcagcuc........................................................................   9       1   7y1
.....................uucguacguagcaCagcagcuc........................................................................   5       1   7y1
.....................uucguacCuagcagagcagcuc........................................................................   7       1   7y1
.....................uCcguacguagcagagcagcuc........................................................................   200     1   7y1
.....................uucguacguagcagUgcagcuc........................................................................   2       1   7y1
.....................uucguacguagcagagUagcuc........................................................................   1       1   7y1
.....................uucguacAuagcagagcagcuc........................................................................   3       1   7y1
.....................uucguacguagcGgagcagcuc........................................................................   1       1   7y1
.....................uucguacguagcagagcaUcuc........................................................................   8       1   7y1
.....................uucguacguagcagagcagcuA........................................................................   8       1   7y1
.....................uucgGacguagcagagcagcuc........................................................................   1       1   7y1
.....................Nucguacguagcagagcagcuc........................................................................   5       1   7y1
.....................uucguacguagcagagcagcGc........................................................................   1       1   7y1
.....................uucguacguagcagagcagcCc........................................................................   1       1   7y1
.....................uucguUcguagcagagcagcuc........................................................................   2       1   7y1
.....................uuAguacguagcagagcagcuc........................................................................   33      1   7y1
.....................uucguacgAagcagagcagcuc........................................................................   15      1   7y1
.....................uucguacguagcagagGagcuc........................................................................   2       1   7y1
.....................uucguacguaAcagagcagcuc........................................................................   2       1   7y1
.....................uucguacguagcagagcagcuG........................................................................   1       1   7y1
.....................Gucguacguagcagagcagcuc........................................................................   29      1   7y1
.....................uucguacguGgcagagcagcuc........................................................................   2       1   7y1
.....................uucguacguagcagagcagUuc........................................................................   1       1   7y1
.....................uucguacguagcagagcagGuc........................................................................   1       1   7y1
.....................uucguacguagcagaAcagcuc........................................................................   1       1   7y1
.....................uucguCcguagcagagcagcuc........................................................................   1       1   7y1
.....................uucguacguagcagaCcagcuc........................................................................   8       1   7y1
.....................uucguacguagcagCgcagcuc........................................................................   16      1   7y1
.....................uucguaAguagcagagcagcuc........................................................................   55      1   7y1
.....................uucguacguagcagCgcagcucc.......................................................................   4       1   7y1
.....................uucguacguagcagagcagAucc.......................................................................   5       1   7y1
.....................uucguacguagcaUagcagcucc.......................................................................   4       1   7y1
.....................uucguacAuagcagagcagcucc.......................................................................   1       1   7y1
.....................uucguacguagcagaCcagcucc.......................................................................   2       1   7y1
.....................uucguacguagcagagcagcCcc.......................................................................   1       1   7y1
.....................uucguacguagcGgagcagcucc.......................................................................   5       1   7y1
.....................uucUuacguagcagagcagcucc.......................................................................   3       1   7y1
.....................uucguacguagcagagcagGucc.......................................................................   1       1   7y1
.....................Gucguacguagcagagcagcucc.......................................................................   17      1   7y1
.....................uucAuacguagcagagcagcucc.......................................................................   5       1   7y1
```

### Star

### Mature

| | | | |
|---|---|---|---|
| gaccugcuucugggucggggu uucguacguagcagagcagcucccucgcugcgaucuauugaaagucagcc cucgacacaaggguuugu ccgcgcgcgcgcgcgcgcgcgcgugc | | | |
| .....................uucguacguagcagagcagcucA....................................................................... | 13 | 1 | 7y1 |
| .....................uucguacguagcUgagcagcucc....................................................................... | 2 | 1 | 7y1 |
| .....................uucguacgGagcagagcagcucc....................................................................... | 1 | 1 | 7y1 |
| .....................uucguacguagcagagcagcuAc....................................................................... | 9 | 1 | 7y1 |
| .....................uucguacguagcagagcagcGcc....................................................................... | 1 | 1 | 7y1 |
| .....................uucguacguaCcagagcagcucc....................................................................... | 3 | 1 | 7y1 |
| .....................uuUguacguagcagagcagcucc....................................................................... | 1 | 1 | 7y1 |
| .....................uucguacguagcagagCCgcucc....................................................................... | 1 | 1 | 7y1 |
| .....................uucguacguagcagagGagcucc....................................................................... | 2 | 1 | 7y1 |
| .....................uucguacguagcagagUagcucc....................................................................... | 1 | 1 | 7y1 |
| .....................Nucguacguagcagagcagcucc....................................................................... | 4 | 1 | 7y1 |
| .....................uucguacCuagcagagcagcucc....................................................................... | 6 | 1 | 7y1 |
| .....................uucguaGguagcagagcagcucc....................................................................... | 1 | 1 | 7y1 |
| .....................uNcguacguagcagagcagcucc....................................................................... | 1 | 1 | 7y1 |
| .....................uucguaAguagcagagcagcucc....................................................................... | 26 | 1 | 7y1 |
| .....................uucguacguagcagagAagcucc....................................................................... | 8 | 1 | 7y1 |
| .....................uucgAacguagcagagcagcucc....................................................................... | 1 | 1 | 7y1 |
| .....................uAcguacguagcagagcagcucc....................................................................... | 17 | 1 | 7y1 |
| .....................uuAguacguagcagagcagcucc....................................................................... | 16 | 1 | 7y1 |
| .....................uucguUcguagcagagcagcucc....................................................................... | 1 | 1 | 7y1 |
| .....................uuGguacguagcagagcagcucc....................................................................... | 2 | 1 | 7y1 |
| .....................uucguacgAagcagagcagcucc....................................................................... | 6 | 1 | 7y1 |
| .....................Aucguacguagcagagcagcucc....................................................................... | 71 | 1 | 7y1 |
| .....................uucguacUuagcagagcagcucc....................................................................... | 2 | 1 | 7y1 |
| .....................uucguacguagcagagcagcucc....................................................................... | 7274 | 0 | 7y1 |
| .....................uucguacguagAagagcagcucc....................................................................... | 11 | 1 | 7y1 |
| .....................uucguacguaUcagagcagcucc....................................................................... | 8 | 1 | 7y1 |
| .....................uucguacguagcagagcagcAcc....................................................................... | 1 | 1 | 7y1 |
| .....................uucguacguGgcagagcagcucc....................................................................... | 2 | 1 | 7y1 |
| .....................uucguCcguagcagagcagcucc....................................................................... | 1 | 1 | 7y1 |
| .....................uucguacguagcagagcaAcucc....................................................................... | 1 | 1 | 7y1 |
| .....................uucguacguagcagagcagcucG....................................................................... | 1 | 1 | 7y1 |
| .....................uucgCacguagcagagcagcucc....................................................................... | 3 | 1 | 7y1 |
| .....................uucguacguagcagagcaUcucc....................................................................... | 3 | 1 | 7y1 |
| .....................uucguacguagcaCagcagcucc....................................................................... | 1 | 1 | 7y1 |
| .....................uucguacguagGagagcagcucc....................................................................... | 1 | 1 | 7y1 |
| .....................uucguacguagcagaUcagcucc....................................................................... | 5 | 1 | 7y1 |
| .....................uucguacguagUagagcagcucc....................................................................... | 1 | 1 | 7y1 |
| .....................uucguacguagcagagcaCcucc....................................................................... | 1 | 1 | 7y1 |
| .....................uucguacguGgcagagcagcuccc...................................................................... | 1 | 1 | 7y1 |
| .....................uucguacguaCcagagcagcuccc...................................................................... | 2 | 1 | 7y1 |
| .....................uucguacguagcagagcCgcuccc...................................................................... | 2 | 1 | 7y1 |
| .....................uucguacUuagcagagcagcuccc...................................................................... | 1 | 1 | 7y1 |
| .....................uucguacguaUcagagcagcuccc...................................................................... | 1 | 1 | 7y1 |
| .....................uucguacguagcagagcagcuAcc...................................................................... | 7 | 1 | 7y1 |
| .....................uucguacguagcagagcagcuccA...................................................................... | 2 | 1 | 7y1 |
| .....................uAcguacguagcagagcagcuccc...................................................................... | 6 | 1 | 7y1 |
| .....................Nucguacguagcagagcagcuccc...................................................................... | 1 | 1 | 7y1 |
| .....................uucguacguagcagagcagcuccc...................................................................... | 3826 | 0 | 7y1 |
| .....................uucguacCuagcagagcagcuccc...................................................................... | 1 | 1 | 7y1 |
| .....................Aucguacguagcagagcagcuccc...................................................................... | 31 | 1 | 7y1 |
| .....................uucguacguagcagagcagcuUcc...................................................................... | 1 | 1 | 7y1 |
| .....................uucguacguagcagagcNgcuccc...................................................................... | 1 | 1 | 7y1 |
| .....................uucguCcguagcagagcagcuccc...................................................................... | 1 | 1 | 7y1 |
| .....................uucguacguagcagagcagcCccc...................................................................... | 1 | 1 | 7y1 |
| .....................uuAguacguagcagagcagcuccc...................................................................... | 10 | 1 | 7y1 |
| .....................uucguacguagcagagcagcucAc...................................................................... | 16 | 1 | 7y1 |
| .....................uucguacguagcGgagcagcuccc...................................................................... | 1 | 1 | 7y1 |
| .....................uucguacguagcagagcagcuGcc...................................................................... | 1 | 1 | 7y1 |
| .....................uucAuacguagcagagcagcuccc...................................................................... | 1 | 1 | 7y1 |
| .....................uucguacgAagcagagcagcuccc...................................................................... | 2 | 1 | 7y1 |
| .....................uucUuacguagcagagcagcuccc...................................................................... | 1 | 1 | 7y1 |
| .....................uucguacguagcagagAagcuccc...................................................................... | 4 | 1 | 7y1 |
| .....................uucgAacguagcagagcagcuccc...................................................................... | 4 | 1 | 7y1 |
| .....................Gucguacguagcagagcagcuccc...................................................................... | 13 | 1 | 7y1 |
| .....................uucguaAguagcagagcagcuccc...................................................................... | 11 | 1 | 7y1 |
| .....................uucguacguagAagagcagcuccc...................................................................... | 1 | 1 | 7y1 |
| .....................uucguacguagcaCagcagcuccc...................................................................... | 1 | 1 | 7y1 |
| .....................uucguacguagcagaCcagcuccc...................................................................... | 1 | 1 | 7y1 |
| .....................uucguacguagcagagcaCcuccc...................................................................... | 4 | 1 | 7y1 |

**Star** **Mature**

| | | | |
|---|---|---|---|
| gaccugcuucugggucgggguuucguacguagcagagcagcucccucgcugcgaucuauugaaagucagcccucgacacaaggguuuguccgcgcgcgcgcgcgcgcgcgugc | | | |
| .....................uucguacguagcaUagcagcuccc.................................................................... | 6 | 1 | 7y1 |
| .....................uucguacguagcagagUagcuccc.................................................................... | 1 | 1 | 7y1 |
| .....................uucguacguagcagagcaUcuccc.................................................................... | 5 | 1 | 7y1 |
| .....................uucguacguagcagagcaAcuccc.................................................................... | 1 | 1 | 7y1 |
| .....................uucguacguagcagaUcagcuccc.................................................................... | 3 | 1 | 7y1 |
| .....................uucguacguagcagagcagcAccc.................................................................... | 2 | 1 | 7y1 |
| .....................uucguacguagcagagcagcucGc.................................................................... | 1 | 1 | 7y1 |
| .....................uucCuacguagcagagcagcuccc.................................................................... | 1 | 1 | 7y1 |
| .....................uucguacguagcagagcagcuUccu................................................................... | 1 | 1 | 7y1 |
| .....................uucUuacguagcagagcagcucccu................................................................... | 4 | 1 | 7y1 |
| .....................uucguacguagcagCgcagcucccu................................................................... | 4 | 1 | 7y1 |
| .....................uucguacguagcagagcagcGcccu................................................................... | 2 | 1 | 7y1 |
| .....................uucguacgCagcagagcagcucccu................................................................... | 2 | 1 | 7y1 |
| .....................uucguacguagcUgagcagcucccu................................................................... | 1 | 1 | 7y1 |
| .....................uucguacguagcagagcUgcucccu................................................................... | 6 | 1 | 7y1 |
| .....................uucguacguagcagagcagcucUcu................................................................... | 5 | 1 | 7y1 |
| .....................Gucguacguagcagagcagcucccu................................................................... | 24 | 1 | 7y1 |
| .....................uucguacguaAcagagcagcucccu................................................................... | 1 | 1 | 7y1 |
| .....................uucguacguagcagGgcagcucccu................................................................... | 1 | 1 | 7y1 |
| .....................Cucguacguagcagagcagcucccu................................................................... | 1 | 1 | 7y1 |
| .....................uucguacguagcagagcagcAcccu................................................................... | 6 | 1 | 7y1 |
| .....................uucguacguagcagagcagcuAccu................................................................... | 23 | 1 | 7y1 |
| .....................uucguacguagcagagAagcucccu................................................................... | 11 | 1 | 7y1 |
| .....................uucguacguagcagagcagcucAcu................................................................... | 24 | 1 | 7y1 |
| .....................uucguacguagcGgagcagcucccu................................................................... | 2 | 1 | 7y1 |
| .....................uucguacguagcagagcagcucccA................................................................... | 13 | 1 | 7y1 |
| .....................uNcguacguagcagagcagcucccu................................................................... | 1 | 1 | 7y1 |
| .....................uucguacguagcagagcCgcucccu................................................................... | 2 | 1 | 7y1 |
| .....................uucguacguagcagaCcagcucccu................................................................... | 3 | 1 | 7y1 |
| .....................uucguUcguagcagagcagcucccu................................................................... | 1 | 1 | 7y1 |
| .....................Nucguacguagcagagcagcucccu................................................................... | 8 | 1 | 7y1 |
| .....................uucguacgGagcagagcagcucccu................................................................... | 1 | 1 | 7y1 |
| .....................uucguacguagAagagcagcucccu................................................................... | 7 | 1 | 7y1 |
| .....................uucguacguagcagagcagcuccGu................................................................... | 2 | 1 | 7y1 |
| .....................uucguacgAagcagagcagcucccu................................................................... | 9 | 1 | 7y1 |
| .....................uuGguacguagcagagcagcucccu................................................................... | 2 | 1 | 7y1 |
| .....................uucCuacguagcagagcagcucccu................................................................... | 1 | 1 | 7y1 |
| .....................uucguaUguagcagagcagcucccu................................................................... | 1 | 1 | 7y1 |
| .....................uucguacguagcagUgcagcucccu................................................................... | 1 | 1 | 7y1 |
| .....................uucgCacguagcagagcagcucccu................................................................... | 2 | 1 | 7y1 |
| .....................uucguacguagcCgagcagcucccu................................................................... | 2 | 1 | 7y1 |
| .....................uucguacguagcagagcagAucccu................................................................... | 5 | 1 | 7y1 |
| .....................uucguacguagcagagcaAcucccu................................................................... | 1 | 1 | 7y1 |
| .....................uucguacguagUagagcagcucccu................................................................... | 1 | 1 | 7y1 |
| .....................uucguacguagcaUagcagcucccu................................................................... | 5 | 1 | 7y1 |
| .....................Aucguacguagcagagcagcucccu................................................................... | 78 | 1 | 7y1 |
| .....................uucguacguagcaCagcagcucccu................................................................... | 3 | 1 | 7y1 |
| .....................uucguacguagcagagcaCcucccu................................................................... | 1 | 1 | 7y1 |
| .....................uucguacguagcagagGagcucccu................................................................... | 2 | 1 | 7y1 |
| .....................uucguacguagcagagUagcucccu................................................................... | 1 | 1 | 7y1 |
| .....................uucguacguGgcagagcagcucccu................................................................... | 1 | 1 | 7y1 |
| .....................uucgAacguagcagagcagcucccu................................................................... | 5 | 1 | 7y1 |
| .....................uucguaGguagcagagcagcucccu................................................................... | 1 | 1 | 7y1 |
| .....................uucguacguagcagagcaUcucccu................................................................... | 5 | 1 | 7y1 |
| .....................uucguacguagcagagcagcCcccu................................................................... | 2 | 1 | 7y1 |
| .....................uucguacguagcagagcagcuGccu................................................................... | 1 | 1 | 7y1 |
| .....................uucguacguagcagagcagcucccG................................................................... | 35 | 1 | 7y1 |
| .....................uucguacguaUcagagcagcucccu................................................................... | 5 | 1 | 7y1 |
| .....................uucguacguaCcagagcagcucccu................................................................... | 3 | 1 | 7y1 |
| .....................uucguacguagcagagcGgcucccu................................................................... | 8 | 1 | 7y1 |
| .....................uucguacguagcagagcagcucccC................................................................... | 1 | 1 | 7y1 |
| .....................uucguacUuagcagagcagcucccu................................................................... | 4 | 1 | 7y1 |
| .....................uucguacguagcagagcagcuccAu................................................................... | 6 | 1 | 7y1 |
| .....................uucguacCuagcagagcagcucccu................................................................... | 5 | 1 | 7y1 |
| .....................uucguacguagGagagcagcucccu................................................................... | 1 | 1 | 7y1 |
| .....................uucguGcguagcagagcagcucccu................................................................... | 1 | 1 | 7y1 |
| .....................uucguaAguagcagagcagcucccu................................................................... | 40 | 1 | 7y1 |
| .....................uucguacAuagcagagcagcucccu................................................................... | 3 | 1 | 7y1 |
| .....................uuAguacguagcagagcagcucccu................................................................... | 23 | 1 | 7y1 |
| .....................uAcguacguagcagagcagcucccu................................................................... | 21 | 1 | 7y1 |

**Star** **Mature**

```
gaccugcuucugggucgggguuucguacguagcagagcagcucccucgcugcgaucuauugaaagucagcccucgacacaaggguuuguccgcgcgcgcgcgcgcgcgcgcgugc
.....................uucguacguagcagaUcagcucccu.....................................................................   2     1   7y1
.....................uucguacguagcaAagcagcucccu.....................................................................   2     1   7y1
.....................uCcguacguagcagagcagcucccu.....................................................................   2     1   7y1
.....................uucguacguagcagagcagcucccu.....................................................................   9826  0   7y1
.....................uucAuacguagcagagcagcucccu.....................................................................   1     1   7y1
.....................uucguacguagcaUagcagcucccuc....................................................................   4     1   7y1
.....................uucguacguagcagagcagcuAccuc....................................................................   12    1   7y1
.....................uucguacguagcGgagcagcucccuc....................................................................   1     1   7y1
.....................uucguacguagcagagcaUcucccuc....................................................................   1     1   7y1
.....................uucCuacguagcagagcagcucccuc....................................................................   2     1   7y1
.....................uucguacguagcagagcagcuccAuc....................................................................   6     1   7y1
.....................uucguacguaUcagagcagcucccuc....................................................................   3     1   7y1
.....................uucguacguagcagGgcagcucccuc....................................................................   1     1   7y1
.....................uucguacguaCcagagcagcucccuc....................................................................   1     1   7y1
.....................uucguacAuagcagagcagcucccuc....................................................................   1     1   7y1
.....................uucguaAguagcagagcagcucccuc....................................................................   33    1   7y1
.....................uucguacguaAcagagcagcucccuc....................................................................   1     1   7y1
.....................uucguacgCagcagagcagcucccuc....................................................................   1     1   7y1
.....................Nucguacguagcagagcagcucccuc....................................................................   5     1   7y1
.....................uucguacguagcagagcagcGcccuc....................................................................   1     1   7y1
.....................uucguacguagcagagcaCcucccuc....................................................................   1     1   7y1
.....................uucguacguagcagagcagcucccuA....................................................................   1     1   7y1
.....................uucguacguagcagagcagcucccAc....................................................................   1     1   7y1
.....................uucguacguagcagagUagcucccuc....................................................................   1     1   7y1
.....................uucguacguagcagagcagcucccuc....................................................................   7376  0   7y1
.....................uucguacguagcagaUcagcucccuc....................................................................   4     1   7y1
.....................uucguacUuagcagagcagcucccuc....................................................................   2     1   7y1
.....................uucguacgAagcagagcagcucccuc....................................................................   6     1   7y1
.....................Gucguacguagcagagcagcucccuc....................................................................   10    1   7y1
.....................uucguacguagcagagcagcuUccuc....................................................................   1     1   7y1
.....................uucguacguagcagagcagcucccCc....................................................................   2     1   7y1
.....................uucguacguagcagUgcagcucccuc....................................................................   2     1   7y1
.....................uucguacguagcagagcUgcucccuc....................................................................   1     1   7y1
.....................uucguacguagcagagcCgcucccuc....................................................................   3     1   7y1
.....................uucguacguagcagagcagcAcccuc....................................................................   5     1   7y1
.....................uucguacguagcagagcaAcucccuc....................................................................   2     1   7y1
.....................uucguGcguagcagagcagcucccuc....................................................................   1     1   7y1
.....................uucguUcguagcagagcagcucccuc....................................................................   1     1   7y1
.....................uucguacCuagcagagcagcucccuc....................................................................   4     1   7y1
.....................uucguacguagcagagcagcucccuG....................................................................   1     1   7y1
.....................uucgAacguagcagagcagcucccuc....................................................................   4     1   7y1
.....................uucguacguagcUgagcagcucccuc....................................................................   2     1   7y1
.....................Cucguacguagcagagcagcucccuc....................................................................   1     1   7y1
.....................uucguacguagAagagcagcucccuc....................................................................   5     1   7y1
.....................uucguacguagcagCgcagcucccuc....................................................................   8     1   7y1
.....................Aucguacguagcagagcagcucccuc....................................................................   59    1   7y1
.....................uuGguacguagcagagcagcucccuc....................................................................   1     1   7y1
.....................uucguacguagcagagAagcucccuc....................................................................   11    1   7y1
.....................uAcguacguagcagagcagcucccuc....................................................................   11    1   7y1
.....................uuUguacguagcagagcagcucccuc....................................................................   1     1   7y1
.....................uucguacguagcagagcagAucccuc....................................................................   5     1   7y1
.....................uucUuacguagcagagcagcucccuc....................................................................   2     1   7y1
.....................uuAguacguagcagagcagcucccuc....................................................................   12    1   7y1
.....................uCcguacguagcagagcagcucccuc....................................................................   1     1   7y1
.....................uucgCacguagcagagcagcucccuc....................................................................   253   1   7y1
.....................uucguacguagcaAagcagcucccuc....................................................................   1     1   7y1
.....................uucguacguagcagaCcagcucccuc....................................................................   2     1   7y1
.....................uucguacguagcagagcagcucccGc....................................................................   1     1   7y1
.....................uucgGacguagcagagcagcucccuc....................................................................   1     1   7y1
.....................uucguacguagcagagcGgcucccuc....................................................................   2     1   7y1
.....................uucguacguagcagagcagcuGccuc....................................................................   4     1   7y1
.....................uucguacguagcagagcagcucAcuc....................................................................   19    1   7y1
.....................uucCuacguagcagagcagcucccucg...................................................................   1     1   7y1
.....................uucguCcguagcagagcagcucccucg...................................................................   1     1   7y1
.....................uAcguacguagcagagcagcucccucg...................................................................   2     1   7y1
.....................uucguacguagcagagcagcucccucC...................................................................   1     1   7y1
.....................uucguacguagcagGgcagcucccucg...................................................................   2     1   7y1
.....................uucguacguagcagagcagcucccucA...................................................................   1     1   7y1
.....................uucguacguagcagaUcagcucccucg...................................................................   3     1   7y1
.....................Nucguacguagcagagcagcucccucg...................................................................   1     1   7y1
```

```
  Star                                            Mature
gaccugcuucugggucggggguuucguacguagcagagcagcucccucgcugcgaucuauugaaagucagcccucgacacaaggguuuguccgcgcgcgcgcgcgcgcgugc
.....................uucguacguagcagagcagcucAcucg...............................................................    4      1    7y1
.....................uucguacguagcagagcaCcucccucg...............................................................    1      1    7y1
.....................uucguacCuagcagagcagcucccucg...............................................................    3      1    7y1
.....................uucgAacguagcagagcagcucccucg...............................................................    2      1    7y1
.....................uucguacguaCcagagcagcucccucg...............................................................    1      1    7y1
.....................uucguacguagcagagcagcucccGcg...............................................................    1      1    7y1
.....................uuAguacguagcagagcagcucccucg...............................................................    3      1    7y1
.....................Gucguacguagcagagcagcucccucg...............................................................    1      1    7y1
.....................uucguaAguagcagagcagcucccucg...............................................................    5      1    7y1
.....................uucguacguagcagCgcagcucccucg...............................................................    2      1    7y1
.....................uucguGcguagcagagcagcucccucg...............................................................    1      1    7y1
.....................uucguacgCagcagagcagcucccucg...............................................................    1      1    7y1
.....................uucguacguagcagagcagcuAccucg...............................................................    1      1    7y1
.....................uucguacguagcagagcagcuccAucg...............................................................    1      1    7y1
.....................uucguacguagcagagcagcucccucg...............................................................    1431   0    7y1
.....................Aucguacguagcagagcagcucccucg...............................................................    13     1    7y1
.....................uucguacguagcagagcagcucccAcg...............................................................    1      1    7y1
.....................uucgCacguagcagagcagcucccucg...............................................................    1      1    7y1
.....................uucguacguagcagagAagcucccucg...............................................................    1      1    7y1
.....................uucguacguagcagagUagcucccucg...............................................................    1      1    7y1
.....................uucguacguagcagagcagAucccucgc..............................................................    1      1    7y1
.....................uuAguacguagcagagcagcucccucgc..............................................................    2      1    7y1
.....................uucguacguagcagagcaAcucccucgc..............................................................    1      1    7y1
.....................uucguacguagcagagcaCcucccucgc..............................................................    1      1    7y1
.....................uucguaAguagcagagcagcucccucgc..............................................................    3      1    7y1
.....................uucguacguagcagagcagcucccucgA..............................................................    1      1    7y1
.....................Cucguacguagcagagcagcucccucgc..............................................................    1      1    7y1
.....................uucguacguagcagagcagcucccucgc..............................................................    923    0    7y1
.....................uucgCacguagcagagcagcucccucgc..............................................................    1      1    7y1
.....................uucguacguagcagagcagGucccucgc..............................................................    1      1    7y1
.....................uAcguacguagcagagcagcucccucgc..............................................................    1      1    7y1
.....................uucguacguagcagagcagcuGccucgc..............................................................    1      1    7y1
.....................uucguacguaCcagagcagcucccucgc..............................................................    1      1    7y1
.....................uucguacguagcagagcagcuAccucgc..............................................................    2      1    7y1
.....................Aucguacguagcagagcagcucccucgc..............................................................    15     1    7y1
.....................uucguacgAagcagagcagcucccucgc..............................................................    1      1    7y1
.....................Gucguacguagcagagcagcucccucgc..............................................................    1      1    7y1
.....................uucguacguagcagagcGgcucccucgc..............................................................    1      1    7y1
.....................uucguacUuagcagagcagcucccucgc..............................................................    1      1    7y1
.....................uucguacguagcagagcagcucAcucgc..............................................................    1      1    7y1
.....................Nucguacguagcagagcagcucccucgc..............................................................    1      1    7y1
.....................uucguacguagcagagcagcucccuAgc..............................................................    1      1    7y1
.....................uucguacguagcagagcagcucccucgcA.............................................................    2      1    7y1
.....................uucguacUuagcagagcagcucccucgcu.............................................................    3      1    7y1
.....................uucguacguagcagagcagcCcccucgcu.............................................................    1      1    7y1
.....................uCcguacguagcagagcagcucccucgcu.............................................................    2      1    7y1
.....................uucguacguagcagagAagcucccucgcu.............................................................    3      1    7y1
.....................uucguacguagcagagcUgcucccucgcu.............................................................    2      1    7y1
.....................uucguacguagcagagcagcucAcucgcu.............................................................    9      1    7y1
.....................uucguacguagcagagcaUcucccucgcu.............................................................    2      1    7y1
.....................uucguaAguagcagagcagcucccucgcu.............................................................    13     1    7y1
.....................uucguacguagcagagcGgcucccucgcu.............................................................    2      1    7y1
.....................uucguacguagcagagcagcucGcucgcu.............................................................    1      1    7y1
.....................uucguacguagAagagcagcucccucgcu.............................................................    6      1    7y1
.....................uucguacCuagcagagcagcucccucgcu.............................................................    1      1    7y1
.....................uucguacguagcagagcagcucccucgGu.............................................................    1      1    7y1
.....................uucAuacguagcagagcagcucccucgcu.............................................................    1      1    7y1
.....................uucguGcguagcagagcagcucccucgcu.............................................................    1      1    7y1
.....................uucCuacguagcagagcagcucccucgcu.............................................................    4      1    7y1
.....................uucguacguagcagagcagcucccucgcu.............................................................    4044   0    7y1
.....................Gucguacguagcagagcagcucccucgcu.............................................................    6      1    7y1
.....................uucguacguagcagagcagcuUccucgcu.............................................................    1      1    7y1
.....................uucguacguagcagagcagAucccucgcu.............................................................    1      1    7y1
.....................uucguacguagcagagcagcucccAcgcu.............................................................    2      1    7y1
.....................uGcguacguagcagagcagcucccucgcu.............................................................    1      1    7y1
.....................uucguacguagcGgagcagcucccucgcu.............................................................    2      1    7y1
.....................uucguacguagcagagcagcucccucUcu.............................................................    1      1    7y1
.....................uucguacguagcagagcagcucccucgAu.............................................................    3      1    7y1
.....................uucguacguagcaUagcagcucccucgcu.............................................................    4      1    7y1
.....................uucguacAuagcagagcagcucccucgcu.............................................................    2      1    7y1
```

**Star** **Mature**

gaccugcuucugggucgggguuucguacguagcagagcagcucccucgcugcgaucuauugaaagucagcccucgacacaaggguuuguccgcgcgcgcgcgcgcgcgcgugc

| | | | |
|---|---|---|---|
| .....................uucguacgAagcagagcagcucccucgcu............................................................... | 4 | 1 | 7y1 |
| .....................uucguacguagcagagcagcuAccucgcu............................................................... | 5 | 1 | 7y1 |
| .....................uAcguacguagcagagcagcucccucgcu............................................................... | 6 | 1 | 7y1 |
| .....................uuGguacguagcagagcagcucccucgcu............................................................... | 2 | 1 | 7y1 |
| .....................uucguacguagcUgagcagcucccucgcu............................................................... | 1 | 1 | 7y1 |
| .....................uucguacguaCcagagcagcucccucgcu............................................................... | 1 | 1 | 7y1 |
| .....................uucguacguagcagagcagcucccucCcu............................................................... | 4 | 1 | 7y1 |
| .....................uucguacguagcagagcCgcucccucgcu............................................................... | 1 | 1 | 7y1 |
| .....................Aucguacguagcagagcagcucccucgcu............................................................... | 36 | 1 | 7y1 |
| .....................uNcguacguagcagagcagcucccucgcu............................................................... | 1 | 1 | 7y1 |
| .....................uuAguacguagcagagcagcucccucgcu............................................................... | 8 | 1 | 7y1 |
| .....................uucguacguagcagagcagcucccucgcG............................................................... | 23 | 1 | 7y1 |
| .....................uucguaUguagcagagcagcucccucgcu............................................................... | 1 | 1 | 7y1 |
| .....................uucguacguagcagagcagcuccAucgcu............................................................... | 2 | 1 | 7y1 |
| .....................uucguacguagcagagcagcucccuAgcu............................................................... | 1 | 1 | 7y1 |
| .....................uucguacguagcagagcagcAcccucgcu............................................................... | 1 | 1 | 7y1 |
| .....................uucgAacguagcagagcagcucccucgcu............................................................... | 1 | 1 | 7y1 |
| .....................uucgGacguagcagagcagcucccucgcu............................................................... | 1 | 1 | 7y1 |
| .....................Nucguacguagcagagcagcucccucgcu............................................................... | 2 | 1 | 7y1 |
| .....................uucCuacguagcagagcagcucccucgcug.............................................................. | 1 | 1 | 7y1 |
| .....................uucguacguagcagagcagcucccuAgcug.............................................................. | 2 | 1 | 7y1 |
| .....................uucguacgAagcagagcagcucccucgcug.............................................................. | 1 | 1 | 7y1 |
| .....................uucguaAguagcagagcagcucccucgcug.............................................................. | 4 | 1 | 7y1 |
| .....................uucguacUuagcagagcagcucccucgcug.............................................................. | 1 | 1 | 7y1 |
| .....................Nucguacguagcagagcagcucccucgcug.............................................................. | 1 | 1 | 7y1 |
| .....................uucguacguagcagaCcagcucccucgcug.............................................................. | 1 | 1 | 7y1 |
| .....................Aucguacguagcagagcagcucccucgcug.............................................................. | 5 | 1 | 7y1 |
| .....................uucguacguagcagagcagcucccucgcGg.............................................................. | 1 | 1 | 7y1 |
| .....................uucguacguagcagagcagAucccucgcug.............................................................. | 1 | 1 | 7y1 |
| .....................uucguacguagcagagcagcucccucgcug.............................................................. | 459 | 0 | 7y1 |
| .....................uAcguacguagcagagcagcucccucgcug.............................................................. | 2 | 1 | 7y1 |
| .....................uucguacCuagcagagcagcucccucgcug.............................................................. | 2 | 1 | 7y1 |
| .....................Gucguacguagcagagcagcucccucgcugc............................................................. | 1 | 1 | 7y1 |
| .....................uucguGcguagcagagcagcucccucgcugc............................................................. | 1 | 1 | 7y1 |
| .....................uucguacguagcagagcagcucccucgcuCc............................................................. | 1 | 1 | 7y1 |
| .....................uucguacguagcaUagcagcucccucgcugc............................................................. | 1 | 1 | 7y1 |
| .....................uucguacUuagcagagcagcucccucgcugc............................................................. | 1 | 1 | 7y1 |
| .....................Aucguacguagcagagcagcucccucgcugc............................................................. | 9 | 1 | 7y1 |
| .....................uucguacguaCcagagcagcucccucgcugc............................................................. | 1 | 1 | 7y1 |
| .....................uucguacguagcagagcagcucAcucgcugc............................................................. | 2 | 1 | 7y1 |
| .....................uucguaAguagcagagcagcucccucgcugc............................................................. | 1 | 1 | 7y1 |
| .....................uucguacguagcagagcCgcucccucgcugc............................................................. | 1 | 1 | 7y1 |
| .....................uucguacguagAagagcagcucccucgcugc............................................................. | 2 | 1 | 7y1 |
| .....................uuAguacguagcagagcagcucccucgcugc............................................................. | 1 | 1 | 7y1 |
| .....................uucguacguagcagaCcagcucccucgcugc............................................................. | 1 | 1 | 7y1 |
| .....................uucguacguagcagagcagcucccucgcugA............................................................. | 1 | 1 | 7y1 |
| .....................uucguacguagcagagcagcuccAucgcugc............................................................. | 1 | 1 | 7y1 |
| .....................uAcguacguagcagagcagcucccucgcugc............................................................. | 3 | 1 | 7y1 |
| .....................uucguacguagcagGgcagcucccucgcugc............................................................. | 1 | 1 | 7y1 |
| .....................uucguacguagcagagcagcucccucgcugc............................................................. | 924 | 0 | 7y1 |
| ......................ucguacCuagcagagcagc........................................................................ | 1 | 1 | 7y1 |
| ......................ucguacguagcagagcagc........................................................................ | 1251 | 0 | 7y1 |
| ......................ucguacguagAagagcagc........................................................................ | 3 | 1 | 7y1 |
| ......................ucguacguagcagagcagA........................................................................ | 1 | 1 | 7y1 |
| ......................ucguacgAagcagagcagc........................................................................ | 3 | 1 | 7y1 |
| ......................Acguacguagcagagcagc........................................................................ | 13 | 1 | 7y1 |
| ......................uAguacguagcagagcagc........................................................................ | 2 | 1 | 7y1 |
| ......................ucguaAguagcagagcagc........................................................................ | 2 | 1 | 7y1 |
| ......................ucguacguagcagCgcagc........................................................................ | 4 | 1 | 7y1 |
| ......................Gcguacguagcagagcagc........................................................................ | 1 | 1 | 7y1 |
| ......................ucguacUuagcagagcagc........................................................................ | 1 | 1 | 7y1 |
| ......................ucguacguagcagagcagG........................................................................ | 1 | 1 | 7y1 |
| ......................ucguacguagcagagcaUc........................................................................ | 2 | 1 | 7y1 |
| ......................ucguacguagcaCagcagc........................................................................ | 1 | 1 | 7y1 |
| ......................ucguacguagcGgagcagc........................................................................ | 1 | 1 | 7y1 |
| ......................ucguacguagcaCagcagcu....................................................................... | 2 | 1 | 7y1 |
| ......................uNguacguagcagagcagcu....................................................................... | 1 | 1 | 7y1 |
| ......................uAguacguagcagagcagcu....................................................................... | 6 | 1 | 7y1 |
| ......................ucUuacguagcagagcagcu....................................................................... | 2 | 1 | 7y1 |
| ......................ucguacguagcagagcagcG....................................................................... | 20 | 1 | 7y1 |

**Star** **Mature**

```
gaccugcuucugggucggggguuucguacguagcagagcagcucccucgcugcgaucuauugaaagucagcccucgacacaaggguuuguccgcgcgcgcgcgcgcgcgcgcgugc
......................ucguacguagcagagAagcu.........................................................................  6     1  7y1
......................ucguacguagAagagcagcu.........................................................................  5     1  7y1
......................ucguacguagcagagcaUcu.........................................................................  4     1  7y1
......................ucguacguagcagCgcagcu.........................................................................  4     1  7y1
......................Gcguacguagcagagcagcu.........................................................................  6     1  7y1
......................ucguacguagcagagcagcu.........................................................................  2186  0  7y1
......................ucguacguagcagagcagcA.........................................................................  1     1  7y1
......................uUguacguagcagagcagcu.........................................................................  1     1  7y1
......................ucguacguagcagaUcagcu.........................................................................  2     1  7y1
......................ucguacguaCcagagcagcu.........................................................................  1     1  7y1
......................ucguaAguagcagagcagcu.........................................................................  6     1  7y1
......................ucguacguagcagagcGgcu.........................................................................  1     1  7y1
......................ucguacguagcagUgcagcu.........................................................................  1     1  7y1
......................Acguacguagcagagcagcu.........................................................................  36    1  7y1
......................ucguacguaUcagagcagcu.........................................................................  2     1  7y1
......................ucguGcguagcagagcagcu.........................................................................  1     1  7y1
......................ucguacguagcagagcUgcu.........................................................................  1     1  7y1
......................ucguacgAagcagagcagcu.........................................................................  2     1  7y1
......................uAguacguagcagagcagcuc........................................................................  19    1  7y1
......................ucguacguagcagaUcagcuc........................................................................  6     1  7y1
......................ucgAacguagcagagcagcuc........................................................................  1     1  7y1
......................ucguacguaCcagagcagcuc........................................................................  6     1  7y1
......................ucguacCuagcagagcagcuc........................................................................  4     1  7y1
......................ucguacguagcagagcaCcuc........................................................................  6     1  7y1
......................ucUuacguagcagagcagcuc........................................................................  5     1  7y1
......................ucguacguagcagagcagcuc........................................................................  6627  0  7y1
......................Acguacguagcagagcagcuc........................................................................  79    1  7y1
......................ucguacguagcagagGagcuc........................................................................  1     1  7y1
......................ucguacguagAagagcagcuc........................................................................  9     1  7y1
......................ucguacguagcagGgcagcuc........................................................................  2     1  7y1
......................ucguacguagcagagcagcuA........................................................................  7     1  7y1
......................ucguacguagcagagUagcuc........................................................................  1     1  7y1
......................ucguacguagcaCagcagcuc........................................................................  238   1  7y1
......................ucguacguagcagUgcagcuc........................................................................  1     1  7y1
......................ucguacguagUagagcagcuc........................................................................  1     1  7y1
......................ucguacgAagcagagcagcuc........................................................................  9     1  7y1
......................Gcguacguagcagagcagcuc........................................................................  7     1  7y1
......................ucguacUuagcagagcagcuc........................................................................  6     1  7y1
......................Ncguacguagcagagcagcuc........................................................................  4     1  7y1
......................ucguacgCagcagagcagcuc........................................................................  2     1  7y1
......................ucguacguagcagagcGgcuc........................................................................  1     1  7y1
......................uGguacguagcagagcagcuc........................................................................  1     1  7y1
......................uUguacguagcagagcagcuc........................................................................  1     1  7y1
......................ucguacguagcagagcagGuc........................................................................  1     1  7y1
......................ucguacguagcagCgcagcuc........................................................................  5     1  7y1
......................ucguacguaAcagagcagcuc........................................................................  1     1  7y1
......................ucguacguagcagaCcagcuc........................................................................  1     1  7y1
......................ucguacguagcagagcagcGc........................................................................  3     1  7y1
......................ucguacAuagcagagcagcuc........................................................................  1     1  7y1
......................ucguacguagcagagcagcAc........................................................................  4     1  7y1
......................ucCuacguagcagagcagcuc........................................................................  5     1  7y1
......................ucguacguagcagagcCgcuc........................................................................  1     1  7y1
......................ucguacguaUcagagcagcuc........................................................................  3     1  7y1
......................ucguacguagcagagcUgcuc........................................................................  2     1  7y1
......................ucguaAguagcagagcagcuc........................................................................  15    1  7y1
......................ucguacguagcagagcaUcuc........................................................................  3     1  7y1
......................ucguUcguagcagagcagcuc........................................................................  1     1  7y1
......................ucguacguagcGgagcagcuc........................................................................  2     1  7y1
......................ucguacguagcagagAagcuc........................................................................  15    1  7y1
......................ucguaUguagcagagcagcuc........................................................................  2     1  7y1
......................ucguaGguagcagagcagcuc........................................................................  1     1  7y1
......................ucguacguagcagagcagcuG........................................................................  1     1  7y1
......................ucAuacguagcagagcagcuc........................................................................  1     1  7y1
......................ucguacguagcagagcagAuc........................................................................  3     1  7y1
......................ucguacguGgcagagcagcucc.......................................................................  1     1  7y1
......................ucguacgAagcagagcagcucc.......................................................................  1     1  7y1
......................ucguacguagcagaUcagcucc.......................................................................  1     1  7y1
......................ucguacguagAagagcagcucc.......................................................................  1     1  7y1
......................ucguacguagcGgagcagcucc.......................................................................  1     1  7y1
......................uGguacguagcagagcagcucc.......................................................................  1     1  7y1
```

```
 Star                                                                  Mature
gaccugcuucugggucgggguuucguacguagcagagcagcucccucgcugcgaucuauugaaagucagcccucgacacaaggguuuguccgcgcgcgcgcgcgcgcgcgcgugc
......................ucguacguagcagagUagcucc.......................................................................    1     1    7y1
......................Ncguacguagcagagcagcucc.......................................................................    1     1    7y1
......................ucguacguagcagagcagcucA.......................................................................    6     1    7y1
......................ucguacguagcagagcagcucU.......................................................................    1     1    7y1
......................ucUuacguagcagagcagcucc.......................................................................    2     1    7y1
......................ucguacguagcagagAagcucc.......................................................................    2     1    7y1
......................ucguacguagcagagcagAucc.......................................................................    1     1    7y1
......................Acguacguagcagagcagcucc.......................................................................    15    1    7y1
......................Gcguacguagcagagcagcucc.......................................................................    5     1    7y1
......................ucguacguagcagaCcagcucc.......................................................................    1     1    7y1
......................ucguacguagcagagcagcucc.......................................................................    1758  0    7y1
......................ucguacguagcagagcagcCcc.......................................................................    1     1    7y1
......................ucguaAguagcagagcagcucc.......................................................................    4     1    7y1
......................ucguacguaUcagagcagcucc.......................................................................    2     1    7y1
......................ucguaUguagcagagcagcucc.......................................................................    1     1    7y1
......................ucguacguagcagagcagcuAc.......................................................................    4     1    7y1
......................ucguacguagcagagcagcuUc.......................................................................    1     1    7y1
......................uAguacguagcagagcagcucc.......................................................................    1     1    7y1
......................ucguacguagcagGgcagcucc.......................................................................    1     1    7y1
......................ucguGcguagcagagcagcucc.......................................................................    1     1    7y1
......................ucguacguaCcagagcagcucc.......................................................................    4     1    7y1
......................ucguacguagcagaAcagcucc.......................................................................    1     1    7y1
......................ucCuacguagcagagcagcucc.......................................................................    2     1    7y1
......................ucguacguagcagUgcagcuccc......................................................................    1     1    7y1
......................ucguacguagcagagcagcuAcc......................................................................    3     1    7y1
......................ucguacguagcagagcagcuccc......................................................................    2876  0    7y1
......................Gcguacguagcagagcagcuccc......................................................................    4     1    7y1
......................ucguacguagcagagcagcAccc......................................................................    1     1    7y1
......................ucguacguagcagagcagcucAc......................................................................    5     1    7y1
......................ucguacguagAagagcagcuccc......................................................................    3     1    7y1
......................ucguaAguagcagagcagcuccc......................................................................    8     1    7y1
......................uAguacguagcagagcagcuccc......................................................................    2     1    7y1
......................ucguacguagcagagcaUcuccc......................................................................    1     1    7y1
......................ucguacguaUcagagcagcuccc......................................................................    1     1    7y1
......................uUguacguagcagagcagcuccc......................................................................    2     1    7y1
......................ucguacguagcagagcaCcuccc......................................................................    2     1    7y1
......................ucguacguagcagagAagcuccc......................................................................    4     1    7y1
......................ucguacguaCcagagcagcuccc......................................................................    2     1    7y1
......................ucguacguagcagCgcagcuccc......................................................................    7     1    7y1
......................ucguacguagcagagcagcuccA......................................................................    2     1    7y1
......................ucAuacguagcagagcagcuccc......................................................................    2     1    7y1
......................ucUuacguagcagagcagcuccc......................................................................    1     1    7y1
......................ucgAacguagcagagcagcuccc......................................................................    1     1    7y1
......................ucguacguagcagagcagcuGcc......................................................................    1     1    7y1
......................ucguUcguagcagagcagcuccc......................................................................    1     1    7y1
......................ucguacguagcagagcagAuccc......................................................................    4     1    7y1
......................Acguacguagcagagcagcuccc......................................................................    26    1    7y1
......................uGguacguagcagagcagcuccc......................................................................    1     1    7y1
......................ucguacCuagcagagcagcuccc......................................................................    1     1    7y1
......................ucguacgAagcagagcagcuccc......................................................................    1     1    7y1
......................ucguacguagcagagGagcuccc......................................................................    1     1    7y1
......................ucguacguagcagaAcagcuccc......................................................................    1     1    7y1
......................ucguacguagcaCagcagcuccc......................................................................    2     1    7y1
......................ucguacgAagcagagcagcucccu.....................................................................    3     1    7y1
......................Ccguacguagcagagcagcucccu.....................................................................    1     1    7y1
......................ucguacguagcagCgcagcucccu.....................................................................    6     1    7y1
......................uUguacguagcagagcagcucccu.....................................................................    1     1    7y1
......................ucguacguaUcagagcagcucccu.....................................................................    1     1    7y1
......................ucguacguagcagagcagcucGcu.....................................................................    1     1    7y1
......................ucguacguagcUgagcagcucccu.....................................................................    1     1    7y1
......................ucguacguagcagagcagcuGccu.....................................................................    1     1    7y1
......................ucguacguagcagagcagUucccu.....................................................................    1     1    7y1
......................ucguacguagcagagcagcuAccu.....................................................................    2     1    7y1
......................Gcguacguagcagagcagcucccu.....................................................................    5     1    7y1
......................ucguacguagcagaCcagcucccu.....................................................................    2     1    7y1
......................ucguacguagcaUagcagcucccu.....................................................................    2     1    7y1
......................uAguacguagcagagcagcucccu.....................................................................    6     1    7y1
......................ucguacguagcagagAagcucccu.....................................................................    1     1    7y1
......................ucguacguagcagagcagcCcccu.....................................................................    1     1    7y1
......................ucguacguagcagagcCgcucccu.....................................................................    2     1    7y1
```

Star                                      Mature

```
gaccugcuucugggucgggguuucguacguagcagagcagcucccucgcugcgaucuauugaaagucagcccucgacacaaggguuuguccgcgcgcgcgcgcgcgcgugc
......................ucguacguGgcagagcagcucccu.................................................................   2     1   7y1
......................ucguGcguagcagagcagcucccu.................................................................   1     1   7y1
......................ucguacguaCcagagcagcucccu.................................................................   1     1   7y1
......................ucguaAguagcagagcagcucccu.................................................................   3     1   7y1
......................ucguacguagcagagcagcucccG.................................................................   9     1   7y1
......................ucguacguagcGgagcagcucccu.................................................................   2     1   7y1
......................ucgAacguagcagagcagcucccu.................................................................   1     1   7y1
......................ucguacguagcagagcagAucccu.................................................................   1     1   7y1
......................ucguacguagcagagcagcucccu.................................................................   2486  0   7y1
......................ucguacguagcagagcagcucAcu.................................................................   4     1   7y1
......................ucCuacguagcagagcagcucccu.................................................................   2     1   7y1
......................ucguacguagcagagUagcucccu.................................................................   1     1   7y1
......................ucguacguagcagagcagcuUccu.................................................................   1     1   7y1
......................ucguacguagcagagcagcucccA.................................................................   5     1   7y1
......................ucguacguagcagagcagGucccu.................................................................   1     1   7y1
......................ucguacguagAagagcagcucccu.................................................................   1     1   7y1
......................Acguacguagcagagcagcucccu.................................................................   21    1   7y1
......................ucguacguagcagagcagcuccAu.................................................................   6     1   7y1
......................ucguacUuagcagagcagcucccu.................................................................   3     1   7y1
......................ucguacCuagcagagcagcucccu.................................................................   1     1   7y1
......................uUguacguagcagagcagcucccuc................................................................   1     1   7y1
......................ucguacCuagcagagcagcucccuc................................................................   1     1   7y1
......................ucgAacguagcagagcagcucccuc................................................................   1     1   7y1
......................ucguacguagcagagcagcuAccuc................................................................   1     1   7y1
......................ucguacguagcagagcagcuUccuc................................................................   1     1   7y1
......................ucguacguagAagagcagcucccuc................................................................   1     1   7y1
......................ucCuacguagcagagcagcucccuc................................................................   1     1   7y1
......................ucguacguagcagagcagcucccuG................................................................   1     1   7y1
......................ucguacguagcagagcagcucccuc................................................................   555   0   7y1
......................Acguacguagcagagcagcucccuc................................................................   7     1   7y1
......................ucguacguaCcagagcagcucccuc................................................................   1     1   7y1
......................ucguaAguagcagagcagcucccuc................................................................   1     1   7y1
......................Ncguacguagcagagcagcucccuc................................................................   1     1   7y1
......................ucguacguagcagagcagcAcccuc................................................................   1     1   7y1
......................ucguacguagcagagcGgcucccuc................................................................   1     1   7y1
......................ucguacguagcagagcagcucAcuc................................................................   1     1   7y1
......................ucguacguagcagCgcagcucccucg...............................................................   1     1   7y1
......................Ncguacguagcagagcagcucccucg...............................................................   1     1   7y1
......................ucguacguagcagagcagcuccAucg...............................................................   1     1   7y1
......................ucguacguaCcagagcagcucccucg...............................................................   1     1   7y1
......................ucguacguagcCgagcagcucccucg...............................................................   1     1   7y1
......................ucguacguagcagaUcagcucccucg...............................................................   1     1   7y1
......................ucguacgAagcagagcagcucccucg...............................................................   1     1   7y1
......................ucCuacguagcagagcagcucccucg...............................................................   1     1   7y1
......................ucguacguagcagagcaCcucccucg...............................................................   1     1   7y1
......................ucguacguagcagagcagcucccucg...............................................................   219   0   7y1
......................ucguacAuagcagagcagcucccucg...............................................................   1     1   7y1
......................uAguacguagcagagcagcucccucg...............................................................   1     1   7y1
......................ucguaAguagcagagcagcucccucgc..............................................................   2     1   7y1
......................ucguacguagcagagcagcucccuAgc..............................................................   1     1   7y1
......................ucguacguagcagagcagcuAccucgc..............................................................   1     1   7y1
......................ucguacguagcagagcagcucccucgc..............................................................   270   0   7y1
......................Acguacguagcagagcagcucccucgc..............................................................   4     1   7y1
......................uAguacguagcagagcagcucccucgc..............................................................   2     1   7y1
......................ucguacguagcagagAagcucccucgc..............................................................   1     1   7y1
......................ucguacguagcagagcagcucccucAcu.............................................................   1     1   7y1
......................ucguacguagcagaAcagcucccucgcu.............................................................   1     1   7y1
......................ucguacguaAcagagcagcucccucgcu.............................................................   1     1   7y1
......................ucguacguagcagagcagcCcccucgcu.............................................................   1     1   7y1
......................ucguacguaCcagagcagcucccucgcu.............................................................   1     1   7y1
......................ucguacguagcagagcagcucccucgcA.............................................................   1     1   7y1
......................ucguacguagcagagcagcuAccucgcu.............................................................   1     1   7y1
......................ucguacUuagcagagcagcucccucgcu.............................................................   1     1   7y1
......................ucguacguagcagUgcagcucccucgcu.............................................................   1     1   7y1
......................ucguacguagcagagcagcucccAcgcu.............................................................   1     1   7y1
......................ucguacguagAagagcagcucccucgcu.............................................................   1     1   7y1
......................ucguacguagcagagcagcucccucgcu.............................................................   1342  0   7y1
......................uUguacguagcagagcagcucccucgcu.............................................................   1     1   7y1
......................ucguacguagcagagAagcucccucgcu.............................................................   3     1   7y1
......................ucguacAuagcagagcagcucccucgcu.............................................................   2     1   7y1
```

**Star** **Mature**

```
gaccugcuucugggucgggguuucguacguagcagagcagcucccucgcugcgaucuauugaaagucagcccucgacacaaggguuuguccgcgcgcgcgcgcgcgcgcgugc
......................ucguacgAagcagagcagcucccucgcu...............................................................    4     1    7y1
......................ucguacguagcagagcagcucccucUcu...............................................................    1     1    7y1
......................ucguacguagcagagcagcuUccucgcu...............................................................    1     1    7y1
......................ucguaUguagcagagcagcucccucgcu...............................................................    1     1    7y1
......................ucguacguagcagagcagcucccucgcG...............................................................    16    1    7y1
......................ucguacguaUcagagcagcucccucgcu...............................................................    1     1    7y1
......................ucguacguagcagagcagcucGcucgcu...............................................................    1     1    7y1
......................ucguacCuagcagagcagcucccucgcu...............................................................    1     1    7y1
......................ucguacguagcagagcagcucAcucgcu...............................................................    1     1    7y1
......................ucguacguagcagagcagAucccucgcu...............................................................    2     1    7y1
......................ucguacguagcagagcagcucccuAgcu...............................................................    3     1    7y1
......................ucguacguagcagagcagcucccucCcu...............................................................    2     1    7y1
......................ucguaAguagcagagcagcucccucgcu...............................................................    1     1    7y1
......................ucguacguagcagagcagcuccAucgcu...............................................................    1     1    7y1
......................Acguacguagcagagcagcucccucgcu...............................................................    11    1    7y1
......................Ncguacguagcagagcagcucccucgcu...............................................................    3     1    7y1
......................uAguacguagcagagcagcucccucgcu...............................................................    2     1    7y1
......................ucguacguagcGgagcagcucccucgcu...............................................................    1     1    7y1
......................ucguacguagcagagcagcuAccucgcug..............................................................    3     1    7y1
......................ucguacguagcagagcagUucccucgcug..............................................................    1     1    7y1
......................uAguacguagcagagcagcucccucgcug..............................................................    3     1    7y1
......................Acguacguagcagagcagcucccucgcug..............................................................    17    1    7y1
......................ucguacguagcagagcagcuccAucgcug..............................................................    4     1    7y1
......................ucguacguagcagagcagcucccucUcug..............................................................    1     1    7y1
......................ucguacguagGagagcagcucccucgcug..............................................................    1     1    7y1
......................ucguacguaUcagagcagcucccucgcug..............................................................    1     1    7y1
......................ucguacguagAagagcagcucccucgcug..............................................................    3     1    7y1
......................ucgAacguagcagagcagcucccucgcug..............................................................    1     1    7y1
......................ucguacguagcagagcagcucccucgcuC..............................................................    1     1    7y1
......................ucguGcguagcagagcagcucccucgcug..............................................................    1     1    7y1
......................ucguacguGgcagagcagcucccucgcug..............................................................    1     1    7y1
......................Gcguacguagcagagcagcucccucgcug..............................................................    3     1    7y1
......................ucguacCuagcagagcagcucccucgcug..............................................................    3     1    7y1
......................uGguacguagcagagcagcucccucgcug..............................................................    2     1    7y1
......................ucguacguaCcagagcagcucccucgcug..............................................................    1     1    7y1
......................ucguacguagcagagcagcuUccucgcug..............................................................    1     1    7y1
......................ucUuacguagcagagcagcucccucgcug..............................................................    1     1    7y1
......................ucguacguagcagagcagcucccAcgcug..............................................................    4     1    7y1
......................ucguaAguagcagagcagcucccucgcug..............................................................    2     1    7y1
......................ucguacguagcagagcagcucAcucgcug..............................................................    3     1    7y1
......................ucguacguagcaCagcagcucccucgcug..............................................................    1     1    7y1
......................ucguacguagcagagcagcucccuUgcug..............................................................    1     1    7y1
......................ucCuacguagcagagcagcucccucgcug..............................................................    3     1    7y1
......................ucguacguagcagagcagcucccucCcug..............................................................    1     1    7y1
......................ucguacguagcagagcagcucccucgcug..............................................................    2333  0    7y1
......................ucguacgAagcagagcagcucccucgcug..............................................................    4     1    7y1
......................Ncguacguagcagagcagcucccucgcug..............................................................    1     1    7y1
......................ucguacguagcagagcagcucccucgcuU..............................................................    6     1    7y1
......................Ccguacguagcagagcagcucccucgcug..............................................................    1     1    7y1
......................ucguacguagcagagcagcCcccucgcug..............................................................    2     1    7y1
......................ucguacguagcagagcagcucccucgcGg..............................................................    8     1    7y1
......................ucguacguagUagagcagcucccucgcug..............................................................    1     1    7y1
......................ucguacguagcagagcagcucccucgAug..............................................................    1     1    7y1
......................ucguacguagcagCgcagcucccucgcug..............................................................    2     1    7y1
.......................cguacguagUagagcagc........................................................................    1     1    7y1
.......................cguacguagcaUagcagc........................................................................    1     1    7y1
.......................cguacguagcagaUcagc........................................................................    2     1    7y1
.......................cguacguaAcagagcagc........................................................................    1     1    7y1
.......................cguacguagcagagcCgc........................................................................    1     1    7y1
.......................cguacguagcagagcagc........................................................................    1761  0    7y1
.......................cguacguaUcagagcagc........................................................................    3     1    7y1
.......................cguacguagcUgagcagc........................................................................    1     1    7y1
.......................cguacguagcGgagcagc........................................................................    2     1    7y1
.......................cguacAuagcagagcagc........................................................................    1     1    7y1
.......................cguacguagcagagAagc........................................................................    2     1    7y1
.......................cgAacguagcagagcagc........................................................................    2     1    7y1
.......................cUuacguagcagagcagc........................................................................    3     1    7y1
.......................cguacguagcagagcaUc........................................................................    1     1    7y1
.......................Gguacguagcagagcagc........................................................................    2     1    7y1
.......................cguacgCagcagagcagc........................................................................    2     1    7y1
```

**Star** **Mature**

```
gaccugcuucugggucgggguuucguacguagcagagcagcucccucgcugcgaucuauugaaagucagcccucgacacaaggguuuguccgcgcgcgcgcgcgcgcgugc
.......................cguacguagcagCgcagc......................................................................    2      1    7y1
.......................cguacguagAagagcagc......................................................................    2      1    7y1
.......................cguacgAagcagagcagc......................................................................    1      1    7y1
.......................cguCcguagcagagcagc......................................................................    1      1    7y1
.......................cguacguagcagagcGgc......................................................................    1      1    7y1
.......................cguacNuagcagagcagc......................................................................    1      1    7y1
.......................cguacguagcaAagcagc......................................................................    1      1    7y1
.......................cCuacguagcagagcagc......................................................................    1      1    7y1
.......................cguacguagcagagcagA......................................................................    2      1    7y1
.......................Aguacguagcagagcagc......................................................................    6      1    7y1
.......................Nguacguagcagagcagc......................................................................    2      1    7y1
.......................cAuacguagcagagcagc......................................................................    1      1    7y1
.......................cguaAguagcagagcagc......................................................................    1      1    7y1
.......................cCuacguagcagagcagcu.....................................................................    1      1    7y1
.......................Aguacguagcagagcagcu.....................................................................    5      1    7y1
.......................cguacguagUagagcagcu.....................................................................    1      1    7y1
.......................cguacguagcagagGagcu.....................................................................    2      1    7y1
.......................cUuacguagcagagcagcu.....................................................................    5      1    7y1
.......................cguacguagcagagcagcA.....................................................................    1      1    7y1
.......................cgAacguagcagagcagcu.....................................................................    1      1    7y1
.......................cAuacguagcagagcagcu.....................................................................    2      1    7y1
.......................cguacguagcagaAcagcu.....................................................................    1      1    7y1
.......................cguacguagcagaCcagcu.....................................................................    1      1    7y1
.......................cguacUuagcagagcagcu.....................................................................    1      1    7y1
.......................cguacguagcagagcagcu.....................................................................    1200   0    7y1
.......................cguacguagcagagcagAu.....................................................................    3      1    7y1
.......................cgCacguagcagagcagcu.....................................................................    1      1    7y1
.......................cguacguagAagagcagcu.....................................................................    3      1    7y1
.......................cguaAguagcagagcagcu.....................................................................    2      1    7y1
.......................cguacguagcagagcagcG.....................................................................    4      1    7y1
.......................cguacguagcaUagcagcu.....................................................................    1      1    7y1
.......................Gguacguagcagagcagcu.....................................................................    1      1    7y1
.......................cguacguagcagagAagcu.....................................................................    3      1    7y1
.......................cguacguagcagagcaUcu.....................................................................    1      1    7y1
.......................cguacguagcagCgcagcu.....................................................................    1      1    7y1
.......................Nguacguagcagagcagcu.....................................................................    1      1    7y1
.......................cguacguagcagagcaCcu.....................................................................    2      1    7y1
.......................cguacgAagcagagcagcu.....................................................................    3      1    7y1
.......................cguacguagcagagcaUcuc....................................................................    5      1    7y1
.......................cguacguagcagagcaCcuc....................................................................    4      1    7y1
.......................cguacguaAcagagcagcuc....................................................................    1      1    7y1
.......................cguacguagUagagcagcuc....................................................................    1      1    7y1
.......................cguacguagcagagcagUuc....................................................................    1      1    7y1
.......................cguacguagcagagcagcAc....................................................................    5      1    7y1
.......................cguacguagAagagcagcuc....................................................................    22     1    7y1
.......................cguGcguagcagagcagcuc....................................................................    2      1    7y1
.......................cguacguagcagUgcagcuc....................................................................    1      1    7y1
.......................cUuacguagcagagcagcuc....................................................................    5      1    7y1
.......................cguacguagcagagcagcGc....................................................................    1      1    7y1
.......................cguacguaUcagagcagcuc....................................................................    4      1    7y1
.......................cguacguagcagagUagcuc....................................................................    1      1    7y1
.......................cguacguaCcagagcagcuc....................................................................    2      1    7y1
.......................cguacguagcaUagcagcuc....................................................................    3      1    7y1
.......................cguacguagcagagcagcuA....................................................................    8      1    7y1
.......................cguacguagcagagAagcuc....................................................................    29     1    7y1
.......................cguaGguagcagagcagcuc....................................................................    1      1    7y1
.......................cguacguagcagagGagcuc....................................................................    2      1    7y1
.......................cguacguagcagGgcagcuc....................................................................    3      1    7y1
.......................cguacguagcagagcUgcuc....................................................................    1      1    7y1
.......................cgAacguagcagagcagcuc....................................................................    12     1    7y1
.......................Aguacguagcagagcagcuc....................................................................    25     1    7y1
.......................cguacguagGagagcagcuc....................................................................    1      1    7y1
.......................cguacAuagcagagcagcuc....................................................................    4      1    7y1
.......................cguacguagcagagcagcuG....................................................................    1      1    7y1
.......................cguacgAagcagagcagcuc....................................................................    23     1    7y1
.......................cguacguagcaCagcagcuc....................................................................    2      1    7y1
.......................cguacCuagcagagcagcuc....................................................................    2      1    7y1
.......................Nguacguagcagagcagcuc....................................................................    2      1    7y1
.......................cguacguagcagaCcagcuc....................................................................    4      1    7y1
.......................cCuacguagcagagcagcuc....................................................................    11     1    7y1
```

**Star** **Mature**

| | | | |
|---|---|---|---|
| gaccugcuucugggucgggguuucguacguagcagagcagcucccucgcugcgaucuauugaaagucagcccucgacacaaggguuuguccgcgcgcgcgcgcgcgcgcgcgugc | | | |
| .......................cguacguagcagagcagcuc........................................................................ | 7571 | 0 | 7y1 |
| .......................cguacguagcagagcGgcuc........................................................................ | 1 | 1 | 7y1 |
| .......................cguacgCagcagagcagcuc........................................................................ | 2 | 1 | 7y1 |
| .......................Gguacguagcagagcagcuc........................................................................ | 3 | 1 | 7y1 |
| .......................cguaAguagcagagcagcuc........................................................................ | 10 | 1 | 7y1 |
| .......................cguacUuagcagagcagcuc........................................................................ | 5 | 1 | 7y1 |
| .......................cguacguagcagagcagAuc........................................................................ | 9 | 1 | 7y1 |
| .......................Uguacguagcagagcagcuc........................................................................ | 1 | 1 | 7y1 |
| .......................cguacguagcagCgcagcuc........................................................................ | 4 | 1 | 7y1 |
| .......................cAuacguagcagagcagcuc........................................................................ | 1 | 1 | 7y1 |
| .......................cguacguagcagaUcagcuc........................................................................ | 4 | 1 | 7y1 |
| .......................cguacguagcGgagcagcuc........................................................................ | 2 | 1 | 7y1 |
| .......................cguacguagcagagcagcAcc....................................................................... | 1 | 1 | 7y1 |
| .......................cguacguagcagagAagcucc....................................................................... | 1 | 1 | 7y1 |
| .......................cguacguagcagagcagcuAc....................................................................... | 1 | 1 | 7y1 |
| .......................cguacguagcagCgcagcucc....................................................................... | 1 | 1 | 7y1 |
| .......................Aguacguagcagagcagcucc....................................................................... | 1 | 1 | 7y1 |
| .......................cguacguagcagagcagcucc....................................................................... | 434 | 0 | 7y1 |
| .......................cUuacguagcagagcagcucc....................................................................... | 1 | 1 | 7y1 |
| .......................cguacguagcagagNagcucc....................................................................... | 1 | 1 | 7y1 |
| .......................cguacguagcagaUcagcucc....................................................................... | 1 | 1 | 7y1 |
| .......................cguacguagcagagGagcucc....................................................................... | 1 | 1 | 7y1 |
| .......................cguacguaCcagagcagcucc....................................................................... | 1 | 1 | 7y1 |
| .......................cAuacguagcagagcagcucc....................................................................... | 1 | 1 | 7y1 |
| .......................cguaAguagcagagcagcucc....................................................................... | 1 | 1 | 7y1 |
| .......................cguacguagcagagAagcuccc...................................................................... | 2 | 1 | 7y1 |
| .......................cUuacguagcagagcagcuccc...................................................................... | 3 | 1 | 7y1 |
| .......................cguacguagcagagcagcAccc...................................................................... | 1 | 1 | 7y1 |
| .......................cguacguagcagagcagcuccA...................................................................... | 2 | 1 | 7y1 |
| .......................cguacgAagcagagcagcuccc...................................................................... | 1 | 1 | 7y1 |
| .......................cguacguagcagagcagcuccc...................................................................... | 660 | 0 | 7y1 |
| .......................Aguacguagcagagcagcuccc...................................................................... | 3 | 1 | 7y1 |
| .......................cguacguagcagagcaUcuccc...................................................................... | 2 | 1 | 7y1 |
| .......................cguacguagAagagcagcuccc...................................................................... | 1 | 1 | 7y1 |
| .......................cguacguagcagUgcagcuccc...................................................................... | 1 | 1 | 7y1 |
| .......................cguaAguagcagagcagcuccc...................................................................... | 1 | 1 | 7y1 |
| .......................cguacguagcagagcagcuGcc...................................................................... | 1 | 1 | 7y1 |
| .......................cguacguagcagagcagcuAcc...................................................................... | 1 | 1 | 7y1 |
| .......................cguacguagcagagcaUcucccu..................................................................... | 2 | 1 | 7y1 |
| .......................cguacguagcagagcagcucccA..................................................................... | 1 | 1 | 7y1 |
| .......................cguaUguagcagagcagcucccu..................................................................... | 1 | 1 | 7y1 |
| .......................cguacguagcagagcagcucccG..................................................................... | 6 | 1 | 7y1 |
| .......................cguacguagcagagcagcuccAu..................................................................... | 2 | 1 | 7y1 |
| .......................cguacguagcagagcGgcucccu..................................................................... | 2 | 1 | 7y1 |
| .......................cguacguagcagagcagcuAccu..................................................................... | 2 | 1 | 7y1 |
| .......................cguacguagcagagAagcucccu..................................................................... | 1 | 1 | 7y1 |
| .......................cAuacguagcagagcagcucccu..................................................................... | 1 | 1 | 7y1 |
| .......................cgAacguagcagagcagcucccu..................................................................... | 1 | 1 | 7y1 |
| .......................cguacguagcagagcagcAcccu..................................................................... | 1 | 1 | 7y1 |
| .......................cUuacguagcagagcagcucccu..................................................................... | 2 | 1 | 7y1 |
| .......................cCuacguagcagagcagcucccu..................................................................... | 1 | 1 | 7y1 |
| .......................cguacguagcagagcagcucccu..................................................................... | 943 | 0 | 7y1 |
| .......................cguacguagcagaUcagcucccu..................................................................... | 1 | 1 | 7y1 |
| .......................cguacguagcagagcUgcucccu..................................................................... | 1 | 1 | 7y1 |
| .......................cguacguagcaUagcagcucccu..................................................................... | 1 | 1 | 7y1 |
| .......................cguacguagcagagcagcucGcu..................................................................... | 1 | 1 | 7y1 |
| .......................cguacguagAagagcagcucccu..................................................................... | 1 | 1 | 7y1 |
| .......................cguacguagcagagcCgcucccu..................................................................... | 1 | 1 | 7y1 |
| .......................cguaAguagcagagcagcucccu..................................................................... | 1 | 1 | 7y1 |
| .......................cguacguagcagagcagGucccuc.................................................................... | 1 | 1 | 7y1 |
| .......................cguacCuagcagagcagcucccuc.................................................................... | 2 | 1 | 7y1 |
| .......................cguacguagcagagcagcucccAc.................................................................... | 2 | 1 | 7y1 |
| .......................cguacAuagcagagcagcucccuc.................................................................... | 1 | 1 | 7y1 |
| .......................cguacgAagcagagcagcucccuc.................................................................... | 4 | 1 | 7y1 |
| .......................cguacguagcagaUcagcucccuc.................................................................... | 2 | 1 | 7y1 |
| .......................cguacguagcagagcagcucAcuc.................................................................... | 2 | 1 | 7y1 |
| .......................Nguacguagcagagcagcucccuc.................................................................... | 1 | 1 | 7y1 |
| .......................cguacguagcagGgcagcucccuc.................................................................... | 1 | 1 | 7y1 |
| .......................cUuacguagcagagcagcucccuc.................................................................... | 3 | 1 | 7y1 |
| .......................cguacguagcagCgcagcucccuc.................................................................... | 2 | 1 | 7y1 |

**Star** **Mature**

| | | | |
|---|---|---|---|
| gaccugcuucugggucgggguuucguacguagcagagcagcucccucgcugcgaucuauugaaagucagcccucgacacaaggguuuguccgcgcgcgcgcgcgcgcgcgugc | | | |
| .......................cCuacguagcagagcagcucccuc.................................................................. | 1 | 1 | 7y1 |
| .......................cguacguaCcagagcagcucccuc.................................................................. | 1 | 1 | 7y1 |
| .......................cguacguagcagagcagcucccuc.................................................................. | 744 | 0 | 7y1 |
| .......................cguacguagcagagcagcAcccuc.................................................................. | 1 | 1 | 7y1 |
| .......................cguacguagcaCagcagcucccuc.................................................................. | 1 | 1 | 7y1 |
| .......................cguacguagcagagAagcucccuc.................................................................. | 4 | 1 | 7y1 |
| .......................cguacguagcagaAcagcucccuc.................................................................. | 2 | 1 | 7y1 |
| .......................cguacgGagcagagcagcucccuc.................................................................. | 1 | 1 | 7y1 |
| .......................cguacguagcagagcagcuAccuc.................................................................. | 1 | 1 | 7y1 |
| .......................cguacgAagcagagcagcucccucg................................................................. | 1 | 1 | 7y1 |
| .......................cguacguaUcagagcagcucccucg................................................................. | 1 | 1 | 7y1 |
| .......................cguacguagcagaUcagcucccucg................................................................. | 1 | 1 | 7y1 |
| .......................cguacguagcagGgcagcucccucg................................................................. | 1 | 1 | 7y1 |
| .......................cguacguagcagagcagcucccucg................................................................. | 297 | 0 | 7y1 |
| .......................cguacguagcagagAagcucccucg................................................................. | 1 | 1 | 7y1 |
| .......................cguacUuagcagagcagcucccucg................................................................. | 1 | 1 | 7y1 |
| .......................cUuacguagcagagcagcucccucg................................................................. | 1 | 1 | 7y1 |
| .......................Aguacguagcagagcagcucccucg................................................................. | 2 | 1 | 7y1 |
| .......................cguaAguagcagagcagcucccucgc................................................................ | 1 | 1 | 7y1 |
| .......................cguacguagcagUgcagcucccucgc................................................................ | 1 | 1 | 7y1 |
| .......................cguacCuagcagagcagcucccucgc................................................................ | 1 | 1 | 7y1 |
| .......................cguacguagcagagcagcucccucCc................................................................ | 1 | 1 | 7y1 |
| .......................cguacguagcagagcagAucccucgc................................................................ | 1 | 1 | 7y1 |
| .......................cguacguagcagagcagcucccucgc................................................................ | 264 | 0 | 7y1 |
| .......................cguacguagcagagAagcucccucgc................................................................ | 1 | 1 | 7y1 |
| .......................cguacgAagcagagcagcucccucgc................................................................ | 1 | 1 | 7y1 |
| .......................cguacguagcagagcagAucccucgcu............................................................... | 1 | 1 | 7y1 |
| .......................cguacguagcagagcagcucccGcgcu............................................................... | 1 | 1 | 7y1 |
| .......................cguacguagcaUagcagcucccucgcu............................................................... | 1 | 1 | 7y1 |
| .......................cguacguagAagagcagcucccucgcu............................................................... | 1 | 1 | 7y1 |
| .......................cUuacguagcagagcagcucccucgcu............................................................... | 3 | 1 | 7y1 |
| .......................cguacguagcagagcagcucccucgcu............................................................... | 407 | 0 | 7y1 |
| .......................Aguacguagcagagcagcucccucgcu............................................................... | 2 | 1 | 7y1 |
| .......................cguacgAagcagagcagcucccucgcu............................................................... | 2 | 1 | 7y1 |
| .......................cguaUguagcagagcagcucccucgcu............................................................... | 264 | 1 | 7y1 |
| .......................cguaAguagcagagcagcucccucgcu............................................................... | 2 | 1 | 7y1 |
| .......................cCuacguagcagagcagcucccucgcu............................................................... | 1 | 1 | 7y1 |
| .......................cguacguagcagagcagcucccucgcG............................................................... | 3 | 1 | 7y1 |
| .......................cguacguagcagagcagcuGccucgcu............................................................... | 1 | 1 | 7y1 |
| .......................cAuacguagcagagcagcucccucgcug.............................................................. | 2 | 1 | 7y1 |
| .......................cguacguagcagagcagcucAcucgcug.............................................................. | 1 | 1 | 7y1 |
| .......................cguacguagcagagcagcucccucgcuC.............................................................. | 1 | 1 | 7y1 |
| .......................cguacguaCcagagcagcucccucgcug.............................................................. | 2 | 1 | 7y1 |
| .......................Aguacguagcagagcagcucccucgcug.............................................................. | 7 | 1 | 7y1 |
| .......................cguacguagcagagcagcucccucUcug.............................................................. | 2 | 1 | 7y1 |
| .......................cguacguagcagagcagcucccucgcGg.............................................................. | 8 | 1 | 7y1 |
| .......................cCuacguagcagagcagcucccucgcug.............................................................. | 1 | 1 | 7y1 |
| .......................cUuacguagcagagcagcucccucgcug.............................................................. | 2 | 1 | 7y1 |
| .......................Nguacguagcagagcagcucccucgcug.............................................................. | 2 | 1 | 7y1 |
| .......................cguacguagcagagcagcuAccucgcug.............................................................. | 2 | 1 | 7y1 |
| .......................cguacguagcGgagcagcucccucgcug.............................................................. | 1 | 1 | 7y1 |
| .......................Gguacguagcagagcagcucccucgcug.............................................................. | 1 | 1 | 7y1 |
| .......................cgCacguagcagagcagcucccucgcug.............................................................. | 1 | 1 | 7y1 |
| .......................cguacguagcagagcagcucccucgcuA.............................................................. | 1 | 1 | 7y1 |
| .......................cguacguagcagagcagcucccucgcug.............................................................. | 1992 | 0 | 7y1 |
| .......................cguaAguagcagagcagcucccucgcug.............................................................. | 1 | 1 | 7y1 |
| .......................cguacguagcagagcagcuccAucgcug.............................................................. | 1 | 1 | 7y1 |
| .......................cguacguagcagCgcagcucccucgcug.............................................................. | 1 | 1 | 7y1 |
| .......................cguacgAagcagagcagcucccucgcug.............................................................. | 2 | 1 | 7y1 |
| .......................cguacUuagcagagcagcucccucgcug.............................................................. | 1 | 1 | 7y1 |
| .......................cguacguagcagagAagcucccucgcug.............................................................. | 2 | 1 | 7y1 |
| .......................cguacguagcagagcagcucccucgcuU.............................................................. | 1 | 1 | 7y1 |
| .......................cguacCuagcagagcagcucccucgcug.............................................................. | 2 | 1 | 7y1 |
| .......................cguacguagcagagcagcuGccucgcug.............................................................. | 1 | 1 | 7y1 |
| .......................cgGacguagcagagcagcucccucgcug.............................................................. | 1 | 1 | 7y1 |
| .......................cguacguagcagagcagcucccucgcAg.............................................................. | 4 | 1 | 7y1 |
| .......................cgAacguagcagagcagcucccucgcug.............................................................. | 2 | 1 | 7y1 |
| .......................cguacguagcagagcagcucGcucgcug.............................................................. | 1 | 1 | 7y1 |
| .......................cguacguagcagagcagcucccuAgcug.............................................................. | 1 | 1 | 7y1 |
| .......................cguacguagcaUagcagcucccucgcug.............................................................. | 1 | 1 | 7y1 |

**Star** **Mature**

```
gaccugcuucugggucggggguucguacguagcagagcagcucccucgcugcgaucuauugaaagucagcccucgacacaaggguuuguccgcgcgcgcgcgcgcgcgcgcgugc
.......................cguacguagAagagcagcucccucgcug................................................................ 3 1 7y1
.......................cguacguagcagagcagcucccucgAug................................................................ 2 1 7y1
........................gAacguagcagagcagcu......................................................................... 2 1 7y1
........................guaAguagcagagcagcu......................................................................... 1 1 7y1
........................guacguagcagagcagcu......................................................................... 154 0 7y1
........................Cuacguagcagagcagcuc........................................................................ 2 1 7y1
........................guacguagcagagcagAuc........................................................................ 1 1 7y1
........................guacguagcagagcagcuc........................................................................ 173 0 7y1
........................guacgAagcagagcagcuc........................................................................ 1 1 7y1
........................Cuacguagcagagcagcucccucgc.................................................................. 1 1 7y1
........................guacguagcagagcagcucccucgcu................................................................. 164 0 7y1
........................guacguagcagagcagAucccucgcu................................................................. 2 1 7y1
........................guacguagcagagAagcucccucgcu................................................................. 1 1 7y1
........................Uuacguagcagagcagcucccucgcu................................................................. 1 1 7y1
........................guacguagcagagcagcucccucgcG................................................................. 3 1 7y1
........................guacgAagcagagcagcucccucgcu................................................................. 1 1 7y1
........................guacguaUcagagcagcucccucgcu................................................................. 1 1 7y1
........................guGcguagcagagcagcucccucgcu................................................................. 1 1 7y1
........................guacguagcagagcagGucccucgcu................................................................. 1 1 7y1
........................guacguagcagagcGgcucccucgcug................................................................ 1 1 7y1
........................guaAguagcagagcagcucccucgcug................................................................ 1 1 7y1
........................guacguagcagagcagcucccucgcug................................................................ 61 0 7y1
.........................uacguagcagagcagcuA........................................................................ 2 1 7y1
.........................uacguagcagagcaAcuc........................................................................ 1 1 7y1
.........................uacguagcagagAagcuc........................................................................ 1 1 7y1
.........................uacguagcagagcagcuc........................................................................ 616 0 7y1
.........................uacguaUcagagcagcuc........................................................................ 2 1 7y1
.........................Aacguagcagagcagcuc........................................................................ 15 1 7y1
.........................uaAguagcagagcagcuc........................................................................ 1 1 7y1
.........................uacguagcagagGagcuc........................................................................ 1 1 7y1
.........................uacguagcagagcagcAc........................................................................ 2 1 7y1
.........................uacguagcagagcCgcuc........................................................................ 1 1 7y1
.........................uacguagcagagcagcuG........................................................................ 1 1 7y1
.........................Gacguagcagagcagcuccc...................................................................... 1 1 7y1
.........................uacguagcGgagcagcuccc...................................................................... 1 1 7y1
.........................uacguagcagCgcagcuccc...................................................................... 1 1 7y1
.........................uacguagcagagcagcAccc...................................................................... 1 1 7y1
.........................Aacguagcagagcagcuccc...................................................................... 3 1 7y1
.........................uacguagcagagcagcuccc...................................................................... 193 0 7y1
.........................uacguagcagagcaCcucccucgcu................................................................. 1 1 7y1
.........................uaAguagcagagcagcucccucgcu................................................................. 1 1 7y1
.........................uacguagcagagcUgcucccucgcu................................................................. 1 1 7y1
.........................uacguagcagagcagcucccucgcu................................................................. 493 0 7y1
.........................Nacguagcagagcagcucccucgcu................................................................. 1 1 7y1
.........................uacguagcagagcagcuAccucgcu................................................................. 1 1 7y1
.........................uacguagcagagAagcucccucgcu................................................................. 2 1 7y1
.........................Aacguagcagagcagcucccucgcu................................................................. 5 1 7y1
.........................uacguagcagagcagcucccucUcu................................................................. 1 1 7y1
.........................uacguagcagagcagcucccucgcG................................................................. 4 1 7y1
.........................uacguagcaCagcagcucccucgcu................................................................. 2 1 7y1
.........................uacguagcagagcagcucccuUgcu................................................................. 1 1 7y1
.........................uacguaCcagagcagcucccucgcu................................................................. 2 1 7y1
.........................uacguagcagagcagcucccucgGu................................................................. 1 1 7y1
.........................uacguagcagagcaAcucccucgcug................................................................ 1 1 7y1
.........................uGcguagcagagcagcucccucgcug................................................................ 1 1 7y1
.........................uacguagcagagcagcucccAcgcug................................................................ 1 1 7y1
.........................uacguagcagagcagcucAcucgcug................................................................ 4 1 7y1
.........................uacguagcagagcagcucccucgcuA................................................................ 1 1 7y1
.........................uacguagcagagcagcucccucgcuC................................................................ 2 1 7y1
.........................uacguGgcagagcagcucccucgcug................................................................ 1 1 7y1
.........................uacgAagcagagcagcucccucgcug................................................................ 2 1 7y1
.........................uacguagcagagcagcAcccucgcug................................................................ 1 1 7y1
.........................uacguagcagagcagcucccuAgcug................................................................ 1 1 7y1
.........................uacguagcagagcagcuccGucgcug................................................................ 1 1 7y1
.........................uacguagcagagcaUcucccucgcug................................................................ 2 1 7y1
.........................uacguagcagagcagcucGcucgcug................................................................ 1 1 7y1
.........................uacguagcagagcagcuAccucgcug................................................................ 2 1 7y1
.........................uacguagcUgagcagcucccucgcug................................................................ 1 1 7y1
.........................uacguagcagagcagcucccucgcGg................................................................ 19 1 7y1
.........................uacguagcagagcaNcucccucgcug................................................................ 1 1 7y1
```

**Star** **Mature**

```
gaccugcuucugggucgggguuucguacguagcagagcagcucccucgcugcgaucuauugaaagucagcccucgacacaaggguuuguccgcgcgcgcgcgcgcgcgcgcgugc
.........................uacguagcagagcagcucccucgcAg................................................................ 1 1 7y1
.........................uacguagcagagcagcucccucgAug................................................................ 2 1 7y1
.........................Aacguagcagagcagcucccucgcug................................................................ 17 1 7y1
.........................uacguUgcagagcagcucccucgcug................................................................ 1 1 7y1
.........................uacguagcagagcagcucccucgcCg................................................................ 1 1 7y1
.........................uUcguagcagagcagcucccucgcug................................................................ 1 1 7y1
.........................uacguagcagagcagcucccucgcug................................................................ 1768 0 7y1
.........................uacguaCcagagcagcucccucgcug................................................................ 1 1 7y1
.........................uacguagAagagcagcucccucgcug................................................................ 3 1 7y1
.........................uaAguagcagagcagcucccucgcug................................................................ 5 1 7y1
.........................uacguagcagagcCgcucccucgcug................................................................ 1 1 7y1
.........................uaGguagcagagcagcucccucgcug................................................................ 3 1 7y1
.........................uacguagcagagcagcucccucgcuU................................................................ 1 1 7y1
.........................Gacguagcagagcagcucccucgcug................................................................ 2 1 7y1
.........................uacguaUcagagcagcucccucgcug................................................................ 1 1 7y1
..........................acguagcagaUcagcucccucgcu................................................................. 1 1 7y1
..........................acguagcagagcagcucccucgcu................................................................. 416 0 7y1
..........................acguagcagagcagcucccucgcG................................................................. 9 1 7y1
..........................Gcguagcagagcagcucccucgcu................................................................. 5 1 7y1
..........................acguagcagagcagAucccucgcu................................................................. 1 1 7y1
..........................acguagcGgagcagcucccucgcu................................................................. 2 1 7y1
..........................acguagcagagcagcucccAcgcu................................................................. 1 1 7y1
..........................acguagcagagcaUcucccucgcu................................................................. 3 1 7y1
..........................acguaCcagagcagcucccucgcu................................................................. 1 1 7y1
..........................acgAagcagagcagcucccucgcu................................................................. 1 1 7y1
..........................acguaUcagagcagcucccucgcu................................................................. 1 1 7y1
..........................acguagcagagAagcucccucgcu................................................................. 1 1 7y1
..........................aAguagcagagcagcucccucgcu................................................................. 2 1 7y1
..........................acguagcagagcagcuAccucgcu................................................................. 1 1 7y1
..........................acguagAagagcagcucccucgcu................................................................. 2 1 7y1
..........................acguagcagagcCgcucccucgcu................................................................. 1 1 7y1
..........................acguagcaUagcagcucccucgcug................................................................ 1 1 7y1
..........................acguagAagagcagcucccucgcug................................................................ 2 1 7y1
..........................acguagcagaUcagcucccucgcug................................................................ 2 1 7y1
..........................acguagcagagAagcucccucgcug................................................................ 1 1 7y1
..........................acguagcagagcagcucccucgcGg................................................................ 5 1 7y1
..........................acguagcagagcGgcucccucgcug................................................................ 1 1 7y1
..........................acguagcagagcagcucccucgcug................................................................ 537 0 7y1
..........................acgAagcagagcagcucccucgcug................................................................ 1 1 7y1
..........................acCuagcagagcagcucccucgcug................................................................ 1 1 7y1
..........................acguagcagagcagcucccucgcuC................................................................ 1 1 7y1
..........................acguagcagagcagcuccAucgcug................................................................ 2 1 7y1
..........................Gcguagcagagcagcucccucgcug................................................................ 1 1 7y1
...........................cguagcagagcagcuccc...................................................................... 199 0 7y1
...........................cguagcagagcagcucGc...................................................................... 1 1 7y1
...........................cguagcagagcagcucAcu..................................................................... 1 1 7y1
...........................cguagAagagcagcucccu..................................................................... 3 1 7y1
...........................cguagcagagcagcuAccu..................................................................... 2 1 7y1
...........................cguagcagaUcagcucccu..................................................................... 1 1 7y1
...........................cgGagcagagcagcucccu..................................................................... 1 1 7y1
...........................cguagcagagcagcucccu..................................................................... 204 0 7y1
...........................cguagcagaCcagcucccu..................................................................... 1 1 7y1
...........................cguaCcagagcagcucccucgcug................................................................ 1 1 7y1
...........................cCuagcagagcagcucccucgcug................................................................ 1 1 7y1
...........................cguagcagagcagcuUccucgcug................................................................ 1 1 7y1
...........................cguagcagagcagcucccucAcug................................................................ 1 1 7y1
...........................cguagcagCgcagcucccucgcug................................................................ 1 1 7y1
...........................cguagcagagcagcucccucgAug................................................................ 1 1 7y1
...........................cguagcagagcagcucccucgcuU................................................................ 2 1 7y1
...........................cguagcagagcagcucccucgcug................................................................ 541 0 7y1
...........................cguaUcagagcagcucccucgcug................................................................ 1 1 7y1
...........................cguagcagagcagAucccucgcug................................................................ 1 1 7y1
...........................Nguagcagagcagcucccucgcug................................................................ 2 1 7y1
...........................cguagcagagcagcucccAcgcug................................................................ 1 1 7y1
...........................cgAagcagagcagcucccucgcug................................................................ 1 1 7y1
...........................Gguagcagagcagcucccucgcug................................................................ 1 1 7y1
...........................cguagcagagcagcuGccucgcug................................................................ 1 1 7y1
...........................cguagcagagcagcAcccucgcug................................................................ 1 1 7y1
............................guagcagagcagcucccucgc.................................................................. 2 0 7y1
............................guagcagagcagcucccucgcug................................................................ 73 0 7y1
```

Star                                                                    Mature

```
gaccugcuucugggucgggguuucguacguagcagagcagcucccucgcugcgaucuauugaaagucagcccucgacacaaggguuuguccgcgcgcgcgcgcgcgcgcgcgugc
..............................agcagagcagcucGcucgcug................................................................   1     1    7y1
..............................aUcagagcagcucccucgcug................................................................   1     1    7y1
..............................agcagagcagcuccAucgcug................................................................   2     1    7y1
..............................agcagagcagcucccucgcug................................................................   273   0    7y1
..............................agcagagAagcucccucgcug................................................................   1     1    7y1
..............................agcagagcagcuAccucgcug................................................................   1     1    7y1
..............................aAcagagcagcucccucgcug................................................................   1     1    7y1
..............................agcagagcagcucAcucgcug................................................................   1     1    7y1
..............................agcagagcagcucccucgAug................................................................   1     1    7y1
................................cagagcaUcucccucgcu.................................................................   1     1    7y1
................................cagagcagcucccucgcu.................................................................   129   0    7y1
................................cagCgcagcucccucgcu.................................................................   1     1    7y1
................................cagagcagcucAcucgcu.................................................................   1     1    7y1
................................cagagcagcucccucgcG.................................................................   11    1    7y1
................................cagagcagcuccGucgcu.................................................................   1     1    7y1
................................cagagcagcAcccucgcu.................................................................   1     1    7y1
....................................gcagcuUccucgcugcgaucu..........................................................   1     1    7y1
....................................gcagcucAcucgcugcgaucu..........................................................   1     1    7y1
....................................gcagcuAccucgcugcgaucu..........................................................   3     1    7y1
....................................gAagcucccucgcugcgaucu..........................................................   1     1    7y1
....................................gcagcucccucgcuCcgaucu..........................................................   1     1    7y1
....................................gcagcucccucgcugcgaucu..........................................................   394   0    7y1
....................................gcagcucccucCcugcgaucu..........................................................   1     1    7y1
....................................gcagcucccucgAugcgaucu..........................................................   1     1    7y1
....................................gcagcucccucgcugcgaucA..........................................................   1     1    7y1
....................................gcagcucccucgcAgcgaucu..........................................................   2     1    7y1
....................................gcagcucccucgcugcgGucu..........................................................   1     1    7y1
....................................gcagAucccucgcugcgaucu..........................................................   1     1    7y1
....................................gcagcucccucgGugcgaucu..........................................................   1     1    7y1
....................................gcagcucccucgcCgcgaucu..........................................................   1     1    7y1
....................................gcagcucccucgcugcgaucuauugaaagu.................................................   224   0    7y1
....................................Ccagcucccucgcugcgaucuauugaaagu.................................................   1     1    7y1
....................................gcagcucccucgcugcgaucuauugaaagG.................................................   1     1    7y1
....................................gcagcucccuAgcugcgaucuauugaaagu.................................................   1     1    7y1
....................................gcagcucccucUcugcgaucuauugaaagu.................................................   1     1    7y1
....................................gcagcucccucgcugcgaucuaAugaaagu.................................................   1     1    7y1
....................................gcagcuUccucgcugcgaucuauugaaagu.................................................   1     1    7y1
....................................Ncagcucccucgcugcgaucuauugaaaguca...............................................   1     1    7y1
....................................gcagcucccucgcugcgaucuauugaaaguca...............................................   182   0    7y1
....................................gcagcucccucgcugcgaucAauugaaaguca...............................................   1     1    7y1
....................................gcagcucAcucgcugcgaucuauugaaaguca...............................................   2     1    7y1
....................................gcagcucccuAgcugcgaucuauugaaaguca...............................................   1     1    7y1
....................................gcagcucccucgcugcgauAuauugaaaguca...............................................   1     1    7y1
....................................gcagcucccucgcugcgaucuauAgaaaguca...............................................   1     1    7y1
....................................gcagGucccucgcugcgaucuauugaaaguca...............................................   1     1    7y1
....................................gcagcucccuUgcugcgaucuauugaaaguca...............................................   1     1    7y1
....................................gcagcucccucgcugAgaucuauugaaaguca...............................................   2     1    7y1
....................................gcagcuAccucgcugcgaucuauugaaaguca...............................................   3     1    7y1
.....................................cagcucccucgcugcgaucAauugaaagucag..............................................   1     1    7y1
.....................................cagcucccucgcugcgaucuaGugaaagucag..............................................   1     1    7y1
.....................................cagcucccucgcugcgaucuauugaaagucag..............................................   188   0    7y1
.....................................cagcucccucgcugcgaAcuauugaaagucag..............................................   1     1    7y1
.....................................cagcAcccucgcugcgaucuauugaaagucag..............................................   2     1    7y1
.....................................cagcucccucgcugcgaucuauugaaagAcag..............................................   1     1    7y1
.....................................cagcuAccucgcugcgaucuauugaaagucag..............................................   1     1    7y1
.....................................cagcucccucgcugAgaucuauugaaagucag..............................................   1     1    7y1
.....................................cagUucccucgcugcgaucuauugaaagucag..............................................   1     1    7y1
.....................................cagcucccuAgcugcgaucuauugaaagucag..............................................   1     1    7y1
.....................................cagcucccucAcugcgaucuauugaaagucag..............................................   1     1    7y1
.....................................cagcuccAucgcugcgaucuauugaaagucag..............................................   2     1    7y1
.....................................Aagcucccucgcugcgaucuauugaaagucag..............................................   2     1    7y1
.....................................cagcuUccucgcugcgaucuauugaaagucag..............................................   1     1    7y1
......................................agcucccucgcugcNaucu..........................................................   1     1    7y1
......................................agcucccucgcugcgUucu..........................................................   1     1    7y1
......................................agcucccucgcugcgaucG..........................................................   1     1    7y1
......................................agcuccAucgcugcgaucu..........................................................   1     1    7y1
......................................agcucccucgcugcgaucu..........................................................   330   0    7y1
......................................agcuAccucgcugcgaucu..........................................................   1     1    7y1
......................................agAucccucgcugcgaucu..........................................................   1     1    7y1
......................................agcucccuAgcugcgaucu..........................................................   1     1    7y1
```

**Star** **Mature**

| | | | |
|---|---|---|---|
| gaccugcuucugggucgggguuucguacguagcagagcagcucccucgcugcgaucuauugaaagucagcccucgacacaaggguuuguccgcgcgcgcgcgcgcgcgcgcgugc | | | |
| ......................................agcucccucgcugAgaucu.......................................................... | 1 | 1 | 7y1 |
| ......................................Cgcucccucgcugcgaucu.......................................................... | 2 | 1 | 7y1 |
| ......................................agcucAcucgcugcgaucu.......................................................... | 1 | 1 | 7y1 |
| ......................................Cgcucccucgcugcgaucuauugaaaguc................................................ | 1 | 1 | 7y1 |
| ......................................agcucccucgcAgcgaucuauugaaaguc................................................ | 1 | 1 | 7y1 |
| ......................................agcucccucgcugcgaucuauugaaaguc................................................ | 185 | 0 | 7y1 |
| ......................................agcucAcucgcugcgaucuauugaaaguc................................................ | 2 | 1 | 7y1 |
| ......................................agcCcccucgcugcgaucuauugaaaguc................................................ | 1 | 1 | 7y1 |
| ......................................agcuAccucgcugcgaucuauugaaaguc................................................ | 2 | 1 | 7y1 |
| ......................................agcuccAucgcugcgaucuauugaaaguc................................................ | 1 | 1 | 7y1 |
| ......................................agcucccucgcugAgaucuauugaaaguc................................................ | 1 | 1 | 7y1 |
| ......................................agcucccucgcugcgaucuauugaaagucag.............................................. | 1 | 0 | 7y1 |
| .......................................gcucccucgcugcgaucG.......................................................... | 1 | 1 | 7y1 |
| .......................................Ccucccucgcugcgaucu.......................................................... | 1 | 1 | 7y1 |
| .......................................gcucccucgcugcgaucu.......................................................... | 194 | 0 | 7y1 |
| .......................................gcucccAcgcugcgaucu.......................................................... | 1 | 1 | 7y1 |
| .......................................gcucccucUcugcgaucu.......................................................... | 1 | 1 | 7y1 |
| .......................................gcucccucgAugcgaucu.......................................................... | 1 | 1 | 7y1 |
| .......................................gcuAccucgcugcgaucu.......................................................... | 1 | 1 | 7y1 |
| .......................................gcucAcucgcugcgaucuauug...................................................... | 3 | 1 | 7y1 |
| .......................................gcuAccucgcugcgaucuauug...................................................... | 1 | 1 | 7y1 |
| .......................................gcuccAucgcugcgaucuauug...................................................... | 2 | 1 | 7y1 |
| .......................................Ccucccucgcugcgaucuauug...................................................... | 1 | 1 | 7y1 |
| .......................................gcucccucgcugcgaucuauAg...................................................... | 1 | 1 | 7y1 |
| .......................................gcucccucgcugcgaucuauug...................................................... | 515 | 0 | 7y1 |
| .......................................gcucccucgcugcgaucAauug...................................................... | 1 | 1 | 7y1 |
| .......................................gcucUcucgcugcgaucuauug...................................................... | 1 | 1 | 7y1 |
| .......................................gcucccuAgcugcgaucuauug...................................................... | 1 | 1 | 7y1 |
| .......................................gcucccucgcugcgaucuauCg...................................................... | 1 | 1 | 7y1 |
| .......................................gcucccucgcugcgaucuauuU...................................................... | 1 | 1 | 7y1 |
| .......................................gcucccucgcugcgUucuauug...................................................... | 1 | 1 | 7y1 |
| .......................................gcucccucgcugcgaucuauugaGag.................................................. | 1 | 1 | 7y1 |
| .......................................gcucccucCcugcgaucuauugaaag.................................................. | 1 | 1 | 7y1 |
| .......................................gcucccucgcugcgaucuUuugaaag.................................................. | 1 | 1 | 7y1 |
| .......................................gcucccucgcugcgaucuauugaaGg.................................................. | 1 | 1 | 7y1 |
| .......................................gcucAcucgcugcgaucuauugaaag.................................................. | 2 | 1 | 7y1 |
| .......................................gcucccucgcugcgaucuauGgaaag.................................................. | 1 | 1 | 7y1 |
| .......................................gcucccucgcugcgauAuauugaaag.................................................. | 1 | 1 | 7y1 |
| .......................................gcucccucgcugcUaucuauugaaag.................................................. | 1 | 1 | 7y1 |
| .......................................gcuUccucgcugcgaucuauugaaag.................................................. | 1 | 1 | 7y1 |
| .......................................gcucccucAcugcgaucuauugaaag.................................................. | 1 | 1 | 7y1 |
| .......................................gcucccucgcGgcgaucuauugaaag.................................................. | 1 | 1 | 7y1 |
| .......................................gcucccucgcugcgaucuauugGaag.................................................. | 1 | 1 | 7y1 |
| .......................................gcucccucgcuCcgaucuauugaaag.................................................. | 1 | 1 | 7y1 |
| .......................................gcuAccucgcugcgaucuauugaaag.................................................. | 2 | 1 | 7y1 |
| .......................................gcucccucgcugAgaucuauugaaag.................................................. | 1 | 1 | 7y1 |
| .......................................gNucccucgcugcgaucuauugaaag.................................................. | 1 | 1 | 7y1 |
| .......................................gcucccuAgcugcgaucuauugaaag.................................................. | 1 | 1 | 7y1 |
| .......................................gcucccucgcugcgaucuauugaaaC.................................................. | 2 | 1 | 7y1 |
| .......................................gcucccucgAugcgaucuauugaaag.................................................. | 1 | 1 | 7y1 |
| .......................................gcucccucgcugcgaucuauugaaag.................................................. | 1199 | 0 | 7y1 |
| .......................................Ncucccucgcugcgaucuauugaaag.................................................. | 1 | 1 | 7y1 |
| .......................................gcuccAucgcugcgaucuauugaaag.................................................. | 1 | 1 | 7y1 |
| .......................................gcucccucgcAgcgaucuauugaaag.................................................. | 1 | 1 | 7y1 |
| .......................................gcucccAcgcugcgaucuauugaaag.................................................. | 2 | 1 | 7y1 |
| .......................................Ccucccucgcugcgaucuauugaaag.................................................. | 2 | 1 | 7y1 |
| .......................................gAucccucgcugcgaucuauugaaag.................................................. | 3 | 1 | 7y1 |
| .......................................gcucccucgcugcgaAcuauugaaagu................................................. | 1 | 1 | 7y1 |
| .......................................gcucccucgcCgcgaucuauugaaagu................................................. | 1 | 1 | 7y1 |
| .......................................gcucccucgcAgcgaucuauugaaagu................................................. | 1 | 1 | 7y1 |
| .......................................gcucAcucgcugcgaucuauugaaagu................................................. | 2 | 1 | 7y1 |
| .......................................gcucccucgcugcgaucCauugaaagu................................................. | 1 | 1 | 7y1 |
| .......................................gcucccucgcuCcgaucuauugaaagu................................................. | 1 | 1 | 7y1 |
| .......................................gcucccucgcugcgaucAauugaaagu................................................. | 1 | 1 | 7y1 |
| .......................................gcuccAucgcugcgaucuauugaaagu................................................. | 1 | 1 | 7y1 |
| .......................................gcucccucgcugcgaucuauAgaaagu................................................. | 1 | 1 | 7y1 |
| .......................................gcucccucgcugcgaucuauugaaaUu................................................. | 1 | 1 | 7y1 |
| .......................................gcucccucgcugcgaucuauugaaagu................................................. | 906 | 0 | 7y1 |
| .......................................gcuAccucgcugcgaucuauugaaagu................................................. | 1 | 1 | 7y1 |
| .......................................gcucccucgcugcUaucuauugaaagu................................................. | 1 | 1 | 7y1 |

**Star** **Mature**

| | | | |
|---|---|---|---|
| gaccugcuucugggucgggguuucguacguagcagagcagcucccucgcugcgaucuauugaaagucagcccucgacacaaggguuuguccgcgcgcgcgcgcgcgcgcgugc | | | |
| .......................................gcucccucgcugcgauGuauugaaagu............................................... | 1 | 1 | 7y1 |
| .......................................gcucccuAgcugcgaucuauugaaagu............................................... | 2 | 1 | 7y1 |
| .......................................gcucccucgcugcgaucuauugaaagA............................................... | 1 | 1 | 7y1 |
| .......................................Ccucccucgcugcgaucuauugaaagu............................................... | 2 | 1 | 7y1 |
| .......................................Ucucccucgcugcgaucuauugaaagu............................................... | 1 | 1 | 7y1 |
| .......................................gcucccucgcugcgaucuaGugaaagu............................................... | 1 | 1 | 7y1 |
| .......................................gcucccucgcugcgaucuauugaaagG............................................... | 1 | 1 | 7y1 |
| .......................................gcucccucgAugcgaucuauugaaagu............................................... | 2 | 1 | 7y1 |
| .......................................gcucccucgcugcgaucuauugaaaguU.............................................. | 1 | 1 | 7y1 |
| .......................................gcuGccucgcugcgaucuauugaaaguc.............................................. | 1 | 1 | 7y1 |
| .......................................gcucccucgcugcgaucuauugaaaguc.............................................. | 229 | 0 | 7y1 |
| .......................................gcucccucgcugAgaucuauugaaaguc.............................................. | 1 | 1 | 7y1 |
| .......................................gcucAcucgcugcgaucuauugaaaguc.............................................. | 1 | 1 | 7y1 |
| .......................................gcuccAucgcugcgaucuauugaaaguc.............................................. | 1 | 1 | 7y1 |
| .......................................Ucucccucgcugcgaucuauugaaaguc.............................................. | 1 | 1 | 7y1 |
| ........................................cucccucgcugcgaucuaAugaa.................................................. | 1 | 1 | 7y1 |
| ........................................cucccAcgcugcgaucuauugaa.................................................. | 1 | 1 | 7y1 |
| ........................................cucccucgcugAgaucuauugaa.................................................. | 2 | 1 | 7y1 |
| ........................................cucccucgcugcgauAuauugaa.................................................. | 1 | 1 | 7y1 |
| ........................................cucccucgcAgcgaucuauugaa.................................................. | 1 | 1 | 7y1 |
| ........................................cucccucgcugcgaAcuauugaa.................................................. | 1 | 1 | 7y1 |
| ........................................cucccucgcugcCaucuauugaa.................................................. | 1 | 1 | 7y1 |
| ........................................cucccucgcugcgaucuauugaa.................................................. | 144 | 0 | 7y1 |
| ........................................cucccucgcugcgaucuauugaaa................................................. | 244 | 0 | 7y1 |
| ........................................cucccucgcugcgaucuauGgaaa................................................. | 1 | 1 | 7y1 |
| ........................................cucccucgcugAgaucuauugaaa................................................. | 1 | 1 | 7y1 |
| ........................................cuAccucgcugcgaucuauugaaa................................................. | 3 | 1 | 7y1 |
| ........................................Aucccucgcugcgaucuauugaaa................................................. | 2 | 1 | 7y1 |
| ........................................cucccucgcuCcgaucuauugaaa................................................. | 1 | 1 | 7y1 |
| ........................................cucccuAgcugcgaucuauugaaag................................................ | 1 | 1 | 7y1 |
| ........................................cucccucgcuCcgaucuauugaaag................................................ | 1 | 1 | 7y1 |
| ........................................cucccucgcuUcgaucuauugaaag................................................ | 1 | 1 | 7y1 |
| ........................................cuccAucgcugcgaucuauugaaag................................................ | 1 | 1 | 7y1 |
| ........................................cucAcucgcugcgaucuauugaaag................................................ | 1 | 1 | 7y1 |
| ........................................cucccucgcugcgaucuauugaaag................................................ | 185 | 0 | 7y1 |
| ........................................cucccucgcuAcgaucuauugaaagu............................................... | 1 | 1 | 7y1 |
| ........................................Aucccucgcugcgaucuauugaaagu............................................... | 6 | 1 | 7y1 |
| ........................................cucccucgAugcgaucuauugaaagu............................................... | 2 | 1 | 7y1 |
| ........................................cucAcucgcugcgaucuauugaaagu............................................... | 1 | 1 | 7y1 |
| ........................................cucccucgcugcgaucuauAgaaagu............................................... | 1 | 1 | 7y1 |
| ........................................cucccucgcuCcgaucuauugaaagu............................................... | 2 | 1 | 7y1 |
| ........................................cucccucCcugcgaucuauugaaagu............................................... | 2 | 1 | 7y1 |
| ........................................cucccucgcAgcgaucuauugaaagu............................................... | 2 | 1 | 7y1 |
| ........................................cucccucgcugcUaucuauugaaagu............................................... | 1 | 1 | 7y1 |
| ........................................cucccucgcugcgaucuauugaaaUu............................................... | 1 | 1 | 7y1 |
| ........................................cucccucgcugcgaucuauugaaagA............................................... | 5 | 1 | 7y1 |
| ........................................cucccucUcugcgaucuauugaaagu............................................... | 3 | 1 | 7y1 |
| ........................................cucccucgcugcgaucuaAugaaagu............................................... | 1 | 1 | 7y1 |
| ........................................cuccUucgcugcgaucuauugaaagu............................................... | 1 | 1 | 7y1 |
| ........................................cucccucgcugcgCucuauugaaagu............................................... | 1 | 1 | 7y1 |
| ........................................cucccucgcugcgaucuauugaaagG............................................... | 1 | 1 | 7y1 |
| ........................................cuAccucgcugcgaucuauugaaagu............................................... | 9 | 1 | 7y1 |
| ........................................cucccucgcugAgaucuauugaaagu............................................... | 2 | 1 | 7y1 |
| ........................................cucccuAgcugcgaucuauugaaagu............................................... | 4 | 1 | 7y1 |
| ........................................cucccucgcugcgaucuauugGaagu............................................... | 1 | 1 | 7y1 |
| ........................................cucccucgcugcgaucuauuAaaagu............................................... | 1 | 1 | 7y1 |
| ........................................cAcccucgcugcgaucuauugaaagu............................................... | 7 | 1 | 7y1 |
| ........................................cucccucgcugcgaucuauugaUagu............................................... | 1 | 1 | 7y1 |
| ........................................cucccucgcugcgaucAauugaaagu............................................... | 2 | 1 | 7y1 |
| ........................................cucccucgcugcgaucuauugaaagu............................................... | 1893 | 0 | 7y1 |
| ........................................cucccucgcugcgaucuauugaaaCu............................................... | 1 | 1 | 7y1 |
| ........................................cucccAcgcugcgaucuauugaaagu............................................... | 5 | 1 | 7y1 |
| ........................................cucccucgcugcgaucuauugaUaguc.............................................. | 1 | 1 | 7y1 |
| ........................................cucccucgcugcUaucuauugaaaguc.............................................. | 3 | 1 | 7y1 |
| ........................................cucccucgcugcgaucuauAgaaaguc.............................................. | 8 | 1 | 7y1 |
| ........................................cucccucgcugcgaucuauuCaaaguc.............................................. | 1 | 1 | 7y1 |
| ........................................cucccucgcuCcgaucuauugaaaguc.............................................. | 3 | 1 | 7y1 |
| ........................................cuAccucgcugcgaucuauugaaaguc.............................................. | 10 | 1 | 7y1 |
| ........................................cucccucgcugcgaAcuauugaaaguc.............................................. | 3 | 1 | 7y1 |
| ........................................Nucccucgcugcgaucuauugaaaguc.............................................. | 3 | 1 | 7y1 |

**Star** **Mature**

| | | | |
|---|---|---|---|
| gaccugcuucugggucgggguuucguacguagcagagcagcucccucgcugcgaucuauugaaagucagcccucgacacaaggguuuguccgcgcgcgcgcgcgcgcgcgcgugc | | | |
| ........................................cucccucgcugcgaucuauugaaaguA................................................ | 4 | 1 | 7y1 |
| ........................................cucccuUgcugcgaucuauugaaaguc................................................ | 1 | 1 | 7y1 |
| ........................................cuGccucgcugcgaucuauugaaaguc................................................ | 1 | 1 | 7y1 |
| ........................................cucUcucgcugcgaucuauugaaaguc................................................ | 2 | 1 | 7y1 |
| ........................................cucccucgcugAgaucuauugaaaguc................................................ | 12 | 1 | 7y1 |
| ........................................cucAcucgcugcgaucuauugaaaguc................................................ | 5 | 1 | 7y1 |
| ........................................cucccucgcugcgaucuauugGaaguc................................................ | 2 | 1 | 7y1 |
| ........................................cucccucgAugcgaucuauugaaaguc................................................ | 1 | 1 | 7y1 |
| ........................................cucccucgcuUcgaucuauugaaaguc................................................ | 6 | 1 | 7y1 |
| ........................................cucccucgcugcgaGcuauugaaaguc................................................ | 2 | 1 | 7y1 |
| ........................................cucccucUcugcgaucuauugaaaguc................................................ | 7 | 1 | 7y1 |
| ........................................cucccucgcugcgaucuaCugaaaguc................................................ | 1 | 1 | 7y1 |
| ........................................cucccucgcugcgaucuauugaaagAc................................................ | 3 | 1 | 7y1 |
| ........................................cucccucgcugcgaucuauugaaaAuc................................................ | 1 | 1 | 7y1 |
| ........................................cucccucgcCgcgaucuauugaaaguc................................................ | 1 | 1 | 7y1 |
| ........................................cAcccucgcugcgaucuauugaaaguc................................................ | 18 | 1 | 7y1 |
| ........................................cucccucgcugcgauAuauugaaaguc................................................ | 3 | 1 | 7y1 |
| ........................................cucccucgcugcgaucuaGugaaaguc................................................ | 1 | 1 | 7y1 |
| ........................................cucccucgcugcgaucuauuAaaaguc................................................ | 1 | 1 | 7y1 |
| ........................................cucccAcgcugcgaucuauugaaaguc................................................ | 9 | 1 | 7y1 |
| ........................................cucccucgGugcgaucuauugaaaguc................................................ | 1 | 1 | 7y1 |
| ........................................cucccucgcAgcgaucuauugaaaguc................................................ | 3 | 1 | 7y1 |
| ........................................cucccucgcugcgaucuaAugaaaguc................................................ | 7 | 1 | 7y1 |
| ........................................cucccucgcugcgaucuauugaaaCuc................................................ | 1 | 1 | 7y1 |
| ........................................cucccucgcugcgaucuauuUaaaguc................................................ | 2 | 1 | 7y1 |
| ........................................cucccucgcugcgaucuauugaaaguc................................................ | 4185 | 0 | 7y1 |
| ........................................cucccucgcuAcgaucuauugaaaguc................................................ | 4 | 1 | 7y1 |
| ........................................cucccucgcugcgaucuauugaaaguG................................................ | 1 | 1 | 7y1 |
| ........................................cucccucCcugcgaucuauugaaaguc................................................ | 4 | 1 | 7y1 |
| ........................................Aucccucgcugcgaucuauugaaaguc................................................ | 8 | 1 | 7y1 |
| ........................................cucccucgcugcgCucuauugaaaguc................................................ | 1 | 1 | 7y1 |
| ........................................cucccucgcugcgaucuGuugaaaguc................................................ | 1 | 1 | 7y1 |
| ........................................cucccucgcugcgaucuauugaaaUuc................................................ | 1 | 1 | 7y1 |
| ........................................cucccuAgcugcgaucuauugaaaguc................................................ | 6 | 1 | 7y1 |
| ........................................cuccAucgcugcgaucuauugaaaguc................................................ | 2 | 1 | 7y1 |
| ........................................cucccucgcugcgaucuauuCaaaguca............................................... | 1 | 1 | 7y1 |
| ........................................cucccucgcugcgaucuauAgaaaguca............................................... | 3 | 1 | 7y1 |
| ........................................cucccucgcugcgaucAauugaaaguca............................................... | 1 | 1 | 7y1 |
| ........................................cucccucgcAgcgaucuauugaaaguca............................................... | 5 | 1 | 7y1 |
| ........................................cucccAcgcugcgaucuauugaaaguca............................................... | 4 | 1 | 7y1 |
| ........................................cucccucgcugcgaucuauugaaaguca............................................... | 1280 | 0 | 7y1 |
| ........................................cucccucgcugcgaucuauugaGaguca............................................... | 1 | 1 | 7y1 |
| ........................................cAcccucgcugcgaucuauugaaaguca............................................... | 1 | 1 | 7y1 |
| ........................................cuAccucgcugcgaucuauugaaaguca............................................... | 4 | 1 | 7y1 |
| ........................................cucccuAgcugcgaucuauugaaaguca............................................... | 1 | 1 | 7y1 |
| ........................................cucccucgcugcgaucuauugaaagucG............................................... | 2 | 1 | 7y1 |
| ........................................cucAcucgcugcgaucuauugaaaguca............................................... | 1 | 1 | 7y1 |
| ........................................Aucccucgcugcgaucuauugaaaguca............................................... | 3 | 1 | 7y1 |
| ........................................cucccucgcugcgaucuauugaaaguGa............................................... | 1 | 1 | 7y1 |
| ........................................cucccucgcuCcgaucuauugaaaguca............................................... | 1 | 1 | 7y1 |
| ........................................cucccucgcugAgaucuauugaaaguca............................................... | 2 | 1 | 7y1 |
| ........................................cucccucgcugcgauAuauugaaaguca............................................... | 2 | 1 | 7y1 |
| ........................................cuAccucgcugcgaucuauugaaagucag.............................................. | 2 | 1 | 7y1 |
| ........................................cucccAcgcugcgaucuauugaaagucag.............................................. | 1 | 1 | 7y1 |
| ........................................cucccucgcAgcgaucuauugaaagucag.............................................. | 1 | 1 | 7y1 |
| ........................................Aucccucgcugcgaucuauugaaagucag.............................................. | 1 | 1 | 7y1 |
| ........................................cucccuAgcugcgaucuauugaaagucag.............................................. | 3 | 1 | 7y1 |
| ........................................cucAcucgcugcgaucuauugaaagucag.............................................. | 1 | 1 | 7y1 |
| ........................................cucccucgcugcgaucuauugaaaCucag.............................................. | 1 | 1 | 7y1 |
| ........................................Gucccucgcugcgaucuauugaaagucag.............................................. | 1 | 1 | 7y1 |
| ........................................cucccucgcugcgaucuauugaaagGcag.............................................. | 1 | 1 | 7y1 |
| ........................................cucccucgcugAgaucuauugaaagucag.............................................. | 4 | 1 | 7y1 |
| ........................................cucccucgcugcgaucuauugaaagucag.............................................. | 378 | 0 | 7y1 |
| ........................................cucccucgcugcgaucuauAgaaagucag.............................................. | 1 | 1 | 7y1 |
| ........................................cuccGucgcugcgaucuauugaaagucagcc............................................ | 1 | 1 | 7y1 |
| ........................................cucccucgAugcgaucuauugaaagucagcc............................................ | 1 | 1 | 7y1 |
| ........................................cucccucgcugcgaucuauugaaaguAagcc............................................ | 1 | 1 | 7y1 |
| ........................................Nucccucgcugcgaucuauugaaagucagcc............................................ | 1 | 1 | 7y1 |
| ........................................cuAccucgcugcgaucuauugaaagucagcc............................................ | 1 | 1 | 7y1 |
| ........................................cucccucgcugcgaucuauugaaagAcagcc............................................ | 1 | 1 | 7y1 |

**Star** **Mature**

```
gaccugcuucugggucgggguuucguacguagcagagcagcucccucgcugcgaucuauugaaagucagcccucgacacaaggguuuguccgcgcgcgcgcgcgcgcgcgugc
........................................cucccucgcugcgaucuauugaaagucagcc..........................................  172  0  7y1
.........................................ucccucgcugcUaucuau......................................................  1    1  7y1
.........................................ucccucgcugcgaAcuau......................................................  1    1  7y1
.........................................Acccucgcugcgaucuau......................................................  1    1  7y1
.........................................ucccucgcugcgaCcuau......................................................  1    1  7y1
.........................................ucccucgcugcgaucuau......................................................  200  0  7y1
.........................................Ncccucgcugcgaucuau......................................................  1    1  7y1
.........................................ucccucgcugcgaucuaG......................................................  1    1  7y1
.........................................uAccucgcugcgaucuauu.....................................................  1    1  7y1
.........................................ucccucgcugcgaucuauu.....................................................  189  0  7y1
.........................................uUccucgcugcgaucuauu.....................................................  1    1  7y1
.........................................Acccucgcugcgaucuauu.....................................................  1    1  7y1
.........................................ucAcucgcugcgaucuauu.....................................................  1    1  7y1
.........................................Gcccucgcugcgaucuauu.....................................................  1    1  7y1
.........................................ucccucgcugcUaucuauu.....................................................  2    1  7y1
.........................................ucccucgcugcgaAcuauug....................................................  1    1  7y1
.........................................ucccucgcugcUaucuauug....................................................  1    1  7y1
.........................................ucccucgcugcgaucuauug....................................................  266  0  7y1
.........................................Acccucgcugcgaucuauug....................................................  4    1  7y1
.........................................ucccucgcugGgaucuauug....................................................  1    1  7y1
.........................................ucccuAgcugcgaucuauug....................................................  1    1  7y1
.........................................Acccucgcugcgaucuauugaaa.................................................  2    1  7y1
.........................................ucccucgcugcgaucuauugaaa.................................................  212  0  7y1
.........................................ucccucgcugcgaucuauuUaaa.................................................  1    1  7y1
.........................................ucccucUcugcgaucuauugaaa.................................................  1    1  7y1
.........................................Gcccucgcugcgaucuauugaaa.................................................  1    1  7y1
.........................................Ccccucgcugcgaucuauugaaa.................................................  1    1  7y1
.........................................ucccuAgcugcgaucuauugaaag................................................  1    1  7y1
.........................................ucccucgcugcgauAuauugaaag................................................  1    1  7y1
.........................................ucccucgcugcgaucAauugaaag................................................  1    1  7y1
.........................................Acccucgcugcgaucuauugaaag................................................  7    1  7y1
.........................................ucccucgcugcgaucuauAgaaag................................................  1    1  7y1
.........................................ucccucgcugcgauUuauugaaag................................................  1    1  7y1
.........................................ucccucgcugcgaucuauuAaaag................................................  1    1  7y1
.........................................ucccucgcuUcgaucuauugaaag................................................  1    1  7y1
.........................................ucAcucgcugcgaucuauugaaag................................................  3    1  7y1
.........................................Ncccucgcugcgaucuauugaaag................................................  3    1  7y1
.........................................Gcccucgcugcgaucuauugaaag................................................  1    1  7y1
.........................................uAccucgcugcgaucuauugaaag................................................  2    1  7y1
.........................................ucccucgcugcgaucuauugCaag................................................  1    1  7y1
.........................................ucccucgcugcgaucuauugaaag................................................  935  0  7y1
.........................................ucccucgcugcgaucuauugGaag................................................  1    1  7y1
.........................................ucccucgcugAgaucuauugaaag................................................  1    1  7y1
.........................................ucccucUcugcgaucuauugaaag................................................  2    1  7y1
.........................................ucccucgcAgcgaucuauugaaag................................................  2    1  7y1
.........................................ucccucgcuCcgaucuauugaaag................................................  3    1  7y1
.........................................ucccucgcugcgaucuauugaaagu...............................................  682  0  7y1
.........................................ucccucgcAgcgaucuauugaaagu...............................................  2    1  7y1
.........................................ucccucgcugAgaucuauugaaagu...............................................  4    1  7y1
.........................................Acccucgcugcgaucuauugaaagu...............................................  12   1  7y1
.........................................ucccucUcugcgaucuauugaaagu...............................................  1    1  7y1
.........................................uccAucgcugcgaucuauugaaagu...............................................  1    1  7y1
.........................................ucAcucgcugcgaucuauugaaagu...............................................  1    1  7y1
.........................................ucccucgcugcgauAuauugaaagu...............................................  2    1  7y1
.........................................ucccucgcugcUaucuauugaaagu...............................................  1    1  7y1
.........................................ucccucgcugcgaucAauugaaagu...............................................  1    1  7y1
.........................................ucccucgcugcgaucuauugaaagA...............................................  1    1  7y1
.........................................Gcccucgcugcgaucuauugaaagu...............................................  1    1  7y1
.........................................uAccucgcugcgaucuauugaaagu...............................................  1    1  7y1
.........................................ucccucgcugcgaucuaAugaaagu...............................................  1    1  7y1
.........................................ucccucgcuUcgaucuauugaaagu...............................................  1    1  7y1
.........................................Gcccucgcugcgaucuauugaaaguc..............................................  2    1  7y1
.........................................ucccucgcugcgaucuauugaaaAuc..............................................  1    1  7y1
.........................................ucccucgcugcgaucuaAugaaaguc..............................................  1    1  7y1
.........................................ucccucgcuCcgaucuauugaaaguc..............................................  2    1  7y1
.........................................ucccucgcugcgaucuauugaaaCuc..............................................  3    1  7y1
.........................................ucccucgcugcgaucAauugaaaguc..............................................  3    1  7y1
.........................................ucccucgcAgcgaucuauugaaaguc..............................................  4    1  7y1
.........................................ucccucgcugcgaucuauAgaaaguc..............................................  2    1  7y1
.........................................ucAcucgcugcgaucuauugaaaguc..............................................  10   1  7y1
```

**Star** **Mature**

```
gaccugcuucugggucgggguuucguacguagcagagcagcucccucgcugcgaucuauugaaagucagcccucgacacaaggguuuguccgcgcgcgcgcgcgcgcgugc
.........................................ucccucgcugcgaucuauugGaaguc............................................ 1 1 7y1
.........................................uAccucgcugcgaucuauugaaaguc............................................ 3 1 7y1
.........................................ucccucgcugcUaucuauugaaaguc............................................ 1 1 7y1
.........................................ucccucgAugcgaucuauugaaaguc............................................ 3 1 7y1
.........................................ucccucgcugcgaucuauugaaaguc............................................ 2692 0 7y1
.........................................ucccuAgcugcgaucuauugaaaguc............................................ 7 1 7y1
.........................................ucccucgcugcgaucuauugaaaguA............................................ 1 1 7y1
.........................................ucccucgcGgcgaucuauugaaaguc............................................ 1 1 7y1
.........................................ucccucgcuUcgaucuauugaaaguc............................................ 2 1 7y1
.........................................ucccucAcugcgaucuauugaaaguc............................................ 1 1 7y1
.........................................ucccucgcugcgaucuGuugaaaguc............................................ 2 1 7y1
.........................................Acccucgcugcgaucuauugaaaguc............................................ 38 1 7y1
.........................................ucccucgcugAgaucuauugaaaguc............................................ 2 1 7y1
.........................................ucccAcgcugcgaucuauugaaaguc............................................ 1 1 7y1
.........................................ucccucgcugcCaucuauugaaaguc............................................ 3 1 7y1
.........................................ucccucgcugcgaucuauugaaagAc............................................ 2 1 7y1
.........................................ucccucUcugcgaucuauugaaaguc............................................ 2 1 7y1
.........................................ucccucgcugcgauAuauugaaaguc............................................ 4 1 7y1
.........................................uccAucgcugcgaucuauugaaaguc............................................ 2 1 7y1
.........................................Ncccucgcugcgaucuauugaaaguc............................................ 1 1 7y1
.........................................ucccucgGugcgaucuauugaaaguca........................................... 1 1 7y1
.........................................ucccucgcugcgaucuauugaaaguca........................................... 586 0 7y1
.........................................Ncccucgcugcgaucuauugaaaguca........................................... 2 1 7y1
.........................................ucccucgcugcgaucCauugaaaguca........................................... 1 1 7y1
.........................................ucccucgcugcgauAuauugaaaguca........................................... 1 1 7y1
.........................................Gcccucgcugcgaucuauugaaaguca........................................... 1 1 7y1
.........................................ucccucgcugAgaucuauugaaaguca........................................... 1 1 7y1
.........................................ucccucgcugcgaucuauugaaagucU........................................... 1 1 7y1
.........................................ucccucUcugcgaucuauugaaaguca........................................... 1 1 7y1
.........................................ucccucgcuAcgaucuauugaaaguca........................................... 1 1 7y1
.........................................ucccucgcuCcgaucuauugaaaguca........................................... 1 1 7y1
.........................................uccAucgcugcgaucuauugaaaguca........................................... 1 1 7y1
.........................................ucccucgcugcCaucuauugaaaguca........................................... 1 1 7y1
.........................................ucccucgAugcgaucuauugaaaguca........................................... 2 1 7y1
.........................................ucccucCcugcgaucuauugaaaguca........................................... 1 1 7y1
.........................................Acccucgcugcgaucuauugaaaguca........................................... 6 1 7y1
.........................................ucccucgcuUcgaucuauugaaagucag.......................................... 1 1 7y1
.........................................ucAcucgcugcgaucuauugaaagucag.......................................... 1 1 7y1
.........................................ucccucgcugcgaucuauAgaaagucag.......................................... 1 1 7y1
.........................................ucccucgcugcgaucuauugaaagucag.......................................... 159 0 7y1
.........................................Acccucgcugcgaucuauugaaagucag.......................................... 2 1 7y1
.........................................ucccucgcugcgUucuauugaaagucag.......................................... 1 1 7y1
.........................................ucccAcgcugcgaucuauugaaagucag.......................................... 1 1 7y1
..........................................cccucgcugcgaucuauu................................................... 172 0 7y1
..........................................cccucgcugcgaucuaAu................................................... 1 1 7y1
..........................................Gccucgcugcgaucuauu................................................... 1 1 7y1
..........................................ccAucgcugcgaucuauu................................................... 1 1 7y1
..........................................cAcucgcugcgaucuauu................................................... 1 1 7y1
..........................................cccucgcugcCaucuauu................................................... 2 1 7y1
..........................................cccucgcAgcgaucuauu................................................... 1 1 7y1
..........................................cccucgcugcgaucuauuga................................................. 157 0 7y1
..........................................Accucgcugcgaucuauuga................................................. 1 1 7y1
..........................................ccAucgcugcgaucuauuga................................................. 1 1 7y1
..........................................cccucgAugcgaucuauuga................................................. 1 1 7y1
..........................................cAcucgcugcgaucuauuga................................................. 2 1 7y1
..........................................cccucgcAgcgaucuauuga................................................. 1 1 7y1
..........................................cccucgcugcgaucuauugG................................................. 1 1 7y1
..........................................Nccucgcugcgaucuauuga................................................. 1 1 7y1
..........................................cccucgcugcgaucuauuAaa................................................ 1 1 7y1
..........................................cAcucgcugcgaucuauugaaa............................................... 3 1 7y1
..........................................cccucgAugcgaucuauugaaa............................................... 3 1 7y1
..........................................cccucgcugcUaucuauugaaa............................................... 1 1 7y1
..........................................cccucgcugcgaucuaAugaaa............................................... 3 1 7y1
..........................................cccucgcuCcgaucuauugaaa............................................... 2 1 7y1
..........................................cccucgcugcgaucuauAgaaa............................................... 1 1 7y1
..........................................cccucgcuUcgaucuauugaaa............................................... 2 1 7y1
..........................................cccucgcugcCaucuauugaaa............................................... 1 1 7y1
..........................................cUcucgcugcgaucuauugaaa............................................... 1 1 7y1
..........................................ccGucgcugcgaucuauugaaa............................................... 1 1 7y1
..........................................cccucgcugcgaucuauugaaa............................................... 1120 0 7y1
```

**Star** **Mature**

gaccugcuucugggucgggguuucguacguagcagagcagcucccucgcugcgaucuauugaaagucagcccucgacacaaggguuuguccgcgcgcgcgcgcgcgcgugc

| | | | |
|---|---|---|---|
| ..........................................cccucgcugcgaAcuauugaaa................................................. | 2 | 1 | 7y1 |
| ..........................................Acccucgcugcgaucuauugaaa................................................. | 3 | 1 | 7y1 |
| ..........................................cccucgcugAgaucuauugaaa................................................. | 3 | 1 | 7y1 |
| ..........................................cccucgcugcgauUuauugaaa................................................. | 1 | 1 | 7y1 |
| ..........................................cccucgcugcgaucuauuCaaa................................................. | 1 | 1 | 7y1 |
| ..........................................cccuAgcugcgaucuauugaaa................................................. | 1 | 1 | 7y1 |
| ..........................................cccAcgcugcgaucuauugaaa................................................. | 1 | 1 | 7y1 |
| ..........................................cccucgcugcgauAuauugaaa................................................. | 1 | 1 | 7y1 |
| ..........................................cccucgcugcgaucGauugaaa................................................. | 1 | 1 | 7y1 |
| ..........................................Ncccucgcugcgaucuauugaaa................................................. | 1 | 1 | 7y1 |
| ..........................................cccucgcCgcgaucuauugaaa................................................. | 1 | 1 | 7y1 |
| ..........................................cccucgcAgcgaucuauugaaa................................................. | 3 | 1 | 7y1 |
| ..........................................cccucgcugcgaucuaCugaaag................................................ | 1 | 1 | 7y1 |
| ..........................................cccucgcAgcgaucuauugaaag................................................ | 1 | 1 | 7y1 |
| ..........................................cccucgcugcgaucuauAgaaag................................................ | 1 | 1 | 7y1 |
| ..........................................cccucgcuUcgaucuauugaaag................................................ | 1 | 1 | 7y1 |
| ..........................................Gccucgcugcgaucuauugaaag................................................ | 1 | 1 | 7y1 |
| ..........................................cccAcgcugcgaucuauugaaag................................................ | 2 | 1 | 7y1 |
| ..........................................cAcucgcugcgaucuauugaaag................................................ | 6 | 1 | 7y1 |
| ..........................................cccucgcugcCaucuauugaaag................................................ | 3 | 1 | 7y1 |
| ..........................................cccucgcugcgauAuauugaaag................................................ | 1 | 1 | 7y1 |
| ..........................................cccucgcugAgaucuauugaaag................................................ | 2 | 1 | 7y1 |
| ..........................................ccUucgcugcgaucuauugaaag................................................ | 1 | 1 | 7y1 |
| ..........................................ccGucgcugcgaucuauugaaag................................................ | 1 | 1 | 7y1 |
| ..........................................cccucgcugcgaucuaAugaaag................................................ | 3 | 1 | 7y1 |
| ..........................................cccucCcugcgaucuauugaaag................................................ | 1 | 1 | 7y1 |
| ..........................................Ncccucgcugcgaucuauugaaag................................................ | 1 | 1 | 7y1 |
| ..........................................cccucgcugcgaucuauugaaag................................................ | 1154 | 0 | 7y1 |
| ..........................................cNcucgcugcgaucuauugaaag................................................ | 1 | 1 | 7y1 |
| ..........................................cccucgcugcgaucuauuAaaag................................................ | 1 | 1 | 7y1 |
| ..........................................Acccucgcugcgaucuauugaaag................................................ | 1 | 1 | 7y1 |
| ..........................................cccucgcugcgaucuauugUaag................................................ | 1 | 1 | 7y1 |
| ..........................................cccucgGugcgaucuauugaaag................................................ | 1 | 1 | 7y1 |
| ..........................................cccucgcugcgaucuauGgaaag................................................ | 1 | 1 | 7y1 |
| ..........................................ccAucgcugcgaucuauugaaag................................................ | 5 | 1 | 7y1 |
| ..........................................ccAucgcugcgaucuauugaaagu............................................... | 4 | 1 | 7y1 |
| ..........................................cccucgAugcgaucuauugaaagu............................................... | 5 | 1 | 7y1 |
| ..........................................cccucgcugcgaucuauuCaaagu............................................... | 1 | 1 | 7y1 |
| ..........................................cccucgcugcgaucCauugaaagu............................................... | 1 | 1 | 7y1 |
| ..........................................cccucgcugcgauAuauugaaagu............................................... | 2 | 1 | 7y1 |
| ..........................................cccucgcugcgaucAauugaaagu............................................... | 1 | 1 | 7y1 |
| ..........................................cAcucgcugcgaucuauugaaagu............................................... | 7 | 1 | 7y1 |
| ..........................................cccucgcugAgaucuauugaaagu............................................... | 6 | 1 | 7y1 |
| ..........................................cccucgcugcCaucuauugaaagu............................................... | 1 | 1 | 7y1 |
| ..........................................Acccucgcugcgaucuauugaaagu............................................... | 2 | 1 | 7y1 |
| ..........................................cccucgcugcgaucuauugaaagu............................................... | 1043 | 0 | 7y1 |
| ..........................................cccucUcugcgaucuauugaaagu............................................... | 2 | 1 | 7y1 |
| ..........................................cccucgcugcgaucuauugaaagG............................................... | 11 | 1 | 7y1 |
| ..........................................cccAcgcugcgaucuauugaaagu............................................... | 1 | 1 | 7y1 |
| ..........................................cccucgcuCcgaucuauugaaagu............................................... | 1 | 1 | 7y1 |
| ..........................................cccucgcugcgaucuauugaaagA............................................... | 2 | 1 | 7y1 |
| ..........................................cccucgcugcgaAcuauugaaagu............................................... | 1 | 1 | 7y1 |
| ..........................................cccucgcugcgaucuauAgaaagu............................................... | 1 | 1 | 7y1 |
| ..........................................Ncccucgcugcgaucuauugaaaguc.............................................. | 3 | 1 | 7y1 |
| ..........................................cccucgcugcgaucuauugaaaguA.............................................. | 1 | 1 | 7y1 |
| ..........................................cccucgcugcUaucuauugaaaguc.............................................. | 1 | 1 | 7y1 |
| ..........................................cccAcgcugcgaucuauugaaaguc.............................................. | 1 | 1 | 7y1 |
| ..........................................cccucgcuUcgaucuauugaaaguc.............................................. | 1 | 1 | 7y1 |
| ..........................................cccucgcugcgCucuauugaaaguc.............................................. | 1 | 1 | 7y1 |
| ..........................................cccucgcugcgaucuauugaGaguc.............................................. | 2 | 1 | 7y1 |
| ..........................................cccucgcugcgaucuaAugaaaguc.............................................. | 2 | 1 | 7y1 |
| ..........................................cccucgcugcgaucuauugaaaUuc.............................................. | 1 | 1 | 7y1 |
| ..........................................cccuAgcugcgaucuauugaaaguc.............................................. | 1 | 1 | 7y1 |
| ..........................................cccucgcugcgaucAauugaaaguc.............................................. | 3 | 1 | 7y1 |
| ..........................................cccucgcugcgaucuauugaaaCuc.............................................. | 1 | 1 | 7y1 |
| ..........................................cccucgcugAgaucuauugaaaguc.............................................. | 6 | 1 | 7y1 |
| ..........................................cNcucgcugcgaucuauugaaaguc.............................................. | 1 | 1 | 7y1 |
| ..........................................cccucgcCgcgaucuauugaaaguc.............................................. | 1 | 1 | 7y1 |
| ..........................................ccGucgcugcgaucuauugaaaguc.............................................. | 1 | 1 | 7y1 |
| ..........................................cccucgcugcgauAuauugaaaguc.............................................. | 2 | 1 | 7y1 |

Star Mature

```
gaccugcuucugggucgggguuucguacguagcagagcagcucccucgcugcgaucuauugaaagucagcccucgacacaaggguuuguccgcgcgcgcgcgcgcgcgcgcgugc
..........................................cccucgcugcgaAcuauugaaaguc................................................ 2 1 7y1
..........................................cccucgUugcgaucuauugaaaguc................................................ 1 1 7y1
..........................................cccucgcAgcgaucuauugaaaguc................................................ 1 1 7y1
..........................................cAcucgcugcgaucuauugaaaguc................................................ 7 1 7y1
..........................................Accucgcugcgaucuauugaaaguc................................................ 7 1 7y1
..........................................cccucgcuCcgaucuauugaaaguc................................................ 2 1 7y1
..........................................cccucUcugcgaucuauugaaaguc................................................ 1 1 7y1
..........................................cccucgcugcgaucuauugaaaguc................................................ 1855 0 7y1
..........................................cccucgcugcgaucuauugaaagAc................................................ 2 1 7y1
..........................................cccucgcugcgaucuauuUaaaguc................................................ 1 1 7y1
..........................................cccucgAugcgaucuauugaaaguc................................................ 8 1 7y1
..........................................ccAucgcugcgaucuauugaaaguc................................................ 8 1 7y1
..........................................cccucgcugcgaucuauAgaaaguc................................................ 1 1 7y1
..........................................Uccucgcugcgaucuauugaaaguc................................................ 2 1 7y1
..........................................cccucgcugcgaucuauuAaaaguc................................................ 1 1 7y1
..........................................cccucgcugcgaucuauCgaaaguc................................................ 1 1 7y1
..........................................cccucgcugGgaucuauugaaaguc................................................ 1 1 7y1
..........................................cccucgcugcgaucuauugaaaguAa............................................... 1 1 7y1
..........................................cccuAgcugcgaucuauugaaaguca............................................... 3 1 7y1
..........................................cccucgcugAgaucuauugaaaguca............................................... 2 1 7y1
..........................................ccAucgcugcgaucuauugaaaguca............................................... 2 1 7y1
..........................................cccuUgcugcgaucuauugaaaguca............................................... 2 1 7y1
..........................................Nccucgcugcgaucuauugaaaguca............................................... 3 1 7y1
..........................................cccucgcugcgaucuauugaaagAca............................................... 3 1 7y1
..........................................Accucgcugcgaucuauugaaaguca............................................... 5 1 7y1
..........................................cccucgcugcgauAuauugaaaguca............................................... 2 1 7y1
..........................................cccucUcugcgaucuauugaaaguca............................................... 2 1 7y1
..........................................cccucgcugcgaAcuauugaaaguca............................................... 1 1 7y1
..........................................cccucgcugcgaucuauugaaaguca............................................... 592 0 7y1
..........................................cccAcgcugcgaucuauugaaaguca............................................... 1 1 7y1
..........................................cccucAcugcgaucuauugaaaguca............................................... 1 1 7y1
..........................................cccucgcAgcgaucuauugaaaguca............................................... 2 1 7y1
..........................................cccucgcugcgaucuauugaaaUuca............................................... 1 1 7y1
..........................................cccucgcugcUaucuauugaaaguca............................................... 1 1 7y1
...........................................ccucgAugcgaucuauug...................................................... 1 1 7y1
...........................................cAucgcugcgaucuauug...................................................... 2 1 7y1
...........................................ccucgcCgcgaucuauug...................................................... 1 1 7y1
...........................................ccucgcugcgaucuauug...................................................... 209 0 7y1
...........................................ccucgcuUcgaucuauug...................................................... 1 1 7y1
...........................................ccucgcugcgaucuauuC...................................................... 1 1 7y1
...........................................ccucgcGgcgaucuauug...................................................... 1 1 7y1
...........................................Ncucgcugcgaucuauug...................................................... 2 1 7y1
...........................................ccucgcugcgaucGauugaa.................................................... 1 1 7y1
...........................................ccucgcugcUaucuauugaa.................................................... 2 1 7y1
...........................................ccucgcugcgaucuauAgaa.................................................... 1 1 7y1
...........................................ccucgcugcgaucuauugaa.................................................... 602 0 7y1
...........................................Ncucgcugcgaucuauugaa.................................................... 1 1 7y1
...........................................ccuAgcugcgaucuauugaa.................................................... 1 1 7y1
...........................................cAucgcugcgaucuauugaa.................................................... 3 1 7y1
...........................................ccAcgcugcgaucuauugaa.................................................... 1 1 7y1
...........................................ccucgcAgcgaucuauugaa.................................................... 2 1 7y1
...........................................ccucCcugcgaucuauugaa.................................................... 1 1 7y1
...........................................ccucgcugcgaucuauuCaa.................................................... 1 1 7y1
...........................................ccucgcugcgaucuauuAaa.................................................... 1 1 7y1
...........................................Acucgcugcgaucuauugaa.................................................... 1 1 7y1
...........................................ccucgcugcgauAuauugaaa................................................... 4 1 7y1
...........................................Acucgcugcgaucuauugaaa................................................... 6 1 7y1
...........................................ccuAgcugcgaucuauugaaa................................................... 1 1 7y1
...........................................ccucgAugcgaucuauugaaa................................................... 6 1 7y1
...........................................ccucgcugcgaucuaAugaaa................................................... 1 1 7y1
...........................................ccucgcAgcgaucuauugaaa................................................... 9 1 7y1
...........................................ccucgcugcUaucuauugaaa................................................... 5 1 7y1
...........................................ccucgcugAgaucuauugaaa................................................... 1 1 7y1
...........................................cAucgcugcgaucuauugaaa................................................... 2 1 7y1
...........................................Gcucgcugcgaucuauugaaa................................................... 1 1 7y1
...........................................ccucgcugcgaucuauugaGa................................................... 1 1 7y1
...........................................ccucgcuUcgaucuauugaaa................................................... 1 1 7y1
...........................................ccucgcugcgaucAauugaaa................................................... 2 1 7y1
...........................................ccucgcugcgaucuauugaaa................................................... 1604 0 7y1
...........................................ccucgcugcgaucuauugaUa................................................... 1 1 7y1
```

**Star** **Mature**

| | | | |
|---|---|---|---|
| gaccugcuucugggucggggguucguacguagcagagcagcucccucgcugcgaucuauugaaagucagcccucgacacaaggguuuguccgcgcgcgcgcgcgcgcgcgugc | | | |
| ...........................................ccucgcugcgUucuauugaaa................................................. | 1 | 1 | 7y1 |
| ...........................................ccucgcugAgaucuauugaaag................................................ | 5 | 1 | 7y1 |
| ...........................................ccucgcugcgaucuauCgaaag................................................ | 1 | 1 | 7y1 |
| ...........................................ccuGgcugcgaucuauugaaag................................................ | 1 | 1 | 7y1 |
| ...........................................ccucgcugcCaucuauugaaag................................................ | 2 | 1 | 7y1 |
| ...........................................Acucgcugcgaucuauugaaag................................................ | 2 | 1 | 7y1 |
| ...........................................ccucgcugcgaAcuauugaaag................................................ | 1 | 1 | 7y1 |
| ...........................................ccucgcugcgaucAauugaaag................................................ | 1 | 1 | 7y1 |
| ...........................................ccucgcuCcgaucuauugaaag................................................ | 1 | 1 | 7y1 |
| ...........................................cAucgcugcgaucuauugaaag................................................ | 4 | 1 | 7y1 |
| ...........................................ccucgcugcgaucuauugaaag................................................ | 1408 | 0 | 7y1 |
| ...........................................ccucgcugcgaucuauugaaaU................................................ | 1 | 1 | 7y1 |
| ...........................................ccucgcugcgaucuauugaaGg................................................ | 1 | 1 | 7y1 |
| ...........................................ccAcgcugcgaucuauugaaag................................................ | 2 | 1 | 7y1 |
| ...........................................ccuAgcugcgaucuauugaaag................................................ | 1 | 1 | 7y1 |
| ...........................................ccucUcugcgaucuauugaaag................................................ | 1 | 1 | 7y1 |
| ...........................................ccucgAugcgaucuauugaaag................................................ | 2 | 1 | 7y1 |
| ...........................................ccucgcugcgaucCauugaaag................................................ | 1 | 1 | 7y1 |
| ...........................................ccucgcugcgauAuauugaaag................................................ | 2 | 1 | 7y1 |
| ...........................................ccucgcugcgaucuaAugaaag................................................ | 1 | 1 | 7y1 |
| ...........................................ccucCcugcgaucuauugaaag................................................ | 2 | 1 | 7y1 |
| ...........................................ccucgcAgcgaucuauugaaag................................................ | 3 | 1 | 7y1 |
| ...........................................Ncucgcugcgaucuauugaaag................................................ | 1 | 1 | 7y1 |
| ...........................................Gcucgcugcgaucuauugaaag................................................ | 2 | 1 | 7y1 |
| ...........................................ccucgcuUcgaucuauugaaag................................................ | 1 | 1 | 7y1 |
| ...........................................ccucgcugcgaucuauAgaaag................................................ | 2 | 1 | 7y1 |
| ...........................................ccucgcuAcgaucuauugaaag................................................ | 1 | 1 | 7y1 |
| ...........................................ccucUcugcgaucuauugaaagu............................................... | 1 | 1 | 7y1 |
| ...........................................ccucgcugcgaucuauugaaagu............................................... | 1291 | 0 | 7y1 |
| ...........................................ccucgcugcgaucuauAgaaagu............................................... | 1 | 1 | 7y1 |
| ...........................................ccuAgcugcgaucuauugaaagu............................................... | 1 | 1 | 7y1 |
| ...........................................Gcucgcugcgaucuauugaaagu............................................... | 1 | 1 | 7y1 |
| ...........................................ccucgcugcgaucuauugaaaUu............................................... | 1 | 1 | 7y1 |
| ...........................................ccucgcuUcgaucuauugaaagu............................................... | 1 | 1 | 7y1 |
| ...........................................ccucgcugcgaucuauugaaagG............................................... | 1 | 1 | 7y1 |
| ...........................................ccucgcugcgaucuaAugaaagu............................................... | 1 | 1 | 7y1 |
| ...........................................ccucgcugcgaucAauugaaagu............................................... | 1 | 1 | 7y1 |
| ...........................................cUucgcugcgaucuauugaaagu............................................... | 1 | 1 | 7y1 |
| ...........................................ccucgcugcUaucuauugaaagu............................................... | 2 | 1 | 7y1 |
| ...........................................ccucgcCgcgaucuauugaaagu............................................... | 1 | 1 | 7y1 |
| ...........................................ccAcgcugcgaucuauugaaagu............................................... | 3 | 1 | 7y1 |
| ...........................................ccucgAugcgaucuauugaaagu............................................... | 2 | 1 | 7y1 |
| ...........................................ccucgcugcgaAcuauugaaagu............................................... | 1 | 1 | 7y1 |
| ...........................................ccucgcugAgaucuauugaaagu............................................... | 6 | 1 | 7y1 |
| ...........................................ccucgcuCcgaucuauugaaagu............................................... | 1 | 1 | 7y1 |
| ...........................................Ncucgcugcgaucuauugaaagu............................................... | 1 | 1 | 7y1 |
| ...........................................ccucgcugcCaucuauugaaagu............................................... | 1 | 1 | 7y1 |
| ...........................................ccucgcugcgauAuauugaaagu............................................... | 1 | 1 | 7y1 |
| ...........................................ccucgcugcgaucuauugaGagu............................................... | 1 | 1 | 7y1 |
| ...........................................Acucgcugcgaucuauugaaagu............................................... | 2 | 1 | 7y1 |
| ...........................................ccucgcAgcgaucuauugaaagu............................................... | 1 | 1 | 7y1 |
| ...........................................cAucgcugcgaucuauugaaagu............................................... | 2 | 1 | 7y1 |
| ...........................................ccucAcugcgaucuauugaaagu............................................... | 2 | 1 | 7y1 |
| ...........................................ccucgcugcgaucuauugaaagA............................................... | 3 | 1 | 7y1 |
| ...........................................ccucgcugcgauGuauugaaagu............................................... | 1 | 1 | 7y1 |
| ...........................................ccuGgcugcgaucuauugaaagu............................................... | 1 | 1 | 7y1 |
| ...........................................ccucgcugcgaucuauugaaagAc.............................................. | 4 | 1 | 7y1 |
| ...........................................ccGcgcugcgaucuauugaaaguc.............................................. | 1 | 1 | 7y1 |
| ...........................................ccucgcugcgaucuUuugaaaguc.............................................. | 1 | 1 | 7y1 |
| ...........................................ccuAgcugcgaucuauugaaaguc.............................................. | 4 | 1 | 7y1 |
| ...........................................ccucgcugcgaucuauAgaaaguc.............................................. | 4 | 1 | 7y1 |
| ...........................................ccucgcugcgaucuauuCaaaguc.............................................. | 1 | 1 | 7y1 |
| ...........................................ccucgcugAgaucuauugaaaguc.............................................. | 9 | 1 | 7y1 |
| ...........................................ccucgcugcUaucuauugaaaguc.............................................. | 2 | 1 | 7y1 |
| ...........................................ccucgcugcgaGcuauugaaaguc.............................................. | 1 | 1 | 7y1 |
| ...........................................ccucgcugcgaucuauCgaaaguc.............................................. | 3 | 1 | 7y1 |
| ...........................................ccucCcugcgaucuauugaaaguc.............................................. | 2 | 1 | 7y1 |
| ...........................................ccucgcugcgaucuauugaaaguc.............................................. | 4155 | 0 | 7y1 |
| ...........................................ccucgcuUcgaucuauugaaaguc.............................................. | 5 | 1 | 7y1 |
| ...........................................ccucgNugcgaucuauugaaaguc.............................................. | 1 | 1 | 7y1 |

**Star** **Mature**

```
gaccugcuucugggucgggguuucguacguagcagagcagcucccucgcugcgaucuauugaaagucagcccucgacacaaggguuuguccgcgcgcgcgcgcgcgcgcgcgugc
...........................................ccucgcugcgaucuaAugaaaguc................................................   3    1    7y1
...........................................ccAcgcugcgaucuauugaaaguc................................................   2    1    7y1
...........................................ccucgcugcgaucuauuUaaaguc................................................   1    1    7y1
...........................................ccucgcugcgaucCauugaaaguc................................................   1    1    7y1
...........................................ccucgcugcgauAuauugaaaguc................................................   7    1    7y1
...........................................ccucgcugcgUucuauugaaaguc................................................   1    1    7y1
...........................................ccucgcugcgaucuauugaGaguc................................................   1    1    7y1
...........................................ccucgcugcgaAcuauugaaaguc................................................   1    1    7y1
...........................................Gcucgcugcgaucuauugaaaguc................................................   2    1    7y1
...........................................ccucgcuCcgaucuauugaaaguc................................................   1    1    7y1
...........................................ccCcgcugcgaucuauugaaaguc................................................   1    1    7y1
...........................................Acucgcugcgaucuauugaaaguc................................................   7    1    7y1
...........................................ccucgcugcCaucuauugaaaguc................................................   4    1    7y1
...........................................ccucgcugcgaucuauugaaaCuc................................................   5    1    7y1
...........................................ccucgcugcgaucuauGgaaaguc................................................   1    1    7y1
...........................................ccucgcugcgaucuauugaaaUuc................................................   1    1    7y1
...........................................ccucgcugcgaucuauugGaaguc................................................   1    1    7y1
...........................................ccucgcAgcgaucuauugaaaguc................................................   16   1    7y1
...........................................ccucgcugcgaucAauugaaaguc................................................   3    1    7y1
...........................................ccucgcCgcgaucuauugaaaguc................................................   1    1    7y1
...........................................ccucgAugcgaucuauugaaaguc................................................   6    1    7y1
...........................................cAucgcugcgaucuauugaaaguc................................................   9    1    7y1
...........................................ccucgcugcgaucuauugaaGguc................................................   1    1    7y1
...........................................ccucgcugcgGucuauugaaaguc................................................   1    1    7y1
...........................................ccucUcugcgaucuauugaaaguc................................................   3    1    7y1
...........................................Ncucgcugcgaucuauugaaaguc................................................   1    1    7y1
...........................................ccucgcugcgaucuauugaaaguA................................................   2    1    7y1
...........................................ccucgcugcgaucuauuAaaaguc................................................   1    1    7y1
...........................................cAucgcugcgaucuauugaaaguca...............................................   2    1    7y1
...........................................Ncucgcugcgaucuauugaaaguca...............................................   1    1    7y1
...........................................ccucgcAgcgaucuauugaaaguca...............................................   1    1    7y1
...........................................ccucgcugAgaucuauugaaaguca...............................................   2    1    7y1
...........................................Acucgcugcgaucuauugaaaguca...............................................   3    1    7y1
...........................................Gcucgcugcgaucuauugaaaguca...............................................   1    1    7y1
...........................................ccucgcugcgaucuauugaaagAca...............................................   1    1    7y1
...........................................ccucgcugcgaucuauugaaaCuca...............................................   1    1    7y1
...........................................ccucgcugcgaucuauugaaaAuca...............................................   1    1    7y1
...........................................ccucgcugcgaucuauugaaaguca...............................................   761  0    7y1
...........................................ccucgcuUcgaucuauugaaaguca...............................................   1    1    7y1
...........................................ccucUcugcgaucuauugaaaguca...............................................   1    1    7y1
...........................................ccucgAugcgaucuauugaaaguca...............................................   1    1    7y1
...........................................ccucgcugcgauAuauugaaaguca...............................................   1    1    7y1
...........................................ccucgcugcgaucuaAugaaaguca...............................................   1    1    7y1
...........................................ccucgcugcgaucuauugaaaUuca...............................................   1    1    7y1
...........................................ccucgcugcgaucuauugaaagucG...............................................   1    1    7y1
...........................................Acucgcugcgaucuauugaaagucag..............................................   1    1    7y1
...........................................ccucgcuCcgaucuauugaaagucag..............................................   1    1    7y1
...........................................cAucgcugcgaucuauugaaagucag..............................................   1    1    7y1
...........................................ccucgcugcgaucuauAgaaagucag..............................................   1    1    7y1
...........................................ccucgcugcUaucuauugaaagucag..............................................   1    1    7y1
...........................................Ncucgcugcgaucuauugaaagucag..............................................   1    1    7y1
...........................................ccucgcugcgaucAauugaaagucag..............................................   1    1    7y1
...........................................ccucgcAgcgaucuauugaaagucag..............................................   2    1    7y1
...........................................ccucAcugcgaucuauugaaagucag..............................................   1    1    7y1
...........................................ccucgcugAgaucuauugaaagucag..............................................   1    1    7y1
...........................................ccucgcugcgaucuauugaaagucag..............................................   163  0    7y1
...........................................ccucgcugcgaucuauugaaagucagUcc...........................................   1    1    7y1
...........................................ccucgcugcgaucuauugaaagucagUccu..........................................   253  1    7y1
............................................cucgAugcgaucuauugaaa...................................................   1    1    7y1
............................................cucgcugcgaucuauAgaaa...................................................   2    1    7y1
............................................cucgcugcgauAuauugaaa...................................................   1    1    7y1
............................................cucgcugcgaucuauugaaa...................................................   413  0    7y1
............................................cucUcugcgaucuauugaaa...................................................   2    1    7y1
............................................cucgcugAgaucuauugaaa...................................................   2    1    7y1
............................................cucgcugcUaucuauugaaa...................................................   1    1    7y1
............................................Aucgcugcgaucuauugaaa...................................................   2    1    7y1
............................................cuAgcugcgaucuauugaaa...................................................   3    1    7y1
............................................cucgcugcCaucuauugaaa...................................................   1    1    7y1
............................................cAcgcugcgaucuauugaaa...................................................   2    1    7y1
............................................cuUgcugcgaucuauugaaag..................................................   1    1    7y1
```

**Star** **Mature**

```
gaccugcuucugggucggggguucguacguagcagagcagcucccucgcugcgaucuauugaaagucagcccucgacacaaggguuuguccgcgcgcgcgcgcgcgcgugc
............................................cucgcugcUaucuauugaaag..............................................  2     1  7y1
............................................cucgcugcgaucuauugUaag..............................................  1     1  7y1
............................................cAcgcugcgaucuauugaaag..............................................  12    1  7y1
............................................cucAcugcgaucuauugaaag..............................................  2     1  7y1
............................................Gucgcugcgaucuauugaaag..............................................  4     1  7y1
............................................cucgcugcgaucuauuUaaag..............................................  1     1  7y1
............................................cucgcAgcgaucuauugaaag..............................................  12    1  7y1
............................................cucgAugcgaucuauugaaag..............................................  3     1  7y1
............................................cuAgcugcgaucuauugaaag..............................................  10    1  7y1
............................................cucgcugcgaucuauAgaaag..............................................  2     1  7y1
............................................cucgcugcgaucAauugaaag..............................................  1     1  7y1
............................................cucgcugcgaucuauugaaaC..............................................  2     1  7y1
............................................cucgcugcgaucuauugaaag..............................................  2182  0  7y1
............................................cucgcugcgaucuauugGaag..............................................  1     1  7y1
............................................cucCcugcgaucuauugaaag..............................................  1     1  7y1
............................................cucgcugcgaucuaAugaaag..............................................  1     1  7y1
............................................cucgcugcgauUuauugaaag..............................................  1     1  7y1
............................................cucgcugcgaucuauugaGag..............................................  167   1  7y1
............................................cucgcugAgaucuauugaaag..............................................  8     1  7y1
............................................cucgcugcgaucuauugaaaU..............................................  2     1  7y1
............................................cucgcugcgaAcuauugaaag..............................................  2     1  7y1
............................................Aucgcugcgaucuauugaaag..............................................  2     1  7y1
............................................cucgcugcgauAuauugaaag..............................................  6     1  7y1
............................................cucgcuUcgaucuauugaaag..............................................  2     1  7y1
............................................cucgcugcgaucuauGgaaag..............................................  1     1  7y1
............................................cucgcugGgaucuauugaaag..............................................  2     1  7y1
............................................cucgcuCcgaucuauugaaagu.............................................  1     1  7y1
............................................cucgcCgcgaucuauugaaagu.............................................  1     1  7y1
............................................cucgcugcgauAuauugaaagu.............................................  1     1  7y1
............................................cAcgcugcgaucuauugaaagu.............................................  7     1  7y1
............................................cucgcugcCaucuauugaaagu.............................................  1     1  7y1
............................................cucgcugcgaucuauugaaagu.............................................  1949  0  7y1
............................................cucUcugcgaucuauugaaagu.............................................  1     1  7y1
............................................cucgcuUcgaucuauugaaagu.............................................  1     1  7y1
............................................cucgcNgcgaucuauugaaagu.............................................  1     1  7y1
............................................cucgcugcgaucuUuugaaagu.............................................  1     1  7y1
............................................cucgcugcUaucuauugaaagu.............................................  2     1  7y1
............................................cucgUugcgaucuauugaaagu.............................................  1     1  7y1
............................................cucgcugcgaucuauugaaagG.............................................  1     1  7y1
............................................cucgcugcgaucCauugaaagu.............................................  1     1  7y1
............................................cucgcugcgaucuauugaaaUu.............................................  1     1  7y1
............................................cuAgcugcgaucuauugaaagu.............................................  4     1  7y1
............................................cucgcAgcgaucuauugaaagu.............................................  4     1  7y1
............................................cucgcugcgaucAauugaaagu.............................................  1     1  7y1
............................................cucgcugcgaucuauuUaaagu.............................................  1     1  7y1
............................................cucgcugcgaAcuauugaaagu.............................................  3     1  7y1
............................................Nucgcugcgaucuauugaaagu.............................................  1     1  7y1
............................................cucgcugcgaucuauAgaaagu.............................................  2     1  7y1
............................................cucgcugcgaucuauuCaaagu.............................................  1     1  7y1
............................................cucgcugAgaucuauugaaagu.............................................  3     1  7y1
............................................Aucgcugcgaucuauugaaagu.............................................  6     1  7y1
............................................cucgcugcgauGuauugaaagu.............................................  1     1  7y1
............................................cNcgcugcgaucuauugaaagu.............................................  1     1  7y1
............................................cucgGugcgaucuauugaaaguc............................................  1     1  7y1
............................................cucgcuAcgaucuauugaaaguc............................................  1     1  7y1
............................................cAcgcugcgaucuauugaaaguc............................................  7     1  7y1
............................................cucgcuUcgaucuauugaaaguc............................................  1     1  7y1
............................................cucgcugcgaucuauuUaaaguc............................................  1     1  7y1
............................................cucgcugcgaucuauAgaaaguc............................................  1     1  7y1
............................................Nucgcugcgaucuauugaaaguc............................................  1     1  7y1
............................................Gucgcugcgaucuauugaaaguc............................................  1     1  7y1
............................................cucgcugcgaucuaAugaaaguc............................................  5     1  7y1
............................................cucgcugcgaucuauugaaaCuc............................................  1     1  7y1
............................................cucCcugcgaucuauugaaaguc............................................  2     1  7y1
............................................cuAgcugcgaucuauugaaaguc............................................  5     1  7y1
............................................cucgcugcgaCcuauugaaaguc............................................  1     1  7y1
............................................cucgcugcUaucuauugaaaguc............................................  1     1  7y1
............................................cucgAugcgaucuauugaaaguc............................................  3     1  7y1
............................................cucgcugcgaucuaCugaaaguc............................................  1     1  7y1
............................................cucgcugcgaGcuauugaaaguc............................................  1     1  7y1
```

### Star Mature

```
gaccugcuucugggucgggguuucguacguagcagagcagcucccucgcugcgaucuauugaaagucagcccucgacacaaggguuugucgcgcgcgcgcgcgcgcgcgcgugc
............................................cucgcugcgaucuauugaaaguc................................................ 2329 0 7y1
............................................cucgcugcgGucuauugaaaguc................................................ 1 1 7y1
............................................cucUcugcgaucuauugaaaguc................................................ 1 1 7y1
............................................cuGgcugcgaucuauugaaaguc................................................ 2 1 7y1
............................................cucgcugcgaucuauuCaaaguc................................................ 4 1 7y1
............................................cucgcAgcgaucuauugaaaguc................................................ 12 1 7y1
............................................cucgcugAgaucuauugaaaguc................................................ 8 1 7y1
............................................cucgcuCcgaucuauugaaaguc................................................ 2 1 7y1
............................................cucgcugcgauAuauugaaaguc................................................ 4 1 7y1
............................................Aucgcugcgaucuauugaaaguc................................................ 10 1 7y1
............................................cucgcugcgaucuauugaaaguA................................................ 2 1 7y1
............................................cucgcugcgaucuauugaaaguca............................................... 238 0 7y1
............................................cucgcugcgaAcuauugaaaguca............................................... 1 1 7y1
............................................cucgcugAgaucuauugaaaguca............................................... 1 1 7y1
............................................Aucgcugcgaucuauugaaaguca............................................... 1 1 7y1
............................................cucgcugcgaucuauugaaaguAa............................................... 3 1 7y1
............................................cuAgcugcgaucuauugaaaguca............................................... 1 1 7y1
.............................................ucgcAgcgaucuauugaa.................................................... 1 1 7y1
.............................................ucgcugAgaucuauugaa.................................................... 1 1 7y1
.............................................Acgcugcgaucuauugaa.................................................... 3 1 7y1
.............................................ucgcugcgaucuauAgaa.................................................... 1 1 7y1
.............................................ucgcugcgaucuauugaa.................................................... 152 0 7y1
.............................................ucgcugcgaucuauugaaG................................................... 2 1 7y1
.............................................ucgcugcgaucuaCugaaag.................................................. 1 1 7y1
.............................................ucgcugcgauGuauugaaag.................................................. 1 1 7y1
.............................................Ncgcugcgaucuauugaaag.................................................. 2 1 7y1
.............................................ucgcAgcgaucuauugaaag.................................................. 3 1 7y1
.............................................ucgcugcgNucuauugaaag.................................................. 1 1 7y1
.............................................ucgcugcgaucuUuugaaag.................................................. 1 1 7y1
.............................................ucgcugcgaucuauAgaaag.................................................. 4 1 7y1
.............................................Gcgcugcgaucuauugaaag.................................................. 1 1 7y1
.............................................ucgcugAgaucuauugaaag.................................................. 17 1 7y1
.............................................ucgcugcgaucuauugaaaC.................................................. 3 1 7y1
.............................................ucgcugcgaucuauugaaUg.................................................. 1 1 7y1
.............................................ucgcCgcgaucuauugaaag.................................................. 2 1 7y1
.............................................ucgcuCcgaucuauugaaag.................................................. 1 1 7y1
.............................................ucgcugcgaucuauuCaaag.................................................. 2 1 7y1
.............................................ucgcugcgaucAauugaaag.................................................. 6 1 7y1
.............................................ucgAugcgaucuauugaaag.................................................. 4 1 7y1
.............................................ucgcugcgCucuauugaaag.................................................. 1 1 7y1
.............................................ucgcugcgaucGauugaaag.................................................. 1 1 7y1
.............................................ucgcugcgaucuaAugaaag.................................................. 3 1 7y1
.............................................ucgcugcgaucuauugaaGg.................................................. 3 1 7y1
.............................................ucgcugcgaucuauugaaag.................................................. 3906 0 7y1
.............................................ucgcuUcgaucuauugaaag.................................................. 5 1 7y1
.............................................ucgcugcgaucuauugaaCg.................................................. 1 1 7y1
.............................................uAgcugcgaucuauugaaag.................................................. 12 1 7y1
.............................................ucgcugcCaucuauugaaag.................................................. 1 1 7y1
.............................................ucgcugcgaucuauuUaaag.................................................. 2 1 7y1
.............................................ucgGugcgaucuauugaaag.................................................. 2 1 7y1
.............................................ucgcugGgaucuauugaaag.................................................. 2 1 7y1
.............................................ucgcugcgauNuauugaaag.................................................. 1 1 7y1
.............................................ucUcugcgaucuauugaaag.................................................. 1 1 7y1
.............................................ucgcugcgauAuauugaaag.................................................. 4 1 7y1
.............................................uGgcugcgaucuauugaaag.................................................. 2 1 7y1
.............................................ucgcugcgaAcuauugaaag.................................................. 11 1 7y1
.............................................Acgcugcgaucuauugaaag.................................................. 60 1 7y1
.............................................ucgcuAcgaucuauugaaag.................................................. 2 1 7y1
.............................................ucgcGgcgaucuauugaaag.................................................. 1 1 7y1
.............................................ucgcugcUaucuauugaaag.................................................. 2 1 7y1
.............................................ucgcugcgaucuGuugaaag.................................................. 1 1 7y1
.............................................Acgcugcgaucuauugaaagu................................................. 4 1 7y1
.............................................ucgcugcCaucuauugaaagu................................................. 1 1 7y1
.............................................ucgcugcgaAcuauugaaagu................................................. 1 1 7y1
.............................................ucgcugcgaucuauugaaagu................................................. 138 0 7y1
.............................................ucgcugcgaucAauugaaagu................................................. 1 1 7y1
.............................................ucgcugcgaucuauugaaagucagcccucgaG...................................... 1 1 7y1
.............................................ucgcugcgaucuauugaaagucagcUcucgac...................................... 1 1 7y1
.............................................ucgcugcgaucuauugaaagucagccAucgac...................................... 1 1 7y1
.............................................Ncgcugcgaucuauugaaagucagcccucgac...................................... 1 1 7y1
```

| Star | Mature | | |
|---|---|---|---|
| gaccugcuucugggucgggguuucguacguagcagagcagcucccucgcugcgaucuauugaaagucagcccucgacacaaggguuuguccgcgcgcgcgcgcgcgcgcgcgugc | | | |
| .............................................ucgcugcgaucuaGugaaagucagcccucgac...................................... | 1 | 1 | 7y1 |
| .............................................ucgcugcgaucuaAugaaagucagcccucgac...................................... | 1 | 1 | 7y1 |
| .............................................ucgcugcgaucuauugaaagucagcccucgac...................................... | 506 | 0 | 7y1 |
| .............................................ucgcugcgaucuauuCaaagucagcccucgac...................................... | 1 | 1 | 7y1 |
| .............................................ucgcugcgaucuauugaaagucagcccucgaA...................................... | 1 | 1 | 7y1 |
| .............................................ucgcugcUaucuauugaaagucagcccucgac...................................... | 1 | 1 | 7y1 |
| .............................................Acgcugcgaucuauugaaagucagcccucgac...................................... | 7 | 1 | 7y1 |
| ..............................................cgcugcgaucuaAugaaa................................................... | 1 | 1 | 7y1 |
| ..............................................cUcugcgaucuauugaaa................................................... | 1 | 1 | 7y1 |
| ..............................................cgcugAgaucuauugaaa................................................... | 1 | 1 | 7y1 |
| ..............................................cCcugcgaucuauugaaa................................................... | 2 | 1 | 7y1 |
| ..............................................cgcugcgaucuauugaaa................................................... | 357 | 0 | 7y1 |
| ..............................................cgcuAcgaucuauugaaa................................................... | 1 | 1 | 7y1 |
| ..............................................cgcugcgaAcuauugaaa................................................... | 1 | 1 | 7y1 |
| ..............................................cgcugcUaucuauugaaa................................................... | 1 | 1 | 7y1 |
| ..............................................cgcugcgaucuauugaaG................................................... | 1 | 1 | 7y1 |
| ..............................................cgcugcgaGcuauugaaag.................................................. | 2 | 1 | 7y1 |
| ..............................................cgcugcgaucuUuugaaag.................................................. | 1 | 1 | 7y1 |
| ..............................................cgcugcgaucuauuCaaag.................................................. | 1 | 1 | 7y1 |
| ..............................................cgAugcgaucuauugaaag.................................................. | 20 | 1 | 7y1 |
| ..............................................cgcugcgaucuauugaUag.................................................. | 4 | 1 | 7y1 |
| ..............................................cgcugcgaucuauCgaaag.................................................. | 1 | 1 | 7y1 |
| ..............................................cgcugcgaucuauugaaaC.................................................. | 1 | 1 | 7y1 |
| ..............................................cgcGgcgaucuauugaaag.................................................. | 2 | 1 | 7y1 |
| ..............................................cgcugcgaucuauuUaaag.................................................. | 150 | 1 | 7y1 |
| ..............................................Agcugcgaucuauugaaag.................................................. | 16 | 1 | 7y1 |
| ..............................................cgGugcgaucuauugaaag.................................................. | 3 | 1 | 7y1 |
| ..............................................cgcugcgaucuauAgaaag.................................................. | 9 | 1 | 7y1 |
| ..............................................Ugcugcgaucuauugaaag.................................................. | 2 | 1 | 7y1 |
| ..............................................cgUugcgaucuauugaaag.................................................. | 1 | 1 | 7y1 |
| ..............................................cgcugAgaucuauugaaag.................................................. | 29 | 1 | 7y1 |
| ..............................................cgcugcgauGuauugaaag.................................................. | 2 | 1 | 7y1 |
| ..............................................cCcugcgaucuauugaaag.................................................. | 4 | 1 | 7y1 |
| ..............................................cgcugcgUucuauugaaag.................................................. | 2 | 1 | 7y1 |
| ..............................................cgcCgcgaucuauugaaag.................................................. | 1 | 1 | 7y1 |
| ..............................................cgcugcgaucuauugaGag.................................................. | 2 | 1 | 7y1 |
| ..............................................cgcugcgaucuauugaaag.................................................. | 6574 | 0 | 7y1 |
| ..............................................cgcugcgaucAauugaaag.................................................. | 4 | 1 | 7y1 |
| ..............................................Ngcugcgaucuauugaaag.................................................. | 1 | 1 | 7y1 |
| ..............................................cgcugGgaucuauugaaag.................................................. | 2 | 1 | 7y1 |
| ..............................................cUcugcgaucuauugaaag.................................................. | 8 | 1 | 7y1 |
| ..............................................cgcugcCaucuauugaaag.................................................. | 4 | 1 | 7y1 |
| ..............................................cgcuCcgaucuauugaaag.................................................. | 2 | 1 | 7y1 |
| ..............................................cgcugcgaucGauugaaag.................................................. | 1 | 1 | 7y1 |
| ..............................................cgcAgcgaucuauugaaag.................................................. | 11 | 1 | 7y1 |
| ..............................................cgcugcgauAuauugaaag.................................................. | 14 | 1 | 7y1 |
| ..............................................cgcugcgaucuauugaaaU.................................................. | 4 | 1 | 7y1 |
| ..............................................cgcugcgaucuauugUaag.................................................. | 2 | 1 | 7y1 |
| ..............................................cAcugcgaucuauugaaag.................................................. | 1 | 1 | 7y1 |
| ..............................................cgcuUcgaucuauugaaag.................................................. | 7 | 1 | 7y1 |
| ..............................................cgcugcgaAcuauugaaag.................................................. | 9 | 1 | 7y1 |
| ..............................................cgcugcgaucuauugaaUg.................................................. | 1 | 1 | 7y1 |
| ..............................................Ggcugcgaucuauugaaag.................................................. | 4 | 1 | 7y1 |
| ..............................................cgcugcgaucuaAugaaag.................................................. | 7 | 1 | 7y1 |
| ..............................................cgcuAcgaucuauugaaag.................................................. | 1 | 1 | 7y1 |
| ..............................................cgcugcUaucuauugaaag.................................................. | 10 | 1 | 7y1 |
| ..............................................cgcugcgaucuauugaaaA.................................................. | 1 | 1 | 7y1 |
| ..............................................cUcugcgaucuauugaaagu................................................. | 3 | 1 | 7y1 |
| ..............................................cgcugcgaucuauugaaaCu................................................. | 1 | 1 | 7y1 |
| ..............................................cgcugcgaucuauugaaagu................................................. | 160 | 0 | 7y1 |
| ..............................................Ggcugcgaucuauugaaagu................................................. | 1 | 1 | 7y1 |
| ..............................................cgcugcgaucuauAgaaagu................................................. | 1 | 1 | 7y1 |
| ..............................................cgcugAgaucuauugaaagu................................................. | 2 | 1 | 7y1 |
| ..............................................cgcugcgaucuaAugaaagu................................................. | 1 | 1 | 7y1 |
| ..............................................cgcugcgaucuauugaaagAc................................................ | 1 | 1 | 7y1 |
| ..............................................cgcugcgaucuauAgaaaguc................................................ | 2 | 1 | 7y1 |
| ..............................................cgcugcgauAuauugaaaguc................................................ | 2 | 1 | 7y1 |
| ..............................................cUcugcgaucuauugaaaguc................................................ | 4 | 1 | 7y1 |
| ..............................................cgcugcgaAcuauugaaaguc................................................ | 1 | 1 | 7y1 |
| ..............................................Agcugcgaucuauugaaaguc................................................ | 2 | 1 | 7y1 |

**Star** **Mature**

```
gaccugcuucugggucgggguuucguacguagcagagcagcucccucgcugcgaucuauugaaagucagcccucgacacaaggguuuguccgcgcgcgcgcgcgcgcgcgcgugc
..............................................cgcugcgaucuauugGaaguc................................................  1     1  7y1
..............................................cgcugcgaucuauugaGaguc................................................  1     1  7y1
..............................................cgcugcgaucuUuugaaaguc................................................  1     1  7y1
..............................................cgcugAgaucuauugaaaguc................................................  2     1  7y1
..............................................cgcugcgauUuauugaaaguc................................................  1     1  7y1
..............................................cgcugcgaucuauugaaaguA................................................  1     1  7y1
..............................................cgcugcgaucuauugaaaguc................................................  553   0  7y1
..............................................cgcAgcgaucuauugaaaguc................................................  1     1  7y1
..............................................cgcugcgaucuauugaaGgucag..............................................  2     1  7y1
..............................................cgcugcgaucuauugaaagucag..............................................  184   0  7y1
..............................................cgcugcgaucuauugaaagucaU..............................................  1     1  7y1
..............................................cgcugcgaucuauAgaaagucag..............................................  1     1  7y1
..............................................cgcugcgaucuaAugaaagucag..............................................  1     1  7y1
..............................................cgcugAgaucuauugaaagucag..............................................  1     1  7y1
..............................................cUcugcgaucuauugaaagucag..............................................  1     1  7y1
..............................................Agcugcgaucuauugaaagucagcccucgac......................................  1     1  7y1
..............................................cgAugcgaucuauugaaagucagcccucgac......................................  1     1  7y1
..............................................cgcugAgaucuauugaaagucagcccucgac......................................  1     1  7y1
..............................................cgcugcgaucuauugaaagucagcccucgCc......................................  1     1  7y1
..............................................cgcugcgaucuauugaaagucagcccucgac......................................  162   0  7y1
...............................................gcuUcgaucuauugaaag..................................................  4     1  7y1
...............................................gcugcgaucAauugaaag..................................................  7     1  7y1
...............................................Ucugcgaucuauugaaag..................................................  12    1  7y1
...............................................gcGgcgaucuauugaaag..................................................  2     1  7y1
...............................................gcugcgaucuauuUaaag..................................................  1     1  7y1
...............................................gcugcUaucuauugaaag..................................................  5     1  7y1
...............................................gcugcgauAuauugaaag..................................................  3     1  7y1
...............................................gcugcgaucuauugaaaC..................................................  2     1  7y1
...............................................gcugcgaucuaAugaaag..................................................  4     1  7y1
...............................................gcugcCaucuauugaaag..................................................  2     1  7y1
...............................................gAugcgaucuauugaaag..................................................  6     1  7y1
...............................................gcugcgaucuauugaaag..................................................  2699  0  7y1
...............................................gcuCcgaucuauugaaag..................................................  1     1  7y1
...............................................gcugcgaucuGuugaaag..................................................  1     1  7y1
...............................................gcugcgaucuauuCaaag..................................................  1     1  7y1
...............................................gcugcgaucuauugGaag..................................................  1     1  7y1
...............................................gcugcgaAcuauugaaag..................................................  1     1  7y1
...............................................gcAgcgaucuauugaaag..................................................  4     1  7y1
...............................................gcugcgaucuauugaUag..................................................  2     1  7y1
...............................................gGugcgaucuauugaaag..................................................  2     1  7y1
...............................................Ccugcgaucuauugaaag..................................................  8     1  7y1
...............................................gcugcgCucuauugaaag..................................................  1     1  7y1
...............................................gcugGgaucuauugaaag..................................................  1     1  7y1
...............................................gcugcgaucuauAgaaag..................................................  8     1  7y1
...............................................gcugcgaucuUuugaaag..................................................  1     1  7y1
...............................................gcugcgaucuauugaGag..................................................  2     1  7y1
...............................................Acugcgaucuauugaaag..................................................  1     1  7y1
...............................................gcugcgaucuaCugaaag..................................................  1     1  7y1
...............................................Ncugcgaucuauugaaag..................................................  2     1  7y1
...............................................gcugAgaucuauugaaag..................................................  6     1  7y1
...............................................gcuAcgaucuauugaaag..................................................  1     1  7y1
...............................................gcugcgaucuauugaaaU..................................................  1     1  7y1
...............................................gcugcgaucuauugaaUg..................................................  1     1  7y1
...............................................gcAgcgaucuauugaaagu.................................................  2     1  7y1
...............................................gcugcgaucuauugaaagu.................................................  111   0  7y1
...............................................gcugcgaucuauugaaagA.................................................  1     1  7y1
...............................................gcugcgaAcuauugaaagu.................................................  1     1  7y1
...............................................gAugcgaucuauugaaagu.................................................  2     1  7y1
...............................................gcugcgaucuauAgaaagu.................................................  1     1  7y1
...............................................gcugcgaucuauugaaagAc................................................  2     1  7y1
...............................................gcugAgaucuauugaaaguc................................................  1     1  7y1
...............................................gcugcgaAcuauugaaaguc................................................  1     1  7y1
...............................................gcugcgaucuauugaaaguU................................................  1     1  7y1
...............................................gcugcgaucuauAgaaaguc................................................  1     1  7y1
...............................................gcuAcgaucuauugaaaguc................................................  1     1  7y1
...............................................gAugcgaucuauugaaaguc................................................  1     1  7y1
...............................................gcugcgaucuauugaaaUuc................................................  1     1  7y1
...............................................Ucugcgaucuauugaaaguc................................................  2     1  7y1
...............................................gcugcgaucuaAugaaaguc................................................  1     1  7y1
...............................................gcugcgaucGauugaaaguc................................................  1     1  7y1
```

**Star** **Mature**

gaccugcuucugggucgggguuucguacguagcagagcagcucccucgcugcgaucuauugaaagucagcccucgacacaaggguuuguccgcgcgcgcgcgcgcgcgugc

| | | | |
|---|---|---|---|
| .............................................gcugcgaucuauuNaaaguc.............................................. | 1 | 1 | 7y1 |
| .............................................gcugcgaucuaCugaaaguc.............................................. | 1 | 1 | 7y1 |
| .............................................gcugcgaucAauugaaaguc.............................................. | 1 | 1 | 7y1 |
| .............................................gcugcgaGcuauugaaaguc.............................................. | 1 | 1 | 7y1 |
| .............................................gcugcgaucuauugaaaguA.............................................. | 1 | 1 | 7y1 |
| .............................................gcuCcgaucuauugaaaguc.............................................. | 1 | 1 | 7y1 |
| .............................................gcugcgaucuauugaaaguc.............................................. | 562 | 0 | 7y1 |
| .............................................Acugcgaucuauugaaagucag............................................ | 1 | 1 | 7y1 |
| .............................................gcugcgaucuauugaaagucag............................................ | 181 | 0 | 7y1 |
| .............................................gAugcgaucuauugaaagucag............................................ | 2 | 1 | 7y1 |
| ..............................................cugcgaAcuauugaaagu............................................... | 2 | 1 | 7y1 |
| ..............................................cugcgaucuauugaaagA............................................... | 1 | 1 | 7y1 |
| ..............................................Nugcgaucuauugaaagu............................................... | 1 | 1 | 7y1 |
| ..............................................cugcgaucuaAugaaagu............................................... | 1 | 1 | 7y1 |
| ..............................................cuAcgaucuauugaaagu............................................... | 1 | 1 | 7y1 |
| ..............................................cugcgaucuauugaaagu............................................... | 301 | 0 | 7y1 |
| ..............................................cugcgaucuauugaaaUu............................................... | 1 | 1 | 7y1 |
| ..............................................Gugcgaucuauugaaagu............................................... | 1 | 1 | 7y1 |
| ..............................................cAgcgaucuauugaaagu............................................... | 1 | 1 | 7y1 |
| ..............................................Augcgaucuauugaaagu............................................... | 1 | 1 | 7y1 |
| ..............................................cugAgaucuauugaaaguc.............................................. | 1 | 1 | 7y1 |
| ..............................................cugcgaucuauugaaaguU.............................................. | 1 | 1 | 7y1 |
| ..............................................cugcgaucuauAgaaaguc.............................................. | 1 | 1 | 7y1 |
| ..............................................cuUcgaucuauugaaaguc.............................................. | 1 | 1 | 7y1 |
| ..............................................cugcgaAcuauugaaaguc.............................................. | 1 | 1 | 7y1 |
| ..............................................cAgcgaucuauugaaaguc.............................................. | 1 | 1 | 7y1 |
| ..............................................cugcgauAuauugaaaguc.............................................. | 1 | 1 | 7y1 |
| ..............................................cugcgaGcuauugaaaguc.............................................. | 1 | 1 | 7y1 |
| ..............................................Augcgaucuauugaaaguc.............................................. | 3 | 1 | 7y1 |
| ..............................................cugcgaucAauugaaaguc.............................................. | 1 | 1 | 7y1 |
| ..............................................cugcgaucuauugaaagAc.............................................. | 1 | 1 | 7y1 |
| ..............................................cugGgaucuauugaaaguc.............................................. | 1 | 1 | 7y1 |
| ..............................................cugcUaucuauugaaaguc.............................................. | 1 | 1 | 7y1 |
| ..............................................cugcgaucuauugaaaguc.............................................. | 436 | 0 | 7y1 |
| ..............................................cugcgaucAauugaaagucagc........................................... | 3 | 1 | 7y1 |
| ..............................................cugcgaucCauugaaagucagc........................................... | 337 | 1 | 7y1 |
| ..............................................cugcgaucuauugaaagucagcccA........................................ | 1 | 1 | 7y1 |
| ..............................................cuAcgaucuauugaaagucagcccu........................................ | 1 | 1 | 7y1 |
| ..............................................Augcgaucuauugaaagucagcccu........................................ | 1 | 1 | 7y1 |
| ..............................................cugcgaucuauugaaagucagcccu........................................ | 172 | 0 | 7y1 |
| ..............................................cuCcgaucuauugaaagucagcccu........................................ | 1 | 1 | 7y1 |
| ..............................................cugcgaucAauugaaagucagcccu........................................ | 1 | 1 | 7y1 |
| ..............................................cuUcgaucuauugaaagucagcccu........................................ | 1 | 1 | 7y1 |
| ..............................................cuUcgaucuauugaaagucagcccuc....................................... | 169 | 1 | 7y1 |
| ..............................................cugUgaucuauugaaagucagcccucgac.................................... | 1 | 1 | 7y1 |
| ..............................................cugcgaucuauugaaagucagcccucgac.................................... | 218 | 0 | 7y1 |
| ..............................................cugcgaucuauugaaaCucagcccucgac.................................... | 1 | 1 | 7y1 |
| ..............................................cugcgaucuauugaaagAcagcccucgac.................................... | 1 | 1 | 7y1 |
| ..............................................cugcgaucuauugaaagucagcccucgaA.................................... | 1 | 1 | 7y1 |
| ...............................................ugAgaucuauugaaagucagc........................................... | 1 | 1 | 7y1 |
| ...............................................ugcgaucuauugaaagucagc........................................... | 702 | 0 | 7y1 |
| ...............................................ugcgaucuauugaaaguAagc........................................... | 1 | 1 | 7y1 |
| ...............................................uCcgaucuauugaaagucagc........................................... | 2 | 1 | 7y1 |
| ...............................................ugcgaucuauugaCagucagc........................................... | 1 | 1 | 7y1 |
| ...............................................ugcgauAuauugaaagucagc........................................... | 2 | 1 | 7y1 |
| ...............................................ugcgaucuauugaaagucagG........................................... | 1 | 1 | 7y1 |
| ...............................................Ggcgaucuauugaaagucagc........................................... | 2 | 1 | 7y1 |
| ...............................................ugcgaucuauugaaagucagU........................................... | 1 | 1 | 7y1 |
| ...............................................ugcgaucuauCgaaagucagc........................................... | 1 | 1 | 7y1 |
| ...............................................ugcgaucuauugaaagAcagc........................................... | 1 | 1 | 7y1 |
| ...............................................Agcgaucuauugaaagucagc........................................... | 10 | 1 | 7y1 |
| ...............................................ugcgaucuauuAaaagucagc........................................... | 1 | 1 | 7y1 |
| ...............................................ugcgaucuauuCaaagucagc........................................... | 1 | 1 | 7y1 |
| ...............................................ugcgaucuauugaaagucagcAcucgacac.................................. | 1 | 1 | 7y1 |
| ...............................................ugcgaucuauugaaagucagcccucgacac.................................. | 154 | 0 | 7y1 |
| ...............................................ugcgaucuauugaaagucagcccuUgacac.................................. | 1 | 1 | 7y1 |
| ...............................................Agcgaucuauugaaagucagcccucgacac.................................. | 1 | 1 | 7y1 |
| ...............................................ugAgaucuauugaaagucagcccucgacac.................................. | 1 | 1 | 7y1 |
| ...............................................ugcgaucuauugaaagucagcccucgUcac.................................. | 1 | 1 | 7y1 |
| ...............................................ugcgaucuauugaaagucagcccucgacaA.................................. | 1 | 1 | 7y1 |

**Star** **Mature**

```
gaccugcuucugggucgggguuucguacguagcagagcagcucccucgcugcgaucuauugaaagucagcccucgacacaaggguuuguccgcgcgcgcgcgcgcgcgcgugc
..................................................gcgaucuauAgaaagucagc...........................................   1     1   7y1
..................................................gcgaucuauuCaaagucagc...........................................   1     1   7y1
..................................................gcgaucAauugaaagucagc...........................................   1     1   7y1
..................................................gcgaucuauugaaagCcagc...........................................   1     1   7y1
..................................................gcgaucCauugaaagucagc...........................................   157   1   7y1
..................................................gcgaucuauugaaagucagc...........................................   259   0   7y1
..................................................gcgaucuauugaGagucagc...........................................   390   1   7y1
..................................................gcgaucuauAgaaagucagcc..........................................   2     1   7y1
..................................................gcgaucuauuAaaagucagcc..........................................   1     1   7y1
..................................................gcgaucuauugaaaCucagcc..........................................   1     1   7y1
..................................................gcgaucuauugaaagucagcc..........................................   149   0   7y1
...................................................cgaucuauugaaagucagA...........................................   3     1   7y1
...................................................cgaucuauAgaaagucagc...........................................   4     1   7y1
...................................................cgaucuauugaaGgucagc...........................................   1     1   7y1
...................................................cgaucuauuUaaagucagc...........................................   1     1   7y1
...................................................cgauAuauugaaagucagc...........................................   1     1   7y1
...................................................Agaucuauugaaagucagc...........................................   1     1   7y1
...................................................cgaucuauugUaagucagc...........................................   1     1   7y1
...................................................cgaucuauugaaagucagc...........................................   565   0   7y1
...................................................cgaucAauugaaagucagc...........................................   2     1   7y1
...................................................cgaucuauGgaaagucagc...........................................   1     1   7y1
...................................................cgaucuauugaaaguGagc...........................................   1     1   7y1
...................................................cgaucuauugaaaguAagc...........................................   1     1   7y1
...................................................cUaucuauugaaagucagc...........................................   1     1   7y1
...................................................cgaucuauCgaaagucagc...........................................   1     1   7y1
...................................................cgaucuauugaaagucCgcccu........................................   1     1   7y1
...................................................cgaucuauugaaaUucagcccu........................................   1     1   7y1
...................................................cgaucuauugaaagucagcUcu........................................   1     1   7y1
...................................................cgaucAauugaaagucagcccu........................................   1     1   7y1
...................................................cgaucuauAgaaagucagcccu........................................   1     1   7y1
...................................................cgaucuauugaaagucagcccu........................................   243   0   7y1
...................................................cgaucuauugaaaguGagcccu........................................   1     1   7y1
...................................................cUaucuauugaaagucagcccu........................................   2     1   7y1
...................................................cgaucuaAugaaagucagcccu........................................   1     1   7y1
...................................................Ggaucuauugaaagucagcccucg......................................   1     1   7y1
...................................................cgauAuauugaaagucagcccucg......................................   1     1   7y1
...................................................Agaucuauugaaagucagcccucg......................................   2     1   7y1
...................................................cgaucAauugaaagucagcccucg......................................   2     1   7y1
...................................................cgaucuauugaaagAcagcccucg......................................   1     1   7y1
...................................................cgaucuauCgaaagucagcccucg......................................   1     1   7y1
...................................................cgaAcuauugaaagucagcccucg......................................   1     1   7y1
...................................................cgaucuauAgaaagucagcccucg......................................   1     1   7y1
...................................................cgaucuaGugaaagucagcccucg......................................   1     1   7y1
...................................................cgaucuauugaGagucagcccucg......................................   1     1   7y1
...................................................cCaucuauugaaagucagcccucg......................................   1     1   7y1
...................................................cgaucuauugaaagucagAccucg......................................   1     1   7y1
...................................................cgaucuauugaaagucagcccucg......................................   478   0   7y1
...................................................Ngaucuauugaaagucagcccucg......................................   1     1   7y1
...................................................cgUucuauugaaagucagcccucg......................................   1     1   7y1
...................................................Ngaucuauugaaagucagcccucgacacaa................................   1     1   7y1
...................................................Agaucuauugaaagucagcccucgacacaa................................   1     1   7y1
...................................................cgaucuauugaaagucagcccucgacacaG................................   1     1   7y1
...................................................cgaucAauugaaagucagcccucgacacaa................................   1     1   7y1
...................................................cgaucuauugaaagucagcccucgacacaa................................   237   0   7y1
...................................................cgaucuauugaaagucagcccucgCcacaa................................   1     1   7y1
...................................................cgauAuauugaaagucagcccucgacacaa................................   2     1   7y1
...................................................cgaucuauugaaagAcagcccucgacacaa................................   1     1   7y1
....................................................gaucuauuUaaagucagc...........................................   1     1   7y1
....................................................gaucCauugaaagucagc...........................................   142   1   7y1
....................................................gaucuauugaaagucagc...........................................   390   0   7y1
....................................................gaAcuauugaaagucagc...........................................   1     1   7y1
....................................................gaucAauugaaagucagc...........................................   2     1   7y1
....................................................gaucuauugaaagucagG...........................................   1     1   7y1
....................................................gauAuauugaaagucagc...........................................   1     1   7y1
....................................................gaucAauugaaagucagcc..........................................   2     1   7y1
....................................................gaucuauugaaaUucagcc..........................................   1     1   7y1
....................................................gaucuauuCaaagucagcc..........................................   2     1   7y1
....................................................gaucuaAugaaagucagcc..........................................   1     1   7y1
....................................................gaucuauugaaagucaCcc..........................................   1     1   7y1
....................................................gaucuauugaaagucGgcc..........................................   1     1   7y1
```

Star Mature

```
gaccugcuucugggucgggguuucguacguagcagagcagcucccucgcugcgaucuauugaaagucagcccucgacacaaggguuuguccgcgcgcgcgcgcgcgcgcgugc
....................................................gauGuauugaaagucagcc............................................ 1 1 7y1
....................................................gauAuauugaaagucagcc............................................ 3 1 7y1
....................................................gaucuauugaaagucagcA............................................ 1 1 7y1
....................................................gaucuauugaaagucagcG............................................ 1 1 7y1
....................................................gGucuauugaaagucagcc............................................ 1 1 7y1
....................................................gaucuauugaaaCucagcc............................................ 2 1 7y1
....................................................Caucuauugaaagucagcc............................................ 1 1 7y1
....................................................gaucuauugaaagucagcc............................................ 683 0 7y1
....................................................gaucuUuugaaagucagcc............................................ 1 1 7y1
....................................................gaucAauugaaagucagcccucg........................................ 2 1 7y1
....................................................gaucCauugaaagucagcccucg........................................ 193 1 7y1
.....................................................aucuauugaaagucagcA............................................ 13 1 7y1
.....................................................aucuauugaaagucagcc............................................ 5120 0 7y1
.....................................................aucuauugaaagucaCcc............................................ 5 1 7y1
.....................................................aucuauugaaaAucagcc............................................ 1 1 7y1
.....................................................aucCauugaaagucagcc............................................ 1243 1 7y1
.....................................................aucuauugaaagucaUcc............................................ 2 1 7y1
.....................................................aucuUuugaaagucagcc............................................ 1 1 7y1
.....................................................aucuauugaaagGcagcc............................................ 1 1 7y1
.....................................................aucuauugaaaCucagcc............................................ 4 1 7y1
.....................................................aucuauugaaagucagUc............................................ 2 1 7y1
.....................................................aucuauugaaagAcagcc............................................ 6 1 7y1
.....................................................aucuaAugaaagucagcc............................................ 23 1 7y1
.....................................................auGuauugaaagucagcc............................................ 1 1 7y1
.....................................................aAcuauugaaagucagcc............................................ 19 1 7y1
.....................................................aucuauugaUagucagcc............................................ 1 1 7y1
.....................................................auAuauugaaagucagcc............................................ 27 1 7y1
.....................................................aucuauAgaaagucagcc............................................ 21 1 7y1
.....................................................aucuaCugaaagucagcc............................................ 1 1 7y1
.....................................................aucAauugaaagucagcc............................................ 10 1 7y1
.....................................................aucuauugaaagucagcG............................................ 2 1 7y1
.....................................................aucuauuCaaagucagcc............................................ 4 1 7y1
.....................................................aucuauugaaGgucagcc............................................ 1 1 7y1
.....................................................aucuauuUaaagucagcc............................................ 3 1 7y1
.....................................................Uucuauugaaagucagcc............................................ 3 1 7y1
.....................................................Nucuauugaaagucagcc............................................ 5 1 7y1
.....................................................aucuauugaaaUucagcc............................................ 2 1 7y1
.....................................................Gucuauugaaagucagcc............................................ 4 1 7y1
.....................................................aucuauugaaUgucagcc............................................ 1 1 7y1
.....................................................aucuauuAaaagucagcc............................................ 2 1 7y1
.....................................................aucuauugaaaguAagcc............................................ 12 1 7y1
.....................................................aucuauAgaaagucagccc........................................... 8 1 7y1
.....................................................aucuUuugaaagucagccc........................................... 1 1 7y1
.....................................................aucuauugaaagucaUccc........................................... 1 1 7y1
.....................................................aucuauugaaagucagccU........................................... 237 1 7y1
.....................................................aucuaCugaaagucagccc........................................... 1 1 7y1
.....................................................aucuauugaaagucagccc........................................... 1694 0 7y1
.....................................................aucuauugaaagAcagccc........................................... 1 1 7y1
.....................................................aucuaAugaaagucagccc........................................... 5 1 7y1
.....................................................auGuauugaaagucagccc........................................... 2 1 7y1
.....................................................aucGauugaaagucagccc........................................... 2 1 7y1
.....................................................aucuauugaaaUucagccc........................................... 2 1 7y1
.....................................................aucuauuUaaagucagccc........................................... 1 1 7y1
.....................................................aucuauugaaagucagccA........................................... 6 1 7y1
.....................................................aucuauugaaagucagcAc........................................... 4 1 7y1
.....................................................aucuauugaUagucagccc........................................... 1 1 7y1
.....................................................aAcuauugaaagucagccc........................................... 10 1 7y1
.....................................................aucuauugUaagucagccc........................................... 1 1 7y1
.....................................................aucuauugaaaCucagccc........................................... 1 1 7y1
.....................................................aucAauugaaagucagccc........................................... 1 1 7y1
.....................................................aucCauugaaagucagccc........................................... 209 1 7y1
.....................................................Nucuauugaaagucagccc........................................... 1 1 7y1
.....................................................auAuauugaaagucagccc........................................... 7 1 7y1
.....................................................aucuauugaaaguAagccc........................................... 5 1 7y1
.....................................................Gucuauugaaagucagccc........................................... 2 1 7y1
.....................................................aucuauugaaagucagccG........................................... 3 1 7y1
.....................................................aucuauugaaagucagcGc........................................... 2 1 7y1
.....................................................Nucuauugaaagucagcccu.......................................... 1 1 7y1
.....................................................aucuauugaaagAcagcccu.......................................... 1 1 7y1
.....................................................aucuauugaaUgucagcccu.......................................... 1 1 7y1
```

# Star          Mature

```
gaccugcuucugggucggggguuucguacguagcagagcagcucccucgcugcgaucuauugaaagucagcccucgacacaaggguuuguccgcgcgcgcgcgcgcgcgcgugc
.....................................................aucuauugaaagucagcAcu........................................    2      1      7y1
.....................................................aucuauuCaaagucagcccu........................................    1      1      7y1
.....................................................aucuauugaaagucaUcccu........................................    1      1      7y1
.....................................................auAuauugaaagucagcccu........................................    2      1      7y1
.....................................................aucuauugaaagucagcccG........................................    2      1      7y1
.....................................................aucuaAugaaagucagcccu........................................    4      1      7y1
.....................................................aucuauugaaagucagcccA........................................    1      1      7y1
.....................................................aucuauugaaagucagcccu........................................    755    0      7y1
.....................................................aucuauAgaaagucagcccu........................................    3      1      7y1
.....................................................aAcuauugaaagucagcccu........................................    2      1      7y1
.....................................................aucuauugaaagucagccAu........................................    1      1      7y1
.....................................................aucCauugaaagucagcccu........................................    223    1      7y1
.....................................................aucuauugaaaguAagcccu........................................    1      1      7y1
.....................................................aucCauugaaagucagcccuc.......................................    1285   1      7y1
.....................................................aucAauugaaagucagcccuc.......................................    2      1      7y1
.....................................................aucuauAgaaagucagcccucg......................................    4      1      7y1
.....................................................aucuauugaaagucagcccucg......................................    1785   0      7y1
.....................................................aucuauugaaagucagcccucU......................................    1      1      7y1
.....................................................aucuauugaaagucagcAcucg......................................    2      1      7y1
.....................................................aucuauugaaagucagAccucg......................................    6      1      7y1
.....................................................aCcuauugaaagucagcccucg......................................    1      1      7y1
.....................................................aucuauuUaaagucagcccucg......................................    1      1      7y1
.....................................................aucuauugaaaguAagcccucg......................................    5      1      7y1
.....................................................aucuauugNaagucagcccucg......................................    1      1      7y1
.....................................................aucuauugaaagAcagcccucg......................................    2      1      7y1
.....................................................aucuaAugaaagucagcccucg......................................    6      1      7y1
.....................................................aAcuauugaaagucagcccucg......................................    5      1      7y1
.....................................................auAuauugaaagucagcccucg......................................    8      1      7y1
.....................................................aucuauugaaagucagcccAcg......................................    1      1      7y1
.....................................................aucuauugaaagucagUccucg......................................    2      1      7y1
.....................................................aucuauugaaagucagGccucg......................................    1      1      7y1
.....................................................aucuauugaaaUucagcccucg......................................    1      1      7y1
.....................................................Gucuauugaaagucagcccucg......................................    1      1      7y1
.....................................................aucuauugaUagucagcccucg......................................    1      1      7y1
.....................................................aucuauugaaagucGgcccucg......................................    2      1      7y1
.....................................................aucCauugaaagucagcccucg......................................    914    1      7y1
.....................................................aucuauuCaaagucagcccucg......................................    2      1      7y1
.....................................................aucuauugaaagucagccAucg......................................    2      1      7y1
.....................................................aucuauugaaagucaCcccucg......................................    1      1      7y1
.....................................................auUuauugaaagucagcccucg......................................    249    1      7y1
.....................................................aucuauugGaagucagcccucg......................................    1      1      7y1
.....................................................aucuauugaaagucagcccucC......................................    2      1      7y1
.....................................................Nucuauugaaagucagcccucg......................................    1      1      7y1
.....................................................aucuauugaaagucagcccuAg......................................    4      1      7y1
.....................................................aucuauugaaaguGagcccucg......................................    1      1      7y1
.....................................................aucuauugaGagucagcccucg......................................    1      1      7y1
.....................................................aucAauugaaagucagcccucg......................................    1      1      7y1
.....................................................aucuauugaaagucaUcccucg......................................    1      1      7y1
.....................................................aucuauugaaagucagcAcucga.....................................    2      1      7y1
.....................................................aucuauugaaaguAagcccucga.....................................    1      1      7y1
.....................................................aucuauugaaagucagccAucga.....................................    2      1      7y1
.....................................................aucuauugaaagucagcUcucga.....................................    1      1      7y1
.....................................................aucuauugaaagucagcccucga.....................................    460    0      7y1
.....................................................auAuauugaaagucagcccucga.....................................    2      1      7y1
.....................................................aucuauugaaagucagcccuAga.....................................    1      1      7y1
.....................................................aucuauugaaagucagccGucga.....................................    1      1      7y1
.....................................................aucuauAgaaagucagcccucga.....................................    6      1      7y1
.....................................................aucuaAugaaagucagcccucga.....................................    2      1      7y1
.....................................................aAcuauugaaagucagcccucga.....................................    3      1      7y1
.....................................................aucuauCgaaagucagcccucga.....................................    1      1      7y1
.....................................................aucuauugUaagucagcccucga.....................................    1      1      7y1
.....................................................aucuauugaaagucagcccucgac....................................    272    0      7y1
.....................................................Uucuauugaaagucagcccucgac....................................    2      1      7y1
.....................................................aucuauAgaaagucagcccucgac....................................    2      1      7y1
.....................................................aucuaAugaaagucagcccucgac....................................    2      1      7y1
.....................................................aucuauugaaagucagcccucgaA....................................    1      1      7y1
.....................................................aAcuauugaaagucagcccucgac....................................    1      1      7y1
.....................................................aucuauugaaagucaAcccucgacacaag...............................    1      1      7y1
.....................................................aucuauugaaagucagcccucUacacaag...............................    1      1      7y1
.....................................................aucuauugaaagucagccAucgacacaag...............................    1      1      7y1
```

```
  Star                                     Mature
gaccugcuucugggucgggguuucguacguagcagagcagcucccucgcugcgaucuauugaaagucagcccucgacacaaggguuuguccgcgcgcgcgcgcgcgcgugc
.....................................................Nucuauugaaagucagcccucgacacaag.............................    1     1    7y1
.....................................................aAcuauugaaagucagcccucgacacaag.............................    1     1    7y1
.....................................................aucuauugaaagucagcccAcgacacaag.............................    1     1    7y1
.....................................................aucuauugaaagucagcccucgCcacaag.............................    1     1    7y1
.....................................................aucuaAugaaagucagcccucgacacaag.............................    1     1    7y1
.....................................................aucuauugaaagucCgcccucgacacaag.............................    1     1    7y1
.....................................................Uucuauugaaagucagcccucgacacaag.............................    1     1    7y1
.....................................................aucuauugaaagucagcccucgacUcaag.............................    1     1    7y1
.....................................................auAuauugaaagucagcccucgacacaag.............................    4     1    7y1
.....................................................aucuauugaaagucagcccucgacacaag.............................    494   0    7y1
.....................................................aucAauugaaagucagcccucgacacaag.............................    1     1    7y1
.....................................................aucuauugaaagucagcGcucgacacaag.............................    1     1    7y1
.....................................................aucuauugaaagucagcccucgacaGaag.............................    1     1    7y1
.....................................................aucuauuUaaagucagcccucgacacaag.............................    1     1    7y1
......................................................ucuauugaaagucagGcc.......................................    1     1    7y1
......................................................ucuauugaaagucagccA.......................................    6     1    7y1
......................................................ucuauugaaagucaUccc.......................................    2     1    7y1
......................................................ucuauugaaagucagccc.......................................    1308  0    7y1
......................................................ucuauugaaaCucagccc.......................................    1     1    7y1
......................................................ucuauugaaagucaAccc.......................................    1     1    7y1
......................................................ucuauugaaagucCgccc.......................................    1     1    7y1
......................................................ucuauuUaaagucagccc.......................................    3     1    7y1
......................................................ucuauugaaaguAagccc.......................................    3     1    7y1
......................................................ucuauuAaaagucagccc.......................................    2     1    7y1
......................................................ucuaAugaaagucagccc.......................................    3     1    7y1
......................................................Gcuauugaaagucagccc.......................................    2     1    7y1
......................................................ucuauugaaagucagcAc.......................................    1     1    7y1
......................................................ucuGuugaaagucagccc.......................................    1     1    7y1
......................................................ucuauugaaagucagUcc.......................................    4     1    7y1
......................................................ucuauAgaaagucagccc.......................................    3     1    7y1
......................................................ucuauugaGagucagccc.......................................    227   1    7y1
......................................................uAuauugaaagucagccc.......................................    5     1    7y1
......................................................ucuauugaaagucagccG.......................................    1     1    7y1
......................................................ucAauugaaagucagccc.......................................    3     1    7y1
......................................................Acuauugaaagucagccc.......................................    16    1    7y1
......................................................ucuauugaaagucagccAu......................................    5     1    7y1
......................................................ucuauugaaagGcagcccu......................................    1     1    7y1
......................................................ucuaAugaaagucagcccu......................................    1     1    7y1
......................................................ucuauugaaagucagcAcu......................................    3     1    7y1
......................................................Ncuauugaaagucagcccu......................................    1     1    7y1
......................................................ucuauugaaagucCgcccu......................................    1     1    7y1
......................................................ucGauugaaagucagcccu......................................    1     1    7y1
......................................................Acuauugaaagucagcccu......................................    6     1    7y1
......................................................ucuauAgaaagucagcccu......................................    3     1    7y1
......................................................ucuauugaaagucagcccG......................................    1     1    7y1
......................................................ucAauugaaagucagcccu......................................    1     1    7y1
......................................................ucuauugaaagucagUccu......................................    1     1    7y1
......................................................ucuauugaaagucagcccA......................................    1     1    7y1
......................................................ucuauuAaaagucagcccu......................................    1     1    7y1
......................................................ucuauugaaagucagcccu......................................    861   0    7y1
......................................................ucuauugaaaCucagcccu......................................    1     1    7y1
......................................................ucAauugaaagucagcccuc.....................................    1     1    7y1
......................................................ucCauugaaagucagcccuc.....................................    278   1    7y1
......................................................ucuauugaaagucCgcccucg....................................    1     1    7y1
......................................................ucAauugaaagucagcccucg....................................    1     1    7y1
......................................................ucuauugaaagucagccGucg....................................    1     1    7y1
......................................................ucuauAgaaagucagcccucg....................................    3     1    7y1
......................................................ucuauCgaaagucagcccucg....................................    1     1    7y1
......................................................ucuauugaaaUucagcccucg....................................    1     1    7y1
......................................................ucuauugaaagucagUccucg....................................    1     1    7y1
......................................................Gcuauugaaagucagcccucg....................................    1     1    7y1
......................................................ucCauugaaagucagcccucg....................................    282   1    7y1
......................................................ucuaAugaaagucagcccucg....................................    2     1    7y1
......................................................uAuauugaaagucagcccucg....................................    1     1    7y1
......................................................Ccuauugaaagucagcccucg....................................    1     1    7y1
......................................................ucuauugaaagucagcccucU....................................    1     1    7y1
......................................................ucuauugaaaguAagcccucg....................................    2     1    7y1
......................................................ucuauugaaagucagcAcucg....................................    3     1    7y1
......................................................ucuauugaaagucaCcccucg....................................    1     1    7y1
......................................................ucuauugaaagucagcUcucg....................................    2     1    7y1
```

### Star Mature

```
gaccugcuucugggucgggguuucguacguagcagagcagcucccucgcugcgaucuauugaaagucagccccucgacacaaggguuuguccgcgcgcgcgcgcgcgcgcgugc
......................................................Acuauugaaagucagcccucg.......................................   9     1   7y1
......................................................ucuauugaaagucagccAucg.......................................   2     1   7y1
......................................................ucuauugaaagucagcccucA.......................................   1     1   7y1
......................................................ucuauugaaagucagcccucg.......................................   1220  0   7y1
......................................................ucuauugaaagucagcUcucga......................................   1     1   7y1
......................................................Gcuauugaaagucagcccucga......................................   2     1   7y1
......................................................ucuauugaaGgucagcccucga......................................   1     1   7y1
......................................................ucAauugaaagucagcccucga......................................   1     1   7y1
......................................................uAuauugaaagucagcccucga......................................   3     1   7y1
......................................................Acuauugaaagucagcccucga......................................   3     1   7y1
......................................................ucuaCugaaagucagcccucga......................................   1     1   7y1
......................................................ucuaAugaaagucagcccucga......................................   1     1   7y1
......................................................ucuauuUaaagucagcccucga......................................   1     1   7y1
......................................................ucuauugaaagucagcccucga......................................   428   0   7y1
......................................................ucuauugaaagucagcccucgG......................................   2     1   7y1
......................................................ucuauugaaaguAagcccucga......................................   1     1   7y1
......................................................ucuauAgaaagucagcccucga......................................   2     1   7y1
......................................................Gcuauugaaagucagcccucgac.....................................   1     1   7y1
......................................................ucuauugaUagucagcccucgac.....................................   1     1   7y1
......................................................uAuauugaaagucagcccucgac.....................................   1     1   7y1
......................................................ucuauugaaagucagcccucgac.....................................   299   0   7y1
......................................................ucuauugaaagucagccAucgac.....................................   1     1   7y1
......................................................ucAauugaaagucagcccucgac.....................................   1     1   7y1
......................................................ucuauugaaagucagcccAcgac.....................................   1     1   7y1
......................................................Acuauugaaagucagcccucgac.....................................   2     1   7y1
.......................................................cuauCgaaagucagcccu.........................................   1     1   7y1
.......................................................cuauugaaagucagcccG.........................................   1     1   7y1
.......................................................cuauugaaagucagcccu.........................................   416   0   7y1
.......................................................cuaAugaaagucagcccu.........................................   1     1   7y1
.......................................................cAauugaaagucagcccu.........................................   2     1   7y1
.......................................................cuauugaaaAucagcccu.........................................   1     1   7y1
.......................................................Auauugaaagucagcccu.........................................   1     1   7y1
.......................................................cuauugaaaguAagcccu.........................................   1     1   7y1
.......................................................cuauugaaaCucagcccu.........................................   1     1   7y1
.......................................................cuauugaaagAcagcccuc........................................   2     1   7y1
.......................................................cuauugaaagucagcccuc........................................   186   0   7y1
.......................................................cCauugaaagucagcccuc........................................   488   1   7y1
.......................................................cuauugaUagucagcccuc........................................   1     1   7y1
.......................................................cAauugaaagucagcccuc........................................   5     1   7y1
.......................................................Guauugaaagucagcccuc........................................   1     1   7y1
.......................................................cCauugaaagucagcccucg.......................................   701   1   7y1
.......................................................cAauugaaagucagcccucg.......................................   4     1   7y1
.......................................................cAauugaaagucagcccucgac.....................................   1     1   7y1
.......................................................cuauugaaagucagcccucgac.....................................   273   0   7y1
.......................................................cuauugaaagucagcAcucgac.....................................   1     1   7y1
.......................................................cuauugaaagucGgcccucgac.....................................   1     1   7y1
.......................................................cuauugaaagucagccAucgac.....................................   3     1   7y1
.......................................................Auauugaaagucagcccucgac.....................................   1     1   7y1
.......................................................cuauugaaagAcagcccucgac.....................................   1     1   7y1
.......................................................cuauugaaagucagcccucgacacaaggguuu...........................   1     0   7y1
........................................................uauugaaaUucagcccucg.......................................   1     1   7y1
........................................................Aauugaaagucagcccucg.......................................   6     1   7y1
........................................................uauugaaagucagcccucC.......................................   1     1   7y1
........................................................uauugaaagucagcGcucg.......................................   1     1   7y1
........................................................uauugaaagucagcccAcg.......................................   2     1   7y1
........................................................uauugaaagucagcccucg.......................................   490   0   7y1
........................................................uauugaaGgucagcccucg.......................................   1     1   7y1
........................................................uauugaaagucagcccuAg.......................................   1     1   7y1
........................................................uauugaaagAcagcccucg.......................................   1     1   7y1
........................................................uauugCaagucagcccucga......................................   1     1   7y1
........................................................uauugaaagucagcccucgG......................................   2     1   7y1
........................................................uauugaaagucaCcccucga......................................   1     1   7y1
........................................................uauugaaUgucagcccucga......................................   1     1   7y1
........................................................Aauugaaagucagcccucga......................................   2     1   7y1
........................................................Gauugaaagucagcccucga......................................   1     1   7y1
........................................................uauugaaagucagcccucga......................................   197   0   7y1
........................................................uauugaaagucagcccucgac.....................................   549   0   7y1
........................................................uaAugaaagucagcccucgac.....................................   4     1   7y1
........................................................uauugaaagAcagcccucgac.....................................   1     1   7y1
........................................................uauugaaaUucagcccucgac.....................................   1     1   7y1
```

**Star** **Mature**

```
gaccugcuucugggucgggguuucguacguagcagagcagcucccucgcugcgaucuauugaaagucagcccucgacacaaggguuuguccgcgcgcgcgcgcgcgcgugc
.........................................................uauugaaagucagcAcucgac..................................  1    1  7y1
.........................................................uauAgaaagucagcccucgac..................................  2    1  7y1
.........................................................Aauugaaagucagcccucgac..................................  11   1  7y1
.........................................................uauugaaagucagcccucUac..................................  1    1  7y1
.........................................................uauugaaagucagcccCcgac..................................  1    1  7y1
.........................................................uauugaaagucaCcccucgac..................................  1    1  7y1
.........................................................uauugaaagucaUcccucgac..................................  1    1  7y1
.........................................................uauugaaagucagAccucgac..................................  1    1  7y1
.........................................................uauugaaagucagcccucgaGacaaggguu.........................  1    1  7y1
.........................................................uauuCaaagucagcccucgacacaaggguu.........................  1    1  7y1
.........................................................uauugaaaguAagcccucgacacaaggguu.........................  1    1  7y1
.........................................................uauugaaagucagcAcucgacacaaggguu.........................  2    1  7y1
.........................................................uauugaaagucagcccucgacacaaggAuu.........................  1    1  7y1
.........................................................Aauugaaagucagcccucgacacaaggguu.........................  1    1  7y1
.........................................................uauugaaagucagcccucgacacaaggguu.........................  155  0  7y1
.........................................................uauugaaagucagcccAcgacacaaggguu.........................  1    1  7y1
.........................................................uauugaaaUucagcccucgacacaaggguu.........................  1    1  7y1
.........................................................uauugaaagucagcccucgacacaaggCuu.........................  1    1  7y1
.........................................................uauugaaagucagcccucgacacaaggguG.........................  1    1  7y1
.........................................................uauugaaagucagcccucgacaUaaggguuu........................  1    1  7y1
.........................................................uauugaaagucagcccucgacacaagggAuu........................  1    1  7y1
.........................................................uauugaaagucagAccucgacacaaggguuu........................  1    1  7y1
.........................................................uauugaaagucagcccucgacacaaggguuu........................  220  0  7y1
.........................................................uauugaaagucagcccAcgacacaaggguuu........................  1    1  7y1
.........................................................Aauugaaagucagcccucgacacaaggguuu........................  7    1  7y1
.........................................................uauugaaagucagccAucgacacaaggguuug.......................  1    1  7y1
.........................................................uauugaaagucagcccuAgacacaaggguuug.......................  3    1  7y1
.........................................................uauugaaagucagcccucgacacaaCgguuug.......................  1    1  7y1
.........................................................uauugaaagucagcccucgacacaagggAuug.......................  2    1  7y1
.........................................................uauugaaagucagcccucgacaAaaggguuug.......................  3    1  7y1
.........................................................Nauugaaagucagcccucgacacaaggguuug.......................  1    1  7y1
.........................................................uauugaaagucagcccucgacacaagggGuug.......................  1    1  7y1
.........................................................Gauugaaagucagcccucgacacaaggguuug.......................  2    1  7y1
.........................................................uauugaaaguAagcccucgacacaaggguuug.......................  2    1  7y1
.........................................................uauugaaagucagcccucgacacaaggguuAg.......................  1    1  7y1
.........................................................uauugaaagucagcccucgacacaagCguuug.......................  1    1  7y1
.........................................................uauugaaagucagGccucgacacaaggguuug.......................  1    1  7y1
.........................................................uauugaaagucGgcccucgacacaaggguuug.......................  1    1  7y1
.........................................................uauuCaaagucagcccucgacacaaggguuug.......................  1    1  7y1
.........................................................uauugaaagucagcccCcgacacaaggguuug.......................  1    1  7y1
.........................................................uauugaaagucagcccAcgacacaaggguuug.......................  2    1  7y1
.........................................................uauugaaagucagcGcucgacacaaggguuug.......................  1    1  7y1
.........................................................uauugaaagucaCcccucgacacaaggguuug.......................  2    1  7y1
.........................................................uauugaaagucagcccucgacacaagUguuug.......................  1    1  7y1
.........................................................uauugaaagucagcAcucgacacaaggguuug.......................  6    1  7y1
.........................................................uauugaaagucagAccucgacacaaggguuug.......................  1    1  7y1
.........................................................uauugaaaUucagcccucgacacaaggguuug.......................  1    1  7y1
.........................................................uauugaaagucagcccucgacacaaggguuug.......................  684  0  7y1
.........................................................uauuAaaagucagcccucgacacaaggguuug.......................  1    1  7y1
.........................................................Aauugaaagucagcccucgacacaaggguuug.......................  13   1  7y1
..........................................................auugaaagucagccAucg....................................  1    1  7y1
..........................................................auuUaaagucagcccucg....................................  1    1  7y1
..........................................................auAgaaagucagcccucg....................................  1    1  7y1
..........................................................auuCaaagucagcccucg....................................  1    1  7y1
..........................................................auugaaagucagcccucg....................................  487  0  7y1
..........................................................aAugaaagucagcccucg....................................  2    1  7y1
..........................................................auugaGagucagcccucg....................................  1    1  7y1
..........................................................auugaaaguAagcccucg....................................  2    1  7y1
..........................................................auugaaagucagUccucg....................................  1    1  7y1
..........................................................Uuugaaagucagcccucg....................................  1    1  7y1
..........................................................auugaaagucagcAcucg....................................  2    1  7y1
..........................................................auugaaagucagAccucg....................................  3    1  7y1
..........................................................auugaaagucagcccucC....................................  1    1  7y1
..........................................................auugaaagucagcccuAga...................................  2    1  7y1
..........................................................auugaaagucaUcccucga...................................  1    1  7y1
..........................................................auugaaagucagcccucgC...................................  1    1  7y1
..........................................................auugaaagucagAccucga...................................  1    1  7y1
..........................................................auugaaagucagcccAcga...................................  1    1  7y1
..........................................................auugaaagucagccAucga...................................  1    1  7y1
..........................................................auugaaagAcagcccucga...................................  1    1  7y1
```

**Star** **Mature**

```
gaccugcuucugggucggggguucguacguagcagagcagcucccucgcugcgaucuauugaaagucagcccucgacacaaggguuuguccgcgcgcgcgcgcgcgcgcgcgugc
.........................................................auugaaaguGagcccucga.......................................  1    1  7y1
.........................................................auugaaagucagcccucga.......................................  452  0  7y1
.........................................................aAugaaagucagcccucga.......................................  1    1  7y1
.........................................................auugaaagucagAccucgac......................................  1    1  7y1
.........................................................auugaaagucagccAucgac......................................  3    1  7y1
.........................................................aAugaaagucagcccucgac......................................  1    1  7y1
.........................................................auugaaagucagcccucgac......................................  419  0  7y1
.........................................................auugaaagucagUccucgac......................................  2    1  7y1
.........................................................auugaaagucagcccucgCc......................................  2    1  7y1
.........................................................auugaaagAcagcccucgac......................................  2    1  7y1
.........................................................auugaaaguAagcccucgac......................................  2    1  7y1
.........................................................auugaaagucagcccucgaca.....................................  189  0  7y1
.........................................................aAugaaagucagcccucgaca.....................................  2    1  7y1
.........................................................auugaaagucagcccucgaGa.....................................  1    1  7y1
.........................................................auugaaaguAagcccucgaca.....................................  2    1  7y1
.........................................................auugaaagucagcccucgacG.....................................  1    1  7y1
.........................................................auugaaagucagcAcucgacaca...................................  1    1  7y1
.........................................................auugaaaguAagcccucgacaca...................................  1    1  7y1
.........................................................auugaaagucagcccucgacaca...................................  165  0  7y1
.........................................................auugaaagucagcccucgCcaca...................................  4    1  7y1
.........................................................aAugaaagucagcccucgacaca...................................  1    1  7y1
.........................................................auugaaagucagcccucgUcaca...................................  1    1  7y1
.........................................................Uuugaaagucagcccucgacaca...................................  1    1  7y1
.........................................................auugaaagucagccAucgacaca...................................  2    1  7y1
.........................................................auugaaagucagcccAcgacacaag.................................  1    1  7y1
.........................................................auuCaaagucagcccucgacacaag.................................  1    1  7y1
.........................................................auugaaagucagccAucgacacaag.................................  2    1  7y1
.........................................................Uuugaaagucagcccucgacacaag.................................  1    1  7y1
.........................................................auugaaagucagcccucgCcacaag.................................  1    1  7y1
.........................................................auugaaaguAagcccucgacacaag.................................  1    1  7y1
.........................................................auugaaagucUgcccucgacacaag.................................  1    1  7y1
.........................................................auugaaaCucagcccucgacacaag.................................  1    1  7y1
.........................................................auugaaagucagcccucgacacaag.................................  209  0  7y1
.........................................................auugaaagucagcccucgacacaaC.................................  1    1  7y1
.........................................................Guugaaagucagcccucgacacaag.................................  2    1  7y1
.........................................................auugaaagucagcccucCacacaag.................................  1    1  7y1
.........................................................auugaaagucagcccucgacacaaggguu.............................  1    0  7y1
.........................................................auugaaaUucagcccucgacacaaggguuug...........................  1    1  7y1
.........................................................auugaaagucagccAucgacacaaggguuug...........................  1    1  7y1
.........................................................auugaaagucagcccucgacacaaggguAug...........................  1    1  7y1
.........................................................auugaaagucagcccuAgacacaaggguuug...........................  1    1  7y1
.........................................................auugaaagucagcccucgacacaaggguuAg...........................  2    1  7y1
.........................................................auugaaagucagcccAcgacacaaggguuug...........................  1    1  7y1
.........................................................auugaaagucagcccucgacacaaggguuug...........................  187  0  7y1
.........................................................auugaaagucagcccucgacacaagggGuug...........................  2    1  7y1
.........................................................auugaaagucagAccucgacacaaggguuug...........................  2    1  7y1
.........................................................auugaaagucagcccucgacGcaaggguuugu..........................  3    1  7y1
.........................................................auugaaagucagcccucgacaGaaggguuugu..........................  1    1  7y1
.........................................................auugaaagucagcccucgacacaaCgguuugu..........................  2    1  7y1
.........................................................auugaaagucagcccucUacacaaggguuugu..........................  1    1  7y1
.........................................................auugaaagucagccGucgacacaaggguuugu..........................  1    1  7y1
.........................................................auugaaagucaUcccucgacacaaggguuugu..........................  4    1  7y1
.........................................................auugaGagucagcccucgacacaaggguuugu..........................  2    1  7y1
.........................................................auugaaagucagcccucgacacaagCguuugu..........................  11   1  7y1
.........................................................auugaaagucagcUcucgacacaaggguuugu..........................  3    1  7y1
.........................................................Nuugaaagucagcccucgacacaaggguuugu..........................  4    1  7y1
.........................................................auugaaagAcagcccucgacacaaggguuugu..........................  6    1  7y1
.........................................................auugaaagucagcccucgacacaaggguuugG..........................  12   1  7y1
.........................................................auugaaaguUagcccucgacacaaggguuugu..........................  2    1  7y1
.........................................................auugaaagucagcccucgacacaGggguuugu..........................  2    1  7y1
.........................................................auugaaagucagcccucgacacaaggguGugu..........................  5    1  7y1
.........................................................auugaaagucCgcccucgacacaaggguuugu..........................  2    1  7y1
.........................................................aAugaaagucagcccucgacacaaggguuugu..........................  23   1  7y1
.........................................................auugaaagucagcccucgaAacaaggguuugu..........................  2    1  7y1
.........................................................auugaaagucagcccucgaUacaaggguuugu..........................  211  1  7y1
.........................................................auuAaaagucagcccucgacacaaggguuugu..........................  1    1  7y1
.........................................................auugaaagucagcccuAgacacaaggguuugu..........................  9    1  7y1
.........................................................auugaaagucagcccucgacaAaaggguuugu..........................  2    1  7y1
.........................................................Guugaaagucagcccucgacacaaggguuugu..........................  6    1  7y1
.........................................................auugaaagucagcccucgacacaaggguuugC..........................  1    1  7y1
```

Star Mature

gaccugcuucugggucgggguuucguacguagcagagcagcucccucgcugcgaucuauugaaagucagcccucgacacaaggguuuguccgcgcgcgcgcgcgcgcgugc

| | | | |
|---|---|---|---|
| .........................................................auugGaagucagcccucgacacaaggguuugu...................... | 1 | 1 | 7y1 |
| .........................................................auugaaagucagcccucgacacUaggguuugu...................... | 1 | 1 | 7y1 |
| .........................................................auugaaagucagcccucgaGacaaggguuugu...................... | 2 | 1 | 7y1 |
| .........................................................auuUaaagucagcccucgacacaaggguuugu...................... | 1 | 1 | 7y1 |
| .........................................................auugaaagucagcccucgCcacaaggguuugu...................... | 19 | 1 | 7y1 |
| .........................................................Uuugaaagucagcccucgacacaaggguuugu...................... | 8 | 1 | 7y1 |
| .........................................................auAgaaagucagcccucgacacaaggguuugu...................... | 4 | 1 | 7y1 |
| .........................................................auugaaagucagcccucgacacaaggUuuugu...................... | 3 | 1 | 7y1 |
| .........................................................auugaaagucagcccucgacacaaggguuAgu...................... | 10 | 1 | 7y1 |
| .........................................................auugaaagucagcccucgacacaaggguuugA...................... | 19 | 1 | 7y1 |
| .........................................................auugaaagucagcccucgacacaagggAuugu...................... | 28 | 1 | 7y1 |
| .........................................................auugaaagucagcccucgacacCaggguuugu...................... | 1 | 1 | 7y1 |
| .........................................................auugaaagucagcccucgacacaaggguuuCu...................... | 11 | 1 | 7y1 |
| .........................................................auugaaagucagcccucgacacaaUgguuugu...................... | 5 | 1 | 7y1 |
| .........................................................auugaaagucagcccucgacCcaaggguuugu...................... | 1 | 1 | 7y1 |
| .........................................................auugaaagucagccUucgacacaaggguuugu...................... | 1 | 1 | 7y1 |
| .........................................................auugaaagCcagcccucgacacaaggguuugu...................... | 1 | 1 | 7y1 |
| .........................................................auugaaagucagcAcucgacacaaggguuugu...................... | 22 | 1 | 7y1 |
| .........................................................auugaaagucagAccucgacacaaggguuugu...................... | 16 | 1 | 7y1 |
| .........................................................auugaaagucagcccAcgacacaaggguuugu...................... | 13 | 1 | 7y1 |
| .........................................................auugaaagucagcccucgacacaaggguAugu...................... | 18 | 1 | 7y1 |
| .........................................................auugaaagucagcccucgacacaagggCuugu...................... | 1 | 1 | 7y1 |
| .........................................................auugaaagucagcccucgacacaaggAuuugu...................... | 1 | 1 | 7y1 |
| .........................................................auugaaagucagcccucgacacaaggguuugu...................... | 7347 | 0 | 7y1 |
| .........................................................auugaaaCucagcccucgacacaaggguuugu...................... | 4 | 1 | 7y1 |
| .........................................................auGgaaagucagcccucgacacaaggguuugu...................... | 1 | 1 | 7y1 |
| .........................................................auugCaagucagcccucgacacaaggguuugu...................... | 1 | 1 | 7y1 |
| .........................................................auugaaagucagcccucgacacaaggCuuugu...................... | 13 | 1 | 7y1 |
| .........................................................auugaaagucagcccucgacacaaggguuuUu...................... | 7 | 1 | 7y1 |
| .........................................................auugaaagucagcGcucgacacaaggguuugu...................... | 1 | 1 | 7y1 |
| .........................................................auuCaaagucagcccucgacacaaggguuugu...................... | 1 | 1 | 7y1 |
| .........................................................auugaaaguAagcccucgacacaaggguuugu...................... | 13 | 1 | 7y1 |
| .........................................................auugaaagucagcccucgUcacaaggguuugu...................... | 1 | 1 | 7y1 |
| .........................................................auugaaagucagcccucgacacaagUguuugu...................... | 3 | 1 | 7y1 |
| .........................................................auugaaagucGgcccucgacacaaggguuugu...................... | 1 | 1 | 7y1 |
| .........................................................auugaaagucaAcccucgacacaaggguuugu...................... | 1 | 1 | 7y1 |
| .........................................................auugaaagucagcccucgacacaagggGuugu...................... | 12 | 1 | 7y1 |
| .........................................................auugaaagucagcccucCacacaaggguuugu...................... | 2 | 1 | 7y1 |
| .........................................................auugaaagucUgcccucgacacaaggguuugu...................... | 1 | 1 | 7y1 |
| .........................................................auCgaaagucagcccucgacacaaggguuugu...................... | 1 | 1 | 7y1 |
| .........................................................auugaaagucagccAucgacacaaggguuugu...................... | 15 | 1 | 7y1 |
| .........................................................auugaaaguGagcccucgacacaaggguuugu...................... | 4 | 1 | 7y1 |
| .........................................................auugaaagucagcccucgacacGaggguuugu...................... | 2 | 1 | 7y1 |
| .........................................................aCugaaagucagcccucgacacaaggguuugu...................... | 1 | 1 | 7y1 |
| ..........................................................uugaaagucagcccucga................................... | 208 | 0 | 7y1 |
| ..........................................................Augaaagucagcccucga................................... | 4 | 1 | 7y1 |
| ..........................................................uugaaagucagccAucga................................... | 3 | 1 | 7y1 |
| ..........................................................uAgaaagucagcccucga................................... | 1 | 1 | 7y1 |
| ..........................................................uugaaaguAagcccucgac.................................. | 1 | 1 | 7y1 |
| ..........................................................Augaaagucagcccucgac.................................. | 4 | 1 | 7y1 |
| ..........................................................uugaaagucagcAcucgac.................................. | 1 | 1 | 7y1 |
| ..........................................................uuUaaagucagcccucgac.................................. | 1 | 1 | 7y1 |
| ..........................................................uugaaagucagcccucgac.................................. | 211 | 0 | 7y1 |
| ..........................................................uugaaagucagcAcucgacacaaggguuug....................... | 1 | 1 | 7y1 |
| ..........................................................uugaaagucagcccucgacacaagUguuug....................... | 1 | 1 | 7y1 |
| ..........................................................uugaaagucagcccuAgacacaaggguuug....................... | 1 | 1 | 7y1 |
| ..........................................................uugaaagucagcccucgacacaaggguuug....................... | 182 | 0 | 7y1 |
| ..........................................................uugaaagucagGccucgacacaaggguuug....................... | 1 | 1 | 7y1 |
| ..........................................................uugaaagucagccAucgacacaaggguuug....................... | 1 | 1 | 7y1 |
| ..........................................................uugaaaguAagcccucgacacaaggguuug....................... | 1 | 1 | 7y1 |
| ..........................................................uAgaaagucagcccucgacacaaggguuug....................... | 1 | 1 | 7y1 |
| ..........................................................uugaaagucagcccucgacacaaCgguuug....................... | 1 | 1 | 7y1 |
| ..........................................................uugaaagucagcccucgacUcaaggguuugu...................... | 1 | 1 | 7y1 |
| ..........................................................uugaaagucagcccucgacacaagggAuugu...................... | 6 | 1 | 7y1 |
| ..........................................................uugaaagucagAccucgacacaaggguuugu...................... | 2 | 1 | 7y1 |
| ..........................................................uugaaagucagcccucgacacaaggguuuCu...................... | 3 | 1 | 7y1 |
| ..........................................................uugaaagucagcccucgacacaaggguuugG...................... | 3 | 1 | 7y1 |
| ..........................................................uugaaagucagcccucgCcacaaggguuugu...................... | 4 | 1 | 7y1 |
| ..........................................................uugaaaguAagcccucgacacaaggguuugu...................... | 4 | 1 | 7y1 |
| ..........................................................uugaaagucagcccucgacacaaggguuAgu...................... | 6 | 1 | 7y1 |

**Star** **Mature**

```
gaccugcuucugggucgggguuucguacguagcagagcagcucccucgcugcgaucuauugaaagucagcccucgacacaaggguuuguccgcgcgcgcgcgcgcgcgugc
..........................................................uugaaagucagcccucgacacaagCguuugu......................    1      1      7y1
..........................................................uugaaagucagcAcucgacacaaggguuugu......................    1      1      7y1
..........................................................Gugaaagucagcccucgacacaaggguuugu......................    3      1      7y1
..........................................................uugaaagucagccAucgacacaaggguuugu......................    3      1      7y1
..........................................................uugaaagucagcccucgacacaaggguuugu......................    1439   0      7y1
..........................................................uugaaagucagcccuAgacacaaggguuugu......................    2      1      7y1
..........................................................uugaaagucagcccucgacacaagUguuugu......................    1      1      7y1
..........................................................Augaaagucagcccucgacacaaggguuugu......................    10     1      7y1
..........................................................uugaaagucagcccucgacacaaggguuuUu......................    2      1      7y1
..........................................................uugaaagucagcccucgacacaaggCuuugu......................    1      1      7y1
..........................................................uugaaagucagGccucgacacaaggguuugu......................    1      1      7y1
..........................................................uugaaagucagcccucgacacaaUgguuugu......................    1      1      7y1
..........................................................uugaaagucagccUucgacacaaggguuugu......................    1      1      7y1
..........................................................uugaaagucagcccucgacacaaggguAugu......................    4      1      7y1
..........................................................uugaaagucGgcccucgacacaaggguuugu......................    1      1      7y1
..........................................................uuCaaagucagcccucgacacaaggguuugu......................    2      1      7y1
..........................................................uugaaagucagcccucgacacaaggguGugu......................    1      1      7y1
..........................................................uugaaaCucagcccucgacacaaggguuugu......................    1      1      7y1
..........................................................uugaaagAcagcccucgacacaaggguuugu......................    1      1      7y1
..........................................................uugaaagucagcccucgacacaaggguuugA......................    4      1      7y1
..........................................................uAgaaagucagcccucgacacaaggguuugu......................    2      1      7y1
..........................................................uugaaagucagcccucgacacaagggGuugu......................    3      1      7y1
..........................................................uugaaagucagcccucgacacaaCgguuugu......................    2      1      7y1
..........................................................uCgaaagucagcccucgacacaaggguuugu......................    2      1      7y1
...........................................................Agaaagucagcccucgac..................................    4      1      7y1
...........................................................ugaaagAcagcccucgac..................................    1      1      7y1
...........................................................ugaaaguAagcccucgac..................................    2      1      7y1
...........................................................ugaaagucagcccucgac..................................    191    0      7y1
...........................................................ugaaagucagUccucgac..................................    1      1      7y1
...........................................................ugaaagucagcccucgCc..................................    1      1      7y1
...........................................................ugaaagucagcccucCacacaaggguuug.......................    1      1      7y1
...........................................................ugaaagucagcccucgacacaagggAuug.......................    2      1      7y1
...........................................................ugaaagucagcccGcgacacaaggguuug.......................    1      1      7y1
...........................................................ugaaagAcagcccucgacacaaggguuug.......................    1      1      7y1
...........................................................ugaaagucagccGucgacacaaggguuug.......................    1      1      7y1
...........................................................ugaaagucagcccucgacacaaggUuuug.......................    1      1      7y1
...........................................................ugaaagucagcccucgCcacaaggguuug.......................    1      1      7y1
...........................................................ugaaagucagcccucgacacaaUgguuug.......................    1      1      7y1
...........................................................uAaaagucagcccucgacacaaggguuug.......................    1      1      7y1
...........................................................ugaaagucagcccucgacacGaggguuug.......................    1      1      7y1
...........................................................ugaaagucagcccucgacacaaggguuug.......................    462    0      7y1
...........................................................ugaaagucagAccucgacacaaggguuug.......................    1      1      7y1
...........................................................ugaaagucagcccucgacacaaggguAug.......................    1      1      7y1
...........................................................ugaaagucagcccucgacacaaggguGug.......................    1      1      7y1
...........................................................Ggaaagucagcccucgacacaaggguuug.......................    1      1      7y1
...........................................................ugaaagucUgcccucgacacaaggguuug.......................    1      1      7y1
...........................................................ugaaagucagcccucgacacaaggguuGg.......................    2      1      7y1
............................................................gaaagucagcccucgacaUaaggguuugu......................    1      1      7y1
............................................................gaaagucagcccucgacCcaaggguuugu......................    1      1      7y1
............................................................gaaagucagAccucgacacaaggguuugu......................    1      1      7y1
............................................................gaaagucagcccAcgacacaaggguuugu......................    1      1      7y1
............................................................gaaagucagcccucgacacaaggguuAgu......................    1      1      7y1
............................................................Caaagucagcccucgacacaaggguuugu......................    1      1      7y1
............................................................gaaagucagcccucgacacaaggguuugu......................    170    0      7y1
............................................................gaGagucagcccucgacacaaggguuugu......................    1      1      7y1
............................................................gaaagucagcccucgacacaagggGuugu......................    1      1      7y1
..............................................................aagucagcccucgacacaaggguuuC.......................    2      1      7y1
..............................................................aagucagcccucgacacaaggguuug.......................    211    0      7y1
..............................................................aagucagcccucgacUcaaggguuug.......................    2      1      7y1
..............................................................aagucagccAucgacacaaggguuug.......................    2      1      7y1
..............................................................aagucagcccucgacacaaggUuuug.......................    1      1      7y1
.................................................................uAagcccucgacacaaggguuug.......................    1      1      7y1
.................................................................ucagcccucgGcacaaggguuug.......................    1      1      7y1
.................................................................Acagcccucgacacaaggguuug.......................    3      1      7y1
.................................................................Ncagcccucgacacaaggguuug.......................    1      1      7y1
.................................................................ucagccAucgacacaaggguuug.......................    1      1      7y1
.................................................................ucagcccucgacacaaggguuug.......................    244    0      7y1
.................................................................ucagcccucgacacaagggAuug.......................    1      1      7y1
.................................................................Gcagcccucgacacaaggguuug.......................    1      1      7y1
..................................................................cagcUcucgacacaaggguuugu......................    211    1      7y1
```

**Star** **Mature**

```
gaccugcuucugggucgggguuucguacguagcagagcagcucccucgcugcgaucuauugaaagucagcccucgacacaaggguuuguccgcgcgcgcgcgcgcgcgugc
...................................................................agccUucgacacaaggguuug.......................  132   1  7y1
....................................................................gcAcucgacacaaggguuug.......................  1     1  7y1
....................................................................gccAucgacacaaggguuug.......................  1     1  7y1
....................................................................Ncccucgacacaaggguuug.......................  1     1  7y1
....................................................................gAccucgacacaaggguuug.......................  2     1  7y1
....................................................................gcccucgaAacaaggguuug.......................  2     1  7y1
....................................................................gcccucgacacaaggguuug.......................  188   0  7y1
....................................................................gcccucAacacaaggguuug.......................  1     1  7y1
.....................................................................cccucgacacaaggguuuU.......................  1     1  7y1
.....................................................................ccAucgacacaaggguuugu......................  2     1  7y1
.....................................................................Nccucgacacaaggguuugu......................  1     1  7y1
.....................................................................cccucgacacaaggguuugu......................  289   0  7y1
.....................................................................cccucgacacaagggAuugu......................  2     1  7y1
.....................................................................cccucgacacaaggCuuugu......................  2     1  7y1
.....................................................................cccucgacaAaaggguuugu......................  1     1  7y1
.....................................................................cccucgacacaaggguuugG......................  2     1  7y1
.....................................................................cccucgacUcaaggguuugu......................  1     1  7y1
.....................................................................cccucgacacaaUgguuugu......................  1     1  7y1
......................................................................Gcucgacacaaggguuug.......................  4     1  7y1
......................................................................ccucgacacaagAguuug.......................  1     1  7y1
......................................................................ccucgacaUaaggguuug.......................  1     1  7y1
......................................................................ccucUacacaaggguuug.......................  3     1  7y1
......................................................................cAucgacacaaggguuug.......................  6     1  7y1
......................................................................ccucgacUcaaggguuug.......................  2     1  7y1
......................................................................ccCcgacacaaggguuug.......................  1     1  7y1
......................................................................ccucgacacaaggguAug.......................  4     1  7y1
......................................................................ccucgacacaaggguuGg.......................  1     1  7y1
......................................................................ccucgaAacaaggguuug.......................  3     1  7y1
......................................................................ccAcgacacaaggguuug.......................  1     1  7y1
......................................................................ccucgacacaagCguuug.......................  2     1  7y1
......................................................................ccucgacacaaUgguuug.......................  1     1  7y1
......................................................................Ncucgacacaaggguuug.......................  2     1  7y1
......................................................................ccuGgacacaaggguuug.......................  1     1  7y1
......................................................................Acucgacacaaggguuug.......................  8     1  7y1
......................................................................ccucgacacaaggguuAg.......................  1     1  7y1
......................................................................ccucgacacaaggguuuA.......................  1     1  7y1
......................................................................ccucgacacaaggguuuC.......................  1     1  7y1
......................................................................ccucgacacaUggguuug.......................  1     1  7y1
......................................................................ccucgacacaaggCuuug.......................  1     1  7y1
......................................................................ccucgacacaaggguuug.......................  1954  0  7y1
......................................................................ccucgCcacaaggguuug.......................  2     1  7y1
......................................................................ccucgacacGaggguuug.......................  1     1  7y1
......................................................................ccucgacacaaUgguuugu......................  5     1  7y1
......................................................................ccucgacacNaggguuugu......................  1     1  7y1
......................................................................ccucgacacaaggguAugu......................  8     1  7y1
......................................................................ccucgacacaGggguuugu......................  1     1  7y1
......................................................................ccucUacacaaggguuugu......................  7     1  7y1
......................................................................ccucgacacaaggguuuAu......................  1     1  7y1
......................................................................ccucgacacaaggguuugA......................  12    1  7y1
......................................................................ccucgacacaagUguuugu......................  6     1  7y1
......................................................................ccucgUcacaaggguuugu......................  1     1  7y1
......................................................................ccucgacaUaaggguuugu......................  153   1  7y1
......................................................................Ucucgacacaaggguuugu......................  358   1  7y1
......................................................................Acucgacacaaggguuugu......................  24    1  7y1
......................................................................ccucgacacaaggguuuCu......................  3     1  7y1
......................................................................ccucgacGcaaggguuugu......................  3     1  7y1
......................................................................ccucgacacaaggUuuugu......................  9     1  7y1
......................................................................cAucgacacaaggguuugu......................  17    1  7y1
......................................................................ccucgacacaaCgguuugu......................  3     1  7y1
......................................................................ccucgacacaaggCuuugu......................  13    1  7y1
......................................................................ccucgCcacaaggguuugu......................  7     1  7y1
......................................................................ccucgacacaaggguuugG......................  17    1  7y1
......................................................................ccuGgacacaaggguuugu......................  1     1  7y1
......................................................................Ncucgacacaaggguuugu......................  3     1  7y1
......................................................................ccucgacacaagCguuugu......................  4     1  7y1
......................................................................ccucgaUacaaggguuugu......................  1     1  7y1
......................................................................ccucgacacaaggguuugC......................  1     1  7y1
......................................................................ccAcgacacaaggguuugu......................  14    1  7y1
......................................................................ccucgaAacaaggguuugu......................  9     1  7y1
......................................................................ccucgacaAaaggguuugu......................  8     1  7y1
```

Star Mature

| | | | |
|---|---|---|---|
| gaccugcuucugggucgggguuucguacguagcagagcagcucccucgcugcgaucuauugaaagucagcccucgacacaaggguuuguccgcgcgcgcgcgcgcgcgcgcgugc | | | |
| ......................................................................ccucgacacGaggguuugu.......................... | 1 | 1 | 7y1 |
| ......................................................................ccucgacacaaggguuugu.......................... | 6344 | 0 | 7y1 |
| ......................................................................ccucgacacaaggguuuUu.......................... | 5 | 1 | 7y1 |
| ......................................................................ccucgacacaagggGuugu.......................... | 4 | 1 | 7y1 |
| ......................................................................ccucgGcacaaggguuugu.......................... | 1 | 1 | 7y1 |
| ......................................................................Gcucgacacaaggguuugu.......................... | 5 | 1 | 7y1 |
| ......................................................................ccucAacacaaggguuugu.......................... | 4 | 1 | 7y1 |
| ......................................................................ccucgacacaagggAuugu.......................... | 8 | 1 | 7y1 |
| ......................................................................ccucgacacaaggguuAgu.......................... | 9 | 1 | 7y1 |
| ......................................................................ccucgacacaaggguCugu.......................... | 1 | 1 | 7y1 |
| ......................................................................cNucgacacaaggguuuguccgcgc.................... | 1 | 1 | 7y1 |
| ......................................................................ccucgacacaaggguuuguccgcgA.................... | 1 | 1 | 7y1 |
| ......................................................................ccAcgacacaaggguuuguccgcgc.................... | 1 | 1 | 7y1 |
| ......................................................................ccucgacacaaggguuuguccgcgc.................... | 277 | 0 | 7y1 |
| ......................................................................ccucgacacaaggguAuguccgcgc.................... | 1 | 1 | 7y1 |
| ......................................................................ccucgacacaaggCuuuguccgcgc.................... | 1 | 1 | 7y1 |
| ......................................................................ccucgacacaaggUuuuguccgcgc.................... | 1 | 1 | 7y1 |
| ......................................................................ccucgacacaagggAuuguccgcgc.................... | 1 | 1 | 7y1 |
| ......................................................................ccuUgacacaaggguuuguccgcgc.................... | 1 | 1 | 7y1 |
| ......................................................................cAucgacacaaggguuuguccgcgc.................... | 1 | 1 | 7y1 |
| .......................................................................cucgacacaaggguuAgu.......................... | 30 | 1 | 7y1 |
| .......................................................................cucgacacGaggguuugu.......................... | 5 | 1 | 7y1 |
| .......................................................................cucgacacaagggGuugu.......................... | 22 | 1 | 7y1 |
| .......................................................................cucgacacaaggguCugu.......................... | 2 | 1 | 7y1 |
| .......................................................................Gucgacacaaggguuugu.......................... | 18 | 1 | 7y1 |
| .......................................................................cNcgacacaaggguuugu.......................... | 2 | 1 | 7y1 |
| .......................................................................cucAacacaaggguuugu.......................... | 13 | 1 | 7y1 |
| .......................................................................Nucgacacaaggguuugu.......................... | 10 | 1 | 7y1 |
| .......................................................................cucgacUcaaggguuugu.......................... | 9 | 1 | 7y1 |
| .......................................................................cucgacacaaggguuugA.......................... | 39 | 1 | 7y1 |
| .......................................................................cucUacacaaggguuugu.......................... | 14 | 1 | 7y1 |
| .......................................................................cucgacacaaggguuCgu.......................... | 1 | 1 | 7y1 |
| .......................................................................cucgacacaaggguuuUu.......................... | 19 | 1 | 7y1 |
| .......................................................................cucgaUacaaggguuugu.......................... | 3 | 1 | 7y1 |
| .......................................................................cucgacacaaggguuugu.......................... | 25979 | 0 | 7y1 |
| .......................................................................cuGgacacaaggguuugu.......................... | 197 | 1 | 7y1 |
| .......................................................................cucgacacaaggCuuugu.......................... | 17 | 1 | 7y1 |
| .......................................................................cucgacacaagCguuugu.......................... | 18 | 1 | 7y1 |
| .......................................................................cucgacaUaaggguuugu.......................... | 3 | 1 | 7y1 |
| .......................................................................cucgacacaagUguuugu.......................... | 175 | 1 | 7y1 |
| .......................................................................cucgacacaaggguuugC.......................... | 1 | 1 | 7y1 |
| .......................................................................cucgacacaaCgguuugu.......................... | 17 | 1 | 7y1 |
| .......................................................................cucgacGcaaggguuugu.......................... | 6 | 1 | 7y1 |
| .......................................................................cucgacacaGggguuugu.......................... | 1 | 1 | 7y1 |
| .......................................................................cucgacacaaAgguuugu.......................... | 197 | 1 | 7y1 |
| .......................................................................cucgacacaaggguuugG.......................... | 34 | 1 | 7y1 |
| .......................................................................cucgacacaaggguuGgu.......................... | 1 | 1 | 7y1 |
| .......................................................................cucgacacaaUgguuugu.......................... | 17 | 1 | 7y1 |
| .......................................................................cucgaAacaaggguuugu.......................... | 34 | 1 | 7y1 |
| .......................................................................cucgacCcaaggguuugu.......................... | 1 | 1 | 7y1 |
| .......................................................................cucgacacCaggguuugu.......................... | 1 | 1 | 7y1 |
| .......................................................................cAcgacacaaggguuugu.......................... | 75 | 1 | 7y1 |
| .......................................................................cucgacacaaggguuuCu.......................... | 23 | 1 | 7y1 |
| .......................................................................cucgaNacaaggguuugu.......................... | 1 | 1 | 7y1 |
| .......................................................................cGcgacacaaggguuugu.......................... | 2 | 1 | 7y1 |
| .......................................................................cucgacacaaggguuuAu.......................... | 7 | 1 | 7y1 |
| .......................................................................cucgacacaaggguGugu.......................... | 5 | 1 | 7y1 |
| .......................................................................cucgacacaaggguAugu.......................... | 35 | 1 | 7y1 |
| .......................................................................cCcgacacaaggguuugu.......................... | 2 | 1 | 7y1 |
| .......................................................................cucgacacaagggAuugu.......................... | 43 | 1 | 7y1 |
| .......................................................................cucgacacaaggUuuugu.......................... | 13 | 1 | 7y1 |
| .......................................................................cucgacacaaggAuuugu.......................... | 3 | 1 | 7y1 |
| .......................................................................cucgacacUaggguuugu.......................... | 2 | 1 | 7y1 |
| .......................................................................cucgaGacaaggguuugu.......................... | 2 | 1 | 7y1 |
| .......................................................................cucgCcacaaggguuugu.......................... | 4 | 1 | 7y1 |
| .......................................................................cucgacaAaaggguuugu.......................... | 21 | 1 | 7y1 |
| .......................................................................cucgacacaUggguuugu.......................... | 2 | 1 | 7y1 |
| .......................................................................cucgacaGaaggguuugu.......................... | 1 | 1 | 7y1 |
| .......................................................................cucgUcacaaggguuugu.......................... | 2 | 1 | 7y1 |
| .......................................................................cucgacacaaggguuuguccgcgc.................... | 1 | 0 | 7y1 |

**Star** **Mature**

gaccugcuucugggucgggguuucguacguagcagagcagcucccucgcugcgaucuauugaaagucagcccucgacacaaggguuuguccgcgcgcgcgcgcgcgcgcgugc
