## Supplementary material for "SEA: The small RNA Expression Atlas": p-hsa-miR-235-2

| | | |
|---|---|---|
| Provisional ID | : | chr22_KI270733v1_random_12920 |
| Score total | : | 144078.7 |
| Score for star read(s) | : | 3.9 |
| Score for read counts | : | 144071.6 |
| Score for mfe | : | 2.2 |
| Score for randfold | : | 1.6 |
| Score for cons. seed | : | -0.6 |
| Total read count | : | 282602 |
| Mature read count | : | 36506 |
| Loop read count | : | 245911 |
| Star read count | : | 185 |

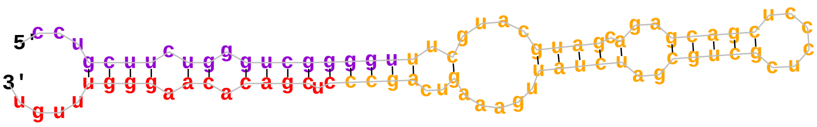


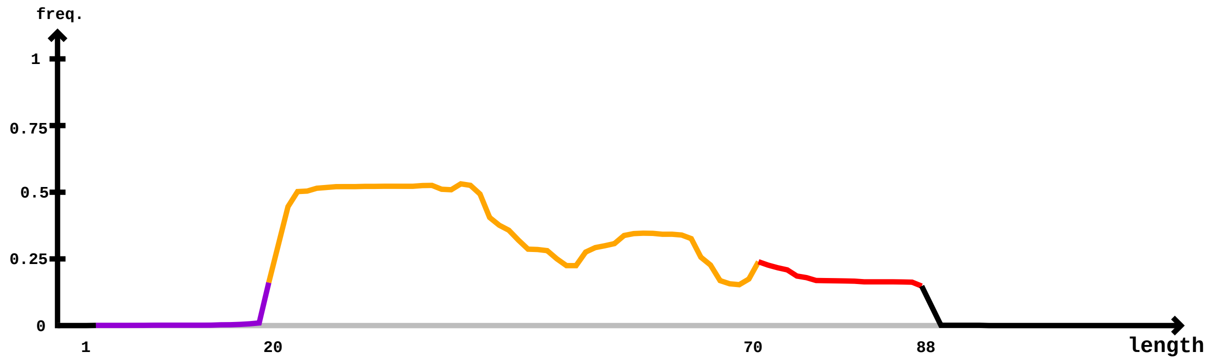


```
      Star                                                                 Mature
5'- gaccugcuucugggucgggguuucguacguagcagagcagcucccucgcugcgaucuauugaaagucagcccucgacacaaggguuuguccgcgcgcgcgcgcgcgcgugcgu   -3'    obs
    gaccugcuucugggucgggguuucguacguagcagagcagcucccucgcugcgaucuauugaaagucagcccucgacacaaggguuuguccgcgcgcgcgcgcgcgcgugcgu          exp
    (((..((((.((.(((((((((.(....((((.(..(((((......)))))..))))).....)..))))).)))).)).)))).)))((((((((((....)))))))))).   reads     mm       sample
    ..ccugcuucugggucgggg...............................................................................................   7         0        7y1
    ..ccugcuucugggucggggA..............................................................................................   1         1        7y1
    ..ccugcuucugggucggggG..............................................................................................   6         1        7y1
    ..cAugcuucugggucggggu..............................................................................................   1         1        7y1
    ..ccugcuucugggucggggu..............................................................................................   164       0        7y1
    ...cugcuucuggguAgggguuucgu.........................................................................................   2         1        7y1
    ...cugcuucugggucgggguAucgu.........................................................................................   1         1        7y1
    ...cugcuucugUgucgggguuucgu.........................................................................................   1         1        7y1
    ...cugcuucugggucgggUuuucgu.........................................................................................   2         1        7y1
    ....ugcuucAgggucgggguu.............................................................................................   1         1        7y1
    .....gcuucuggCucgggguuucgu.........................................................................................   1         1        7y1
    .....gcuucugggAcgggguuucgu.........................................................................................   1         1        7y1
    .....gcuucuggguAgggguuucgu.........................................................................................   1         1        7y1
    .....gcuucuggUucgggguuucgu.........................................................................................   1         1        7y1
    ......cuucugggucgggguuuG...........................................................................................   1         1        7y1
    ......cuucugggAcgggguuuc...........................................................................................   1         1        7y1
    ......cuucugggucgggguGuc...........................................................................................   1         1        7y1
    ......cuucuAggucgggguuuc...........................................................................................   1         1        7y1
    ......cuucugggucggCguuuc...........................................................................................   1         1        7y1
    ......cuucuUggucgggguuucgu.........................................................................................   1         1        7y1
    ......cuucugggAcgggguuucgu.........................................................................................   2         1        7y1
    ......cuucuCggucgggguuucgu.........................................................................................   1         1        7y1
    ......cuucuggguAgggguuucgu.........................................................................................   4         1        7y1
    ......cuucugggucCggguuucgu.........................................................................................   1         1        7y1
    ......cuucugggucggCguuucgu.........................................................................................   1         1        7y1
    ......cuucugggucUggguuucgu.........................................................................................   1         1        7y1
    ......cuucugggucggggGuucgu.........................................................................................   2         1        7y1
    ......cuucugggucgggAuuucgu.........................................................................................   1         1        7y1
    ......cuucugggucgggUuuucgu.........................................................................................   1         1        7y1
    ......cuucugggucgggguuucUu.........................................................................................   3         1        7y1
    ......cuucugggucgggguCucgu.........................................................................................   1         1        7y1
    ......cuucugggucgggguAucgu.........................................................................................   1         1        7y1
    ......cuucugggucggggAuucgu.........................................................................................   1         1        7y1
```

```
      Star                                                                       Mature
gaccugcuucugggucgggguuucguacguagcagagcagcucccucgcugcgaucuauugaaagucagcccucgacacaaggguuuguccgcgcgcgcgcgcgcgcgcgugcgu
......cuucuggguGgggguuucgu.........................................................................................    1    1    7y1
......cuucugggucggUguuucgu.........................................................................................    1    1    7y1
.......uucugggAcgggguuucgu.........................................................................................    3    1    7y1
.......uucugggucgUgguuucgu.........................................................................................    2    1    7y1
.......uucuggguAgggguuucgu.........................................................................................    6    1    7y1
.......uucugCgucgggguuucgu.........................................................................................    4    1    7y1
.......uucugggucUgguuucgu..........................................................................................    2    1    7y1
.......uucugggucggggCuucgu.........................................................................................    1    1    7y1
.......uucugggucggCguuucgu.........................................................................................    1    1    7y1
.......uucuUggucgggguuucgu.........................................................................................    2    1    7y1
.......uucugggucggUguuucgu.........................................................................................    1    1    7y1
.......uucugggucgggCuuucgu.........................................................................................    4    1    7y1
.......uucugggucgggguAucgu.........................................................................................    3    1    7y1
.......uucugggucggggGuucgu.........................................................................................    1    1    7y1
.......uucuggUucgggguuucgu.........................................................................................    2    1    7y1
.......uucugggucgggguCucgu.........................................................................................    2    1    7y1
.......uucugUgucgggguuucgu.........................................................................................    1    1    7y1
........ucuggCucgggguuucgu.........................................................................................    4    1    7y1
........ucugggucgggCuuucgu.........................................................................................    1    1    7y1
........ucugCgucgggguuucgu.........................................................................................    1    1    7y1
........ucugggucgUgguuucgu.........................................................................................    1    1    7y1
........ucuggguAgggguuucgu.........................................................................................    4    1    7y1
........ucugggucgCgguuucgu.........................................................................................    8    1    7y1
........ucuggAucgggguuucgu.........................................................................................    1    1    7y1
........ucugggucggggAuucgu.........................................................................................    1    1    7y1
........ucugggucggCguuucgu.........................................................................................    6    1    7y1
........ucugggucgggguAucgu.........................................................................................    4    1    7y1
........ucugggucCggguuucgu.........................................................................................    2    1    7y1
........ucugggucgggguCucgu.........................................................................................    1    1    7y1
........ucugUgucgggguuucgu.........................................................................................    2    1    7y1
........ucugggucgggUuuucgu.........................................................................................    1    1    7y1
........ucuAggucgggguuucgu.........................................................................................    2    1    7y1
........ucuggUucgggguuucgu.........................................................................................    7    1    7y1
........ucugggucggUguuucgu.........................................................................................    3    1    7y1
........ucugggucUgguuucgu..........................................................................................    2    1    7y1
........ucugggAcgggguuucgu.........................................................................................    4    1    7y1
........ucuggguGgggguuucgu.........................................................................................    1    1    7y1
........ucugggucggggguuAcgu........................................................................................    1    1    7y1
........ucugggucgggUuuucgua........................................................................................    2    1    7y1
........ucugggucggggAuucgua........................................................................................    1    1    7y1
........ucugggucgggguuAcgua........................................................................................    1    1    7y1
........ucugggucUgguuucgua.........................................................................................    3    1    7y1
........ucuggguAgggguuucgua........................................................................................    4    1    7y1
........ucugggucgggCuuucgua........................................................................................    1    1    7y1
........ucugggucgUgguuucgua........................................................................................    3    1    7y1
........ucuggguGgggguuucgua........................................................................................    1    1    7y1
........ucugggucgggguuGcgua........................................................................................    1    1    7y1
........ucugggucggUguuucgua........................................................................................    4    1    7y1
........ucugggucggCguuucgua........................................................................................    2    1    7y1
........ucugUgucgggguuucgua........................................................................................    2    1    7y1
........ucugggucgCgguuucgua........................................................................................    6    1    7y1
........ucuggAucgggguuucgua........................................................................................    1    1    7y1
........ucugggAcgggguuucgua........................................................................................    12   1    7y1
........ucugggucggAguuucgua........................................................................................    1    1    7y1
........ucugggucCggguuucgua........................................................................................    2    1    7y1
........ucugCgucgggguuucgua........................................................................................    2    1    7y1
........ucuggUucgggguuucgua........................................................................................    5    1    7y1
........ucugggucgggguAucgua........................................................................................    2    1    7y1
.........cugggucgggguuuAgua........................................................................................    5    1    7y1
.........cugggucgggUuuucgua........................................................................................    1    1    7y1
.........cugggucgCgguuucgua........................................................................................    1    1    7y1
.........cugggucgAgguuucgua........................................................................................    1    1    7y1
.........cugggucgggguuuGgua........................................................................................    1    1    7y1
.........cugCgucgggguuucgua........................................................................................    1    1    7y1
.........cuggguAgggguuucgua........................................................................................    5    1    7y1
.........cugggucUgguuucgua.........................................................................................    1    1    7y1
.........cugggucggUguuucgua........................................................................................    1    1    7y1
.........cugggucggggGuucgua........................................................................................    1    1    7y1
.........cugggucgggguuAcguac.......................................................................................    1    1    7y1
.........cugggucgggguuuGguac.......................................................................................    1    1    7y1
```

**Star** **Mature**

```
gaccugcuucugggucgggguuucguacguagcagagcagcucccucgcugcgaucuauugaaagucagcccucgacacaaggguuuguccgcgcgcgcgcgcgcgcgugcgu
.........cugggucgggguuuAguac.....................................................................................     1     1     7y1
.........cuggguAgggguuucguac.....................................................................................     1     1     7y1
.........cugggucCggguuucguac.....................................................................................     1     1     7y1
.........cugggucggCguuucguac.....................................................................................     1     1     7y1
.........cugggAcgggguuucguac.....................................................................................     1     1     7y1
.........cugggucgggUuuucguac.....................................................................................     1     1     7y1
..............ucgggguAucguacguag.................................................................................     1     1     7y1
..............ucggggAuucguacguag.................................................................................     1     1     7y1
..............ucggggAuucguacguagc................................................................................     1     1     7y1
..............ucgggUuuucguacguagc................................................................................     1     1     7y1
..............ucggggCuucguacguagc................................................................................     1     1     7y1
..............ucgggguAucguacguagc................................................................................     1     1     7y1
..............ucggggGuucguacguagc................................................................................     1     1     7y1
..............ucgggguuucguaUguagc................................................................................     1     1     7y1
..............ucgggguuucAuacguagcagagc...........................................................................     2     1     7y1
..............ucgggguGucguacguagcagagcagcuc......................................................................     1     1     7y1
..............ucgggguCucguacguagcagagcagcuc......................................................................     1     1     7y1
..............ucgggCuuucguacguagcagagcagcuc......................................................................     6     1     7y1
..............ucgggUuuucguacguagcagagcagcuc......................................................................     4     1     7y1
..............ucggggCuucguacguagcagagcagcuc......................................................................     2     1     7y1
..............ucggggAuucguacguagcagagcagcuc......................................................................     23    1     7y1
...............cgggguuucgCacguagc................................................................................     1     1     7y1
...............cggAguuucguacguagc................................................................................     1     1     7y1
...............cggggAuucguacguagc................................................................................     2     1     7y1
...............cgggguuucguaAguagc................................................................................     1     1     7y1
...............cgggguuAcguacguagc................................................................................     1     1     7y1
...............cgggguuucUuacguagc................................................................................     1     1     7y1
...............cggCguuucguacguagc................................................................................     1     1     7y1
...............cgggguAucguacguagc................................................................................     3     1     7y1
...............cgggguuucguacAuagca...............................................................................     1     1     7y1
...............cggggAuucguacguagca...............................................................................     2     1     7y1
...............cgggguuucUuacguagca...............................................................................     1     1     7y1
...............cgggguuucguaAguagca...............................................................................     1     1     7y1
...............cgggguuucCuacguagcag..............................................................................     1     1     7y1
...............cgggguuucguacCuagcag..............................................................................     1     1     7y1
...............Agggguuucguacguagcag..............................................................................     1     1     7y1
...............cgggguuucguacgAagcag..............................................................................     1     1     7y1
...............cggggAuucguacguagcag..............................................................................     1     1     7y1
...............cgggguuuAguacguagcag..............................................................................     1     1     7y1
...............cgggUuuucguacguagcag..............................................................................     1     1     7y1
...............cgggguGucguacguagcaga.............................................................................     1     1     7y1
...............cgggguuAcguacguagcaga.............................................................................     2     1     7y1
...............cgggguuuAguacguagcaga.............................................................................     1     1     7y1
...............cgggguuucUuacguagcaga.............................................................................     1     1     7y1
...............cgggguuucCuacguagcaga.............................................................................     1     1     7y1
...............cgggguuucguaGguagcaga.............................................................................     1     1     7y1
...............cgggguuucguacUuagcaga.............................................................................     4     1     7y1
...............cgggCuuucguacguagcaga.............................................................................     3     1     7y1
...............cgggguuucguaAguagcaga.............................................................................     3     1     7y1
...............cgggguuuGguacguagcaga.............................................................................     1     1     7y1
...............cgggguuucguacgCagcaga.............................................................................     1     1     7y1
...............cgggguAucguacguagcaga.............................................................................     4     1     7y1
...............cggggAuucguacguagcaga.............................................................................     5     1     7y1
...............cgggguuuUguacguagcaga.............................................................................     1     1     7y1
...............cggggGuucguacguagcaga.............................................................................     1     1     7y1
...............cgggguuucAuacguagcagag............................................................................     1     1     7y1
...............cgggguuuAguacguagcagag............................................................................     1     1     7y1
...............cgggguuucguacUuagcagag............................................................................     1     1     7y1
...............cgggUuuucguacguagcagag............................................................................     1     1     7y1
...............cgggguuucUuacguagcagag............................................................................     1     1     7y1
...............cgggguAucguacguagcagag............................................................................     2     1     7y1
...............cgggguuucguaAguagcagag............................................................................     3     1     7y1
...............cgggguuucgAacguagcagag............................................................................     1     1     7y1
...............cgggguuucguacUuagcagagc...........................................................................     2     1     7y1
...............cgggCuuucguacguagcagagc...........................................................................     3     1     7y1
...............cgggguuuAguacguagcagagc...........................................................................     7     1     7y1
...............cgggguuucgGacguagcagagc...........................................................................     1     1     7y1
...............cgggguuucCuacguagcagagc...........................................................................     2     1     7y1
...............cggggGuucguacguagcagagc...........................................................................     3     1     7y1
...............cgggguuucgAacguagcagagc...........................................................................     3     1     7y1
```

### Star

### Mature

| | | | |
|---|---|---|---|
| gaccugcuucugggucggggguuucguacguagcagagcagcucccucgcugcgaucuauugaaagucagcccucgacacaaggguuuguccgcgcgcgcgcgcgcgcgugcgu | | | |
| ...............cgggguuucguacCuagcagagc........................................................................... | 1 | 1 | 7y1 |
| ...............cgggguuucguaAguagcagagc........................................................................... | 11 | 1 | 7y1 |
| ...............cggggAuucguacguagcagagc........................................................................... | 27 | 1 | 7y1 |
| ...............cgggguuucUuacguagcagagc........................................................................... | 1 | 1 | 7y1 |
| ...............cgggguuCcguacguagcagagc........................................................................... | 2 | 1 | 7y1 |
| ...............cgggguuucAuacguagcagagc........................................................................... | 1 | 1 | 7y1 |
| ...............cgggguuucguacgAagcagagc........................................................................... | 3 | 1 | 7y1 |
| ...............cgggguuucgCacguagcagagc........................................................................... | 3 | 1 | 7y1 |
| ...............cgggguAucguacguagcagagc........................................................................... | 14 | 1 | 7y1 |
| ...............cgggUuuucguacguagcagagc........................................................................... | 2 | 1 | 7y1 |
| ...............cgggguuucguGcguagcagagc........................................................................... | 1 | 1 | 7y1 |
| ...............cgggguuucguacAuagcagagc........................................................................... | 1 | 1 | 7y1 |
| ...............cgggguuuGguacguagcagagc........................................................................... | 1 | 1 | 7y1 |
| ...............cgggguCucguacguagcagagc........................................................................... | 1 | 1 | 7y1 |
| ...............cgggguuAcguacguagcagagc........................................................................... | 3 | 1 | 7y1 |
| ...............cgggguuucguaUguagcagagc........................................................................... | 1 | 1 | 7y1 |
| ...............cgggguuucguacgAagcagagca.......................................................................... | 3 | 1 | 7y1 |
| ...............cgggguuucgAacguagcagagca.......................................................................... | 3 | 1 | 7y1 |
| ...............cgggguuucCuacguagcagagca.......................................................................... | 2 | 1 | 7y1 |
| ...............cgggguuucguaAguagcagagca.......................................................................... | 4 | 1 | 7y1 |
| ...............cgggUuuucguacguagcagagca.......................................................................... | 1 | 1 | 7y1 |
| ...............cgggguuucgCacguagcagagca.......................................................................... | 1 | 1 | 7y1 |
| ...............cgggguuucguacUuagcagagca.......................................................................... | 2 | 1 | 7y1 |
| ...............cgggguuucguacgGagcagagca.......................................................................... | 1 | 1 | 7y1 |
| ...............cgggguuucguGcguagcagagca.......................................................................... | 2 | 1 | 7y1 |
| ...............cgggguuAcguacguagcagagca.......................................................................... | 6 | 1 | 7y1 |
| ...............cgggCuuucguacguagcagagca.......................................................................... | 3 | 1 | 7y1 |
| ...............cgggguuuAguacguagcagagca.......................................................................... | 5 | 1 | 7y1 |
| ...............cggggAuucguacguagcagagca.......................................................................... | 10 | 1 | 7y1 |
| ...............cgggguuucAuacguagcagagca.......................................................................... | 1 | 1 | 7y1 |
| ...............cggggGuucguacguagcagagca.......................................................................... | 1 | 1 | 7y1 |
| ...............cgggguuucgGacguagcagagca.......................................................................... | 1 | 1 | 7y1 |
| ...............cgggguuuUguacguagcagagca.......................................................................... | 1 | 1 | 7y1 |
| ...............cgggguuucUuacguagcagagca.......................................................................... | 4 | 1 | 7y1 |
| ...............cgggguAucguacguagcagagca.......................................................................... | 10 | 1 | 7y1 |
| ...............cgggAuuucguacguagcagagca.......................................................................... | 1 | 1 | 7y1 |
| ...............cgggguCucguacguagcagagca.......................................................................... | 1 | 1 | 7y1 |
| ...............cgggCuuucguacguagcagagcag......................................................................... | 1 | 1 | 7y1 |
| ...............cgggguuucguacUuagcagagcag......................................................................... | 1 | 1 | 7y1 |
| ...............cgggguuucgAacguagcagagcag......................................................................... | 1 | 1 | 7y1 |
| ...............cgggguuucCuacguagcagagcag......................................................................... | 1 | 1 | 7y1 |
| ...............cgggguuuAguacguagcagagcag......................................................................... | 4 | 1 | 7y1 |
| ...............cgggguAucguacguagcagagcag......................................................................... | 1 | 1 | 7y1 |
| ...............cgggguuAcguacguagcagagcag......................................................................... | 1 | 1 | 7y1 |
| ...............cgggUuuucguacguagcagagcag......................................................................... | 2 | 1 | 7y1 |
| ...............cgggguuuGguacguagcagagcag......................................................................... | 1 | 1 | 7y1 |
| ...............cgggguuucguaAguagcagagcag......................................................................... | 1 | 1 | 7y1 |
| ...............cggggCuucguacguagcagagcagc........................................................................ | 1 | 1 | 7y1 |
| ...............cgggguuuAguacguagcagagcagc........................................................................ | 2 | 1 | 7y1 |
| ...............cgggguuucguaAguagcagagcagc........................................................................ | 5 | 1 | 7y1 |
| ...............cgggguuucguacCuagcagagcagc........................................................................ | 1 | 1 | 7y1 |
| ...............cgggguAucguacguagcagagcagc........................................................................ | 9 | 1 | 7y1 |
| ...............cgggguGucguacguagcagagcagc........................................................................ | 1 | 1 | 7y1 |
| ...............cgggguuAcguacguagcagagcagc........................................................................ | 2 | 1 | 7y1 |
| ...............cgggguuucguaGguagcagagcagc........................................................................ | 1 | 1 | 7y1 |
| ...............cgggguuucgAacguagcagagcagc........................................................................ | 4 | 1 | 7y1 |
| ...............cgggguuCcguacguagcagagcagc........................................................................ | 1 | 1 | 7y1 |
| ...............cgggguuucAuacguagcagagcagc........................................................................ | 2 | 1 | 7y1 |
| ...............cgggguuucguacgAagcagagcagc........................................................................ | 2 | 1 | 7y1 |
| ...............cggggGuucguacguagcagagcagc........................................................................ | 1 | 1 | 7y1 |
| ...............cggggAuucguacguagcagagcagc........................................................................ | 10 | 1 | 7y1 |
| ...............cgggguuucgGacguagcagagcagc........................................................................ | 1 | 1 | 7y1 |
| ...............cgggguuucguacUuagcagagcagc........................................................................ | 2 | 1 | 7y1 |
| ...............cgggguuuUguacguagcagagcagc........................................................................ | 1 | 1 | 7y1 |
| ...............cgggguuucguacUuagcagagcagcu....................................................................... | 2 | 1 | 7y1 |
| ...............cgggguuuAguacguagcagagcagcu....................................................................... | 1 | 1 | 7y1 |
| ...............cggggAuucguacguagcagagcagcu....................................................................... | 10 | 1 | 7y1 |
| ...............cgggguuucguGcguagcagagcagcu....................................................................... | 1 | 1 | 7y1 |
| ...............cgggUuuucguacguagcagagcagcu....................................................................... | 2 | 1 | 7y1 |
| ...............cgggguAucguacguagcagagcagcu....................................................................... | 12 | 1 | 7y1 |

```
Star                                                                   Mature
gaccugcuucugggucgggguuucguacguagcagagcagcucccucgcugcgaucuauugaaagucagcccucgacacaagguuuguccgcgcgcgcgcgcgcgcgugcgu
...............cgggguuucguacgAagcagagcagcu......................................................................  1   1  7y1
...............cgggguuucgAacguagcagagcagcu......................................................................  4   1  7y1
...............cgggguuucCuacguagcagagcagcu......................................................................  1   1  7y1
...............cgggguuucAuacguagcagagcagcu......................................................................  2   1  7y1
...............cgggguuucgGacguagcagagcagcu......................................................................  1   1  7y1
...............cgggguuuUguacguagcagagcagcu......................................................................  1   1  7y1
...............cgggguCucguacguagcagagcagcu......................................................................  1   1  7y1
...............cgggguuucguaAguagcagagcagcu......................................................................  2   1  7y1
...............cgggguuAcguacguagcagagcagcu......................................................................  4   1  7y1
...............cggggGuucguacguagcagagcagcu......................................................................  3   1  7y1
...............cggggCuucguacguagcagagcagcu......................................................................  1   1  7y1
...............cgggguuucUuacguagcagagcagcu......................................................................  1   1  7y1
...............cgggguGucguacguagcagagcagcuc.....................................................................  2   1  7y1
...............cggggGuucguacguagcagagcagcuc.....................................................................  4   1  7y1
...............cgggguuucUuacguagcagagcagcuc.....................................................................  1   1  7y1
...............cgggguuuGguacguagcagagcagcuc.....................................................................  1   1  7y1
...............cgggguuucguacUuagcagagcagcuc.....................................................................  3   1  7y1
...............cgggguuucCuacguagcagagcagcuc.....................................................................  1   1  7y1
...............cggggAuucguacguagcagagcagcuc.....................................................................  12  1  7y1
...............cgggguuucguacCuagcagagcagcuc.....................................................................  1   1  7y1
...............cgggguAucguacguagcagagcagcuc.....................................................................  16  1  7y1
...............cgggguuucguacgAagcagagcagcuc.....................................................................  1   1  7y1
...............cgggguuucAuacguagcagagcagcuc.....................................................................  2   1  7y1
...............cgggUuuucguacguagcagagcagcuc.....................................................................  2   1  7y1
...............cgggguuAcguacguagcagagcagcuc.....................................................................  7   1  7y1
...............cgggguuucgAacguagcagagcagcuc.....................................................................  2   1  7y1
...............cgggCuuucguacguagcagagcagcuc.....................................................................  2   1  7y1
...............cgggguuucgCacguagcagagcagcuc.....................................................................  4   1  7y1
...............cgggguuucguaAguagcagagcagcuc.....................................................................  1   1  7y1
...............cgggguuucguaUguagcagagcagcuc.....................................................................  1   1  7y1
...............cgggguuuAguacguagcagagcagcuc.....................................................................  5   1  7y1
...............cggggAuucguacguagcagagcagcucccuc.................................................................  1   1  7y1
................gggguuAcguacguagcaga............................................................................  2   1  7y1
................gggguuucguaAguagcaga............................................................................  1   1  7y1
................gggguuucgAacguagcagag...........................................................................  1   1  7y1
................gggguuuAguacguagcagag...........................................................................  1   1  7y1
................gggguuuAguacguagcagagc..........................................................................  2   1  7y1
................gggguuucUuacguagcagagc..........................................................................  1   1  7y1
................gggguuAcguacguagcagagc..........................................................................  3   1  7y1
................gggUuuucguacguagcagagc..........................................................................  3   1  7y1
................gggguAucguacguagcagagc..........................................................................  5   1  7y1
................gggguuucgAacguagcagagca.........................................................................  1   1  7y1
................gggguuucguaAguagcagagca.........................................................................  1   1  7y1
................gggguuuAguacguagcagagca.........................................................................  1   1  7y1
................gggguAucguacguagcagagca.........................................................................  2   1  7y1
................gggUuuucguacguagcagagca.........................................................................  1   1  7y1
................gggguuAcguacguagcagagca.........................................................................  2   1  7y1
................gggguuAcguacguagcagagcagc.......................................................................  3   1  7y1
................gggguuucgAacguagcagagcagc.......................................................................  1   1  7y1
................gggguuucguUcguagcagagcagc.......................................................................  2   1  7y1
................gggguuucguacgCagcagagcagc.......................................................................  1   1  7y1
................ggggGuucguacguagcagagcagc.......................................................................  1   1  7y1
................gggguAucguacguagcagagcagc.......................................................................  3   1  7y1
................gggguGucguacguagcagagcagc.......................................................................  1   1  7y1
................gggguuCcguacguagcagagcagc.......................................................................  1   1  7y1
................gggguuuAguacguagcagagcagc.......................................................................  3   1  7y1
................gggguuucguacgAagcagagcagc.......................................................................  1   1  7y1
................gggguuucguacguGgcagagcagc.......................................................................  1   1  7y1
................gggguuucCuacguagcagagcagcu......................................................................  1   1  7y1
................gggguuucguacgCagcagagcagcu......................................................................  2   1  7y1
................gggguuucguacUuagcagagcagcu......................................................................  1   1  7y1
................ggggAuucguacguagcagagcagcu......................................................................  2   1  7y1
................gggguuucguGcguagcagagcagcu......................................................................  1   1  7y1
................gggCuuucguacguagcagagcagcu......................................................................  1   1  7y1
................ggggGuucguacguagcagagcagcu......................................................................  1   1  7y1
................gggguuucguaAguagcagagcagcu......................................................................  1   1  7y1
................gggguAucguacguagcagagcagcu......................................................................  6   1  7y1
................gggguuAcguacguagcagagcagcu......................................................................  1   1  7y1
................gggguuuAguacguagcagagcagcu......................................................................  1   1  7y1
................gggguuucguacAuagcagagcagcuc.....................................................................  2   1  7y1
```

```
   Star                                                                   Mature
gaccugcuucugggucggggguuucguacguagcagagcagcucccucgcugcgaucuauugaaagucagcccucgacacaaggguuuguccgcgcgcgcgcgcgcgcgugcgu
................gggguuucAuacguagcagagcagcuc......................................................................   1     1     7y1
................gggguAucguacguagcagagcagcuc......................................................................   9     1     7y1
................gggguuucguacUuagcagagcagcuc......................................................................   3     1     7y1
................gggguCucguacguagcagagcagcuc......................................................................   2     1     7y1
................gggguuGcguacguagcagagcagcuc......................................................................   1     1     7y1
................gggguuucguacguGgcagagcagcuc......................................................................   1     1     7y1
................gggguuucguaAguagcagagcagcuc......................................................................   3     1     7y1
................gggUuuucguacguagcagagcagcuc......................................................................   1     1     7y1
................gggguuuAguacguagcagagcagcuc......................................................................   8     1     7y1
................gggguuucCuacguagcagagcagcuc......................................................................   1     1     7y1
................gggguuucguacguUgcagagcagcuc......................................................................   1     1     7y1
................gggguuucguGcguagcagagcagcuc......................................................................   1     1     7y1
................ggggAuucguacguagcagagcagcuc......................................................................   1     1     7y1
................gggguuAcguacguagcagagcagcuc......................................................................   4     1     7y1
................gggguuucgAacguagcagagcagcuc......................................................................   2     1     7y1
.................ggguuucguaAguagcaga.............................................................................   1     1     7y1
.................ggguuAcguacguagcaga.............................................................................   1     1     7y1
.................ggguuucguacgCagcaga.............................................................................   1     1     7y1
.................ggguuuAguacguagcaga.............................................................................   1     1     7y1
.................ggguuucAuacguagcagagc...........................................................................   1     1     7y1
.................ggguAucguacguagcagagc...........................................................................   2     1     7y1
.................gggGuucguacguagcagagc...........................................................................   5     1     7y1
.................ggguuAcguacguagcagagc...........................................................................   1     1     7y1
.................ggguuucguaAguagcagagc...........................................................................   1     1     7y1
.................ggguuuAguacguagcagagc...........................................................................   5     1     7y1
.................gggAuucguacguagcagagc...........................................................................   1     1     7y1
.................ggguuucgAacguagcagagc...........................................................................   2     1     7y1
.................ggguuucguaUguagcagagc...........................................................................   1     1     7y1
.................ggguuucgGacguagcagagca..........................................................................   1     1     7y1
.................ggguuucUuacguagcagagca..........................................................................   1     1     7y1
.................gggGuucguacguagcagagca..........................................................................   1     1     7y1
.................ggCuuucguacguagcagagca..........................................................................   269   1     7y1
.................ggguuucgAacguagcagagca..........................................................................   1     1     7y1
.................ggguAucguacguagcagagca..........................................................................   1     1     7y1
.................ggguuucguaAguagcagagca..........................................................................   1     1     7y1
.................ggguCucguacguagcagagca..........................................................................   1     1     7y1
.................ggguuuGguacguagcagagca..........................................................................   1     1     7y1
.................ggguuucguacguGgcagagca..........................................................................   1     1     7y1
.................ggguuCcguacguagcagagca..........................................................................   1     1     7y1
.................ggguuuAguacguagcagagca..........................................................................   3     1     7y1
.................ggguuAcguacguagcagagca..........................................................................   1     1     7y1
.................ggguGucguacguagcagagca..........................................................................   1     1     7y1
.................ggguuucguacguaCcagagca..........................................................................   1     1     7y1
.................ggguuucguacguaUcagagca..........................................................................   1     1     7y1
.................ggguuucguaAguagcagagcagc........................................................................   1     1     7y1
.................ggguuucgAacguagcagagcagc........................................................................   1     1     7y1
.................ggguuuAguacguagcagagcagc........................................................................   2     1     7y1
.................gggGuucguacguagcagagcagc........................................................................   1     1     7y1
.................ggguuucguacCuagcagagcagcu.......................................................................   1     1     7y1
.................ggguAucguacguagcagagcagcu.......................................................................   2     1     7y1
.................ggguuuAguacguagcagagcagcu.......................................................................   6     1     7y1
.................ggCuuucguacguagcagagcagcu.......................................................................   1     1     7y1
.................gggGuucguacguagcagagcagcu.......................................................................   2     1     7y1
.................ggguuucguacguaUcagagcagcu.......................................................................   1     1     7y1
.................ggguuucCuacguagcagagcagcu.......................................................................   1     1     7y1
.................ggguuucgAacguagcagagcagcu.......................................................................   1     1     7y1
.................ggguuAcguacguagcagagcagcu.......................................................................   5     1     7y1
.................ggguuucguaUguagcagagcagcu.......................................................................   1     1     7y1
.................ggguuucguaAguagcagagcagcu.......................................................................   1     1     7y1
.................gggGuucguacguagcagagcagcuc......................................................................   2     1     7y1
.................ggguuucguacUuagcagagcagcuc......................................................................   1     1     7y1
.................ggguuAcguacguagcagagcagcuc......................................................................   6     1     7y1
.................ggguAucguacguagcagagcagcuc......................................................................   5     1     7y1
.................ggguuucguacgAagcagagcagcuc......................................................................   2     1     7y1
.................ggguuCcguacguagcagagcagcuc......................................................................   1     1     7y1
.................ggguuuAguacguagcagagcagcuc......................................................................   9     1     7y1
.................ggguuucguaUguagcagagcagcuc......................................................................   1     1     7y1
.................gggCuucguacguagcagagcagcuc......................................................................   1     1     7y1
.................ggguuucguacCuagcagagcagcuc......................................................................   1     1     7y1
.................ggguuucguaAguagcagagcagcuc......................................................................   2     1     7y1
```

| | | | |
|---|---|---|---|
| Star / Mature | | | |
| gaccugcuucugggucggggu uucguacguagcagagcagcucccucgcugcgaucuauugaaagucagcc cucgacacaaggguuugu ccgcgcgcgcgcgcgcgugcgu | | | |
| .................ggguuucUuacguagcagagcagcuc.................................................................... | 1 | 1 | 7y1 |
| .................ggguuucAuacguagcagagcagcuc.................................................................... | 2 | 1 | 7y1 |
| .................ggguuucguacguCgcagagcagcuc.................................................................... | 1 | 1 | 7y1 |
| .................ggguuucgAacguagcagagcagcuc.................................................................... | 1 | 1 | 7y1 |
| .................ggguuucguacgAagcagagcagcucc................................................................... | 1 | 1 | 7y1 |
| .................gggGuucguacguagcagagcagcucccu................................................................. | 1 | 1 | 7y1 |
| .................ggguuucgAacguagcagagcagcucccu................................................................. | 1 | 1 | 7y1 |
| .................ggguuucguacguGgcagagcagcucccu................................................................. | 1 | 1 | 7y1 |
| .................ggguuAcguacguagcagagcagcucccu................................................................. | 1 | 1 | 7y1 |
| .................ggguuuAguacguagcagagcagcucccu................................................................. | 1 | 1 | 7y1 |
| ..................gguuAcguacguagcagagc......................................................................... | 1 | 1 | 7y1 |
| ..................gguuucCuacguagcagagc......................................................................... | 4 | 1 | 7y1 |
| ..................gguuucguacguaCcagagc......................................................................... | 3 | 1 | 7y1 |
| ..................gguuucAuacguagcagagc......................................................................... | 3 | 1 | 7y1 |
| ..................gguuucguacUuagcagagc......................................................................... | 1 | 1 | 7y1 |
| ..................gUuuucguacguagcagagc......................................................................... | 3 | 1 | 7y1 |
| ..................gguuucguacguagAagagc......................................................................... | 5 | 1 | 7y1 |
| ..................gguuucguUcguagcagagc......................................................................... | 1 | 1 | 7y1 |
| ..................gguuucguaGguagcagagc......................................................................... | 1 | 1 | 7y1 |
| ..................gguuucgAacguagcagagc......................................................................... | 9 | 1 | 7y1 |
| ..................gguuucguacguGgcagagc......................................................................... | 1 | 1 | 7y1 |
| ..................gguuucguacCuagcagagc......................................................................... | 2 | 1 | 7y1 |
| ..................ggGuucguacguagcagagc......................................................................... | 1 | 1 | 7y1 |
| ..................ggAuucguacguagcagagc......................................................................... | 4 | 1 | 7y1 |
| ..................gguGucguacguagcagagc......................................................................... | 1 | 1 | 7y1 |
| ..................gguuucguacguaUcagagc......................................................................... | 2 | 1 | 7y1 |
| ..................gguuucguaAguagcagagc......................................................................... | 16 | 1 | 7y1 |
| ..................gguCucguacguagcagagc......................................................................... | 1 | 1 | 7y1 |
| ..................gguuuAguacguagcagagc......................................................................... | 11 | 1 | 7y1 |
| ..................gguuucguacguagGagagc......................................................................... | 1 | 1 | 7y1 |
| ..................gguuucguacguUgcagagc......................................................................... | 1 | 1 | 7y1 |
| ..................gguAucguacguagcagagc......................................................................... | 3 | 1 | 7y1 |
| ..................gguuucguacgGagcagagc......................................................................... | 1 | 1 | 7y1 |
| ..................gguuuGguacguagcagagc......................................................................... | 2 | 1 | 7y1 |
| ..................gCuuucguacguagcagagc......................................................................... | 2 | 1 | 7y1 |
| ..................gguuucguacgAagcagagc......................................................................... | 5 | 1 | 7y1 |
| ..................gguuAcguacguagcagagca........................................................................ | 1 | 1 | 7y1 |
| ..................gguuucguacCuagcagagca........................................................................ | 1 | 1 | 7y1 |
| ..................gCuuucguacguagcagagca........................................................................ | 1 | 1 | 7y1 |
| ..................gguuucguaUguagcagagca........................................................................ | 1 | 1 | 7y1 |
| ..................gguuuAguacguagcagagca........................................................................ | 3 | 1 | 7y1 |
| ..................gguuucgGacguagcagagca........................................................................ | 1 | 1 | 7y1 |
| ..................gguuucguacguaAcagagca........................................................................ | 1 | 1 | 7y1 |
| ..................gguuucgAacguagcagagca........................................................................ | 1 | 1 | 7y1 |
| ..................gguCucguacguagcagagca........................................................................ | 1 | 1 | 7y1 |
| ..................gguuucguacguaCcagagca........................................................................ | 2 | 1 | 7y1 |
| ..................gguuucCuacguagcagagca........................................................................ | 2 | 1 | 7y1 |
| ..................gguuucguacguagUagagca........................................................................ | 1 | 1 | 7y1 |
| ..................gguuucguaAguagcagagca........................................................................ | 3 | 1 | 7y1 |
| ..................ggAuucguacguagcagagca........................................................................ | 1 | 1 | 7y1 |
| ..................gUuuucguacguagcagagcag....................................................................... | 1 | 1 | 7y1 |
| ..................gguAucguacguagcagagcag....................................................................... | 1 | 1 | 7y1 |
| ..................gguuucCuacguagcagagcag....................................................................... | 1 | 1 | 7y1 |
| ..................gguuucgAacguagcagagcag....................................................................... | 2 | 1 | 7y1 |
| ..................gguuuAguacguagcagagcag....................................................................... | 1 | 1 | 7y1 |
| ..................ggAuucguacguagcagagcag....................................................................... | 1 | 1 | 7y1 |
| ..................gguuucguacguagGagagcagc...................................................................... | 3 | 1 | 7y1 |
| ..................gCuuucguacguagcagagcagc...................................................................... | 3 | 1 | 7y1 |
| ..................gguuucguacCuagcagagcagc...................................................................... | 1 | 1 | 7y1 |
| ..................gguuucguaAguagcagagcagc...................................................................... | 5 | 1 | 7y1 |
| ..................gguuucguacgAagcagagcagc...................................................................... | 6 | 1 | 7y1 |
| ..................gguuucguacguagUagagcagc...................................................................... | 2 | 1 | 7y1 |
| ..................ggAuucguacguagcagagcagc...................................................................... | 7 | 1 | 7y1 |
| ..................gguuucgAacguagcagagcagc...................................................................... | 8 | 1 | 7y1 |
| ..................gguuucUuacguagcagagcagc...................................................................... | 4 | 1 | 7y1 |
| ..................gguuucguaGguagcagagcagc...................................................................... | 2 | 1 | 7y1 |
| ..................gguuucguacguaCcagagcagc...................................................................... | 2 | 1 | 7y1 |
| ..................gAuuucguacguagcagagcagc...................................................................... | 1 | 1 | 7y1 |
| ..................gguuucguacUuagcagagcagc...................................................................... | 3 | 1 | 7y1 |
| ..................gguuucguacgCagcagagcagc...................................................................... | 1 | 1 | 7y1 |

| Star Mature | | | |
|---|---|---|---|
| gaccugcuucugggucgggguuucguacguagcagagcagcucccucgcugcgaucuauugaaagucagcccucgacacaaggguuuguccgcgcgcgcgcgcgcgcgcgugcgu | | | |
| ..................gguuucguacguaUcagagcagc.......................................................................... | 4 | 1 | 7y1 |
| ..................gguuucguacguagAagagcagc.......................................................................... | 4 | 1 | 7y1 |
| ..................ggGuucguacguagcagagcagc.......................................................................... | 1 | 1 | 7y1 |
| ..................gguuucCuacguagcagagcagc.......................................................................... | 3 | 1 | 7y1 |
| ..................gguuucguaUguagcagagcagc.......................................................................... | 1 | 1 | 7y1 |
| ..................gUuuucguacguagcagagcagc.......................................................................... | 6 | 1 | 7y1 |
| ..................gguAucguacguagcagagcagc.......................................................................... | 2 | 1 | 7y1 |
| ..................gguuuAguacguagcagagcagc.......................................................................... | 7 | 1 | 7y1 |
| ..................gguCucguacguagcagagcagc.......................................................................... | 2 | 1 | 7y1 |
| ..................gguuAcguacguagcagagcagcu......................................................................... | 5 | 1 | 7y1 |
| ..................gUuuucguacguagcagagcagcu......................................................................... | 4 | 1 | 7y1 |
| ..................gguuucguacgCagcagagcagcu......................................................................... | 1 | 1 | 7y1 |
| ..................gguuuUguacguagcagagcagcu......................................................................... | 1 | 1 | 7y1 |
| ..................gguuucguUcguagcagagcagcu......................................................................... | 2 | 1 | 7y1 |
| ..................gguAucguacguagcagagcagcu......................................................................... | 5 | 1 | 7y1 |
| ..................gCuuucguacguagcagagcagcu......................................................................... | 2 | 1 | 7y1 |
| ..................ggAuucguacguagcagagcagcu......................................................................... | 6 | 1 | 7y1 |
| ..................gguuucguaGguagcagagcagcu......................................................................... | 1 | 1 | 7y1 |
| ..................gguuucguacAuagcagagcagcu......................................................................... | 4 | 1 | 7y1 |
| ..................gguuucguacUuagcagagcagcu......................................................................... | 1 | 1 | 7y1 |
| ..................gguuucgAacguagcagagcagcu......................................................................... | 10 | 1 | 7y1 |
| ..................gguuGcguacguagcagagcagcu......................................................................... | 1 | 1 | 7y1 |
| ..................gguuucUuacguagcagagcagcu......................................................................... | 4 | 1 | 7y1 |
| ..................gguuucguacguagGagagcagcu......................................................................... | 2 | 1 | 7y1 |
| ..................gguuucguacguaUcagagcagcu......................................................................... | 4 | 1 | 7y1 |
| ..................gguuucguacguGgcagagcagcu......................................................................... | 3 | 1 | 7y1 |
| ..................gguuucguGcguagcagagcagcu......................................................................... | 1 | 1 | 7y1 |
| ..................gguuucAuacguagcagagcagcu......................................................................... | 2 | 1 | 7y1 |
| ..................gguuucCuacguagcagagcagcu......................................................................... | 2 | 1 | 7y1 |
| ..................gguuuGguacguagcagagcagcu......................................................................... | 2 | 1 | 7y1 |
| ..................gguuucguaAguagcagagcagcu......................................................................... | 20 | 1 | 7y1 |
| ..................gguuucguacgAagcagagcagcu......................................................................... | 6 | 1 | 7y1 |
| ..................ggCuucguacguagcagagcagcu......................................................................... | 1 | 1 | 7y1 |
| ..................gguuCcguacguagcagagcagcu......................................................................... | 3 | 1 | 7y1 |
| ..................gguuucguacguagAagagcagcu......................................................................... | 10 | 1 | 7y1 |
| ..................gguuuAguacguagcagagcagcu......................................................................... | 20 | 1 | 7y1 |
| ..................gguuucguacCuagcagagcagcu......................................................................... | 7 | 1 | 7y1 |
| ..................gguuucguacguaAcagagcagcu......................................................................... | 1 | 1 | 7y1 |
| ..................gguuucguaUguagcagagcagcuc........................................................................ | 1 | 1 | 7y1 |
| ..................gguuucAuacguagcagagcagcuc........................................................................ | 1 | 1 | 7y1 |
| ..................gguuuAguacguagcagagcagcuc........................................................................ | 3 | 1 | 7y1 |
| ..................ggGuucguacguagcagagcagcuc........................................................................ | 2 | 1 | 7y1 |
| ..................gguuucguacgAagcagagcagcuc........................................................................ | 4 | 1 | 7y1 |
| ..................gguuucUuacguagcagagcagcuc........................................................................ | 1 | 1 | 7y1 |
| ..................gguuucguacCuagcagagcagcuc........................................................................ | 2 | 1 | 7y1 |
| ..................gCuuucguacguagcagagcagcuc........................................................................ | 3 | 1 | 7y1 |
| ..................gguuucCuacguagcagagcagcuc........................................................................ | 1 | 1 | 7y1 |
| ..................gguuucguaAguagcagagcagcuc........................................................................ | 6 | 1 | 7y1 |
| ..................gguuucguacguagUagagcagcuc........................................................................ | 1 | 1 | 7y1 |
| ..................gguuucguacguaUcagagcagcuc........................................................................ | 1 | 1 | 7y1 |
| ..................gguuucguacAuagcagagcagcuc........................................................................ | 2 | 1 | 7y1 |
| ..................gguuucgCacguagcagagcagcuc........................................................................ | 1 | 1 | 7y1 |
| ..................gguuucUuacguagcagagcagcucc....................................................................... | 1 | 1 | 7y1 |
| ..................gguuucguacguaUcagagcagcucc....................................................................... | 1 | 1 | 7y1 |
| ..................gUuuucguacguagcagagcagcucc....................................................................... | 1 | 1 | 7y1 |
| ..................gguAucguacguagcagagcagcucc....................................................................... | 1 | 1 | 7y1 |
| ..................ggAuucguacguagcagagcagcucc....................................................................... | 4 | 1 | 7y1 |
| ..................gguuucguacgAagcagagcagcucc....................................................................... | 2 | 1 | 7y1 |
| ..................gguuGcguacguagcagagcagcucc....................................................................... | 1 | 1 | 7y1 |
| ..................gguuucguCcguagcagagcagcucc....................................................................... | 1 | 1 | 7y1 |
| ..................gguuucgCacguagcagagcagcucc....................................................................... | 1 | 1 | 7y1 |
| ..................gguuAcguacguagcagagcagcucc....................................................................... | 2 | 1 | 7y1 |
| ..................gguuucguacguagAagagcagcucc....................................................................... | 1 | 1 | 7y1 |
| ..................gguuucguacAuagcagagcagcucc....................................................................... | 1 | 1 | 7y1 |
| ..................gguuucguNcguagcagagcagcucc....................................................................... | 1 | 1 | 7y1 |
| ..................gguuucCuacguagcagagcagcucc....................................................................... | 1 | 1 | 7y1 |
| ..................gguuucguaAguagcagagcagcucc....................................................................... | 2 | 1 | 7y1 |
| ..................gguuucguacCuagcagagcagcucc....................................................................... | 2 | 1 | 7y1 |
| ..................ggGuucguacguagcagagcagcucc....................................................................... | 2 | 1 | 7y1 |
| ..................gguuuAguacguagcagagcagcucc....................................................................... | 5 | 1 | 7y1 |

**Star** **Mature**

| | | | |
|---|---|---|---|
| gaccugcuucugggucgggguuucguacguagcagagcagcucccucgcugcgaucuauugaaagucagcccucgacacaaggguuuguccgcgcgcgcgcgcgcgcgugcgu | | | |
| ..................gCuuucguacguagcagagcagcucc..................................................................... | 1 | 1 | 7y1 |
| ..................gguuucguUcguagcagagcagcucc..................................................................... | 1 | 1 | 7y1 |
| ..................gguAucguacguagcagagcagcuccc.................................................................... | 2 | 1 | 7y1 |
| ..................gguuucUuacguagcagagcagcuccc.................................................................... | 4 | 1 | 7y1 |
| ..................gguuucguacguagGagagcagcuccc.................................................................... | 1 | 1 | 7y1 |
| ..................gguuucguacguaCcagagcagcuccc.................................................................... | 3 | 1 | 7y1 |
| ..................gguuucguacguaUcagagcagcuccc.................................................................... | 1 | 1 | 7y1 |
| ..................gguuucguacgAagcagagcagcuccc.................................................................... | 2 | 1 | 7y1 |
| ..................gguuuUguacguagcagagcagcuccc.................................................................... | 1 | 1 | 7y1 |
| ..................gguuucCuacguagcagagcagcuccc.................................................................... | 3 | 1 | 7y1 |
| ..................gguuucguacUuagcagagcagcuccc.................................................................... | 1 | 1 | 7y1 |
| ..................ggGuucguacguagcagagcagcuccc.................................................................... | 3 | 1 | 7y1 |
| ..................gguuucgGacguagcagagcagcuccc.................................................................... | 2 | 1 | 7y1 |
| ..................gguuucguacguagAagagcagcuccc.................................................................... | 5 | 1 | 7y1 |
| ..................gguuucguaGguagcagagcagcuccc.................................................................... | 2 | 1 | 7y1 |
| ..................ggCuucguacguagcagagcagcuccc.................................................................... | 1 | 1 | 7y1 |
| ..................gCuuucguacguagcagagcagcuccc.................................................................... | 1 | 1 | 7y1 |
| ..................gguuucguacgGagcagagcagcuccc.................................................................... | 1 | 1 | 7y1 |
| ..................gguuuAguacguagcagagcagcuccc.................................................................... | 15 | 1 | 7y1 |
| ..................ggAuucguacguagcagagcagcuccc.................................................................... | 3 | 1 | 7y1 |
| ..................gguuucguaAguagcagagcagcuccc.................................................................... | 11 | 1 | 7y1 |
| ..................gguuucguacCuagcagagcagcuccc.................................................................... | 4 | 1 | 7y1 |
| ..................gguuucgAacguagcagagcagcuccc.................................................................... | 2 | 1 | 7y1 |
| ..................gguuucguacAuagcagagcagcuccc.................................................................... | 1 | 1 | 7y1 |
| ..................gUuuucguacguagcagagcagcuccc.................................................................... | 2 | 1 | 7y1 |
| ..................gguuGcguacguagcagagcagcuccc.................................................................... | 1 | 1 | 7y1 |
| ..................gguuucguacgCagcagagcagcuccc.................................................................... | 1 | 1 | 7y1 |
| ..................gguuucguacguCgcagagcagcucccu................................................................... | 1 | 1 | 7y1 |
| ..................gguuucguacgAagcagagcagcucccu................................................................... | 5 | 1 | 7y1 |
| ..................gguuucguacCuagcagagcagcucccu................................................................... | 2 | 1 | 7y1 |
| ..................gguuucguacguGgcagagcagcucccu................................................................... | 2 | 1 | 7y1 |
| ..................gguuucguacguagAagagcagcucccu................................................................... | 5 | 1 | 7y1 |
| ..................gguuucguacUuagcagagcagcucccu................................................................... | 1 | 1 | 7y1 |
| ..................gguuuAguacguagcagagcagcucccu................................................................... | 9 | 1 | 7y1 |
| ..................ggAuucguacguagcagagcagcucccu................................................................... | 4 | 1 | 7y1 |
| ..................gCuuucguacguagcagagcagcucccu................................................................... | 2 | 1 | 7y1 |
| ..................ggCuucguacguagcagagcagcucccu................................................................... | 1 | 1 | 7y1 |
| ..................gguuucguacguagGagagcagcucccu................................................................... | 1 | 1 | 7y1 |
| ..................gguuucguacguaUcagagcagcucccu................................................................... | 3 | 1 | 7y1 |
| ..................ggGuucguacguagcagagcagcucccu................................................................... | 2 | 1 | 7y1 |
| ..................gguuucguGcguagcagagcagcucccu................................................................... | 1 | 1 | 7y1 |
| ..................gguuucguaAguagcagagcagcucccu................................................................... | 12 | 1 | 7y1 |
| ..................gguuAcguacguagcagagcagcucccu................................................................... | 2 | 1 | 7y1 |
| ..................gAuuucguacguagcagagcagcucccu................................................................... | 1 | 1 | 7y1 |
| ..................gguuucgAacguagcagagcagcucccu................................................................... | 7 | 1 | 7y1 |
| ..................gguuucgCacguagcagagcagcucccu................................................................... | 1 | 1 | 7y1 |
| ..................gguuuUguacguagcagagcagcucccu................................................................... | 1 | 1 | 7y1 |
| ..................gguuucguacguaAcagagcagcucccu................................................................... | 2 | 1 | 7y1 |
| ..................gguuucguCcguagcagagcagcucccu................................................................... | 2 | 1 | 7y1 |
| ..................gguuucUuacguagcagagcagcucccu................................................................... | 5 | 1 | 7y1 |
| ..................gguuucguaUguagcagagcagcucccu................................................................... | 1 | 1 | 7y1 |
| ..................gguuucAuacguagcagagcagcucccu................................................................... | 1 | 1 | 7y1 |
| ..................gUuuucguacguagcagagcagcucccu................................................................... | 1 | 1 | 7y1 |
| ..................gguAucguacguagcagagcagcucccu................................................................... | 3 | 1 | 7y1 |
| ..................gguuucUuacguagcagagcagcucccuc.................................................................. | 1 | 1 | 7y1 |
| ..................gguuAcguacguagcagagcagcucccuc.................................................................. | 1 | 1 | 7y1 |
| ..................gguuucguGcguagcagagcagcucccuc.................................................................. | 1 | 1 | 7y1 |
| ..................gguuuGguacguagcagagcagcucccuc.................................................................. | 1 | 1 | 7y1 |
| ..................gguuuAguacguagcagagcagcucccuc.................................................................. | 1 | 1 | 7y1 |
| ..................gguuucguacAuagcagagcagcucccuc.................................................................. | 2 | 1 | 7y1 |
| ..................gguuucguaAguagcagagcagcucccuc.................................................................. | 1 | 1 | 7y1 |
| ..................ggCuucguacguagcagagcagcucccuc.................................................................. | 1 | 1 | 7y1 |
| ..................gguuucgAacguagcagagcagcucccuc.................................................................. | 2 | 1 | 7y1 |
| ..................ggAuucguacguagcagagcagcucccuc.................................................................. | 2 | 1 | 7y1 |
| ..................gguuucguacUuagcagagcagcucccuc.................................................................. | 2 | 1 | 7y1 |
| ..................gguuucguacguagAagagcagcucccuc.................................................................. | 1 | 1 | 7y1 |
| ..................gguuucCuacguagcagagcagcucccuc.................................................................. | 2 | 1 | 7y1 |
| ..................gguuucguacgAagcagagcagcucccuc.................................................................. | 1 | 1 | 7y1 |
| ..................gguuucguacguaCcagagcagcucccuc.................................................................. | 1 | 1 | 7y1 |
| ..................gguuucguacguaUcagagcagcucccuc.................................................................. | 1 | 1 | 7y1 |

```
   Star                                                                  Mature
gaccugcuucugggucgggguuucguacguagcagagcagcucccucgcugcgaucuauugaaagucagcccucgacacaaggguuuguccgcgcgcgcgcgcgcgcgugcgu
..................gguuucguaGguagcagagcagcucccuc..................................................................   1   1   7y1
..................gguuucgGacguagcagagcagcucccuc..................................................................   1   1   7y1
..................gguuucguacguGgcagagcagcucccuc..................................................................   1   1   7y1
..................gCuuucguacguagcagagcagcucccuc..................................................................   1   1   7y1
..................gguuucguacguaCcagagcagcucccucg.................................................................   1   1   7y1
..................gguuucguacgAagcagagcagcucccucg.................................................................   3   1   7y1
..................gguuAcguacguagcagagcagcucccucg.................................................................   1   1   7y1
..................ggAuucguacguagcagagcagcucccucg.................................................................   3   1   7y1
..................gguuucguUcguagcagagcagcucccucg.................................................................   1   1   7y1
..................gguuucgGacguagcagagcagcucccucg.................................................................   1   1   7y1
..................gguuucguacguagUagagcagcucccucg.................................................................   1   1   7y1
..................gguuuAguacguagcagagcagcucccucg.................................................................   3   1   7y1
..................gCuuucguacguagcagagcagcucccucg.................................................................   1   1   7y1
..................gguuucguacguagAagagcagcucccucg.................................................................   2   1   7y1
..................gguuucCuacguagcagagcagcucccucg.................................................................   2   1   7y1
..................gguuucguacguagGagagcagcucccucg.................................................................   1   1   7y1
..................gguuucguaAguagcagagcagcucccucg.................................................................   1   1   7y1
..................gguuucguaUguagcagagcagcucccucg.................................................................   2   1   7y1
..................gguAucguacguagcagagcagcucccucg.................................................................   2   1   7y1
..................gguuucguacguaAcagagcagcucccucgc................................................................   1   1   7y1
..................gguCucguacguagcagagcagcucccucgc................................................................   1   1   7y1
..................gCuuucguacguagcagagcagcucccucgc................................................................   1   1   7y1
..................gguuucAuacguagcagagcagcucccucgc................................................................   1   1   7y1
..................ggGuucguacguagcagagcagcucccucgc................................................................   2   1   7y1
..................gguuucguacUuagcagagcagcucccucgc................................................................   1   1   7y1
..................gguuucguacguagUagagcagcucccucgc................................................................   1   1   7y1
..................gguuucguacgAagcagagcagcucccucgc................................................................   2   1   7y1
..................gguuuAguacguagcagagcagcucccucgc................................................................   1   1   7y1
..................gguuucguacguGgcagagcagcucccucgc................................................................   1   1   7y1
..................gguGucguacguagcagagcagcucccucgc................................................................   1   1   7y1
..................ggAuucguacguagcagagcagcucccucgc................................................................   2   1   7y1
..................gguAucguacguagcagagcagcucccucgc................................................................   1   1   7y1
..................gguuucCuacguagcagagcagcucccucgc................................................................   4   1   7y1
..................gUuuucguacguagcagagcagcucccucgc................................................................   2   1   7y1
..................gguuucgAacguagcagagcagcucccucgc................................................................   4   1   7y1
..................gguuCcguacguagcagagcagcucccucgc................................................................   1   1   7y1
..................gguuucguaAguagcagagcagcucccucgc................................................................   5   1   7y1
..................gguuucguacguagAagagcagcucccucgcu...............................................................   1   1   7y1
..................gguuucUuacguagcagagcagcucccucgcu...............................................................   1   1   7y1
..................ggAuucguacguagcagagcagcucccucgcu...............................................................   1   1   7y1
..................gguuucguacCuagcagagcagcucccucgcu...............................................................   1   1   7y1
..................gguuucguGcguagcagagcagcucccucgcu...............................................................   1   1   7y1
..................gguuucguacguaUcagagcagcucccucgcu...............................................................   1   1   7y1
..................gguuucguacguaCcagagcagcucccucgcu...............................................................   1   1   7y1
..................gUuuucguacguagcagagcagcucccucgcu...............................................................   1   1   7y1
..................gguuucAuacguagcagagcagcucccucgcu...............................................................   1   1   7y1
...................guuGcguacguagcagagc...........................................................................   1   1   7y1
...................guuucguaAguagcagagc...........................................................................   5   1   7y1
...................Nuuucguacguagcagagc...........................................................................   1   1   7y1
...................Uuuucguacguagcagagc...........................................................................   1   1   7y1
...................guuuAguacguagcagagc...........................................................................   1   1   7y1
...................guuucCuacguagcagagc...........................................................................   1   1   7y1
...................Cuuucguacguagcagagc...........................................................................   3   1   7y1
...................guuucgAacguagcagagc...........................................................................   3   1   7y1
...................guuucUuacguagcagagc...........................................................................   1   1   7y1
...................guuAcguacguagcagagca..........................................................................   1   1   7y1
...................guuuAguacguagcagagca..........................................................................   1   1   7y1
...................guuucguacguagAagagca..........................................................................   2   1   7y1
...................gAuucguacguagcagagca..........................................................................   1   1   7y1
...................guAucguacguagcagagca..........................................................................   1   1   7y1
...................guuCcguacguagcagagca..........................................................................   1   1   7y1
...................guuucCuacguagcagagca..........................................................................   2   1   7y1
...................guuucguacCuagcagagca..........................................................................   2   1   7y1
...................Uuuucguacguagcagagca..........................................................................   1   1   7y1
...................guuucgAacguagcagagca..........................................................................   5   1   7y1
...................Cuuucguacguagcagagca..........................................................................   2   1   7y1
...................guuucguacgAagcagagcagc........................................................................   1   1   7y1
...................Auuucguacguagcagagcagc........................................................................   1   1   7y1
...................guuucguaAguagcagagcagc........................................................................   2   1   7y1
...................gAuucguacguagcagagcagc........................................................................   5   1   7y1
```

**Star** **Mature**

| | | | |
|---|---|---|---|
| gaccugcuucugggucgggguuucguacguagcagagcagcucccucgcugcgaucuauugaaagucagcccucgacacaaggguuuguccgcgcgcgcgcgcgcgcgugcgu | | | |
| ...................guAucguacguagcagagcagc...................................................................... | 3 | 1 | 7y1 |
| ...................Uuuucguacguagcagagcagc...................................................................... | 286 | 1 | 7y1 |
| ...................guuucgAacguagcagagcagc...................................................................... | 2 | 1 | 7y1 |
| ...................guuucguUcguagcagagcagc...................................................................... | 1 | 1 | 7y1 |
| ...................guAucguacguagcagagcagcu..................................................................... | 4 | 1 | 7y1 |
| ...................guuucguacgAagcagagcagcu..................................................................... | 3 | 1 | 7y1 |
| ...................Uuuucguacguagcagagcagcu..................................................................... | 3 | 1 | 7y1 |
| ...................gGuucguacguagcagagcagcu..................................................................... | 1 | 1 | 7y1 |
| ...................Cuuucguacguagcagagcagcu..................................................................... | 2 | 1 | 7y1 |
| ...................guuucgAacguagcagagcagcu..................................................................... | 3 | 1 | 7y1 |
| ...................guuucguacUuagcagagcagcu..................................................................... | 1 | 1 | 7y1 |
| ...................Nuuucguacguagcagagcagcu..................................................................... | 3 | 1 | 7y1 |
| ...................gAuucguacguagcagagcagcu..................................................................... | 6 | 1 | 7y1 |
| ...................guuucguacguGgcagagcagcu..................................................................... | 1 | 1 | 7y1 |
| ...................guuAcguacguagcagagcagcu..................................................................... | 2 | 1 | 7y1 |
| ...................guuucguacguagAagagcagcu..................................................................... | 1 | 1 | 7y1 |
| ...................guuucUuacguagcagagcagcu..................................................................... | 1 | 1 | 7y1 |
| ...................guuuAguacguagcagagcagcu..................................................................... | 1 | 1 | 7y1 |
| ...................gAuucguacguagcagagcagcuc.................................................................... | 2 | 1 | 7y1 |
| ...................guuuUguacguagcagagcagcuc.................................................................... | 1 | 1 | 7y1 |
| ...................guAucguacguagcagagcagcuc.................................................................... | 3 | 1 | 7y1 |
| ...................guCucguacguagcagagcagcuc.................................................................... | 1 | 1 | 7y1 |
| ...................gGuucguacguagcagagcagcuc.................................................................... | 1 | 1 | 7y1 |
| ...................Uuuucguacguagcagagcagcucc................................................................... | 1 | 1 | 7y1 |
| ...................guuucguacgAagcagagcagcucc................................................................... | 1 | 1 | 7y1 |
| ...................guAucguacguagcagagcagcucc................................................................... | 1 | 1 | 7y1 |
| ...................guuucguacguaAcagagcagcucc................................................................... | 1 | 1 | 7y1 |
| ...................guuucguacguaUcagagcagcucc................................................................... | 179 | 1 | 7y1 |
| ...................guuucguGcguagcagagcagcucc................................................................... | 1 | 1 | 7y1 |
| ...................gAuucguacguagcagagcagcuccc.................................................................. | 4 | 1 | 7y1 |
| ...................guuucguacguagAagagcagcuccc.................................................................. | 5 | 1 | 7y1 |
| ...................guuucguacguCgcagagcagcuccc.................................................................. | 1 | 1 | 7y1 |
| ...................guAucguacguagcagagcagcuccc.................................................................. | 2 | 1 | 7y1 |
| ...................Cuuucguacguagcagagcagcuccc.................................................................. | 2 | 1 | 7y1 |
| ...................guuucguacCuagcagagcagcuccc.................................................................. | 3 | 1 | 7y1 |
| ...................guuucguaAguagcagagcagcuccc.................................................................. | 2 | 1 | 7y1 |
| ...................guuAcguacguagcagagcagcuccc.................................................................. | 1 | 1 | 7y1 |
| ...................guuucgAacguagcagagcagcuccc.................................................................. | 2 | 1 | 7y1 |
| ...................guuucCuacguagcagagcagcuccc.................................................................. | 2 | 1 | 7y1 |
| ...................gCuucguacguagcagagcagcuccc.................................................................. | 2 | 1 | 7y1 |
| ...................guuucguacguaAcagagcagcuccc.................................................................. | 2 | 1 | 7y1 |
| ...................guuucguacgGagcagagcagcuccc.................................................................. | 1 | 1 | 7y1 |
| ...................guuucguacgAagcagagcagcuccc.................................................................. | 1 | 1 | 7y1 |
| ...................guuuAguacguagcagagcagcuccc.................................................................. | 3 | 1 | 7y1 |
| ...................guuuUguacguagcagagcagcuccc.................................................................. | 1 | 1 | 7y1 |
| ...................guuucguacguaCcagagcagcuccc.................................................................. | 3 | 1 | 7y1 |
| ...................guuucguacguaUcagagcagcuccc.................................................................. | 1 | 1 | 7y1 |
| ...................guuuAguacguagcagagcagcucccu................................................................. | 1 | 1 | 7y1 |
| ...................guuAcguacguagcagagcagcucccu................................................................. | 2 | 1 | 7y1 |
| ...................guuucCuacguagcagagcagcucccu................................................................. | 1 | 1 | 7y1 |
| ...................guuucguacgGagcagagcagcucccu................................................................. | 1 | 1 | 7y1 |
| ...................guuucguacAuagcagagcagcucccu................................................................. | 2 | 1 | 7y1 |
| ...................gGuucguacguagcagagcagcucccu................................................................. | 295 | 1 | 7y1 |
| ...................guuucguacgCagcagagcagcucccu................................................................. | 1 | 1 | 7y1 |
| ...................guAucguacguagcagagcagcucccu................................................................. | 2 | 1 | 7y1 |
| ...................guuucguacguagcGgagcagcucccu................................................................. | 2 | 1 | 7y1 |
| ...................Cuuucguacguagcagagcagcucccu................................................................. | 3 | 1 | 7y1 |
| ...................guuucguacCuagcagagcagcucccu................................................................. | 1 | 1 | 7y1 |
| ...................guuucgAacguagcagagcagcucccu................................................................. | 7 | 1 | 7y1 |
| ...................gAuucguacguagcagagcagcucccu................................................................. | 7 | 1 | 7y1 |
| ...................guuucguacguagAagagcagcucccu................................................................. | 3 | 1 | 7y1 |
| ...................Uuuucguacguagcagagcagcucccu................................................................. | 2 | 1 | 7y1 |
| ...................guuucguacguagcGgagcagcucccuc................................................................ | 1 | 1 | 7y1 |
| ...................guuucguGcguagcagagcagcucccuc................................................................ | 2 | 1 | 7y1 |
| ...................gGuucguacguagcagagcagcucccuc................................................................ | 1 | 1 | 7y1 |
| ...................guuucguacgAagcagagcagcucccuc................................................................ | 1 | 1 | 7y1 |
| ...................guuucguaAguagcagagcagcucccuc................................................................ | 1 | 1 | 7y1 |
| ...................guuucguacUuagcagagcagcucccuc................................................................ | 3 | 1 | 7y1 |
| ...................guuucgAacguagcagagcagcucccuc................................................................ | 2 | 1 | 7y1 |
| ...................guuucguacAuagcagagcagcucccuc................................................................ | 1 | 1 | 7y1 |

```
  Star                                                                    Mature
gaccugcuucugggucgggguuucguacguagcagagcagcucccucgcugcgaucuauugaaagucagcccucgacacaaggguuuguccgcgcgcgcgcgcgcgcgugcgu
...................guuCcguacguagcagagcagcucccuc..................................................................    1      1      7y1
...................guAucguacguagcagagcagcucccuc..................................................................    1      1      7y1
...................Cuuucguacguagcagagcagcucccuc..................................................................    1      1      7y1
...................guuucguacguagAagagcagcucccuc..................................................................    2      1      7y1
...................guuucguacguaUcagagcagcucccuc..................................................................    1      1      7y1
...................guuucguacCuagcagagcagcucccuc..................................................................    1      1      7y1
...................Uuuucguacguagcagagcagcucccuc..................................................................    3      1      7y1
...................guuucguacguaUcagagcagcucccucg.................................................................    1      1      7y1
...................guuucgAacguagcagagcagcucccucg.................................................................    4      1      7y1
...................gAuucguacguagcagagcagcucccucg.................................................................    1      1      7y1
...................guAucguacguagcagagcagcucccucg.................................................................    1      1      7y1
...................guuucguacAuagcagagcagcucccucg.................................................................    1      1      7y1
...................guuucguacguagcGgagcagcucccucg.................................................................    1      1      7y1
...................guuucguacgCagcagagcagcucccucgc................................................................    1      1      7y1
...................guuucCuacguagcagagcagcucccucgc................................................................    1      1      7y1
...................guuucguacCuagcagagcagcucccucgc................................................................    1      1      7y1
...................guuucguacUuagcagagcagcucccucgc................................................................    2      1      7y1
...................Uuuucguacguagcagagcagcucccucgc................................................................    1      1      7y1
...................guuucguaAguagcagagcagcucccucgc................................................................    3      1      7y1
...................gAuucguacguagcagagcagcucccucgc................................................................    1      1      7y1
...................guuucgGacguagcagagcagcucccucgc................................................................    1      1      7y1
...................gGuucguacguagcagagcagcucccucgc................................................................    1      1      7y1
...................guuucguacguaUcagagcagcucccucgc................................................................    2      1      7y1
...................Nuuucguacguagcagagcagcucccucgc................................................................    1      1      7y1
...................guAucguacguagcagagcagcucccucgc................................................................    2      1      7y1
...................guuucguacgAagcagagcagcucccucgc................................................................    1      1      7y1
...................guuucgAacguagcagagcagcucccucgc................................................................    4      1      7y1
...................guuAcguacguagcagagcagcucccucgc................................................................    2      1      7y1
...................Cuuucguacguagcagagcagcucccucgc................................................................    1      1      7y1
...................guuucUuacguagcagagcagcucccucgc................................................................    1      1      7y1
...................guuucguacguagAagagcagcucccucgc................................................................    1      1      7y1
...................guuucguacguagAagagcagcucccucgcu...............................................................    1      1      7y1
...................guuucguaGguagcagagcagcucccucgcu...............................................................    1      1      7y1
...................guuucUuacguagcagagcagcucccucgcu...............................................................    1      1      7y1
...................gAuucguacguagcagagcagcucccucgcu...............................................................    2      1      7y1
...................Cuuucguacguagcagagcagcucccucgcu...............................................................    2      1      7y1
...................gAuucguacguagcagagcagcucccucgcug..............................................................    1      1      7y1
...................guuucguacguagAagagcagcucccucgcug..............................................................    1      1      7y1
...................guuucguacUuagcagagcagcucccucgcug..............................................................    1      1      7y1
....................uuucguacgCagcagagc...........................................................................    1      1      7y1
....................uuucguacguagcagaUc...........................................................................    5      1      7y1
....................Auucguacguagcagagc...........................................................................    52     1      7y1
....................uuAcguacguagcagagc...........................................................................    2      1      7y1
....................uuucCuacguagcagagc...........................................................................    192    1      7y1
....................uuucguacguaCcagagc...........................................................................    1      1      7y1
....................uuucguacgAagcagagc...........................................................................    11     1      7y1
....................uuucguacguagcGgagc...........................................................................    2      1      7y1
....................uuucgGacguagcagagc...........................................................................    1      1      7y1
....................Guucguacguagcagagc...........................................................................    9      1      7y1
....................uuucguacguaUcagagc...........................................................................    2      1      7y1
....................Nuucguacguagcagagc...........................................................................    3      1      7y1
....................uuucguacguagcagagc...........................................................................    4344   0      7y1
....................uuucAuacguagcagagc...........................................................................    1      1      7y1
....................uuucguacguagcUgagc...........................................................................    2      1      7y1
....................uuucUuacguagcagagc...........................................................................    1      1      7y1
....................uuucgAacguagcagagc...........................................................................    8      1      7y1
....................Cuucguacguagcagagc...........................................................................    2      1      7y1
....................uuucguacCuagcagagc...........................................................................    4      1      7y1
....................uuucguacguagcaUagc...........................................................................    1      1      7y1
....................uAucguacguagcagagc...........................................................................    9      1      7y1
....................uuuAguacguagcagagc...........................................................................    5      1      7y1
....................uuucguacguagcagaCc...........................................................................    2      1      7y1
....................uuucguacUuagcagagc...........................................................................    1      1      7y1
....................uuucguacguGgcagagc...........................................................................    2      1      7y1
....................uuucguaUguagcagagc...........................................................................    1      1      7y1
....................uuucguacguagcagagA...........................................................................    7      1      7y1
....................uuucguacguagAagagc...........................................................................    4      1      7y1
....................uuuGguacguagcagagc...........................................................................    3      1      7y1
....................uuucguaAguagcagagc...........................................................................    8      1      7y1
....................uuucguaGguagcagagca..........................................................................    1      1      7y1
```

# Star    Mature

| | | | |
|---|---|---|---|
| gaccugcuucugggucgggguuucguacguagcagagcagcucccucgcugcgaucuauugaaagucagcccucgacacaaggguuuguccgcgcgcgcgcgcgcgcgugcgu | | | |
| ....................uuucguacguagcagagAa.......................................................................... | 4 | 1 | 7y1 |
| ....................uuucguacguagAagagca.......................................................................... | 3 | 1 | 7y1 |
| ....................uCucguacguagcagagca.......................................................................... | 1 | 1 | 7y1 |
| ....................uuucguacguagcagCgca.......................................................................... | 3 | 1 | 7y1 |
| ....................uuuAguacguagcagagca.......................................................................... | 2 | 1 | 7y1 |
| ....................uuucguaAguagcagagca.......................................................................... | 2 | 1 | 7y1 |
| ....................Nuucguacguagcagagca.......................................................................... | 3 | 1 | 7y1 |
| ....................Auucguacguagcagagca.......................................................................... | 24 | 1 | 7y1 |
| ....................uuucguacguagcagaUca.......................................................................... | 1 | 1 | 7y1 |
| ....................uuucguacguaCcagagca.......................................................................... | 1 | 1 | 7y1 |
| ....................uuucguacguagcagagca.......................................................................... | 1926 | 0 | 7y1 |
| ....................uuucguGcguagcagagca.......................................................................... | 1 | 1 | 7y1 |
| ....................uuucguacguUgcagagca.......................................................................... | 2 | 1 | 7y1 |
| ....................uuucguacguagcagaCca.......................................................................... | 3 | 1 | 7y1 |
| ....................uuucguacguagcagagGa.......................................................................... | 1 | 1 | 7y1 |
| ....................uuucguacguagUagagca.......................................................................... | 1 | 1 | 7y1 |
| ....................uuucgAacguagcagagca.......................................................................... | 2 | 1 | 7y1 |
| ....................uuucguacguaUcagagca.......................................................................... | 2 | 1 | 7y1 |
| ....................uuucguacCuagcagagca.......................................................................... | 2 | 1 | 7y1 |
| ....................uuucguacgGagcagagca.......................................................................... | 1 | 1 | 7y1 |
| ....................uuucguacguagcGgagca.......................................................................... | 1 | 1 | 7y1 |
| ....................uuucCuacguagcagagca.......................................................................... | 1 | 1 | 7y1 |
| ....................uuucguCcguagcagagca.......................................................................... | 1 | 1 | 7y1 |
| ....................uuucguacUuagcagagca.......................................................................... | 1 | 1 | 7y1 |
| ....................uuucguacgCagcagagca.......................................................................... | 1 | 1 | 7y1 |
| ....................uuucguacguagcUgagca.......................................................................... | 1 | 1 | 7y1 |
| ....................Guucguacguagcagagca.......................................................................... | 4 | 1 | 7y1 |
| ....................uuucgCacguagcagagca.......................................................................... | 1 | 1 | 7y1 |
| ....................uAucguacguagcagagca.......................................................................... | 5 | 1 | 7y1 |
| ....................uuucguacguagcaUagca.......................................................................... | 1 | 1 | 7y1 |
| ....................uuucguacguagcaUagcag......................................................................... | 1 | 1 | 7y1 |
| ....................uuucguacguagcagagcaU......................................................................... | 2 | 1 | 7y1 |
| ....................uuCcguacguagcagagcag......................................................................... | 1 | 1 | 7y1 |
| ....................uuucguacgAagcagagcag......................................................................... | 1 | 1 | 7y1 |
| ....................uuAcguacguagcagagcag......................................................................... | 1 | 1 | 7y1 |
| ....................uAucguacguagcagagcag......................................................................... | 4 | 1 | 7y1 |
| ....................uuucguacguagcagagAag......................................................................... | 1 | 1 | 7y1 |
| ....................uuucguacguagcGgagcag......................................................................... | 1 | 1 | 7y1 |
| ....................uuucguacguagcagaUcag......................................................................... | 1 | 1 | 7y1 |
| ....................uuucguacguagcagagcaC......................................................................... | 2 | 1 | 7y1 |
| ....................Auucguacguagcagagcag......................................................................... | 11 | 1 | 7y1 |
| ....................uuucgCacguagcagagcag......................................................................... | 1 | 1 | 7y1 |
| ....................uuucguacguagcagagcag......................................................................... | 641 | 0 | 7y1 |
| ....................uuucguacguagcagagGagc........................................................................ | 2 | 1 | 7y1 |
| ....................uuucgAacguagcagagcagc........................................................................ | 3 | 1 | 7y1 |
| ....................uuucgGacguagcagagcagc........................................................................ | 3 | 1 | 7y1 |
| ....................uuucguGcguagcagagcagc........................................................................ | 1 | 1 | 7y1 |
| ....................uuucguaAguagcagagcagc........................................................................ | 5 | 1 | 7y1 |
| ....................uuucguaGguagcagagcagc........................................................................ | 1 | 1 | 7y1 |
| ....................uuucguacguagcagagcaAc........................................................................ | 1 | 1 | 7y1 |
| ....................uuucguaUguagcagagcagc........................................................................ | 1 | 1 | 7y1 |
| ....................uuucguacguagAagagcagc........................................................................ | 2 | 1 | 7y1 |
| ....................uuucguCcguagcagagcagc........................................................................ | 1 | 1 | 7y1 |
| ....................uuucguacguagcagagcagc........................................................................ | 2127 | 0 | 7y1 |
| ....................Cuucguacguagcagagcagc........................................................................ | 237 | 1 | 7y1 |
| ....................uuucgCacguagcagagcagc........................................................................ | 1 | 1 | 7y1 |
| ....................Auucguacguagcagagcagc........................................................................ | 18 | 1 | 7y1 |
| ....................uuucguacgAagcagagcagc........................................................................ | 2 | 1 | 7y1 |
| ....................uuucguacguagUagagcagc........................................................................ | 1 | 1 | 7y1 |
| ....................uuucguacguagcGgagcagc........................................................................ | 1 | 1 | 7y1 |
| ....................uuucguacUuagcagagcagc........................................................................ | 1 | 1 | 7y1 |
| ....................uAucguacguagcagagcagc........................................................................ | 5 | 1 | 7y1 |
| ....................Nuucguacguagcagagcagc........................................................................ | 1 | 1 | 7y1 |
| ....................uuucguacguagcagagcaCc........................................................................ | 1 | 1 | 7y1 |
| ....................uuucguacguagcagagcaUc........................................................................ | 1 | 1 | 7y1 |
| ....................uuucguacguagGagagcagc........................................................................ | 1 | 1 | 7y1 |
| ....................uuucguacguagcagagAagc........................................................................ | 3 | 1 | 7y1 |
| ....................uuucguacguagcaUagcagc........................................................................ | 1 | 1 | 7y1 |
| ....................uuucguacguagcagagcagA........................................................................ | 1 | 1 | 7y1 |
| ....................uuuGguacguagcagagcagc........................................................................ | 2 | 1 | 7y1 |

### Star — Mature

```
gaccugcuucugggucgggguuucguacguagcagagcagcucccucgcugcgaucuauugaaagucagcccucgacacaaggguuuguccgcgcgcgcgcgcgcgugcgu
....................Guucguacguagcagagcagc..........................................................................  3     1  7y1
....................uuucguacguagcagCgcagc..........................................................................  5     1  7y1
....................uuucguacguagcUgagcagcu.........................................................................  1     1  7y1
....................uuucguacUuagcagagcagcu.........................................................................  1     1  7y1
....................uuucguacguagcagGgcagcu.........................................................................  1     1  7y1
....................uuucguacguagcaUagcagcu.........................................................................  272   1  7y1
....................uuucguacguagcagagcagAu.........................................................................  1     1  7y1
....................Nuucguacguagcagagcagcu.........................................................................  2     1  7y1
....................uuucguacguagcagagcagcC.........................................................................  1     1  7y1
....................uuucguacguagcagagGagcu.........................................................................  2     1  7y1
....................uAucguacguagcagagcagcu.........................................................................  10    1  7y1
....................uuucguacguaAcagagcagcu.........................................................................  1     1  7y1
....................uuAcguacguagcagagcagcu.........................................................................  3     1  7y1
....................uuucAuacguagcagagcagcu.........................................................................  2     1  7y1
....................uuucguaUguagcagagcagcu.........................................................................  1     1  7y1
....................uuucguacguagcagagAagcu.........................................................................  9     1  7y1
....................uuuUguacguagcagagcagcu.........................................................................  2     1  7y1
....................uuucguaAguagcagagcagcu.........................................................................  8     1  7y1
....................uuucguacAuagcagagcagcu.........................................................................  1     1  7y1
....................uuucguacguaCcagagcagcu.........................................................................  1     1  7y1
....................uuuAguacguagcagagcagcu.........................................................................  7     1  7y1
....................Cuucguacguagcagagcagcu.........................................................................  3     1  7y1
....................uuucguacguagAagagcagcu.........................................................................  2     1  7y1
....................uuucguacguagcagagcagcG.........................................................................  22    1  7y1
....................Auucguacguagcagagcagcu.........................................................................  37    1  7y1
....................uuucguacguagcagaCcagcu.........................................................................  1     1  7y1
....................uuucgAacguagcagagcagcu.........................................................................  4     1  7y1
....................uuucguacguagcagagcagcA.........................................................................  3     1  7y1
....................uuucguacCuagcagagcagcu.........................................................................  4     1  7y1
....................uuucguacguagcagCgcagcu.........................................................................  4     1  7y1
....................uuucguacguagcagagcagcu.........................................................................  4088  0  7y1
....................uuucguacguagcagagcaUcu.........................................................................  4     1  7y1
....................uuucguacguagcagagcCgcu.........................................................................  3     1  7y1
....................uuucguacguagUagagcagcu.........................................................................  1     1  7y1
....................uuucgCacguagcagagcagcu.........................................................................  1     1  7y1
....................uuucguacguagcagagcaCcu.........................................................................  1     1  7y1
....................uuucguGcguagcagagcagcu.........................................................................  1     1  7y1
....................uuucguacgAagcagagcagcu.........................................................................  11    1  7y1
....................Guucguacguagcagagcagcu.........................................................................  7     1  7y1
....................uuucguacguagcagUgcagcu.........................................................................  2     1  7y1
....................uuucguacguagcaCagcagcu.........................................................................  2     1  7y1
....................uuucCuacguagcagagcagcu.........................................................................  2     1  7y1
....................uuucguacguagcagagcagGuc........................................................................  1     1  7y1
....................uuucguacguagcagagcagcuc........................................................................  7935  0  7y1
....................uuucAuacguagcagagcagcuc........................................................................  1     1  7y1
....................uuucguacguagcagagAagcuc........................................................................  10    1  7y1
....................uuucguacguagcagagcUgcuc........................................................................  1     1  7y1
....................uuucUuacguagcagagcagcuc........................................................................  6     1  7y1
....................uuucguacguagcUgagcagcuc........................................................................  1     1  7y1
....................uuucguacguagcagUgcagcuc........................................................................  1     1  7y1
....................uuucgAacguagcagagcagcuc........................................................................  16    1  7y1
....................uuucguacguagcagagcCgcuc........................................................................  3     1  7y1
....................uuucguacguagcagagcagcuA........................................................................  8     1  7y1
....................uuucguacguagcagaUcagcuc........................................................................  3     1  7y1
....................uuucguacguagcagaCcagcuc........................................................................  4     1  7y1
....................Auucguacguagcagagcagcuc........................................................................  89    1  7y1
....................uuucguacguagcagagcGgcuc........................................................................  2     1  7y1
....................uuucguacguagcagagcaCcuc........................................................................  4     1  7y1
....................uuuUguacguagcagagcagcuc........................................................................  1     1  7y1
....................uuucguacguaCcagagcagcuc........................................................................  2     1  7y1
....................uuucguacguagcagagGagcuc........................................................................  1     1  7y1
....................uuucguacgCagcagagcagcuc........................................................................  1     1  7y1
....................uuucguacguagcagGgcagcuc........................................................................  1     1  7y1
....................uuucguacguagAagagcagcuc........................................................................  9     1  7y1
....................uuGcguacguagcagagcagcuc........................................................................  1     1  7y1
....................uuucguacguagcaCagcagcuc........................................................................  1     1  7y1
....................uuuAguacguagcagagcagcuc........................................................................  12    1  7y1
....................uuucCuacguagcagagcagcuc........................................................................  2     1  7y1
....................uuucguacCuagcagagcagcuc........................................................................  4     1  7y1
....................uuucguacguagcagagcagcGc........................................................................  1     1  7y1
```

**Star** **Mature**

```
gaccugcuucugggucggggguuucguacguagcagagcagcucccucgcugcgaucuauugaaagucagcccucgacacaaggguuuguccgcgcgcgcgcgcgcgcgugcgu
....................uuucguacguagcagagcagcuG......................................................................   1      1      7y1
....................uuucguacguagcCgagcagcuc......................................................................   1      1      7y1
....................uuucguacAuagcagagcagcuc......................................................................   1      1      7y1
....................Guucguacguagcagagcagcuc......................................................................   13     1      7y1
....................uuucguaAguagcagagcagcuc......................................................................   9      1      7y1
....................uuucguacguagcagagcagcAc......................................................................   2      1      7y1
....................uuCcguacguagcagagcagcuc......................................................................   1      1      7y1
....................uuucguacguagcagagcaUcuc......................................................................   7      1      7y1
....................uuucguacguGgcagagcagcuc......................................................................   2      1      7y1
....................uuucguacguagcGgagcagcuc......................................................................   1      1      7y1
....................uuucguacguagcagagcagcuU......................................................................   1      1      7y1
....................uuucgCacguagcagagcagcuc......................................................................   1      1      7y1
....................uuucguacgAagcagagcagcuc......................................................................   11     1      7y1
....................uuucguacUuagcagagcagcuc......................................................................   3      1      7y1
....................uuucguaUguagcagagcagcuc......................................................................   1      1      7y1
....................uuucguacguagcagCgcagcuc......................................................................   7      1      7y1
....................uuuGguacguagcagagcagcuc......................................................................   3      1      7y1
....................uuucguCcguagcagagcagcuc......................................................................   2      1      7y1
....................uuucguacguagcagagcagAuc......................................................................   5      1      7y1
....................Nuucguacguagcagagcagcuc......................................................................   4      1      7y1
....................uAucguacguagcagagcagcuc......................................................................   11     1      7y1
....................uuucguacguaUcagagcagcuc......................................................................   1      1      7y1
....................Cuucguacguagcagagcagcuc......................................................................   175    1      7y1
....................uuucguGcguagcagagcagcuc......................................................................   1      1      7y1
....................uuucguacguagUagagcagcuc......................................................................   2      1      7y1
....................uuucguacguagcaUagcagcuc......................................................................   2      1      7y1
....................uuAcguacguagcagagcagcuc......................................................................   3      1      7y1
....................uCucguacguagcagagcagcuc......................................................................   1      1      7y1
....................uuucguacguagcagagcagcucG.....................................................................   1      1      7y1
....................uuucguacguagcagGgcagcucc.....................................................................   4      1      7y1
....................uuucguacgGagcagagcagcucc.....................................................................   1      1      7y1
....................uuucguacguagcagagcagcuUc.....................................................................   1      1      7y1
....................uuucguaAguagcagagcagcucc.....................................................................   7      1      7y1
....................uuucguacguagcagagcagcAcc.....................................................................   1      1      7y1
....................uuucguacguagcaUagcagcucc.....................................................................   1      1      7y1
....................uuucguacCuagcagagcagcucc.....................................................................   4      1      7y1
....................uuucguacguagcaCagcagcucc.....................................................................   1      1      7y1
....................uuucguacguaUcagagcagcucc.....................................................................   3      1      7y1
....................uuucguacguagcagagcCgcucc.....................................................................   1      1      7y1
....................uuucguacguagcagagcaUcucc.....................................................................   3      1      7y1
....................uuucCuacguagcagagcagcucc.....................................................................   1      1      7y1
....................uuucguacguagcagagcagcucc.....................................................................   5981   0      7y1
....................uuucguacguagcagaCcagcucc.....................................................................   3      1      7y1
....................uuucguacguagcagagcagcucU.....................................................................   2      1      7y1
....................Auucguacguagcagagcagcucc.....................................................................   67     1      7y1
....................uuucguacguagcagagcGgcucc.....................................................................   1      1      7y1
....................uuucguacguagcagagcagcCcc.....................................................................   2      1      7y1
....................Nuucguacguagcagagcagcucc.....................................................................   3      1      7y1
....................uuuGguacguagcagagcagcucc.....................................................................   2      1      7y1
....................uuucguacUuagcagagcagcucc.....................................................................   2      1      7y1
....................uuucguacguagcagCgcagcucc.....................................................................   5      1      7y1
....................uuucguacguagcagagcagcucA.....................................................................   2      1      7y1
....................uuucguacgAagcagagcagcucc.....................................................................   7      1      7y1
....................uuucUuacguagcagagcagcucc.....................................................................   2      1      7y1
....................uuucgCacguagcagagcagcucc.....................................................................   2      1      7y1
....................uuAcguacguagcagagcagcucc.....................................................................   1      1      7y1
....................uuCcguacguagcagagcagcucc.....................................................................   2      1      7y1
....................uuGcguacguagcagagcagcucc.....................................................................   1      1      7y1
....................Guucguacguagcagagcagcucc.....................................................................   14     1      7y1
....................uuucgAacguagcagagcagcucc.....................................................................   11     1      7y1
....................uuucguacguagcagagcagAucc.....................................................................   4      1      7y1
....................uuucguacguagcagagcaCcucc.....................................................................   3      1      7y1
....................uuucguacguUgcagagcagcucc.....................................................................   1      1      7y1
....................uAucguacguagcagagcagcucc.....................................................................   9      1      7y1
....................uuucguacgCagcagagcagcucc.....................................................................   1      1      7y1
....................uuucguacguagAagagcagcucc.....................................................................   9      1      7y1
....................uuucguacguagcGgagcagcucc.....................................................................   1      1      7y1
....................uuucguacguagcagagAagcucc.....................................................................   7      1      7y1
....................uuucguaGguagcagagcagcucc.....................................................................   2      1      7y1
....................uuucguaUguagcagagcagcucc.....................................................................   1      1      7y1
```

Star Mature

```
gaccugcuucugggucgggguuucguacguagcagagcagcucccucgcugcgaucuauugaaagucagcccucgacacaaggguuuguccgcgcgcgcgcgcgcgcgugcgu
....................uuucguGcguagcagagcagcucccu...................................................................  1     1  7y1
....................uuucguacguagcagagcagcCcccu...................................................................  1     1  7y1
....................uuucguacAuagcagagcagcucccu...................................................................  3     1  7y1
....................Guucguacguagcagagcagcucccu...................................................................  14    1  7y1
....................uuucguacguagcaAagcagcucccu...................................................................  1     1  7y1
....................uuucguaUguagcagagcagcucccu...................................................................  2     1  7y1
....................uuucguacguagcagagcagcuccAu...................................................................  4     1  7y1
....................uuucguacguagcagagcagcucccG...................................................................  17    1  7y1
....................uuucgAacguagcagagcagcucccu...................................................................  9     1  7y1
....................uuGcguacguagcagagcagcucccu...................................................................  1     1  7y1
....................uuucguacguagcagagcagcucccA...................................................................  8     1  7y1
....................uuucguaAguagcagagcagcucccu...................................................................  4     1  7y1
....................uuucguacguagUagagcagcucccu...................................................................  225   1  7y1
....................uuucguacguagcagagcagcucAcu...................................................................  3     1  7y1
....................uuucguacguagAagagcagcucccu...................................................................  2     1  7y1
....................Nuucguacguagcagagcagcucccu...................................................................  1     1  7y1
....................uuucguacguagcagCgcagcucccu...................................................................  10    1  7y1
....................Cuucguacguagcagagcagcucccu...................................................................  2     1  7y1
....................uuucguacguagcagaCcagcucccu...................................................................  2     1  7y1
....................uuucguacguagcagagcagcuAccu...................................................................  10    1  7y1
....................uuAcguacguagcagagcagcucccuc..................................................................  3     1  7y1
....................uuucCuacguagcagagcagcucccuc..................................................................  2     1  7y1
....................uuucguacguagcagagAagcucccuc..................................................................  6     1  7y1
....................uuucguacguagcagagcaCcucccuc..................................................................  1     1  7y1
....................Cuucguacguagcagagcagcucccuc..................................................................  1     1  7y1
....................uuucguacguagcagCgcagcucccuc..................................................................  10    1  7y1
....................uuucguacguagcagagcagcuAccuc..................................................................  7     1  7y1
....................uuucguacguagcagagcCgcucccuc..................................................................  1     1  7y1
....................Nuucguacguagcagagcagcucccuc..................................................................  6     1  7y1
....................uuucguacguagcagagcagcCcccuc..................................................................  1     1  7y1
....................uuucguacguaAcagagcagcucccuc..................................................................  1     1  7y1
....................uuucguacguagcagUgcagcucccuc..................................................................  2     1  7y1
....................uuucguacguagcagagcaUcucccuc..................................................................  2     1  7y1
....................uuucguacguagcagagcagcuccAuc..................................................................  5     1  7y1
....................uuucguacguagcaUagcagcucccuc..................................................................  2     1  7y1
....................uuucgCacguagcagagcagcucccuc..................................................................  1     1  7y1
....................uuucguacguagcagagcGgcucccuc..................................................................  2     1  7y1
....................uuucguaUguagcagagcagcucccuc..................................................................  1     1  7y1
....................uuucguacguagcagagcagcuGccuc..................................................................  1     1  7y1
....................uuucguacguagcagagcagcucAcuc..................................................................  6     1  7y1
....................uuucguacguaUcagagcagcucccuc..................................................................  1     1  7y1
....................uuucguacguagcagagcagAucccuc..................................................................  4     1  7y1
....................uuucguacguagcagagcagcucccuc..................................................................  4379  0  7y1
....................uuucUuacguagcagagcagcucccuc..................................................................  2     1  7y1
....................uuucguacguagcagagcUgcucccuc..................................................................  1     1  7y1
....................uuucguacguagAagagcagcucccuc..................................................................  5     1  7y1
....................uAucguacguagcagagcagcucccuc..................................................................  9     1  7y1
....................uuucguacguagUagagcagcucccuc..................................................................  1     1  7y1
....................Guucguacguagcagagcagcucccuc..................................................................  12    1  7y1
....................Auucguacguagcagagcagcucccuc..................................................................  39    1  7y1
....................uuucgAacguagcagagcagcucccuc..................................................................  9     1  7y1
....................uuucguacgAagcagagcagcucccuc..................................................................  7     1  7y1
....................uuuGguacguagcagagcagcucccuc..................................................................  1     1  7y1
....................uuucguaAguagcagagcagcucccuc..................................................................  3     1  7y1
....................uuucguacguagcagagcagcucccuA..................................................................  3     1  7y1
....................uuucguacguagcagaGgagcucccuc..................................................................  1     1  7y1
....................uuucguacguagcagagcagcucUcuc..................................................................  1     1  7y1
....................uuucguacguagcaCagcagcucccuc..................................................................  2     1  7y1
....................uuucguacguagcagagcagcucccAc..................................................................  4     1  7y1
....................uuucguacguagcagagcagcAcccuc..................................................................  2     1  7y1
....................uuuAguacguagcagagcagcucccuc..................................................................  11    1  7y1
....................uuucguacguagcagagcagcuUccuc..................................................................  1     1  7y1
....................uuucguacguagcagagcagcucGcuc..................................................................  2     1  7y1
....................uuucguacCuagcagagcagcucccuc..................................................................  1     1  7y1
....................uuucguacUuagcagagcagcucccuc..................................................................  2     1  7y1
....................uuAcguacguagcagagcagcucccucgc................................................................  1     1  7y1
....................uuucguacguagcagCgcagcucccucgc................................................................  1     1  7y1
....................uuuAguacguagcagagcagcucccucgc................................................................  1     1  7y1
....................uuucguacguagUagagcagcucccucgc................................................................  1     1  7y1
....................uuucguacguagcagagcagcucAcucgc................................................................  1     1  7y1
```

```
 Star                                                                Mature
gaccugcuucugggucggggguuucguacguagcagagcagcucccucgcugcgaucuauugaaagucagcccucgacacaaggguuuguccgcgcgcgcgcgcgcgcgugcgu
....................uuucguacguagcagaUcagcucccucgc................................................................   1     1    7y1
....................uuucguacguagcagagAagcucccucgc................................................................   1     1    7y1
....................uuuGguacguagcagagcagcucccucgc................................................................   1     1    7y1
....................uuucguacguagcagagcagcucccAcgc................................................................   1     1    7y1
....................uuucguacguagcagagcUgcucccucgc................................................................   1     1    7y1
....................Guucguacguagcagagcagcucccucgc................................................................   2     1    7y1
....................uuucguacguagcagagcaCcucccucgc................................................................   1     1    7y1
....................uuucguacguagcagagcagcucccucgc................................................................   614   0    7y1
....................uuucguacgAagcagagcagcucccucgc................................................................   1     1    7y1
....................uAucguacguagcagagcagcucccucgc................................................................   1     1    7y1
....................uuucgAacguagcagagcagcucccucgc................................................................   1     1    7y1
....................uuucguacCuagcagagcagcucccucgc................................................................   1     1    7y1
....................uuucguacguagcagagcagcuAccucgc................................................................   1     1    7y1
....................Auucguacguagcagagcagcucccucgc................................................................   4     1    7y1
....................uuuAguacguagcagagcagcucccucgcu..............................................................   4     1    7y1
....................uuucguacguagGagagcagcucccucgcu..............................................................   1     1    7y1
....................uuucguacguagcagagcagcucccuGgcu..............................................................   1     1    7y1
....................uuucguacguagcagaCcagcucccucgcu..............................................................   1     1    7y1
....................Auucguacguagcagagcagcucccucgcu..............................................................   20    1    7y1
....................uuucguacguagcagagcagcCcccucgcu..............................................................   1     1    7y1
....................uuucguacguagAagagcagcucccucgcu..............................................................   3     1    7y1
....................uuucguacCuagcagagcagcucccucgcu..............................................................   1     1    7y1
....................uuucguacguagcCgagcagcucccucgcu..............................................................   1     1    7y1
....................uuucguacguagcagagcCgcucccucgcu..............................................................   2     1    7y1
....................uuucguacguaCcagagcagcucccucgcu..............................................................   1     1    7y1
....................uuucguacguagcagagcagcucUcucgcu..............................................................   2     1    7y1
....................uuucguUcguagcagagcagcucccucgcu..............................................................   1     1    7y1
....................uuucguacguagcagagAagcucccucgcu..............................................................   2     1    7y1
....................uuucguacguagcagagcGgcucccucgcu..............................................................   1     1    7y1
....................uuucguacguagcagagcaUcucccucgcu..............................................................   1     1    7y1
....................uuucguacguagcagUgcagcucccucgcu..............................................................   1     1    7y1
....................uuucguacguagcagagcUgcucccucgcu..............................................................   1     1    7y1
....................uuucguacguagcagaUcagcucccucgcu..............................................................   1     1    7y1
....................uuucguacguagcagagcagcuAccucgcu..............................................................   7     1    7y1
....................uuucguacguagcagagcagcucccuAgcu..............................................................   3     1    7y1
....................uuucguacguagcagGgcagcucccucgcu..............................................................   2     1    7y1
....................uuucguGcguagcagagcagcucccucgcu..............................................................   1     1    7y1
....................uuucgAacguagcagagcagcucccucgcu..............................................................   4     1    7y1
....................uuucguacguagcagagcagcucccucgcu..............................................................   2277  0    7y1
....................uuucguaAguagcagagcagcucccucgcu..............................................................   2     1    7y1
....................Guucguacguagcagagcagcucccucgcu..............................................................   178   1    7y1
....................uAucguacguagcagagcagcucccucgcu..............................................................   1     1    7y1
....................uuucguacguagcagagcagcucGcucgcu..............................................................   1     1    7y1
....................uuGcguacguagcagagcagcucccucgcu..............................................................   1     1    7y1
....................uuAcguacguagcagagcagcucccucgcu..............................................................   1     1    7y1
....................uuucguacguagcagCgcagcucccucgcu..............................................................   2     1    7y1
....................uuucguacgCagcagagcagcucccucgcu..............................................................   1     1    7y1
....................uuucguacgAagcagagcagcucccucgcu..............................................................   6     1    7y1
....................uuucguacguagcaCagcagcucccucgcu..............................................................   2     1    7y1
....................uuucguacUuagcagagcagcucccucgcu..............................................................   1     1    7y1
....................uuucguacguagcagagcagUucccucgcu..............................................................   1     1    7y1
....................uuucguacguagcagagcagcucccucAcu..............................................................   2     1    7y1
....................uuucAuacguagcagagcagcucccucgcu..............................................................   2     1    7y1
....................uuucguacguagcagagcagcucccucgAu..............................................................   3     1    7y1
....................uuucguacguagcagagcagcucccucgcG..............................................................   20    1    7y1
....................Nuucguacguagcagagcagcucccucgcu..............................................................   2     1    7y1
....................uuucguacguagcagagcagcucccAcgcu..............................................................   2     1    7y1
....................uuucguacguagcagagcagcucAcucgcu..............................................................   1     1    7y1
....................uuucguacCuagcagagcagcucccucgcug............................................................   1     1    7y1
....................uuucAuacguagcagagcagcucccucgcug............................................................   1     1    7y1
....................uuucguacguagcUgagcagcucccucgcug............................................................   1     1    7y1
....................uuucguacguagcagagcGgcucccucgcug............................................................   1     1    7y1
....................uuucguacguagcagagcagcucccucCcug............................................................   1     1    7y1
....................uuucguacguagcagagcagcucccucgcGg............................................................   9     1    7y1
....................Auucguacguagcagagcagcucccucgcug............................................................   6     1    7y1
....................uuucguacguagcagagcagcucccAcgcug............................................................   1     1    7y1
....................uuucguUcguagcagagcagcucccucgcug............................................................   1     1    7y1
....................uuucguacguagcagagcagcucccucgcug............................................................   855   0    7y1
....................uuucguacguagcagagcagcucccucUcug............................................................   1     1    7y1
....................uuucguacgAagcagagcagcucccucgcug............................................................   1     1    7y1
```

Star                                                                     Mature

```
gaccugcuucugggucggggguuucguacguagcagagcagcucccucgcugcgaucuauugaaagucagcccucgacacaaggguuuguccgcgcgcgcgcgcgcgcgugcgu
```

```
....................uuucguacguagcagagcaAcucccucgcug..............................................................   1      1   7y1
....................uuucUuacguagcagagcagcucccucgcug..............................................................   1      1   7y1
....................uuucguacguagcagagcagcuAccucgcug..............................................................   2      1   7y1
....................uuucguacguagcagagAagcucccucgcug..............................................................   2      1   7y1
....................uAucguacguagcagagcagcucccucgcug..............................................................   2      1   7y1
....................uuucguCcguagcagagcagcucccucgcug..............................................................   1      1   7y1
....................uuucguacguagcagagcagcucccucgGug..............................................................   1      1   7y1
....................uuucgAacguagcagagcagcucccucgcug..............................................................   2      1   7y1
....................uuucguacguagAagagcagcucccucgcug..............................................................   1      1   7y1
....................uuucguaAguagcagagcagcucccucgcug..............................................................   1      1   7y1
....................uuucguacguagcagagcagAucccucgcugc.............................................................   1      1   7y1
....................uAucguacguagcagagcagcucccucgcugc.............................................................   1      1   7y1
....................uuucguacguagcagagcagcAcccucgcugc.............................................................   1      1   7y1
....................uuucguacCuagcagagcagcucccucgcugc.............................................................   1      1   7y1
....................Guucguacguagcagagcagcucccucgcugc.............................................................   1      1   7y1
....................uuucguacguagcagagcagcucccucgcugc.............................................................   499    0   7y1
....................uuucguacguagcagagcagcucccucgcuCc.............................................................   1      1   7y1
....................uuucguacgAagcagagcagcucccucgcugc.............................................................   1      1   7y1
....................Auucguacguagcagagcagcucccucgcugc.............................................................   5      1   7y1
....................uuuAguacguagcagagcagcucccucgcugc.............................................................   1      1   7y1
....................uuucguacguagcagagcagcuAccucgcugc.............................................................   1      1   7y1
....................Nuucguacguagcagagcagcucccucgcugc.............................................................   2      1   7y1
....................uuucguacguagcagagcagcucccucgAugc.............................................................   1      1   7y1
.....................uucguacguagcaUagca..........................................................................   1      1   7y1
.....................uucguacguagcaCagca..........................................................................   1      1   7y1
.....................uucguacguagcagagAa..........................................................................   3      1   7y1
.....................uucguacguaCcagagca..........................................................................   1      1   7y1
.....................uucguacguaAcagagca..........................................................................   1      1   7y1
.....................uucguacguagcagGgca..........................................................................   1      1   7y1
.....................uucguacguagcagagGa..........................................................................   1      1   7y1
.....................uucguacguagAagagca..........................................................................   1      1   7y1
.....................uucguacgAagcagagca..........................................................................   3      1   7y1
.....................uucAuacguagcagagca..........................................................................   1      1   7y1
.....................uAcguacguagcagagca..........................................................................   4      1   7y1
.....................Nucguacguagcagagca..........................................................................   1      1   7y1
.....................uucguaAguagcagagca..........................................................................   8      1   7y1
.....................Gucguacguagcagagca..........................................................................   2      1   7y1
.....................uuAguacguagcagagca..........................................................................   4      1   7y1
.....................uucguacguagcagaAca..........................................................................   1      1   7y1
.....................uucguacguagcagagcG..........................................................................   1      1   7y1
.....................uNcguacguagcagagca..........................................................................   1      1   7y1
.....................Aucguacguagcagagca..........................................................................   12     1   7y1
.....................uucguacCuagcagagca..........................................................................   2      1   7y1
.....................uucguacguagcagagca..........................................................................   1212   0   7y1
.....................Cucguacguagcagagca..........................................................................   1      1   7y1
.....................uucguacguagcagagcC..........................................................................   1      1   7y1
.....................uucguacguagcagaUca..........................................................................   2      1   7y1
.....................uucguacAuagcagagcag.........................................................................   1      1   7y1
.....................uucguacguagcaCagcag.........................................................................   1      1   7y1
.....................Aucguacguagcagagcag.........................................................................   16     1   7y1
.....................uAcguacguagcagagcag.........................................................................   3      1   7y1
.....................Gucguacguagcagagcag.........................................................................   2      1   7y1
.....................uucguacguagcagagcag.........................................................................   1378   0   7y1
.....................uucguacguagAagagcag.........................................................................   1      1   7y1
.....................uucguaAguagcagagcag.........................................................................   4      1   7y1
.....................uucgCacguagcagagcag.........................................................................   1      1   7y1
.....................uucguacgAagcagagcag.........................................................................   2      1   7y1
.....................uucguacguagcagagAag.........................................................................   1      1   7y1
.....................uucguacguGgcagagcag.........................................................................   1      1   7y1
.....................uucguacguagcagagcaU.........................................................................   2      1   7y1
.....................uucguacCuagcagagcag.........................................................................   1      1   7y1
.....................uucguacguagcagagcUg.........................................................................   1      1   7y1
.....................uucguacguaUcagagcag.........................................................................   1      1   7y1
.....................uuAguacguagcagagcag.........................................................................   4      1   7y1
.....................uucguUcguagcagagcagc........................................................................   2      1   7y1
.....................uucguacgAagcagagcagc........................................................................   1      1   7y1
.....................uucguacAuagcagagcagc........................................................................   1      1   7y1
.....................Nucguacguagcagagcagc........................................................................   1      1   7y1
.....................uucguacUuagcagagcagc........................................................................   2      1   7y1
.....................uucguaGguagcagagcagc........................................................................   2      1   7y1
```

```
  Star                                                                    Mature
gaccugcuucugggucgggguuucguacguagcagagcagcucccucgcugcgaucuauugaaagucagcccucgacacaaggguuuguccgcgcgcgcgcgcgcgcgcgugcgu
.....................uAcguacguagcagagcagc..........................................................................   2      1   7y1
.....................uucguacguagcaCagcagc..........................................................................   1      1   7y1
.....................uucCuacguagcagagcagc..........................................................................   1      1   7y1
.....................uucguacCuagcagagcagc..........................................................................   1      1   7y1
.....................uucguacguagcagagcaUc..........................................................................   3      1   7y1
.....................uucguacguagcagaUcagc..........................................................................   1      1   7y1
.....................uucguacguagcagaCcagc..........................................................................   1      1   7y1
.....................Aucguacguagcagagcagc..........................................................................   33     1   7y1
.....................uucguacguGgcagagcagc..........................................................................   2      1   7y1
.....................uucguacguagcagagcagA..........................................................................   2      1   7y1
.....................uuUguacguagcagagcagc..........................................................................   1      1   7y1
.....................uucguacguagcaUagcagc..........................................................................   1      1   7y1
.....................uucgAacguagcagagcagc..........................................................................   1      1   7y1
.....................uucguacguagcUgagcagc..........................................................................   1      1   7y1
.....................uucguaAguagcagagcagc..........................................................................   2      1   7y1
.....................uucguacguaAcagagcagc..........................................................................   1      1   7y1
.....................uCcguacguagcagagcagc..........................................................................   1      1   7y1
.....................uucguacgCagcagagcagc..........................................................................   1      1   7y1
.....................uucguacguagcaAagcagc..........................................................................   1      1   7y1
.....................uucguacguagcagagcagc..........................................................................   1790   0   7y1
.....................uucguacguagcagGgcagc..........................................................................   1      1   7y1
.....................uuAguacguagcagagcagc..........................................................................   5      1   7y1
.....................uucguacguagcagagcGgcu.........................................................................   5      1   7y1
.....................uucguacguagcagagAagcu.........................................................................   9      1   7y1
.....................uucguacguagUagagcagcu.........................................................................   185    1   7y1
.....................uucguacguagcagagcagcA.........................................................................   4      1   7y1
.....................uucguacguagcagagcagAu.........................................................................   4      1   7y1
.....................uucguacguagcGgagcagcu.........................................................................   1      1   7y1
.....................Nucguacguagcagagcagcu.........................................................................   2      1   7y1
.....................uucguGcguagcagagcagcu.........................................................................   1      1   7y1
.....................uucguacguaUcagagcagcu.........................................................................   3      1   7y1
.....................uucguacguagcUgagcagcu.........................................................................   2      1   7y1
.....................uucguacguagcagagcUgcu.........................................................................   1      1   7y1
.....................uucCuacguagcagagcagcu.........................................................................   1      1   7y1
.....................uucguacgAagcagagcagcu.........................................................................   8      1   7y1
.....................uucUuacguagcagagcagcu.........................................................................   5      1   7y1
.....................uucguacguagcagagcaUcu.........................................................................   3      1   7y1
.....................uucguacguagcagagcaAcu.........................................................................   1      1   7y1
.....................uucguacguagcagaCcagcu.........................................................................   5      1   7y1
.....................uuGguacguagcagagcagcu.........................................................................   2      1   7y1
.....................uucguacguagcagagcagGu.........................................................................   2      1   7y1
.....................uucguacguagcagGgcagcu.........................................................................   1      1   7y1
.....................uucguacguagcagagcaCcu.........................................................................   3      1   7y1
.....................uucgGacguagcagagcagcu.........................................................................   1      1   7y1
.....................uucguacguagcagagcagcu.........................................................................   7198   0   7y1
.....................Aucguacguagcagagcagcu.........................................................................   80     1   7y1
.....................uucguacguagcagagcagcG.........................................................................   40     1   7y1
.....................uucgCacguagcagagcagcu.........................................................................   3      1   7y1
.....................uucguacguagAagagcagcu.........................................................................   3      1   7y1
.....................uucguacguagcagCgcagcu.........................................................................   7      1   7y1
.....................uucguacguGgcagagcagcu.........................................................................   2      1   7y1
.....................uucgAacguagcagagcagcu.........................................................................   9      1   7y1
.....................uucguacguagcagagGagcu.........................................................................   4      1   7y1
.....................uucguacguagcagagcagcC.........................................................................   1      1   7y1
.....................uucguacguUgcagagcagcu.........................................................................   1      1   7y1
.....................uucguaUguagcagagcagcu.........................................................................   3      1   7y1
.....................Gucguacguagcagagcagcu.........................................................................   8      1   7y1
.....................uucguacCuagcagagcagcu.........................................................................   4      1   7y1
.....................uucguacguagcaAagcagcu.........................................................................   1      1   7y1
.....................uucguUcguagcagagcagcu.........................................................................   1      1   7y1
.....................uuAguacguagcagagcagcu.........................................................................   24     1   7y1
.....................uucguacguaCcagagcagcu.........................................................................   3      1   7y1
.....................uAcguacguagcagagcagcu.........................................................................   18     1   7y1
.....................uucguacguagcagagUagcu.........................................................................   1      1   7y1
.....................uucguacguagcaUagcagcu.........................................................................   5      1   7y1
.....................uucguaGguagcagagcagcu.........................................................................   1      1   7y1
.....................uucguacguagcaCagcagcu.........................................................................   1      1   7y1
.....................uucguacAuagcagagcagcu.........................................................................   1      1   7y1
.....................uucguaAguagcagagcagcu.........................................................................   22     1   7y1
.....................uucguacguagcagagcCgcu.........................................................................   1      1   7y1
```

Star                                                          Mature

```
gaccugcuucugggucgggguuucguacguagcagagcagcucccucgcugcgaucuauugaaagucagcccucgacacaaggguuuguccgcgcgcgcgcgcgcgugcgu
.....................uucguacguagcagagcagcucA...................................................................	13	1	7y1
.....................uucguacguagcUgagcagcucc...................................................................	2	1	7y1
.....................uucguacgGagcagagcagcucc...................................................................	1	1	7y1
.....................uucguacguagcagagcagcuAc...................................................................	9	1	7y1
.....................uucguacguagcagagcagcGcc...................................................................	1	1	7y1
.....................uucguacguaCcagagcagcucc...................................................................	3	1	7y1
.....................uuUguacguagcagagcagcucc...................................................................	1	1	7y1
.....................uucguacguagcagagcCgcucc...................................................................	1	1	7y1
.....................uucguacguagcagagGagcucc...................................................................	2	1	7y1
.....................uucguacguagcagagUagcucc...................................................................	1	1	7y1
.....................Nucguacguagcagagcagcucc...................................................................	4	1	7y1
.....................uucguacCuagcagagcagcucc...................................................................	6	1	7y1
.....................uucguaGguagcagagcagcucc...................................................................	1	1	7y1
.....................uNcguacguagcagagcagcucc...................................................................	1	1	7y1
.....................uucgAacguagcagagcagcucc...................................................................	1	1	7y1
.....................uAcguacguagcagagcagcucc...................................................................	17	1	7y1
.....................uuAguacguagcagagcagcucc...................................................................	16	1	7y1
.....................uucguUcguagcagagcagcucc...................................................................	1	1	7y1
.....................uucguacguagcagagAagcucc...................................................................	8	1	7y1
.....................uuGguacguagcagagcagcucc...................................................................	2	1	7y1
.....................uucguacgAagcagagcagcucc...................................................................	6	1	7y1
.....................Aucguacguagcagagcagcucc...................................................................	71	1	7y1
.....................uucguacUuagcagagcagcucc...................................................................	2	1	7y1
.....................uucguacguagcagagcagcucc...................................................................	7274	0	7y1
.....................uucguaAguagcagagcagcucc...................................................................	26	1	7y1
.....................uucguacguagAagagcagcucc...................................................................	11	1	7y1
.....................uucguacguaUcagagcagcucc...................................................................	8	1	7y1
.....................uucguacguagcagagcagcAcc...................................................................	1	1	7y1
.....................uucguacguGgcagagcagcucc...................................................................	2	1	7y1
.....................uucguCcguagcagagcagcucc...................................................................	1	1	7y1
.....................uucguacguagcagagcaAcucc...................................................................	1	1	7y1
.....................uucguacguagcagagcagcucG...................................................................	1	1	7y1
.....................uucgCacguagcagagcagcucc...................................................................	3	1	7y1
.....................uucguacguagcagagcaUcucc...................................................................	3	1	7y1
.....................uucguacguagcaCagcagcucc...................................................................	1	1	7y1
.....................uucguacguagGagagcagcucc...................................................................	1	1	7y1
.....................uucguacguagcagaUcagcucc...................................................................	5	1	7y1
.....................uucguacguagUagagcagcucc...................................................................	1	1	7y1
.....................uucguacguagcagagcaCcucc...................................................................	1	1	7y1
.....................uucguacguGgcagagcagcuccc..................................................................	1	1	7y1
.....................uucguacguaCcagagcagcuccc..................................................................	2	1	7y1
.....................uucguacguagcagagcCgcuccc..................................................................	2	1	7y1
.....................uucguacUuagcagagcagcuccc..................................................................	1	1	7y1
.....................uucguacguaUcagagcagcuccc..................................................................	1	1	7y1
.....................uucguacguagcagagcagcuAcc..................................................................	7	1	7y1
.....................uucguacguagcagagcagcuccA..................................................................	2	1	7y1
.....................uAcguacguagcagagcagcuccc..................................................................	6	1	7y1
.....................Nucguacguagcagagcagcuccc..................................................................	1	1	7y1
.....................uucguacguagcagagcagcuccc..................................................................	3826	0	7y1
.....................uucguacCuagcagagcagcuccc..................................................................	1	1	7y1
.....................Aucguacguagcagagcagcuccc..................................................................	31	1	7y1
.....................uucguacguagcagagcagcuUcc..................................................................	1	1	7y1
.....................uucguacguagcagagcNgcuccc..................................................................	1	1	7y1
.....................uucguCcguagcagagcagcuccc..................................................................	1	1	7y1
.....................uucguacguagcagagcagcCccc..................................................................	1	1	7y1
.....................uuAguacguagcagagcagcuccc..................................................................	10	1	7y1
.....................uucguacguagcagagcagcucAc..................................................................	16	1	7y1
.....................uucguacguagcGgagcagcuccc..................................................................	1	1	7y1
.....................uucguacguagcagagcagcuGcc..................................................................	1	1	7y1
.....................uucAuacguagcagagcagcuccc..................................................................	1	1	7y1
.....................uucguacgAagcagagcagcuccc..................................................................	2	1	7y1
.....................uucUuacguagcagagcagcuccc..................................................................	1	1	7y1
.....................uucguacguagcagagAagcuccc..................................................................	4	1	7y1
.....................uucgAacguagcagagcagcuccc..................................................................	4	1	7y1
.....................Gucguacguagcagagcagcuccc..................................................................	13	1	7y1
.....................uucguaAguagcagagcagcuccc..................................................................	11	1	7y1
.....................uucguacguagAagagcagcuccc..................................................................	1	1	7y1
.....................uucguacguagcaCagcagcuccc..................................................................	1	1	7y1
.....................uucguacguagcagaCcagcuccc..................................................................	1	1	7y1
.....................uucguacguagcagagcaCcuccc..................................................................	4	1	7y1
```

**Star** **Mature**

```
gaccugcuucugggucggggguuucguacguagcagagcagcucccucgcugcgaucuauugaaagucagcccucgacacaaggguuuguccgcgcgcgcgcgcgcgcgcgugcgu
.....................uucguacguagcaUagcagcuccc......................................................................   6    1    7y1
.....................uucguacguagcagagUagcuccc......................................................................   1    1    7y1
.....................uucguacguagcagagcaUcuccc......................................................................   5    1    7y1
.....................uucguacguagcagagcaAcuccc......................................................................   1    1    7y1
.....................uucguacguagcagaUcagcuccc......................................................................   3    1    7y1
.....................uucguacguagcagagcagcAccc......................................................................   2    1    7y1
.....................uucguacguagcagagcagcucGc......................................................................   1    1    7y1
.....................uucCuacguagcagagcagcuccc......................................................................   1    1    7y1
.....................uucguacguagcagagcagcuUccu.....................................................................   1    1    7y1
.....................uucUuacguagcagagcagcucccu.....................................................................   4    1    7y1
.....................uucguacguagcagCgcagcucccu.....................................................................   4    1    7y1
.....................uucguacguagcagagcagcGcccu.....................................................................   2    1    7y1
.....................uucguacgCagcagagcagcucccu.....................................................................   2    1    7y1
.....................uucguacguagcUgagcagcucccu.....................................................................   1    1    7y1
.....................uucguacguagcagagcUgcucccu.....................................................................   6    1    7y1
.....................uucguacguagcagagcagcucUcu.....................................................................   5    1    7y1
.....................Gucguacguagcagagcagcucccu.....................................................................   24   1    7y1
.....................uucguacguaAcagagcagcucccu.....................................................................   1    1    7y1
.....................uucguacguagcagGgcagcucccu.....................................................................   1    1    7y1
.....................Cucguacguagcagagcagcucccu.....................................................................   1    1    7y1
.....................uucguacguagcagagcagcAcccu.....................................................................   6    1    7y1
.....................uucguacguagcagagcagcuAccu.....................................................................   23   1    7y1
.....................uucguacguagcagagAagcucccu.....................................................................   11   1    7y1
.....................uucguacguagcagagcagcucAcu.....................................................................   24   1    7y1
.....................uucguacguagcGgagcagcucccu.....................................................................   2    1    7y1
.....................uucguacguagcagagcagcucccA.....................................................................   13   1    7y1
.....................uNcguacguagcagagcagcucccu.....................................................................   1    1    7y1
.....................uucguacguagcagagcCgcucccu.....................................................................   2    1    7y1
.....................uucguacguagcagaCcagcucccu.....................................................................   3    1    7y1
.....................uucguUcguagcagagcagcucccu.....................................................................   1    1    7y1
.....................Nucguacguagcagagcagcucccu.....................................................................   8    1    7y1
.....................uucguacgGagcagagcagcucccu.....................................................................   1    1    7y1
.....................uucguacguagAagagcagcucccu.....................................................................   7    1    7y1
.....................uucguacguagcagagcagcuccGu.....................................................................   2    1    7y1
.....................uucguacgAagcagagcagcucccu.....................................................................   9    1    7y1
.....................uuGguacguagcagagcagcucccu.....................................................................   2    1    7y1
.....................uucCuacguagcagagcagcucccu.....................................................................   1    1    7y1
.....................uucguaUguagcagagcagcucccu.....................................................................   1    1    7y1
.....................uucguacguagcagUgcagcucccu.....................................................................   1    1    7y1
.....................uucgCacguagcagagcagcucccu.....................................................................   2    1    7y1
.....................uucguacguagcCgagcagcucccu.....................................................................   2    1    7y1
.....................uucguacguagcagagcagAucccu.....................................................................   5    1    7y1
.....................uucguacguagcagagcaAcucccu.....................................................................   1    1    7y1
.....................uucguacguagcaUagcagcucccu.....................................................................   5    1    7y1
.....................uucguacguagUagagcagcucccu.....................................................................   1    1    7y1
.....................Aucguacguagcagagcagcucccu.....................................................................   78   1    7y1
.....................uucguacguagcaCagcagcucccu.....................................................................   3    1    7y1
.....................uucguacguagcagagcaCcucccu.....................................................................   1    1    7y1
.....................uucguacguagcagagGagcucccu.....................................................................   2    1    7y1
.....................uucguacguagcagagUagcucccu.....................................................................   1    1    7y1
.....................uucguacguGgcagagcagcucccu.....................................................................   1    1    7y1
.....................uucgAacguagcagagcagcucccu.....................................................................   5    1    7y1
.....................uucguaGguagcagagcagcucccu.....................................................................   1    1    7y1
.....................uucguacguagcagagcaUcucccu.....................................................................   5    1    7y1
.....................uucguacguagcagagcagcCcccu.....................................................................   2    1    7y1
.....................uucguacguagcagagcagcuGccu.....................................................................   1    1    7y1
.....................uucguacguagcagagcagcucccG.....................................................................   35   1    7y1
.....................uucguacguaUcagagcagcucccu.....................................................................   5    1    7y1
.....................uucguacguaCcagagcagcucccu.....................................................................   3    1    7y1
.....................uucguacguagcagagcGgcucccu.....................................................................   8    1    7y1
.....................uucguacguagcagagcagcucccC.....................................................................   1    1    7y1
.....................uucguacUuagcagagcagcucccu.....................................................................   4    1    7y1
.....................uucguacguagcagagcagcuccAu.....................................................................   6    1    7y1
.....................uucguacCuagcagagcagcucccu.....................................................................   5    1    7y1
.....................uucguacguagGagagcagcucccu.....................................................................   1    1    7y1
.....................uucguGcguagcagagcagcucccu.....................................................................   1    1    7y1
.....................uucguaAguagcagagcagcucccu.....................................................................   40   1    7y1
.....................uucguacAuagcagagcagcucccu.....................................................................   3    1    7y1
.....................uuAguacguagcagagcagcucccu.....................................................................   23   1    7y1
.....................uAcguacguagcagagcagcucccu.....................................................................   21   1    7y1
```

**Star** **Mature**

```
gaccugcuucugggucgggguuucguacguagcagagcagcucccucgcugcgaucuauugaaagucagcccucgacacaaggguuuguccgcgcgcgcgcgcgcgcgugcgu
.....................uucguacguagcagaUcagcucccu...................................................................  2     1  7y1
.....................uucguacguagcaAagcagcucccu...................................................................  2     1  7y1
.....................uCcguacguagcagagcagcucccu...................................................................  2     1  7y1
.....................uucguacguagcagagcagcucccu...................................................................  9826  0  7y1
.....................uucAuacguagcagagcagcucccu...................................................................  1     1  7y1
.....................uucguacguagcaUagcagcucccuc..................................................................  4     1  7y1
.....................uucguacguagcagagcagcuAccuc..................................................................  12    1  7y1
.....................uucguacguagcGgagcagcucccuc..................................................................  1     1  7y1
.....................uucguacguagcagagcaUcucccuc..................................................................  1     1  7y1
.....................uucCuacguagcagagcagcucccuc..................................................................  2     1  7y1
.....................uucguacguagcagagcagcuccAuc..................................................................  6     1  7y1
.....................uucguacguaUcagagcagcucccuc..................................................................  3     1  7y1
.....................uucguacguagcagGgcagcucccuc..................................................................  1     1  7y1
.....................uucguacguaCcagagcagcucccuc..................................................................  1     1  7y1
.....................uucguacAuagcagagcagcucccuc..................................................................  1     1  7y1
.....................uucguaAguagcagagcagcucccuc..................................................................  33    1  7y1
.....................uucguacguaAcagagcagcucccuc..................................................................  1     1  7y1
.....................uucguacgCagcagagcagcucccuc..................................................................  1     1  7y1
.....................Nucguacguagcagagcagcucccuc..................................................................  5     1  7y1
.....................uucguacguagcagagcagcGcccuc..................................................................  1     1  7y1
.....................uucguacguagcagagcaCcucccuc..................................................................  1     1  7y1
.....................uucguacguagcagagcagcucccuA..................................................................  1     1  7y1
.....................uucguacguagcagagcagcucccAc..................................................................  1     1  7y1
.....................uucguacguagcagagUagcucccuc..................................................................  1     1  7y1
.....................uucguacguagcagagcagcucccuc..................................................................  7376  0  7y1
.....................uucguacguagcagaUcagcucccuc..................................................................  4     1  7y1
.....................uucguacUuagcagagcagcucccuc..................................................................  2     1  7y1
.....................uucguacgAagcagagcagcucccuc..................................................................  6     1  7y1
.....................Gucguacguagcagagcagcucccuc..................................................................  10    1  7y1
.....................uucguacguagcagagcagcuUccuc..................................................................  1     1  7y1
.....................uucguacguagcagagcagcucccCc..................................................................  2     1  7y1
.....................uucguacguagcagUgcagcucccuc..................................................................  2     1  7y1
.....................uucguacguagcagagcUgcucccuc..................................................................  1     1  7y1
.....................uucguacguagcagagcCgcucccuc..................................................................  3     1  7y1
.....................uucguacguagcagagcagcAcccuc..................................................................  5     1  7y1
.....................uucguacguagcagagcaAcucccuc..................................................................  2     1  7y1
.....................uucguGcguagcagagcagcucccuc..................................................................  1     1  7y1
.....................uucguUcguagcagagcagcucccuc..................................................................  1     1  7y1
.....................uucguacCuagcagagcagcucccuc..................................................................  4     1  7y1
.....................uucguacguagcagagcagcucccuG..................................................................  1     1  7y1
.....................uucgAacguagcagagcagcucccuc..................................................................  4     1  7y1
.....................uucguacguagcUgagcagcucccuc..................................................................  2     1  7y1
.....................Cucguacguagcagagcagcucccuc..................................................................  1     1  7y1
.....................uucguacguagcagCgcagcucccuc..................................................................  8     1  7y1
.....................uucguacguagAagagcagcucccuc..................................................................  5     1  7y1
.....................Aucguacguagcagagcagcucccuc..................................................................  59    1  7y1
.....................uuGguacguagcagagcagcucccuc..................................................................  1     1  7y1
.....................uAcguacguagcagagcagcucccuc..................................................................  11    1  7y1
.....................uucguacguagcagagAagcucccuc..................................................................  11    1  7y1
.....................uuUguacguagcagagcagcucccuc..................................................................  1     1  7y1
.....................uucguacguagcagagcagAucccuc..................................................................  5     1  7y1
.....................uuAguacguagcagagcagcucccuc..................................................................  12    1  7y1
.....................uucUuacguagcagagcagcucccuc..................................................................  2     1  7y1
.....................uCcguacguagcagagcagcucccuc..................................................................  1     1  7y1
.....................uucgCacguagcagagcagcucccuc..................................................................  253   1  7y1
.....................uucguacguagcaAagcagcucccuc..................................................................  1     1  7y1
.....................uucguacguagcagaCcagcucccuc..................................................................  2     1  7y1
.....................uucguacguagcagagcagcucccGc..................................................................  1     1  7y1
.....................uucgGacguagcagagcagcucccuc..................................................................  1     1  7y1
.....................uucguacguagcagagcGgcucccuc..................................................................  2     1  7y1
.....................uucguacguagcagagcagcuGccuc..................................................................  4     1  7y1
.....................uucguacguagcagagcagcucAcuc..................................................................  19    1  7y1
.....................uucCuacguagcagagcagcucccucg.................................................................  1     1  7y1
.....................uucguCcguagcagagcagcucccucg.................................................................  1     1  7y1
.....................uAcguacguagcagagcagcucccucg.................................................................  2     1  7y1
.....................uucguacguagcagagcagcucccucC.................................................................  1     1  7y1
.....................uucguacguagcagGgcagcucccucg.................................................................  2     1  7y1
.....................uucguacguagcagagcagcucccucA.................................................................  1     1  7y1
.....................uucguacguagcagaUcagcucccucg.................................................................  3     1  7y1
.....................Nucguacguagcagagcagcucccucg.................................................................  1     1  7y1
```

**Star** **Mature**

```
gaccugcuucugggucggggguuucguacguagcagagcagcucccucgcugcgaucuauugaaagucagcccucgacacaaggguuuguccgcgcgcgcgcgcgcgcgugcgu
.....................uucguacguagcagagcagcucAcucg.................................................................  4     1   7y1
.....................uucguacguagcagagcaCcucccucg.................................................................  1     1   7y1
.....................uucguacCuagcagagcagcucccucg.................................................................  3     1   7y1
.....................uucgAacguagcagagcagcucccucg.................................................................  2     1   7y1
.....................uucguacguaCcagagcagcucccucg.................................................................  1     1   7y1
.....................uucguacguagcagagcagcucccGcg.................................................................  1     1   7y1
.....................uuAguacguagcagagcagcucccucg.................................................................  3     1   7y1
.....................Gucguacguagcagagcagcucccucg.................................................................  1     1   7y1
.....................uucguaAguagcagagcagcucccucg.................................................................  5     1   7y1
.....................uucguacguagcagCgcagcucccucg.................................................................  2     1   7y1
.....................uucguGcguagcagagcagcucccucg.................................................................  1     1   7y1
.....................uucguacgCagcagagcagcucccucg.................................................................  1     1   7y1
.....................uucguacguagcagagcagcuAccucg.................................................................  1     1   7y1
.....................uucguacguagcagagcagcuccAucg.................................................................  1     1   7y1
.....................uucguacguagcagagcagcucccucg.................................................................  1431  0   7y1
.....................Aucguacguagcagagcagcucccucg.................................................................  13    1   7y1
.....................uucguacguagcagagcagcucccAcg.................................................................  1     1   7y1
.....................uucgCacguagcagagcagcucccucg.................................................................  1     1   7y1
.....................uucguacguagcagagAagcucccucg.................................................................  1     1   7y1
.....................uucguacguagcagagUagcucccucg.................................................................  1     1   7y1
.....................uucguacguagcagagcagAucccucgc................................................................  1     1   7y1
.....................uuAguacguagcagagcagcucccucgc................................................................  2     1   7y1
.....................uucguacguagcagagcaAcucccucgc................................................................  1     1   7y1
.....................uucguacguagcagagcaCcucccucgc................................................................  1     1   7y1
.....................uucguaAguagcagagcagcucccucgc................................................................  3     1   7y1
.....................uucguacguagcagagcagcucccucgA................................................................  1     1   7y1
.....................Cucguacguagcagagcagcucccucgc................................................................  1     1   7y1
.....................uucguacguagcagagcagcucccucgc................................................................  923   0   7y1
.....................uucgCacguagcagagcagcucccucgc................................................................  1     1   7y1
.....................uucguacguagcagagcagGucccucgc................................................................  1     1   7y1
.....................uAcguacguagcagagcagcucccucgc................................................................  1     1   7y1
.....................uucguacguagcagagcagcuGccucgc................................................................  1     1   7y1
.....................uucguacguaCcagagcagcucccucgc................................................................  1     1   7y1
.....................uucguacguagcagagcagcuAccucgc................................................................  2     1   7y1
.....................Aucguacguagcagagcagcucccucgc................................................................  15    1   7y1
.....................uucguacgAagcagagcagcucccucgc................................................................  1     1   7y1
.....................Gucguacguagcagagcagcucccucgc................................................................  1     1   7y1
.....................uucguacguagcagagcGgcucccucgc................................................................  1     1   7y1
.....................uucguacUuagcagagcagcucccucgc................................................................  1     1   7y1
.....................uucguacguagcagagcagcucAcucgc................................................................  1     1   7y1
.....................Nucguacguagcagagcagcucccucgc................................................................  1     1   7y1
.....................uucguacguagcagagcagcucccuAgc................................................................  1     1   7y1
.....................uucguacguagcagagcagcucccucgcA...............................................................  2     1   7y1
.....................uucguacUuagcagagcagcucccucgcu...............................................................  3     1   7y1
.....................uucguacguagcagagcagcCcccucgcu...............................................................  1     1   7y1
.....................uCcguacguagcagagcagcucccucgcu...............................................................  2     1   7y1
.....................uucguacguagcagagAagcucccucgcu...............................................................  3     1   7y1
.....................uucguacguagcagagcUgcucccucgcu...............................................................  2     1   7y1
.....................uucguacguagcagagcagcucAcucgcu...............................................................  9     1   7y1
.....................uucguacguagcagagcaUcucccucgcu...............................................................  2     1   7y1
.....................uucguaAguagcagagcagcucccucgcu...............................................................  13    1   7y1
.....................uucguacguagcagagcGgcucccucgcu...............................................................  2     1   7y1
.....................uucguacguagcagagcagcucGcucgcu...............................................................  1     1   7y1
.....................uucguacguagAagagcagcucccucgcu...............................................................  6     1   7y1
.....................uucguacCuagcagagcagcucccucgcu...............................................................  1     1   7y1
.....................uucguacguagcagagcagcucccucgGu...............................................................  1     1   7y1
.....................uucAuacguagcagagcagcucccucgcu...............................................................  1     1   7y1
.....................uucguGcguagcagagcagcucccucgcu...............................................................  1     1   7y1
.....................uucCuacguagcagagcagcucccucgcu...............................................................  4     1   7y1
.....................uucguacguagcagagcagcucccucgcu...............................................................  4044  0   7y1
.....................Gucguacguagcagagcagcucccucgcu...............................................................  6     1   7y1
.....................uucguacguagcagagcagcuUccucgcu...............................................................  1     1   7y1
.....................uucguacguagcagagcagAucccucgcu...............................................................  1     1   7y1
.....................uucguacguagcagagcagcucccAcgcu...............................................................  2     1   7y1
.....................uGcguacguagcagagcagcucccucgcu...............................................................  1     1   7y1
.....................uucguacguagcGgagcagcucccucgcu...............................................................  2     1   7y1
.....................uucguacguagcagagcagcucccucUcu...............................................................  1     1   7y1
.....................uucguacguagcagagcagcucccucgAu...............................................................  3     1   7y1
.....................uucguacguagcaUagcagcucccucgcu...............................................................  4     1   7y1
.....................uucguacAuagcagagcagcucccucgcu...............................................................  2     1   7y1
```

Star                                    Mature

```
gaccugcuucugggucgggguuucguacguagcagagcagcucccucgcugcgaucuauugaaagucagcccucgacacaaggguuuguccgcgcgcgcgcgcgcgcgugcgu
......................Ncguacguagcagagcagcucc.....................................................................  1     1     7y1
......................ucguacguagcagaUcagcucc.....................................................................  1     1     7y1
......................ucguacguagcagagcagcucA.....................................................................  6     1     7y1
......................ucguacguagcagagcagcucU.....................................................................  1     1     7y1
......................ucUuacguagcagagcagcucc.....................................................................  2     1     7y1
......................ucguacguagcagagAagcucc.....................................................................  2     1     7y1
......................ucguacguagcagagcagAucc.....................................................................  1     1     7y1
......................Acguacguagcagagcagcucc.....................................................................  15    1     7y1
......................Gcguacguagcagagcagcucc.....................................................................  5     1     7y1
......................ucguacguagcagaCcagcucc.....................................................................  1     1     7y1
......................ucguacguagcagagcagcucc.....................................................................  1758  0     7y1
......................ucguacguagcagagcagcCcc.....................................................................  1     1     7y1
......................ucguaAguagcagagcagcucc.....................................................................  4     1     7y1
......................ucguacguaUcagagcagcucc.....................................................................  2     1     7y1
......................ucguaUguagcagagcagcucc.....................................................................  1     1     7y1
......................ucguacguagcagagcagcuAc.....................................................................  4     1     7y1
......................ucguacguagcagagcagcuUc.....................................................................  1     1     7y1
......................uAguacguagcagagcagcucc.....................................................................  1     1     7y1
......................ucguacguagcagGgcagcucc.....................................................................  1     1     7y1
......................ucguGcguagcagagcagcucc.....................................................................  1     1     7y1
......................ucguacguaCcagagcagcucc.....................................................................  4     1     7y1
......................ucguacguagcagaAcagcucc.....................................................................  1     1     7y1
......................ucCuacguagcagagcagcucc.....................................................................  2     1     7y1
......................ucguacguagcagUgcagcuccc....................................................................  1     1     7y1
......................Gcguacguagcagagcagcuccc....................................................................  4     1     7y1
......................ucguacguagcagagcagcuAcc....................................................................  3     1     7y1
......................ucguacguagcagagcagcuccc....................................................................  2876  0     7y1
......................ucguacguagcagagcagcAccc....................................................................  1     1     7y1
......................ucguacguagcagagcagcucAc....................................................................  5     1     7y1
......................ucguacguagAagagcagcuccc....................................................................  3     1     7y1
......................ucguaAguagcagagcagcuccc....................................................................  8     1     7y1
......................uAguacguagcagagcagcuccc....................................................................  2     1     7y1
......................ucguacguagcagagcaUcuccc....................................................................  1     1     7y1
......................ucguacguaUcagagcagcuccc....................................................................  1     1     7y1
......................uUguacguagcagagcagcuccc....................................................................  2     1     7y1
......................ucguacguagcagagcaCcuccc....................................................................  2     1     7y1
......................ucguacguagcagagAagcuccc....................................................................  4     1     7y1
......................ucguacguaCcagagcagcuccc....................................................................  2     1     7y1
......................ucguacguagcagCgcagcuccc....................................................................  7     1     7y1
......................ucguacguagcagagcagcuccA....................................................................  2     1     7y1
......................ucAuacguagcagagcagcuccc....................................................................  2     1     7y1
......................ucUuacguagcagagcagcuccc....................................................................  1     1     7y1
......................ucgAacguagcagagcagcuccc....................................................................  1     1     7y1
......................ucguacguagcagagcagcuGcc....................................................................  1     1     7y1
......................ucguUcguagcagagcagcuccc....................................................................  1     1     7y1
......................ucguacguagcagagcagAuccc....................................................................  4     1     7y1
......................Acguacguagcagagcagcuccc....................................................................  26    1     7y1
......................uGguacguagcagagcagcuccc....................................................................  1     1     7y1
......................ucguacCuagcagagcagcuccc....................................................................  1     1     7y1
......................ucguacgAagcagagcagcuccc....................................................................  1     1     7y1
......................ucguacguagcagagGagcuccc....................................................................  1     1     7y1
......................ucguacguagcagaAcagcuccc....................................................................  1     1     7y1
......................ucguacguagcaCagcagcuccc....................................................................  2     1     7y1
......................ucguacgAagcagagcagcucccu...................................................................  3     1     7y1
......................Ccguacguagcagagcagcucccu...................................................................  1     1     7y1
......................ucguacguagcagCgcagcucccu...................................................................  6     1     7y1
......................uUguacguagcagagcagcucccu...................................................................  1     1     7y1
......................ucguacguagcagagcagcucGcu...................................................................  1     1     7y1
......................ucguacguagcUgagcagcucccu...................................................................  1     1     7y1
......................ucguacguagcagagcagcuGccu...................................................................  1     1     7y1
......................ucguacguagcagagcagUucccu...................................................................  1     1     7y1
......................ucguacguaUcagagcagcucccu...................................................................  1     1     7y1
......................ucguacguagcagagcagcuAccu...................................................................  2     1     7y1
......................Gcguacguagcagagcagcucccu...................................................................  5     1     7y1
......................ucguacguagcagaCcagcucccu...................................................................  2     1     7y1
......................ucguacguagcaUagcagcucccu...................................................................  2     1     7y1
......................uAguacguagcagagcagcucccu...................................................................  6     1     7y1
......................ucguacguagcagagAagcucccu...................................................................  1     1     7y1
......................ucguacguagcagagcagcCcccu...................................................................  1     1     7y1
......................ucguacguagcagagcCgcucccu...................................................................  2     1     7y1
```

```
   Star                                                      Mature
gaccugcuucugggucgggguuucguacguagcagagcagcucccucgcugcgaucuauugaaagucagcccucgacacaaggguuuguccgcgcgcgcgcgcgcgcgugcgu
......................ucguacguGgcagagcagcucccu...................................................................    2      1     7y1
......................ucguacguagcagagcagAucccu...................................................................    1      1     7y1
......................ucguGcguagcagagcagcucccu...................................................................    1      1     7y1
......................ucguacguaCcagagcagcucccu...................................................................    1      1     7y1
......................ucguaAguagcagagcagcucccu...................................................................    3      1     7y1
......................ucguacguagcagagcagcucccG...................................................................    9      1     7y1
......................ucguacguagcGgagcagcucccu...................................................................    2      1     7y1
......................ucgAacguagcagagcagcucccu...................................................................    1      1     7y1
......................ucguacguagcagagcagcucccu...................................................................    2486   0     7y1
......................ucguacguagcagagcagcucAcu...................................................................    4      1     7y1
......................ucCuacguagcagagcagcucccu...................................................................    2      1     7y1
......................ucguacguagcagagUagcucccu...................................................................    1      1     7y1
......................ucguacguagcagagcagcuUccu...................................................................    1      1     7y1
......................ucguacguagcagagcagcucccA...................................................................    5      1     7y1
......................ucguacguagcagagcagGucccu...................................................................    1      1     7y1
......................ucguacguagAagagcagcucccu...................................................................    1      1     7y1
......................Acguacguagcagagcagcucccu...................................................................    21     1     7y1
......................ucguacguagcagagcagcuccAu...................................................................    6      1     7y1
......................ucguacUuagcagagcagcucccu...................................................................    3      1     7y1
......................ucguacCuagcagagcagcucccu...................................................................    1      1     7y1
......................uUguacguagcagagcagcucccuc..................................................................    1      1     7y1
......................ucguacCuagcagagcagcucccuc..................................................................    1      1     7y1
......................ucgAacguagcagagcagcucccuc..................................................................    1      1     7y1
......................ucguacguagcagagcagcuAccuc..................................................................    1      1     7y1
......................ucguacguagcagagcagcuUccuc..................................................................    1      1     7y1
......................ucguacguagAagagcagcucccuc..................................................................    1      1     7y1
......................ucCuacguagcagagcagcucccuc..................................................................    1      1     7y1
......................ucguacguagcagagcagcucccuG..................................................................    1      1     7y1
......................ucguacguagcagagcagcucccuc..................................................................    555    0     7y1
......................Acguacguagcagagcagcucccuc..................................................................    7      1     7y1
......................ucguacguaCcagagcagcucccuc..................................................................    1      1     7y1
......................ucguaAguagcagagcagcucccuc..................................................................    1      1     7y1
......................Ncguacguagcagagcagcucccuc..................................................................    1      1     7y1
......................ucguacguagcagagcagcAcccuc..................................................................    1      1     7y1
......................ucguacguagcagagcGgcucccuc..................................................................    1      1     7y1
......................ucguacguagcagagcagcucAcuc..................................................................    1      1     7y1
......................ucguacguagcagCgcagcucccucg.................................................................    1      1     7y1
......................Ncguacguagcagagcagcucccucg.................................................................    1      1     7y1
......................ucguacguagcagagcagcuccAucg.................................................................    1      1     7y1
......................ucguacguaCcagagcagcucccucg.................................................................    1      1     7y1
......................ucguacguagcagaUcagcucccucg.................................................................    1      1     7y1
......................ucguacgAagcagagcagcucccucg.................................................................    1      1     7y1
......................ucCuacguagcagagcagcucccucg.................................................................    1      1     7y1
......................ucguacguagcagagcaCcucccucg.................................................................    1      1     7y1
......................ucguacguagcCgagcagcucccucg.................................................................    1      1     7y1
......................ucguacguagcagagcagcucccucg.................................................................    219    0     7y1
......................ucguacAuagcagagcagcucccucg.................................................................    1      1     7y1
......................uAguacguagcagagcagcucccucg.................................................................    1      1     7y1
......................ucguaAguagcagagcagcucccucgc................................................................    2      1     7y1
......................ucguacguagcagagcagcucccuAgc................................................................    1      1     7y1
......................ucguacguagcagagcagcuAccucgc................................................................    1      1     7y1
......................ucguacguagcagagcagcucccucgc................................................................    270    0     7y1
......................Acguacguagcagagcagcucccucgc................................................................    4      1     7y1
......................uAguacguagcagagcagcucccucgc................................................................    2      1     7y1
......................ucguacguagcagagAagcucccucgc................................................................    1      1     7y1
......................ucguacguagcagagcagcucccucAcu...............................................................    1      1     7y1
......................ucguacguagcagaAcagcucccucgcu...............................................................    1      1     7y1
......................ucguacguaAcagagcagcucccucgcu...............................................................    1      1     7y1
......................ucguacguagcagagcagcCcccucgcu...............................................................    1      1     7y1
......................ucguacguaCcagagcagcucccucgcu...............................................................    1      1     7y1
......................ucguacguagcagagcagcucccucgcA...............................................................    1      1     7y1
......................ucguacguagcagagcagcuAccucgcu...............................................................    1      1     7y1
......................ucguacUuagcagagcagcucccucgcu...............................................................    1      1     7y1
......................ucguacguagcagUgcagcucccucgcu...............................................................    1      1     7y1
......................ucguacguagcagagcagcucccAcgcu...............................................................    1      1     7y1
......................ucguacguagAagagcagcucccucgcu...............................................................    1      1     7y1
......................ucguacguagcagagcagcucccucgcu...............................................................    1342   0     7y1
......................uUguacguagcagagcagcucccucgcu...............................................................    1      1     7y1
......................ucguacguagcagagAagcucccucgcu...............................................................    3      1     7y1
......................ucguacAuagcagagcagcucccucgcu...............................................................    2      1     7y1
```

**Star** **Mature**

```
gaccugcuucugggucgggguuucguacguagcagagcagcucccucgcugcgaucuauugaaagucagcccucgacacaaggguuuguccgcgcgcgcgcgcgcgcgugcgu
......................ucguacgAagcagagcagcucccucgcu...............................................................   4     1   7y1
......................ucguacguagcagagcagcucccucUcu...............................................................   1     1   7y1
......................ucguacguagcagagcagcuUccucgcu...............................................................   1     1   7y1
......................ucguaUguagcagagcagcucccucgcu...............................................................   1     1   7y1
......................ucguacguagcagagcagcucccucgcG...............................................................   16    1   7y1
......................ucguacguaUcagagcagcucccucgcu...............................................................   1     1   7y1
......................ucguacguagcagagcagcucGcucgcu...............................................................   1     1   7y1
......................ucguacCuagcagagcagcucccucgcu...............................................................   1     1   7y1
......................ucguacguagcagagcagcucAcucgcu...............................................................   1     1   7y1
......................ucguacguagcagagcagAucccucgcu...............................................................   2     1   7y1
......................ucguacguagcagagcagcucccuAgcu...............................................................   3     1   7y1
......................ucguacguagcagagcagcucccucCcu...............................................................   2     1   7y1
......................ucguaAguagcagagcagcucccucgcu...............................................................   1     1   7y1
......................ucguacguagcagagcagcuccAucgcu...............................................................   1     1   7y1
......................Acguacguagcagagcagcucccucgcu...............................................................   11    1   7y1
......................Ncguacguagcagagcagcucccucgcu...............................................................   3     1   7y1
......................uAguacguagcagagcagcucccucgcu...............................................................   2     1   7y1
......................ucguacguagcGgagcagcucccucgcu...............................................................   1     1   7y1
......................ucguacguagcagagcagcuAccucgcug..............................................................   3     1   7y1
......................ucguacguagcagagcagUucccucgcug..............................................................   1     1   7y1
......................uAguacguagcagagcagcucccucgcug..............................................................   3     1   7y1
......................Acguacguagcagagcagcucccucgcug..............................................................   17    1   7y1
......................ucguacguagcagagcagcuccAucgcug..............................................................   4     1   7y1
......................ucguacguagcagagcagcucccucUcug..............................................................   1     1   7y1
......................ucguacguagGagagcagcucccucgcug..............................................................   1     1   7y1
......................ucguacguaUcagagcagcucccucgcug..............................................................   1     1   7y1
......................ucguacguagcagagcagcucccuUgcug..............................................................   1     1   7y1
......................ucguacguagAagagcagcucccucgcug..............................................................   3     1   7y1
......................ucgAacguagcagagcagcucccucgcug..............................................................   1     1   7y1
......................ucguacguagcagagcagcucccucgcuC..............................................................   1     1   7y1
......................ucguGcguagcagagcagcucccucgcug..............................................................   1     1   7y1
......................ucguacguGgcagagcagcucccucgcug..............................................................   1     1   7y1
......................Gcguacguagcagagcagcucccucgcug..............................................................   3     1   7y1
......................ucguacCuagcagagcagcucccucgcug..............................................................   3     1   7y1
......................uGguacguagcagagcagcucccucgcug..............................................................   2     1   7y1
......................ucguacguaCcagagcagcucccucgcug..............................................................   1     1   7y1
......................ucguacguagcagagcagcuUccucgcug..............................................................   1     1   7y1
......................ucUuacguagcagagcagcucccucgcug..............................................................   1     1   7y1
......................ucguacguagcagagcagcucccAcgcug..............................................................   4     1   7y1
......................ucguaAguagcagagcagcucccucgcug..............................................................   2     1   7y1
......................ucguacguagcagagcagcucAcucgcug..............................................................   3     1   7y1
......................ucguacguagcaCagcagcucccucgcug..............................................................   1     1   7y1
......................ucCuacguagcagagcagcucccucgcug..............................................................   3     1   7y1
......................ucguacguagcagagcagcucccucCcug..............................................................   1     1   7y1
......................ucguacguagcagagcagcucccucgcug..............................................................   2333  0   7y1
......................ucguacgAagcagagcagcucccucgcug..............................................................   4     1   7y1
......................Ncguacguagcagagcagcucccucgcug..............................................................   1     1   7y1
......................ucguacguagcagagcagcucccucgcuU..............................................................   6     1   7y1
......................Ccguacguagcagagcagcucccucgcug..............................................................   1     1   7y1
......................ucguacguagcagagcagcCcccucgcug..............................................................   2     1   7y1
......................ucguacguagcagagcagcucccucgcGg..............................................................   8     1   7y1
......................ucguacguagUagagcagcucccucgcug..............................................................   1     1   7y1
......................ucguacguagcagagcagcucccucgAug..............................................................   1     1   7y1
......................ucguacguagcagCgcagcucccucgcug..............................................................   2     1   7y1
.......................cguacguagUagagcagc........................................................................   1     1   7y1
.......................cguacguagcaUagcagc........................................................................   1     1   7y1
.......................cguacguagcagaUcagc........................................................................   2     1   7y1
.......................cguacguaAcagagcagc........................................................................   1     1   7y1
.......................cguacguagcagagcCgc........................................................................   1     1   7y1
.......................cguacguagcagagcagc........................................................................   1761  0   7y1
.......................cguacguaUcagagcagc........................................................................   3     1   7y1
.......................cguacguagcUgagcagc........................................................................   1     1   7y1
.......................cguacguagcGgagcagc........................................................................   2     1   7y1
.......................cguacAuagcagagcagc........................................................................   1     1   7y1
.......................cguacguagcagagAagc........................................................................   2     1   7y1
.......................cgAacguagcagagcagc........................................................................   2     1   7y1
.......................cUuacguagcagagcagc........................................................................   3     1   7y1
.......................cguacguagcagagcaUc........................................................................   1     1   7y1
.......................Gguacguagcagagcagc........................................................................   2     1   7y1
.......................cguacgCagcagagcagc........................................................................   2     1   7y1
```

```
 Star                                                                   Mature
gaccugcuucugggucgggguuucguacguagcagagcagcucccucgcugcgaucuauugaaagucagcccucgacacaaggguuuguccgcgcgcgcgcgcgcgugcgu
.........................uacguagcagagcagcucccucgcAg............................................................ 1     1    7y1
.........................uacguagcagagcagcucccucgAug............................................................ 2     1    7y1
.........................Aacguagcagagcagcucccucgcug............................................................ 17    1    7y1
.........................uacguUgcagagcagcucccucgcug............................................................ 1     1    7y1
.........................uacguagcagagcagcucccucgcCg............................................................ 1     1    7y1
.........................uUcguagcagagcagcucccucgcug............................................................ 1     1    7y1
.........................uacguagcagagcagcucccucgcug............................................................ 1768  0    7y1
.........................uacguaCcagagcagcucccucgcug............................................................ 1     1    7y1
.........................uacguagAagagcagcucccucgcug............................................................ 3     1    7y1
.........................uaAguagcagagcagcucccucgcug............................................................ 5     1    7y1
.........................uacguagcagagcCgcucccucgcug............................................................ 1     1    7y1
.........................uaGguagcagagcagcucccucgcug............................................................ 3     1    7y1
.........................uacguagcagagcagcucccucgcuU............................................................ 1     1    7y1
.........................Gacguagcagagcagcucccucgcug............................................................ 2     1    7y1
.........................uacguaUcagagcagcucccucgcug............................................................ 1     1    7y1
..........................acguagcagaUcagcucccucgcu............................................................. 1     1    7y1
..........................acguagcagagcagcucccucgcu............................................................. 416   0    7y1
..........................acguagcagagcagcucccucgcG............................................................. 9     1    7y1
..........................Gcguagcagagcagcucccucgcu............................................................. 5     1    7y1
..........................acguagcagagcagAucccucgcu............................................................. 1     1    7y1
..........................acguagcGgagcagcucccucgcu............................................................. 2     1    7y1
..........................acguagcagagcagcucccAcgcu............................................................. 1     1    7y1
..........................acguagcagagcaUcucccucgcu............................................................. 3     1    7y1
..........................acguaCcagagcagcucccucgcu............................................................. 1     1    7y1
..........................acgAagcagagcagcucccucgcu............................................................. 1     1    7y1
..........................acguaUcagagcagcucccucgcu............................................................. 1     1    7y1
..........................acguagcagagAagcucccucgcu............................................................. 1     1    7y1
..........................aAguagcagagcagcucccucgcu............................................................. 2     1    7y1
..........................acguagcagagcagcuAccucgcu............................................................. 1     1    7y1
..........................acguagAagagcagcucccucgcu............................................................. 2     1    7y1
..........................acguagcagagcCgcucccucgcu............................................................. 1     1    7y1
..........................acguagcaUagcagcucccucgcug............................................................ 1     1    7y1
..........................acguagAagagcagcucccucgcug............................................................ 2     1    7y1
..........................acguagcagaUcagcucccucgcug............................................................ 2     1    7y1
..........................acguagcagagAagcucccucgcug............................................................ 1     1    7y1
..........................acguagcagagcagcucccucgcGg............................................................ 5     1    7y1
..........................acguagcagagcGgcucccucgcug............................................................ 1     1    7y1
..........................acguagcagagcagcucccucgcug............................................................ 537   0    7y1
..........................acgAagcagagcagcucccucgcug............................................................ 1     1    7y1
..........................acCuagcagagcagcucccucgcug............................................................ 1     1    7y1
..........................acguagcagagcagcucccucgcuC............................................................ 1     1    7y1
..........................acguagcagagcagcuccAucgcug............................................................ 2     1    7y1
..........................Gcguagcagagcagcucccucgcug............................................................ 1     1    7y1
...........................cguagcagagcagcuccc.................................................................. 199   0    7y1
...........................cguagcagagcagcucGc.................................................................. 1     1    7y1
...........................cguagcagagcagcucAcu................................................................. 1     1    7y1
...........................cguagAagagcagcucccu................................................................. 3     1    7y1
...........................cguagcagagcagcuAccu................................................................. 2     1    7y1
...........................cguagcagaUcagcucccu................................................................. 1     1    7y1
...........................cgGagcagagcagcucccu................................................................. 1     1    7y1
...........................cguagcagagcagcucccu................................................................. 204   0    7y1
...........................cguagcagaCcagcucccu................................................................. 1     1    7y1
...........................cguaCcagagcagcucccucgcug............................................................ 1     1    7y1
...........................cCuagcagagcagcucccucgcug............................................................ 1     1    7y1
...........................cguagcagagcagcuUccucgcug............................................................ 1     1    7y1
...........................cguagcagagcagcucccucAcug............................................................ 1     1    7y1
...........................cguagcagCgcagcucccucgcug............................................................ 1     1    7y1
...........................cguagcagagcagcucccucgAug............................................................ 1     1    7y1
...........................cguagcagagcagcucccucgcuU............................................................ 2     1    7y1
...........................cguagcagagcagcucccucgcug............................................................ 541   0    7y1
...........................cguaUcagagcagcucccucgcug............................................................ 1     1    7y1
...........................cguagcagagcagAucccucgcug............................................................ 1     1    7y1
...........................Nguagcagagcagcucccucgcug............................................................ 2     1    7y1
...........................cguagcagagcagcucccAcgcug............................................................ 1     1    7y1
...........................cgAagcagagcagcucccucgcug............................................................ 1     1    7y1
...........................Gguagcagagcagcucccucgcug............................................................ 1     1    7y1
...........................cguagcagagcagcuGccucgcug............................................................ 1     1    7y1
...........................cguagcagagcagcAcccucgcug............................................................ 1     1    7y1
............................guagcagagcagcucccucgc.............................................................. 2     0    7y1
............................guagcagagcagcucccucgcug............................................................ 73    0    7y1
```

**Star** **Mature**

```
gaccugcuucugggucgggguuucguacguagcagagcagcucccucgcugcgaucuauugaaagucagcccucgacacaaggguuuguccgcgcgcgcgcgcgcgcgugcgu
......................................agcucccucgcugAgaucu......................................................     1     1     7y1
......................................Cgcucccucgcugcgaucu......................................................     2     1     7y1
......................................agcucAcucgcugcgaucu......................................................     1     1     7y1
......................................Cgcucccucgcugcgaucuauugaaaguc............................................     1     1     7y1
......................................agcucccucgcAgcgaucuauugaaaguc............................................     1     1     7y1
......................................agcucccucgcugcgaucuauugaaaguc............................................     185   0     7y1
......................................agcucAcucgcugcgaucuauugaaaguc............................................     2     1     7y1
......................................agcCcccucgcugcgaucuauugaaaguc............................................     1     1     7y1
......................................agcuAccucgcugcgaucuauugaaaguc............................................     2     1     7y1
......................................agcuccAucgcugcgaucuauugaaaguc............................................     1     1     7y1
......................................agcucccucgcugAgaucuauugaaaguc............................................     1     1     7y1
......................................agcucccucgcugcgaucuauugaaagucag..........................................     1     0     7y1
.......................................gcucccucgcugcgaucG......................................................     1     1     7y1
.......................................Ccucccucgcugcgaucu......................................................     1     1     7y1
.......................................gcucccucgcugcgaucu......................................................     194   0     7y1
.......................................gcucccAcgcugcgaucu......................................................     1     1     7y1
.......................................gcucccucUcugcgaucu......................................................     1     1     7y1
.......................................gcucccucgAugcgaucu......................................................     1     1     7y1
.......................................gcuAccucgcugcgaucu......................................................     1     1     7y1
.......................................gcucAcucgcugcgaucuauug..................................................     3     1     7y1
.......................................gcuAccucgcugcgaucuauug..................................................     1     1     7y1
.......................................gcuccAucgcugcgaucuauug..................................................     2     1     7y1
.......................................Ccucccucgcugcgaucuauug..................................................     1     1     7y1
.......................................gcucccucgcugcgaucuauAg..................................................     1     1     7y1
.......................................gcucccucgcugcgaucuauug..................................................     515   0     7y1
.......................................gcucccucgcugcgaucAauug..................................................     1     1     7y1
.......................................gcucUcucgcugcgaucuauug..................................................     1     1     7y1
.......................................gcucccuAgcugcgaucuauug..................................................     1     1     7y1
.......................................gcucccucgcugcgaucuauCg..................................................     1     1     7y1
.......................................gcucccucgcugcgaucuauuU..................................................     1     1     7y1
.......................................gcucccucgcugcgUucuauug..................................................     1     1     7y1
.......................................gcucccucgcugcgaucuauugaGag..............................................     1     1     7y1
.......................................gcucccucCcugcgaucuauugaaag..............................................     1     1     7y1
.......................................gcucccucgcugcgaucuUuugaaag..............................................     1     1     7y1
.......................................gcucccucgcugcgaucuauugaaGg..............................................     1     1     7y1
.......................................gcucAcucgcugcgaucuauugaaag..............................................     2     1     7y1
.......................................gcucccucgcugcgaucuauGgaaag..............................................     1     1     7y1
.......................................gcucccucgcugcgauAuauugaaag..............................................     1     1     7y1
.......................................gcucccucgcugcUaucuauugaaag..............................................     1     1     7y1
.......................................gcuUccucgcugcgaucuauugaaag..............................................     1     1     7y1
.......................................gcucccucAcugcgaucuauugaaag..............................................     1     1     7y1
.......................................gcucccucgcGgcgaucuauugaaag..............................................     1     1     7y1
.......................................gcucccucgcugcgaucuauugGaag..............................................     1     1     7y1
.......................................gcucccucgcuCcgaucuauugaaag..............................................     1     1     7y1
.......................................gcuAccucgcugcgaucuauugaaag..............................................     2     1     7y1
.......................................gcucccucgcugAgaucuauugaaag..............................................     1     1     7y1
.......................................gNucccucgcugcgaucuauugaaag..............................................     1     1     7y1
.......................................gcucccuAgcugcgaucuauugaaag..............................................     1     1     7y1
.......................................gcucccucgcugcgaucuauugaaaC..............................................     2     1     7y1
.......................................gcucccucgAugcgaucuauugaaag..............................................     1     1     7y1
.......................................gcucccucgcugcgaucuauugaaag..............................................     1199  0     7y1
.......................................Ncucccucgcugcgaucuauugaaag..............................................     1     1     7y1
.......................................gcuccAucgcugcgaucuauugaaag..............................................     1     1     7y1
.......................................gcucccucgcAgcgaucuauugaaag..............................................     1     1     7y1
.......................................gcucccAcgcugcgaucuauugaaag..............................................     2     1     7y1
.......................................Ccucccucgcugcgaucuauugaaag..............................................     2     1     7y1
.......................................gAucccucgcugcgaucuauugaaag..............................................     3     1     7y1
.......................................gcucccucgcugcgaAcuauugaaagu.............................................     1     1     7y1
.......................................gcucccucgcCgcgaucuauugaaagu.............................................     1     1     7y1
.......................................gcucccucgcAgcgaucuauugaaagu.............................................     1     1     7y1
.......................................gcucAcucgcugcgaucuauugaaagu.............................................     2     1     7y1
.......................................gcucccucgcugcgaucCauugaaagu.............................................     1     1     7y1
.......................................gcucccucgcuCcgaucuauugaaagu.............................................     1     1     7y1
.......................................gcucccucgcugcgaucAauugaaagu.............................................     1     1     7y1
.......................................gcuccAucgcugcgaucuauugaaagu.............................................     1     1     7y1
.......................................gcucccucgcugcgaucuauAgaaagu.............................................     1     1     7y1
.......................................gcucccucgcugcgaucuauugaaaUu.............................................     1     1     7y1
.......................................gcucccucgcugcgaucuauugaaagu.............................................     906   0     7y1
.......................................gcuAccucgcugcgaucuauugaaagu.............................................     1     1     7y1
.......................................gcucccucgcugcUaucuauugaaagu.............................................     1     1     7y1
```

Star Mature

```
gaccugcuucugggucggggguucguacguagcagagcagcucccucgcugcgaucuauugaaagucagcccucgacacaaggguuuguccgcgcgcgcgcgcgcgcgugcgu
.......................................gcucccucgcugcgauGuauugaaagu...............................................  1     1  7y1
.......................................gcucccuAgcugcgaucuauugaaagu...............................................  2     1  7y1
.......................................gcucccucgcugcgaucuauugaaagA...............................................  1     1  7y1
.......................................Ccucccucgcugcgaucuauugaaagu...............................................  2     1  7y1
.......................................Ucucccucgcugcgaucuauugaaagu...............................................  1     1  7y1
.......................................gcucccucgcugcgaucuaGugaaagu...............................................  1     1  7y1
.......................................gcucccucgcugcgaucuauugaaagG...............................................  1     1  7y1
.......................................gcucccucgAugcgaucuauugaaagu...............................................  2     1  7y1
.......................................gcucccucgcugcgaucuauugaaaguU..............................................  1     1  7y1
.......................................gcuGccucgcugcgaucuauugaaaguc..............................................  1     1  7y1
.......................................gcucccucgcugcgaucuauugaaaguc..............................................  229   0  7y1
.......................................gcucccucgcugAgaucuauugaaaguc..............................................  1     1  7y1
.......................................gcucAcucgcugcgaucuauugaaaguc..............................................  1     1  7y1
.......................................gcuccAucgcugcgaucuauugaaaguc..............................................  1     1  7y1
.......................................Ucucccucgcugcgaucuauugaaaguc..............................................  1     1  7y1
........................................cucccucgcugcgaucuaAugaa..................................................  1     1  7y1
........................................cucccAcgcugcgaucuauugaa..................................................  1     1  7y1
........................................cucccucgcugAgaucuauugaa..................................................  2     1  7y1
........................................cucccucgcugcgauAuauugaa..................................................  1     1  7y1
........................................cucccucgcAgcgaucuauugaa..................................................  1     1  7y1
........................................cucccucgcugcgaAcuauugaa..................................................  1     1  7y1
........................................cucccucgcugcCaucuauugaa..................................................  1     1  7y1
........................................cucccucgcugcgaucuauugaa..................................................  144   0  7y1
........................................cucccucgcugcgaucuauugaaa.................................................  244   0  7y1
........................................cucccucgcugcgaucuauGgaaa.................................................  1     1  7y1
........................................cucccucgcugAgaucuauugaaa.................................................  1     1  7y1
........................................cuAccucgcugcgaucuauugaaa.................................................  3     1  7y1
........................................Aucccucgcugcgaucuauugaaa.................................................  2     1  7y1
........................................cucccucgcuCcgaucuauugaaa.................................................  1     1  7y1
........................................cucccuAgcugcgaucuauugaaag................................................  1     1  7y1
........................................cucccucgcuCcgaucuauugaaag................................................  1     1  7y1
........................................cucccucgcuUcgaucuauugaaag................................................  1     1  7y1
........................................cuccAucgcugcgaucuauugaaag................................................  1     1  7y1
........................................cucAcucgcugcgaucuauugaaag................................................  1     1  7y1
........................................cucccucgcugcgaucuauugaaag................................................  185   0  7y1
........................................cucccucgcuAcgaucuauugaaagu...............................................  1     1  7y1
........................................Aucccucgcugcgaucuauugaaagu...............................................  6     1  7y1
........................................cucccucgAugcgaucuauugaaagu...............................................  2     1  7y1
........................................cucAcucgcugcgaucuauugaaagu...............................................  1     1  7y1
........................................cucccucgcugcgaucuauAgaaagu...............................................  1     1  7y1
........................................cucccucgcuCcgaucuauugaaagu...............................................  2     1  7y1
........................................cucccucCcugcgaucuauugaaagu...............................................  2     1  7y1
........................................cucccucgcAgcgaucuauugaaagu...............................................  2     1  7y1
........................................cucccucgcugcUaucuauugaaagu...............................................  1     1  7y1
........................................cucccucgcugcgaucuauugaaaUu...............................................  1     1  7y1
........................................cucccucgcugcgaucuauugaaagA...............................................  5     1  7y1
........................................cucccucUcugcgaucuauugaaagu...............................................  3     1  7y1
........................................cucccucgcugcgaucuaAugaaagu...............................................  1     1  7y1
........................................cuccUucgcugcgaucuauugaaagu...............................................  1     1  7y1
........................................cucccucgcugcgCucuauugaaagu...............................................  1     1  7y1
........................................cucccucgcugcgaucuauugaaagG...............................................  1     1  7y1
........................................cuAccucgcugcgaucuauugaaagu...............................................  9     1  7y1
........................................cucccucgcugAgaucuauugaaagu...............................................  2     1  7y1
........................................cucccuAgcugcgaucuauugaaagu...............................................  4     1  7y1
........................................cucccucgcugcgaucuauuGGaagu...............................................  1     1  7y1
........................................cucccucgcugcgaucuauuAaaagu...............................................  1     1  7y1
........................................cAcccucgcugcgaucuauugaaagu...............................................  7     1  7y1
........................................cucccucgcugcgaucuauugaUagu...............................................  1     1  7y1
........................................cucccucgcugcgaucAauugaaagu...............................................  2     1  7y1
........................................cucccucgcugcgaucuauugaaagu...............................................  1893  0  7y1
........................................cucccucgcugcgaucuauugaaaCu...............................................  1     1  7y1
........................................cucccAcgcugcgaucuauugaaagu...............................................  5     1  7y1
........................................cucccucgcugcgaucuauugaUaguc..............................................  1     1  7y1
........................................cucccucgcugcUaucuauugaaaguc..............................................  3     1  7y1
........................................cucccucgcugcgaucuauAgaaaguc..............................................  8     1  7y1
........................................cucccucgcugcgaucuauuCaaaguc..............................................  1     1  7y1
........................................cucccucgcuCcgaucuauugaaaguc..............................................  3     1  7y1
........................................cuAccucgcugcgaucuauugaaaguc..............................................  10    1  7y1
........................................cucccucgcugcgaAcuauugaaaguc..............................................  3     1  7y1
........................................Nucccucgcugcgaucuauugaaaguc..............................................  3     1  7y1
```

**Star** **Mature**

```
gaccugcuucugggucgggguuucguacguagcagagcagcucccucgcugcgaucuauugaaagucagcccucgacacaaggguuuguccgcgcgcgcgcgcgcgcgcgugcgu
........................................cucccucgcugcgaucuauugaaagucagcc............................................   172   0   7y1
.........................................ucccucgcugcUaucuau........................................................   1     1   7y1
.........................................ucccucgcugcgaAcuau........................................................   1     1   7y1
.........................................Acccucgcugcgaucuau........................................................   1     1   7y1
.........................................ucccucgcugcgaCcuau........................................................   1     1   7y1
.........................................ucccucgcugcgaucuau........................................................   200   0   7y1
.........................................Ncccucgcugcgaucuau........................................................   1     1   7y1
.........................................ucccucgcugcgaucuaG........................................................   1     1   7y1
.........................................uAccucgcugcgaucuauu.......................................................   1     1   7y1
.........................................ucccucgcugcgaucuauu.......................................................   189   0   7y1
.........................................uUccucgcugcgaucuauu.......................................................   1     1   7y1
.........................................Acccucgcugcgaucuauu.......................................................   1     1   7y1
.........................................ucAcucgcugcgaucuauu.......................................................   1     1   7y1
.........................................Gcccucgcugcgaucuauu.......................................................   1     1   7y1
.........................................ucccucgcugcUaucuauu.......................................................   2     1   7y1
.........................................ucccucgcugcgaAcuauug......................................................   1     1   7y1
.........................................ucccucgcugcUaucuauug......................................................   1     1   7y1
.........................................ucccucgcugcgaucuauug......................................................   266   0   7y1
.........................................Acccucgcugcgaucuauug......................................................   4     1   7y1
.........................................ucccucgcugGgaucuauug......................................................   1     1   7y1
.........................................ucccuAgcugcgaucuauug......................................................   1     1   7y1
.........................................Acccucgcugcgaucuauugaaa...................................................   2     1   7y1
.........................................ucccucgcugcgaucuauugaaa...................................................   212   0   7y1
.........................................ucccucgcugcgaucuauuUaaa...................................................   1     1   7y1
.........................................ucccucUcugcgaucuauugaaa...................................................   1     1   7y1
.........................................Gcccucgcugcgaucuauugaaa...................................................   1     1   7y1
.........................................Ccccucgcugcgaucuauugaaa...................................................   1     1   7y1
.........................................ucccuAgcugcgaucuauugaaag..................................................   1     1   7y1
.........................................ucccucgcugcgauAuauugaaag..................................................   1     1   7y1
.........................................ucccucgcugcgaucAauugaaag..................................................   1     1   7y1
.........................................Acccucgcugcgaucuauugaaag..................................................   7     1   7y1
.........................................ucccucgcugcgaucuauAgaaag..................................................   1     1   7y1
.........................................ucccucgcugcgauUuauugaaag..................................................   1     1   7y1
.........................................ucccucgcugcgaucuauuAaaag..................................................   1     1   7y1
.........................................ucccucgcuUcgaucuauugaaag..................................................   1     1   7y1
.........................................ucAcucgcugcgaucuauugaaag..................................................   3     1   7y1
.........................................Ncccucgcugcgaucuauugaaag..................................................   3     1   7y1
.........................................Gcccucgcugcgaucuauugaaag..................................................   1     1   7y1
.........................................uAccucgcugcgaucuauugaaag..................................................   2     1   7y1
.........................................ucccucgcugcgaucuauugCaag..................................................   1     1   7y1
.........................................ucccucgcugcgaucuauugaaag..................................................   935   0   7y1
.........................................ucccucgcugcgaucuauugGaag..................................................   1     1   7y1
.........................................ucccucgcugAgaucuauugaaag..................................................   1     1   7y1
.........................................ucccucUcugcgaucuauugaaag..................................................   2     1   7y1
.........................................ucccucgcAgcgaucuauugaaag..................................................   2     1   7y1
.........................................ucccucgcuCcgaucuauugaaag..................................................   3     1   7y1
.........................................ucccucgcugcgaucuauugaaagu.................................................   682   0   7y1
.........................................ucccucgcAgcgaucuauugaaagu.................................................   2     1   7y1
.........................................ucccucgcugAgaucuauugaaagu.................................................   4     1   7y1
.........................................Acccucgcugcgaucuauugaaagu.................................................   12    1   7y1
.........................................ucccucUcugcgaucuauugaaagu.................................................   1     1   7y1
.........................................uccAucgcugcgaucuauugaaagu.................................................   1     1   7y1
.........................................ucAcucgcugcgaucuauugaaagu.................................................   1     1   7y1
.........................................ucccucgcugcgauAuauugaaagu.................................................   2     1   7y1
.........................................ucccucgcugcUaucuauugaaagu.................................................   1     1   7y1
.........................................ucccucgcugcgaucAauugaaagu.................................................   1     1   7y1
.........................................ucccucgcugcgaucuauugaaagA.................................................   1     1   7y1
.........................................Gcccucgcugcgaucuauugaaagu.................................................   1     1   7y1
.........................................uAccucgcugcgaucuauugaaagu.................................................   1     1   7y1
.........................................ucccucgcugcgaucuaAugaaagu.................................................   1     1   7y1
.........................................ucccucgcuUcgaucuauugaaagu.................................................   1     1   7y1
.........................................Gcccucgcugcgaucuauugaaaguc................................................   2     1   7y1
.........................................ucccucgcugcgaucuauugaaaAuc................................................   1     1   7y1
.........................................ucccucgcugcgaucuaAugaaaguc................................................   1     1   7y1
.........................................ucccucgcuCcgaucuauugaaaguc................................................   2     1   7y1
.........................................ucccucgcugcgaucuauugaaaCuc................................................   3     1   7y1
.........................................ucccucgcugcgaucAauugaaaguc................................................   3     1   7y1
.........................................ucccucgcAgcgaucuauugaaaguc................................................   4     1   7y1
.........................................ucccucgcugcgaucuauAgaaaguc................................................   2     1   7y1
.........................................ucAcucgcugcgaucuauugaaaguc................................................   10    1   7y1
```

**Star** **Mature**

```
gaccugcuucugggucggggguucguacguagcagagcagcucccucgcugcgaucuauugaaagucagcccucgacacaaggguuuguccgcgcgcgcgcgcgcgugcgu
.........................................ucccucgcugcgaucuauugGaaguc............................................ 1 1 7y1
.........................................uAccucgcugcgaucuauugaaaguc............................................ 3 1 7y1
.........................................ucccucgcugcUaucuauugaaaguc............................................ 1 1 7y1
.........................................ucccucgAugcgaucuauugaaaguc............................................ 3 1 7y1
.........................................ucccucgcugcgaucuauugaaaguc............................................ 2692 0 7y1
.........................................ucccuAgcugcgaucuauugaaaguc............................................ 7 1 7y1
.........................................ucccucgcugcgaucuauugaaaguA............................................ 1 1 7y1
.........................................ucccucgcGgcgaucuauugaaaguc............................................ 1 1 7y1
.........................................ucccucgcuUcgaucuauugaaaguc............................................ 2 1 7y1
.........................................ucccucAcugcgaucuauugaaaguc............................................ 1 1 7y1
.........................................ucccucgcugcgaucuGuugaaaguc............................................ 2 1 7y1
.........................................Acccucgcugcgaucuauugaaaguc............................................ 38 1 7y1
.........................................ucccucgcugAgaucuauugaaaguc............................................ 2 1 7y1
.........................................ucccAcgcugcgaucuauugaaaguc............................................ 1 1 7y1
.........................................ucccucgcugcCaucuauugaaaguc............................................ 3 1 7y1
.........................................ucccucgcugcgaucuauugaaagAc............................................ 2 1 7y1
.........................................ucccucUcugcgaucuauugaaaguc............................................ 2 1 7y1
.........................................ucccucgcugcgauAuauugaaaguc............................................ 4 1 7y1
.........................................uccAucgcugcgaucuauugaaaguc............................................ 2 1 7y1
.........................................Ncccucgcugcgaucuauugaaaguc............................................ 1 1 7y1
.........................................ucccucgGugcgaucuauugaaaguca........................................... 1 1 7y1
.........................................ucccucgcugcgaucuauugaaaguca........................................... 586 0 7y1
.........................................Ncccucgcugcgaucuauugaaaguca........................................... 2 1 7y1
.........................................ucccucgcugcgaucCauugaaaguca........................................... 1 1 7y1
.........................................ucccucgcugcgauAuauugaaaguca........................................... 1 1 7y1
.........................................Gcccucgcugcgaucuauugaaaguca........................................... 1 1 7y1
.........................................ucccucgcugAgaucuauugaaaguca........................................... 1 1 7y1
.........................................ucccucgcugcgaucuauugaaagucU........................................... 1 1 7y1
.........................................ucccucUcugcgaucuauugaaaguca........................................... 1 1 7y1
.........................................ucccucgcuAcgaucuauugaaaguca........................................... 1 1 7y1
.........................................ucccucgcuCcgaucuauugaaaguca........................................... 1 1 7y1
.........................................uccAucgcugcgaucuauugaaaguca........................................... 1 1 7y1
.........................................ucccucgcugcCaucuauugaaaguca........................................... 1 1 7y1
.........................................ucccucgAugcgaucuauugaaaguca........................................... 2 1 7y1
.........................................ucccucCcugcgaucuauugaaaguca........................................... 1 1 7y1
.........................................Acccucgcugcgaucuauugaaaguca........................................... 6 1 7y1
.........................................ucccucgcuUcgaucuauugaaagucag.......................................... 1 1 7y1
.........................................ucAcucgcugcgaucuauugaaagucag.......................................... 1 1 7y1
.........................................ucccucgcugcgaucuauAgaaagucag.......................................... 1 1 7y1
.........................................ucccucgcugcgaucuauugaaagucag.......................................... 159 0 7y1
.........................................Acccucgcugcgaucuauugaaagucag.......................................... 2 1 7y1
.........................................ucccucgcugcgUucuauugaaagucag.......................................... 1 1 7y1
.........................................ucccAcgcugcgaucuauugaaagucag.......................................... 1 1 7y1
..........................................cccucgcugcgaucuauu................................................... 172 0 7y1
..........................................cccucgcugcgaucuaAu................................................... 1 1 7y1
..........................................Gccucgcugcgaucuauu................................................... 1 1 7y1
..........................................ccAucgcugcgaucuauu................................................... 1 1 7y1
..........................................cAcucgcugcgaucuauu................................................... 1 1 7y1
..........................................cccucgcugcCaucuauu................................................... 2 1 7y1
..........................................cccucgcAgcgaucuauu................................................... 1 1 7y1
..........................................cccucgcugcgaucuauuga................................................. 157 0 7y1
..........................................Accucgcugcgaucuauuga................................................. 1 1 7y1
..........................................ccAucgcugcgaucuauuga................................................. 1 1 7y1
..........................................cccucgAugcgaucuauuga................................................. 1 1 7y1
..........................................cAcucgcugcgaucuauuga................................................. 2 1 7y1
..........................................cccucgcAgcgaucuauuga................................................. 1 1 7y1
..........................................cccucgcugcgaucuauugG................................................. 1 1 7y1
..........................................Nccucgcugcgaucuauuga................................................. 1 1 7y1
..........................................cccucgcugcgaucuauuAaa................................................ 1 1 7y1
..........................................cAcucgcugcgaucuauugaaa............................................... 3 1 7y1
..........................................cccucgAugcgaucuauugaaa............................................... 3 1 7y1
..........................................cccucgcugcUaucuauugaaa............................................... 1 1 7y1
..........................................cccucgcugcgaucuaAugaaa............................................... 3 1 7y1
..........................................cccucgcugcgaucuauAgaaa............................................... 1 1 7y1
..........................................cccucgcuUcgaucuauugaaa............................................... 2 1 7y1
..........................................cccucgcugcCaucuauugaaa............................................... 1 1 7y1
..........................................cUcucgcugcgaucuauugaaa............................................... 1 1 7y1
..........................................ccGucgcugcgaucuauugaaa............................................... 1 1 7y1
..........................................cccucgcugcgaucuauugaaa............................................... 1120 0 7y1
..........................................cccucgcugcgaAcuauugaaa............................................... 2 1 7y1
```

```
   Star                                    Mature
gaccugcuucugggucgggguuucguacguagcagagcagcucccucgcugcgaucuauugaaagucagcccucgacacaaggguuuguccgcgcgcgcgcgcgcgugcgu
..........................................cccucgcugcgaAcuauugaaaguc............................................    2      1     7y1
..........................................cccucgUugcgaucuauugaaaguc............................................    1      1     7y1
..........................................cccucgcAgcgaucuauugaaaguc............................................    1      1     7y1
..........................................cAcucgcugcgaucuauugaaaguc............................................    7      1     7y1
..........................................Accucgcugcgaucuauugaaaguc............................................    7      1     7y1
..........................................cccucgcuCcgaucuauugaaaguc............................................    2      1     7y1
..........................................cccucUcugcgaucuauugaaaguc............................................    1      1     7y1
..........................................cccucgcugcgaucuauugaaaguc............................................    1855   0     7y1
..........................................cccucgcugcgaucuauugaaagAc............................................    2      1     7y1
..........................................cccucgcugcgaucuauuUaaaguc............................................    1      1     7y1
..........................................cccucgAugcgaucuauugaaaguc............................................    8      1     7y1
..........................................ccAucgcugcgaucuauugaaaguc............................................    8      1     7y1
..........................................cccucgcugcgaucuauAgaaaguc............................................    1      1     7y1
..........................................Uccucgcugcgaucuauugaaaguc............................................    2      1     7y1
..........................................cccucgcugcgaucuauuAaaaguc............................................    1      1     7y1
..........................................cccucgcugcgaucuauCgaaaguc............................................    1      1     7y1
..........................................cccucgcugGgaucuauugaaaguc............................................    1      1     7y1
..........................................cccucgcugcgaucuauugaaaguAa...........................................    1      1     7y1
..........................................cccuAgcugcgaucuauugaaaguca...........................................    3      1     7y1
..........................................cccucgcugAgaucuauugaaaguca...........................................    2      1     7y1
..........................................ccAucgcugcgaucuauugaaaguca...........................................    2      1     7y1
..........................................cccuUgcugcgaucuauugaaaguca...........................................    2      1     7y1
..........................................Nccucgcugcgaucuauugaaaguca...........................................    3      1     7y1
..........................................cccucgcugcgaucuauugaaagAca...........................................    3      1     7y1
..........................................Accucgcugcgaucuauugaaaguca...........................................    5      1     7y1
..........................................cccucgcugcgauAuauugaaaguca...........................................    2      1     7y1
..........................................cccucUcugcgaucuauugaaaguca...........................................    2      1     7y1
..........................................cccucgcugcgaAcuauugaaaguca...........................................    1      1     7y1
..........................................cccucgcugcgaucuauugaaaguca...........................................    592    0     7y1
..........................................cccAcgcugcgaucuauugaaaguca...........................................    1      1     7y1
..........................................cccucAcugcgaucuauugaaaguca...........................................    1      1     7y1
..........................................cccucgcAgcgaucuauugaaaguca...........................................    2      1     7y1
..........................................cccucgcugcgaucuauugaaaUuca...........................................    1      1     7y1
..........................................cccucgcugcUaucuauugaaaguca...........................................    1      1     7y1
...........................................ccucgAugcgaucuauug..................................................    1      1     7y1
...........................................cAucgcugcgaucuauug..................................................    2      1     7y1
...........................................ccucgcCgcgaucuauug..................................................    1      1     7y1
...........................................ccucgcugcgaucuauug..................................................    209    0     7y1
...........................................ccucgcuUcgaucuauug..................................................    1      1     7y1
...........................................ccucgcugcgaucuauuC..................................................    1      1     7y1
...........................................ccucgcGgcgaucuauug..................................................    1      1     7y1
...........................................Ncucgcugcgaucuauug..................................................    2      1     7y1
...........................................ccucgcugcgaucGauugaa................................................    1      1     7y1
...........................................ccucgcugcUaucuauugaa................................................    2      1     7y1
...........................................ccucgcugcgaucuauAgaa................................................    1      1     7y1
...........................................ccucgcugcgaucuauugaa................................................    602    0     7y1
...........................................Ncucgcugcgaucuauugaa................................................    1      1     7y1
...........................................ccuAgcugcgaucuauugaa................................................    1      1     7y1
...........................................cAucgcugcgaucuauugaa................................................    3      1     7y1
...........................................ccAcgcugcgaucuauugaa................................................    1      1     7y1
...........................................ccucgcAgcgaucuauugaa................................................    2      1     7y1
...........................................ccucCcugcgaucuauugaa................................................    1      1     7y1
...........................................ccucgcugcgaucuauuCaa................................................    1      1     7y1
...........................................ccucgcugcgaucuauuAaa................................................    1      1     7y1
...........................................Acucgcugcgaucuauugaa................................................    1      1     7y1
...........................................ccucgcugcgauAuauugaaa...............................................    4      1     7y1
...........................................Acucgcugcgaucuauugaaa...............................................    6      1     7y1
...........................................ccuAgcugcgaucuauugaaa...............................................    1      1     7y1
...........................................ccucgAugcgaucuauugaaa...............................................    6      1     7y1
...........................................ccucgcugcgaucuaAugaaa...............................................    1      1     7y1
...........................................ccucgcAgcgaucuauugaaa...............................................    9      1     7y1
...........................................ccucgcugcUaucuauugaaa...............................................    5      1     7y1
...........................................ccucgcugAgaucuauugaaa...............................................    1      1     7y1
...........................................cAucgcugcgaucuauugaaa...............................................    2      1     7y1
...........................................Gcucgcugcgaucuauugaaa...............................................    1      1     7y1
...........................................ccucgcugcgaucuauugaGa...............................................    1      1     7y1
...........................................ccucgcuUcgaucuauugaaa...............................................    1      1     7y1
...........................................ccucgcugcgaucAauugaaa...............................................    2      1     7y1
...........................................ccucgcugcgaucuauugaaa...............................................    1604   0     7y1
...........................................ccucgcugcgaucuauugaUa...............................................    1      1     7y1
```

**Star** **Mature**

```
gaccugcuucugggucgggguuucguacguagcagagcagcucccucgcugcgaucuauugaaagucagcccucgacacaaggguuuguccgcgcgcgcgcgcgcgcgugcgu
...........................................ccucgcugcgaucuaAugaaaguc............................................ 3    1  7y1
...........................................ccAcgcugcgaucuauugaaaguc............................................ 2    1  7y1
...........................................ccucgcugcgaucuauuUaaaguc............................................ 1    1  7y1
...........................................ccucgcugcgaucCauugaaaguc............................................ 1    1  7y1
...........................................ccucgcugcgauAuauugaaaguc............................................ 7    1  7y1
...........................................ccucgcugcgUucuauugaaaguc............................................ 1    1  7y1
...........................................ccucgcugcgaucuauugaGaguc............................................ 1    1  7y1
...........................................ccucgcugcgaAcuauugaaaguc............................................ 1    1  7y1
...........................................Gcucgcugcgaucuauugaaaguc............................................ 2    1  7y1
...........................................ccucgcuCcgaucuauugaaaguc............................................ 1    1  7y1
...........................................ccCcgcugcgaucuauugaaaguc............................................ 1    1  7y1
...........................................Acucgcugcgaucuauugaaaguc............................................ 7    1  7y1
...........................................ccucgcugcCaucuauugaaaguc............................................ 4    1  7y1
...........................................ccucgcugcgaucuauugaaaCuc............................................ 5    1  7y1
...........................................ccucgcugcgaucuauGgaaaguc............................................ 1    1  7y1
...........................................ccucgcugcgaucuauugaaaUuc............................................ 1    1  7y1
...........................................ccucgcugcgaucuauugGaaguc............................................ 1    1  7y1
...........................................ccucgcAgcgaucuauugaaaguc............................................ 16   1  7y1
...........................................ccucgcugcgaucAauugaaaguc............................................ 3    1  7y1
...........................................ccucgcCgcgaucuauugaaaguc............................................ 1    1  7y1
...........................................ccucgAugcgaucuauugaaaguc............................................ 6    1  7y1
...........................................cAucgcugcgaucuauugaaaguc............................................ 9    1  7y1
...........................................ccucgcugcgaucuauugaaGguc............................................ 1    1  7y1
...........................................ccucgcugcgGucuauugaaaguc............................................ 1    1  7y1
...........................................ccucUcugcgaucuauugaaaguc............................................ 3    1  7y1
...........................................Ncucgcugcgaucuauugaaaguc............................................ 1    1  7y1
...........................................ccucgcugcgaucuauugaaaguA............................................ 2    1  7y1
...........................................ccucgcugcgaucuauuAaaaguc............................................ 1    1  7y1
...........................................cAucgcugcgaucuauugaaaguca........................................... 2    1  7y1
...........................................Ncucgcugcgaucuauugaaaguca........................................... 1    1  7y1
...........................................ccucgcAgcgaucuauugaaaguca........................................... 1    1  7y1
...........................................ccucgcugAgaucuauugaaaguca........................................... 2    1  7y1
...........................................Acucgcugcgaucuauugaaaguca........................................... 3    1  7y1
...........................................Gcucgcugcgaucuauugaaaguca........................................... 1    1  7y1
...........................................ccucgcugcgaucuauugaaagAca........................................... 1    1  7y1
...........................................ccucgcugcgaucuauugaaaCuca........................................... 1    1  7y1
...........................................ccucgcugcgaucuauugaaaAuca........................................... 1    1  7y1
...........................................ccucgcugcgaucuauugaaaguca........................................... 761  0  7y1
...........................................ccucgcuUcgaucuauugaaaguca........................................... 1    1  7y1
...........................................ccucUcugcgaucuauugaaaguca........................................... 1    1  7y1
...........................................ccucgAugcgaucuauugaaaguca........................................... 1    1  7y1
...........................................ccucgcugcgauAuauugaaaguca........................................... 1    1  7y1
...........................................ccucgcugcgaucuaAugaaaguca........................................... 1    1  7y1
...........................................ccucgcugcgaucuauugaaaUuca........................................... 1    1  7y1
...........................................ccucgcugcgaucuauugaaagucG........................................... 1    1  7y1
...........................................Acucgcugcgaucuauugaaagucag.......................................... 1    1  7y1
...........................................ccucgcuCcgaucuauugaaagucag.......................................... 1    1  7y1
...........................................cAucgcugcgaucuauugaaagucag.......................................... 1    1  7y1
...........................................ccucgcugcgaucuauAgaaagucag.......................................... 1    1  7y1
...........................................ccucgcugcUaucuauugaaagucag.......................................... 1    1  7y1
...........................................Ncucgcugcgaucuauugaaagucag.......................................... 1    1  7y1
...........................................ccucgcugcgaucAauugaaagucag.......................................... 1    1  7y1
...........................................ccucgcAgcgaucuauugaaagucag.......................................... 2    1  7y1
...........................................ccucAcugcgaucuauugaaagucag.......................................... 1    1  7y1
...........................................ccucgcugAgaucuauugaaagucag.......................................... 1    1  7y1
...........................................ccucgcugcgaucuauugaaagucag.......................................... 163  0  7y1
...........................................ccucgcugcgaucuauugaaagucagUcc....................................... 1    1  7y1
...........................................ccucgcugcgaucuauugaaagucagUccu...................................... 253  1  7y1
............................................cucgAugcgaucuauugaaa............................................... 1    1  7y1
............................................cucgcugcgaucuauAgaaa............................................... 2    1  7y1
............................................cucgcugcgauAuauugaaa............................................... 1    1  7y1
............................................cucgcugcgaucuauugaaa............................................... 413  0  7y1
............................................cucUcugcgaucuauugaaa............................................... 2    1  7y1
............................................cucgcugAgaucuauugaaa............................................... 2    1  7y1
............................................cucgcugcUaucuauugaaa............................................... 1    1  7y1
............................................Aucgcugcgaucuauugaaa............................................... 2    1  7y1
............................................cuAgcugcgaucuauugaaa............................................... 3    1  7y1
............................................cucgcugcCaucuauugaaa............................................... 1    1  7y1
............................................cAcgcugcgaucuauugaaa............................................... 2    1  7y1
............................................cuUgcugcgaucuauugaaag.............................................. 1    1  7y1
```

**Star** **Mature**

```
gaccugcuucugggucgggguuucguacguagcagagcagcucccucgcugcgaucuauugaaagucagcccucgacacaaggguuugucccgcgcgcgcgcgcgcgcgugcgu
............................................cucgcugcgaucuauugaaaguc..............................................  2329  0  7y1
............................................cucgcugcgGucuauugaaaguc..............................................  1     1  7y1
............................................cucUcugcgaucuauugaaaguc..............................................  1     1  7y1
............................................cuGgcugcgaucuauugaaaguc..............................................  2     1  7y1
............................................cucgcugcgaucuauuCaaaguc..............................................  4     1  7y1
............................................cucgcAgcgaucuauugaaaguc..............................................  12    1  7y1
............................................cucgcugAgaucuauugaaaguc..............................................  8     1  7y1
............................................cucgcuCcgaucuauugaaaguc..............................................  2     1  7y1
............................................cucgcugcgauAuauugaaaguc..............................................  4     1  7y1
............................................Aucgcugcgaucuauugaaaguc..............................................  10    1  7y1
............................................cucgcugcgaucuauugaaaguA..............................................  2     1  7y1
............................................cucgcugcgaucuauugaaaguca.............................................  238   0  7y1
............................................cucgcugcgaAcuauugaaaguca.............................................  1     1  7y1
............................................cucgcugAgaucuauugaaaguca.............................................  1     1  7y1
............................................Aucgcugcgaucuauugaaaguca.............................................  1     1  7y1
............................................cucgcugcgaucuauugaaaguAa.............................................  3     1  7y1
............................................cuAgcugcgaucuauugaaaguca.............................................  1     1  7y1
.............................................ucgcAgcgaucuauugaa..................................................  1     1  7y1
.............................................ucgcugAgaucuauugaa..................................................  1     1  7y1
.............................................Acgcugcgaucuauugaa..................................................  3     1  7y1
.............................................ucgcugcgaucuauAgaa..................................................  1     1  7y1
.............................................ucgcugcgaucuauugaa..................................................  152   0  7y1
.............................................ucgcugcgaucuauugaaG.................................................  2     1  7y1
.............................................ucgcugcgaucuaCugaaag................................................  1     1  7y1
.............................................ucgcugcgauGuauugaaag................................................  1     1  7y1
.............................................Ncgcugcgaucuauugaaag................................................  2     1  7y1
.............................................ucgcAgcgaucuauugaaag................................................  3     1  7y1
.............................................ucgcugcgNucuauugaaag................................................  1     1  7y1
.............................................ucgcugcgaucuUuugaaag................................................  1     1  7y1
.............................................ucgcugcgaucuauAgaaag................................................  4     1  7y1
.............................................Gcgcugcgaucuauugaaag................................................  1     1  7y1
.............................................ucgcugAgaucuauugaaag................................................  17    1  7y1
.............................................ucgcugcgaucuauugaaaC................................................  3     1  7y1
.............................................ucgcugcgaucuauugaaUg................................................  1     1  7y1
.............................................ucgcCgcgaucuauugaaag................................................  2     1  7y1
.............................................ucgcuCcgaucuauugaaag................................................  1     1  7y1
.............................................ucgcugcgaucuauuCaaag................................................  2     1  7y1
.............................................ucgcugcgaucAauugaaag................................................  6     1  7y1
.............................................ucgAugcgaucuauugaaag................................................  4     1  7y1
.............................................ucgcugcgCucuauugaaag................................................  1     1  7y1
.............................................ucgcugcgaucGauugaaag................................................  1     1  7y1
.............................................ucgcugcgaucuaAugaaag................................................  3     1  7y1
.............................................ucgcugcgaucuauugaaGg................................................  3     1  7y1
.............................................ucgcugcgaucuauugaaag................................................  3906  0  7y1
.............................................ucgcuUcgaucuauugaaag................................................  5     1  7y1
.............................................ucgcugcgaucuauugaaCg................................................  1     1  7y1
.............................................uAgcugcgaucuauugaaag................................................  12    1  7y1
.............................................ucgcugcCaucuauugaaag................................................  1     1  7y1
.............................................ucgcugcgaucuauuUaaag................................................  2     1  7y1
.............................................ucgGugcgaucuauugaaag................................................  2     1  7y1
.............................................ucgcugGgaucuauugaaag................................................  2     1  7y1
.............................................ucgcugcgauNuauugaaag................................................  1     1  7y1
.............................................ucUcugcgaucuauugaaag................................................  1     1  7y1
.............................................ucgcugcgauAuauugaaag................................................  4     1  7y1
.............................................uGgcugcgaucuauugaaag................................................  2     1  7y1
.............................................ucgcugcgaAcuauugaaag................................................  11    1  7y1
.............................................Acgcugcgaucuauugaaag................................................  60    1  7y1
.............................................ucgcuAcgaucuauugaaag................................................  2     1  7y1
.............................................ucgcGgcgaucuauugaaag................................................  1     1  7y1
.............................................ucgcugcUaucuauugaaag................................................  2     1  7y1
.............................................ucgcugcgaucuGuugaaag................................................  1     1  7y1
.............................................Acgcugcgaucuauugaaagu...............................................  4     1  7y1
.............................................ucgcugcCaucuauugaaagu...............................................  1     1  7y1
.............................................ucgcugcgaAcuauugaaagu...............................................  1     1  7y1
.............................................ucgcugcgaucuauugaaagu...............................................  138   0  7y1
.............................................ucgcugcgaucAauugaaagu...............................................  1     1  7y1
.............................................ucgcugcgaucuauugaaagucagcccucgaG....................................  1     1  7y1
.............................................ucgcugcgaucuauugaaagucagcUcucgac....................................  1     1  7y1
.............................................ucgcugcgaucuauugaaagucagccAucgac....................................  1     1  7y1
.............................................Ncgcugcgaucuauugaaagucagcccucgac....................................  1     1  7y1
```

**Star** **Mature**

```
gaccugcuucugggucgggguuucguacguagcagagcagcucccucgcugcgaucuauugaaagucagcccucgacacaaggguuuguccgcgcgcgcgcgcgcgcgugcgu
..............................................cgcugcgaucuauugGaaguc..............................................	1	1	7y1
..............................................cgcugcgaucuauugaGaguc..............................................	1	1	7y1
..............................................cgcugcgaucuUuugaaaguc..............................................	1	1	7y1
..............................................cgcugAgaucuauugaaaguc..............................................	2	1	7y1
..............................................cgcugcgauUuauugaaaguc..............................................	1	1	7y1
..............................................cgcugcgaucuauugaaaguA..............................................	1	1	7y1
..............................................cgcugcgaucuauugaaaguc..............................................	553	0	7y1
..............................................cgcAgcgaucuauugaaaguc..............................................	1	1	7y1
..............................................cgcugcgaucuauugaaGgucag............................................	2	1	7y1
..............................................cgcugcgaucuauugaaagucag............................................	184	0	7y1
..............................................cgcugcgaucuauugaaagucaU............................................	1	1	7y1
..............................................cgcugcgaucuauAgaaagucag............................................	1	1	7y1
..............................................cgcugcgaucuaAugaaagucag............................................	1	1	7y1
..............................................cgcugAgaucuauugaaagucag............................................	1	1	7y1
..............................................cUcugcgaucuauugaaagucag............................................	1	1	7y1
..............................................Agcugcgaucuauugaaagucagcccucgac....................................	1	1	7y1
..............................................cgAugcgaucuauugaaagucagcccucgac....................................	1	1	7y1
..............................................cgcugAgaucuauugaaagucagcccucgac....................................	1	1	7y1
..............................................cgcugcgaucuauugaaagucagcccucgCc....................................	1	1	7y1
..............................................cgcugcgaucuauugaaagucagcccucgac....................................	162	0	7y1
...............................................gcuUcgaucuauugaaag................................................	4	1	7y1
...............................................gcugcgaucAauugaaag................................................	7	1	7y1
...............................................Ucugcgaucuauugaaag................................................	12	1	7y1
...............................................gcGgcgaucuauugaaag................................................	2	1	7y1
...............................................gcugcgaucuauuUaaag................................................	1	1	7y1
...............................................gcugcUaucuauugaaag................................................	5	1	7y1
...............................................gcugcgauAuauugaaag................................................	3	1	7y1
...............................................gcugcgaucuauugaaaC................................................	2	1	7y1
...............................................gcugcgaucuaAugaaag................................................	4	1	7y1
...............................................gcugcCaucuauugaaag................................................	2	1	7y1
...............................................gAugcgaucuauugaaag................................................	6	1	7y1
...............................................gcugcgaucuauugaaag................................................	2699	0	7y1
...............................................gcuCcgaucuauugaaag................................................	1	1	7y1
...............................................gcugcgaucuGuugaaag................................................	1	1	7y1
...............................................gcugcgaucuauuCaaag................................................	1	1	7y1
...............................................gcugcgaucuauugGaag................................................	1	1	7y1
...............................................gcugcgaAcuauugaaag................................................	1	1	7y1
...............................................gcAgcgaucuauugaaag................................................	4	1	7y1
...............................................gcugcgaucuauugaUag................................................	2	1	7y1
...............................................gGugcgaucuauugaaag................................................	2	1	7y1
...............................................Ccugcgaucuauugaaag................................................	8	1	7y1
...............................................gcugcgCucuauugaaag................................................	1	1	7y1
...............................................gcugGgaucuauugaaag................................................	1	1	7y1
...............................................gcugcgaucuauAgaaag................................................	8	1	7y1
...............................................gcugcgaucuUuugaaag................................................	1	1	7y1
...............................................gcugcgaucuauugaGag................................................	2	1	7y1
...............................................Acugcgaucuauugaaag................................................	1	1	7y1
...............................................gcugcgaucuaCugaaag................................................	1	1	7y1
...............................................Ncugcgaucuauugaaag................................................	2	1	7y1
...............................................gcugAgaucuauugaaag................................................	6	1	7y1
...............................................gcuAcgaucuauugaaag................................................	1	1	7y1
...............................................gcugcgaucuauugaaaU................................................	1	1	7y1
...............................................gcugcgaucuauugaaUg................................................	1	1	7y1
...............................................gcAgcgaucuauugaaagu...............................................	2	1	7y1
...............................................gcugcgaucuauugaaagu...............................................	111	0	7y1
...............................................gcugcgaucuauugaaagA...............................................	1	1	7y1
...............................................gcugcgaAcuauugaaagu...............................................	1	1	7y1
...............................................gAugcgaucuauugaaagu...............................................	2	1	7y1
...............................................gcugcgaucuauAgaaagu...............................................	1	1	7y1
...............................................gcugcgaucuauugaaagAc..............................................	2	1	7y1
...............................................gcugAgaucuauugaaaguc..............................................	1	1	7y1
...............................................gcugcgaAcuauugaaaguc..............................................	1	1	7y1
...............................................gcugcgaucuauugaaaguU..............................................	1	1	7y1
...............................................gcugcgaucuauAgaaaguc..............................................	1	1	7y1
...............................................gcuAcgaucuauugaaaguc..............................................	1	1	7y1
...............................................gAugcgaucuauugaaaguc..............................................	1	1	7y1
...............................................gcugcgaucuauugaaaUuc..............................................	1	1	7y1
...............................................Ucugcgaucuauugaaaguc..............................................	2	1	7y1
...............................................gcugcgaucuaAugaaaguc..............................................	1	1	7y1
...............................................gcugcgaucGauugaaaguc..............................................	1	1	7y1
```

**Star** **Mature**

```
gaccugcuucugggucgggguuucguacguagcagagcagcucccucgcugcgaucuauugaaagucagcccucgacacaaggguuuguccgcgcgcgcgcgcgcgcgugcgu
.....................................................aucuauugaaagucagcAcu........................................ 2 1 7y1
.....................................................aucuauuCaaagucagcccu........................................ 1 1 7y1
.....................................................aucuauugaaagucaUcccu........................................ 1 1 7y1
.....................................................auAuauugaaagucagcccu........................................ 2 1 7y1
.....................................................aucuauugaaagucagcccG........................................ 2 1 7y1
.....................................................aucuaAugaaagucagcccu........................................ 4 1 7y1
.....................................................aucuauugaaagucagcccA........................................ 1 1 7y1
.....................................................aucuauugaaagucagcccu........................................ 755 0 7y1
.....................................................aucuauAgaaagucagcccu........................................ 3 1 7y1
.....................................................aAcuauugaaagucagcccu........................................ 2 1 7y1
.....................................................aucuauugaaagucagccAu........................................ 1 1 7y1
.....................................................aucCauugaaagucagcccu........................................ 223 1 7y1
.....................................................aucuauugaaaguAagcccu........................................ 1 1 7y1
.....................................................aucCauugaaagucagcccuc....................................... 1285 1 7y1
.....................................................aucAauugaaagucagcccuc....................................... 2 1 7y1
.....................................................aucuauAgaaagucagcccucg...................................... 4 1 7y1
.....................................................aucuauugaaagucagcccucg...................................... 1785 0 7y1
.....................................................aucuauugaaagucagcccucU...................................... 1 1 7y1
.....................................................aucuauugaaagucagcAcucg...................................... 2 1 7y1
.....................................................aucuauugaaagucagAccucg...................................... 6 1 7y1
.....................................................aCcuauugaaagucagcccucg...................................... 1 1 7y1
.....................................................aucuauuUaaagucagcccucg...................................... 1 1 7y1
.....................................................aucuauugaaaguAagcccucg...................................... 5 1 7y1
.....................................................aucuauugNaagucagcccucg...................................... 1 1 7y1
.....................................................aucuauugaaagAcagcccucg...................................... 2 1 7y1
.....................................................aucuaAugaaagucagcccucg...................................... 6 1 7y1
.....................................................aAcuauugaaagucagcccucg...................................... 5 1 7y1
.....................................................auAuauugaaagucagcccucg...................................... 8 1 7y1
.....................................................aucuauugaaagucagcccAcg...................................... 1 1 7y1
.....................................................aucuauugaaagucagUccucg...................................... 2 1 7y1
.....................................................aucuauugaaagucagGccucg...................................... 1 1 7y1
.....................................................aucuauugaaaUucagcccucg...................................... 1 1 7y1
.....................................................Gucuauugaaagucagcccucg...................................... 1 1 7y1
.....................................................aucuauugaUagucagcccucg...................................... 1 1 7y1
.....................................................aucuauugaaagucGgcccucg...................................... 2 1 7y1
.....................................................aucCauugaaagucagcccucg...................................... 914 1 7y1
.....................................................aucuauuCaaagucagcccucg...................................... 2 1 7y1
.....................................................aucuauugaaagucagccAucg...................................... 2 1 7y1
.....................................................aucuauugaaagucaCcccucg...................................... 1 1 7y1
.....................................................auUuauugaaagucagcccucg...................................... 249 1 7y1
.....................................................aucuauugGaagucagcccucg...................................... 1 1 7y1
.....................................................aucuauugaaagucagcccucC...................................... 2 1 7y1
.....................................................Nucuauugaaagucagcccucg...................................... 1 1 7y1
.....................................................aucuauugaaagucagcccuAg...................................... 4 1 7y1
.....................................................aucuauugaaaguGagcccucg...................................... 1 1 7y1
.....................................................aucuauugaGagucagcccucg...................................... 1 1 7y1
.....................................................aucAauugaaagucagcccucg...................................... 1 1 7y1
.....................................................aucuauugaaagucaUcccucg...................................... 1 1 7y1
.....................................................aucuauugaaagucagcAcucga..................................... 2 1 7y1
.....................................................aucuauugaaaguAagcccucga..................................... 1 1 7y1
.....................................................aucuauugaaagucagccAucga..................................... 2 1 7y1
.....................................................aucuauugaaagucagcUcucga..................................... 1 1 7y1
.....................................................aucuauugaaagucagcccucga..................................... 460 0 7y1
.....................................................auAuauugaaagucagcccucga..................................... 2 1 7y1
.....................................................aucuauugaaagucagcccuAga..................................... 1 1 7y1
.....................................................aucuauugaaagucagccGucga..................................... 1 1 7y1
.....................................................aucuauAgaaagucagcccucga..................................... 6 1 7y1
.....................................................aucuaAugaaagucagcccucga..................................... 2 1 7y1
.....................................................aAcuauugaaagucagcccucga..................................... 3 1 7y1
.....................................................aucuauCgaaagucagcccucga..................................... 1 1 7y1
.....................................................aucuauugUaagucagcccucga..................................... 1 1 7y1
.....................................................aucuauugaaagucagcccucgac.................................... 272 0 7y1
.....................................................Uucuauugaaagucagcccucgac.................................... 2 1 7y1
.....................................................aucuauAgaaagucagcccucgac.................................... 2 1 7y1
.....................................................aucuaAugaaagucagcccucgac.................................... 2 1 7y1
.....................................................aucuauugaaagucagcccucgaA.................................... 1 1 7y1
.....................................................aAcuauugaaagucagcccucgac.................................... 1 1 7y1
.....................................................aucuauugaaagucaAcccucgacacaag............................... 1 1 7y1
.....................................................aucuauugaaagucagcccucUacacaag............................... 1 1 7y1
.....................................................aucuauugaaagucagccAucgacacaag............................... 1 1 7y1
```

**Star** **Mature**

```
gaccugcuucugggucgggguuucguacguagcagagcagcucccucgcugcgaucuauugaaagucagcccucgacacaaggguuuguccgcgcgcgcgcgcgcgcgcgugcgu
.....................................................Nucuauugaaagucagcccucgacacaag................................	1	1	7y1
.....................................................aAcuauugaaagucagcccucgacacaag................................	1	1	7y1
.....................................................aucuauugaaagucagcccAcgacacaag................................	1	1	7y1
.....................................................aucuauugaaagucagcccucgCcacaag................................	1	1	7y1
.....................................................aucuaAugaaagucagcccucgacacaag................................	1	1	7y1
.....................................................aucuauugaaagucCgcccucgacacaag................................	1	1	7y1
.....................................................Uucuauugaaagucagcccucgacacaag................................	1	1	7y1
.....................................................aucuauugaaagucagcccucgacUcaag................................	1	1	7y1
.....................................................auAuauugaaagucagcccucgacacaag................................	4	1	7y1
.....................................................aucuauugaaagucagcccucgacacaag................................	494	0	7y1
.....................................................aucAauugaaagucagcccucgacacaag................................	1	1	7y1
.....................................................aucuauugaaagucagcGcucgacacaag................................	1	1	7y1
.....................................................aucuauugaaagucagcccucgacaGaag................................	1	1	7y1
.....................................................aucuauuUaaagucagcccucgacacaag................................	1	1	7y1
......................................................ucuauugaaagucagGcc..........................................	1	1	7y1
......................................................ucuauugaaagucagccA..........................................	6	1	7y1
......................................................ucuauugaaagucaUccc..........................................	2	1	7y1
......................................................ucuauugaaagucagccc..........................................	1308	0	7y1
......................................................ucuauugaaaCucagccc..........................................	1	1	7y1
......................................................ucuauugaaagucaAccc..........................................	1	1	7y1
......................................................ucuauugaaagucCgccc..........................................	1	1	7y1
......................................................ucuauuUaaagucagccc..........................................	3	1	7y1
......................................................ucuauugaaaguAagccc..........................................	3	1	7y1
......................................................ucuauuAaaagucagccc..........................................	2	1	7y1
......................................................ucuaAugaaagucagccc..........................................	3	1	7y1
......................................................Gcuauugaaagucagccc..........................................	2	1	7y1
......................................................ucuauugaaagucagcAc..........................................	1	1	7y1
......................................................ucuGuugaaagucagccc..........................................	1	1	7y1
......................................................ucuauugaaagucagUcc..........................................	4	1	7y1
......................................................ucuauAgaaagucagccc..........................................	3	1	7y1
......................................................ucuauugaGagucagccc..........................................	227	1	7y1
......................................................uAuauugaaagucagccc..........................................	5	1	7y1
......................................................ucuauugaaagucagccG..........................................	1	1	7y1
......................................................ucAauugaaagucagccc..........................................	3	1	7y1
......................................................Acuauugaaagucagccc..........................................	16	1	7y1
......................................................ucuauugaaagucagccAu.........................................	5	1	7y1
......................................................ucuauugaaagGcagcccu.........................................	1	1	7y1
......................................................ucuaAugaaagucagcccu.........................................	1	1	7y1
......................................................ucuauugaaagucagcAcu.........................................	3	1	7y1
......................................................Ncuauugaaagucagcccu.........................................	1	1	7y1
......................................................ucuauugaaagucCgcccu.........................................	1	1	7y1
......................................................ucGauugaaagucagcccu.........................................	1	1	7y1
......................................................Acuauugaaagucagcccu.........................................	6	1	7y1
......................................................ucuauAgaaagucagcccu.........................................	3	1	7y1
......................................................ucuauugaaagucagcccG.........................................	1	1	7y1
......................................................ucAauugaaagucagcccu.........................................	1	1	7y1
......................................................ucuauugaaagucagUccu.........................................	1	1	7y1
......................................................ucuauugaaagucagcccA.........................................	1	1	7y1
......................................................ucuauuAaaagucagcccu.........................................	1	1	7y1
......................................................ucuauugaaagucagcccu.........................................	861	0	7y1
......................................................ucuauugaaaCucagcccu.........................................	1	1	7y1
......................................................ucAauugaaagucagcccuc........................................	1	1	7y1
......................................................ucCauugaaagucagcccuc........................................	278	1	7y1
......................................................ucuauugaaagucCgcccucg.......................................	1	1	7y1
......................................................ucAauugaaagucagcccucg.......................................	1	1	7y1
......................................................ucuauugaaagucagccGucg.......................................	1	1	7y1
......................................................ucuauAgaaagucagcccucg.......................................	3	1	7y1
......................................................ucuauCgaaagucagcccucg.......................................	1	1	7y1
......................................................ucuauugaaaUucagcccucg.......................................	1	1	7y1
......................................................ucuauugaaagucagUccucg.......................................	1	1	7y1
......................................................Gcuauugaaagucagcccucg.......................................	1	1	7y1
......................................................ucCauugaaagucagcccucg.......................................	282	1	7y1
......................................................ucuaAugaaagucagcccucg.......................................	2	1	7y1
......................................................uAuauugaaagucagcccucg.......................................	1	1	7y1
......................................................Ccuauugaaagucagcccucg.......................................	1	1	7y1
......................................................ucuauugaaagucagcccucU.......................................	1	1	7y1
......................................................ucuauugaaaguAagcccucg.......................................	2	1	7y1
......................................................ucuauugaaagucagcAcucg.......................................	3	1	7y1
......................................................ucuauugaaagucaCcccucg.......................................	1	1	7y1
......................................................ucuauugaaagucagcUcucg.......................................	2	1	7y1
```

```
       Star                                                                    Mature
gaccugcuucugggucggggguuucguacguagcagagcagcucccucgcugcgaucuauugaaagucagcccucgacacaaggguuuguccgcgcgcgcgcgcgcgcgugcgu
......................................................Acuauugaaagucagcccucg......................................    9       1    7y1
......................................................ucuauugaaagucagccAucg......................................    2       1    7y1
......................................................ucuauugaaagucagcccucA......................................    1       1    7y1
......................................................ucuauugaaagucagcccucg......................................    1220    0    7y1
......................................................ucuauugaaagucagcUcucga.....................................    1       1    7y1
......................................................Gcuauugaaagucagcccucga.....................................    2       1    7y1
......................................................ucuauugaaGgucagcccucga.....................................    1       1    7y1
......................................................ucAauugaaagucagcccucga.....................................    1       1    7y1
......................................................uAuauugaaagucagcccucga.....................................    3       1    7y1
......................................................Acuauugaaagucagcccucga.....................................    3       1    7y1
......................................................ucuaCugaaagucagcccucga.....................................    1       1    7y1
......................................................ucuaAugaaagucagcccucga.....................................    1       1    7y1
......................................................ucuauuUaaagucagcccucga.....................................    1       1    7y1
......................................................ucuauugaaagucagcccucga.....................................    428     0    7y1
......................................................ucuauugaaagucagcccucgG.....................................    2       1    7y1
......................................................ucuauugaaaguAagcccucga.....................................    1       1    7y1
......................................................ucuauAgaaagucagcccucga.....................................    2       1    7y1
......................................................Gcuauugaaagucagcccucgac....................................    1       1    7y1
......................................................ucuauugaUagucagcccucgac....................................    1       1    7y1
......................................................uAuauugaaagucagcccucgac....................................    1       1    7y1
......................................................ucuauugaaagucagcccucgac....................................    299     0    7y1
......................................................ucuauugaaagucagccAucgac....................................    1       1    7y1
......................................................ucAauugaaagucagcccucgac....................................    1       1    7y1
......................................................ucuauugaaagucagcccAcgac....................................    1       1    7y1
......................................................Acuauugaaagucagcccucgac....................................    2       1    7y1
.......................................................cuauCgaaagucagcccu........................................    1       1    7y1
.......................................................cuauugaaagucagcccG........................................    1       1    7y1
.......................................................cuauugaaagucagcccu........................................    416     0    7y1
.......................................................cuaAugaaagucagcccu........................................    1       1    7y1
.......................................................cAauugaaagucagcccu........................................    2       1    7y1
.......................................................cuauugaaaAucagcccu........................................    1       1    7y1
.......................................................Auauugaaagucagcccu........................................    1       1    7y1
.......................................................cuauugaaaguAagcccu........................................    1       1    7y1
.......................................................cuauugaaaCucagcccu........................................    1       1    7y1
.......................................................cuauugaaagAcagcccuc.......................................    2       1    7y1
.......................................................cuauugaaagucagcccuc.......................................    186     0    7y1
.......................................................cCauugaaagucagcccuc.......................................    488     1    7y1
.......................................................cuauugaUagucagcccuc.......................................    1       1    7y1
.......................................................cAauugaaagucagcccuc.......................................    5       1    7y1
.......................................................Guauugaaagucagcccuc.......................................    1       1    7y1
.......................................................cCauugaaagucagcccucg......................................    701     1    7y1
.......................................................cAauugaaagucagcccucg......................................    4       1    7y1
.......................................................cAauugaaagucagcccucgac....................................    1       1    7y1
.......................................................cuauugaaagucagcccucgac....................................    273     0    7y1
.......................................................cuauugaaagucagcAcucgac....................................    1       1    7y1
.......................................................cuauugaaagucGgcccucgac....................................    1       1    7y1
.......................................................cuauugaaagucagccAucgac....................................    3       1    7y1
.......................................................Auauugaaagucagcccucgac....................................    1       1    7y1
.......................................................cuauugaaagAcagcccucgac....................................    1       1    7y1
.......................................................cuauugaaagucagcccucgacacaaggguuu..........................    1       0    7y1
........................................................uauugaaaUucagcccucg......................................    1       1    7y1
........................................................Aauugaaagucagcccucg......................................    6       1    7y1
........................................................uauugaaagucagcccucC......................................    1       1    7y1
........................................................uauugaaagucagcGcucg......................................    1       1    7y1
........................................................uauugaaagucagcccAcg......................................    2       1    7y1
........................................................uauugaaagucagcccucg......................................    490     0    7y1
........................................................uauugaaGgucagcccucg......................................    1       1    7y1
........................................................uauugaaagucagcccuAg......................................    1       1    7y1
........................................................uauugaaagAcagcccucg......................................    1       1    7y1
........................................................uauugCaagucagcccucga.....................................    1       1    7y1
........................................................uauugaaagucagcccucgG.....................................    2       1    7y1
........................................................uauugaaagucaCcccucga.....................................    1       1    7y1
........................................................uauugaaUgucagcccucga.....................................    1       1    7y1
........................................................Aauugaaagucagcccucga.....................................    2       1    7y1
........................................................Gauugaaagucagcccucga.....................................    1       1    7y1
........................................................uauugaaagucagcccucga.....................................    197     0    7y1
........................................................uauugaaagucagcccucgac....................................    549     0    7y1
........................................................uaAugaaagucagcccucgac....................................    4       1    7y1
........................................................uauugaaagAcagcccucgac....................................    1       1    7y1
........................................................uauugaaaUucagcccucgac....................................    1       1    7y1
```

**Star** **Mature**

```
gaccugcuucugggucggggguucguacguagcagagcagcucccucgcugcgaucuauugaaagucagcccucgacacaaggguuuguccgcgcgcgcgcgcgcgcgugcgu
........................................................uauugaaagucagcAcucgac....................................    1    1    7y1
........................................................uauAgaaagucagcccucgac....................................    2    1    7y1
........................................................Aauugaaagucagcccucgac....................................    11   1    7y1
........................................................uauugaaagucagcccucUac....................................    1    1    7y1
........................................................uauugaaagucagcccCcgac....................................    1    1    7y1
........................................................uauugaaagucaCcccucgac....................................    1    1    7y1
........................................................uauugaaagucaUcccucgac....................................    1    1    7y1
........................................................uauugaaagucagAccucgac....................................    1    1    7y1
........................................................uauugaaagucagcccucgaGacaaggguu...........................    1    1    7y1
........................................................uauuCaaagucagcccucgacacaaggguu...........................    1    1    7y1
........................................................uauugaaaguAagcccucgacacaaggguu...........................    1    1    7y1
........................................................uauugaaagucagcAcucgacacaaggguu...........................    2    1    7y1
........................................................uauugaaagucagcccucgacacaaggAuu...........................    1    1    7y1
........................................................Aauugaaagucagcccucgacacaaggguu...........................    1    1    7y1
........................................................uauugaaagucagcccucgacacaaggguu...........................    155  0    7y1
........................................................uauugaaagucagcccAcgacacaaggguu...........................    1    1    7y1
........................................................uauugaaaUucagcccucgacacaaggguu...........................    1    1    7y1
........................................................uauugaaagucagcccucgacacaaggCuu...........................    1    1    7y1
........................................................uauugaaagucagcccucgacacaaggguG...........................    1    1    7y1
........................................................uauugaaagucagcccucgacaUaaggguuu..........................    1    1    7y1
........................................................uauugaaagucagcccucgacacaagggAuu..........................    1    1    7y1
........................................................uauugaaagucagAccucgacacaaggguuu..........................    1    1    7y1
........................................................uauugaaagucagcccucgacacaaggguuu..........................    220  0    7y1
........................................................uauugaaagucagcccAcgacacaaggguuu..........................    1    1    7y1
........................................................Aauugaaagucagcccucgacacaaggguuu..........................    7    1    7y1
........................................................uauugaaagucagccAucgacacaaggguuug.........................    1    1    7y1
........................................................uauugaaagucagcccuAgacacaaggguuug.........................    3    1    7y1
........................................................uauugaaagucagcccucgacacaaCgguuug.........................    1    1    7y1
........................................................uauugaaagucagcccucgacacaagggAuug.........................    2    1    7y1
........................................................uauugaaagucagcccucgacaAaaggguuug.........................    3    1    7y1
........................................................Nauugaaagucagcccucgacacaaggguuug.........................    1    1    7y1
........................................................uauugaaagucagcccucgacacaagggGuug.........................    1    1    7y1
........................................................Gauugaaagucagcccucgacacaaggguuug.........................    2    1    7y1
........................................................uauugaaaguAagcccucgacacaaggguuug.........................    2    1    7y1
........................................................uauugaaagucagcccucgacacaaggguuAg.........................    1    1    7y1
........................................................uauugaaagucagcccucgacacaagCguuug.........................    1    1    7y1
........................................................uauugaaagucagGccucgacacaaggguuug.........................    1    1    7y1
........................................................uauugaaagucGgcccucgacacaaggguuug.........................    1    1    7y1
........................................................uauuCaaagucagcccucgacacaaggguuug.........................    1    1    7y1
........................................................uauugaaagucagcccCcgacacaaggguuug.........................    1    1    7y1
........................................................uauugaaagucagcccAcgacacaaggguuug.........................    2    1    7y1
........................................................uauugaaagucagcGcucgacacaaggguuug.........................    1    1    7y1
........................................................uauugaaagucaCcccucgacacaaggguuug.........................    2    1    7y1
........................................................uauugaaagucagcccucgacacaagUguuug.........................    1    1    7y1
........................................................uauugaaagucagcAcucgacacaaggguuug.........................    6    1    7y1
........................................................uauugaaagucagAccucgacacaaggguuug.........................    1    1    7y1
........................................................uauugaaaUucagcccucgacacaaggguuug.........................    1    1    7y1
........................................................uauugaaagucagcccucgacacaaggguuug.........................    684  0    7y1
........................................................uauuAaaagucagcccucgacacaaggguuug.........................    1    1    7y1
........................................................Aauugaaagucagcccucgacacaaggguuug.........................    13   1    7y1
.........................................................auugaaagucagccAucg......................................    1    1    7y1
.........................................................auuUaaagucagcccucg......................................    1    1    7y1
.........................................................auAgaaagucagcccucg......................................    1    1    7y1
.........................................................auuCaaagucagcccucg......................................    1    1    7y1
.........................................................auugaaagucagcccucg......................................    487  0    7y1
.........................................................aAugaaagucagcccucg......................................    2    1    7y1
.........................................................auugaGagucagcccucg......................................    1    1    7y1
.........................................................auugaaaguAagcccucg......................................    2    1    7y1
.........................................................auugaaagucagUccucg......................................    1    1    7y1
.........................................................Uuugaaagucagcccucg......................................    1    1    7y1
.........................................................auugaaagucagcAcucg......................................    2    1    7y1
.........................................................auugaaagucagAccucg......................................    3    1    7y1
.........................................................auugaaagucagcccucC......................................    1    1    7y1
.........................................................auugaaagucagcccuAga.....................................    2    1    7y1
.........................................................auugaaagucaUcccucga.....................................    1    1    7y1
.........................................................auugaaagucagcccucgC.....................................    1    1    7y1
.........................................................auugaaagucagAccucga.....................................    1    1    7y1
.........................................................auugaaagucagcccAcga.....................................    1    1    7y1
.........................................................auugaaagucagccAucga.....................................    1    1    7y1
.........................................................auugaaagAcagcccucga.....................................    1    1    7y1
```

**Star** **Mature**

```
gaccugcuucugggucgggguuucguacguagcagagcagcucccucgcugcgaucuauugaaagucagcccucgacacaaggguuuguccgcgcgcgcgcgcgcgcgugcgu
..........................................................uugaaagucagcccucgacacaagCguuugu......................    1     1    7y1
..........................................................uugaaagucagcAcucgacacaaggguuugu......................    1     1    7y1
..........................................................Gugaaagucagcccucgacacaaggguuugu......................    3     1    7y1
..........................................................uugaaagucagccAucgacacaaggguuugu......................    3     1    7y1
..........................................................uugaaagucagcccucgacacaaggguuugu......................    1439  0    7y1
..........................................................uugaaagucagcccuAgacacaaggguuugu......................    2     1    7y1
..........................................................uugaaagucagcccucgacacaagUguuugu......................    1     1    7y1
..........................................................Augaaagucagcccucgacacaaggguuugu......................    10    1    7y1
..........................................................uugaaagucagcccucgacacaaggguuuUu......................    2     1    7y1
..........................................................uugaaagucagcccucgacacaaggCuuugu......................    1     1    7y1
..........................................................uugaaagucagGccucgacacaaggguuugu......................    1     1    7y1
..........................................................uugaaagucagcccucgacacaaUgguuugu......................    1     1    7y1
..........................................................uugaaagucagccUucgacacaaggguuugu......................    1     1    7y1
..........................................................uugaaagucagcccucgacacaaggguAugu......................    4     1    7y1
..........................................................uugaaagucGgcccucgacacaaggguuugu......................    1     1    7y1
..........................................................uuCaaagucagcccucgacacaaggguuugu......................    2     1    7y1
..........................................................uugaaagucagcccucgacacaaggguGugu......................    1     1    7y1
..........................................................uugaaaCucagcccucgacacaaggguuugu......................    1     1    7y1
..........................................................uugaaaguAagcccucgacacaaggguuugu......................    4     1    7y1
..........................................................uugaaagucagcccucgacacaaggguuugA......................    4     1    7y1
..........................................................uAgaaagucagcccucgacacaaggguuugu......................    2     1    7y1
..........................................................uugaaagucagcccucgacacaagggGuugu......................    3     1    7y1
..........................................................uugaaagucagcccucgacacaaCgguuugu......................    2     1    7y1
..........................................................uCgaaagucagcccucgacacaaggguuugu......................    2     1    7y1
...........................................................Agaaagucagcccucgac..................................    4     1    7y1
...........................................................ugaaagAcagcccucgac..................................    1     1    7y1
...........................................................ugaaaguAagcccucgac..................................    2     1    7y1
...........................................................ugaaagucagcccucgac..................................    191   0    7y1
...........................................................ugaaagucagUccucgac..................................    1     1    7y1
...........................................................ugaaagucagcccucgCc..................................    1     1    7y1
...........................................................ugaaagucagcccucCacacaaggguuug.......................    1     1    7y1
...........................................................ugaaagucagcccucgacacaagggAuug.......................    2     1    7y1
...........................................................ugaaagucagcccGcgacacaaggguuug.......................    1     1    7y1
...........................................................ugaaagAcagcccucgacacaaggguuug.......................    1     1    7y1
...........................................................ugaaagucagccGucgacacaaggguuug.......................    1     1    7y1
...........................................................ugaaagucagcccucgacacaaggUuuug.......................    1     1    7y1
...........................................................ugaaagucagcccucgCcacaaggguuug.......................    1     1    7y1
...........................................................ugaaagucagcccucgacacaaUgguuug.......................    1     1    7y1
...........................................................uAaaagucagcccucgacacaaggguuug.......................    1     1    7y1
...........................................................ugaaagucagcccucgacacGaggguuug.......................    1     1    7y1
...........................................................ugaaagucagcccucgacacaaggguuug.......................    462   0    7y1
...........................................................ugaaagucagAccucgacacaaggguuug.......................    1     1    7y1
...........................................................ugaaagucagcccucgacacaaggguAug.......................    1     1    7y1
...........................................................ugaaagucagcccucgacacaaggguGug.......................    1     1    7y1
...........................................................Ggaaagucagcccucgacacaaggguuug.......................    1     1    7y1
...........................................................ugaaagucUgcccucgacacaaggguuug.......................    1     1    7y1
...........................................................ugaaagucagcccucgacacaaggguuGg.......................    2     1    7y1
............................................................gaaagucagcccucgacaUaaggguuugu......................    1     1    7y1
............................................................gaaagucagcccucgacCcaaggguuugu......................    1     1    7y1
............................................................gaaagucagAccucgacacaaggguuugu......................    1     1    7y1
............................................................gaaagucagcccAcgacacaaggguuugu......................    1     1    7y1
............................................................gaaagucagcccucgacacaaggguuAgu......................    1     1    7y1
............................................................Caaagucagcccucgacacaaggguuugu......................    1     1    7y1
............................................................gaaagucagcccucgacacaaggguuugu......................    170   0    7y1
............................................................gaGagucagcccucgacacaaggguuugu......................    1     1    7y1
............................................................gaaagucagcccucgacacaagggGuugu......................    1     1    7y1
..............................................................aagucagcccucgacacaaggguuuC.......................    2     1    7y1
..............................................................aagucagcccucgacacaaggguuug.......................    211   0    7y1
..............................................................aagucagcccucgacUcaaggguuug.......................    2     1    7y1
..............................................................aagucagccAucgacacaaggguuug.......................    2     1    7y1
..............................................................aagucagcccucgacacaaggUuuug.......................    1     1    7y1
.................................................................uAagcccucgacacaaggguuug.......................    1     1    7y1
.................................................................ucagcccucgGcacaaggguuug.......................    1     1    7y1
.................................................................Acagcccucgacacaaggguuug.......................    3     1    7y1
.................................................................Ncagcccucgacacaaggguuug.......................    1     1    7y1
.................................................................ucagccAucgacacaaggguuug.......................    1     1    7y1
.................................................................ucagcccucgacacaaggguuug.......................    244   0    7y1
.................................................................ucagcccucgacacaagggAuug.......................    1     1    7y1
.................................................................Gcagcccucgacacaaggguuug.......................    1     1    7y1
..................................................................cagcUcucgacacaaggguuugu......................    211   1    7y1
```

```
Star                                                                  Mature
gaccugcuucugggucgggguuucguacguagcagagcagcucccucgcugcgaucuauugaaagucagcccucgacacaaggguuuguccgcgcgcgcgcgcgcgcgugcgu
...................................................................agccUucgacacaaggguuug.........................   132    1   7y1
....................................................................gcAcucgacacaaggguuug.........................   1      1   7y1
....................................................................gccAucgacacaaggguuug.........................   1      1   7y1
....................................................................Ncccucgacacaaggguuug.........................   1      1   7y1
....................................................................gAccucgacacaaggguuug.........................   2      1   7y1
....................................................................gcccucgaAacaaggguuug.........................   2      1   7y1
....................................................................gcccucgacacaaggguuug.........................   188    0   7y1
....................................................................gcccucAacacaaggguuug.........................   1      1   7y1
.....................................................................cccucgacacaaggguuuU.........................   1      1   7y1
.....................................................................ccAucgacacaaggguuugu........................   2      1   7y1
.....................................................................Nccucgacacaaggguuugu........................   1      1   7y1
.....................................................................cccucgacacaaggguuugu........................   289    0   7y1
.....................................................................cccucgacacaagggAuugu........................   2      1   7y1
.....................................................................cccucgacacaaggCuuugu........................   2      1   7y1
.....................................................................cccucgacaAaaggguuugu........................   1      1   7y1
.....................................................................cccucgacUcaaggguuugu........................   1      1   7y1
.....................................................................cccucgacacaaggguuugG........................   2      1   7y1
.....................................................................cccucgacacaaUgguuugu........................   1      1   7y1
......................................................................Gcucgacacaaggguuug.........................   4      1   7y1
......................................................................ccucgacacaagAguuug.........................   1      1   7y1
......................................................................ccucgacaUaaggguuug.........................   1      1   7y1
......................................................................ccucUacacaaggguuug.........................   3      1   7y1
......................................................................cAucgacacaaggguuug.........................   6      1   7y1
......................................................................ccucgacUcaaggguuug.........................   2      1   7y1
......................................................................ccCcgacacaaggguuug.........................   1      1   7y1
......................................................................ccucgacacaaggguAug.........................   4      1   7y1
......................................................................ccucgacacaaggguuGg.........................   1      1   7y1
......................................................................ccucgaAacaaggguuug.........................   3      1   7y1
......................................................................ccAcgacacaaggguuug.........................   1      1   7y1
......................................................................ccucgacacaagCguuug.........................   2      1   7y1
......................................................................ccucgacacaaUgguuug.........................   1      1   7y1
......................................................................Ncucgacacaaggguuug.........................   2      1   7y1
......................................................................ccuGgacacaaggguuug.........................   1      1   7y1
......................................................................Acucgacacaaggguuug.........................   8      1   7y1
......................................................................ccucgacacaaggguuAg.........................   1      1   7y1
......................................................................ccucgacacaaggguuuA.........................   1      1   7y1
......................................................................ccucgacacaaggguuuC.........................   1      1   7y1
......................................................................ccucgacacaUggguuug.........................   1      1   7y1
......................................................................ccucgacacaaggCuuug.........................   1      1   7y1
......................................................................ccucgacacaaggguuug.........................   1954   0   7y1
......................................................................ccucgCcacaaggguuug.........................   2      1   7y1
......................................................................ccucgacacGaggguuug.........................   1      1   7y1
......................................................................ccucgacacaaUgguuugu........................   5      1   7y1
......................................................................ccucgacacNaggguuugu........................   1      1   7y1
......................................................................ccucgacacaaggguAugu........................   8      1   7y1
......................................................................ccucgacacaGggguuugu........................   1      1   7y1
......................................................................ccucUacacaaggguuugu........................   7      1   7y1
......................................................................ccucgacacaaggguuuAu........................   1      1   7y1
......................................................................ccucgacacaaggguuugA........................   12     1   7y1
......................................................................ccucgacacaagUguuugu........................   6      1   7y1
......................................................................ccucgUcacaaggguuugu........................   1      1   7y1
......................................................................ccucgacaUaaggguuugu........................   153    1   7y1
......................................................................Ucucgacacaaggguuugu........................   358    1   7y1
......................................................................Acucgacacaaggguuugu........................   24     1   7y1
......................................................................ccucgacacaaggguuuCu........................   3      1   7y1
......................................................................ccucgacGcaaggguuugu........................   3      1   7y1
......................................................................ccucgacacaaggUuuugu........................   9      1   7y1
......................................................................cAucgacacaaggguuugu........................   17     1   7y1
......................................................................ccucgacacaaCgguuugu........................   3      1   7y1
......................................................................ccucgacacaaggCuuugu........................   13     1   7y1
......................................................................ccucgCcacaaggguuugu........................   7      1   7y1
......................................................................ccucgacacaaggguuugG........................   17     1   7y1
......................................................................ccuGgacacaaggguuugu........................   1      1   7y1
......................................................................Ncucgacacaaggguuugu........................   3      1   7y1
......................................................................ccucgacacaagCguuugu........................   4      1   7y1
......................................................................ccucgaUacaaggguuugu........................   1      1   7y1
......................................................................ccucgacacaaggguuugC........................   1      1   7y1
......................................................................ccAcgacacaaggguuugu........................   14     1   7y1
......................................................................ccucgaAacaaggguuugu........................   9      1   7y1
......................................................................ccucgacaAaaggguuugu........................   8      1   7y1
```

```
  Star                                                                 Mature
gaccugcuucugggucgggguuucguacguagcagagcagcucccucgcugcgaucuauugaaagucagcccucgacacaaggguuuguccgcgcgcgcgcgcgcgcgcgugcgu
.....................................................................ccucgacacGaggguuugu..........................    1      1    7y1
.....................................................................ccucgacacaaggguuugu..........................    6344   0    7y1
.....................................................................ccucgacacaaggguuuUu..........................    5      1    7y1
.....................................................................ccucgacacaagggGuugu..........................    4      1    7y1
.....................................................................ccucgGcacaaggguuugu..........................    1      1    7y1
.....................................................................Gcucgacacaaggguuugu..........................    5      1    7y1
.....................................................................ccucAacacaaggguuugu..........................    4      1    7y1
.....................................................................ccucgacacaagggAuugu..........................    8      1    7y1
.....................................................................ccucgacacaaggguuAgu..........................    9      1    7y1
.....................................................................ccucgacacaaggguCugu..........................    1      1    7y1
.....................................................................cNucgacacaaggguuuguccgcgc....................    1      1    7y1
.....................................................................ccucgacacaaggguuuguccgcgA....................    1      1    7y1
.....................................................................ccAcgacacaaggguuuguccgcgc....................    1      1    7y1
.....................................................................ccucgacacaaggguuuguccgcgc....................    277    0    7y1
.....................................................................ccucgacacaaggguAuguccgcgc....................    1      1    7y1
.....................................................................ccucgacacaaggCuuuguccgcgc....................    1      1    7y1
.....................................................................ccucgacacaaggUuuuguccgcgc....................    1      1    7y1
.....................................................................ccucgacacaagggAuuguccgcgc....................    1      1    7y1
.....................................................................ccuUgacacaaggguuuguccgcgc....................    1      1    7y1
.....................................................................cAucgacacaaggguuuguccgcgc....................    1      1    7y1
......................................................................cucgacacaaggguuAgu..........................    30     1    7y1
......................................................................cucgacacGaggguuugu..........................    5      1    7y1
......................................................................cucgacacaagggGuugu..........................    22     1    7y1
......................................................................cucgacacaaggguCugu..........................    2      1    7y1
......................................................................Gucgacacaaggguuugu..........................    18     1    7y1
......................................................................cNcgacacaaggguuugu..........................    2      1    7y1
......................................................................cucAacacaaggguuugu..........................    13     1    7y1
......................................................................Nucgacacaaggguuugu..........................    10     1    7y1
......................................................................cucgacUcaaggguuugu..........................    9      1    7y1
......................................................................cucgacacaaggguuugA..........................    39     1    7y1
......................................................................cucUacacaaggguuugu..........................    14     1    7y1
......................................................................cucgacacaaggguuCgu..........................    1      1    7y1
......................................................................cucgacacaaggguuuUu..........................    19     1    7y1
......................................................................cucgaUacaaggguuugu..........................    3      1    7y1
......................................................................cucgacacaaggguuugu..........................    25979  0    7y1
......................................................................cuGgacacaaggguuugu..........................    197    1    7y1
......................................................................cucgacacaaggCuuugu..........................    17     1    7y1
......................................................................cucgacacaagCguuugu..........................    18     1    7y1
......................................................................cucgacaUaaggguuugu..........................    3      1    7y1
......................................................................cucgacacaagUguuugu..........................    175    1    7y1
......................................................................cucgacacaaggguuugC..........................    1      1    7y1
......................................................................cucgacacaaCgguuugu..........................    17     1    7y1
......................................................................cucgacGcaaggguuugu..........................    6      1    7y1
......................................................................cucgacacaGggguuugu..........................    1      1    7y1
......................................................................cucgacacaaAgguuugu..........................    197    1    7y1
......................................................................cucgacacaaggguuugG..........................    34     1    7y1
......................................................................cucgacacaaggguuGgu..........................    1      1    7y1
......................................................................cucgacacaaUgguuugu..........................    17     1    7y1
......................................................................cucgaAacaaggguuugu..........................    34     1    7y1
......................................................................cucgacCcaaggguuugu..........................    1      1    7y1
......................................................................cucgacacCaggguuugu..........................    1      1    7y1
......................................................................cAcgacacaaggguuugu..........................    75     1    7y1
......................................................................cucgacacaaggguuuCu..........................    23     1    7y1
......................................................................cucgaNacaaggguuugu..........................    1      1    7y1
......................................................................cGcgacacaaggguuugu..........................    2      1    7y1
......................................................................cucgacacaaggguuuAu..........................    7      1    7y1
......................................................................cucgacacaaggguGugu..........................    5      1    7y1
......................................................................cucgacacaaggguAugu..........................    35     1    7y1
......................................................................cCcgacacaaggguuugu..........................    2      1    7y1
......................................................................cucgacacaagggAuugu..........................    43     1    7y1
......................................................................cucgacacaaggUuuugu..........................    13     1    7y1
......................................................................cucgacacaaggAuuugu..........................    3      1    7y1
......................................................................cucgacacUaggguuugu..........................    2      1    7y1
......................................................................cucgaGacaaggguuugu..........................    2      1    7y1
......................................................................cucgCcacaaggguuugu..........................    4      1    7y1
......................................................................cucgacaAaaggguuugu..........................    21     1    7y1
......................................................................cucgacacaUggguuugu..........................    2      1    7y1
......................................................................cucgacaGaaggguuugu..........................    1      1    7y1
......................................................................cucgUcacaaggguuugu..........................    2      1    7y1
......................................................................cucgacacaaggguuuguccgcgc....................    1      0    7y1
```

Star Mature

gaccugcuucugggucggggguuucguacguagcagagcagcucccucgcugcgaucuauugaaagucagcccucgacacaaggguuuguccgcgcgcgcgcgcgcgcgugcgu
