## Supplementary material for "SEA: The small RNA Expression Atlas": p-hsa-miR-235-3

| | | |
|---|---|---|
| Provisional ID | : | chr21_12210 |
| Score total | : | 144078.6 |
| Score for star read(s) | : | 3.9 |
| Score for read counts | : | 144071.6 |
| Score for mfe | : | 2.1 |
| Score for randfold | : | 1.6 |
| Score for cons. seed | : | -0.6 |
| Total read count | : | 282602 |
| Mature read count | : | 36506 |
| Loop read count | : | 245911 |
| Star read count | : | 185 |

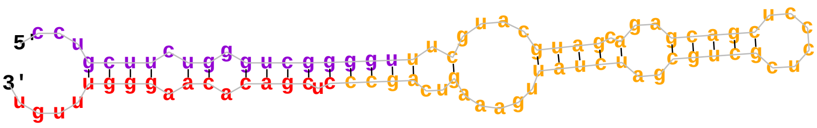

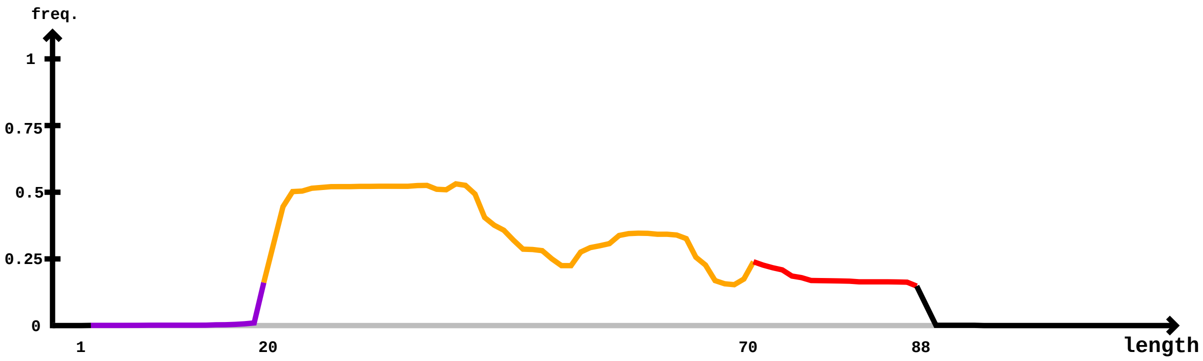


Star Mature

```
5'- gaccugcuucugggucgggguuucguacguagcagagcagcucccucgcugcgaucuauugaaagucagcccucgacacaaggguuuguccgcgcgcgcgcgcgcgugcgugc  -3'  obs
    gaccugcuucugggucgggguuucguacguagcagagcagcucccucgcugcgaucuauugaaagucagcccucgacacaaggguuuguccgcgcgcgcgcgcgcgugcgugc       exp
    (((..((((.((.(((((((((.(....((((.(..(((((......)))))..))))).....)..))))).)))).)).)))).))).((((((((((....))))))))))  reads  mm  sample
    ..ccugcuucugggucgggg.............................................................................................  7      0   7y1
    ..ccugcuucugggucggggA............................................................................................  1      1   7y1
    ..ccugcuucugggucggggG............................................................................................  6      1   7y1
    ..cAugcuucugggucggggu............................................................................................  1      1   7y1
    ..ccugcuucugggucggggu............................................................................................  164    0   7y1
    ...cugcuucuggguAgggguuucgu.......................................................................................  2      1   7y1
    ...cugcuucugggucgggguAucgu.......................................................................................  1      1   7y1
    ...cugcuucugUgucgggguuucgu.......................................................................................  1      1   7y1
    ...cugcuucugggucgggUuuucgu.......................................................................................  2      1   7y1
    ....ugcuucAgggucgggguu...........................................................................................  1      1   7y1
    .....gcuucuggCucgggguuucgu.......................................................................................  1      1   7y1
    .....gcuucugggAcgggguuucgu.......................................................................................  1      1   7y1
    .....gcuucuggguAgggguuucgu.......................................................................................  1      1   7y1
    .....gcuucuggUucgggguuucgu.......................................................................................  1      1   7y1
    ......cuucugggucgggguuuG.........................................................................................  1      1   7y1
    ......cuucugggAcgggguuuc.........................................................................................  1      1   7y1
    ......cuucugggucgggguGuc.........................................................................................  1      1   7y1
    ......cuucuAggucgggguuuc.........................................................................................  1      1   7y1
    ......cuucugggucggCguuuc.........................................................................................  1      1   7y1
    ......cuucuUggucgggguuucgu.......................................................................................  1      1   7y1
    ......cuucugggAcgggguuucgu.......................................................................................  2      1   7y1
    ......cuucuCggucgggguuucgu.......................................................................................  1      1   7y1
    ......cuucuggguAgggguuucgu.......................................................................................  4      1   7y1
    ......cuucugggucCggguuucgu.......................................................................................  1      1   7y1
    ......cuucugggucggCguuucgu.......................................................................................  1      1   7y1
    ......cuucugggucUggguuucgu.......................................................................................  1      1   7y1
    ......cuucugggucggggGuucgu.......................................................................................  2      1   7y1
    ......cuucugggucgggAuuucgu.......................................................................................  1      1   7y1
    ......cuucugggucgggUuuucgu.......................................................................................  1      1   7y1
    ......cuucugggucgggguuucUu.......................................................................................  3      1   7y1
    ......cuucugggucgggguCucgu.......................................................................................  1      1   7y1
    ......cuucugggucgggguAucgu.......................................................................................  1      1   7y1
    ......cuucugggucggggAuucgu.......................................................................................  1      1   7y1
```

```
  Star                                                                     Mature
gaccugcuucugggucgggguuucguacguagcagagcagcucccucgcugcgaucuauugaaagucagcccucgacacaaggguuuguccgcgcgcgcgcgcgcgcgugcgugc
......cuucuggguGgggguuucgu.........................................................................................   1   1   7y1
......cuucugggucggUguuucgu.........................................................................................   1   1   7y1
.......uucugggAcgggguuucgu.........................................................................................   3   1   7y1
.......uucugggucgUgguuucgu.........................................................................................   2   1   7y1
.......uucuggguAgggguuucgu.........................................................................................   6   1   7y1
.......uucugCgucgggguuucgu.........................................................................................   4   1   7y1
.......uucugggucUggguuucgu.........................................................................................   2   1   7y1
.......uucugggucggggCuucgu.........................................................................................   1   1   7y1
.......uucugggucggCguuucgu.........................................................................................   1   1   7y1
.......uucuUggucgggguuucgu.........................................................................................   2   1   7y1
.......uucugggucggUguuucgu.........................................................................................   1   1   7y1
.......uucugggucgggCuuucgu.........................................................................................   4   1   7y1
.......uucugggucgggguAucgu.........................................................................................   3   1   7y1
.......uucugggucggggGuucgu.........................................................................................   1   1   7y1
.......uucuggUucgggguuucgu.........................................................................................   2   1   7y1
.......uucugggucgggguCucgu.........................................................................................   2   1   7y1
.......uucugUgucgggguuucgu.........................................................................................   1   1   7y1
........ucuggCucgggguuucgu.........................................................................................   4   1   7y1
........ucugggucgggCuuucgu.........................................................................................   1   1   7y1
........ucugCgucgggguuucgu.........................................................................................   1   1   7y1
........ucugggucgUgguuucgu.........................................................................................   1   1   7y1
........ucuggguAgggguuucgu.........................................................................................   4   1   7y1
........ucugggucgCgguuucgu.........................................................................................   8   1   7y1
........ucuggAucgggguuucgu.........................................................................................   1   1   7y1
........ucugggucggggAuucgu.........................................................................................   1   1   7y1
........ucugggucggCguuucgu.........................................................................................   6   1   7y1
........ucugggucgggguAucgu.........................................................................................   4   1   7y1
........ucugggucCggguuucgu.........................................................................................   2   1   7y1
........ucugggucgggguCucgu.........................................................................................   1   1   7y1
........ucugUgucgggguuucgu.........................................................................................   2   1   7y1
........ucugggucgggUuuucgu.........................................................................................   1   1   7y1
........ucuAggucgggguuucgu.........................................................................................   2   1   7y1
........ucuggUucgggguuucgu.........................................................................................   7   1   7y1
........ucugggucggUguuucgu.........................................................................................   3   1   7y1
........ucugggucUggguuucgu.........................................................................................   2   1   7y1
........ucugggAcgggguuucgu.........................................................................................   4   1   7y1
........ucuggguGgggguuucgu.........................................................................................   1   1   7y1
........ucugggucgggguuAcgu.........................................................................................   1   1   7y1
........ucugggucgggUuuucgua.......................................................................................   2   1   7y1
........ucugggucggggAuucgua.......................................................................................   1   1   7y1
........ucugggucgggguuAcgua.......................................................................................   1   1   7y1
........ucugggucUggguuucgua.......................................................................................   3   1   7y1
........ucuggguAgggguuucgua.......................................................................................   4   1   7y1
........ucugggucgggCuuucgua.......................................................................................   1   1   7y1
........ucugggucgUgguuucgua.......................................................................................   3   1   7y1
........ucuggguGgggguuucgua.......................................................................................   1   1   7y1
........ucugggucgggguuGcgua.......................................................................................   1   1   7y1
........ucugggucggUguuucgua.......................................................................................   4   1   7y1
........ucugggucggCguuucgua.......................................................................................   2   1   7y1
........ucugUgucgggguuucgua.......................................................................................   2   1   7y1
........ucugggucgCgguuucgua.......................................................................................   6   1   7y1
........ucuggAucgggguuucgua.......................................................................................   1   1   7y1
........ucugggAcgggguuucgua.......................................................................................   12  1   7y1
........ucugggucggAguuucgua.......................................................................................   1   1   7y1
........ucugggucCggguuucgua.......................................................................................   2   1   7y1
........ucugCgucgggguuucgua.......................................................................................   2   1   7y1
........ucuggUucgggguuucgua.......................................................................................   5   1   7y1
........ucugggucgggguAucgua.......................................................................................   2   1   7y1
.........cugggucgggguuuAgua.......................................................................................   5   1   7y1
.........cugggucgggUuuucgua.......................................................................................   1   1   7y1
.........cugggucgCgguuucgua.......................................................................................   1   1   7y1
.........cugggucgAgguuucgua.......................................................................................   1   1   7y1
.........cugggucgggguuuGgua.......................................................................................   1   1   7y1
.........cugCgucgggguuucgua.......................................................................................   1   1   7y1
.........cuggguAgggguuucgua.......................................................................................   5   1   7y1
.........cugggucUggguuucgua.......................................................................................   1   1   7y1
.........cugggucggUguuucgua.......................................................................................   1   1   7y1
.........cugggucggggGuucgua.......................................................................................   1   1   7y1
.........cugggucgggguuAcguac......................................................................................   1   1   7y1
.........cugggucgggguuuGguac......................................................................................   1   1   7y1
```

```
 Star                                                         Mature
gaccugcuucugggucgggguuucguacguagcagagcagcucccucgcugcgaucuauugaaagucagcccucgacacaaggguuuguccgcgcgcgcgcgcgcgugcgugc
.........cugggucgggguuuAguac.....................................................................................    1    1    7y1
.........cuggguAgggguuucguac.....................................................................................    1    1    7y1
.........cugggucCggguuucguac.....................................................................................    1    1    7y1
.........cugggucggCguuucguac.....................................................................................    1    1    7y1
.........cugggAcgggguuucguac.....................................................................................    1    1    7y1
.........cugggucgggUuuucguac.....................................................................................    1    1    7y1
..............ucgggguAucguacguag.................................................................................    1    1    7y1
..............ucggggAuucguacguag.................................................................................    1    1    7y1
..............ucggggAuucguacguagc................................................................................    1    1    7y1
..............ucgggUuuucguacguagc................................................................................    1    1    7y1
..............ucggggCuucguacguagc................................................................................    1    1    7y1
..............ucgggguAucguacguagc................................................................................    1    1    7y1
..............ucggggGuucguacguagc................................................................................    1    1    7y1
..............ucgggguuucguaUguagc................................................................................    1    1    7y1
..............ucgggguuucAuacguagcagagc...........................................................................    2    1    7y1
..............ucgggguGucguacguagcagagcagcuc......................................................................    1    1    7y1
..............ucgggguCucguacguagcagagcagcuc......................................................................    1    1    7y1
..............ucgggCuuucguacguagcagagcagcuc......................................................................    6    1    7y1
..............ucgggUuuucguacguagcagagcagcuc......................................................................    4    1    7y1
..............ucggggCuucguacguagcagagcagcuc......................................................................    2    1    7y1
..............ucggggAuucguacguagcagagcagcuc......................................................................    23   1    7y1
...............cgggguuucgCacguagc................................................................................    1    1    7y1
...............cggAguuucguacguagc................................................................................    1    1    7y1
...............cggggAuucguacguagc................................................................................    2    1    7y1
...............cgggguuucguaAguagc................................................................................    1    1    7y1
...............cgggguuAcguacguagc................................................................................    1    1    7y1
...............cgggguuucUuacguagc................................................................................    1    1    7y1
...............cggCguuucguacguagc................................................................................    1    1    7y1
...............cggggu Aucguacguagc...............................................................................    3    1    7y1
...............cgggguuucguacAuagca...............................................................................    1    1    7y1
...............cggggAuucguacguagca...............................................................................    2    1    7y1
...............cgggguuucUuacguagca...............................................................................    1    1    7y1
...............cgggguuucguaAguagca...............................................................................    1    1    7y1
...............cgggguuucCuacguagcag..............................................................................    1    1    7y1
...............cgggguuucguacCuagcag..............................................................................    1    1    7y1
...............Agggguuucguacguagcag..............................................................................    1    1    7y1
...............cgggguuucguacgAagcag..............................................................................    1    1    7y1
...............cggggAuucguacguagcag..............................................................................    1    1    7y1
...............cgggguuuAguacguagcag..............................................................................    1    1    7y1
...............cgggUuuucguacguagcag..............................................................................    1    1    7y1
...............cgggguGucguacguagcaga.............................................................................    1    1    7y1
...............cgggguuAcguacguagcaga.............................................................................    2    1    7y1
...............cgggguuuAguacguagcaga.............................................................................    1    1    7y1
...............cgggguuucUuacguagcaga.............................................................................    1    1    7y1
...............cgggguuucCuacguagcaga.............................................................................    1    1    7y1
...............cgggguuucguaGguagcaga.............................................................................    1    1    7y1
...............cgggguuucguacUuagcaga.............................................................................    4    1    7y1
...............cgggCuuucguacguagcaga.............................................................................    3    1    7y1
...............cgggguuucguaAguagcaga.............................................................................    3    1    7y1
...............cgggguuuGguacguagcaga.............................................................................    1    1    7y1
...............cgggguuucguacgCagcaga.............................................................................    1    1    7y1
...............cggggu Aucguacguagcaga............................................................................    4    1    7y1
...............cggggAuucguacguagcaga.............................................................................    5    1    7y1
...............cgggguuuUguacguagcaga.............................................................................    1    1    7y1
...............cggggGuucguacguagcaga.............................................................................    1    1    7y1
...............cgggguuucAuacguagcagag............................................................................    1    1    7y1
...............cgggguuuAguacguagcagag............................................................................    1    1    7y1
...............cgggguuucguacUuagcagag............................................................................    1    1    7y1
...............cgggUuuucguacguagcagag............................................................................    1    1    7y1
...............cgggguuucUuacguagcagag............................................................................    1    1    7y1
...............cgggguAucguacguagcagag............................................................................    2    1    7y1
...............cgggguuucguaAguagcagag............................................................................    3    1    7y1
...............cgggguuucgAacguagcagag............................................................................    1    1    7y1
...............cgggguuucguacUuagcagagc...........................................................................    2    1    7y1
...............cgggCuuucguacguagcagagc...........................................................................    3    1    7y1
...............cgggguuuAguacguagcagagc...........................................................................    7    1    7y1
...............cgggguuucgGacguagcagagc...........................................................................    1    1    7y1
...............cgggguuucCuacguagcagagc...........................................................................    2    1    7y1
...............cggggGuucguacguagcagagc...........................................................................    3    1    7y1
...............cgggguuucgAacguagcagagc...........................................................................    3    1    7y1
```

**Star** **Mature**

```
gaccugcuucugggucggggguuucguacguagcagagcagcucccucgcugcgaucuauugaaagucagcccucgacacaaggguuuguccgcgcgcgcgcgcgcgugcgugc
..................gCuuucguacguagcagagcagcucc.....................................................................	1	1	7y1
..................gguuucguUcguagcagagcagcucc.....................................................................	1	1	7y1
..................gguAucguacguagcagagcagcuccc....................................................................	2	1	7y1
..................gguuucUuacguagcagagcagcuccc....................................................................	4	1	7y1
..................gguuucguacguagGagagcagcuccc....................................................................	1	1	7y1
..................gguuucguacguaCcagagcagcuccc....................................................................	3	1	7y1
..................gguuucguacguaUcagagcagcuccc....................................................................	1	1	7y1
..................gguuucguacgAagcagagcagcuccc....................................................................	2	1	7y1
..................gguuuUguacguagcagagcagcuccc....................................................................	1	1	7y1
..................gguuucCuacguagcagagcagcuccc....................................................................	3	1	7y1
..................gguuucguacUuagcagagcagcuccc....................................................................	1	1	7y1
..................ggGuucguacguagcagagcagcuccc....................................................................	3	1	7y1
..................gguuucgGacguagcagagcagcuccc....................................................................	2	1	7y1
..................gguuucguacguagAagagcagcuccc....................................................................	5	1	7y1
..................gguuucguaGguagcagagcagcuccc....................................................................	2	1	7y1
..................ggCuucguacguagcagagcagcuccc....................................................................	1	1	7y1
..................gCuuucguacguagcagagcagcuccc....................................................................	1	1	7y1
..................gguuucguacgGagcagagcagcuccc....................................................................	1	1	7y1
..................gguuuAguacguagcagagcagcuccc....................................................................	15	1	7y1
..................ggAuucguacguagcagagcagcuccc....................................................................	3	1	7y1
..................gguuucguaAguagcagagcagcuccc....................................................................	11	1	7y1
..................gguuucguacCuagcagagcagcuccc....................................................................	4	1	7y1
..................gguuucgAacguagcagagcagcuccc....................................................................	2	1	7y1
..................gguuucguacAuagcagagcagcuccc....................................................................	1	1	7y1
..................gUuuucguacguagcagagcagcuccc....................................................................	2	1	7y1
..................gguuGcguacguagcagagcagcuccc....................................................................	1	1	7y1
..................gguuucguacgCagcagagcagcuccc....................................................................	1	1	7y1
..................gguuucguacguCgcagagcagcucccu...................................................................	1	1	7y1
..................gguuucguacgAagcagagcagcucccu...................................................................	5	1	7y1
..................gguuucguacCuagcagagcagcucccu...................................................................	2	1	7y1
..................gguuucguacguGgcagagcagcucccu...................................................................	2	1	7y1
..................gguuucguacguagAagagcagcucccu...................................................................	5	1	7y1
..................gguuucguacUuagcagagcagcucccu...................................................................	1	1	7y1
..................gguuuAguacguagcagagcagcucccu...................................................................	9	1	7y1
..................ggAuucguacguagcagagcagcucccu...................................................................	4	1	7y1
..................gCuuucguacguagcagagcagcucccu...................................................................	2	1	7y1
..................ggCuucguacguagcagagcagcucccu...................................................................	1	1	7y1
..................gguuucguacguagGagagcagcucccu...................................................................	1	1	7y1
..................gguuucguacguaUcagagcagcucccu...................................................................	3	1	7y1
..................ggGuucguacguagcagagcagcucccu...................................................................	2	1	7y1
..................gguuucguGcguagcagagcagcucccu...................................................................	1	1	7y1
..................gguuucguaAguagcagagcagcucccu...................................................................	12	1	7y1
..................gguuAcguacguagcagagcagcucccu...................................................................	2	1	7y1
..................gAuuucguacguagcagagcagcucccu...................................................................	1	1	7y1
..................gguuucgAacguagcagagcagcucccu...................................................................	7	1	7y1
..................gguuucgCacguagcagagcagcucccu...................................................................	1	1	7y1
..................gguuuUguacguagcagagcagcucccu...................................................................	1	1	7y1
..................gguuucguacguaAcagagcagcucccu...................................................................	2	1	7y1
..................gguuucguCcguagcagagcagcucccu...................................................................	2	1	7y1
..................gguuucUuacguagcagagcagcucccu...................................................................	5	1	7y1
..................gguuucguaUguagcagagcagcucccu...................................................................	1	1	7y1
..................gguuucAuacguagcagagcagcucccu...................................................................	1	1	7y1
..................gUuuucguacguagcagagcagcucccu...................................................................	1	1	7y1
..................gguAucguacguagcagagcagcucccu...................................................................	3	1	7y1
..................gguuucUuacguagcagagcagcucccuc..................................................................	1	1	7y1
..................gguuAcguacguagcagagcagcucccuc..................................................................	1	1	7y1
..................gguuucguGcguagcagagcagcucccuc..................................................................	1	1	7y1
..................gguuuGguacguagcagagcagcucccuc..................................................................	1	1	7y1
..................gguuuAguacguagcagagcagcucccuc..................................................................	1	1	7y1
..................gguuucguacAuagcagagcagcucccuc..................................................................	2	1	7y1
..................gguuucguaAguagcagagcagcucccuc..................................................................	1	1	7y1
..................ggCuucguacguagcagagcagcucccuc..................................................................	1	1	7y1
..................gguuucgAacguagcagagcagcucccuc..................................................................	2	1	7y1
..................ggAuucguacguagcagagcagcucccuc..................................................................	2	1	7y1
..................gguuucguacUuagcagagcagcucccuc..................................................................	2	1	7y1
..................gguuucguacguagAagagcagcucccuc..................................................................	1	1	7y1
..................gguuucCuacguagcagagcagcucccuc..................................................................	2	1	7y1
..................gguuucguacgAagcagagcagcucccuc..................................................................	1	1	7y1
..................gguuucguacguaCcagagcagcucccuc..................................................................	1	1	7y1
..................gguuucguacguaUcagagcagcucccuc..................................................................	1	1	7y1
```

```
   Star                                                               Mature
gaccugcuucugggucggggguuucguacguagcagagcagcagcucccucgcugcgaucuauugaaagucagcccucgacacaaggguuuguccgcgcgcgcgcgcgcgugcgugc
..................gguuucguaGguagcagagcagcucccuc..................................................................... 1 1 7y1
..................gguuucgGacguagcagagcagcucccuc..................................................................... 1 1 7y1
..................gguuucguacguGgcagagcagcucccuc..................................................................... 1 1 7y1
..................gCuuucguacguagcagagcagcucccuc..................................................................... 1 1 7y1
..................gguuucguacguaCcagagcagcucccucg.................................................................... 1 1 7y1
..................gguuucguacgAagcagagcagcucccucg.................................................................... 3 1 7y1
..................gguuAcguacguagcagagcagcucccucg.................................................................... 1 1 7y1
..................ggAuucguacguagcagagcagcucccucg.................................................................... 3 1 7y1
..................gguuucguUcguagcagagcagcucccucg.................................................................... 1 1 7y1
..................gguuucgGacguagcagagcagcucccucg.................................................................... 1 1 7y1
..................gguuucguacguagUagagcagcucccucg.................................................................... 1 1 7y1
..................gguuuAguacguagcagagcagcucccucg.................................................................... 3 1 7y1
..................gCuuucguacguagcagagcagcucccucg.................................................................... 1 1 7y1
..................gguuucguacguagAagagcagcucccucg.................................................................... 2 1 7y1
..................gguuucCuacguagcagagcagcucccucg.................................................................... 2 1 7y1
..................gguuucguacguagGagagcagcucccucg.................................................................... 1 1 7y1
..................gguuucguaAguagcagagcagcucccucg.................................................................... 1 1 7y1
..................gguuucguaUguagcagagcagcucccucg.................................................................... 2 1 7y1
..................gguAucguacguagcagagcagcucccucg.................................................................... 2 1 7y1
..................gguuucguacguaAcagagcagcucccucgc................................................................... 1 1 7y1
..................gguCucguacguagcagagcagcucccucgc................................................................... 1 1 7y1
..................gCuuucguacguagcagagcagcucccucgc................................................................... 1 1 7y1
..................gguuucAuacguagcagagcagcucccucgc................................................................... 1 1 7y1
..................ggGuucguacguagcagagcagcucccucgc................................................................... 2 1 7y1
..................gguuucguacUuagcagagcagcucccucgc................................................................... 1 1 7y1
..................gguuucguacguagUagagcagcucccucgc................................................................... 1 1 7y1
..................gguuucguacgAagcagagcagcucccucgc................................................................... 2 1 7y1
..................gguuuAguacguagcagagcagcucccucgc................................................................... 1 1 7y1
..................gguuucguacguGgcagagcagcucccucgc................................................................... 1 1 7y1
..................gguGucguacguagcagagcagcucccucgc................................................................... 1 1 7y1
..................ggAuucguacguagcagagcagcucccucgc................................................................... 2 1 7y1
..................gguAucguacguagcagagcagcucccucgc................................................................... 1 1 7y1
..................gguuucCuacguagcagagcagcucccucgc................................................................... 4 1 7y1
..................gUuuucguacguagcagagcagcucccucgc................................................................... 2 1 7y1
..................gguuucgAacguagcagagcagcucccucgc................................................................... 4 1 7y1
..................gguuCcguacguagcagagcagcucccucgc................................................................... 1 1 7y1
..................gguuucguaAguagcagagcagcucccucgc................................................................... 5 1 7y1
..................gguuucguacguagAagagcagcucccucgcu.................................................................. 1 1 7y1
..................gguuucUuacguagcagagcagcucccucgcu.................................................................. 1 1 7y1
..................ggAuucguacguagcagagcagcucccucgcu.................................................................. 1 1 7y1
..................gguuucguacCuagcagagcagcucccucgcu.................................................................. 1 1 7y1
..................gguuucguGcguagcagagcagcucccucgcu.................................................................. 1 1 7y1
..................gguuucguacguaUcagagcagcucccucgcu.................................................................. 1 1 7y1
..................gguuucguacguaCcagagcagcucccucgcu.................................................................. 1 1 7y1
..................gUuuucguacguagcagagcagcucccucgcu.................................................................. 1 1 7y1
..................gguuucAuacguagcagagcagcucccucgcu.................................................................. 1 1 7y1
...................guuGcguacguagcagagc.............................................................................. 1 1 7y1
...................guuucguaAguagcagagc.............................................................................. 5 1 7y1
...................Nuuucguacguagcagagc.............................................................................. 1 1 7y1
...................Uuuucguacguagcagagc.............................................................................. 1 1 7y1
...................guuuAguacguagcagagc.............................................................................. 1 1 7y1
...................guuucCuacguagcagagc.............................................................................. 1 1 7y1
...................Cuuucguacguagcagagc.............................................................................. 3 1 7y1
...................guuucgAacguagcagagc.............................................................................. 3 1 7y1
...................guuucUuacguagcagagc.............................................................................. 1 1 7y1
...................guuAcguacguagcagagca............................................................................. 1 1 7y1
...................guuuAguacguagcagagca............................................................................. 1 1 7y1
...................guuucguacguagAagagca............................................................................. 2 1 7y1
...................gAuucguacguagcagagca............................................................................. 1 1 7y1
...................guAucguacguagcagagca............................................................................. 1 1 7y1
...................guuCcguacguagcagagca............................................................................. 1 1 7y1
...................guuucCuacguagcagagca............................................................................. 2 1 7y1
...................guuucguacCuagcagagca............................................................................. 2 1 7y1
...................Uuuucguacguagcagagca............................................................................. 1 1 7y1
...................guuucgAacguagcagagca............................................................................. 5 1 7y1
...................Cuuucguacguagcagagca............................................................................. 2 1 7y1
...................guuucguacgAagcagagcagc........................................................................... 1 1 7y1
...................Auuucguacguagcagagcagc........................................................................... 1 1 7y1
...................guuucguaAguagcagagcagc........................................................................... 2 1 7y1
...................gAuucguacguagcagagcagc........................................................................... 5 1 7y1
```

```
Star                                                                   Mature
gaccugcuucugggucgggguuucguacguagcagagcagcucccucgcugcgaucuauugaaagucagcccucgacacaaggguuuguccgcgcgcgcgcgcgcgugcgugc
...................guAucguacguagcagagcagc........................................................................ 3    1  7y1
...................Uuuucguacguagcagagcagc........................................................................ 286  1  7y1
...................guuucgAacguagcagagcagc........................................................................ 2    1  7y1
...................guuucguUcguagcagagcagc........................................................................ 1    1  7y1
...................guAucguacguagcagagcagcu....................................................................... 4    1  7y1
...................guuucguacgAagcagagcagcu....................................................................... 3    1  7y1
...................Uuuucguacguagcagagcagcu....................................................................... 3    1  7y1
...................gGuucguacguagcagagcagcu....................................................................... 1    1  7y1
...................Cuuucguacguagcagagcagcu....................................................................... 2    1  7y1
...................guuucgAacguagcagagcagcu....................................................................... 3    1  7y1
...................guuucguacUuagcagagcagcu....................................................................... 1    1  7y1
...................Nuuucguacguagcagagcagcu....................................................................... 3    1  7y1
...................gAuucguacguagcagagcagcu....................................................................... 6    1  7y1
...................guuucguacguGgcagagcagcu....................................................................... 1    1  7y1
...................guuAcguacguagcagagcagcu....................................................................... 2    1  7y1
...................guuucguacguagAagagcagcu....................................................................... 1    1  7y1
...................guuucUuacguagcagagcagcu....................................................................... 1    1  7y1
...................guuuAguacguagcagagcagcu....................................................................... 1    1  7y1
...................gAuucguacguagcagagcagcuc...................................................................... 2    1  7y1
...................guuuUguacguagcagagcagcuc...................................................................... 1    1  7y1
...................guAucguacguagcagagcagcuc...................................................................... 3    1  7y1
...................guCucguacguagcagagcagcuc...................................................................... 1    1  7y1
...................gGuucguacguagcagagcagcuc...................................................................... 1    1  7y1
...................Uuuucguacguagcagagcagcucc..................................................................... 1    1  7y1
...................guuucguacgAagcagagcagcucc..................................................................... 1    1  7y1
...................guAucguacguagcagagcagcucc..................................................................... 1    1  7y1
...................guuucguacguaAcagagcagcucc..................................................................... 1    1  7y1
...................guuucguacguaUcagagcagcucc..................................................................... 179  1  7y1
...................guuucguGcguagcagagcagcucc..................................................................... 1    1  7y1
...................gAuucguacguagcagagcagcuccc.................................................................... 4    1  7y1
...................guuucguacguagAagagcagcuccc.................................................................... 5    1  7y1
...................guuucguacguCgcagagcagcuccc.................................................................... 1    1  7y1
...................guAucguacguagcagagcagcuccc.................................................................... 2    1  7y1
...................Cuuucguacguagcagagcagcuccc.................................................................... 2    1  7y1
...................guuucguacCuagcagagcagcuccc.................................................................... 3    1  7y1
...................guuucguaAguagcagagcagcuccc.................................................................... 2    1  7y1
...................guuAcguacguagcagagcagcuccc.................................................................... 1    1  7y1
...................guuucgAacguagcagagcagcuccc.................................................................... 2    1  7y1
...................guuucCuacguagcagagcagcuccc.................................................................... 2    1  7y1
...................gCuucguacguagcagagcagcuccc.................................................................... 2    1  7y1
...................guuucguacguaAcagagcagcuccc.................................................................... 2    1  7y1
...................guuucguacgGagcagagcagcuccc.................................................................... 1    1  7y1
...................guuucguacgAagcagagcagcuccc.................................................................... 1    1  7y1
...................guuuAguacguagcagagcagcuccc.................................................................... 3    1  7y1
...................guuuUguacguagcagagcagcuccc.................................................................... 1    1  7y1
...................guuucguacguaCcagagcagcuccc.................................................................... 3    1  7y1
...................guuucguacguaUcagagcagcuccc.................................................................... 1    1  7y1
...................guuuAguacguagcagagcagcucccu................................................................... 1    1  7y1
...................guuAcguacguagcagagcagcucccu................................................................... 2    1  7y1
...................guuucCuacguagcagagcagcucccu................................................................... 1    1  7y1
...................guuucguacgGagcagagcagcucccu................................................................... 1    1  7y1
...................guuucguacAuagcagagcagcucccu................................................................... 2    1  7y1
...................gGuucguacguagcagagcagcucccu................................................................... 295  1  7y1
...................guuucguacgCagcagagcagcucccu................................................................... 1    1  7y1
...................guAucguacguagcagagcagcucccu................................................................... 2    1  7y1
...................guuucguacguagcGgagcagcucccu................................................................... 2    1  7y1
...................Cuuucguacguagcagagcagcucccu................................................................... 3    1  7y1
...................guuucguacCuagcagagcagcucccu................................................................... 1    1  7y1
...................guuucgAacguagcagagcagcucccu................................................................... 7    1  7y1
...................gAuucguacguagcagagcagcucccu................................................................... 7    1  7y1
...................guuucguacguagAagagcagcucccu................................................................... 3    1  7y1
...................Uuuucguacguagcagagcagcucccu................................................................... 2    1  7y1
...................guuucguacguagcGgagcagcucccuc.................................................................. 1    1  7y1
...................guuucguGcguagcagagcagcucccuc.................................................................. 2    1  7y1
...................gGuucguacguagcagagcagcucccuc.................................................................. 1    1  7y1
...................guuucguacgAagcagagcagcucccuc.................................................................. 1    1  7y1
...................guuucguaAguagcagagcagcucccuc.................................................................. 1    1  7y1
...................guuucguacUuagcagagcagcucccuc.................................................................. 3    1  7y1
...................guuucgAacguagcagagcagcucccuc.................................................................. 2    1  7y1
...................guuucguacAuagcagagcagcucccuc.................................................................. 1    1  7y1
```

```
  Star                                                                 Mature
gaccugcuucugggucgggguuucguacguagcagagcagcucccucgcugcgaucuauugaaagucagcccucgacacaaggguuuguccgcgcgcgcgcgcgcgugcgugc
...................guuCcguacguagcagagcagcucccuc..................................................................   1      1    7y1
...................guAucguacguagcagagcagcucccuc..................................................................   1      1    7y1
...................Cuuucguacguagcagagcagcucccuc..................................................................   1      1    7y1
...................guuucguacguagAagagcagcucccuc..................................................................   2      1    7y1
...................guuucguacguaUcagagcagcucccuc..................................................................   1      1    7y1
...................guuucguacCuagcagagcagcucccuc..................................................................   1      1    7y1
...................Uuuucguacguagcagagcagcucccuc..................................................................   3      1    7y1
...................guuucguacguaUcagagcagcucccucg.................................................................   1      1    7y1
...................guuucgAacguagcagagcagcucccucg.................................................................   4      1    7y1
...................gAuucguacguagcagagcagcucccucg.................................................................   1      1    7y1
...................guAucguacguagcagagcagcucccucg.................................................................   1      1    7y1
...................guuucguacAuagcagagcagcucccucg.................................................................   1      1    7y1
...................guuucguacguagcGgagcagcucccucg.................................................................   1      1    7y1
...................guuucguacgCagcagagcagcucccucgc................................................................   1      1    7y1
...................guuucCuacguagcagagcagcucccucgc................................................................   1      1    7y1
...................guuucguacCuagcagagcagcucccucgc................................................................   1      1    7y1
...................guuucguacUuagcagagcagcucccucgc................................................................   2      1    7y1
...................Uuuucguacguagcagagcagcucccucgc................................................................   1      1    7y1
...................guuucguaAguagcagagcagcucccucgc................................................................   3      1    7y1
...................gAuucguacguagcagagcagcucccucgc................................................................   1      1    7y1
...................guuucgGacguagcagagcagcucccucgc................................................................   1      1    7y1
...................gGuucguacguagcagagcagcucccucgc................................................................   1      1    7y1
...................guuucguacguaUcagagcagcucccucgc................................................................   2      1    7y1
...................Nuuucguacguagcagagcagcucccucgc................................................................   1      1    7y1
...................guAucguacguagcagagcagcucccucgc................................................................   2      1    7y1
...................guuucguacgAagcagagcagcucccucgc................................................................   1      1    7y1
...................guuucgAacguagcagagcagcucccucgc................................................................   4      1    7y1
...................guuAcguacguagcagagcagcucccucgc................................................................   2      1    7y1
...................Cuuucguacguagcagagcagcucccucgc................................................................   1      1    7y1
...................guuucUuacguagcagagcagcucccucgc................................................................   1      1    7y1
...................guuucguacguagAagagcagcucccucgc................................................................   1      1    7y1
...................guuucguacguagAagagcagcucccucgcu...............................................................   1      1    7y1
...................guuucguaGguagcagagcagcucccucgcu...............................................................   1      1    7y1
...................guuucUuacguagcagagcagcucccucgcu...............................................................   1      1    7y1
...................gAuucguacguagcagagcagcucccucgcu...............................................................   2      1    7y1
...................Cuuucguacguagcagagcagcucccucgcu...............................................................   2      1    7y1
...................gAuucguacguagcagagcagcucccucgcug..............................................................   1      1    7y1
...................guuucguacguagAagagcagcucccucgcug..............................................................   1      1    7y1
...................guuucguacUuagcagagcagcucccucgcug..............................................................   1      1    7y1
....................uuucguacgCagcagagc...........................................................................   1      1    7y1
....................uuucguacguagcagaUc...........................................................................   5      1    7y1
....................Auucguacguagcagagc...........................................................................   52     1    7y1
....................uuAcguacguagcagagc...........................................................................   2      1    7y1
....................uuucCuacguagcagagc...........................................................................   192    1    7y1
....................uuucguacguaCcagagc...........................................................................   1      1    7y1
....................uuucguacgAagcagagc...........................................................................   11     1    7y1
....................uuucguacguagcGgagc...........................................................................   2      1    7y1
....................uuucgGacguagcagagc...........................................................................   1      1    7y1
....................Guucguacguagcagagc...........................................................................   9      1    7y1
....................uuucguacguaUcagagc...........................................................................   2      1    7y1
....................Nuucguacguagcagagc...........................................................................   3      1    7y1
....................uuucguacguagcagagc...........................................................................   4344   0    7y1
....................uuucAuacguagcagagc...........................................................................   1      1    7y1
....................uuucguacguagcUgagc...........................................................................   2      1    7y1
....................uuucUuacguagcagagc...........................................................................   1      1    7y1
....................uuucgAacguagcagagc...........................................................................   8      1    7y1
....................Cuucguacguagcagagc...........................................................................   2      1    7y1
....................uuucguacCuagcagagc...........................................................................   4      1    7y1
....................uuucguacguagcaUagc...........................................................................   1      1    7y1
....................uAucguacguagcagagc...........................................................................   9      1    7y1
....................uuuAguacguagcagagc...........................................................................   5      1    7y1
....................uuucguacguagcagaCc...........................................................................   2      1    7y1
....................uuucguacUuagcagagc...........................................................................   1      1    7y1
....................uuucguacguGgcagagc...........................................................................   2      1    7y1
....................uuucguaUguagcagagc...........................................................................   1      1    7y1
....................uuucguacguagcagagA...........................................................................   7      1    7y1
....................uuucguacguagAagagc...........................................................................   4      1    7y1
....................uuuGguacguagcagagc...........................................................................   3      1    7y1
....................uuucguaAguagcagagc...........................................................................   8      1    7y1
....................uuucguaGguagcagagca..........................................................................   1      1    7y1
```

```
 Star                                                                   Mature
gaccugcuucugggucggggguuucguacguagcagagcagcucccucgcugcgaucuauugaaagucagcccucgacacaaggguuuguccgcgcgcgcgcgcgcgugcgugc
....................uuucguacguagcagagAa..........................................................................    4      1      7y1
....................uuucguacguagAagagca..........................................................................    3      1      7y1
....................uCucguacguagcagagca..........................................................................    1      1      7y1
....................uuucguacguagcagCgca..........................................................................    3      1      7y1
....................uuuAguacguagcagagca..........................................................................    2      1      7y1
....................uuucguaAguagcagagca..........................................................................    2      1      7y1
....................Nuucguacguagcagagca..........................................................................    3      1      7y1
....................Auucguacguagcagagca..........................................................................    24     1      7y1
....................uuucguacguagcagaUca..........................................................................    1      1      7y1
....................uuucguacguaCcagagca..........................................................................    1      1      7y1
....................uuucguacguagcagagca..........................................................................    1926   0      7y1
....................uuucguGcguagcagagca..........................................................................    1      1      7y1
....................uuucguacguUgcagagca..........................................................................    2      1      7y1
....................uuucguacguagcagaCca..........................................................................    3      1      7y1
....................uuucguacguagcagagGa..........................................................................    1      1      7y1
....................uuucguacguagUagagca..........................................................................    1      1      7y1
....................uuucgAacguagcagagca..........................................................................    2      1      7y1
....................uuucguacguaUcagagca..........................................................................    2      1      7y1
....................uuucguacCuagcagagca..........................................................................    2      1      7y1
....................uuucguacgGagcagagca..........................................................................    1      1      7y1
....................uuucguacguagcGgagca..........................................................................    1      1      7y1
....................uuucCuacguagcagagca..........................................................................    1      1      7y1
....................uuucguCcguagcagagca..........................................................................    1      1      7y1
....................uuucguacUuagcagagca..........................................................................    1      1      7y1
....................uuucguacgCagcagagca..........................................................................    1      1      7y1
....................uuucguacguagcUgagca..........................................................................    1      1      7y1
....................Guucguacguagcagagca..........................................................................    4      1      7y1
....................uuucgCacguagcagagca..........................................................................    1      1      7y1
....................uAucguacguagcagagca..........................................................................    5      1      7y1
....................uuucguacguagcaUagca..........................................................................    1      1      7y1
....................uuucguacguagcaUagcag.........................................................................    1      1      7y1
....................uuucguacguagcagagcaU.........................................................................    2      1      7y1
....................uuCcguacguagcagagcag.........................................................................    1      1      7y1
....................uuucguacgAagcagagcag.........................................................................    1      1      7y1
....................uuAcguacguagcagagcag.........................................................................    1      1      7y1
....................uAucguacguagcagagcag.........................................................................    4      1      7y1
....................uuucguacguagcagagAag.........................................................................    1      1      7y1
....................uuucguacguagcGgagcag.........................................................................    1      1      7y1
....................uuucguacguagcagaUcag.........................................................................    1      1      7y1
....................uuucguacguagcagagcaC.........................................................................    2      1      7y1
....................Auucguacguagcagagcag.........................................................................    11     1      7y1
....................uuucgCacguagcagagcag.........................................................................    1      1      7y1
....................uuucguacguagcagagcag.........................................................................    641    0      7y1
....................uuucguacguagcagagGagc........................................................................    2      1      7y1
....................uuucgAacguagcagagcagc........................................................................    3      1      7y1
....................uuucgGacguagcagagcagc........................................................................    3      1      7y1
....................uuucguGcguagcagagcagc........................................................................    1      1      7y1
....................uuucguaAguagcagagcagc........................................................................    5      1      7y1
....................uuucguaGguagcagagcagc........................................................................    1      1      7y1
....................uuucguacguagcagagcaAc........................................................................    1      1      7y1
....................uuucguaUguagcagagcagc........................................................................    1      1      7y1
....................uuucguacguagAagagcagc........................................................................    2      1      7y1
....................uuucguCcguagcagagcagc........................................................................    1      1      7y1
....................uuucguacguagcagagcagc........................................................................    2127   0      7y1
....................Cuucguacguagcagagcagc........................................................................    237    1      7y1
....................uuucgCacguagcagagcagc........................................................................    1      1      7y1
....................Auucguacguagcagagcagc........................................................................    18     1      7y1
....................uuucguacgAagcagagcagc........................................................................    2      1      7y1
....................uuucguacguagUagagcagc........................................................................    1      1      7y1
....................uuucguacguagcGgagcagc........................................................................    1      1      7y1
....................uuucguacUuagcagagcagc........................................................................    1      1      7y1
....................uAucguacguagcagagcagc........................................................................    5      1      7y1
....................Nuucguacguagcagagcagc........................................................................    1      1      7y1
....................uuucguacguagcagagcaCc........................................................................    1      1      7y1
....................uuucguacguagcagagcaUc........................................................................    1      1      7y1
....................uuucguacguagGagagcagc........................................................................    1      1      7y1
....................uuucguacguagcagagAagc........................................................................    3      1      7y1
....................uuucguacguagcaUagcagc........................................................................    1      1      7y1
....................uuucguacguagcagagcagA........................................................................    1      1      7y1
....................uuuGguacguagcagagcagc........................................................................    2      1      7y1
```

**Star** **Mature**

```
gaccugcuucugggucggggguuucguacguagcagagcagcucccucgcugcgaucuauugaaagucagcccucgacacaaggguuuguccgcgcgcgcgcgcgcgcgugcgugc
....................Guucguacguagcagagcagc..........................................................................  3  1  7y1
....................uuucguacguagcagCgcagc..........................................................................  5  1  7y1
....................uuucguacguagcUgagcagcu.........................................................................  1  1  7y1
....................uuucguacUuagcagagcagcu.........................................................................  1  1  7y1
....................uuucguacguagcagGgcagcu.........................................................................  1  1  7y1
....................uuucguacguagcaUagcagcu.........................................................................  272  1  7y1
....................uuucguacguagcagagcagAu.........................................................................  1  1  7y1
....................Nuucguacguagcagagcagcu.........................................................................  2  1  7y1
....................uuucguacguagcagagcagcC.........................................................................  1  1  7y1
....................uuucguacguagcagagGagcu.........................................................................  2  1  7y1
....................uAucguacguagcagagcagcu.........................................................................  10  1  7y1
....................uuucguacguaAcagagcagcu.........................................................................  1  1  7y1
....................uuAcguacguagcagagcagcu.........................................................................  3  1  7y1
....................uuucAuacguagcagagcagcu.........................................................................  2  1  7y1
....................uuucguaUguagcagagcagcu.........................................................................  1  1  7y1
....................uuucguacguagcagagAagcu.........................................................................  9  1  7y1
....................uuuUguacguagcagagcagcu.........................................................................  2  1  7y1
....................uuucguaAguagcagagcagcu.........................................................................  8  1  7y1
....................uuucguacAuagcagagcagcu.........................................................................  1  1  7y1
....................uuucguacguaCcagagcagcu.........................................................................  1  1  7y1
....................uuuAguacguagcagagcagcu.........................................................................  7  1  7y1
....................Cuucguacguagcagagcagcu.........................................................................  3  1  7y1
....................uuucguacguagAagagcagcu.........................................................................  2  1  7y1
....................uuucguacguagcagagcagcG.........................................................................  22  1  7y1
....................Auucguacguagcagagcagcu.........................................................................  37  1  7y1
....................uuucguacguagcagaCcagcu.........................................................................  1  1  7y1
....................uuucgAacguagcagagcagcu.........................................................................  4  1  7y1
....................uuucguacguagcagagcagcA.........................................................................  3  1  7y1
....................uuucguacCuagcagagcagcu.........................................................................  4  1  7y1
....................uuucguacguagcagCgcagcu.........................................................................  4  1  7y1
....................uuucguacguagcagagcagcu.........................................................................  4088  0  7y1
....................uuucguacguagcagagcaUcu.........................................................................  4  1  7y1
....................uuucguacguagcagagcCgcu.........................................................................  3  1  7y1
....................uuucguacguagUagagcagcu.........................................................................  1  1  7y1
....................uuucgCacguagcagagcagcu.........................................................................  1  1  7y1
....................uuucguacguagcagagcaCcu.........................................................................  1  1  7y1
....................uuucguGcguagcagagcagcu.........................................................................  1  1  7y1
....................uuucguacgAagcagagcagcu.........................................................................  11  1  7y1
....................Guucguacguagcagagcagcu.........................................................................  7  1  7y1
....................uuucguacguagcagUgcagcu.........................................................................  2  1  7y1
....................uuucguacguagcaCagcagcu.........................................................................  2  1  7y1
....................uuucCuacguagcagagcagcu.........................................................................  2  1  7y1
....................uuucguacguagcagagcagGuc........................................................................  1  1  7y1
....................uuucguacguagcagagcagcuc........................................................................  7935  0  7y1
....................uuucAuacguagcagagcagcuc........................................................................  1  1  7y1
....................uuucguacguagcagagAagcuc........................................................................  10  1  7y1
....................uuucguacguagcagagcUgcuc........................................................................  1  1  7y1
....................uuucUuacguagcagagcagcuc........................................................................  6  1  7y1
....................uuucguacguagcUgagcagcuc........................................................................  1  1  7y1
....................uuucguacguagcagUgcagcuc........................................................................  1  1  7y1
....................uuucgAacguagcagagcagcuc........................................................................  16  1  7y1
....................uuucguacguagcagagcCgcuc........................................................................  3  1  7y1
....................uuucguacguagcagagcagcuA........................................................................  8  1  7y1
....................uuucguacguagcagaUcagcuc........................................................................  3  1  7y1
....................uuucguacguagcagaCcagcuc........................................................................  4  1  7y1
....................Auucguacguagcagagcagcuc........................................................................  89  1  7y1
....................uuucguacguagcagagcGgcuc........................................................................  2  1  7y1
....................uuucguacguagcagagcaCcuc........................................................................  4  1  7y1
....................uuuUguacguagcagagcagcuc........................................................................  1  1  7y1
....................uuucguacguaCcagagcagcuc........................................................................  2  1  7y1
....................uuucguacguagcagagGagcuc........................................................................  1  1  7y1
....................uuucguacgCagcagagcagcuc........................................................................  1  1  7y1
....................uuucguacguagcagGgcagcuc........................................................................  1  1  7y1
....................uuucguacguagAagagcagcuc........................................................................  9  1  7y1
....................uuGcguacguagcagagcagcuc........................................................................  1  1  7y1
....................uuucguacguagcaCagcagcuc........................................................................  1  1  7y1
....................uuuAguacguagcagagcagcuc........................................................................  12  1  7y1
....................uuucCuacguagcagagcagcuc........................................................................  2  1  7y1
....................uuucguacCuagcagagcagcuc........................................................................  4  1  7y1
....................uuucguacguagcagagcagcGc........................................................................  1  1  7y1
```

```
                   Star                                                                  Mature
gaccugcuucugggucgggguuucguacguagcagagcagcucccucgcugcgaucuauugaaagucagcccucgacacaaggguuuguccgcgcgcgcgcgcgcgugcgugc
....................uuucguacguagcagagcagcuG......................................................................    1      1     7y1
....................uuucguacguagcCgagcagcuc......................................................................    1      1     7y1
....................uuucguacAuagcagagcagcuc......................................................................    1      1     7y1
....................Guucguacguagcagagcagcuc......................................................................    13     1     7y1
....................uuucguaAguagcagagcagcuc......................................................................    9      1     7y1
....................uuucguacguagcagagcagcAc......................................................................    2      1     7y1
....................uuCcguacguagcagagcagcuc......................................................................    1      1     7y1
....................uuucguacguagcagagcaUcuc......................................................................    7      1     7y1
....................uuucguacguGgcagagcagcuc......................................................................    2      1     7y1
....................uuucguacguagcGgagcagcuc......................................................................    1      1     7y1
....................uuucguacguagcagagcagcuU......................................................................    1      1     7y1
....................uuucgCacguagcagagcagcuc......................................................................    1      1     7y1
....................uuucguacgAagcagagcagcuc......................................................................    11     1     7y1
....................uuucguacUuagcagagcagcuc......................................................................    3      1     7y1
....................uuucguaUguagcagagcagcuc......................................................................    1      1     7y1
....................uuucguacguagcagCgcagcuc......................................................................    7      1     7y1
....................uuuGguacguagcagagcagcuc......................................................................    3      1     7y1
....................uuucguCcguagcagagcagcuc......................................................................    2      1     7y1
....................uuucguacguagcagagcagAuc......................................................................    5      1     7y1
....................Nuucguacguagcagagcagcuc......................................................................    4      1     7y1
....................uAucguacguagcagagcagcuc......................................................................    11     1     7y1
....................uuucguacguaUcagagcagcuc......................................................................    1      1     7y1
....................Cuucguacguagcagagcagcuc......................................................................    175    1     7y1
....................uuucguGcguagcagagcagcuc......................................................................    1      1     7y1
....................uuucguacguagUagagcagcuc......................................................................    2      1     7y1
....................uuucguacguagcaUagcagcuc......................................................................    2      1     7y1
....................uuAcguacguagcagagcagcuc......................................................................    3      1     7y1
....................uCucguacguagcagagcagcuc......................................................................    1      1     7y1
....................uuucguacguagcagagcagcucG.....................................................................    1      1     7y1
....................uuucguacguagcagGgcagcucc.....................................................................    4      1     7y1
....................uuucguacgGagcagagcagcucc.....................................................................    1      1     7y1
....................uuucguacguagcagagcagcuUc.....................................................................    1      1     7y1
....................uuucguaAguagcagagcagcucc.....................................................................    7      1     7y1
....................uuucguacguagcagagcagcAcc.....................................................................    1      1     7y1
....................uuucguacguagcaUagcagcucc.....................................................................    1      1     7y1
....................uuucguacCuagcagagcagcucc.....................................................................    4      1     7y1
....................uuucguacguagcaCagcagcucc.....................................................................    1      1     7y1
....................uuucguacguaUcagagcagcucc.....................................................................    3      1     7y1
....................uuucguacguagcagagcCgcucc.....................................................................    1      1     7y1
....................uuucguacguagcagagcaUcucc.....................................................................    3      1     7y1
....................uuucCuacguagcagagcagcucc.....................................................................    1      1     7y1
....................uuucguacguagcagagcagcucc.....................................................................    5981   0     7y1
....................uuucguacguagcagaCcagcucc.....................................................................    3      1     7y1
....................uuucguacguagcagagcagcucU.....................................................................    2      1     7y1
....................Auucguacguagcagagcagcucc.....................................................................    67     1     7y1
....................uuucguacguagcagagcGgcucc.....................................................................    1      1     7y1
....................uuucguacguagcagagcagcCcc.....................................................................    2      1     7y1
....................Nuucguacguagcagagcagcucc.....................................................................    3      1     7y1
....................uuuGguacguagcagagcagcucc.....................................................................    2      1     7y1
....................uuucguacUuagcagagcagcucc.....................................................................    2      1     7y1
....................uuucguacguagcagCgcagcucc.....................................................................    5      1     7y1
....................uuucguacguagcagagcagcucA.....................................................................    2      1     7y1
....................uuucguacgAagcagagcagcucc.....................................................................    7      1     7y1
....................uuucUuacguagcagagcagcucc.....................................................................    2      1     7y1
....................uuucgCacguagcagagcagcucc.....................................................................    2      1     7y1
....................uuAcguacguagcagagcagcucc.....................................................................    1      1     7y1
....................uuCcguacguagcagagcagcucc.....................................................................    2      1     7y1
....................uuGcguacguagcagagcagcucc.....................................................................    1      1     7y1
....................Guucguacguagcagagcagcucc.....................................................................    14     1     7y1
....................uuucgAacguagcagagcagcucc.....................................................................    11     1     7y1
....................uuucguacguagcagagcagAucc.....................................................................    4      1     7y1
....................uuucguacguagcagagcaCcucc.....................................................................    3      1     7y1
....................uuucguacguUgcagagcagcucc.....................................................................    1      1     7y1
....................uAucguacguagcagagcagcucc.....................................................................    9      1     7y1
....................uuucguacgCagcagagcagcucc.....................................................................    1      1     7y1
....................uuucguacguagAagagcagcucc.....................................................................    9      1     7y1
....................uuucguacguagcGgagcagcucc.....................................................................    1      1     7y1
....................uuucguacguagcagagAagcucc.....................................................................    7      1     7y1
....................uuucguaGguagcagagcagcucc.....................................................................    2      1     7y1
....................uuucguaUguagcagagcagcucc.....................................................................    1      1     7y1
```

**Star** **Mature**

```
gaccugcuucugggucgggguuucguacguagcagagcagcucccucgcugcgaucuauugaaagucagcccucgacacaaggguuuguccgcgcgcgcgcgcgcgugcgugc
....................uuucguacguagcagagcagcuAc.....................................................................  15    1  7y1
....................Cuucguacguagcagagcagcucc.....................................................................  1     1  7y1
....................uuucguUcguagcagagcagcucc.....................................................................  2     1  7y1
....................uuuAguacguagcagagcagcucc.....................................................................  6     1  7y1
....................uNucguacguagcagagcagcucc.....................................................................  1     1  7y1
....................uuucguacguagcagaUcagcucc.....................................................................  1     1  7y1
....................uuuUguacguagcagagcagcucc.....................................................................  1     1  7y1
....................uuucguacguagcagagcaAcucc.....................................................................  1     1  7y1
....................uuucguacguagcagUgcagcucc.....................................................................  3     1  7y1
....................uuucguacguagcagagcagAuccc....................................................................  1     1  7y1
....................uuucUuacguagcagagcagcuccc....................................................................  1     1  7y1
....................uuucguacguagcagagcagGuccc....................................................................  1     1  7y1
....................uuucguacCuagcagagcagcuccc....................................................................  1     1  7y1
....................uuucguacguagcagUgcagcuccc....................................................................  2     1  7y1
....................uuucguacguagcagagcagcuccA....................................................................  4     1  7y1
....................uuucguacguagUagagcagcuccc....................................................................  277   1  7y1
....................uuuGguacguagcagagcagcuccc....................................................................  3     1  7y1
....................uuucguacguagcagaCcagcuccc....................................................................  1     1  7y1
....................uuucguacguagcagagcagcAccc....................................................................  1     1  7y1
....................uuucguacguagcGgagcagcuccc....................................................................  1     1  7y1
....................uuucguaAguagcagagcagcuccc....................................................................  3     1  7y1
....................uuucAuacguagcagagcagcuccc....................................................................  1     1  7y1
....................Auucguacguagcagagcagcuccc....................................................................  17    1  7y1
....................uuucguacguagcagCgcagcuccc....................................................................  4     1  7y1
....................uuucguacguagcagagcaUcuccc....................................................................  3     1  7y1
....................uuucCuacguagcagagcagcuccc....................................................................  1     1  7y1
....................uAucguacguagcagagcagcuccc....................................................................  3     1  7y1
....................uuucguacguagcagagcagcuccG....................................................................  1     1  7y1
....................uuucguacUuagcagagcagcuccc....................................................................  1     1  7y1
....................uuucgCacguagcagagcagcuccc....................................................................  1     1  7y1
....................uuucguacguagcagagcagcuAcc....................................................................  11    1  7y1
....................uuucguacguagcagagAagcuccc....................................................................  2     1  7y1
....................uuucguacguagcagagcaCcuccc....................................................................  2     1  7y1
....................Guucguacguagcagagcagcuccc....................................................................  2     1  7y1
....................uuucguaGguagcagagcagcuccc....................................................................  1     1  7y1
....................uuucguacguagAagagcagcuccc....................................................................  2     1  7y1
....................uuucguacguagcagagcagcuccc....................................................................  2072  0  7y1
....................uuucguacgAagcagagcagcuccc....................................................................  3     1  7y1
....................uuucguacguaUcagagcagcuccc....................................................................  1     1  7y1
....................uuuAguacguagcagagcagcuccc....................................................................  2     1  7y1
....................uuAcguacguagcagagcagcuccc....................................................................  2     1  7y1
....................uuucguacguagGagagcagcuccc....................................................................  1     1  7y1
....................uuucgAacguagcagagcagcuccc....................................................................  4     1  7y1
....................uuucguacguaCcagagcagcucccu...................................................................  1     1  7y1
....................uuucguacguagcagagcCgcucccu...................................................................  1     1  7y1
....................uuucguacguagcagagcagAucccu...................................................................  2     1  7y1
....................uuucUuacguagcagagcagcucccu...................................................................  3     1  7y1
....................uuucguacguagcagaUcagcucccu...................................................................  1     1  7y1
....................uAucguacguagcagagcagcucccu...................................................................  9     1  7y1
....................uuucguacguagcagagcagcucccC...................................................................  152   1  7y1
....................uuucguacguagcagagcGgcucccu...................................................................  1     1  7y1
....................uCucguacguagcagagcagcucccu...................................................................  1     1  7y1
....................uuucguacguagcUgagcagcucccu...................................................................  1     1  7y1
....................uuucguacguagcGgagcagcucccu...................................................................  1     1  7y1
....................uuucguacguagcagagAagcucccu...................................................................  6     1  7y1
....................uGucguacguagcagagcagcucccu...................................................................  1     1  7y1
....................uuucguacUuagcagagcagcucccu...................................................................  7     1  7y1
....................uuucguacguaAcagagcagcucccu...................................................................  2     1  7y1
....................uuucCuacguagcagagcagcucccu...................................................................  1     1  7y1
....................uuucguacCuagcagagcagcucccu...................................................................  1     1  7y1
....................uuucguacguGgcagagcagcucccu...................................................................  2     1  7y1
....................Auucguacguagcagagcagcucccu...................................................................  42    1  7y1
....................uuucguacguaUcagagcagcucccu...................................................................  1     1  7y1
....................uuuAguacguagcagagcagcucccu...................................................................  6     1  7y1
....................uuucguacguagcagGgcagcucccu...................................................................  2     1  7y1
....................uuucguacguagcaUagcagcucccu...................................................................  1     1  7y1
....................uuucguacguagcagagcagcucccu...................................................................  4551  0  7y1
....................uuucguacgAagcagagcagcucccu...................................................................  7     1  7y1
....................uuAcguacguagcagagcagcucccu...................................................................  4     1  7y1
....................uuucguacguagcagagcagcAcccu...................................................................  1     1  7y1
```

**Star** **Mature**

```
gaccugcuucugggucggggguuucguacguagcagagcagcucccucgcugcgaucuauugaaagucagcccucgacacaaggguuuguccgcgcgcgcgcgcgcgugcgugc
....................uuucguGcguagcagagcagcucccu...................................................................   1     1   7y1
....................uuucguacguagcagagcagcCcccu...................................................................   1     1   7y1
....................uuucguacAuagcagagcagcucccu...................................................................   3     1   7y1
....................Guucguacguagcagagcagcucccu...................................................................   14    1   7y1
....................uuucguacguagcaAagcagcucccu...................................................................   1     1   7y1
....................uuucguaUguagcagagcagcucccu...................................................................   2     1   7y1
....................uuucguacguagcagagcagcuccAu...................................................................   4     1   7y1
....................uuucguacguagcagagcagcucccG...................................................................   17    1   7y1
....................uuucgAacguagcagagcagcucccu...................................................................   9     1   7y1
....................uuGcguacguagcagagcagcucccu...................................................................   1     1   7y1
....................uuucguacguagcagagcagcucccA...................................................................   8     1   7y1
....................uuucguaAguagcagagcagcucccu...................................................................   4     1   7y1
....................uuucguacguagUagagcagcucccu...................................................................   225   1   7y1
....................uuucguacguagcagagcagcucAcu...................................................................   3     1   7y1
....................uuucguacguagAagagcagcucccu...................................................................   2     1   7y1
....................Nuucguacguagcagagcagcucccu...................................................................   1     1   7y1
....................uuucguacguagcagCgcagcucccu...................................................................   10    1   7y1
....................Cuucguacguagcagagcagcucccu...................................................................   2     1   7y1
....................uuucguacguagcagaCcagcucccu...................................................................   2     1   7y1
....................uuucguacguagcagagcagcuAccu...................................................................   10    1   7y1
....................uuAcguacguagcagagcagcucccuc..................................................................   3     1   7y1
....................uuucCuacguagcagagcagcucccuc..................................................................   2     1   7y1
....................uuucguacguagcagagAagcucccuc..................................................................   6     1   7y1
....................uuucguacguagcagagcaCcucccuc..................................................................   1     1   7y1
....................Cuucguacguagcagagcagcucccuc..................................................................   1     1   7y1
....................uuucguacguagcagCgcagcucccuc..................................................................   10    1   7y1
....................uuucguacguagcagagcagcuAccuc..................................................................   7     1   7y1
....................uuucguacguagcagagcCgcucccuc..................................................................   1     1   7y1
....................Nuucguacguagcagagcagcucccuc..................................................................   6     1   7y1
....................uuucguacguagcagagcagcCcccuc..................................................................   1     1   7y1
....................uuucguacguaAcagagcagcucccuc..................................................................   1     1   7y1
....................uuucguacguagcagUgcagcucccuc..................................................................   2     1   7y1
....................uuucguacguagcagagcaUcucccuc..................................................................   2     1   7y1
....................uuucguacguagcagagcagcuccAuc..................................................................   5     1   7y1
....................uuucguacguagcaUagcagcucccuc..................................................................   2     1   7y1
....................uuucgCacguagcagagcagcucccuc..................................................................   1     1   7y1
....................uuucguacguagcagagcGgcucccuc..................................................................   2     1   7y1
....................uuucguaUguagcagagcagcucccuc..................................................................   1     1   7y1
....................uuucguacguagcagagcagcuGccuc..................................................................   1     1   7y1
....................uuucguacguagcagagcagcucAcuc..................................................................   6     1   7y1
....................uuucguacguaUcagagcagcucccuc..................................................................   1     1   7y1
....................uuucguacguagcagagcagAucccuc..................................................................   4     1   7y1
....................uuucguacguagcagagcagcucccuc..................................................................   4379  0   7y1
....................uuucUuacguagcagagcagcucccuc..................................................................   2     1   7y1
....................uuucguacguagcagagcUgcucccuc..................................................................   1     1   7y1
....................uuucguacguagAagagcagcucccuc..................................................................   5     1   7y1
....................uAucguacguagcagagcagcucccuc..................................................................   9     1   7y1
....................uuucguacguagUagagcagcucccuc..................................................................   1     1   7y1
....................Guucguacguagcagagcagcucccuc..................................................................   12    1   7y1
....................Auucguacguagcagagcagcucccuc..................................................................   39    1   7y1
....................uuucgAacguagcagagcagcucccuc..................................................................   9     1   7y1
....................uuucguacgAagcagagcagcucccuc..................................................................   7     1   7y1
....................uuuGguacguagcagagcagcucccuc..................................................................   1     1   7y1
....................uuucguaAguagcagagcagcucccuc..................................................................   3     1   7y1
....................uuucguacguagcagagcagcucccuA..................................................................   3     1   7y1
....................uuucguacguagcagagGagcucccuc..................................................................   1     1   7y1
....................uuucguacguagcagagcagcucUcuc..................................................................   1     1   7y1
....................uuucguacguagcaCagcagcucccuc..................................................................   2     1   7y1
....................uuucguacguagcagagcagcucccAc..................................................................   4     1   7y1
....................uuucguacguagcagagcagcAcccuc..................................................................   2     1   7y1
....................uuuAguacguagcagagcagcucccuc..................................................................   11    1   7y1
....................uuucguacguagcagagcagcuUccuc..................................................................   1     1   7y1
....................uuucguacguagcagagcagcucGcuc..................................................................   2     1   7y1
....................uuucguacCuagcagagcagcucccuc..................................................................   1     1   7y1
....................uuucguacUuagcagagcagcucccuc..................................................................   2     1   7y1
....................uuAcguacguagcagagcagcucccucgc................................................................   1     1   7y1
....................uuucguacguagcagCgcagcucccucgc................................................................   1     1   7y1
....................uuuAguacguagcagagcagcucccucgc................................................................   1     1   7y1
....................uuucguacguagUagagcagcucccucgc................................................................   1     1   7y1
....................uuucguacguagcagagcagcucAcucgc................................................................   1     1   7y1
```

```
  Star                                                         Mature
gaccugcuucugggucggggu uucguacguagcagagcagcucccucgcugcgaucuauugaaagucagcccucgacacaagguuuguccgcgcgcgcgcgcgcgugcgugc
....................uuucguacguagcagaUcagcucccucgc...............................................................   1      1   7y1
....................uuucguacguagcagagAagcucccucgc...............................................................   1      1   7y1
....................uuuGguacguagcagagcagcucccucgc...............................................................   1      1   7y1
....................uuucguacguagcagagcagcucccAcgc...............................................................   1      1   7y1
....................uuucguacguagcagagcUgcucccucgc...............................................................   1      1   7y1
....................Guucguacguagcagagcagcucccucgc...............................................................   2      1   7y1
....................uuucguacguagcagagcaCcucccucgc...............................................................   1      1   7y1
....................uuucguacguagcagagcagcucccucgc...............................................................   614    0   7y1
....................uuucguacgAagcagagcagcucccucgc...............................................................   1      1   7y1
....................uAucguacguagcagagcagcucccucgc...............................................................   1      1   7y1
....................uuucgAacguagcagagcagcucccucgc...............................................................   1      1   7y1
....................uuucguacCuagcagagcagcucccucgc...............................................................   1      1   7y1
....................uuucguacguagcagagcagcuAccucgc...............................................................   1      1   7y1
....................Auucguacguagcagagcagcucccucgc...............................................................   4      1   7y1
....................uuuAguacguagcagagcagcucccucgcu..............................................................   4      1   7y1
....................uuucguacguagGagagcagcucccucgcu..............................................................   1      1   7y1
....................uuucguacguagcagagcagcucccuGgcu..............................................................   1      1   7y1
....................uuucguacguagcagaCcagcucccucgcu..............................................................   1      1   7y1
....................Auucguacguagcagagcagcucccucgcu..............................................................   20     1   7y1
....................uuucguacguagcagagcagcCcccucgcu..............................................................   1      1   7y1
....................uuucguacguagAagagcagcucccucgcu..............................................................   3      1   7y1
....................uuucguacCuagcagagcagcucccucgcu..............................................................   1      1   7y1
....................uuucguacguagcCgagcagcucccucgcu..............................................................   1      1   7y1
....................uuucguacguagcagagcCgcucccucgcu..............................................................   2      1   7y1
....................uuucguacguaCcagagcagcucccucgcu..............................................................   1      1   7y1
....................uuucguacguagcagagcagcucUcucgcu..............................................................   2      1   7y1
....................uuucguUcguagcagagcagcucccucgcu..............................................................   1      1   7y1
....................uuucguacguagcagagAagcucccucgcu..............................................................   2      1   7y1
....................uuucguacguagcagagcGgcucccucgcu..............................................................   1      1   7y1
....................uuucguacguagcagagcaUcucccucgcu..............................................................   1      1   7y1
....................uuucguacguagcagUgcagcucccucgcu..............................................................   1      1   7y1
....................uuucguacguagcagagcUgcucccucgcu..............................................................   1      1   7y1
....................uuucguacguagcagaUcagcucccucgcu..............................................................   1      1   7y1
....................uuucguacguagcagagcagcuAccucgcu..............................................................   7      1   7y1
....................uuucguacguagcagagcagcucccuAgcu..............................................................   3      1   7y1
....................uuucguacguagcagGgcagcucccucgcu..............................................................   2      1   7y1
....................uuucguacguGgcagagcagcucccucgcu..............................................................   1      1   7y1
....................uuucgAacguagcagagcagcucccucgcu..............................................................   4      1   7y1
....................uuucguacguagcagagcagcucccucgcu..............................................................   2277   0   7y1
....................uuucguaAguagcagagcagcucccucgcu..............................................................   2      1   7y1
....................Guucguacguagcagagcagcucccucgcu..............................................................   178    1   7y1
....................uAucguacguagcagagcagcucccucgcu..............................................................   1      1   7y1
....................uuucguacguagcagagcagcucGcucgcu..............................................................   1      1   7y1
....................uuGcguacguagcagagcagcucccucgcu..............................................................   1      1   7y1
....................uuAcguacguagcagagcagcucccucgcu..............................................................   1      1   7y1
....................uuucguacguagcagCgcagcucccucgcu..............................................................   2      1   7y1
....................uuucguacgCagcagagcagcucccucgcu..............................................................   1      1   7y1
....................uuucguacgAagcagagcagcucccucgcu..............................................................   6      1   7y1
....................uuucguacguagcaCagcagcucccucgcu..............................................................   2      1   7y1
....................uuucguacUuagcagagcagcucccucgcu..............................................................   1      1   7y1
....................uuucguacguagcagagcagUucccucgcu..............................................................   1      1   7y1
....................uuucguacguagcagagcagcucccucAcu..............................................................   2      1   7y1
....................uuucAuacguagcagagcagcucccucgcu..............................................................   2      1   7y1
....................uuucguacguagcagagcagcucccucgAu..............................................................   3      1   7y1
....................uuucguacguagcagagcagcucccucgcG..............................................................   20     1   7y1
....................Nuucguacguagcagagcagcucccucgcu..............................................................   2      1   7y1
....................uuucguacguagcagagcagcucccAcgcu..............................................................   2      1   7y1
....................uuucguacguagcagagcagcucAcucgcu..............................................................   1      1   7y1
....................uuucguacCuagcagagcagcucccucgcug.............................................................   1      1   7y1
....................uuucAuacguagcagagcagcucccucgcug.............................................................   1      1   7y1
....................uuucguacguagcUgagcagcucccucgcug.............................................................   1      1   7y1
....................uuucguacguagcagagcGgcucccucgcug.............................................................   1      1   7y1
....................uuucguacguagcagagcagcucccucCcug.............................................................   1      1   7y1
....................uuucguacguagcagagcagcucccucgcGg.............................................................   9      1   7y1
....................Auucguacguagcagagcagcucccucgcug.............................................................   6      1   7y1
....................uuucguacguagcagagcagcucccAcgcug.............................................................   1      1   7y1
....................uuucguUcguagcagagcagcucccucgcug.............................................................   1      1   7y1
....................uuucguacguagcagagcagcucccucgcug.............................................................   855    0   7y1
....................uuucguacguagcagagcagcucccucUcug.............................................................   1      1   7y1
....................uuucguacgAagcagagcagcucccucgcug.............................................................   1      1   7y1
```

**Star** **Mature**

```
gaccugcuucugggucgggguuucguacguagcagagcagcucccucgcugcgaucuauugaaagucagcccucgacacaaggguuuguccgcgcgcgcgcgcgcgugcgugc
....................uuucguacguagcagagcaAcucccucgcug..............................................................	1	1	7y1
....................uuucUuacguagcagagcagcucccucgcug..............................................................	1	1	7y1
....................uuucguacguagcagagcagcuAccucgcug..............................................................	2	1	7y1
....................uuucguacguagcagagAagcucccucgcug..............................................................	2	1	7y1
....................uAucguacguagcagagcagcucccucgcug..............................................................	2	1	7y1
....................uuucguCcguagcagagcagcucccucgcug..............................................................	1	1	7y1
....................uuucguacguagcagagcagcucccucgGug..............................................................	1	1	7y1
....................uuucgAacguagcagagcagcucccucgcug..............................................................	2	1	7y1
....................uuucguacguagAagagcagcucccucgcug..............................................................	1	1	7y1
....................uuucguaAguagcagagcagcucccucgcug..............................................................	1	1	7y1
....................uuucguacguagcagagcagAucccucgcugc.............................................................	1	1	7y1
....................uAucguacguagcagagcagcucccucgcugc.............................................................	1	1	7y1
....................uuucguacguagcagagcagcAcccucgcugc.............................................................	1	1	7y1
....................uuucguacCuagcagagcagcucccucgcugc.............................................................	1	1	7y1
....................Guucguacguagcagagcagcucccucgcugc.............................................................	1	1	7y1
....................uuucguacguagcagagcagcucccucgcugc.............................................................	499	0	7y1
....................uuucguacguagcagagcagcucccucgcuCc.............................................................	1	1	7y1
....................uuucguacgAagcagagcagcucccucgcugc.............................................................	1	1	7y1
....................Auucguacguagcagagcagcucccucgcugc.............................................................	5	1	7y1
....................uuuAguacguagcagagcagcucccucgcugc.............................................................	1	1	7y1
....................uuucguacguagcagagcagcuAccucgcugc.............................................................	1	1	7y1
....................Nuucguacguagcagagcagcucccucgcugc.............................................................	2	1	7y1
....................uuucguacguagcagagcagcucccucgAugc.............................................................	1	1	7y1
.....................uucguacguagcaUagca..........................................................................	1	1	7y1
.....................uucguacguagcaCagca..........................................................................	1	1	7y1
.....................uucguacguagcagagAa..........................................................................	3	1	7y1
.....................uucguacguaCcagagca..........................................................................	1	1	7y1
.....................uucguacguaAcagagca..........................................................................	1	1	7y1
.....................uucguacguagcagGgca..........................................................................	1	1	7y1
.....................uucguacguagcagagGa..........................................................................	1	1	7y1
.....................uucguacguagAagagca..........................................................................	1	1	7y1
.....................uucguacgAagcagagca..........................................................................	3	1	7y1
.....................uucAuacguagcagagca..........................................................................	1	1	7y1
.....................uAcguacguagcagagca..........................................................................	4	1	7y1
.....................Nucguacguagcagagca..........................................................................	1	1	7y1
.....................uucguaAguagcagagca..........................................................................	8	1	7y1
.....................Gucguacguagcagagca..........................................................................	2	1	7y1
.....................uuAguacguagcagagca..........................................................................	4	1	7y1
.....................uucguacguagcagaAca..........................................................................	1	1	7y1
.....................uucguacguagcagagcG..........................................................................	1	1	7y1
.....................uNcguacguagcagagca..........................................................................	1	1	7y1
.....................Aucguacguagcagagca..........................................................................	12	1	7y1
.....................uucguacCuagcagagca..........................................................................	2	1	7y1
.....................uucguacguagcagagca..........................................................................	1212	0	7y1
.....................Cucguacguagcagagca..........................................................................	1	1	7y1
.....................uucguacguagcagagcC..........................................................................	1	1	7y1
.....................uucguacguagcagaUca..........................................................................	2	1	7y1
.....................uucguacAuagcagagcag.........................................................................	1	1	7y1
.....................uucguacguagcaCagcag.........................................................................	1	1	7y1
.....................Aucguacguagcagagcag.........................................................................	16	1	7y1
.....................uAcguacguagcagagcag.........................................................................	3	1	7y1
.....................Gucguacguagcagagcag.........................................................................	2	1	7y1
.....................uucguacguagcagagcag.........................................................................	1378	0	7y1
.....................uucguacguagAagagcag.........................................................................	1	1	7y1
.....................uucguaAguagcagagcag.........................................................................	4	1	7y1
.....................uucgCacguagcagagcag.........................................................................	1	1	7y1
.....................uucguacgAagcagagcag.........................................................................	2	1	7y1
.....................uucguacguagcagagAag.........................................................................	1	1	7y1
.....................uucguacguGgcagagcag.........................................................................	1	1	7y1
.....................uucguacguagcagagcaU.........................................................................	2	1	7y1
.....................uucguacCuagcagagcag.........................................................................	1	1	7y1
.....................uucguacguagcagagcUg.........................................................................	1	1	7y1
.....................uucguacguaUcagagcag.........................................................................	1	1	7y1
.....................uuAguacguagcagagcag.........................................................................	4	1	7y1
.....................uucguUcguagcagagcagc........................................................................	2	1	7y1
.....................uucguacgAagcagagcagc........................................................................	1	1	7y1
.....................uucguacAuagcagagcagc........................................................................	1	1	7y1
.....................Nucguacguagcagagcagc........................................................................	1	1	7y1
.....................uucguacUuagcagagcagc........................................................................	2	1	7y1
.....................uucguaGguagcagagcagc........................................................................	2	1	7y1
```

```
 Star                                                                  Mature
gaccugcuucugggucgggguuucguacguagcagagcagcucccucgcugcgaucuauugaaagucagcccucgacacaaggguuuguccgcgcgcgcgcgcgcgugcgugc
.....................uAcguacguagcagagcagc........................................................................    2      1    7y1
.....................uucguacguagcaCagcagc........................................................................    1      1    7y1
.....................uucCuacguagcagagcagc........................................................................    1      1    7y1
.....................uucguacCuagcagagcagc........................................................................    1      1    7y1
.....................uucguacguagcagagcaUc........................................................................    3      1    7y1
.....................uucguacguagcagaUcagc........................................................................    1      1    7y1
.....................uucguacguagcagaCcagc........................................................................    1      1    7y1
.....................Aucguacguagcagagcagc........................................................................    33     1    7y1
.....................uucguacguGgcagagcagc........................................................................    2      1    7y1
.....................uucguacguagcagagcagA........................................................................    2      1    7y1
.....................uuUguacguagcagagcagc........................................................................    1      1    7y1
.....................uucguacguagcaUagcagc........................................................................    1      1    7y1
.....................uucgAacguagcagagcagc........................................................................    1      1    7y1
.....................uucguacguagcUgagcagc........................................................................    1      1    7y1
.....................uucguaAguagcagagcagc........................................................................    2      1    7y1
.....................uucguacguaAcagagcagc........................................................................    1      1    7y1
.....................uCcguacguagcagagcagc........................................................................    1      1    7y1
.....................uucguacgCagcagagcagc........................................................................    1      1    7y1
.....................uucguacguagcaAagcagc........................................................................    1      1    7y1
.....................uucguacguagcagagcagc........................................................................    1790   0    7y1
.....................uucguacguagcagGgcagc........................................................................    1      1    7y1
.....................uuAguacguagcagagcagc........................................................................    5      1    7y1
.....................uucguacguagcagagcGgcu.......................................................................    5      1    7y1
.....................uucguacguagcagagAagcu.......................................................................    9      1    7y1
.....................uucguacguagUagagcagcu.......................................................................    185    1    7y1
.....................uucguacguagcagagcagcA.......................................................................    4      1    7y1
.....................uucguacguagcagagcagAu.......................................................................    4      1    7y1
.....................uucguacguagcGgagcagcu.......................................................................    1      1    7y1
.....................NucguacguagcagagcagcU.......................................................................    2      1    7y1
.....................uucguGcguagcagagcagcu.......................................................................    1      1    7y1
.....................uucguacguaUcagagcagcu.......................................................................    3      1    7y1
.....................uucguacguagcUgagcagcu.......................................................................    2      1    7y1
.....................uucguacguagcagagcUgcu.......................................................................    1      1    7y1
.....................uucCuacguagcagagcagcu.......................................................................    1      1    7y1
.....................uucguacgAagcagagcagcu.......................................................................    8      1    7y1
.....................uucUuacguagcagagcagcu.......................................................................    5      1    7y1
.....................uucguacguagcagagcaUcu.......................................................................    3      1    7y1
.....................uucguacguagcagagcaAcu.......................................................................    1      1    7y1
.....................uucguacguagcagaCcagcu.......................................................................    5      1    7y1
.....................uuGguacguagcagagcagcu.......................................................................    2      1    7y1
.....................uucguacguagcagagcagGu.......................................................................    2      1    7y1
.....................uucguacguagcagGgcagcu.......................................................................    1      1    7y1
.....................uucguacguagcagagcaCcu.......................................................................    3      1    7y1
.....................uucgGacguagcagagcagcu.......................................................................    1      1    7y1
.....................uucguacguagcagagcagcu.......................................................................    7198   0    7y1
.....................Aucguacguagcagagcagcu.......................................................................    80     1    7y1
.....................uucguacguagcagagcagcG.......................................................................    40     1    7y1
.....................uucgCacguagcagagcagcu.......................................................................    3      1    7y1
.....................uucguacguagAagagcagcu.......................................................................    3      1    7y1
.....................uucguacguagcagCgcagcu.......................................................................    7      1    7y1
.....................uucguacguGgcagagcagcu.......................................................................    2      1    7y1
.....................uucgAacguagcagagcagcu.......................................................................    9      1    7y1
.....................uucguacguagcagagGagcu.......................................................................    4      1    7y1
.....................uucguacguagcagagcagcC.......................................................................    1      1    7y1
.....................uucguacguUgcagagcagcu.......................................................................    1      1    7y1
.....................uucguaUguagcagagcagcu.......................................................................    3      1    7y1
.....................Gucguacguagcagagcagcu.......................................................................    8      1    7y1
.....................uucguacCuagcagagcagcu.......................................................................    4      1    7y1
.....................uucguacguagcaAagcagcu.......................................................................    1      1    7y1
.....................uucguUcguagcagagcagcu.......................................................................    1      1    7y1
.....................uuAguacguagcagagcagcu.......................................................................    24     1    7y1
.....................uucguacguaCcagagcagcu.......................................................................    3      1    7y1
.....................uAcguacguagcagagcagcu.......................................................................    18     1    7y1
.....................uucguacguagcagagUagcu.......................................................................    1      1    7y1
.....................uucguacguagcaUagcagcu.......................................................................    5      1    7y1
.....................uucguaGguagcagagcagcu.......................................................................    1      1    7y1
.....................uucguacguagcaCagcagcu.......................................................................    1      1    7y1
.....................uucguacAuagcagagcagcu.......................................................................    1      1    7y1
.....................uucguaAguagcagagcagcu.......................................................................    22     1    7y1
.....................uucguacguagcagagcCgcu.......................................................................    1      1    7y1
```

**Star** **Mature**

```
gaccugcuucugggucgggguuucguacguagcagagcagcucccucgcugcgaucuauugaaagucagcccucgacacaaggguuugucgcgcgcgcgcgcgcgugcgugc
.........................................ucccucgcugcgaucuauugGaaguc............................................ 1 1 7y1
.........................................uAccucgcugcgaucuauugaaaguc............................................ 3 1 7y1
.........................................ucccucgcugcUaucuauugaaaguc............................................ 1 1 7y1
.........................................ucccucgAugcgaucuauugaaaguc............................................ 3 1 7y1
.........................................ucccucgcugcgaucuauugaaaguc............................................ 2692 0 7y1
.........................................ucccuAgcugcgaucuauugaaaguc............................................ 7 1 7y1
.........................................ucccucgcugcgaucuauugaaaguA............................................ 1 1 7y1
.........................................ucccucgcGgcgaucuauugaaaguc............................................ 1 1 7y1
.........................................ucccucgcuUcgaucuauugaaaguc............................................ 2 1 7y1
.........................................ucccucAcugcgaucuauugaaaguc............................................ 1 1 7y1
.........................................ucccucgcugcgaucuGuugaaaguc............................................ 2 1 7y1
.........................................Acccucgcugcgaucuauugaaaguc............................................ 38 1 7y1
.........................................ucccucgcugAgaucuauugaaaguc............................................ 2 1 7y1
.........................................ucccAcgcugcgaucuauugaaaguc............................................ 1 1 7y1
.........................................ucccucgcugcCaucuauugaaaguc............................................ 3 1 7y1
.........................................ucccucgcugcgaucuauugaaagAc............................................ 2 1 7y1
.........................................ucccucUcugcgaucuauugaaaguc............................................ 2 1 7y1
.........................................ucccucgcugcgauAuauugaaaguc............................................ 4 1 7y1
.........................................uccAucgcugcgaucuauugaaaguc............................................ 2 1 7y1
.........................................Ncccucgcugcgaucuauugaaaguc............................................ 1 1 7y1
.........................................ucccucgGugcgaucuauugaaaguca........................................... 1 1 7y1
.........................................ucccucgcugcgaucuauugaaaguca........................................... 586 0 7y1
.........................................Ncccucgcugcgaucuauugaaaguca........................................... 2 1 7y1
.........................................ucccucgcugcgaucCauugaaaguca........................................... 1 1 7y1
.........................................ucccucgcugcgauAuauugaaaguca........................................... 1 1 7y1
.........................................Gcccucgcugcgaucuauugaaaguca........................................... 1 1 7y1
.........................................ucccucgcugAgaucuauugaaaguca........................................... 1 1 7y1
.........................................ucccucgcugcgaucuauugaaagucU........................................... 1 1 7y1
.........................................ucccucUcugcgaucuauugaaaguca........................................... 1 1 7y1
.........................................ucccucgcuAcgaucuauugaaaguca........................................... 1 1 7y1
.........................................ucccucgcuCcgaucuauugaaaguca........................................... 1 1 7y1
.........................................uccAucgcugcgaucuauugaaaguca........................................... 1 1 7y1
.........................................ucccucgcugcCaucuauugaaaguca........................................... 1 1 7y1
.........................................ucccucgAugcgaucuauugaaaguca........................................... 2 1 7y1
.........................................ucccucCcugcgaucuauugaaaguca........................................... 1 1 7y1
.........................................Acccucgcugcgaucuauugaaaguca........................................... 6 1 7y1
.........................................ucccucgcuUcgaucuauugaaagucag.......................................... 1 1 7y1
.........................................ucAcucgcugcgaucuauugaaagucag.......................................... 1 1 7y1
.........................................ucccucgcugcgaucuauAgaaagucag.......................................... 1 1 7y1
.........................................ucccucgcugcgaucuauugaaagucag.......................................... 159 0 7y1
.........................................Acccucgcugcgaucuauugaaagucag.......................................... 2 1 7y1
.........................................ucccucgcugcgUucuauugaaagucag.......................................... 1 1 7y1
.........................................ucccAcgcugcgaucuauugaaagucag.......................................... 1 1 7y1
..........................................cccucgcugcgaucuauu................................................... 172 0 7y1
..........................................cccucgcugcgaucuaAu................................................... 1 1 7y1
..........................................Gccucgcugcgaucuauu................................................... 1 1 7y1
..........................................ccAucgcugcgaucuauu................................................... 1 1 7y1
..........................................cAcucgcugcgaucuauu................................................... 1 1 7y1
..........................................cccucgcugcCaucuauu................................................... 2 1 7y1
..........................................cccucgcAgcgaucuauu................................................... 1 1 7y1
..........................................cccucgcugcgaucuauuga................................................. 157 0 7y1
..........................................Accucgcugcgaucuauuga................................................. 1 1 7y1
..........................................ccAucgcugcgaucuauuga................................................. 1 1 7y1
..........................................cccucgAugcgaucuauuga................................................. 1 1 7y1
..........................................cAcucgcugcgaucuauuga................................................. 2 1 7y1
..........................................cccucgcAgcgaucuauuga................................................. 1 1 7y1
..........................................cccucgcugcgaucuauugG................................................. 1 1 7y1
..........................................Nccucgcugcgaucuauuga................................................. 1 1 7y1
..........................................cccucgcugcgaucuauuAaa................................................ 1 1 7y1
..........................................cAcucgcugcgaucuauugaaa............................................... 3 1 7y1
..........................................cccucgAugcgaucuauugaaa............................................... 3 1 7y1
..........................................cccucgcugcUaucuauugaaa............................................... 1 1 7y1
..........................................cccucgcugcgaucuaAugaaa............................................... 3 1 7y1
..........................................cccucgcugcgaucuauAgaaa............................................... 1 1 7y1
..........................................cccucgcuUcgaucuauugaaa............................................... 2 1 7y1
..........................................cccucgcugcCaucuauugaaa............................................... 1 1 7y1
..........................................cUcucgcugcgaucuauugaaa............................................... 1 1 7y1
..........................................ccGucgcugcgaucuauugaaa............................................... 1 1 7y1
..........................................cccucgcugcgaucuauugaaa............................................... 1120 0 7y1
..........................................cccucgcugcgaAcuauugaaa............................................... 2 1 7y1
```

**Star** **Mature**

gaccugcuucugggucggggguuucguacguagcagagcagcucccucgcugcgaucuauugaaagucagcccucgacacaagguuugu ccgcgcgcgcgcgcgcgugcgugc
